# Crystallographic characterisation and development of bi-substrate inhibitors of coronavirus nsp14 methyltransferase

**DOI:** 10.1101/2025.09.30.679521

**Authors:** Irene Georgiou, Colin Robinson, Sean O’Byrne, Alex Matsuda, Przemysław Grygier, Craig D. Smith, Sandra O’Neill, Shamshad A. Ahmad, Suzanne Norval, John M. Post, G. J. Mirjam Groenewold, Nadya Urakova, Patrick Wanningen, Leanid Kresik, Jacek Plewka, Adrien Delpal, Kexin See, Thomas Eadsforth, Kinga Wierzbicka, Etienne Decroly, Kumar Singh Saikatendu, Edcon Chang, Eric J. Snijder, Krzysztof PyrĆ, Anna Czarna, Duncan Scott, Ian H. Gilbert

## Abstract

SARS-CoV-2 non-structural protein 14 (nsp14) is essential for viral mRNA cap guanine-N7 methylation and represents a promising but underexplored antiviral target. Herein we describe a structure-guided campaign based on a hit from a focussed SAM mimetic library. Systematic SAR exploration guided by six X-ray co-crystal structures in complex with SARS-CoV-2 led to compound **26**, a bi-substrate inhibitor that bridges the SAM and RNA cap binding sites. Compound **26** achieved nanomolar potency against nsp14 from SARS-CoV-2 (IC_50_ = 53 nM), SARS-CoV-1, and two alphacoronaviruses, with excellent selectivity over human RNMT and flaviviral MTase. In general, the compounds demonstrated favourable metabolic stability, passive permeability, and no HepG2 cytotoxicity. However, cellular antiviral activity was limited, revealing disconnects between enzyme inhibition and phenotypic response. These findings provide a structural framework for optimizing bi-substrate methyltransferase inhibitors against coronaviruses with a view for pan-coronaviral activity.

## Introduction

The emergence of severe acute respiratory syndrome coronavirus 2 (SARS-CoV-2) and the resulting COVID-19 pandemic have drawn significant attention to the development of antiviral compounds against this novel virus.1 Extensive efforts have been dedicated to unravelling the life cycle of SARS-CoV-2, particularly its entry into host cells. ^2-4^ The virus employs a receptor-based mechanism, with its spike (S) protein interacting with the human angiotensin-converting enzyme 2 (ACE2). Once inside the cell, the viral RNA is released from its protective protein shell. Subsequent translation of open reading frames (ORF1a and ORF1b) produces precursor polypeptides pp1a and pp1ab, which are cleaved by the viral main protease (Mpro) and papain-like protease (PLpro). In this manner, polyprotein processing yields 16 mature non-structural proteins (nsps) that engage in viral RNA synthesis and a range of interactions with the host cell that serve to facilitate virus reproduction.

Not all of the replicative enzymes of coronaviruses have been validated for drug development, however there is encouraging genetic validation *in vitro* that knocking out the methyltransferase (MTase) activity of nsp14 leads to severely compromised or non-viable virus,^5-7^ underscoring the potential significance of directing therapeutic attention towards this enzyme.

Given their replication in the cytoplasm, coronaviruses have evolved their own enzymatic pathway for mRNA capping, involving the combined activities of nsp9, nsp12, nsp13, nsp14, and nsp16.^2-4, 8, 9^ Nsp14 consists of two domains: the exoribonuclease (ExoN) domain^10^ and the MTase domain. The MTase domain plays a crucial role in viral mRNA capping by transferring a methyl group from the co-factor S-adenosyl methionine (SAM) to the N7-position of the guanosine moiety of RNA cap substrate, resulting in the RNA cap-0 structure and the release of S-adenosyl-L-homocysteine (SAH) by-product.11, 12 This process is followed by a further methylation by nsp16 on the 2’-ribose hydroxyl of the next base, culminating in the formation of cap-1.13 While an interaction with nsp10 strongly promotes the ExoN activity of nsp14, the MTase activity remains barely affected.14 The RNA-cap structure holds major significance in eukaryotic cells, as it facilitates efficient mRNA translation and shields against mRNA degradation.15 Furthermore, viral mRNA lacking a cap structure can trigger the induction of innate immunity responses in the host cell.16 Experiments have demonstrated that mutations in nsp14 leading to reduced N7-MTase activity result in compromised virus performance *in vivo*, particularly observed in an animal model of SARS-CoV-2 infection that employs mice expressing human ACE2.^5^ The essential role of nsp14-mediated N7-methylation has been validated for a number of betacoronaviruses^7^ involving nsp14 mutants, where certain mutants failed to generate viable virus progeny, and engineered mutants displayed decreased pathogenicity in wild-type mice due to reduced N7-MTase activity.

Therefore, the MTase activity of nsp14 presents an appealing avenue for drug discovery. Numerous inhibitors targeting SARS-CoV-2 nsp14 MTase, discovered via screening commercially accessible libraries, have shown relatively modest biochemical potency. ^17-19^ Virtual screenings have been conducted to pinpoint potential inhibitors for nsp14 MTase.^20, 21^ However, the most potent strategy to target this protein has been the design and testing of analogues of the cofactor S-adenosylmethionine (SAM).^19, 20^

Several studies have investigated SAM-based series as potential inhibitors for SARS-CoV-2 nsp14 MTase. Devkota *et al*. employed a radiometric MTase assay to screen a library of SAM competitive inhibitors and analogues, identifying potent compounds, including a weaker one that exhibited bi-substrate inhibition, competing against both SAM and the RNA substrate.^22^ Bobiļeva *et al*. reported a series of thioether SAM-mimetics with strong activity (single-digit nM IC_50_).^23^ Docking models suggested that these molecules exclusively bind within the SAM pocket. These compounds displayed comparable potency against both SARS-CoV-2 nsp14 and nsp16, raising the possibility of dual nsp14/16 inhibitors. However, they were also active against the human guanosine N7-MTase (GNMT or RNMT), highlighting the difficultly of achieving selectivity for this class of molecules. Leveraging the high sequence similarity between SARS-CoV-2 nsp14 and SARS-CoV nsp14, a set of SAM-derived inhibitors was designed which varied the adenosine base with different aromatic groups. The most effective compound demonstrated an IC_50_ of 3.0 ± 0.5 nM against SARS-CoV-2 nsp14.^24^ Adhmed-Belkacem *et al*. reported a series of SAM-mimetic inhibitors, with the best compounds exhibiting an IC_50_ of 19 nM.^25^ Interestingly, computational modelling suggested that these compounds function as bi-substrate inhibitors, anchored in the SAM-binding site while projecting a group into the guanosine-binding pocket that binds the RNA substrate. Building on this work, Hausdorff *et al*. recently described a series of SAM-mimetics with enhanced potency (sub-nM range).^26^

However, despite the description of numerous highly potent inhibitors against nsp14 MTase, these compounds display poor antiviral activity against SARS-CoV-2 replication in infected cells. To date, the exception to this is a SAM-based sulfonamide described by Jung *et al*., which they report has a SARS-CoV-2 antiviral EC_50_ of 0.72 µM in an immunofluorescence antiviral assay.^27^ The poor antiviral activity for this class of molecules has been investigated by Stefek *et al*. ^28^ They have reported a series of potent SAM-based nsp14 inhibitors but observed no significant inhibition of SARS-CoV-2 replication, up to concentrations of 100 µM, in both Vero E6 and Calu-3 cells. The lack of antiviral activity with biochemically potent compounds raised questions pertaining to compound cell permeability and uptake, but subsequent measurement showed that the compounds were present at intracellular levels exceeding their biochemical IC_50_ by>10,000-fold. The poor efficacy of these compounds on virus replication is therefore confusing and intriguing, especially given the genetic validation of the target.^7^ Whilst SARS-CoV-2 is of immediate interest, there are other betacoronaviruses such as Middle East respiratory syndrome coronavirus (MERS-CoV) which represents an ongoing threat to human health. More broadly, human coronaviruses (HCoVs) from the alphacoronavirus genus, such as HCoV-NL63 and -229E also infect humans. For effective preparedness against any future coronavirus outbreaks or pandemics, as broad activity as possible of small-molecule antivirals is highly desirable.

## RESULTS AND DISCUSSION

In the pursuit of effective nsp14 inhibitors, a library comprising 6683 SAM-mimetic compounds, underwent screening against SARS-CoV-2 nsp14 using a mass-directed (RapidFire) biochemical assay.^19^ Following hit validation, compound **1** emerged as a confirmed hit, exhibiting an IC_50_ of 0.42 ± 0.2 µM (n=13) against SARS-CoV-2 nsp14 in the biochemical assay (Figure 1A). To further validate the activity of compound **1**, it underwent evaluation in an orthogonal MTase-Glo assay, measuring a similar IC_50_ of 0.11 ± 0.05 µM. Compound **1** showed comparable activity in RapidFire biochemical assays against nsp14 from both the alphacoronavirus HCoV-229E (IC_50_=0.31 ± 0.008 µM) and HCoV-NL63 (IC_50_=0.19 ± 0.03 µM). Excellent selectivity over the human N7-MTase RNMT (IC_50_ > 100 µM) was also observed, indicating that compound **1** was a good starting point, selective for both alpha and betacoronavirus nsp14 but inactive against human RNMT. The mouse microsomal clearance was low and no toxicity against human HepG2 cells was observed (**Table S2**). The crystal structure of compound **1** bound to SARS-CoV-2 nsp14 is shown in Figure 1B, with some of the key interactions highlighted (pdb code: 9S0M). An overlay of our SARS-CoV-2 nsp14 structure with compound **1** bound and a SARS-CoV nsp14 structure with SAH and/or SAH/GpppA bound (pdb code: 5c8s)^29^ is shown for comparison in Figure 1C and D.

**Figure 1.**
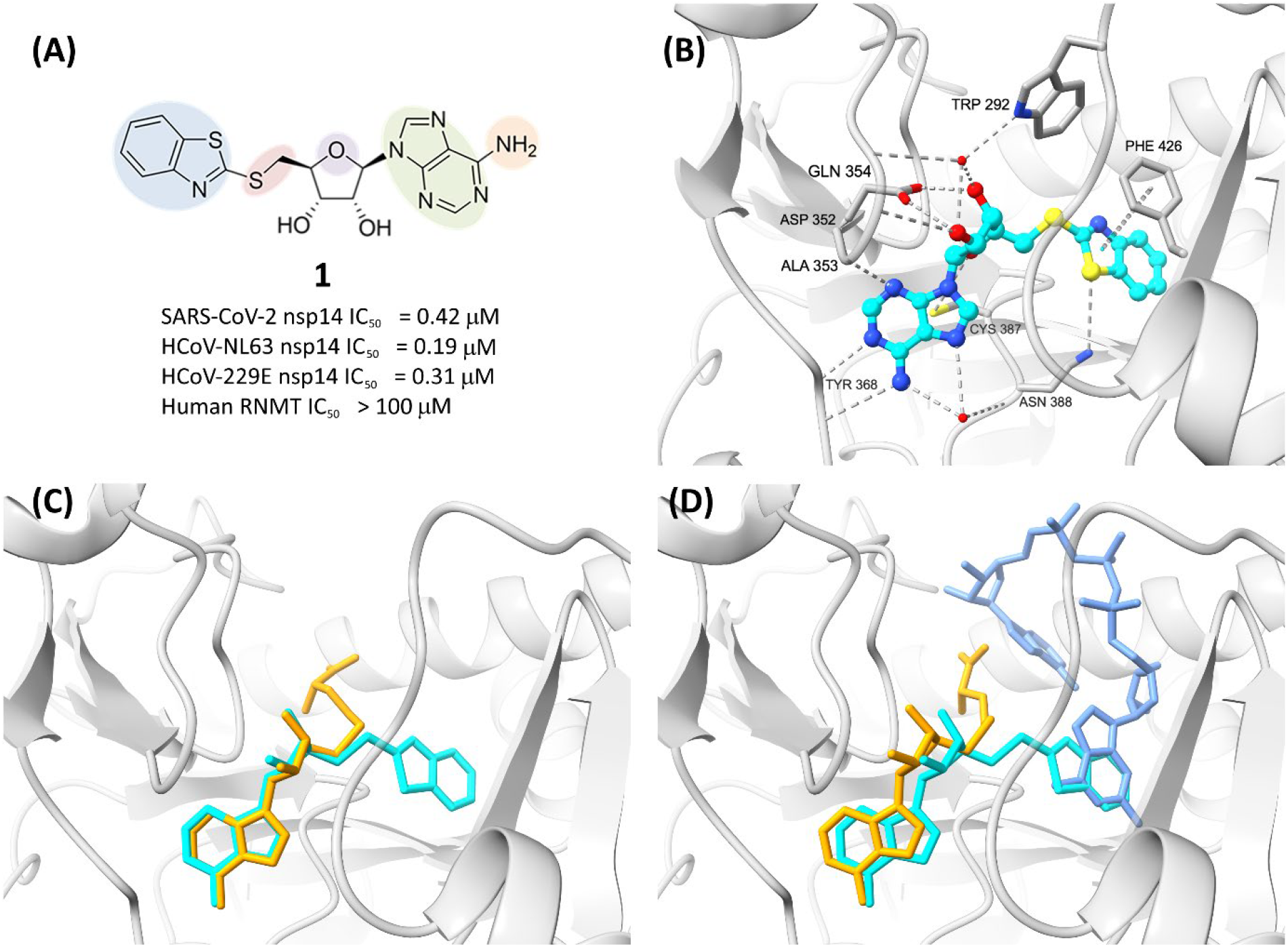
(A) Structure of hit compound **1**, an inhibitor of the methyltransferase activity of SARS-CoV-2 nsp14, IC_50_ = 0.42 ± 0.2 µM (pdb code: 9s0m). (B) Key residues interacting with compound **1** bound to SARS-CoV-2 nsp14. (C) Overlay of X-ray structure of compound **1** (cyan) bound to SARS-CoV-2 nsp14 (grey) and the SARS-CoV-1 nsp14 complex (pdb code: 7r2v, protein not shown) with SAH (orange). (D) Overlay of X-ray structure of compound **1** (cyan) bound to SARS-CoV-2 nsp14 (grey) and the SARS-CoV-1 nsp14 complex (pdb code: 5c8s, protein not shown) with SAH (orange) and GpppA (blue). Hydrogen bonds are shown as dashed lines (grey).

Compound **1** binds within the same cleft as SAH in nsp14, closely mimicking SAH’s hydrogen bonding interactions with the amino-purinyl moiety via Ala353 and Tyr368. Additionally, it forms hydrogen bonds between the ribose moiety and Asp352, along with a water-mediated interaction with Gln354. Compound **1** extends into the GpppA pocket, where the benzothiazole ring engages in a π-π stacking interaction with Phe426 (Figure 1B-D). This binding into both the SAM and RNA binding pocket not only secures compound **1** within the active site but also potentially improves specificity with respect to other MTases.

We systematically explored the structural features of compound **1** with the aim of designing inhibitors possessing enhanced biochemical potency and physicochemical properties. Our approach focused on modifications across five components of the compound highlighted in Figure 1A, left to right; (1) substitution of the benzothiazole ring with alternative aromatic or saturated systems; (2) alterations to the linker connecting the 5’-deoxyadenosine and the benzothiazole; (3) modifications of the 5’-deoxyadenosine; (4) exploration of a different nucleoside base; and (5) functionalization of the amine moiety of the base.

We initiated our structure-activity relationship (SAR) studies by investigating various replacements for the benzothiazole group, compounds **2**-**17**, Table 1. By varying the group in this portion of the molecule, the potency was modulated from inactive (IC_50_ > 100 µM, compound **11**) to IC_50_ around 0.3 µM (**2, 6** and **7**). Some weak regioselectivity for the position of a methyl substitution on the benzothiazole ring was observed in **2** and **3** (IC_50_ = 0.26 ± 0.07 µM and IC_50_ = 1.2 ± 0.15 µM respectively). Interestingly, the least active compound, thiazole **11**, completely lacks the aromatic 6-membered ring of the benzothiazole suggesting an important interaction of the Phe426 ring with aromatic groups. Relatively flat SAR is observed across this subset of compounds, indeed a 6,5 or 6,6 aromatic system with the appropriate connectivity as compound **1** display relatively similar potencies. Of note, analogues **12** and **16**, which both present the pendent aromatic group with a different connectivity, are both much less active. In general, the mouse microsomal clearance was rather low for this set of molecules, with no significant instabilities introduced, with perhaps the exception of compound **4**. With no significant advantages in either potency or mouse microsomal clearance **(Table S2)** identified for this subset, the benzothiazole was taken forward into subsequent SAR studies.

**Table 1.**
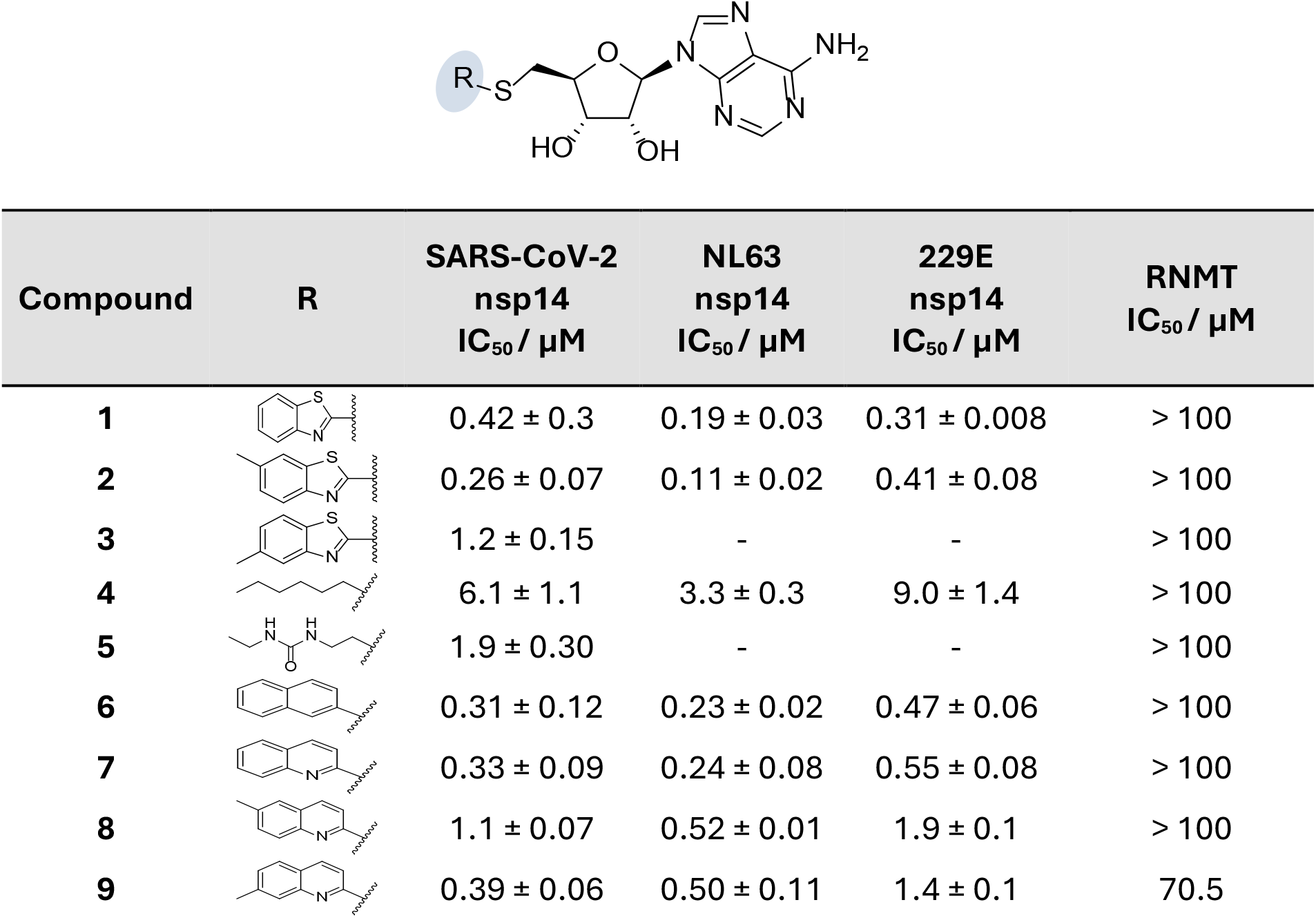
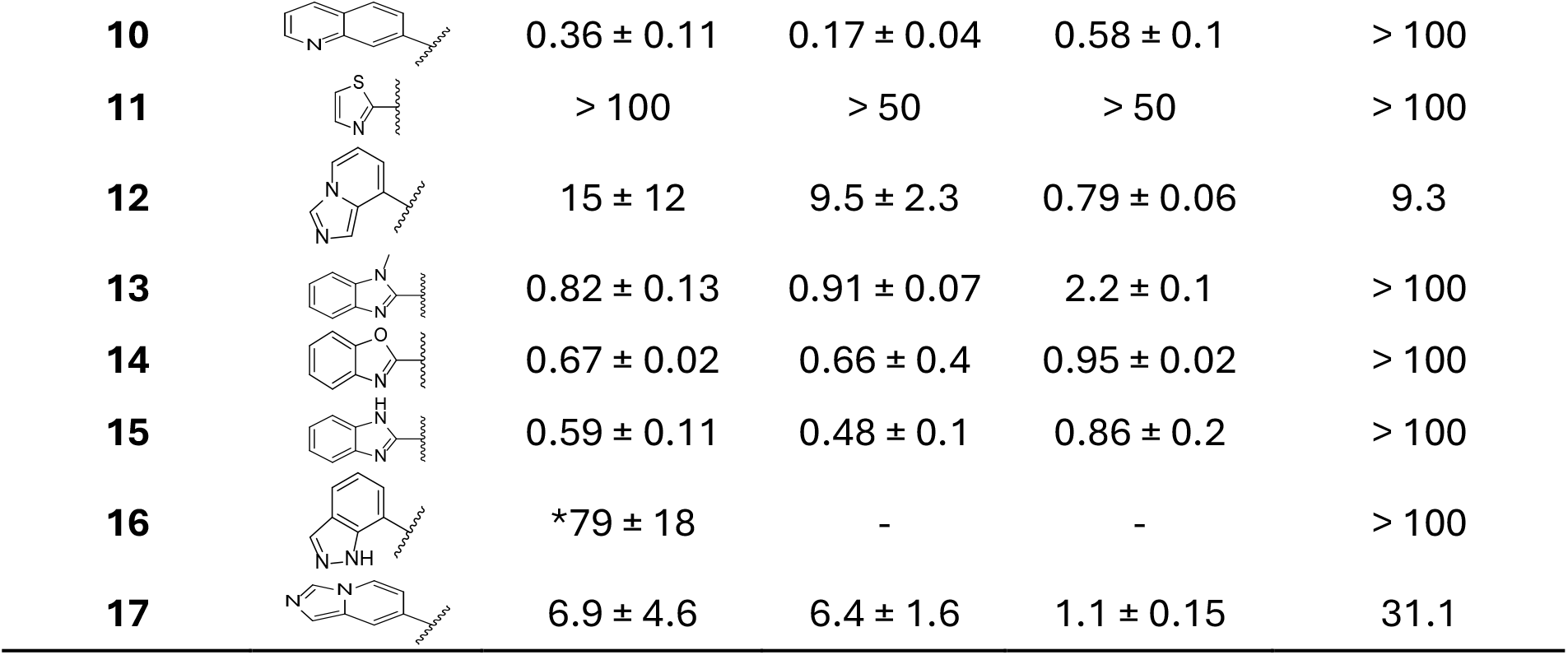
Enzymatic inhibition of compounds against SARS-CoV-2 nsp14, NL63 nsp14, 229E nsp14 and human RNMT. * one IC_50_ measured > 100 µM. Repeat N-values are given in Table S5.

| Compound | R | SARS-CoV-2<br>nsp14<br>$IC_{50} / \mu M$ | NL63<br>nsp14<br>$IC_{50} / \mu M$ | 229E<br>nsp14<br>$IC_{50} / \mu M$ | RNMT<br>$IC_{50} / \mu M$ |
| --- | --- | --- | --- | --- | --- |
| **1** | | $0.42 \pm 0.3$ | $0.19 \pm 0.03$ | $0.31 \pm 0.008$ | $> 100$ |
| **2** | | $0.26 \pm 0.07$ | $0.11 \pm 0.02$ | $0.41 \pm 0.08$ | $> 100$ |
| **3** | | $1.2 \pm 0.15$ | - | - | $> 100$ |
| **4** | | $6.1 \pm 1.1$ | $3.3 \pm 0.3$ | $9.0 \pm 1.4$ | $> 100$ |
| **5** | | $1.9 \pm 0.30$ | - | - | $> 100$ |
| **6** | | $0.31 \pm 0.12$ | $0.23 \pm 0.02$ | $0.47 \pm 0.06$ | $> 100$ |
| **7** | | $0.33 \pm 0.09$ | $0.24 \pm 0.08$ | $0.55 \pm 0.08$ | $> 100$ |
| **8** | | $1.1 \pm 0.07$ | $0.52 \pm 0.01$ | $1.9 \pm 0.1$ | $> 100$ |
| **9** | | $0.39 \pm 0.06$ | $0.50 \pm 0.11$ | $1.4 \pm 0.1$ | 70.5 |
| **10** | | $0.36 \pm 0.11$ | $0.17 \pm 0.04$ | $0.58 \pm 0.1$ | > 100 |
| **11** |  | > 100 | > 50 | > 50 | > 100 |
| **12** | | $15 \pm 12$ | $9.5 \pm 2.3$ | $0.79 \pm 0.06$ | 9.3 |
| **13** | | $0.82 \pm 0.13$ | $0.91 \pm 0.07$ | $2.2 \pm 0.1$ | > 100 |
| **14** | | $0.67 \pm 0.02$ | $0.66 \pm 0.4$ | $0.95 \pm 0.02$ | > 100 |
| **15** | | $0.59 \pm 0.11$ | $0.48 \pm 0.1$ | $0.86 \pm 0.2$ | > 100 |
| **16** | | *$79 \pm 18$ | - | - | > 100 |
| **17** | | $6.9 \pm 4.6$ | $6.4 \pm 1.6$ | $1.1 \pm 0.15$ | 31.1 |

Next, we looked at reducing overall polarity and exploring the potential for enhanced binding by substitution at the N6-adenine position. The structural biology reveals the adenine NH_2_ makes a hydrogen-bond interaction with the backbone carbonyl of Tyr368 but is otherwise relatively solvent-exposed in a narrow channel, suggesting opportunities for modest modification. However, the mono-methylation at this position in compound **18**, resulted in a significant drop of efficacy, while di-methylation in compound **19** led to a complete loss of activity. The addition of larger groups, including cyclopentyl, isopropyl and oxetanyl led to inactive compounds **20**-**22**. Additionally, we decided to replace the potentially labile thioether linker between the 5’-deoxyadenosine and the thiazole ring. Unfortunately, the O-or N-linked variants **23, 24** were found to be inactive, in contrast to the methylene derivative **25**, which exhibited equipotent activity to the parent compound, **Table 2**. From an overlay of compound **1** bound and a structure with SAH bound, the sulfur atom of the thioether of compound **1** binds in the same location as the equivalent sulfur atom of SAM/SAH, in a hydrophobic pocket formed by Trp292 and Phe426. The hydrophobic nature of this interaction may explain the selectivity of thioether **1** and carbon-linked **25** over the more polar ether **23** and amine **24**. Conformational effects on the preferred ground-state of the small molecules may also have additional effect. The low mouse microsomal clearance of the hit was maintained in compound **25**.

**Table 2.** Enzymatic inhibition against SARS-CoV-2 nsp14, human RNMT, CHI-logD and mouse microsomal stability. Repeat N-values are given in Table S5.

| Compound | X | R | SARS-CoV-2 nsp14<br>IC<sub>50</sub> / μM | RNMT<br>IC<sub>50</sub> / μM | CHI-logD | Mouse liver Mic<br>Clint<br>mL/min/g | T<sub>1/2</sub> /<br>min |
| --- | --- | --- | --- | --- | --- | --- | --- |
| **1** | S | NH<sub>2</sub> | 0.42 ± 0.3 | > 100 | 1.2 | 1.0 | > 30 |
| **18** | S | NHMe | 2.1 ± 0.2 | - | 1.3 | 3.4 | 19.7 |
| **19** | S | NMe<sub>2</sub> | > 100 | > 100 | 1.8 | - | - |
| **20** | S | NH-cyclopentyl | > 100 | > 100 | 2.5 | - | - |
| **21** | S | NH-*iso*-<br>propyl | > 100 | > 100 | 2.1 | - | - |
| **22** | S | NH-oxetanyl | 57 ± 18 | > 100 | 1.4 | - | - |
| **23** | O | NH<sub>2</sub> | > 100 | > 100 | 0.5 | - | - |
| **24** | NH | NH<sub>2</sub> | > 100 | 54.6 | 0.7 | - | - |
| **25** | CH<sub>2</sub> | NH<sub>2</sub> | 0.31 ± 0.07 | > 100 | 1.0 | 1.1 | > 30 |

**Table 3.** Alterations to the ribose and adenosine moiety and impact on enzymatic inhibition. Repeat N-values are given in Table S5.

|  | X | Y | R | SARS-CoV-2<br>nsp14<br>IC<sub>50</sub> / µM | NL63<br>nsp14<br>IC<sub>50</sub> / µM | 229E<br>nsp14<br>IC<sub>50</sub> / µM | RNMT<br>IC<sub>50</sub> / µM |
| --- | --- | --- | --- | --- | --- | --- | --- |
| **1** | O | N | - | 0.420 ± 0.3 | 0.19 ± 0.03 | 0.31 ± 0.008 | > 100 |
| **26** | CH<sub>2</sub> | N | - | 0.053 ± 0.03 | 0.008 ± 0.003 | 0.021 ± 0.004 | > 100 |
| **27** | O | C | H | 0.055 ± 0.02 | 0.039 ± 0.006 | 0.041 ± 0.004 | > 100 |
| **28** | CH<sub>2</sub> | C | H | 0.049 ± 0.005 | 0.016 ± 0.007 | 0.051 ± 0.021 | 22.8 |
| **29** | O | C | CN | 0.170 ± 0.02 | 0.073 ± 0.026 | 0.10 ± 0.007 | > 100 |
| **30** | O | C | C≡CH | 0.054 ± 0.01 | 0.029 ± 0.009 | 0.033 ± 0.002 | > 100 |
| **31** | O | C | Et | 0.250 ± 0.2 | 0.052 ± 0.009 | 0.12 ± 0.01 | > 100 |

Two previously implemented strategies in improving SAM mimetic potency involve the replacement of the ribose ring or the adenine moiety.^27^ In our hands both strategies yielded more potent compounds. In particular, replacing the ribose ring with an all-carbon cyclopentyl moiety yielded compound **26** with improved IC_50_ value of 0.05 ± 0.03 µM. The crystal structure of **1** reveals the ribose oxygen packs against the lipophilic sidechain of Cys387, potentially increasing van der Waals interaction with the cyclopentyl compound **26**. In the second strategy, the adenine base was substituted with a 7-deazaadenine to give compound **27** with a IC_50_ of 0.05 ± 0.02 µM. Unfortunately, the combination of both strategies did not achieve the desired additive effect, with compound **28** showing no improvement in potency, IC_50_ = 0.049 ± 0.005 µM. Potentially the two modifications of the ribose and base may alter the ground state conformation in a mutually counteracting manner. Additionally, extension at the 7-position of the base resulted in analogues **29** – **31** with comparable inhibitory activity but reduced metabolic stability, suggesting the introduction of a metabolic soft-spot (**Table S3**). These findings underscore the complexity of optimising both potency and metabolic properties in SAM mimetic analogues. Further X-ray crystal structures were solved, revealing the complex between nsp14 and compounds **5, 6, 18, 26** and **27, Figure 2**.

**Figure 2.**
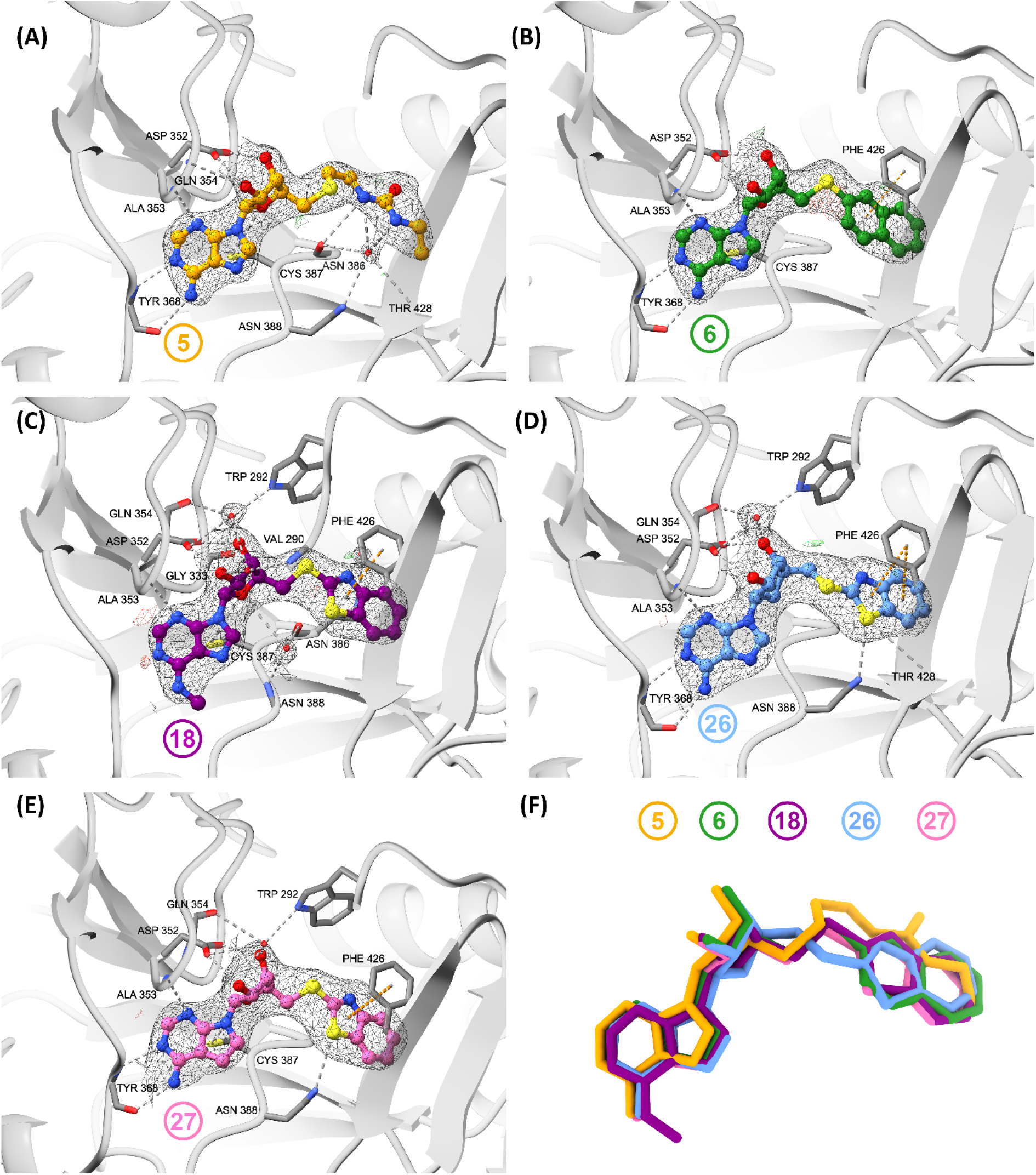
X-ray crystallographic structure of SARS-CoV-2 nsp14 in complex with (A) compound **5**, (B) compound **6**, (C) compound **18**, (D) compound **26** and (E) compound **27**. (F) Superimposition of the five compounds (**5, 6, 18, 26**, and **27**), demonstrating the conservation of the binding modes. PDB codes for structures nsp14 bound to compound **5, 6, 18, 26** and **27** are respectively; 9SAJ, 9SAK, 9SAL, 9SAM and 9SAN. Hydrogen bonds and π-stacking interactions are shown as dashed lines (grey and orange, respectively), and the electron density is shown as a mesh, 2Fo-fc: +1.0σ (grey); Fo-Fc omit map: +3.0σ (green); Fo-Fc omit map: −3.0σ (red).

Jung *et al*.^27^ have previously reported the only antiviral activity of an SAM-binding nsp14 inhibitor to date, where the adenosine base (“Base A”) **32** is replaced with a pyrrolo[2,1-f][1,2,4]-triazin-4-amine ring (“Base B”) **34**, which is the adenine-replacement present in Remdesivir. It is a standout result as the successful translation of biochemical activity to antiviral activity is rarely reported and poorly understood for this family of nsp14 inhibitors against SARS-CoV-2. ^27^ Therefore, we sought to repeat this result and apply the same change of base to our most potent compounds in order to potentially advance their activity in infected cells in a similar way as reported. A selection of compounds with either base A or B were therefore synthesised, **Table 4**.

**Table 4.**
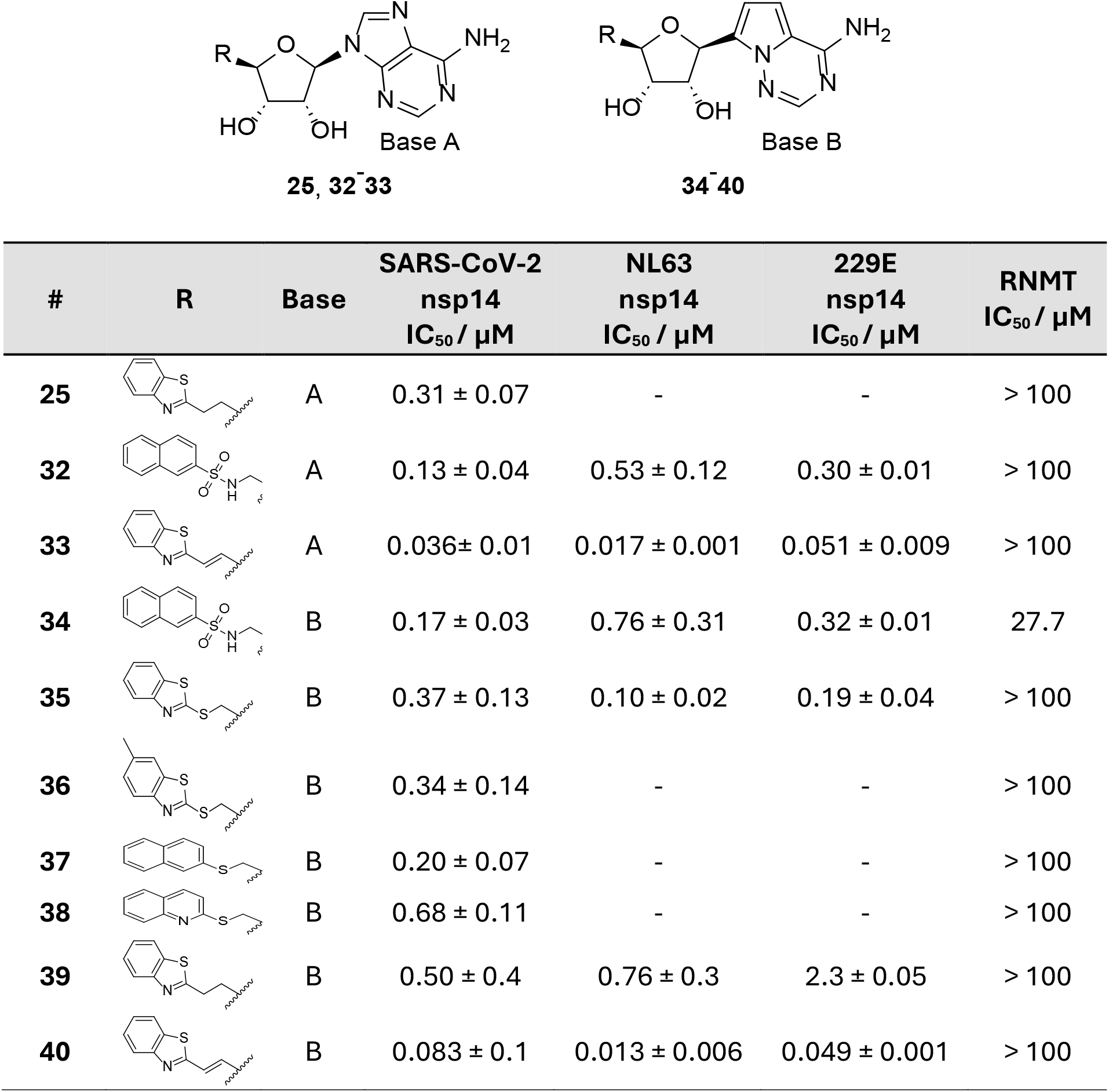
Activity of compounds **25** and **32**-**40**, with variation of the guanosine pocket binding group R and the base A or B. Repeat N-values are given in Table S5.

| # | R | Base | SARS-CoV-2<br>nsp14<br>IC<sub>50</sub> / μM | NL63<br>nsp14<br>IC<sub>50</sub> / μM | 229E<br>nsp14<br>IC<sub>50</sub> / μM | RNMT<br>IC<sub>50</sub> / μM |
| --- | --- | --- | --- | --- | --- | --- |
| **25** |  | A | 0.31 ± 0.07 | - | - | > 100 |
| **32** |  | A | 0.13 ± 0.04 | 0.53 ± 0.12 | 0.30 ± 0.01 | > 100 |
| **33** |  | A | 0.036 ± 0.01 | 0.017 ± 0.001 | 0.051 ± 0.009 | > 100 |
| **34** |  | B | 0.17 ± 0.03 | 0.76 ± 0.31 | 0.32 ± 0.01 | 27.7 |
| **35** |  | B | 0.37 ± 0.13 | 0.10 ± 0.02 | 0.19 ± 0.04 | > 100 |
| **36** |  | B | 0.34 ± 0.14 | - | - | > 100 |
| **37** |  | B | 0.20 ± 0.07 | - | - | > 100 |
| **38** |  | B | 0.68 ± 0.11 | - | - | > 100 |
| **39** |  | B | 0.50 ± 0.4 | 0.76 ± 0.3 | 2.3 ± 0.05 | > 100 |
| **40** |  | B | 0.083 ± 0.1 | 0.013 ± 0.006 | 0.049 ± 0.001 | > 100 |

Interestingly, an unsaturated chain as in compounds **33** and **40** showed increased potency as compared to the saturated chains of **25** and **39**. This may be due to the increased rigidity of the alkene linkage and a lower entropic penalty upon binding. As a matched pair to the previously reported compounds **32** and **34**, we selected potent and stable compound **33** and **40**, with the same change of adenosine base, Base A, to Base B and measure the antiviral activity. Compound **32** was previously reported to be inactive in an immunofluorescence antiviral assay with SARS-CoV-2 virus and A549/ACE2/TMPRSS2 cells, and **34** reported active with an EC_50_ value of 0.72 μM and no cellular toxicity.^27^ In our hands in an antiviral cytopathic effect assay with SARS-CoV-2 in Vero E6 cells, compound **32** had no antiviral activity, but compound **34** had significantly weaker activity than previously reported, with EC_50_ = 30 μM (versus 0.72 μM reported^27^), **Table 5**. One potential reason for this disparity may be attributed to the differing experimental conditions used in the two assays and requires further investigation. Compound **33**, showed some apparent antiviral potency (EC_50_ = 8.7 μM), however compound **33** was cytotoxic with a comparable CC_50_ = 21 μM. Therefore, consistent with the body of the literature, we observed no significant antiviral activity against SARS-CoV-2 for this class of compounds. We hypothesise that as the compounds bind in both the SAM and the RNA pocket of nsp14 and are therefore competitive with both substrates, in the cellular context there is a significant competition with intracellular SAM/SAH and the viral RNA, which is likely present at a higher concentration than used in the biochemical assay. The concentration of SAM used in the SARS-CoV-2 nsp14 assay is 1.0 µM and although measurements of SAM or SAH concentrations in Vero E6 cells haven’t been reported, others have reported intracellular concentrations of SAM of 354 µM in other cell lines.^30^ The concentration of the viral RNA substrate *in cellulo* and increased effective concentrations from the proximity of other nsp proteins involved in viral RNA production may also play a role that is further not captured in the biochemical assay. Further ADME data for compounds **32**-**40** are included in **Table S4**.

**Table 5.** Selected compounds with cellular potency. The apparent EC_50_ values in the cytopathic effect assay are shown, along with the toxicity effect CC_50_.

| Compound | TPSA Å<sup>2</sup> | CHI-logD | Solubility/<br>μM | MDCK<br>nm/sec | SARS-CoV-2<br>CPE assay |  |
| --- | --- | --- | --- | --- | --- | --- |
|  |  |  |  |  | EC<sub>50</sub><br>/ μM | CC<sub>50</sub><br>/ μM |
| **32** | 165 | 1.5 | 212.9 | 12 | >100 | >100 |
| **33** | 132 | 1.3 | 18.2 | 49 | 8.7 | 21 |
| **34** | 152 | 1.6 | 226.2 | 10 | 30 | >100 |
| **40** | 119 | 1.5 | 4.7 | 128 | >10 | >10 |

In the absence of robust antiviral activity, we decided to explore the selectivity against other viral MTase by biochemical assays. We selected one of the most potent analogues, compound **26**, for further biochemical profiling against several viral MTases across the coronavirus family, namely SARS-CoV-1 nsp14, MERS-CoV nsp14, HCoV-NL63 nsp14, HCoV-229E nsp14, SARS2-CoV-2 nsp16 and against Dengue virus NS5, **Table 6**. Consistent with the rest of the series, excellent activity was observed against alphacoronaviruses HCoV-NL63 and HCoV-229E, suggesting opportunities for this class of compounds against alphacoronaviruses in particular, if activity in cellular models of infection can be achieved. The most significant drop-off in activity was observed against MERS-CoV nsp14, with an IC_50_ of 0.73 μM, approximately a ten-fold reduction in activity as compared to SARS-CoV-2 nsp14. There is a high amino acid conservation between the ligand-binding site of MERS nsp14 and SARS-CoV-2 nsp14. As there are no MERS nsp14 structures published, we generated an AlphaFold model of MERS nsp14 to help visualise any mutations that may explain the potency drop-off observed. Whilst there are no differing residues which directly interact with the ligand, there are a couple in proximity of the adenosine binding site that may contribute; Ser369 (MERS Thr366) and Asp390 (MERS Pro386), **Figure S1**. The compound was also tested against the ribose 2’O-MTase nsp10/nsp16 of SARS-CoV-2 and NS5 MTase of dengue virus, where no inhibition was observed.

**Table 6.**
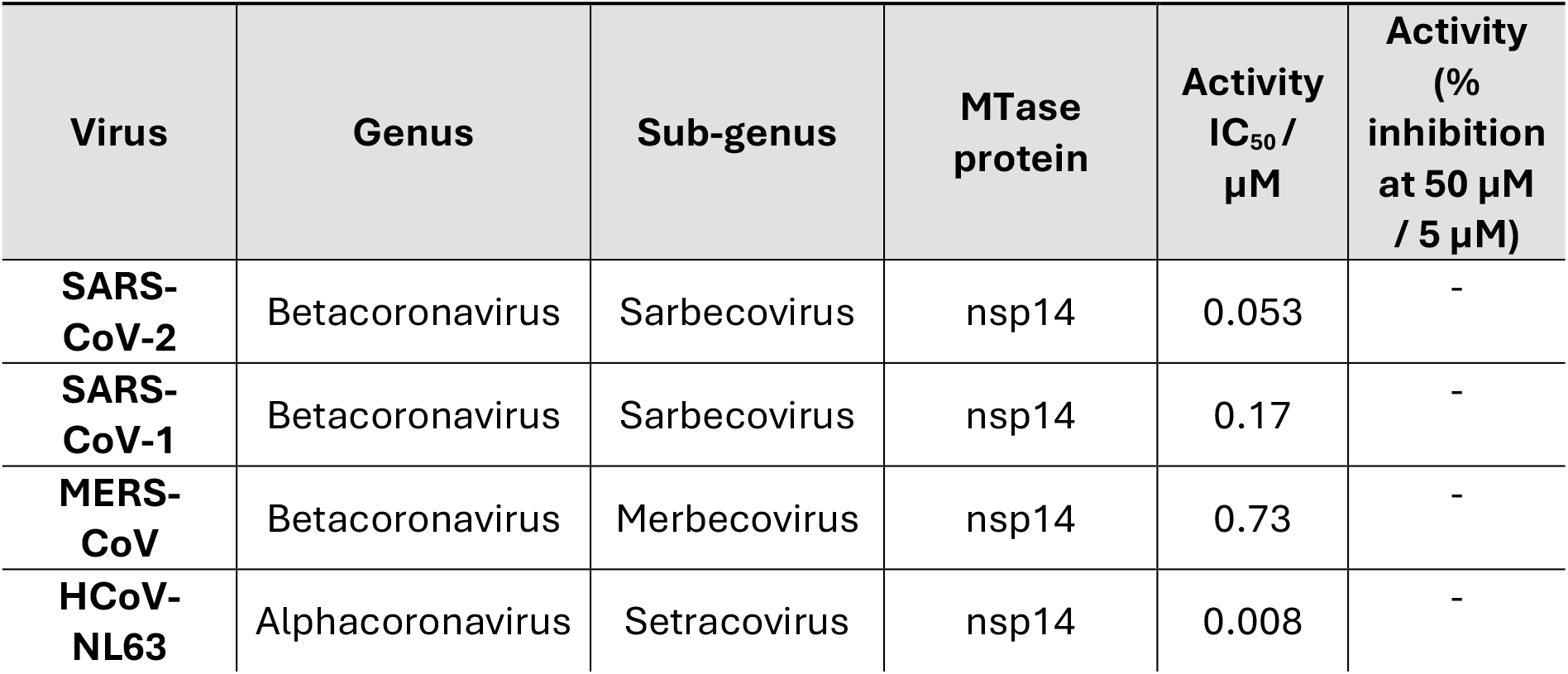
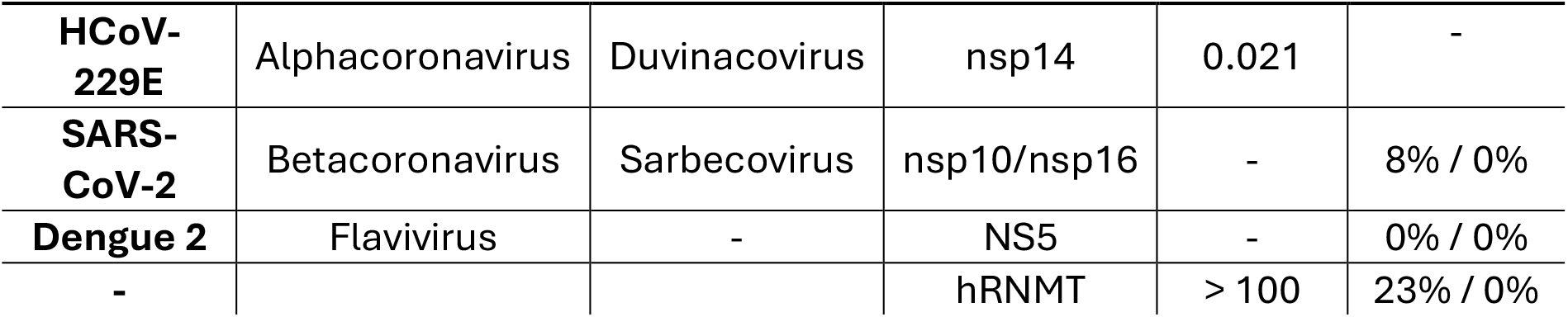
Profiling of compound **26** against various viral methyltransferases and human RNMT.

### Synthesis

**Scheme 1.**
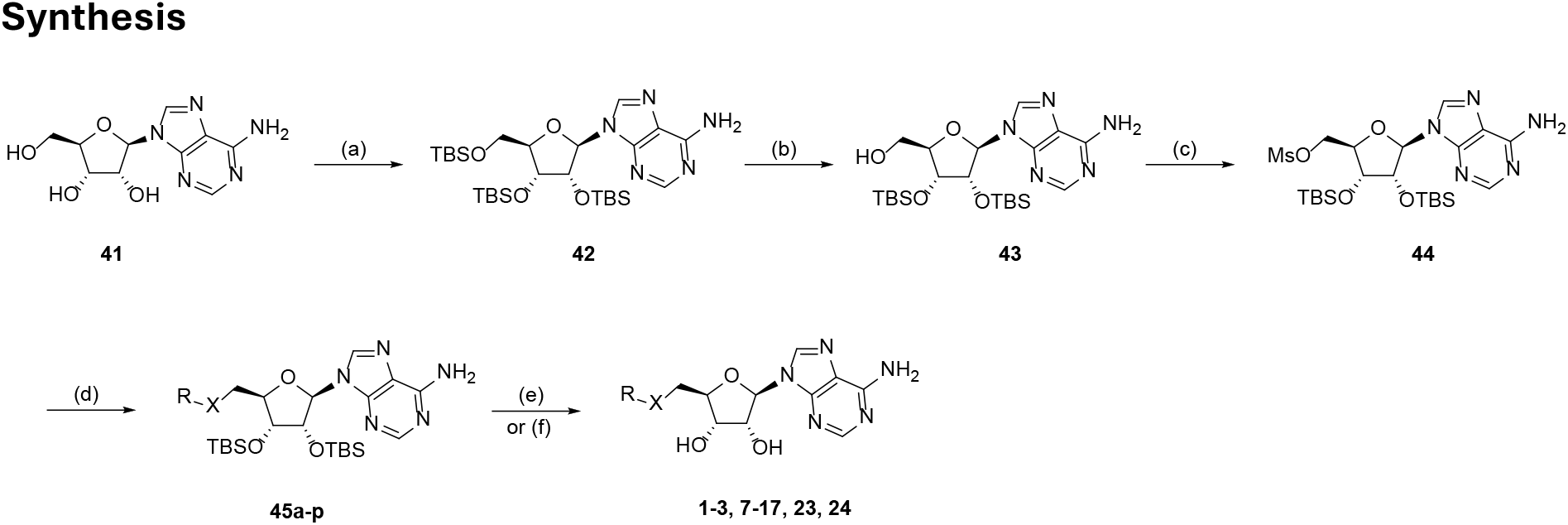
Synthetic route for compounds 1-3, 7-17, 23 and 24. Reagents and conditions: (a) TBS-Cl, 1H-imidazole, DMF, 25 °C, 16 h, 79%; (b) CCl_3_CO_2_H, THF, H_2_O, 0°C, 4 h, 61%; (c) Ms_2_O, TEA, THF, 25°C, 2 h, 17%; (d) R-XH, NaH, THF, 50°C, 16 h, 9-87%; (e) KF, MeOH, rt, 16 h, 7-69% or (f) Et_3_N-3HF, THF, rt, 16 h, 17- 42%.

The synthesis of compounds **1**-**3, 7**-**17** (**Table 1**) and **23, 24** (**Table 2**) was commenced with commercially available adenosine **41**. Protection of the free hydroxyl moieties with TBS-Cl in the presence of imidazole resulted in fully protected nucleoside **42** in 48% yield over two steps. Selective deprotection of the secondary alcohol was achieved using trichloroacetic acid and subsequent mesylation lead to key intermediate **44**. Activated **44** was then reacted with a variety of either commercially available or novel thiols (SI) and through nucleophilic substitution afforded thioethers **45a**-**p**. Finally, deprotection of the TBS-protected hydroxyl groups in the presence of KF or Et_3_N.3HF lead to desired thioethers **1-3** and **7**-**17**.

Mono or dimethylation of intermediate **45a**, followed by deprotection lead to compounds **18** and **19** (SI). As of compounds **4**-**6** (**Table 1**) and **20**-**22** (**Table 2**) bearing different secondary amines on the base region a similar synthetic pathway was explored as outlined in the SI.

Matched pair analogues **23** and **24** where the thioether was replaced with an ether or an amine linker moiety followed the same synthetic route (**Scheme 1**). Key intermediate **44** was reacted in these occasions with the corresponding hydoxybenzothiazole or benzothiazolamine resulting after deprotection in compounds **23** and **24**.

**Scheme 2.**
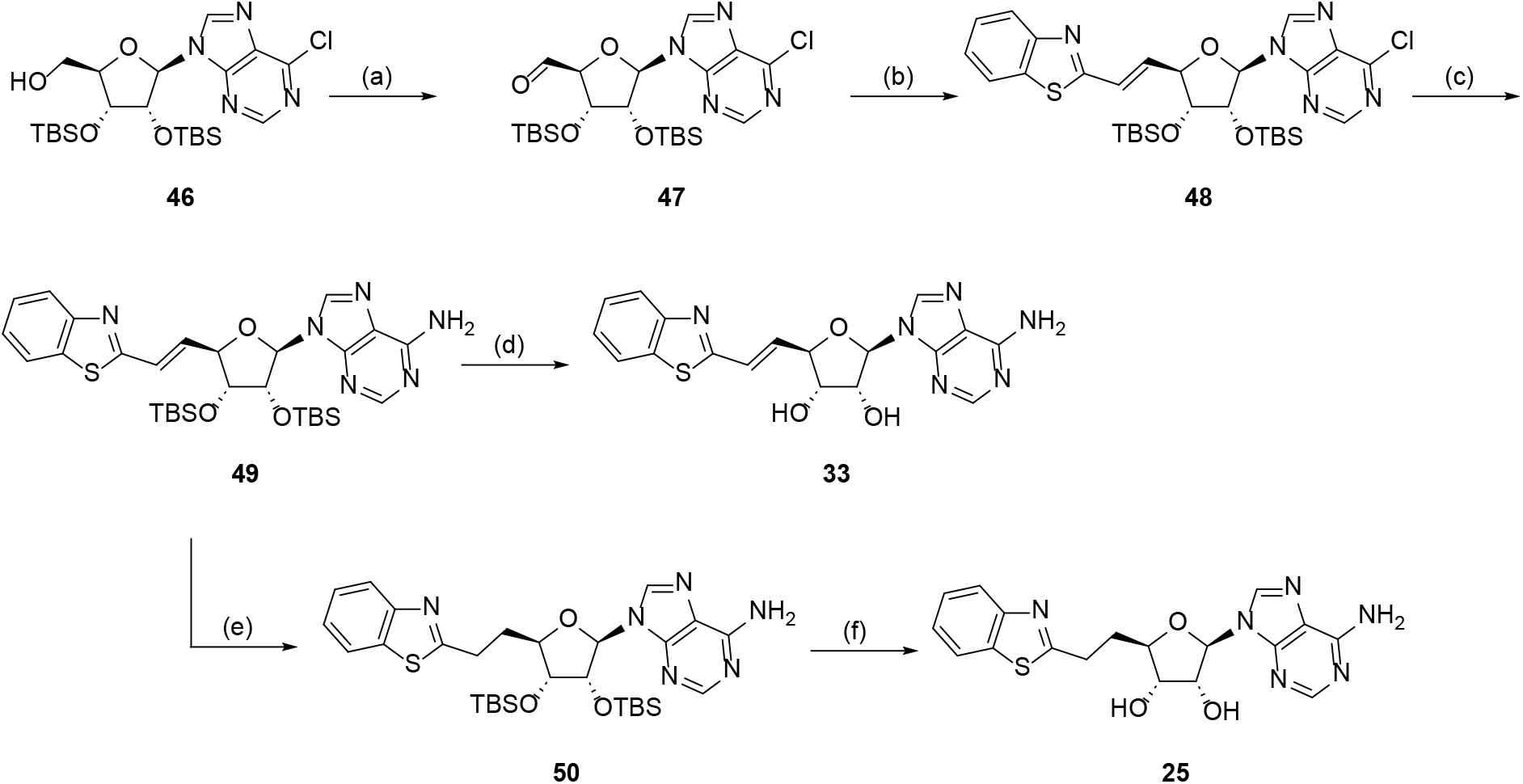
Synthetic route for compounds 25 and 33. (a) (COCl)_2_, DMSO, TEA, DCM, -60 °C, 3 h, 67%; (b)(benzo[d]thiazol-2-ylmethyl)triphenylphosphonium chloride, KHMDS, THF, -78 °C, 2 h, 87%; (c) NH_3_/THF,-70-80 °C, 16 h, 51%; (d) KF, DMF, 60 °C, 16 h, 57%; (e) H_2_, Pd/C, THF, rt, 16 h, 79%; (f) KF, DMF, 60 °C, 16h, 35%.

The synthesis of compounds **25** and **33**, where the backbone is linked through an ethyl or ethylene moiety to the nucleotide is outlined in Scheme 2. Chlorinated derivate **46** (SI) was oxidised under Swern conditions to the corresponding aldehyde **47** in 67% yield. Subsequently a Wittig reaction using the relevant phosphonium ylide allows the insertion of the double bond and isolation of key intermediate **49**. In the first instance **49** was deprotected in the presence of potassium fluoride to result in analogue **33**. Whereas the alkene moiety of **49** was also reduced and further deprotected to allow the formation of the ethyl-linked analogue **25**.

**Scheme 3.**
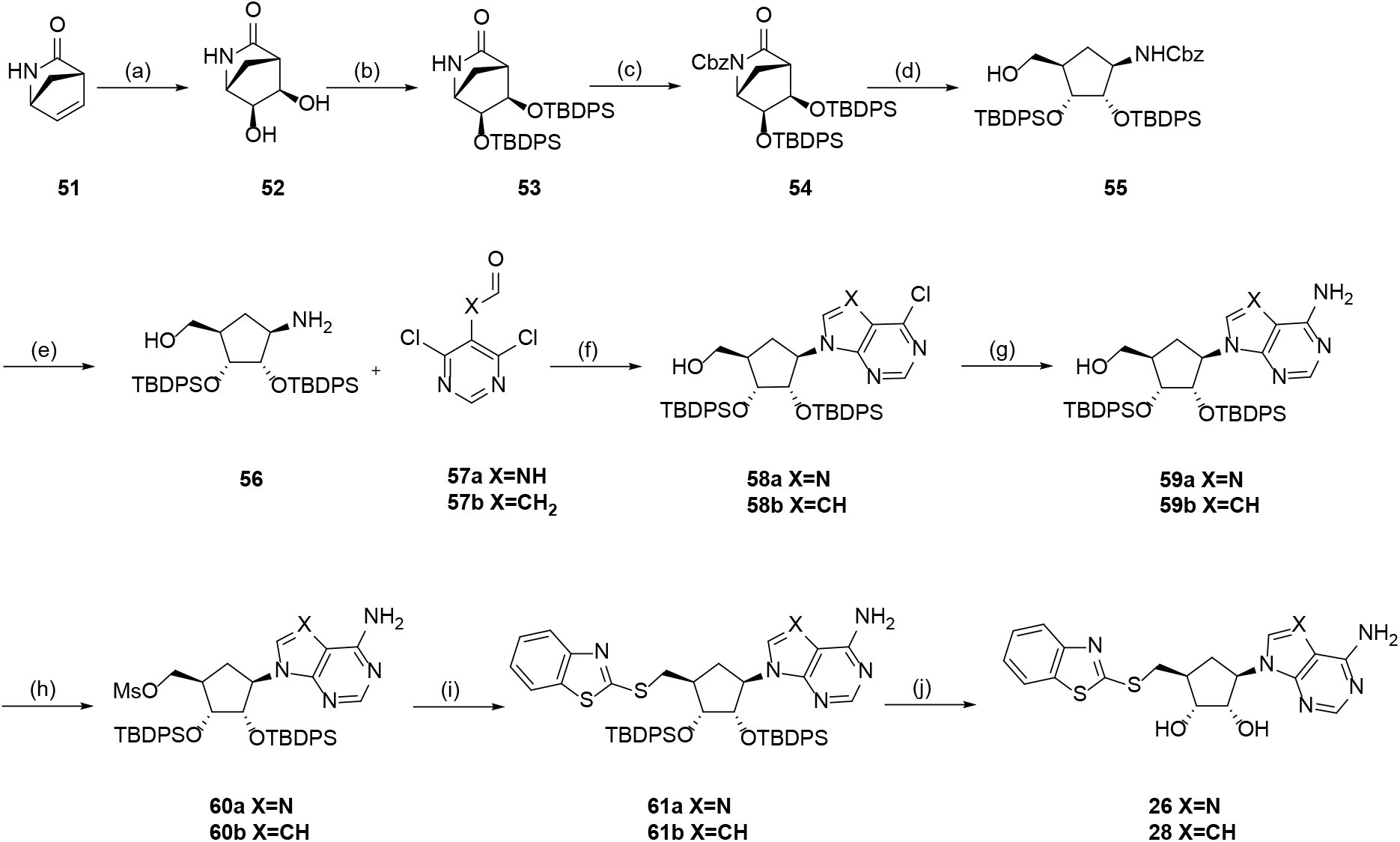
Synthetic Route for Compounds 26 and 28. Reagents and conditions (a) K_2_OsO_4_, NMO, THF, t-BuOH, H_2_O, rt, 16 h, 54%; (b) TBDPSCl, 1H-imidazole, DMF, 25 °C, 24 h, 63%; (c) CbzCl, LiHMDS, THF, - 70 °C, 30 min, 92%; (d) NaBH_4_, THF, MeOH, 0 °C, 2 h, 97%; (e) H_2_, Pd/C, THF, rt, 16 h, 82%; (f) TEA, n-BuOH,130 °C, 24 h, 78%, 77%; (g) NH_3_/MeOH, 90 °C, 16 h, 77%, 92%; (h) Ms_2_O, TEA, THF, 20 °C, 16 h, 72%, 90%;(i) sodium benzo[d]thiazole-2-thiolate, DMF, 50 °C, 16 h, 55%, 50%; (j) KF, MeOH, 50 °C, 16 h, 43%, 55%.

The corresponding analogues where the deoxyribose sugar moiety has been replaced with a cyclopentyl modified nucleotide followed a similar literature reported procedure as depicted in **Scheme 3**.^31^ Dihydroxylation of bicyclic, Vince lactam **51** with catalytic osmium tetroxide in the presence of NMO afforded *cis*-hydroxylated compound **52** in 54% yield.^32^ Sequential protection of the free hydroxyl moieties with TBDPS, followed by Cbz-protection of the lactam lead to compound **54**. Reductive cleavage of the lactam using NaBH_4_ and selective deprotection of the amine under reducing conditions allowed the isolation of key intermediate **56**. Incorporation of the base region was accomplished by employing a literature procedure in which *N*-(4,6-dichloropyrimidin-5-yl)formamide **57a** reacted with **56** under basic conditions leading to **58a**^33^. Nucleophilic displacement of the C-6 chlorine, followed by the reaction of the formyl moiety with the amine results in the formation of the purine base. Utilisation of acetaldehyde **57b** instead results in **58b** bearing a pyrrolopyrimidine ring. Chlorinated analogues **59a** and **59b** were then reacted under SN_Ar_ conditions with NH_3_ to afford **60a** and **60b**. The subsequent steps follow the same synthetic pathway as outlined in **Scheme 1**. Activation of the free hydroxyl moiety by formation of the mesylate, followed by nucleophilic substitution with the relevant thiol and finally deprotection of the hydroxyl moieties leads to **26** and **28**.

**Scheme 4.**
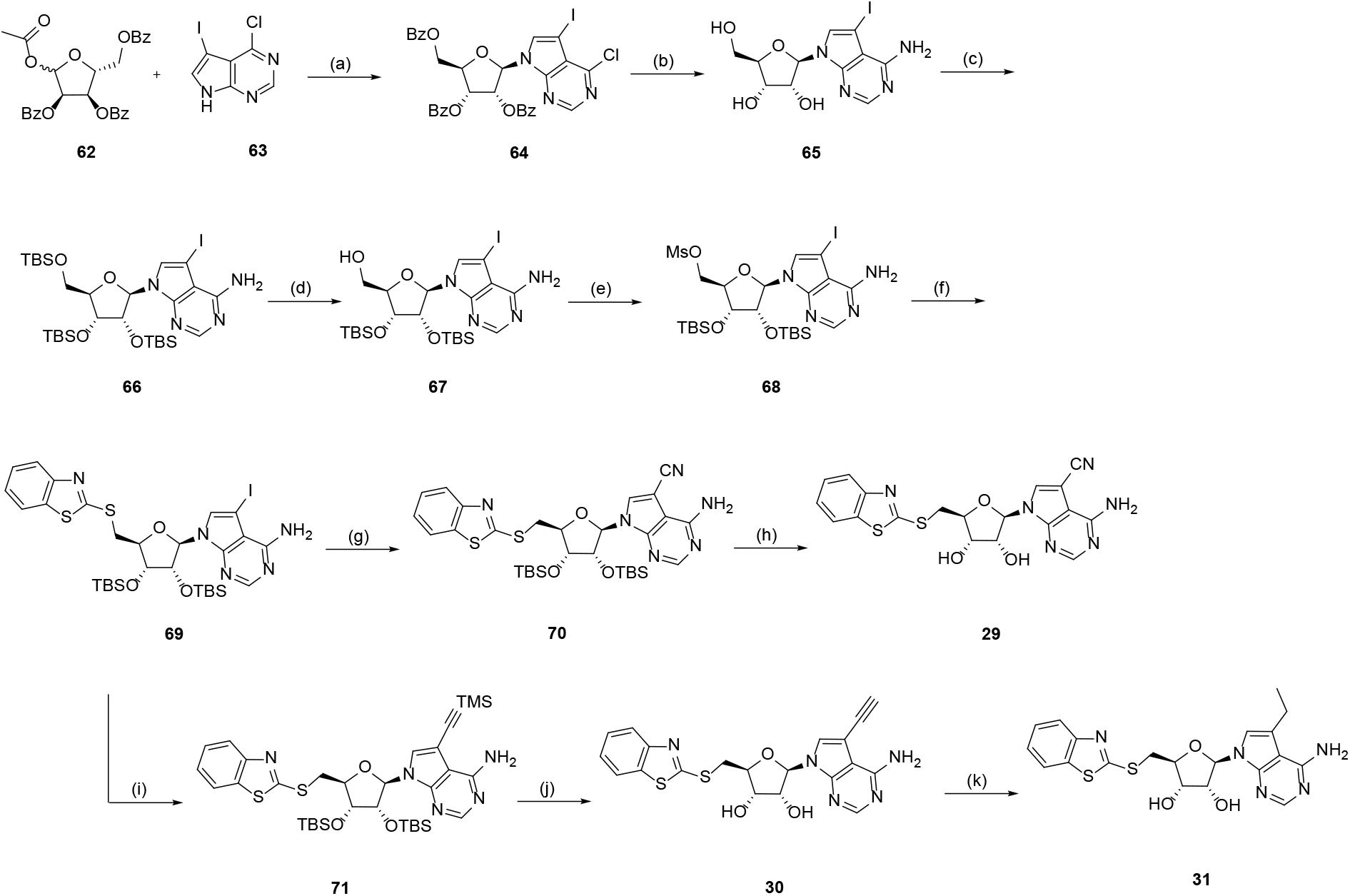
ynthetic route for compounds 29-31. Reagents and conditions: (a) BSA, TMSOTf, MeCN, 25-80 °C, 1.5 h, 49%; (b) NH_3_/MeOH, 90 °C, 16 h, 98%; (c) TBS-Cl, 1H-imidazole, DMF, 25 °C, 16 h, 30%; (d)Cl_3_CCO_2_H, THF/H2O, 0 °C, 1 h, 84%; (e) Ms_2_O, TEA, THF, 0-25 °C, 2 h, 89%; (f) sodium benzo[d]thiazole-2-thiolate, DMF, 50 °C, 12 h, 95%; (g) Pd(PPh_3_)_4_, Zn(CN)_2_, DMF, 90 °C, 16 h, 92%; (h) KF, MeOH, 60 °C, 16 h,12%; (i) TMS-acetylene, Pd(PPh_3_)_4_, DMF, 60 °C, 2 h, 62%; (j) KF, MeOH, 60 °C, 16 h, 74%; (k) H_2_, Pd/C, rt, 2h, 18%.

The synthesis of target compounds substituted at the 7-position of the modified nucleobase **29**-**31** followed a slightly amended route previously reported for structurally similar analogues^24, 28^. 6-Chloro-7-iodo-7-deazapurine **63** was coupled with protected ribose sugar **62** under glycosylation conditions in the presence of BSA and TMSOTf to give **64**^34^. Protecting group swapping allowed the selective deprotection of the primary alcohol to afford nucleoside **67** in 24% over three steps. Transformation of the free hydroxyl moiety to the activated mesylate, followed by nucleophilic displacement using the thiol of interest results in intermediate **69**. Subsequently Pd-catalysed cross-coupling conditions allow either a nitrile or an acetylene moiety to be installed by treating with Zn(CN)_2_ or TMS-acetylene to afford **70** and **71** respectively. Finally, deprotection leads to desired analogues **29** and **30**. In the case, of ethylene substituted **31** it was synthesised by reducing the acetylene moiety of compound **30**.

## Conclusion

We report a medicinal chemistry programme against SARS-CoV-2, focussing on a series of bi-substrate SAM-like molecules which occupy both the SAM/SAH and the RNA binding pocket of SARS-CoV-2 nsp14. Throughout compound development, we evaluated inhibitory effects of the analogues on the MTase activity of nsp14 from betacoronavirus SARS-CoV-2 and alphacoronaviruses HCoV-NL63 and HCoV-229E. Interestingly, activity of the compounds was largely comparable across these three nsp14 proteins tested. Broader screening of one series exemplar in biochemical assays defined some boundaries of the MTase inhibition, showing no activity against SARS-CoV-2 nsp16 or human RNMT, nor against Dengue virus NS5 MTase. Despite excellent stability and permeability for many of these compounds, translation of biochemical activity to antiviral activity remains a challenge to overcome for this class of nsp14 inhibitors.

## Acknowledgements

This work was supported in part by the Gates Foundation [INV-016131]. The conclusions and opinions expressed in this work are those of the author(s) alone and shall not be attributed to the Foundation. Under the grant conditions of the Foundation, a Creative Commons Attribution 4.0 License has already been assigned to the Author Accepted Manuscript version that might arise from this submission. Please note works submitted as a preprint have not undergone a peer review process. This work was also kindly supported by the Corona Accelerated R&D in Europe (CARE) project funded by the Innovative Medicines Initiative two Joint Undertaking (JU) under grant agreement no. 101005077, the Fondation pour la Recherche Médicale (Aide aux Équipes), the Swift COronavirus therapeutics REsponse (SCORE) project (European Union Horizon 2020 Research and Innovation Program Grant Agreement 101003627). This work was supported by a grant from National Science Centre UMO-2022/45/B/NZ7/04269.

The authors kindly thank Vincent Postis, Marilyn Paul, Karolina Wrobel, Jennifer Riley and Hali Joji for useful discussions, Proteros Biostructures GmbH for solving the nsp14 crystal structures, and WuXi AppTec and Nuvisan for chemical synthesis.

## Materials and Methods

### Protein expression and purification

For biochemical assays, SARS-CoV-2 nsp14 protein was expressed and purified as described previously by Pearson *et al*.^19^ Genes encoding nsp14 from HCoV-NL63 (sequence encoding polyprotein 1ab amino acids 5568-6085) and HCoV-229E (sequence encoding polyprotein 1ab 5593 – 6110) were synthesized by Genscript.

Sequences were codon optimised for expression in *E*.*coli*, with 5’ BamHI and 3’ NotI and cloned in fusion with a Tobacco Etch Virus (TEV) cleavable N-terminal His in the first MCS of pETDuet1 vector.

Nsp14 from HCoV-NL63 and HCoV-229E were expressed in *Escherichia coli* BL21 (DE3) Rosetta 2 (Merck) using TB media in Ultra yield Flask (Thomson Scientific) at 250rpm at 37 degrees. Cells were grown to an OD_600_ of ∼1.2-1.4 prior to induction with 1 mM IPTG. Cells were further incubated for 18 hours at 20 degrees, prior to harvesting by centrifugation.

Cells were subsequently lysed by sonication in 50 mM Tris, 500 mM NaCl, 5 mM MgCl2, 5 mM BME, 10% glycerol and 1 0mM imidazole pH 8.5 prior to being purified by batch purification using HisPur Ni NTA resin (Thermo Scientific) and eluted using 50 mM Tris, 250 mM NaCl, 5 mM MgCl_2_, 5 mM BME, 10% glycerol and 100 mM imidazole pH 8.5. Proteins were then buffer exchanged into 50 mM Tris, 250 mM NaCl, 5 mM MgCl_2_, 5 mM BME, 10% glycerol and 10 mM imidazole pH 8.5 prior to gel filtration using a Superdex 200 16/60 column (Cytiva) equilibrated in 50 mM Tris pH 8.5, 150 mM NaCl, 5 mM MgCl_2_ and 1 mM TCEP. The protein was eluted in 50 mM Tris pH 8.5, 150 mM NaCl, 5 mM MgCl_2_ and 1 mM TCEP prior to concentration to ∼ 1 mg/mL.

Expression and purification of the SARS-CoV-2 nsp14 protein for the X-ray crystallography were performed following the methods outlined described by Czarna *et al*.^35^

SARS-CoV-2 nsp16,^36^ SARS-CoV-1 nsp14,^12^ MERS nsp14^37^ and dengue virus NS5 protein^38^ were expressed, purified and assayed as previously described. Human RNMT was purified, expressed and assays were run according to Gray et al.^39^

### SARS-CoV-2 nsp14 Protein crystallization

Crystals of the complex of nsp14 with compounds **1, 5, 6, 18, 26** and **27** were prepared by back-soaking of co-crystals generated with a proprietary ligand by co-crystallization. X-ray diffraction data were collected at either the Diamond Light Source (compounds **1** and **5**; DLS, Oxford, England) or the Swiss Light Source (compounds **6, 18, 26** and **27**; SLS, Villigen, Switzerland) using cryogenic conditions. The diffraction data was indexed and integrated using AutoPROC^40^, XDS^41^ and the data was scaled in AutoPROC^40^, AIMLESS^42^. Following steps were performed in Phenix.^43^ The structures of nsp14 were solved by molecular replacement using PHASER^44^ and previously solved nsp14 structure. Models were refined by interchanging cycles of automated refinement using phenix.refine^45^ and manual building in Coot^46^. Restraints for the inhibitor were created in GRADE Web Server. Ligand-bound structures of nsp14 with compounds **1, 5, 6, 18, 26** and **27** have been uploaded to the PDB repository and the PDB codes are respectively; 9S0M, 9SAJ, 9SAK, 9SAL, 9SAM and 9SAN.

### Methyltransferase biochemical assays

#### Determination of the MTase activity of SARS-CoV-2, HCoV-NL63 and HCoV-229E nsp14 by RapidFire assay

SARS-CoV-2 nsp14 activity and inhibition by compounds was assessed using the protocol previously described by Pearson et al19, which was adjusted for the respective HCoV-NL63 and HCoV-229E proteins to account for differences in activity and substrate K_M_. For each protein, the K_M_ of both the SAM and RNA cap substrates were established, and the initial concentration of each substrate was at the K_M_ concentration in the assay. Repeat n-values for each assay are described in Table S5.

| Protein | Protein conc. / nM | Incubation time | [SAM] substrate | [CAP] substrate |
| --- | --- | --- | --- | --- |
| SARS-CoV-2 nsp14 | 5 nM | 1 hour | 1.0 μM | 0.7 μM |
| HCoV-NL63 nsp14 | 10 nM | 3 hours | 3.5 μM | 1.5 μM |
| HCoV-229E nsp14 | 10 nM | 2 hours | 5.0 μM | 3.0 μM |

#### Determination of the MTase activity of SARS-CoV-2 nsp14 by MTase-Glo assay

Inhibition of MTase activity was also evaluated using an MTase-Glo™ bioluminescence-based assay kit (Promega, Cat. #V7601). The assay was performed in a 10µL final reaction volume containing 5 nM nsp14, 0.7 µM cap analogue ((5′)Gppp(5′)G sodium salt, S1407L, New England Biolabs) and 1 µM SAM. The reaction buffer was 20 mM Tris-HCl, 50 mM NaCl, 3 mM MgCl_2_, 0.1 mg/mL BSA, 0.005% NP40 substitute and 1 mM TCEP. The assay was performed at room temperature in 384-well plates (Greiner #784904, 384 well Microplate, F-bottom, white, nonbinding, low volume). After allowing the reaction to proceed for 1 hour, the product S-adenosyl homocysteine (SAH) was converted to ADP by the addition of 2.5µL MTase-Glo reagent. After an additional 30 minute incubation at room temperature, 12.5µL of MTase-Glo detection buffer was added and further incubated in the dark for 30 minutes. Finally, the bioluminescence signal was read on an BMG PHERAstar FS.

#### Determination of the MTase activity of SARS-CoV-1 nsp14, MERS-CoV nsp14, SARS-CoV-2 nsp16 and dengue NS5 by filter binding assay (FBA)

The FBA MTase assays were carried out in reaction mixture [40 mM Tris-HCl (pH 8.0), 1 mM DTT, 1 mM MgCl_2_, 1.9 μM SAM, and 0.1 μM ^3^H-SAM (Perkin Elmer)] in the presence of 0.7 μM GpppAC_4_ synthetic RNA^47^ and the MTase (10 nM). Briefly, the MTases were first mixed with the compound suspended in 50% DMSO (2.5% final DMSO) before the addition of RNA substrate and SAM and then incubated at 30 °C. Reactions mixtures were stopped after 60 min by their 10-fold dilution in ice-cold water. Samples were transferred to diethylaminoethyl (DEAE) filtermat (Perkin Elmer) using a Filtermat Harvester (Packard Instruments). The RNA-retaining mats were washed twice with 10 mM ammonium formate pH 8.0, twice with water and once with ethanol. They were soaked with scintillation fluid (Perkin Elmer), and ^3^H-methyl transfer to the RNA substrates was determined using a Wallac MicroBeta TriLux liquid scintillation counter (Perkin Elmer). For IC_50_ measurements, values were normalized and fitted with Prism (GraphPad software) using the following equation: *Y* = 100/[1 + ((*X*/IC_50_)^Hillslope)]. IC_50_ is defined as the inhibitory compound concentration that causes 50% reduction in enzyme activity.

#### SARS-CoV-2 CPE reduction assay

Vero E6 cells were maintained in Dulbecco’s modified Eagle’s medium (DMEM; Lonza), supplemented with 8% fetal calf serum (FCS; Bodinco), 2 mM L-glutamine, 100 IU/ml of penicillin, and 100 µg/ml of streptomycin (Sigma-Aldrich). For CPE-based assays, cells were seeded in 96-well cell culture plates at a density of 5*10^3^ cells per well the day before. Cells were incubated with 2-fold serial dilutions of the test compound for 1-2 hours. Subsequently, cells either were mock infected (for analysis of cytotoxicity of the compound) or infected with SARS-CoV-2 (isolate Leiden-0002, derived from a nasopharyngeal sample in March 2020; GenBank accession nr. MT510999) at an MOI of 0.015. Cell viability was assessed 4 days post-infection by MTS assay using a CellTiter 96 aqueous nonradioactive cell proliferation kit (Promega), and absorption was measured using an EnVision multilabel plate reader (PerkinElmer). Every compound concentration was tested in quadruplicates for both toxicity and antiviral effects, viability was normalized to untreated mock-infected control cells, and the average ± standard deviation was calculated. All experiments with infectious SARS-CoV-2 were performed in a biosafety level 3 facility at the Leiden University Medical Center.

#### Determination of Chromatographic Hydrophobicity Index logD (CHI logD)

CHI logD measurements were determined at pH 7.4 using a 5 μL aliquot from a 10 mM DMSO stock which was diluted to 250 μM by adding 195 μL of MeCN:H2O (1:1, v/v) and mixing in a 96-well plate. An aliquot (2 μL) was then injected onto an Acquity UPLC BEH C18 column (Waters, Wilmslow, UK, 2.1 x 50 mm) with 1.7 µm particle size. The mobile phase A (MPA) was 10 mM ammonium acetate pH 7.4 and mobile Phase B (MPB) was acetonitrile. The HPLC system used was a Shimadzu Nexera X2 HPLC, where a linear gradient from 95% to 5% MPA was applied over 4 min and held there for a further 0.5 min. At 4.51 min, the MPA was switched back to 95% and held there up to 5.51 min to re-equilibrate. The flow rate was 0.6 mL/min with detection at 254 nm UV and ambient temperature. Using the retention time of ten calibration compounds (theophylline, phenyltetrazole, benzimidazole, colchicine, phenyltheophylline, acetophenone, indole, propiophenone, butyrophenone, valerophenone) and uracil (unretained component), prepared at 10 μg/ml in MeCN:water (1/1, v/v), the retention factor (k) for each was calculated using the following relationship:

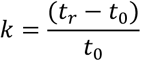

Where tr is the retention time of the calibration compound and t_0_ is the retention time of the unretained component. This covers the CHI log D range from -0.51 to 3.78.

The k value was plotted against literature CHI values of each component and a linear regression performed to produce a line of best fit through the data points.^48^ Using the calibration line, the CHI and CHI log D values of the calibration compounds and test compounds was calculated as follows:

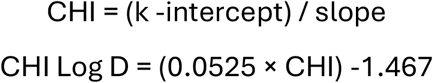

#### Intrinsic clearance (Cl_*i*_)

Test compound (0.5 µM) was incubated with female CD1 mouse liver microsomes or mixed gender human liver microsomes (Xenotech™; 0.5 mg/mL 50 mM potassium phosphate buffer, pH 7.4) and the reaction started with addition of excess NADPH (8 mg/mL 50 mM potassium phosphate buffer, pH7.4). Immediately, at time zero, then at 3, 6, 9, 15 and 30 minutes an aliquot (50 µL) of the incubation mixture was removed and mixed with acetonitrile (100 µL) to stop the reaction. Internal standard was added to all samples, the samples centrifuged to sediment precipitated protein and the plates then sealed prior to UPLC-MS/MS analysis eg. (Xevo TQ-S Micro, Waters ™). XLfit (IDBS, UK) was used to calculate the exponential decay and consequently the rate constant (k) from the ratio of peak area of test compound to internal standard at each timepoint. The rate of intrinsic clearance (CLi) of each test compound was then calculated using the following calculation:

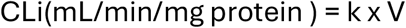

Where V (mL/mg protein) = incubation volume (0.5mL)/mg protein added (0.25 mg protein) Verapamil (0.5µM) was used as a positive control to confirm acceptable assay performance. A scaling factor of 48 mg mouse liver microsomal protein per g liver is applied.

#### Solubility assessment

Test compounds were dissolved in DMSO to obtain 10 mM solutions. Solubility test samples were prepared by adding 5 µL of the 10 mM solution to 195 µL of phosphate buffered saline, pH 7.4 (Sigma-Aldrich, Cat no. P4417, made as per manufacturer’s instructions). This solution was then mixed for 24 hours (rotary mixing, 900 rpm, 25°C), protected from light. After mixing, the solubility test samples were filtered using a proprietary filter (Millipore Multiscreen HTS filter, 96-well format) to remove any undissolved material. Samples were drawn through the filter using vacuum. The filtrate from the above was analysed for dissolved drug compound using a truncated UHPLC methodology. A Shimadzu Nexera X2 UHPLC system equipped with a Shimadzu LCMS 2020 single quadrupole mass spectrometer was used. A reversed-phase methodology was used with a simple formic acid gradient elution. A calibration solution was prepared as follows: The same 10 mM compound solution used to prepare the solubility test sample was diluted in DMSO to give a 500 µM solution. This solution was further diluted with 50:50 acetonitrile:water to obtain a 50 µM solution. Aliquots (0.2, 2.0 and 5.0 µL) of this 50 µM solution were injected onto the UHPLC system and the areas of the resultant peaks integrated to produce a calibration line. Aliquots of the test sample filtrate (0.4 and5.0 µL) were injected onto the UHPLC system and the resultant peak areas for any peaks corresponding to the test compound determined and quantified using the calibration line (the injection volume that gave a peak area closest to the calibrated range was used for determining solubility). The mass spectrometer was not used for quantitative purposes but was solely used to check the identity (nominal mass) of the quantified peak in UV.

#### MDCK passive permeability measurement

MDCK-II cells (ECACC) were maintained in DMEM (Gibco Cat: 61965-026) supplemented with 1% penicillin/streptomycin and 10% FCS. Cells were seeded onto individual transwell ‘Thincerts’ (Greiner, Cat 662610) at a density of 35,000 cells/well. Cells were grown at 37 °C, 5% CO_2_ for 3 days. On day 4, growth media was replaced with fresh media and incubated for 1 hour. After incubation, media was replaced with Dulbecco’s PBS (Gibco, 14287-080) and cell inserts were incubated for a further 1 hour. Dosing solutions containing 3 µM test compound and 10 µM Lucifer Yellow (1% DMSO) were prepared. 1.2 mL of PBS (1% DMSO) was added to wells of a 24-well cell culture plate (Corning, Cat 353504). 0.35 mL of dosing solution was added in duplicate to the apical side of the transwell and transwells transferred into the receiver plate solutions. Transwell plates were incubated for 1 hour and inserts were then transferred to an empty plate to prevent any further permeation of compound. 100 µL of solution from donor wells was transferred to a 96 well plate, alongside 100 µL of dosing solution. 150 µL of acetonitrile containing internal standard (eg 100 ng/mL sulfadimethoxine) was then added to all samples prior to analysis by LC-MS/MS. Bupropion (positive controls) and atenolol (negative control) were run alongside test compounds. To confirm monolayer integrity, a further 100 µL from each compartment was added to the 96 well F-bottomed microtitre plate containing the Lucifer Yellow standard curve for fluorescence determination of Lucifer Yellow concentrations. Papp (apparent permeability) values were calculated using the following equation:

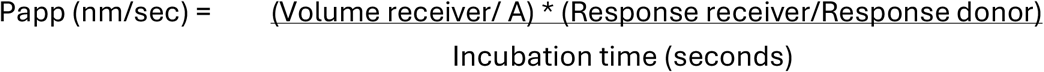

