## Supplemental Information for "Crystallographic characterisation and development of bi-substrate inhibitors of coronavirus nsp14 methyltransferase"

### Supporting Information

### 1. Chemical Synthesis

#### 1.1 Solvents and Reagents

All starting materials and solvents were obtained from standard chemical suppliers. Dry solvents were purchased in sure sealed bottles stored over molecular sieves. Organic solutions were routinely dried over anhydrous sodium sulfate or magnesium sulfate.

#### 1.2 Chromatography

Thin-layer chromatography (TLC) was performed on pre-coated silica gel plates (Kieselgel 60 F254, BDH). Visualization of the developed chromatogram was achieved by using fluorescence quenching and/or by staining with potassium permanganate. Column chromatography purification was performed on prepacked silica gel (230–400 mesh, 40–63 µm) cartridges. Reverse column chromatography was performed on SWPAFLASH® SW040 Bonded Spherical C18 (20-45µm, 100 Å). All compounds are >95% pure by HPLC analysis.

#### 1.3 NMR Spectroscopy

All $^{1}H$ NMR spectra were recorded on a Brucker Avance III 400 MHz. Chemical shifts (δ) are expressed in ppm recorded using the residual undeuterated solvent as the internal reference in all cases. Signal splitting patterns are described as singlet (s), doublet (d), triplet (t), quartet (q), multiplet (m), broad (br), or a combination. Coupling constants (J) are quoted to the nearest 0.1 Hz.

#### 1.4 Reverse Phase HPLC conditions for the LCMS Analytical Methods

**Method 1:** Agilent 1200\G6110A Kinetex EVO C18 2.1 × 30 mm, 5 µm at 40 ℃; Mobile Phase: A: 0.0375% TFA in water (v/v); B: 0.01875% TFA in Acetonitrile; flow rate held at 1.5 mL/min; eluted with the mobile phase over 1.50 min employing UV detection at 220&254 nm. Gradient information: 0.01-0.80 min, ramped from 95% A-5% B to 5% A-95% B; 0.80-1.20 min, held at 5% A-95% B; 1.20-1.21 min, returned to 95% A-5% B, 1.21-1.5 min, held at 95% A-5% B.

**Method 2:** SHIMADZU LCMS-2020 Kinetex EVO C18 2.1 × 30 mm, 5 µm at 40 ℃; Mobile Phase: A: 0.025% $NH_3 \cdot H_2O$ in Water (v/v); B: Acetonitrile; flow rate held at 1.5 mL/min; eluted with the mobile phase over 1.55 min employing UV detection at 220&254 nm. Gradient information: 0-0.80 min, ramped from 95% A-5% B to 5% A-95% B; 0.80-1.20 min, held at 5% A-95% B; 1.20-1.21 min, returned to 95% A-5% B, 1.21-1.55 min, held at 95% A-5% B.

**Method 3:** SHIMADZU LCMS-2020 Kinetex EVO C18 2.1 × 30 mm, 5 µm at 50 ℃; Mobile Phase: A: 0.0375% TFA in water (v/v); B: 0.01875% TFA in Acetonitrile; flow rate held at 1.5 mL/min; eluted with the mobile phase over 1.55 min employing UV detection at 220&254 nm. Gradient information: 0-0.80 min, ramped from 95% A-5% B to 5% A-95% B; 0.80-1.20 min, held at 5%

A-95% B; 1.20-1.21 min, returned to 95% A-5% B, 1.21-1.55 min, held at 95% A-5% B.

**Method 4:** SHIMADZU LCMS-2020 Kinetex EVO C18 2.1 × 30 mm, 5 µm at 50℃; Mobile Phase: A: 0.0375% TFA in water (v/v); B: 0.01875% TFA in Acetonitrile; flow rate held at 1.5 mL/min; eluted with the mobile phase over 1.55 min employing UV detection at 220&254 nm. Gradient information: 0-0.80 min, ramped from 100% A-0% B to 40% A-60% B; 0.80-1.20 min, held at 40% A-60% B; 1.20-1.21 min, returned to 100% A-0% B, 1.21-1.55 min, held at 100% A-0% B.

**Method 5:** SHIMADZU LCMS-2020 Kinetex EVO C18 2.1 × 30 mm, 5 µm at 40℃; Mobile Phase: A: 0.025% $NH_3 \cdot H_2O$ in Water (v/v); B: Acetonitrile; flow rate held at 1.0 mL/min; eluted with the mobile phase over 2.00 min employing UV detection at 220&254 nm. Gradient information: 0-1.20 min, ramped from 100% A-0% B to 40% A-60% B; 1.20-1.60 min, held at 40% A-60% B; 1.60-1.61 min, returned to 100% A-0% B, 1.61-2.00 min, held at 100% A-0% B.

**Method 6:** Agilent 1200\G6110A Kinetex EVO C18 2.1 × 30 mm, 5 µm at 50℃; Mobile Phase: A: 0.0375% TFA in water (v/v); B: 0.01875% TFA in Acetonitrile; flow rate held at 1.5 mL/min; eluted with the mobile phase over 1.50 min employing UV detection at 220&254 nm. Gradient information: 0.01-0.80 min, ramped from 100% A-0% B to 40% A-60% B; 0.80-1.20 min, held at 40% A-60% B; 1.20-1.21 min, returned to 100% A-0% B, 1.21-1.55 min, held at 100% A-0% B.

**Method 7:** Agilent 1200\G6110A XBridge C18 2.1 × 50 mm, 5 µm at 50℃; Mobile Phase: A: 0.025% $NH_3 \cdot H_2O$ in water (v/v); B: Acetonitrile; flow rate held at 1.0 mL/min; eluted with the mobile phase over 2.00 min employing UV detection at 220&254 nm. Gradient information: 0-1.20 min, ramped from 100% A-0% B to 40% A-60% B; 1.20-1.60 min, held at 40% A-60% B; 1.60-1.61 min, returned to 100% A-0% B, 1.61-2.00 min, held at 100% A-0% B.

##### 1.5 Reverse Phase HPLC conditions for the HRMS Analytical Methods

**Method 1:** Agilent 1290 LC & Agilent G6530B Q-TOF Agilent ZORBAX Extend-C18 2.1 × 50 mm, 5 µm at 40 ℃; Mobile Phase: A: 0.0375% TFA in water (v/v); B: 0.0188% TFA in Acetonitrile; flow rate held at 0.8 mL/min; eluted with the mobile phase over 4.50 min employing DAD detection. Gradient information: 0-0.40 min, held at 0% A-100% B; 0.40-3.40 min, ramped from 0% A-100% B to 10% A-90% B; 3.40-3.90 min, ramped from 10% A-90% B to 0% A-100% B; 3.91-4.00 min, returned to 0% A-100% B, 4.00-4.50 min, held at 0% A-100% B (flow rate held at 1.0 mL/min).

##### 1.6 Preparative Reverse Phase HPLC conditions

**Method 1:** column: Waters Xbridge 150 x 25 mm x 5 µm; mobile phase: [water ($NH_4HCO_3$)-ACN]; B%: 18%-48%, 10 min.

**Method 2:** column: Waters Xbridge 150 x 25 mm x 5 µm; mobile phase: [water (ammonia hydroxide v/v)-ACN]; B%: 19%-49%, 9 min.

**<u>Method 3</u>:** column: Waters Xbridge 150 x 25 mm x 5 µm; mobile phase: [water (ammonia hydroxide v/v)-ACN]; B%: 22%-52%, 9 min

**<u>Method 4</u>:** column: Waters Xbridge 150 x 25 mm x 5 µm; mobile phase: [water (ammonia hydroxide v/v)-ACN]; gradient: 23%-53% B, 9 min

**<u>Method 5</u>:** column: Waters Xbridge 150 x 25 mm x 5 µm; mobile phase: [water ($NH_4HCO_3$)-ACN]; B%: 10%-40%, 8 min

**<u>Method 6</u>:** column: Waters Xbridge 150 x 25 mm x 5 µm; mobile phase: [water ($NH_4HCO_3$)-ACN]; B%: 7%-37%, 8 min

**<u>Method 6</u>:** column: Waters Xbridge 150 x 25 mm x 5 µm; mobile phase: [water (ammonia hydroxide v/v)-ACN]; B%: 2%-30%, 9 min

**<u>Method 7</u>:** column: Waters Xbridge 150 x 25 mm x 5 µm; mobile phase: [water ($NH_4HCO_3$)-ACN]; B%: 21%-51%, 10 min

**<u>Method 8</u>:** column: Waters Xbridge 150 x 25 mm x 5 µm; mobile phase: [water (ammonia hydroxide v/v)-ACN]; B%: 15%-45%, 9 min

**<u>Method 9</u>:** column: Waters Xbridge 150 x 25 mm x 5 µm; mobile phase: [water (ammonia hydroxide v/v)-ACN]; B%: 14%-44%, 9 min

**<u>Method 10</u>:** column: Phenomenex luna C18 150 x 25 mm x 10 µm; mobile phase: [water(FA)-ACN]; B%: 20%-50%, 10 min

**<u>Method 11</u>:** column: Waters Xbridge 150 x 25 mm x 5 µm; mobile phase: [water (ammonia hydroxide v/v)-ACN]; B%: 25%-55%, 9 min

**<u>Method 12</u>:** column: Waters Xbridge 150 x 25 mm x 5 µm; mobile phase: [water (ammonia hydroxide v/v)-ACN]; B%: 18%-48%, 9 min

**<u>Method 13</u>:** column: Waters Xbridge 150 x 25 mm x 5 µm; mobile phase: [water ($NH_4HCO_3$)-ACN]; gradient:22%-52% B, 10 min

**<u>Method 14</u>:** column: Waters Xbridge 150 x 25 mm x 5 µm; mobile phase: [water ($NH_4HCO_3$)-ACN]; gradient: 23%-53% B, 9 min

**<u>Method 15</u>:** column: Waters Xbridge 150 x 25 mm x 5 µm; mobile phase: [water ($NH_4HCO_3$)-ACN]; gradient: 26%-56% B, 9 min

**<u>Method 16</u>:** column: Waters Xbridge 150 x 25 mm x 5 µm; mobile phase: [water ($NH_4HCO_3$)-ACN]; gradient: 28%-58% B, 9 min

**<u>Method 17</u>:** column: Waters Xbridge 150 x 25 mm x 5 µm; mobile phase: [water ($NH_4HCO_3$)-ACN]; gradient: 24%-54% B, 9 min

**<u>Method 18</u>:** column: Phenomenex C18 80 x 30 mm x 3 µm; mobile phase: [water ($NH_4HCO_3$)-

ACN]; gradient: 16%-46% B, 10 min

**Method 19:** column: Waters Xbridge 150 x 2 5mm 10 µm; mobile phase: [water ($NH_4HCO_3$)-ACN]; gradient: 15% - 45% B, 10 min.

#### 1.7 Synthetic Procedures

##### 1.7.1 Synthesis of compounds 1-3, 7-17, 23 and 24

**Scheme S1**

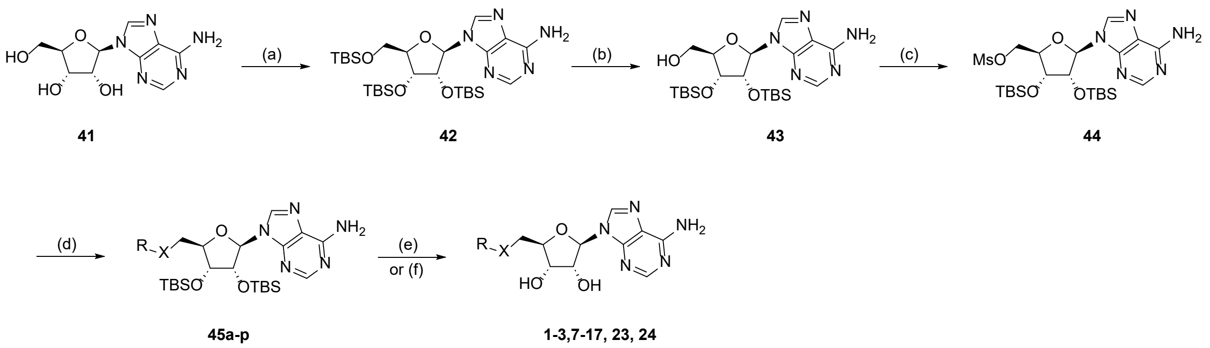

(a) TBS-Cl, 1H-imidazole, DMF, RT, 16 h, 79%; (b) $CCl_3CO_2H$, THF, $H_2O$, 0 °C, 4 h, 61%; (c) $Ms_2O$, TEA, THF, RT, 2 h, 17%; (d) R-XH, NaH, THF, 50 °C, 16 h, 9-87%; (e) KF, MeOH, RT, 16 h, 7-69% or (f) $Et_3N$-3HF, THF, RT, 16 h, 17-42%.

##### 9-[(2*R*,3*R*,4*R*,5*R*)-3,4-Bis[[tert-butyl(dimethyl)silyl]oxy]-5-[[tert-butyl(dimethyl)silyl] oxymethyl]tetrahydrofuran-2-yl]purin-6-amine (42)

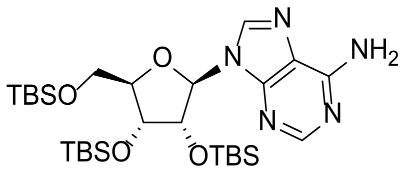

To a solution of (2*R*,3*R*,4*S*,5*R*)-2-(6-aminopurin-9-yl)-5-(hydroxymethyl)tetrahydrofuran -3,4-diol **41** (10.0 g, 37.4 mmol) in DMF (40 mL) was added 1H-imidazole (28.0 g, 411 mmol) and *tert*-butylchlorodimethylsilane (25.2 mL, 206 mmol). The resulting mixture was stirred at RT for 16 h. The reaction was quenched with water (200 mL) and extracted with EtOAc (3 x 100 mL). The combined organic layers were washed with brine (3 x 50 mL), dried over sodium sulfate, filtered, and concentrated under reduced pressure. The residue was purified by flash column chromatography (petroleum ether/ethyl acetate gradient 2-20%) to afford compound **42** (18.0 g, 79% yield) as a white solid. LCMS ($ES^+$, Method 1) *m/z* 610.4 $[M+H]^+$. $^1H$ NMR (400 MHz, $CDCl_3$): $\delta$ 8.34 (s, 1H), 8.16 (s, 1H), 6.02 (d, *J* = 4.8 Hz, 1H), 5.79 (s, 2H), 4.69 (t, *J* = 4.8 Hz, 1H), 4.32 (t, *J* = 4.0 Hz, 1H), 4.15 - 4.13 (m, 1H), 4.03 (dd, *J* = 11.6, 4.0 Hz, 1H), 3.79 (dd, *J* =

11.6, 3.2 Hz, 1H), 0.96 (s, 9H), 0.93 (s, 9H), 0.80 (s, 9H), 0.14 (s, 3H), 0.13 (s, 3H), 0.11 (s, 3H), 0.10 (s, 3H), -0.04 (s, 3H), -0.22 (s, 3H).

**[(2*R*,3*R*,4*R*,5*R*)-5-(6-Aminopurin-9-yl)-3,4-bis[[tert-butyl(dimethyl)silyl]oxy] tetrahydrofuran-2-yl]methanol (43)**

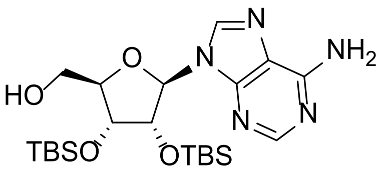

To a solution of compound **42** (10.0 g, 16.4 mmol) in THF (100 mL) was added dropwise a solution of trichloroacetic acid (26.8 g, 164 mmol) in water (30 mL) at 0 °C and stirred for 4 h. The reaction mixture was neutralized with sodium carbonate, diluted with water (100 mL) and extracted with EtOAc (3 x 50 mL). The combined organic layers were dried over sodium sulfate, filtered, and concentrated under reduced pressure. The residue was purified by flash column chromatography (petroleum ether/ethyl acetate gradient 2-20%) to afford compound **43** (5.00 g, 61% yield) as a white solid. LCMS (ES$^+$, Method 2) *m/z* 496.4 [M+H]$^+$. $^1$H NMR (400 MHz, DMSO-$d_6$): $\delta$ 8.40 (s, 1H), 8.14 (s, 1H), 7.38 (s, 2H), 5.90 (d, *J* = 7.2 Hz, 1H), 5.72 (dd, *J* = 8.0, 4.0 Hz, 1H), 4.91 (dd, *J* = 7.2, 4.4 Hz, 1H), 4.30 (d, *J* = 4.4 Hz, 1H), 3.99 (t, *J* = 2.8 Hz, 1H), 3.76 - 3.72 (m, 1H), 3.62 - 3.59 (m, 1H), 0.93 (s, 9H), 0.70 (s, 9H), 0.13 (s, 3H), 0.12 (s, 3H), -0.04 (s, 3H), -0.13 (s, 3H), -0.46 (s, 3H).

**[(2*R*,3*R*,4*R*,5*R*)-5-(6-Aminopurin-9-yl)-3,4-bis[[tert-butyl(dimethyl)silyl] oxy] tetrahydrofuran-2-yl]methyl methanesulfonate (44)**

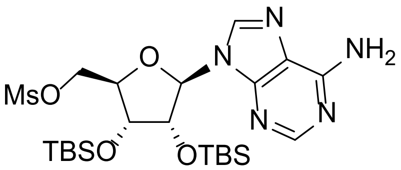

To a solution of compound **43** (1.00 g, 2.02 mmol) and triethylamine (842 µL, 6.05 mmol) in THF (10 mL) was added dropwise a solution of methanesulfonic anhydride (422 mg, 2.42 mmol) in THF (5 mL) under $N_2$ at 0 °C. The resulting mixture was warmed to RT and stirred for 2 h. The reaction was quenched with water (50 mL) and then extracted with EtOAc (3 x 30 mL). The combined organic layers were washed with brine (3 x 20 mL), dried over sodium sulfate, filtered, and concentrated under reduced pressure. The residue was purified by flash column chromatography (petroleum ether/ethyl acetate gradient 2-50%) to afford compound **44** (200 mg, 348 µmol, 17% yield) as a white solid. LCMS (ES$^+$, Method 2) *m/z* 574.2 [M+H]$^+$. $^1$H NMR (400 MHz, $CDCl_3$): *δ* 8.35 (s, 1H), 7.94 (s, 1H), 5.89 (d, *J* = 5.2 Hz, 1H), 5.74 (s, 2H), 5.02 (t, *J* = 4.8 Hz, 1H), 4.61 (dd, *J* = 11.2, 4.4 Hz, 1H), 4.52 (dd, *J* = 11.2, 5.2 Hz, 1H), 4.41 (t, *J* = 4.0 Hz, 1H), 4.33 - 4.32 (m, 1H), 3.00 (s, 3H), 0.95 (s, 9H), 0.82 (s, 9H), 0.14 (s, 3H), 0.13 (s, 3H), -0.02 (s, 3H), -0.23 (s, 3H).

**9-[(2*R*,3*R*,4*R*,5*S*)-5-(1,3-Benzothiazol-2-ylsulfanylmethyl)-3,4-bis[[tert-butyl (dimethyl)silyl] oxy]tetrahydrofuran-2-yl]purin-6-amine (45a)**

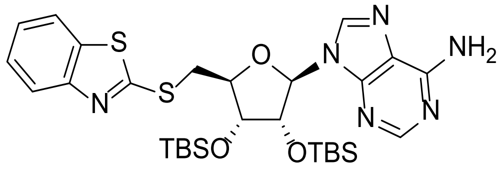

To a solution of compound **44** (100 mg, 174 µmol) in DMF (2 mL) was added 1,3-benzothiazol-2-ylsulfanylsodium (49.5 mg, 261 µmol). The resulting mixture was heated to 50 °C and stirred for 16 h. The reaction was cooled to RT, quenched with water (20 mL) and extracted with EtOAc (3 x 10 mL). The combined organic layers were washed with brine (10 mL), dried over sodium sulfate, filtered, and concentrated under reduced pressure. The residue was purified by flash column chromatography (petroleum ether/ethyl acetate gradient 0-2%) to afford compound **45a** (70.0 mg, 62% yield) as a white solid LCMS (ES$^+$, Method 2) *m/z* 645.5 [M+H]$^+$.

**9-[(2*R*,3*R*,4*R*,5*S*)-3,4-Bis[[tert-butyl(dimethyl)silyl]oxy]-5-[(6- methyl-1,3-benzothiazol-2-yl) sulfanylmethyl]tetrahydrofuran-2-yl]purin-6-amine (45b)**

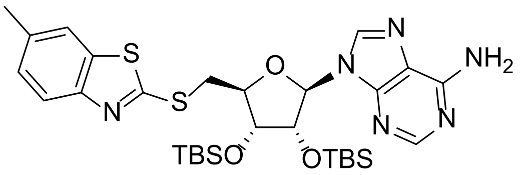

To a solution of 6-methyl-1,3-benzothiazole-2-thiol (63.2 mg, 348 µmol) in THF (2 mL) was added NaH (13.9 mg, 348 µmol, 60% dispersion in mineral oil) under inert atmosphere at 0 °C and the mixture was stirred for 30 min. A solution of compound **44** (100 mg, 174 µmol) in THF

(1 mL) was then added and the resulting mixture was heated to 70 °C and stirred for 16 h. The reaction was cooled to RT, quenched with water (20 mL) and extracted with EtOAc (3 x 10 mL). The combined organic layers were washed with brine (10 mL), dried over sodium sulfate, filtered, and concentrated under reduced pressure. The residue was purified by flash column chromatography (petroleum ether/ethyl acetate gradient 2-50%) to afford compound **45b** (100 mg, 87% yield) as a white solid. LCMS ($ES^+$, Method 3) *m/z* 659.3 $[M+H]^+$. $^1H$ NMR (400 MHz, DMSO-$d_6$): $\delta$ 8.45 (s, 1H), 8.16 (s, 1H), 7.80 (s, 1H), 7.69 (d, *J* = 8.0 Hz, 1H), 7.31 - 7.28 (m, 3H), 5.94 (d, *J* = 7.6 Hz, 1H), 5.24 (dd, *J* = 7.6, 4.4 Hz, 1H), 4.44 (d, *J* = 4.4 Hz, 1H) , 4.30 (t, *J* = 7.2 Hz, 1H), 4.04 - 4.01 (m, 1H), 3.80 - 3.75 (m, 1H), 2.41 (s, 3H), 0.88 (s, 9H), 0.67 (s, 9H), 0.08 (s, 3H), 0.07 (s, 3H), -0.13 (s, 3H), -0.46 (s, 3H).

**9-[(2*R*,3*R*,4*R*,5*S*)-3,4-bis[[tert-butyl(dimethyl)silyl]oxy]-5- [(5-methyl-1,3-benzothiazol-2-yl)sulfanylmethyl]tetrahydrofuran-2-yl]purin-6-amine (45c)**

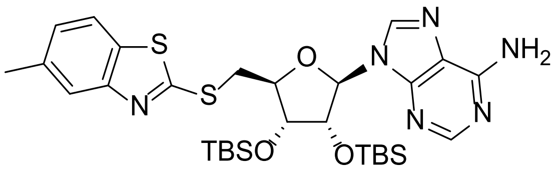

To a solution of 5-methyl-1,3-benzothiazole-2-thiol (63.1 mg, 348 µmol) in THF (2 mL) was added NaH (13.9 mg, 348 µmol, 60% dispersion in mineral oil) under inert atmosphere at 0 °C and the mixture was stirred for 30 min. A solution of compound **44** (100 mg, 174 µmol) in THF (2 mL) was then added and the resulting mixture was heated to 70 °C and stirred for 16 h. The reaction was cooled to RT, quenched with water (20 mL) and extracted with EtOAc (3 x 10 mL). The combined organic layers were washed with brine (10 mL), dried over sodium sulfate, filtered, and concentrated under reduced pressure. The residue was purified by flash column chromatography (petroleum ether/ethyl acetate gradient 2-50%) to afford compound **45c** (50.0 mg, 43.5% yield) as a white solid. LCMS ($ES^+$, Method 2) *m/z* 659.4 $[M+H]^+$.

**9-[(2*R*,3*R*,4*R*,5*S*)-3,4-bis[[tert-butyl(dimethyl)silyl]oxy]-5-(2-quinolylsulfanylmethyl) tetrahydrofuran-2-yl]purin-6-amine (45d)**

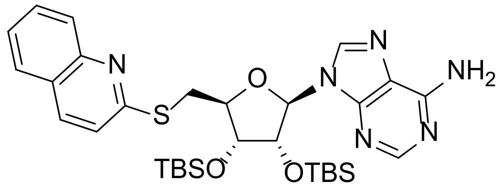

To a solution of quinoline-2-thiol (112 mg, 697 µmol) in THF (2 mL) was added NaH (27.9 mg, 697 µmol, 60% dispersion in mineral oil) under inert atmosphere at 0 °C and the mixture was stirred for 30 min. A solution compound **44** (200 mg, 348 µmol) in THF (2 mL) was then added and the resulting mixture was heated to 70 °C and stirred for 16 h. The reaction was cooled to RT, quenched with water (30 mL) and extracted with EtOAc (3 x 15 mL). The combined organic layers were washed with brine (2 x 10 mL), dried over sodium sulfate, filtered, and concentrated under reduced pressure. The residue was purified by flash column chromatography (petroleum ether/ethyl acetate gradient 2-50%) to afford compound **45d** (180 mg, 80% yield) as colorless oil. $^1$H NMR (400 MHz, DMSO-$d_6$): $\delta$ 8.39 (s, 1H), 8.01 (s, 1H), 7.93 (t, $J$ = 8.8 Hz, 2H), 7.74 (d, $J$ = 8.4 Hz, 1H), 7.68 - 7.64 (m, 1H), 7.45 (t, $J$ = 7.2 Hz, 1H), 7.24 (d, $J$ = 8.4 Hz, 1H), 5.92 (d, $J$ = 6.4 Hz, 1H), 5.73 (s, 2H), 5.26 (dd, $J$ = 6.4, 4.0 Hz, 1H), 4.47 (td, $J$ = 6.8, 1.6 Hz, 1H), 4.36 (dd, $J$ = 4.4, 1.6 Hz, 1H), 4.10 - 4.05 (m, 1H), 3.85 (dd, $J$ = 14.0, 6.4 Hz, 1H), 0.91 (s, 9H), 0.73 (s, 9H), 0.07 (s, 6H), -0.19 (s, 3H), -0.39 (s, 3H).

**9-[(2R,3R,4R,5S)-3,4-Bis[[tert-butyl(dimethyl)silyl]oxy]-5- [(6-methyl-2-quinolyl) sulfanyl methyl]tetrahydrofuran-2-yl]purin-6-amine (45e)**

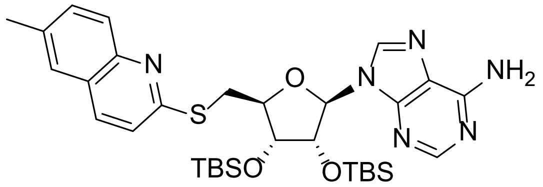

To a solution of 6-methylquinoline-2-thiol **73** (61.1 mg, 348 µmol) in THF (2 mL) was added NaH (13.9 mg, 348 µmol, 60% dispersion in mineral oil) under inert atmosphere at 0 °C and the mixture was stirred for 30 min. A solution of compound **44** (100 mg, 174 µmol) in THF (2 mL) was then added and the resulting mixture was heated to 70 °C for 16 h. The reaction was cooled to RT, quenched with water (20 mL) and extracted with EtOAc (3 x 10 mL). The combined organic layers were washed with brine (10 mL), dried over sodium sulfate, filtered, and the filtrate was concentrated under reduced pressure. The residue was purified by flash column chromatography (petroleum ether/ethyl acetate gradient 2-50%) to afford compound **45e** (90.0 mg, 79% yield) as a white solid. LCMS (ES$^+$, Method 2) *m/z* 653.5 [M+H]$^+$.

**9-[(2*R*,3*R*,4*R*,5*S*)-3,4-Bis[[tert-butyl(dimethyl)silyl]oxy]-5-[(7 -methyl-2-quinolyl)sulfanyl methyl]tetrahydrofuran-2-yl]purin-6-amine (45f)**

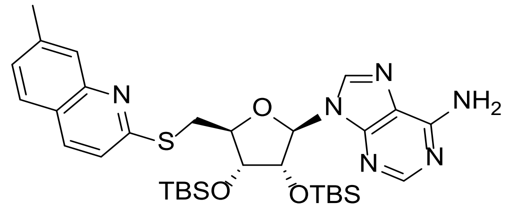

To a solution of 7-methylquinoline-2-thiol **77** (61.1 mg, 348 µmol) in THF (2 mL) was added NaH (13.9 mg, 348 µmol, 60% dispersion in mineral oil) under inert atmosphere at 0 °C and the mixture was stirred for 30 min. A solution of compound **44** (100 mg, 174 µmol) in THF (1 mL) was added and the resulting mixture was heated to 70 °C for 16 h. The reaction was cooled to RT, quenched with water (20 mL) and extracted with EtOAc (3 x 10 mL). The combined organic layers were washed with brine (10 mL), dried over sodium sulfate, filtered and the filtrate was concentrated under reduced pressure. The residue was purified by preparative TLC ($SiO_2$, petroleum ether/ethyl acetate 50%) to afford compound **45f** (60.0 mg, 52% yield) as a white solid. LCMS (ES$^+$, Method 2) *m/z* 635.5 [M+H]$^+$.

**9-((2R,3R,4R,5S)-3,4-Bis((tert-butyldimethylsilyl)oxy)-5-((quinolin-7-ylthio)methyl) tetrahydrofuran-2-yl)-9H-purin-6-amine (45g)**

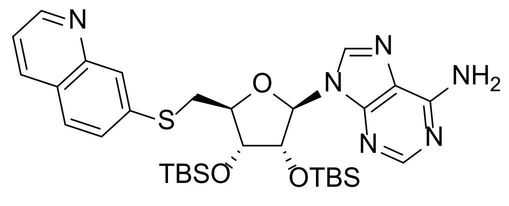

To a solution of imidazo-quinoline-7-thiol (110 mg, 680 µmol) in THF (2 mL) was added NaH (27.2 mg, 680 µmol, 60% dispersion in mineral oil) under inert atmosphere at 0 °C and the mixture was stirred for 30 min. A solution of compound **44** (130 mg, 227 µmol) in THF (1 mL) was then added and the resulting mixture was heated at 70 °C for 16 h. The reaction mixture was quenched with water (10 mL) and extracted with EtOAc (3 × 10 mL). The combined organic layers were washed with brine (10 mL), dried over sodium sulfate, filtered, and concentrated under reduced pressure. The residue was purified by flash column chromatography (petroleum ether/ethyl acetate gradient 0-100%) to afford compound **45g** (120 mg, 83% yield) as colorless oil. LCMS (ES$^+$, Method 1) *m/z* 639.3 [M+H]$^+$. $^1$H NMR (400 MHz, $CDCl_3$): *δ* 8.91 - 8.86 (m, 1H), 8.35 (s, 1H), 8.13 - 8.08 (m, 1H), 8.07 - 8.047 (m, 1H), 7.89 (s, 1H), 7.71 (d, *J* = 8.8 Hz, 1H), 7.51 - 7.46 (m, 1H), 7.38 - 7.34 (m, 1H), 5.86 (d, *J* = 6 Hz, 1H), 5.83 - 5.75 (m, 2H), 5.28 - 5.23 (m, 1H), 4.39 - 4.33 (m, 2H), 3.79 - 3.72 (m, 1H), 3.61 - 3.53 (m, 1H), 0.90 (s, 9H), 0.76 (s, 9H), 0.11 - 0.06 (m, 6H), -0.14 (s, 3H), -0.35 (s, 3H).

**9-[(2*R*,3*R*,4*R*,5*S*)-3,4-Bis[[tert-butyl(dimethyl)silyl]oxy]-5-(thiazol-2-ylsulfanylmethyl)tetrahydrofuran-2-yl]purin-6-amine (45h)**

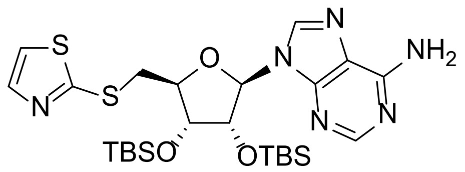

To a solution of thiazole-2-thiol (79.6 mg, 679 µmol) in THF (2 mL) was added NaH (27.2 mg, 679 µmol, 60% dispersion in mineral oil) under inert atmosphere at 0 °C and the mixture was stirred for 30 min. A solution of compound **44** (130 mg, 226 µmol) in THF (2 mL) was then added, and the resulting mixture was heated to 70 °C for 16 h. The reaction was cooled at 0 °C, quenched with water (20 mL), and then extracted with EtOAc (3 x 10 mL). The combined organic layers were washed with brine (2 x 10 mL), dried over sodium sulfate, filtered, and concentrated under reduced pressure. The residue was purified by flash column chromatography (petroleum ether/ethyl acetate gradient 2-100%) to afford compound **45h** (70.0 mg, 52% yield) as a white solid. LCMS (ES$^+$, Method 1) *m/z* 595.2 [M+H]$^+$. $^1$H NMR (400MHz, $CDCl_3$): *δ* 8.35 (s, 1H), 7.91 (s, 1H), 7.65 (d, *J* = 3.2 Hz, 1H), 7.22 (d, *J* = 3.2 Hz, 1H), 5.88 (d, *J* = 6.0 Hz, 1H), 5.59 (s, 2H), 5.31 - 5.27 (m, 1H), 4.39 (td, *J* = 6.4, 1.6 Hz, 1H), 4.35 - 4.33 (m, 1H), 3.93 (dd, *J* = 14.0, 6.8 Hz, 1H), 3.70 (dd, *J* = 14.4, 6.4 Hz, 1H), 0.93 (s, 9H), 0.77 (s, 9H), 0.10 (s, 3H), 0.09 (s, 3H), 0.01 (s, 3H), -0.09 (s, 3H).

**9-[(2*R*,3*R*,4*R*,5*S*)-3,4-Bis[[tert-butyl(dimethyl)silyl]oxy]-5-(imidazo[1,5-a]pyridin-8-ylsulfanylmethyl)tetrahydrofuran-2-yl]purin-6-amine (45i)**

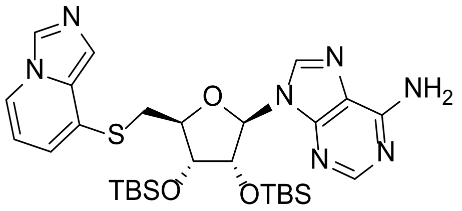

To a solution of compound **80** (47.1 mg, 313 µmol) in THF (2 mL) was added NaH (12.5 mg, 313 µmol, 60% dispersion in mineral oil) under inert atmosphere at 0 °C and the mixture was stirred for 30 min. A solution of compound **44** (120 mg, 209 µmol) in THF (1 mL) was then added and the resulting mixture was heated at 70 °C for 16 h. The reaction mixture was cooled to RT, quenched with water (20 mL) and extracted with EtOAc (3 x 10 mL). The combined organic layers were washed with brine (10 mL), dried over sodium sulfate, filtered, and concentrated under reduced pressure. The residue was purified by flash column chromatography (petroleum ether/ethyl acetate gradient 0-50%) to afford compound **45i** (80.0 mg, 61% yield) as yellow oil. LCMS ($ES^+$, Method 2) *m/z* 628.5 $[M+H]^+$. $^1H$ NMR (400MHz, $CDCl_3$): $\delta$ 8.33 (s, 1H), 8.13 (s, 1H), 7.85 (s, 1H), 7.80 (d, *J* = 6.8 Hz, 1H), 7.55 (s, 1H), 6.71 (d, *J* = 6.8 Hz, 1H), 6.49 (t, *J* = 6.8 Hz, 1H), 5.83 (d, *J* = 6.0 Hz, 1H), 5.66 (s, 2H), 5.28 (dd, *J* = 6.0, 4.4 Hz, 1H), 4.35 - 4.34 (m, 1H), 4.32 - 4.28 (m, 1H), 3.72 - 3.70 (m, 1H), 3.48 - 3.43 (m, 1H), 0.90 (s, 9H), 0.76 (s, 9H), 0.08 (s, 3H), 0.04 (s, 3H), -0.02 (s, 3H), -0.36 (s, 3H).

**9-((2R,3R,4R,5S)-3,4-Bis((tert-butyldimethylsilyl)oxy)-5-(((1-methyl-1H-benzo[d]imidazol-2-yl)thio)methyl)tetrahydrofuran-2-yl)-9H-purin-6-amine (45j)**

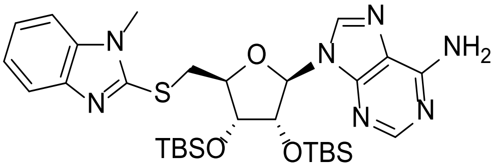

To a solution of 1-methyl-1H-benzo[d]imidazole-2-thiol (57.2 mg, 349 µmol) in THF (4 mL) was added NaH (13.9 mg, 349 µmol, 60% dispersion in mineral oil) under inert atmosphere at 0 °C and the reaction mixture was stirred for 30 min. A solution of compound **44** (100 mg, 174 µmol) in THF (1 mL) was then added and the resulting mixture was heated at 70 °C for 16 h. The reaction mixture was cooled to RT, quenched with water (20 mL) and extracted with EtOAc (3 × 10 mL). The combined organic layers were washed with brine (10 mL), dried over sodium sulphate, filtered and then concentrated under reduced pressure. The residue was purified by preparative TLC ($SiO_2$, petroleum ether/ethyl acetate 50%) to afford compound **45j** (98.0 mg, 87% yield) as a white solid. LCMS ($ES^+$, Method 2) *m/z* 642.4 $[M+H]^+$. $^1H$ NMR (400 MHz, $CDCl_3$): $\delta$ 8.35 (s, 1H), 7.99 (s, 1H), 7.70 - 7.60 (m, 1H), 7.25 - 7.21 (m, 3H). 5.90 (d, $J$ = 6.8 Hz, 1H), 5.79 (brs, 2H), 5.31 - 5.28 (m, 1H), 4.49 - 4.46 (m, 1H), 4.41 - 4.39 (m, 1H), 4.07 - 4.02 (m, Hz, 1H), 3.97 - 3.92 (m, 1H), 3.66 (s, 3H), 0.93 (s, 9H), 0.76 (s, 9H), 0.12 (s, 3H), 0.11(s, 3H), -0.10 (s, 3H), -0.37 (s, 3H).

**9-((2R,3R,4R,5S)-5-((Benzo[d]oxazol-2-ylthio)methyl)-3,4-bis((tert-butyldimethylsilyl) oxy)tetrahydrofuran-2-yl)-9H-purin-6-amine (45k)**

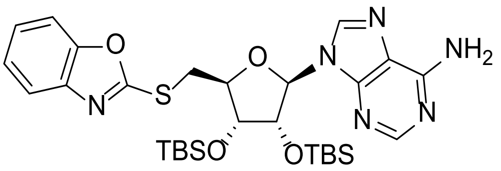

To a solution of 1,3-benzoxazole-2-thiol (105 mg, 697 µmol) in THF (4 mL) was added NaH (27.9 mg, 697 µmol, 60% dispersion in mineral oil) under inert atmosphere at 0 °C and the mixture was stirred for 30 min. A solution of compound **44** (200 mg, 349 µmol) in THF (2 mL) was then added and the resulting mixture was stirred initially at RT for 30 min and then heated at 70 °C for 16 h. The reaction mixture was cooled to RT, quenched with water (20 mL) and extracted with EtOAc (3 × 10 mL). The combined organic layers were washed with brine (10 mL), dried sodium sulphate, filtered, and concentrated under reduced pressure. The residue was purified by reverse column chromatography (0.1% $NH_3{\cdot}H_2O$ in water/MeCN gradient 0-100%) to afford compound **45k** (160 mg, 73% yield) as a yellow oil. LCMS ($ES^+$, Method 2) *m/z* 629.4 $[M+H]^+$.

**9-((2R,3R,4R,5S)-5-(((1H-Benzo[d]imidazol-2-yl)thio)methyl)-3,4-bis((tertbutyldimethyl silyl)oxy)tetrahydrofuran-2-yl)-9H-purin-6-amine (45l)**

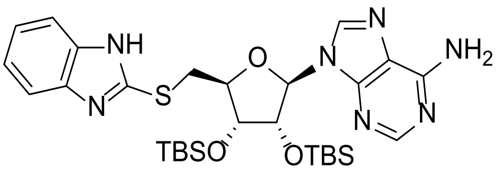

To a solution of 1H-benzimidazole-2-thiol (78.5 mg, 522.8 µmol) in THF (4 mL) was added NaH (20.9 mg, 522.8 µmol, 60% dispersion in mineral oil) under inert atmosphere at 0 °C and the mixture was stirred for 30 min. A solution of compound **44** (200 mg, 349 µmol) in THF (2 mL) was added and the resulting mixture was stirred initially at RT for 30 mins and then heated at 70 °C and for 16 h. The reaction was cooled to RT, quenched with water (20 mL) and extracted with EtOAc (3 × 10 mL). The combined organic layers were washed with brine (10 mL), dried sodium sulphate, filtered, and concentrated under reduced pressure. The residue was purified by reverse column chromatography (0.1% $NH_3 \cdot H_2O$ in water/MeCN gradient 0-100%) to afford compound **45l** (103 mg, 47% yield) as a yellow oil. LCMS (ES$^+$, Method 2) *m/z* 628.4 [M+H]$^+$.

**9-((2R,3R,4R,5S)-5-(((1H-Indazol-7-yl)thio)methyl)-3,4-bis((tert-butyldimethylsilyl)oxy) tetrahydrofuran-2-yl)-9H-purin-6-amine (45m)**

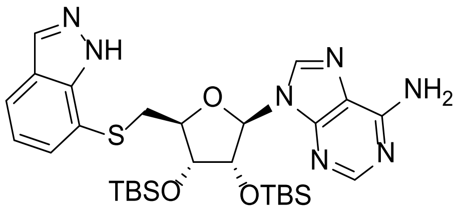

To a solution of compound **85** (52.7 mg, 349 µmol) in DMF (2 mL) was added NaH (13.9 mg, 349 µmol, 60% dispersion in mineral oil) under $N_2$ at 0 °C and the mixture was stirred for 30 min. A solution of compound **44** (100 mg, 174 µmol) in DMF (1 mL) was then added and the resulting mixture was heated at 70 °C for 16 h. The reaction mixture was cooled to RT, quenched with water (15 mL) and then extracted with EtOAc (3 × 20 mL). The combined organic layers were washed with brine (60 mL), dried over sodium sulfate, filtered, and concentrated under reduced pressure. The residue was purified by preparative TLC ($SiO_2$, petroleum ether/ethyl acetate 50%) to afford compound **45m** (10.0 mg, 9% yield) as a colorless oil. LCMS (ES$^+$, Method 3) *m/z* 628.1 [M+H]$^+$.

**9-[(2*R*,3*R*,4*R*,5*S*)-3,4-bis[[tert-butyl(dimethyl)silyl]oxy]-5-(imidazo [1,5-a]pyridin-7-ylsulfanylmethyl)tetrahydrofuran-2-yl]purin-6-amine (45n)**

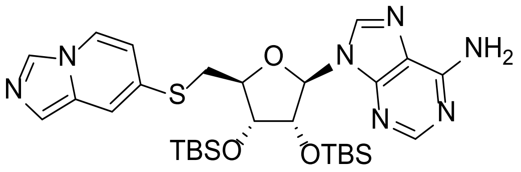

To a solution of imidazo[1,5-a]pyridine-7-thiol **91** (68.1 mg, 453 µmol) in THF (2 mL) was added NaH (18.1 mg, 453 µmol, 60% dispersion in mineral oil) under inert atmosphere at 0 °C and the mixture was stirred for 30 min. A solution of compound **44** (130 mg, 226 µmol) in THF (1 mL) was added and the resulting mixture was heated to 70 °C for 16 h. The reaction was cooled to RT, quenched with water (20 mL) and extracted with EtOAc (3 x 10 mL). The combined organic layers were washed with brine (10 mL), dried over sodium sulfate, filtered, and the filtrate was concentrated under reduced pressure. The residue was purified by flash column chromatography (petroleum ether/ethyl acetate gradient 0-50%) to afford compound **45n** (120 mg, 84% yield) as yellow oil. LCMS (ES$^+$, Method 2) *m/z* 628.3 [M+H]$^+$. $^1$H NMR (400 MHz, $CDCl_3$): *δ* 8.30 (s, 1H), 8.04 (s, 1H), 7.87 (s, 1H), 7.80 (d, *J* = 7.6 Hz, 1H), 7.36 (s, 1H), 7.28 - 7.27 (m, 1H), 6.50 (dd, *J* = 7.6, 2.0 Hz, 1H), 6.06 (s, 2H), 5.83 (d, *J* = 5.6 Hz, 1H), 5.24 (dd, *J* = 5.6, 4.4 Hz, 1H), 4.38 - 4.36 (m, 1H), 4.30 - 4.26 (m, 1H), 3.54 (dd, *J* = 14.4, 6.8 Hz, 1H), 3.70 (dd, *J* = 14.4, 6.0 Hz, 1H), 0.93 (s, 9H), 0.77 (s, 9H), 0.12 (s, 3H), 0.11 (s, 3H), -0.11 (s, 3H), -0.33 (s, 3H).

**9-[(2*R*,3*R*,4*R*,5*R*)-5-(1,3-Benzothiazol-2-yloxymethyl)-3,4-bis[[tert-butyl(dimethyl)silyl] oxy]tetrahydrofuran-2-yl]purin-6-amine (45o)**

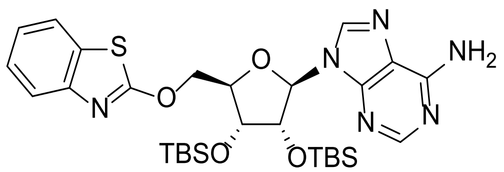

To a solution of 1,3-benzothiazol-2-ol (39.52 mg, 261.39 µmol) in THF (1.5 mL) was added NaH (10.5 mg, 261 µmol, 60% dispersion in mineral oil) under inert atmosphere at 0 °C and the mixture was stirred for 30 min. A solution of compound **44** (100 mg, 174 µmol) in THF (1.5 mL) was then added and the resulting mixture was heated to 50 °C for 2 h. The reaction was cooled to RT, quenched with water (10 mL) and extracted with EtOAc (3 x 5 mL). The combined organic layers were washed with brine (5 mL), dried over sodium sulfate, filtered, and concentrated under reduced pressure. The residue was purified by flash column chromatography (petroleum ether/ethyl acetate gradient 2-100%) to afford compound **45o** (30.0 mg, 27% yield) as a white solid. LCMS ($ES^+$, Method 2) *m/z* 629.2 $[M+H]^+$. $^1H$ NMR (400 MHz, $CDCl_3$): $\delta$ 8.40 (s, 1H), 7.81 (s, 1H), 7.44 (d, *J* = 7.6 Hz, 1H), 7.24 (d, *J* = 7.6 Hz, 1H), 7.18 - 7.16 (m, 2H), 5.86 (d, *J* = 7.2 Hz, 1H), 5.60 (s, 2H), 5.36 (dd, *J* = 6.4, 4.4 Hz, 1H), 4.76 (dd, *J* = 14.4, 7.6 Hz, 1H), 4.39 (d, *J* = 4.4 Hz, 1H), 4.36 (d, *J* = 6.8 Hz, 1H), 4.16 (d, *J* = 5.6 Hz, 1H), 0.87 (s, 9H), 0.75 (s, 9H), 0.04 (s, 3H), -0.06 (d, *J* = 5.6 Hz, 6H), -0.37 (s, 3H).

**N-[[(2R,3R,4R,5R)-5-(6-Aminopurin-9-yl)-3,4-bis[[tert-butyl(dimethyl)silyl]oxy] tetrahydrofuran-2-yl]methyl]-1,3-benzothiazol-2-amine (45p)**

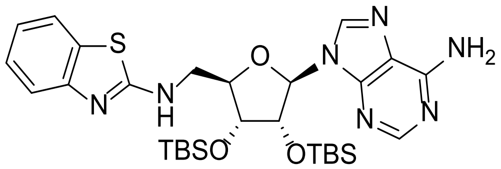

To a solution of 1,3-benzothiazol-2-amine (78.5 mg, 522 µmol) in THF (1 mL) was added NaH (20.9 mg, 522 µmol, 60% dispersion in mineral oil) under inert atmosphere at 0 °C and the mixture was stirred for 30 min. A solution of compound **44** (100 mg, 174 µmol) in THF (1 mL) was then added and the resulting mixture was heated to 70 °C for 16 h. The reaction was cooled to room RT, quenched with water (20 mL) and extracted with EtOAc (3 x 10 mL). The combined organic layers were washed with brine (10 mL), dried over sodium sulfate, filtered, and the filtrate was concentrated under reduced pressure. The residue was purified by flash column chromatography (petroleum ether/ethyl acetate gradient 2-50%) to afford compound **45p** (35.0 mg, 32% yield) as colorless oil. LCMS ($ES^+$, Method 3) *m/z* 628.2 $[M+H]^+$.

**(2R,3R,4S,5S)-2-(6-aminopurin-9-yl)-5-(1,3-benzothiazol-2-ylsulfanylmethyl) tetrahydrofuran-3,4-diol (1)**

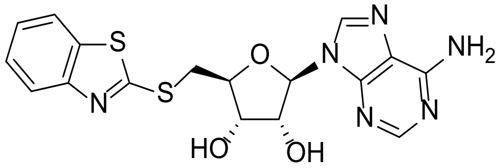

To a solution of **45a** (50.0 mg, 77.5 µmol) in MeOH (2 mL) was added potassium fluoride (13.5

mg, 232 µmol). The reaction mixture was stirred at RT for 16 h and then concentrated under reduced pressure. The residue was purified by prep-HPLC (Method 1) to afford compound **1** (12.9 mg, 40% yield) as a white solid. HRMS (Method 1) *m/z* [M+H]$^+$ calcd for $C_{17}H_{16}N_6O_3S_2$ 417.0804; found 417.0803. $^1$H NMR (400 MHz, DMSO-*d*$_6$): *δ* 8.39 (s, 1H), 8.15 (s, 1H), 8.01 (d, *J* = 8.0 Hz, 1H), 7.86 (d, *J* = 8.0 Hz, 1H), 7.47 (t, *J* = 7.6 Hz, 1H), 7.39 - 7.33 (m, 3H), 5.88 (d, *J* = 5.6 Hz, 1H), 5.92 (d, *J* = 6.0 Hz, 1H), 5.58 (s, 1H), 5.49 (s, 1H), 7.84 (s, 1H), 4.28 - 4.24 (m, 2H), 3.89 - 3.84 (m, 1H), 3.77 - 3.74 (m, 1H).

**(2*R*,3*R*,4*S*,5*S*)-2-(6-aminopurin-9-yl)-5-[(6-methyl-1,3-benzothiazol -2-yl)sulfanylmethyl] tetrahydrofuran-3,4-diol (2)**

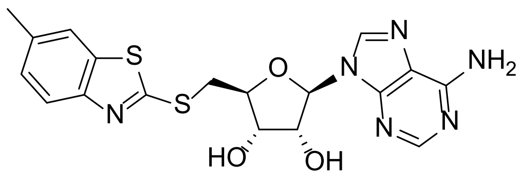

To a solution of **45b** (100 mg, 0.15 mmol) in DMF (2 mL) was added potassium fluoride (88.1 mg, 1.52 mmol). The reaction mixture was stirred at 60 °C for 16 h, cooled to RT and then concentrated under reduced pressure. The residue was purified by prep-HPLC (Method 2) to afford compound **2** (23.4 mg, 35% yield) as an off-white solid. HRMS *m/z* [M+H]$^+$ calcd for $C_{18}H_{18}N_6O_3S_2$ 431.5090; found 431.0964. $^1$H NMR (400 MHz, DMSO-*d*$_6$): *δ* 8.37 (s, 1H), 8.15 (s, 1H), 7.78 (s, 1H), 7.73 (d, *J* = 8.4 Hz, 1H), 7.30 - 7.27 (m, 3H), 5.92 (d, *J* = 5.6 Hz, 1H), 5.51 (brs, 2H), 4.86 (t, *J* = 5.2 Hz, 1H), 4.29 - 4.27 (m, 2H), 4.23 - 4.22 (m, 1H), 3.84 (dd, *J* = 13.6, 5.6 Hz, 1H), 3.72 (dd, *J* = 13.6, 7.6 Hz, 1H), 2.41 (s, 3H).

**2*R*,3*R*,4*S*,5*S*)-2-(6-aminopurin-9-yl)-5-[(5-methyl-1,3-benzothiazol -2-yl)sulfanylmethyl] tetrahydrofuran-3,4-diol (3)**

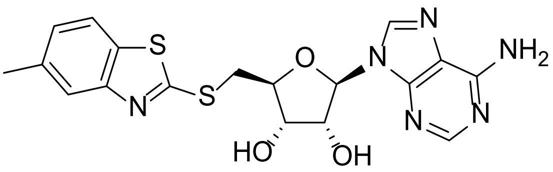

To a solution of compound **45c** (50.0 mg, 75.8 µmol) in DMF (2 mL) was added potassium fluoride (44.1 mg, 758 µmol). The reaction mixture was stirred at 60 °C for 16 h, cooled to RT and then concentrated under reduced pressure. The residue was purified by prep-HPLC (Method 1) to afford compound **3** (11.2 mg, 34% yield) as a white solid. HRMS (Method 3) *m/z* $[M+H]^+$ calcd for $C_{17}H_{17}N_7O_3S$ 431.0960; found 431.0975. $^1H$ NMR (400 MHz, DMSO-$d_6$): $\delta$ 8.45 (s, 1H), 8.22 (s, 1H), 7.86 (d, *J* = 8.4 Hz, 1H), 7.67 (s, 2H), 7.66 - 7.56 (m, 1H), 7.19 (d, *J* = 7.6 Hz, 1H), 5.93 (d, *J* = 5.6 Hz, 1H), 5.65 - 5.38 (s, 1H), 4.84 (t, *J* = 5.2 Hz, 1H), 4.21 - 4.32 (m, 2H) , 3.85 (dd, *J* = 13.6, 5.6 Hz, 1H), 3.73 (dd, *J* = 13.6, 5.6 Hz,1H), 2.42 (s, 3H).

**(2*R*,3*R*,4*S*,5*S*)-2-(6-aminopurin-9-yl)-5-(2-quinolylsulfanylmethyl) tetrahydrofuran-3,4-diol (7)**

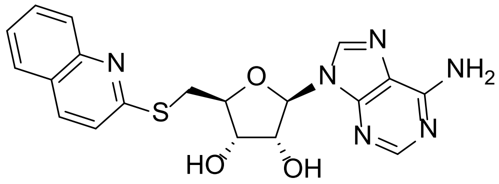

To a solution of compound **45d** (130 mg, 0.20 mmol) in MeOH (5 mL) was added potassium fluoride (118 mg, 2.03 mmol). The reaction mixture was stirred at 60 °C for 16 h, cooled to RT and then concentrated under reduced pressure. The residue was purified by prep-HPLC (Method 1) to compound **7** (13.12 mg, 16% yield) as a white solid. HRMS (Method 1) *m/z* $[M+H]^+$ calcd for $C_{19}H_{18}N_6O_3S$ 411.1239; found: 411.1249. $^1H$ NMR (400 MHz, DMSO-$d_6$): $\delta$ 8.40 (s, 1H), 8.18 - 8.15 (m, 2H), 7.89 (t, *J* = 8.4 Hz, 1H), 7.71 (td, *J* = 8.0, 1.2 Hz, 1H), 7.51 (t, *J* = 7.6 Hz, 1H), 7.39 (d, *J* = 8.8 Hz, 1H), 7.30 (s, 2H), 5.92 (d, *J* = 6.4 Hz, 1H), 4.93 (t, *J* = 5.6 Hz, 1H), 4.25 - 4.23 (m, 1H), 4.19 - 4.18 (m, 1H), 3.84 (dd, *J* = 14.0, 6.8 Hz, 1H), 3.64 (dd, *J* = 14.0, 6.8 Hz, 1H).

**(2*R*,3*R*,4*S*,5*S*)-2-(6-Aminopurin-9-yl)-5-[(6-methyl-2-quinolyl) sulfanylmethyl] tetrahydrofuran-3,4-diol (8)**

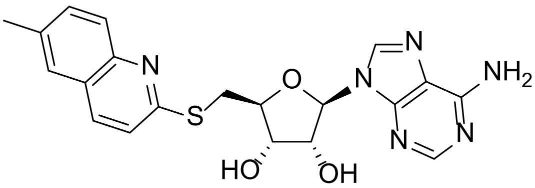

To a solution of compound **45e** (90.0 mg, 0.14 mmol) in DMF (2 mL) was added potassium fluoride (80.1 mg, 1.38 mmol). The reaction mixture was stirred at 60 °C for 16 h, cooled to RT and then concentrated under reduced pressure. The residue was purified by prep-HPLC (Method 1) to afford compound **8** (25.1 mg, 43% yield) as a white solid. HRMS (Method 4) *m/z* $[M+H]^+$ calcd for $C_{20}H_{20}N_6O_3S$ 425.1396; found 425.1416. $^1H$ NMR (400 MHz, DMSO-$d_6$): $\delta$ 8.40 (s, 1H), 8.16 (s, 1H), 8.07 (d, $J$ = 8.8 Hz, 1H), 7.78 (d, $J$ = 8.8 Hz,1H), 7.67 (s, 1H), 7.55 (dd, $J$ = 8.8, 1.6 Hz, 1H), 7.34 (d, $J$ = 8.8 Hz, 1H), 7.30 (s, 1H), 5.91 (d, $J$ = 6.4 Hz, 1H), 5.55 (s, 1H), 5.36 (s, 1H), 4.93 (s, 1H), 4.23 (s, 1H), 4.20 - 4.15 (m,1H), 3.82 (dd, $J$ = 14.0, 6.8 Hz, 1H), 3.62 (dd, $J$ = 14.0, 6.8 Hz, 1H), 2.46 (s, 3H).

**(2*R*,3*R*,4*S*,5*S*)-2-(6-Aminopurin-9-yl)-5-[(7-methyl-2-quinolyl) sulfanylmethyl] tetrahydrofuran-3,4-diol (9)**

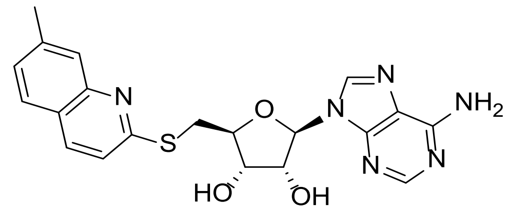

To a solution of compound **45f** (60.0 mg, 91.9 µmol) in DMF (2 mL) was added potassium fluoride (53.3 mg, 918 µmol). The reaction mixture was stirred at 60 °C for 16 h, cooled to RT and then concentrated under reduced pressure. The residue was purified by prep-HPLC (Method 3) to afford compound **9** (14.6 mg, 37% yield) as a white solid. HRMS (Method 1) *m/z* $[M+H]^+$ calcd for $C_{20}H_{20}N_6O_3S$ 425.1396; found 425.1391. $^1H$ NMR (400 MHz, DMSO-$d_6$): $\delta$ 8.32 (s, 1H), 8.08 (s, 1H), 7.90 (d, $J$ = 8.8 Hz, 1H), 7.64 (d, $J$ = 8.4 Hz, 1H), 7.62 (s, 1H), 7.30 (dd, $J$ = 8.0, 0.8 Hz, 1H), 7.22 (d, $J$ = 8.4 Hz, 1H), 6.04 (d, $J$ = 3.6 Hz, 1H), 5.74 (s, 2H), 4.69 - 4.65 (m, 2H), 4.44 (t, $J$ = 4.8 Hz, 1H), 3.97 (dd, $J$ = 14.8, 4.4 Hz, 1H), 3.62 (dd, $J$ = 14.8, 5.2 Hz, 1H), 2.52 (s, 3H).

**(2*R*,3*R*,4*S*,5*S*)-2-(6-Aminopurin-9-yl)-5-(7-quinolylsulfanylmethyl) tetrahydrofuran-3,4-diol (10)**

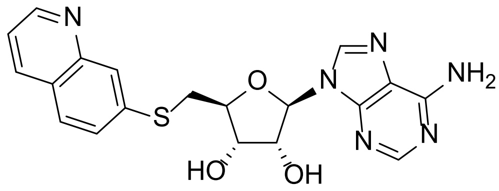

To a solution of compound **45g** (100 mg, 0.16 mmol) in MeOH (3 mL) was added potassium fluoride (90.9 mg, 1.57 mmol). The reaction mixture was stirred at 50 °C for 16 h, cooled to RT and then concentrated under reduced pressure. The residue was purified by reverse column chromatography (0.05% $NH_3 \cdot H_2O$ in water/MeCN gradient 0-25%) to afford compound **10** (29.2 mg, 45% yield) as a white solid. HRMS (Method 1) *m/z* $[M+H]^+$ calcd for $C_{19}H_{18}N_6O_3S$ 411.1239; found 411.1243. $^1H$ NMR (400 MHz, DMSO-$d_6$): $\delta$ 8.85 (dd, $J$ = 4.0, 1.6 Hz, 1H), 8.15 (s, 1H), 7.89 - 7.87 (m, 2H), 7.52 (dd, $J$ = 8.4, 1.6 Hz, 1H), 7.45 (dd, $J$ = 8.4, 4.0 Hz, 1H), 7.27 (s, 1H), 5.90 (d, $J$ = 6.0 Hz, 1H), 4.81 (t, $J$ = 5.2 Hz, 1H), 4.24 (t, $J$ = 4.4 Hz, 1H), 4.12 - 4.08 (m, 1H), 3.62 (dd, $J$ = 13.6, 5.6 Hz, 1H), 3.49 (dd, $J$ = 13.6, 7.2 Hz, 1H).

**(2*R*,3*R*,4*S*,5*S*)-2-(6-Aminopurin-9-yl)-5-(thiazol-2-ylsulfanylmethyl) tetrahydrofuran-3,4-diol (11)**

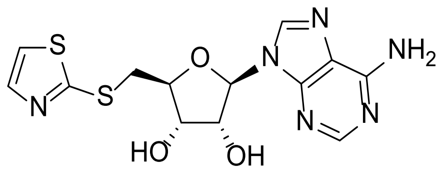

To a solution of **45h** (70.0 mg, 0.12 mmol) in MeOH (5 mL) was added potassium fluoride (68.4 mg, 1.18 mmol). The reaction mixture was stirred at 60 °C for 16 h, cooled to RT and then concentrated under reduced pressure. The residue was purified by reverse column chromatography (0.05% $NH_3 \cdot H_2O$ in water/MeCN gradient 0-45%) to afford compound **11** (18.5 mg, 43% yield) as a white solid. HRMS (Method 1) *m/z* $[M+H]^+$ calcd for $C_{13}H_{14}N_6O_3S_2$ 367.0647; found 367.0656. $^1H$ NMR (400 MHz, DMSO-$d_6$): $\delta$ 8.34 (s, 1H), 8.14 (s, 1H), 7.71 (d, $J$ = 3.2 Hz, 1H), 7.64 (d, $J$ = 3.6 Hz, 1H), 7.30 (s, 2H), 8.06 (s, 1H), 5.89 (d, $J$ = 6.0 Hz, 1H), 4.80 (t, $J$ = 5.6 Hz, 1H), 4.21 (t, $J$ = 3.6 Hz, 1H), 4.15 - 4.13 (m, 1H), 3.70 (dd, $J$ = 14.0, 5.6 Hz, 1H), 3.56 (dd, $J$ = 14.0, 7.6 Hz, 1H ).

**(2*R*,3*R*,4*S*,5*S*)-2-(6-Aminopurin-9-yl)-5-(imidazo[1,5-a]pyridin-8- ylsulfanylmethyl)**

**tetrahydrofuran-3,4-diol (12)**

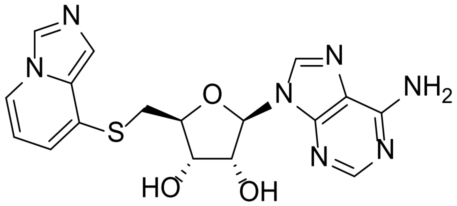

To a solution of compound **45i** (70.0 mg, 0.11 mmol) in MeOH (3 mL) was added potassium fluoride (64.7 mg, 1.11 mmol). The reaction mixture was heated to 60 °C for 16 h, cooled to RT and then concentrated under reduced pressure. The residue was purified by reverse column chromatography (0.1% $NH_3{\cdot}H_2O$ in water/MeCN gradient 0-50%) to afford compound **12** (31.3 mg, 69% yield) as an off-white solid. HRMS (Method 1) *m/z* $[M+H]^+$ calcd for $C_{17}H_{17}N_7O_3S$ 400.1192; found 400.1201. $^1H$ NMR (400 MHz, DMSO-$d_6$): $\delta$ 8.40 (s, 1H), 8.35 (s, 1H), 8.21 (d, *J* = 7.2 Hz, 1H), 8.14 (s, 1H), 7.35 (s, 1H), 7.29 (s, 2H), 6.77 (d, *J* = 6.8 Hz, 1H), 6.60 (t, *J* = 6.8 Hz, 1H), 6.03 (brs, 1H), 5.90 (d, *J* = 5.6 Hz, 1H), 4.80 (t, *J* = 5.6 Hz, 1H), 4.22 (t, *J* = 4.8 Hz, 1H), 4.07 - 4.06 (m, 1H), 3.53 (dd, *J* = 13.6, 5.2 Hz, 1H), 3.42 (dd, *J* = 13.6, 7.6 Hz, 1H).

**(2R,3R,4S,5S)-2-(6-Amino-9H-purin-9-yl)-5-(((1-methyl-1H-benzo[d]imidazol-2-yl)thio) methyl)tetrahydrofuran-3,4-diol (13)**

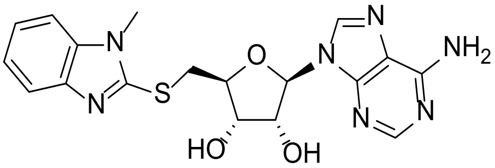

To a solution of compound **45j** (98.0 mg, 0,15 mmol) in MeOH (4 mL) was added potassium fluoride (3.28 mmol). The reaction mixture was stirred at 60 °C for 16 h, cooled to RT and then concentrated under reduced pressure. The residue was purified by reverse column chromatography (0.1% $NH_3{\cdot}H_2O$ in water/MeCN gradient 0-100%) to afford compound **13** (4.67 mg, 7% yield) as a white solid. HRMS (Method 1) *m/z* $[M+H]^+$ calcd for $C_{17}H_{17}N_7O_3S$ 414.1378; found 414.1356. $^1H$ NMR (400 MHz, DMSO-$d_6$): $\delta$ 8.40 (s, 1H), 8.16 (s, 1H), 7.55 - 7.53(m, 1H), 7.50 - 7.44 (m, 1H), 7.30 (brs, 2H), 7.22 - 7.11 (m, 2H), 5.91 (d, *J* = 6.0 M Hz, 1H), 5.54 (d, *J* = 6.0 M Hz, 1H), 5.44 (d, *J* = 4.4 M Hz, 1H), 4.88 - 4.86 (m, 1H), 4.30 - 4.24 (m, 1H), 4.23 - 4.18 (m, 1H), 3.87 - 3.79 (m, 1H), 3.74 - 3.68 (m, 1H ), 3.66 (s, 3H).

**(2R,3R,4S,5S)-2-(6-Amino-9H-purin-9-yl)-5-((benzo[d]oxazol-2-ylthio)methyl) tetrahydrofuran-3,4-diol (14)**

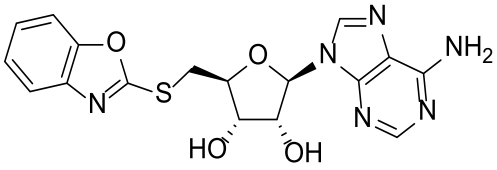

To a solution of compound **45k** (160 mg, 0,25 mmol) in MeOH (4 mL) was added potassium fluoride (119 µL, 5.09 mmol). The reaction mixture was stirred at 60 °C for 16 h, cooled to RT and then concentrated under reduced pressure. The residue was purified by reverse column chromatography (0.1% $NH_3 \cdot H_2O$ in water/MeCN gradient 0-100%) to afford compound **14** (22.0 mg, 21% yield) as a white solid. HRMS (Method 1) *m/z* $[M+H]^+$ calcd for $C_{17}H_{16}N_6O_4S$ 401.1078; found 400.1043. $^1H$ NMR (400 MHz, DMSO-$d_6$): $\delta$ 8.38 (s, 1H), 8.14 (s, 1H), 7.65 - 7.61 (m, 2H), 7.36 - 7.29 (m, 4H), 5.9 (d, *J* = 5.6 Hz, 1H), 5.73 - 5.42 (m, 1H), 4.85 (t, *J* = 5.2 Hz, 1H), 4.24 - 4.24 (m, 2H), 3.86 - 3.78 (m, 1H), 3.75 - 3.66 (m, 1H) .

**(2S,3S,4R,5R)-2-(((1H-Benzo[d]imidazol-2-yl)thio)methyl)-5-(6-amino-9H-purin-9-yl) tetrahydrofuran-3,4-diol (15)**

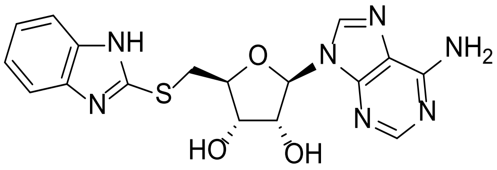

To a solution of compound **45l** (103 mg, 0.16 mmol) in MeOH (4 mL) was added potassium fluoride (77 µL, 3.28 mmol). The reaction mixture was stirred at 60 °C for 16 h, cooled to RT and then concentrated under reduced pressure. The residue was purified by reverse column chromatography (0.1% $NH_3 \cdot H_2O$ in water/MeCN gradient 0-100%) to afford compound **15** (40.5 mg, 62% yield) as a white solid. HRMS (Method 1) *m/z* $[M+H]^+$ calcd for $C_{17}H_{17}N_7O_3S$ 400.1178; found 400.1201. $^1H$ NMR (400 MHz, DMSO-$d_6$): $\delta$ 8.39 (s, 1H), 8.15 (s, 1H), 7.44 - 7.42 (m, 2H), 7.30 (s, 2H), 7.15 - 7.05 (m, 2H), 5.97 (d, *J* = 6 Hz, 1H), 5.53 - 5.52 (m, 1H), 4.84 - 4.83 (m, 1H), 4.25 - 4.19 (m, 2H), 3.80 - 3.73 (m, 1H), 3.67 - 3.62 (m, 1H).

**(2S,3S,4R,5R)-2-(((1H-Indazol-7-yl)thio)methyl)-5-(6-amino-9H-purin-9-yl)tetrahydrofuran-3,4-diol (16)**

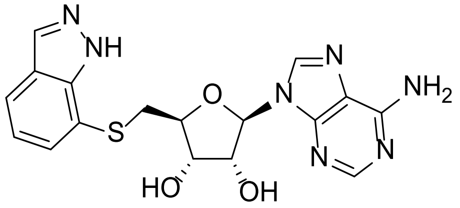

To a solution of compound **45m** (10.0 mg, 15.9 µmol) in MeOH (2 mL) was added potassium fluoride (7.5 µL, 319 µmol). The reaction mixture was stirred at 60 °C for 16 h, cooled to RT and then concentrated under reduced pressure. The residue was purified by reverse column chromatography (0.1% $NH_3 \cdot H_2O$ in water/MeCN gradient 0-100%) to afford compound **16** (3.86 mg, 61%). HRMS (Method 1) *m/z* $[M+H]^+$ calcd for $C_{17}H_{17}N_7O_3S$ 400.4378; found 400.1187. $^1H$ NMR (400 MHz, DMSO-*d*$_6$): *δ* 8.36 (s, 1H), 8.17 - 8.13 (m, 2H), 7.69 - 7.66 (m, 1H), 7.41 - 7.38 (m, 1H), 7.31 - 7.28 (s, 2H), 7.07 (t, *J* = 7.6Hz, 1H), 5.89 (d, *J* = 6Hz, 1H), 4.82 (t, *J* = 5.6Hz, 1H), 4.23 - 4.81 (m, 1H), 4.02 - 3.98 (m, 1H), 3.58 - 3.55 (m, 2H).

**(2*R*,3*R*,4*S*,5*S*)-2-(6-Aminopurin-9-yl)-5-(imidazo[1,5-a]pyridine-7-ylsulfanylmethyl) tetrahydrofuran-3,4-diol (17)**

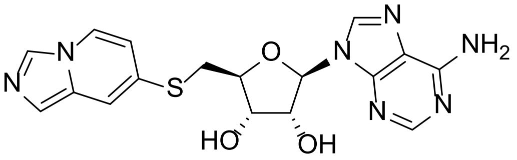

To a solution compound **45n** (100 mg, 0.16 mmol) in MeOH (2 mL) was added potassium fluoride (92.5 mg, 1.59 mmol). The reaction mixture was stirred at 60 °C for 16 h, cooled to RT and concentrated under reduced pressure. The residue was purified by reverse column chromatography (100 Å; 0.05% $NH_3 \cdot H_2O$ in water/MeCN gradient 0-25%) to afford compound **17** (22.2 mg, 33% yield) as a green solid. HRMS (Method 1) *m/z* $[M+H]^+$ calcd for $C_{17}H_{17}N_7O_3S$ 400.1192; found 400.1176. $^1H$ NMR (400 MHz, DMSO-*d*$_6$): *δ* 8.36 (s, 1H), 8.27 (s, 1H), 8.24 (d, *J* = 7.6 Hz, 1H), 8.15 (s, 1H), 7.43 (s, 1H), 7.29 (s, 2H), 7.16 (s, 1H), 6.57 (dd, *J* = 7.2, 1.6 Hz, 1H), 5.90 (d, *J* = 6.0 Hz, 1H), 5.82 (s, 1H), 4.80 (t, *J* = 5.6 Hz, 1H), 4.22 (t, *J* = 4.0 Hz, 1H), 4.06 - 4.02 (m, 1H), 3.43 (dd, *J* = 13.6, 5.6 Hz, 1H), 3.36 - 3.34 (m, 1H).

**(2*R*,3*R*,4*S*,5*R*)-2-(6-aminopurin-9-yl)-5-(1,3-benzothiazol-2-yloxymethyl)tetrahydrofuran-3,4-diol (23)**

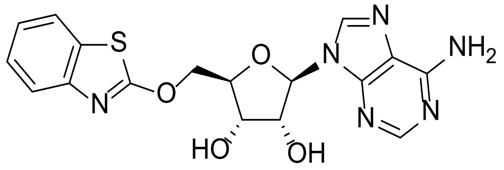

To a solution of compound **45o** (30.0 mg, 47.7 µmol) in THF (3 mL) was added *N*,*N*-diethylethanamine trihydrofluoride (23.1 mg, 143 µmol). The reaction mixture was stirred at RT for 16 h and then concentrated under reduced pressure. The residue was purified by prep-HPLC (Method 5) to afford compound **23** (8.15 mg, 42% yield) as a white solid. HRMS (Method 1) *m/z* $[M+H]^+$ calcd for $C_{17}H_{16}N_6O_4S$ 401.1032; found 401.1031. $^1H$ NMR (400 MHz, DMSO-*d$_6$*): 8.32 (s, 1H), 8.13 (s, 1H), 7.61 (d, *J* = 7.6 Hz, 1H), 7.32 (s, 1H), 7.29 (d, *J* = 5.6 Hz, 1H), 7.17 (t, *J* = 7.6 Hz, 1H), 7.13 (t, *J* = 7.6 Hz, 1H), 6.87 (d, *J* = 5.2 Hz, 1H), 5.57 (d, *J* = 6.0 Hz, 1H), 5.42 (d, *J* = 5.2 Hz, 1H), 4.77 (q, *J* = 5.2 Hz, 1H), 4.35 (t, *J* = 9.2 Hz, 1H), 4.30 - 4.28 (m, 1H), 4.25 - 4.23 (m, 2H).

**2*R*,3*R*,4*S*,5*R*)-2-(6-Aminopurin-9-yl)-5-[(1,3-benzothiazol-2-ylamino)methyl] tetrahydrofuran-3,4-diol (24)**

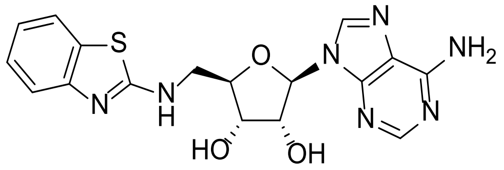

To a solution of **45p** (35.0 mg, 55.7 µmol) in THF (4 mL) was added *N*,*N*-diethylethanamine trihydrofluoride (17.9 mg, 111 µmol). The reaction mixture was stirred at RT for 16 h and then concentrated under reduced pressure. The residue was purified by prep-HPLC (Method 6), followed by (Method 7) to afford compound **24** (4.01 mg, 17% yield) as a white solid. HRMS (Method 1) *m/z* $[M+H]^+$ calcd for $C_{17}H_{17}N_7O_3S$ 400.1192; found 400.1197. $^1H$ NMR (400 MHz, DMSO-*d$_6$*): $\delta$ 9.45 (s, 1H), 8.33 (s, 1H), 8.14 (s, 1H), 7.45 (d, *J* = 7.2 Hz, 1H), 7.32 (d, *J* = 8.4 Hz, 1H), 7.29 (s, 2H), 7.09 (dd, *J* = 8.0, 0.8 Hz, 1H), 6.97 (t, *J* = 7.2 Hz, 1H), 5.88 (d, *J* = 5.6 Hz, 1H), 5.49 (d, *J* = 6.0 Hz, 1H), 5.36 (s, 1H), 4.81 (q, *J* = 5.2 Hz, 1H), 4.17 (t, *J* = 3.6 Hz, 1H), 3.94 - 3.93 (m, 1H), 3.23 - 3.19 (m, 2H).

#### Scheme S2

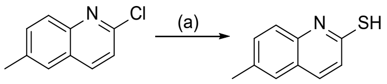

72 73

(a) thiourea, EtOH, 80 °C, 16 h, 63%

##### 6-Methylquinoline-2-thiol (73)

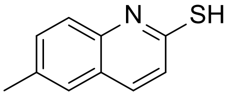

To a solution of 2-chloro-6-methyl-quinoline **72** (200 mg, 1.13 mmol) in EtOH (4 mL) was added thiourea (257 mg, 3.38 mmol) and the reaction mixture was stirred at 80 °C for 16 h. The mixture was cooled to RT, filtered, and the resulting filter-cake was dried under reduced pressure to afford compound **73** (125 mg, 63% yield) as a yellow solid. The isolated material was used in the following step without any further purification. LCMS ($ES^+$, Method 3) *m/z* 176.1 $[M+H]^+$. $^1H$ NMR (400 MHz, DMSO-$d_6$): $\delta$ 13.62 (s, 1H), 7.76 (d, *J* = 8.8 Hz, 1H), 7.57 - 7.45 (m, 3H), 7.23 (d, *J* = 9.2 Hz, 1H), 2.37 (s, 3H).

#### Scheme S3

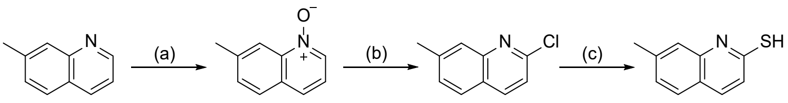

74 75 76 77

(a) m-CPBA, DCM, RT, 16 h, 72%; (b) $POCl_3$, toluene, 70 °C, 16 h, 68%; (c) thiourea, EtOH, 80 °C, 16 h, 76%

##### 7-Methyl-1-oxido-quinolin-1-ium (75)

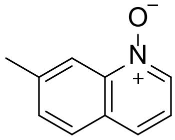

To a solution of 7-methylquinoline **74** (1.00 g, 6.98 mmol) in DCM (15 mL) was added meta-chloroperoxybenzoic acid (2.13 g, 10.5 mmol, 85% purity) and stirred at RT for 16 h. The reaction mixture was quenched with sat. aq. sodium bicarbonate solution (20 mL) and extracted with DCM (3 x 10mL). The combined organic layers were dried over sodium sulfate, filtered, and the filtrate was concentrated under reduced pressure. The residue was purified by reverse column chromatography (0.05% $NH_3{\cdot}H_2O$ in water/MeCN gradient 0-5%) to afford compound **75** (800 mg, 72% yield) as a brown solid. LCMS ($ES^+$, Method 2) *m/z* 160.1 $[M+H]^+$. $^1H$ NMR (400 MHz, DMSO-$d_6$): $\delta$ 8.54 (d, $J$ = 6.0 Hz, 1H), 8.35 (s, 1H), 7.98 (d, $J$ = 8.4 Hz, 1H), 7.88 (d, $J$ = 8.4 Hz, 1H), 7.57 (d, $J$ = 8.4 Hz, 1H), 7.41 - 7.38 (m, 1H), 2.55 (s, 3H).

**2-Chloro-7-methyl-quinoline (76)**

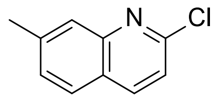

To a solution of compound **75** (500 mg, 3.14 mmol) in toluene (8 mL) was added phosphoryl chloride (583 µL, 6.28 mmol), and stirred at 70 °C for 16 h. The reaction mixture was cooled to 0 °C, quenched with water (20 mL), and the aqueous layer extracted with EtOAc (3 x 10 mL). The combined organic layers were washed with brine (3 x 10 mL), dried over sodium sulfate, filtered, and the filtrate was concentrated under reduced pressure. The residue was purified by flash column chromatography (petroleum ether/ethyl acetate gradient 2-50%) to afford compound **76** (380 mg, 68% yield) as a white solid. LCMS ($ES^+$, Method 2) *m/z* 340.5 $[M+H]^+$. $^1H$ NMR (400 MHz, $CDCl_3$): $\delta$ 8.05 (d, $J$ = 8.4 Hz, 1H), 7.80 (s, 1H), 7.71 (d, $J$ = 8.4 Hz, 1H), 7.40 (dd, $J$ = 8.0, 0.8 Hz, 1H), 7.32 (d, $J$ = 8.8 Hz, 1H), 2.57 (s, 3H).

**7-Methylquinoline-2-thiol (77)**

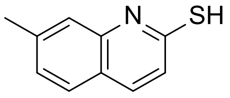

To a solution of compound **76** (200 mg, 1.13 mmol) in EtOH (3 mL) was added thiourea (257 mg, 3.38 mmol) and stirred at 80 °C for 16 h. The reaction mixture was cooled to RT and filtered. The filter-cake was collected and dried under reduced pressure to afford compound **77** (150 mg, 76% yield) as a yellow solid. The isolated material was used in the following step without any further purification. LCMS ($ES^+$, Method 4) *m/z* 176.1 $[M+H]^+$. $^1H$ NMR (400 MHz,

DMSO-$d_6$): $\delta$ 13.58 (s, 1H), 7.78 (d, $J$ = 8.8 Hz, 1H), 7.66 (d, $J$ = 8.0 Hz, 1H), 7.40 (s, 1H), 7.20 - 7.17 (m, 2H), 2.41 (s, 3H).

**Scheme S4**

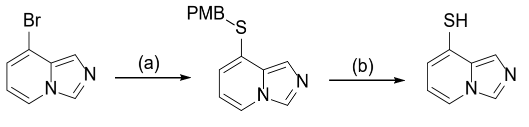

78 79 80

(a) (4-methoxyphenyl) methanethiol, $Pd_2(dba)_3$, Xantphos, $Cs_2CO_3$, DMF, 100 °C, 16 h, 55%; (b) TFA, 70 °C, 16 h, 72%

**8-[(4-methoxyphenyl)methylsulfanyl]imidazo[1,5-a]pyridine (79)**

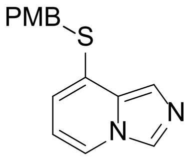

To a solution of 8-bromoimidazo[1,5-a]pyridine **78** (200 mg, 1.02 mmol) in DMF (5 mL) was added (4-methoxyphenyl)methanethiol (172 mg, 1.12 mmol), $Pd_2(dba)_3$ (46.5 mg, 0.05 mmol), Xantphos (58.7 mg, 0.10 mmol) and cesium carbonate (992 mg, 3.05 mmol) under inert atmosphere. The resulting mixture was stirred at 100 °C for 16 h. The reaction mixture was cooled to RT, quenched with water (50 mL) and extracted with EtOAc (3 x 20 mL). The combined organic layers were washed with brine (3 x 20 mL), dried over sodium sulfate, filtered, and concentrated under reduced pressure. The residue was purified by flash column chromatography (petroleum ether/ethyl acetate gradient 2-50%) to afford compound **79** (150 mg, 55% yield) as a brown solid. LCMS (ES$^+$, Method 1) $m/z$ 271.1 [M+H]$^+$. $^1$H NMR (400 MHz, $CDCl_3$): $\delta$ 8.13 (s, 1H), 7.80 (d, $J$ = 6.8 Hz, 1H), 7.54 (s, 1H), 7.24 (d, $J$ = 8.8 Hz, 2H), 6.86 - 6.82 (m, 2H), 6.58 (d, $J$ = 6.4 Hz, 1H), 6.48 (t, $J$ = 7.2 Hz, 1H), 4.18 (s, 2H), 3.80 (s, 3H).

**Imidazo[1,5-a]pyridine-8-thiol (80)**

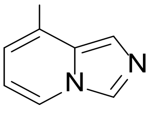

A solution of compound **79** (150 mg, 555 µmol) in trifluoroacetic acid (2 mL) was stirred at 70 °C for 16 h, cooled to RT and then concentrated under reduced pressure. The residue was purified by reverse column chromatography (0.05% TFA in water/MeCN gradient 0%-15%) to afford compound **80** (60.0 mg, 72% yield) as brown solid. $^1$H NMR (400 MHz, $CDCl_3$): $\delta$ 8.25 (s, 1H), 7.90 (d, $J$ = 7.2 Hz, 1H), 7.60 (s, 1H), 6.96 (d, $J$ = 6.8 Hz, 1H), 6.55 (t, $J$ = 6.8 Hz, 1H).

**Scheme S5**

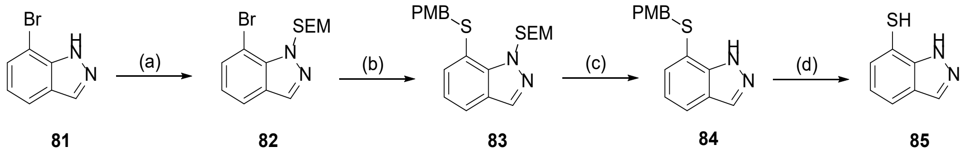

(a) SEM-Cl, NaH, THF, RT, 16 h, 66%; (b) (4-methoxyphenyl) methanethiol, $Pd_2(dba)_3$, Xantphos, $Cs_2CO_3$, dioxane, 100 °C, 16 h, 74%; (c) TBAF, 70 °C, 16 h, 22%; (d) TFA, 70 °C, 16 h, 50%

**7-Bromo-1-((2-(trimethylsilyl)ethoxy)methyl)-1H-indazole (82)**

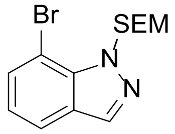

To a solution of 7-bromo-1H-indazole **81** (2.00 g, 10.2 mmol) in THF (20 mL) was added NaH (487 mg, 12.2 mmol, 60% dispersion in mineral oil) under inert atmosphere at 0 °C and stirred for 30 min. 2-(Trimethylsilyl)ehtoxymethyl chloride (2.16 mL, 12.2 mmol) was then added and stirred at RT for 16 h. The reaction mixture was quenched with water (100 mL) and extracted with EtOAc (3 × 50 mL). The combined organic layers were washed with brine (50 mL), dried over sodium sulfate, filtered, and concentrated under reduced pressure. The residue was purified by purified by flash column chromatography (petroleum ether/ethyl acetate gradient 0-30%) to afford compound **82** (2.2 g, 66% yield) as a yellow liquid. LCMS (ES$^+$, Method 2) *m/z* 269.0 [M+H]$^+$. $^1$H NMR (400 MHz, $CD_3OD$): $\delta$ 8.04 (s, 1H), 7.71 - 7.68 (m, 1H), 7.62 - 7.60 (m, 1H), 7.05 (t, $J$ = 7.6 M Hz, 1H), 6.09 (s, 2H), 3.59 (t, $J$ = 8.0 M Hz, 2H), 0.90 (t, $J$ = 8.4 M Hz, 2H), -0.067 (s, 9H).

**7-((4-Methoxybenzyl)thio)-1-((2-(trimethylsilyl)ethoxy) methyl)- 1H-indazole (83)**

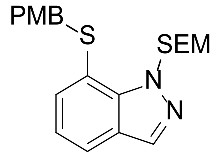

To a solution of compound **82** (2.20 g, 6.72 mmol) in DMF (20 mL) was added (4-methoxyphenyl) methanethiol (1.12 mL, 8.07 mmol), $Pd_2(dba)_3$ (308 mg, 0,34 mmol), Xantphos (389 mg, 0.67 mmol) and cesium carbonate (6.57 g, 20.2 mmol) under inert atmosphere. The resulting mixture was heated at 100 °C for 16 h. The reaction mixture was cooled to RT, quenched with water (20 mL) and then extracted with EtOAc (3 × 20 mL). The combined organic layers were washed with brine (20 mL), dried over sodium sulphate, filtered, and concentrated under reduced pressure. The residue was purified by flash column chromatography (petroleum ether/ethyl acetate gradient 0-20%) to afford compound **83** (2.00 g, 74% yield) as yellow liquid. LCMS ($ES^+$, Method 1) *m/z* 400.9 $[M+H]^+$.

**S7-((4-Methoxybenzyl)thio)-1H-indazole (84)**

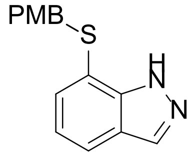

A mixture of compound **83** (1.66 g, 3.25 mmol) and TBAF (15 mL, 10.0 mmol, 1 M in THF) was stirred at 70 °C for 16 h. The reaction mixture was cooled to RT, quenched with water (15 mL) and extracted with EtOAc (3 × 15 mL). The combined organic layers were washed with brine (15 mL), dried over sodium sulphate, filtered and concentrated under reduced pressure. The residue was purified by flash column chromatography (petroleum ether/ethyl acetate gradient 0-20%) to afford compound **84** (360 mg, 22% yield) as a yellow solid. LCMS ($ES^+$, Method 1) *m/z* 271.2 $[M+H]^+$.

**1H-Indazole-7-thiol (85)**

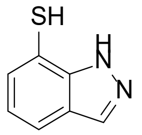

A solution of compound **84** (360 mg, 1.33 mmol) in trifluoroacetic acid (3.00 mL, 40.5 mmol) was stirred at 70 °C for 16 h. The reaction mixture was cooled to RT concentrated under reduced pressure and the residue was purified by flash column chromatography (SWPAFLASH® SW040 Bonded Spherical C18, 20-45µm, 100 Å; 0.1% TFA in water/MeCN gradient 0-100%) to afford compound **85** (100 mg, 50% yield) as a yellow solid. LCMS ($ES^+$, Method 3) *m/z* 151.1 $[M+H]^+$. $^1H$ NMR (400 MHz, $CD_3OD$): $\delta$ 8.14 (s, 1H), 7.81 - 7.79 (m, 1H), 7.11 - 7.09 (m, 1H) 7.00 - 6.96 (m, 1H).

**Scheme S6**

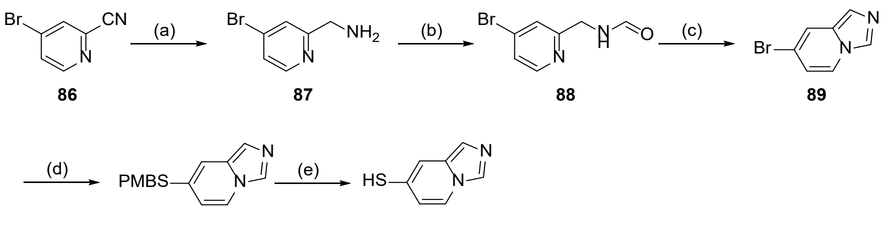

(a) $BH_3$-THF, THF, RT, 16 h, 49%; (b) $HCO_2H$, 100 °C, 3 h; (c) TFAA, DCM, RT, 1 h, 43%; (d) (4-methoxyphenyl)methanethiol, $Pd_2(dba)_3$, Xantphos, $Cs_2CO_3$, dioxane, 100 °C, 16h, 85%, (e) TFA, 70 °C, 16 h, 67%

##### (4-Bromo-2-pyridyl)methanamine (87)

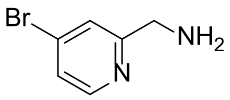

To a solution of 4-bromopyridine-2-carbonitrile **86** (3.00 g, 16.3 mmol) in THF (30 mL) was added dropwise borane (75.0 mL, 1 M in THF) under inert atmosphere and stirred at RT for 16 h. The reaction mixture was cooled to 0 °C, HCl (75.0 mL, 2M aq.) was then added and heated to 80 °C for 30 min. The reaction mixture was cooled to RT, basified to pH = 8 using 2M aq. sodium hydroxide and then extracted with EtOAc (3 x 30 mL). The combined organic layers were washed with brine (30 mL), dried over sodium sulfate, filtered, and the filtrate was concentrated under reduced pressure. The residue was purified by reverse column chromatography (0.05% $NH_3 \cdot H_2O$ in water/MeCN gradient 0-30%) to afford compound **87** (1.50 g, 48.9% yield) as yellow oil. LCMS (ES$^+$, Method 5) *m/z* 189.0 [M+H]$^+$. $^1$H NMR (400 MHz, DMSO-$d_6$): *δ* 8.37 (d, *J* = 5.6 Hz, 1H), 7.74 (d, *J* = 1.6 Hz, 1H), 7.51 (dd, *J* = 5.2, 2.0 Hz, 1H), 3.82 (s, 2H).

**N-[(4-Bromo-2-pyridyl)methyl]formamide (88)**

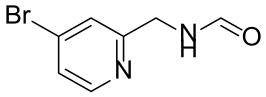

A solution of (4-bromo-2-pyridyl)methanamine **87** (1.5 0g, 8.02 mmol) in formic acid (75.00 mL, 1.99 mol) was heated to 100 °C and stirred for 3 h. The mixture was cooled to RT and concentrated under reduced pressure. The residue was dissolved in sat. aq. sodium bicarbonate solution (50 mL) and extracted with DCM (3 x 50 mL). The combined organic layers were concentrated under reduced pressure to afford crude compound **88** which was used in the following step without any further purification (1.70 g) as a yellow solid. $^1$H NMR (400 MHz, $CDCl_3$): δ 8.37 (d, J = 5.2 Hz, 1H), 8.34 (s, 1H), 7.49 (s, 1H), 7.41 (d, J = 5.2 Hz, 1H), 4.60 (d, J = 5.2 Hz, 2H).

**7-Bromoimidazo[1,5-a]pyridine (89)**

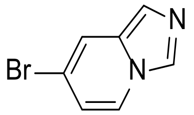

To a solution of compound **88** (1.7 g, 7.91 mmol) in DCM (20 mL) was added trifluoroacetic anhydride (1.21 mL, 8.70 mmol) at 0 °C. The reaction mixture was warmed gradually to RT and stirred for 1 h. The reaction was quenched with sat. aq. sodium bicarbonate solution (2 mL) and water (10 mL), and the mixture was extracted with DCM (3 x 5 mL). The combined organic layers were dried over sodium sulfate, filtered, and the filtrate was concentrated under reduced pressure. The residue was purified by flash column chromatography (petroleum ether/ethyl acetate gradient 2-50%) to afford compound **89** (680 mg, 43% yield) as yellow oil. $^1$H NMR (400 MHz, $CDCl_3$): δ 8.09 (s, 1H), 7.80 (d, J = 7.2 Hz, 1H), 7.64 (s, 1H), 7.37 (s, 1H), 6.62 (dd, J = 7.2, 1.2 Hz, 1H).

**7-[(4-Methoxyphenyl)methylsulfanyl]imidazo[1,5-a]pyridine (90)**

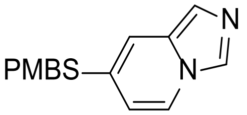

To a solution of compound **89** (680 mg, 3.45 mmol) in DMF (10 mL) was added (4-methoxyphenyl)methanethiol (527 µL, 3.80 mmol), $Pd_2(dba)_3$ (158 mg, 0.17 mmol), Xantphos (199mg, 0.35 mmol) and cesium carbonate (3.37 g, 10.3 mmol) under inert atmosphere and stirred at 100 °C for 16 h. The reaction mixture was cooled to RT, quenched with water (20 mL) and extracted with EtOAc (3 x 10 mL). The combined organic layers were washed with brine (3 x 10 mL), dried over sodium sulfate, filtered and the filtrate was concentrated under reduced pressure. The residue was purified by flash column chromatography (petroleum ether/ethyl acetate gradient 2-50%) to afford compound **90** (800 mg, 85% yield) as yellow oil. LCMS ($ES^+$, Method 6 ) *m/z* 271.1 $[M+H]^+$. $^1H$ NMR (400 MHz, $CDCl_3$): δ 8.02 (d, *J* = 4.4 Hz, 2H), 7.80 (d, *J* = 7.6 Hz, 1H), 7.28 (d, *J* = 5.2 Hz, 1H), 7.25 - 7.22 (m, 2H), 6.85 - 6.83 (m, 2H), 6.44 (dd, *J* = 7.2, 1.6 Hz, 1H), 4.08 (s, 2H), 3.79 (s, 3H).

**Imidazo[1,5-a]pyridine-7-thiol (91)**

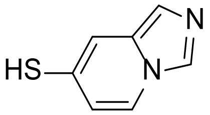

A solution of compound **90** (800 mg, 2.96 mmol) in trifluoroacetic acid (16.00 mL 216.1 mmol) was heated to 70 °C and stirred for 16 h. The reaction mixture was cooled to RT and concentrated under reduced pressure. The residue was purified by reverse column chromatography (0.05% TFA in water/MeCN gradient 0-10%) to afford compound **91** (300 mg, 67% yield) as a yellow solid. $^1H$ NMR (400 MHz, DMSO-$d_6$): *δ* 9.00 (s, 1H), 8.46 (d, *J* = 8.4 Hz, 1H), 7.90 (s, 1H), 7.70 (s, 1H), 7.01 (d, *J* = 7.6 Hz, 1H).

##### 1.7.2 Synthesis of compounds 4 and 6

**Scheme S7**

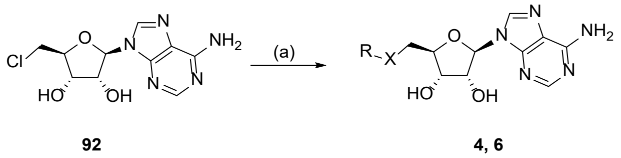

(a) R-XH, NaO$^t$Bu, DMF, 75 °C, 10 h, 44-67%.

**(2R,3R,4S,5S)-2-(6-Amino-9H-purin-9-yl)-5-((hexylthio)methyl)tetrahydrofuran-3,4-diol (4)**

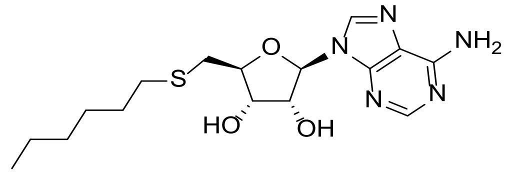

To a solution of (2R,3R,4S,5S)-2-(6-amino-9H-purin-9-yl)-5-(chloromethyl)tetrahydrofuran-3,4-diol **66** (500 mg, 1.75 mmol) in DMF (10 mL) was added hexane-1-thiol (494 µL, 3.50 mmol), potassium t-butoxide (393 mg, 3.50 mmol) and stirred at 75 °C for 10 h . The reaction mixture was cooled to RT and quenched with water. The resulting suspension was filtered off and the isolated solid material was dissolved in DMF. To the solution was then added water and the resulting precipitate was filtered off and washed with MeCN to afford compound **4** (222 mg, 34% yield) as a white solid. LCMS (ES$^+$) *m/z* 368.4 $[M+H]^+$. $^1$H NMR (400 MHz, DMSO-$d_6$): $\delta$ 8.35 (s, 1H) 8.14 (s, 1H) 7.29 (s, 2H) 5.88 (d, *J* = 5.58 Hz, 1H) 5.49 (d, *J* = 5.83 Hz, 1H) 5.31 (d, *J* = 5.07 Hz, 1H) 4.77 (d, *J* = 5.32 Hz, 1H) 4.10 - 4.19 (m, 1H) 3.92 - 4.05 (m, 1H) 2.74 - 2.94 (m, 2H) 2.45 - 2.49 (m, 2H) 1.36 - 1.52 (m, 2H) 1.09 - 1.31 (m, 6H) 0.72 - 0.89 (m, 3H).

**(2R,3R,4S,5S)-2-(6-Amino-9H-purin-9-yl)-5-((naphthalen-2-ylthio)methyl)tetrahydrofuran-3,4-diol (6)**

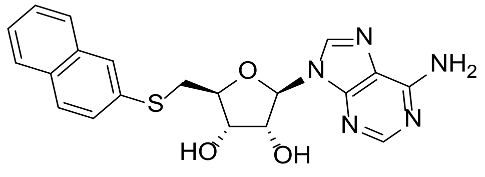

To a solution of (2R,3R,4S,5S)-2-(6-amino-9H-purin-9-yl)-5-(chloromethyl)tetrahydrofuran-3,4-diol **92** (500 mg, 1.75 mmol) in DMF (10 mL) was added 2-naphthalenethiol (561 mg, 3.50 mmol), potassium t-butoxide (393 mg, 3.50 mmol) and stirred at 75 °C for 10 h . The reaction mixture was cooled to RT and quenched with water. The resulting suspension was filtered off and the isolated solid material was dissolved in DMF. To the solution was then added water and the resulting precipitate was filtered off and washed with MeCN to afford compound **6** (480 mg, 67% yield) as a white solid. LCMS (ES$^+$) *m/z* 410.4 [M+H]$^+$. $^1$H NMR (400 MHz, DMSO-$d_6$): $\delta$ 8.39 (s, 1 H) 8.16 (s, 1 H) 7.81 - 7.89 (m, 3 H) 7.75 - 7.80 (m, 1 H) 7.42 - 7.53 (m, 3 H) 7.32 (s, 2 H) 5.90 (d, *J* = 5.83 Hz, 1 H) 5.54 (d, *J* = 6.34 Hz, 1 H) 5.40 (d, *J* = 5.07 Hz, 1 H) 4.80 - 4.89 (m, 1 H) 4.19 - 4.29 (m, 1 H) 3.99 - 4.11 (m, 1 H) 3.50 - 3.59 (m, 1 H) 3.39 - 3.49 (m, 1 H).

##### 1.7.3 Synthesis of compounds 18 and 19

Scheme S8

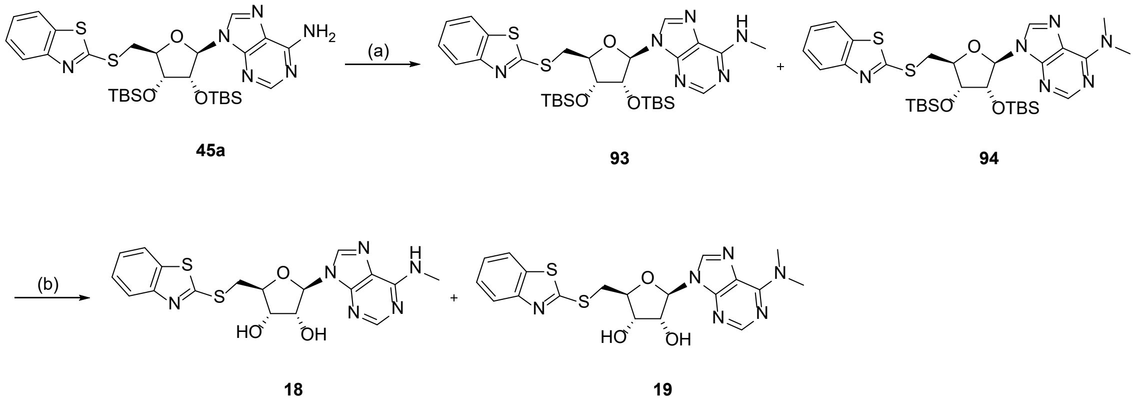

(a) MeI, $Cs_2CO_3$, DMF, RT, 16 h, 58%; (b) $Et_3N{\cdot}3HF$, THF, RT, 16 h, 26-59%.

**9-[(2*R*,3*R*,4*R*,5*S*)-5-(1,3-Benzothiazol-2-ylsulfanylmethyl)-3,4-bis[[tert-butyl(dimethyl) silyl] oxy]tetrahydrofuran-2-yl]-*N*-methyl-purin-6-amine (93) and 9-[(2*R*,3*R*,4*R*,5*S*)-5-(1,3-Benzothiazol-2-ylsulfanylmethyl)-3,4-bis[[tert-butyl(dimethyl) silyl]oxy]tetrahydrofuran-2-yl]-*N*,*N*-dimethyl-purin-6-amine (94)**

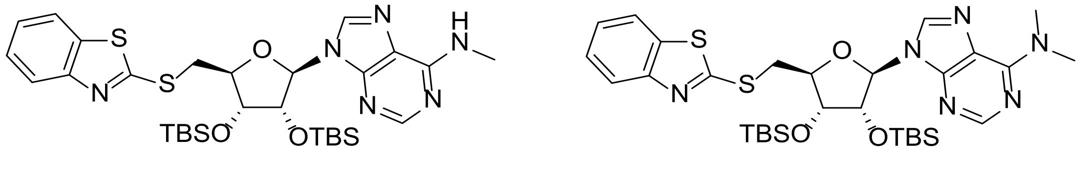

To a solution of compound **45a** (50.0 mg, 77.5 µmol) in DMF (1 mL) was added cesium carbonate (25.2 mg, 77.5 µmol) and methyl iodide (4.83 µL, 77.5 µmol) and stirred at RT for 16 h. The reaction mixture was quenched with water (20 mL) and extracted with EtOAc (3 x 30

mL). The combined organic layers were washed with brine (3 x 20 mL), dried over sodium sulfate, filtered, and the filtrate was concentrated under reduced pressure. The residue was purified by flash column chromatography (petroleum ether/ethyl acetate gradient 2-50%) to afford compound **93** (30.0 mg, 58% yield) as yellow oil and compound **94** (30.0 mg, 58% yield) as yellow oil. Compound **93** LCMS (ES$^+$, Method 2) *m/z* 659.5 $[M+H]^+$. $^1$H NMR (400 MHz, $CDCl_3$): *δ* 8.41 (s, 1H), 7.91 (s, 1H), 7.85 (d, *J* = 7.6 Hz, 1H), 7.76 (d, *J* = 8.0 Hz, 1H), 7.42 (t, *J* = 7.6 Hz, 1H), 7.31 (t, *J* = 7.6 Hz, 1H), 5.87 (d, *J* = 6.8 Hz, 1H), 5.82 (s, 1H), 5.34 (dd, *J* = 5.6, 4.4 Hz, 1H), 4.50 (t, *J* = 6.0 Hz, 1H), 4.39 (d, *J* = 2.8 Hz, 1H), 4.05 (dd, *J* = 14.0, 6.4 Hz, 1H), 3.89 (dd, *J* = 14.0, 6.8 Hz, 1H), 3.23 (s, 3H), 0.93 (s, 9H), 0.76 (s, 9H), 0.12 (s, 6H), -0.12 (s, 3H), -0.38 (s, 3H). Compound **94** LCMS (ES$^+$, Method 3) *m/z* 673.2 $[M+H]^+$. $^1$H NMR (400 MHz, $CDCl_3$): *δ* 8.47 (s, 1H), 8.25 (s, 1H), 8.00 (d, *J* = 7.6 Hz, 1H), 7.81 (d, *J* = 8.0 Hz, 1H), 7.48 (td, *J* = 8.4, 1.2 Hz, 1H), 7.37 (td, *J* = 8.4, 1.2 Hz, 1H), 5.97 (d, *J* = 7.6 Hz, 1H), 5.27 (dd, *J* = 7.6, 4.4 Hz, 1H), 4.45 (d, *J* = 4.4 Hz, 1H), 4.32 (t, *J* = 6.8 Hz, 1H), 4.07 - 4.04 (m, 1H), 3.80 (dd, *J* = 14.0, 7.2 Hz, 1H), 3.51 (s, 6H), 0.88 (s, 9H), 0.68 (s, 9H), 0.08 (s, 6H), -0.13 (s, 3H), -0.45 (s, 3H).

**(2*S*,3*S*,4*R*,5*R*)-2-(1,3-Benzothiazol-2-ylsulfanylmethyl)-5-[6-(methylamino)purin-9-yl]tetrahydrofuran-3,4-diol (18)**

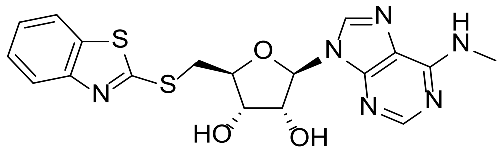

To a solution of compound **93** (30.0 mg, 45.5 µmol) in THF (2 mL) was added N,N-diethylethanamine trihydrofluoride (22.0 mg, 136 µmol) and stirred at 50 °C for 16 h. The reaction mixture was cooled to RT and concentrated under reduced pressure. The residue was purified by prep-HPLC (Method 7) to afford compound **18** (5.10 mg, 26% yield) as a white solid. HRMS (Method 1) *m/z* $[M+H]^+$ calcd for $C_{18}H_{18}N_6O_3S_2$ 431.0960; found 431.0970; $^1$H NMR (400 MHz, DMSO-$d_6$): *δ* 8.37 (s, 1H), 8.24 (s, 1H), 7.99 (d, *J* = 8.0 Hz, 1H), 7.85 (d, *J* = 8.0 Hz, 1H), 7.78 (s, 1H), 7.46 (td, *J* = 8.4, 1.2 Hz, 1H), 7.36 (td, *J* = 7.6, 1.2 Hz, 1H), 5.92 (d, *J* = 5.6 Hz, 1H), 5.56 (d, *J* = 6.0 Hz, 1H), 5. 46 (d, *J* = 4.8 Hz, 1H), 4.86 (q, *J* = 5.2 Hz, 1H), 4.29 - 4.22 (m, 2H), 3.86 (dd, *J* = 13.6, 5.6 Hz, 1H), 3.75 (dd, *J* = 13.6, 7.6 Hz, 1H), 3.31 (s, 3H).

**(2*R*,3*R*,4*S*,5*S*)-2-(6-Aminopurin-9-yl)-5-(1,3-benzothiazol-2-ylsulfanylmethyl) tetrahydrofuran-3,4-diol (19)**

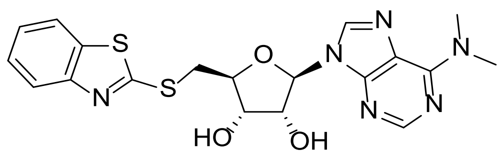

To a solution of compound **94** (200 mg, 297 µmol) in THF (5 mL) was added N,N-diethylethanamine;trihydrofluoride (143 mg, 891 µmol) and stirred at 50 °C for 16 h. The reaction mixture was cooled to RT and concentrated under reduced pressure. The residue was purified by reverse column chromatography (0.05% $NH_3{\cdot}H_2O$ in water/MeCN gradient 0-5%) and then triturated with MeOH (10 mL) to afford compound **19** (80.6 mg, 59% yield) as a white solid. HRMS (Method 1) *m/z* $[M+H]^+$ calcd for $C_{19}H_{20}N_6O_3S_2$ 445.1117; found 445.1135. $^1H$ NMR (400 MHz, DMSO-$d_6$): $\delta$ 8.40 (s, 1H), 8.23 (s, 1H), 8.00 (d, *J* = 7.6 Hz, 1H), 7.85 (d, *J* = 8.0 Hz, 1H), 7.47 (td, *J* = 8.4, 1.2 Hz, 1H), 7.36 (td, *J* = 8.4, 1.2 Hz, 1H), 5.95 (d, *J* = 5.6 Hz, 1H), 5.47 (d, *J* = 5.2 Hz, 1H), 4.84 (q, *J* = 5.2 Hz, 1H), 4.29 - 4.23 (m, 2H), 3.86 (dd, *J* = 13.6, 5.2 Hz, 1H), 3.74 (dd, *J* = 13.6, 7.6 Hz, 1H), 3.46 (brs, 6H).

##### 1.7.4 Synthesis of compounds 20-22

**Scheme S9**

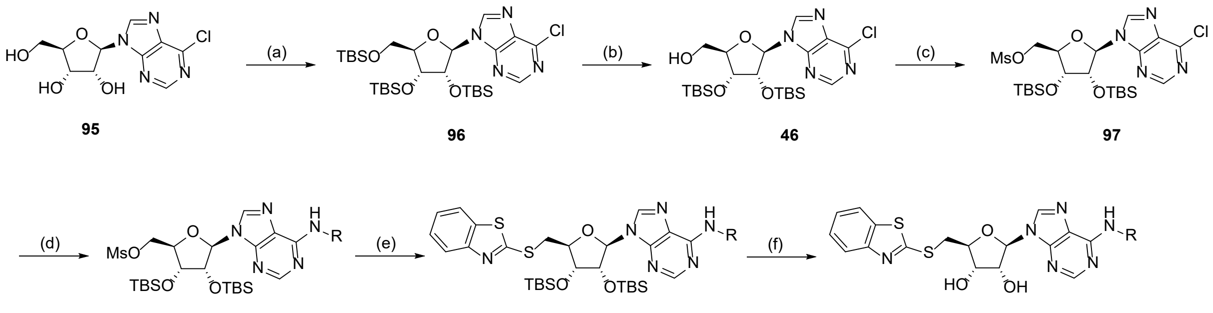

95 96 46 97

98a-c 99a-c 20-22

(α) TBS-Cl, 1H-imidazole, DMF, RT, 16 h, 91%; (b) $CCl_3COOH$, THF, H2O, 0 °C, 56%; (c) $Ms_2O$, TEA, THF, RT, 18%; (d) TEA, KI, THF, RT, 32 h, 75-92%; (e) DMF, 50 °C, 16 h, 70-90%; (f) KF, MeOH, 60 °C, 16 h, 24-61%.

**[(2*R*,3*R*,4*R*,5*R)*-3,4-Bis[[tert-butyl(dimethyl)silyl]oxy]-5-(6-chloropurin-9-yl) tetrahydrofuran-2-yl]methoxy-tert-butyl-dimethyl-silane (96)**

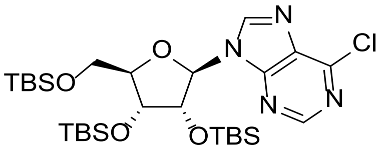

To a solution of (2*R*,3*R*,4*S*,5*R*)-2-(6-chloropurin-9-yl)-5-(hydroxymethyl)tetrahydrofuran- 3,4-diol **95** (10.0 g, 34.9 mmol) in DMF (40 mL) was added imidazole (26.1 g, 383 mmol) and *tert*-butylchlorodimethylsilane (28.9 g, 191 mmol). The resulting mixture was stirred at RT for 16 h. The reaction was quenched with water (200 mL) and extracted with EtOAc (3 x 100 mL). The combined organic layers were washed with brine (3 x 50 mL), dried over sodium sulfate, filtered, and the filtrate was concentrated under reduced pressure. The residue was purified by flash column chromatography (petroleum ether/ethyl acetate gradient 2-20%) to afford compound **96** (20.0 g, 91% yield) as a white solid. $^{1}$H NMR (400 MHz, $CDCl_3$): *δ* 8.75 (s, 1H), 8.56 (s, 1H), 6.13 (d, *J* = 5.2 Hz, 1H), 4.59 (t, *J* = 4.4 Hz, 1H), 4.31 (t, *J* = 4.0 Hz, 1H), 4.18 - 4.15 (m, 1H), 4.03 (dd, *J* = 11.6, 3.6 Hz, 1H), 3.81 (dd, *J* = 11.2, 3.2 Hz, 1H), 0.97 (s, 9H), 0.94 (s, 9H), 0.79 (s, 9H), 0.16 (s, 3H), 0.15 (s, 3H), 0.11 (s, 3H), 0.10 (s, 3H), -0.03 (s, 3H), -0.24 (s, 3H).

**[(2*R*,3*R*,4*R*,5*R*)-3,4-Bis[[tert-butyl(dimethyl)silyl]oxy]-5- (6-chloropurin-9-yl) tetrahydrofuran-2-yl]methanol (46)**

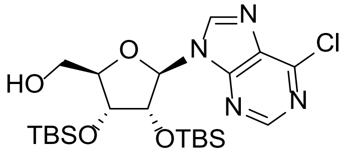

To a solution of compound **96** (5.00 g, 7.94 mmol) in THF (50 mL) was added dropwise a solution of 2,2,2-trichloroacetic acid (12.9 g, 79.4 mmol) in water (15 mL) at 0 °C. The reaction mixture was stirred at 0 °C for 1 h, before being quenched with sat. aq. sodium bicarbonate (50 mL). The aqueous layer was extracted with EtOAc (3 × 50 mL). The combined organic layers were washed with brine (2 × 30 mL), dried over sodium sulfate, filtered, and the filtrate concentrated under reduced pressure. The residue was purified by flash column chromatography (petroleum ether/ethyl acetate 25%) to afford compound **46** (2.30 g, 56% yield) as a white solid. LCMS (ES$^{+}$, Method 7) *m/z* 515.2 [M+H]$^{+}$; $^{1}$H NMR (400 MHz, $CDCl_3$): *δ* 8.79 (s, 1H), 8.21 (s, 1H), 5.89 (d, *J* = 8.0 Hz, 1H), 5.53 (brs, 1H), 4.99 (dd, *J* = 8.0, 0.8 Hz, 1H), 4.35 (d, *J* = 4.4 Hz, 1H), 4.20 (s, 1H), 3.97 (dd, *J* = 13.2, 2.0 Hz, 1H), 3.75 (d, *J* = 13.2 Hz, 1H), 0.96 (s, 9H), 0.75 (s, 9H), 0.14 (d, *J* = 5.2 Hz, 6H), -0.12 (s, 3H), -0.64 (s, 3H).

**[(2R,3R,4R,5R)-3,4-Bis[[tert-butyl(dimethyl)silyl]oxy]-5- (6-chloropurin-9-yl) tetrahydrofuran-2-yl]methyl methanesulfonate (97)**

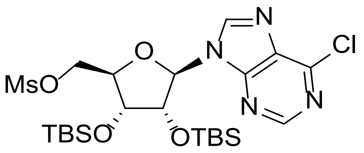

To a solution of compound **46** (2.20 g, 4.27 mmol) and triethylamine (11.9 mL, 85.4 mmol) in THF (10 mL) was added a solution of methanesulfonic anhydride (7.44 g, 42.7 mmol) in THF (10 mL). The reaction mixture was stirred at RT for 30 min before being quenched with sat. aq. sodium bicarbonate solution (20 mL). The aqueous layer was extracted with EtOAc (3 × 20 mL). The combined organic layers were washed with brine (2 × 10 mL), dried over sodium sulfate, filtered, and the filtrate was concentrated under reduce pressure. The residue was purified by flash column chromatography (petroleum ether/ethyl acetate 18%) to afford compound **97** (2.10 g, 83% yield) as a white solid. LCMS ($ES^+$, Method 7) *m/z* 593.2 $[M+H]^+$; $^1H$ NMR (400 MHz, $CDCl_3$): $\delta$ 8.78 (s, 1H), 8.34 (s, 1H), 6.01 (d, *J* = 4.8 Hz, 1H), 4.92 - 4.89 (m, 1H), 4.61 - 4.59 (m, 1H), 4.49 - 4.44 (m, 1H), 4.38 - 4.35 (m, 2H), 3.05 (s, 3H), 0.94 (s, 9H), 0.82 (s, 9H), 0.14 (s, 3H), 0.13 (s, 3H), 0.01 (s, 3H), -0.20 (s, 3H).

**[(2*R*,3*R*,4*R*,5*R*)-3,4-Bis[[tert-butyl(dimethyl)silyl]oxy]-5-[6- (cyclopentylamino)purin-9-yl]tetrahydrofuran-2-yl]methyl methanesulfonate (98a)**

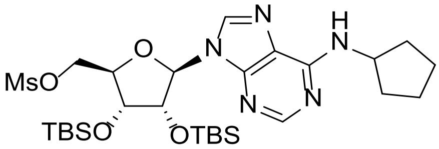

To a solution of compound **97** (500 mg, 0.85 mmol) in THF (5 mL) was added triethylamine (352 µL, 2.53 mmol), potassium iodide (69.9 mg, 0,42 mmol) and cyclopentanamine (332 µL, 3.37 mmol). The reaction mixture was stirred at RT for 32 h and then concentrated under reduced pressure. The residue was purified by flash column chromatography (petroleum ether/ethyl acetate gradient 2-75%) to afford compound **98a** (420 mg, 77% yield) as colorless oil. LCMS ($ES^+$, Method 1) *m/z* 642.4 $[M+H]^+$. $^1H$ NMR (400 MHz, $CDCl_3$): $\delta$ 8.34 (s, 1H), 7.85 (s, 1H), 5.86 (d, J=4.8Hz, 1H), 5.82 (s, 1H), 5.03 (t, *J* = 4.8 Hz, 1H), 4.60 (dd, *J* = 11.2, 4.0 Hz, 1H),

4.53 (dd, $J$ = 11.2, 4.0 Hz, 1H), 4.42 (t, $J$ = 4.0 Hz, 1H), 4.33 - 4.31 (m, 1H), 2.99 (s, 3H), 2.16 - 2.10 (m, 2H), 1.80 - 1.76 (m, 2H), 1.73 - 1.68 (m, 2H), 1.58 - 1.56 (m, 2H), 0.94 (s, 9H), 0.82 (s, 9H), 0.14 (s, 3H), 0.13 (s, 3H), -0.02 (s, 3H), -0.23 (s, 3H).

**((2R,3R,4R,5R)-3,4-Bis((tert-butyldimethylsilyl)oxy)-5-(6-(isopropylamino)-9H-purin-9-yl)tetrahydrofuran-2-yl)methyl methanesulfonate (98b)**

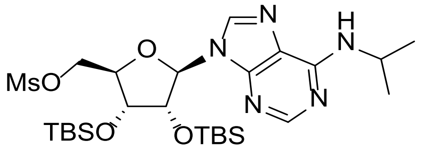

To a solution of compound **97** (250 mg, 0.42 mmol) in THF (2 mL) was added triethylamine (176 µL, 1.26 mmol), potassium iodide (35.0 mg, 0.21 mmol) and propan-2-amine (1.45 µL, 1.69 mmol). The reaction mixture was stirred at RT for 16 h and then concentrated under reduced pressure. The residue was purified by preparative TLC ($SiO_2$, petroleum ether/ethyl acetate 75%) to afford compound **98b** (230 mg, 92% yield) as yellow oil. LCMS ($ES^+$, Method 1) *m/z* 616.3 $[M+H]^+$. $^1H$ NMR (400 MHz, $CDCl_3$): $\delta$ 8.34 (s, 1H), 7.86 (s, H), 5.87 (d, $J$ = 4.8 Hz, 1H), 5.75 - 5.65 (m, 1H), 5.03 (t, $J$ = 4.8Hz, 1H), 4.63 - 4.59 (m, 1H), 4.56 - 4.51 (m, 1H), 4.43 - 4.41 (m, 1H), 4.34 - 4.31 (m, 1H), 2.99 (s, 3H), 1.34 - 1.32 (m, 6H), 1.29 - 1.22 (m, 1H), 0.98 - 0.89 (m, 10H), 0.86 - 0.79 (m, 10H), 0.15 - 0.13 (m, 6H), -0.22 (s, 3H).

**[(2*R*,3*R*,4*R*,5*R*)-3,4-Bis[[tert-butyl(dimethyl)silyl]oxy]-5-[6-(oxetan -3-ylamino)purin-9-yl]tetrahydrofuran-2-yl]methyl methanesulfonate (98c)**

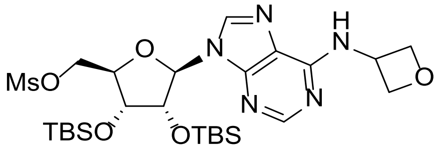

To a solution of compound **97** (500 mg, 0.84 mmol) in THF (5 mL) was added triethylamine (352 µL, 2.53 mmol), potassium iodide (69.9 mg, 0.42 mmol) and oxetan-3-amine (246mg, 3.37 mmol). The reaction mixture was stirred at RT for 16 h and then concentrated under reduced pressure. The residue was purified by flash column chromatography (petroleum ether/ethyl acetate gradient 2-20%) to afford compound **98c** (400 mg, 75% yield) as a colorless oil. LCMS ($ES^+$, Method 3) *m/z* 630.1 $[M+H]^+$. $^1H$ NMR (400 MHz, $CDCl_3$): $\delta$ 8.34 (s, 1H), 7.93 (s, 1H), 5.88 (d, $J$ = 5.2 Hz, 1H), 5.47 (s, 1H), 5.07 (t, $J$ = 7.2 Hz, 2H), 5.01 (t, $J$ = 4.4 Hz,

1H), 4.70 - 4.66 (m, 2H), 4.60 (dd, *J* = 11.2, 4.4 Hz, 1H), 4.52 (dd, *J* = 10.8, 5.2 Hz, 1H), 4.40 (t, *J* = 4.0 Hz, 1H), 4.33 - 4.31 (m, 1H), 3.00 (s, 3H), 0.95 (s, 9H), 0.81 (s, 9H), 0.14 (s, 3H), 0.13 (s, 3H), -0.02 (s, 3H), -0.24 (s, 3H).

**9-[(2*R*,3*R*,4*R*,5*S*)-5-(1,3-Benzothiazol-2-ylsulfanylmethyl)-3,4-bis [[tert-butyl(dimethyl) silyl]oxy]tetrahydrofuran-2-yl]-*N*-cyclopentyl-purin-6-amine (99a)**

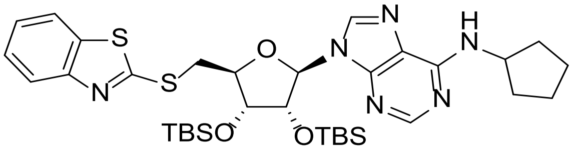

To a solution of compound **98a** (400 mg, 0.62 mmol) in DMF (5 mL) was added 1,3-benzothiazol-2-ylsulfanyl sodium (353mg, 1.87 mmol) and stirred at 50 °C for 16 h. The reaction mixture was cooled to RT, quenched with water (30 mL) and extracted with EtOAc (3 x 15 mL). The combined organic layers were washed with brine (3 x 10 mL), dried over sodium sulfate, filtered and the filtrate was concentrated under reduced pressure. The residue was purified by flash column chromatography (petroleum ether/ethyl acetate gradient 2-100%) to afford compound **99a** (400 mg, 90% yield) as a white solid. LCMS (ES$^+$, Method 3) *m/z* 713.3 $[M+H]^+$. $^1$H NMR (400 MHz, $CDCl_3$): *δ* 8.37 (s, 1H), 7.86 (s, 1H), 7.84 (s, 1H), 7.75 (d, *J* = 8.0 Hz, 1H), 7.44 - 7.40 (m, 1H), 7.32 - 7.28 (m, 1H), 5.87 - 5.84 (m, 2H), 5.33 (dd, *J* = 6.4, 4.4 Hz, 1H), 4.49 (td, *J* = 6.8, 1.6 Hz, 1H), 4.39 (dd, *J* = 4.0, 2.0 Hz, 1H), 4.05 (dd, *J* = 14.0, 6.0 Hz, 1H), 3.87 (dd, *J* = 14.0, 7.2 Hz, 1H), 2.18 - 2.12 (m, 2H), 1.74 - 1.67 (m, 4H), 1.63 - 1.55 (m, 2H), 0.93 (s, 9H), 0.76 (s, 9H), 0.12 (s, 3H), 0.11 (s, 3H), -0.11 (s, 3H), -0.36 (s, 3H).

**9-((2R,3R,4R,5S)-5-((Benzo[d]thiazol-2-ylthio)methyl)-3,4-bis((tert-butyldimethylsilyl) oxy)tetrahydrofuran-2-yl)-N-isopropyl-9H-purin-6-amine (99b)**

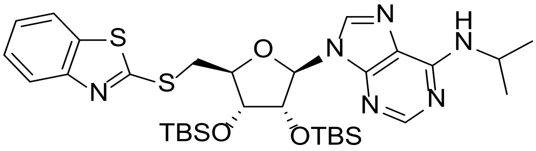

To a solution of compound **98b** (230 mg, 0.37 mmol) in DMF (3 mL) was added 1,3-benzothiazol-2-ylsulfanyl sodium (212 mg, 1.12 mmol) and stirred at 50 °C for 16 h. The reaction mixture was cooled to RT, quenched with water (5 mL) and then extracted with EtOAc (3 × 10 mL). The combined organic layers were washed with brine (15 mL), dried over sodium sulfate, filtered, and the filtrate was concentrated under reduced pressure. The residue was purified by preparative TLC ($SiO_2$, petroleum ether/ethyl acetate 83%) to afford compound **99b** (180 mg, 70% yield) as a colorless oil. LCMS ($ES^+$, Method 3) *m/z* 687.4 $[M+H]^+$. $^1H$ NMR (400 MHz, $CDCl_3$): $\delta$ 8.36 (s, 1H), 7.90-7.83 (m, 2H), 7.78 - 7.74 (m, 1H), 7.49 - 7.43 (m, 1H), 7.42 - 7.38 (m, 1H), 7.37 - 7.32 (m, 1H), 5.89 - 5.85 (d, *J* = 16 Hz, 1H), 5.80 - 5.60 (m, 1H), 5.40 - 5.27 (m, 1H), 4.53 - 4.45 (m, 1H), 4.43 - 4.36 (m, 1H), 4.08 - 4.01 (m, 1H), 3.93 - 3.83 (m, 1H), 1.36 - 1.33 (m, 6H), 0.93 (s, 9H), 0.76 (s, 9H), 0.12 (d, *J* = 1.6 Hz, 6H), -0.11 (s, 3H), -0.36 (s, 3H).

**9-[(2*R*,3*R*,4*R*,5*S*)-5-(1,3-Benzothiazol-2-ylsulfanylmethyl)-3,4-bis [[tert-butyl(dimethyl) silyl]oxy]tetrahydrofuran-2-yl]-*N*-(oxetan-3-yl)purin-6-amine (99c)**

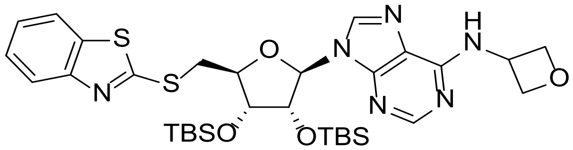

To a solution of compound **98c** (180 mg, 285 µmol) in DMF (3 mL) was added 1,3-benzothiazol-2-ylsulfanyl sodium (162 mg, 857 µmol) and stirred at 50 °C for 16 h. The reaction mixture was cooled to RT, quenched with water (30 mL) and extracted with EtOAc (3 x 10 mL). The combined organic layers were washed with brine (10 mL), dried over sodium sulfate, filtered, and the filtrate was concentrated under reduced pressure. The residue was purified by flash column chromatography (petroleum ether/ethyl acetate gradient 2-100%) to afford compound **99c** (150 mg, 75% yield) as a white solid. LCMS ($ES^+$, Method 3) *m/z* 701.2 $[M+H]^+$. $^1H$ NMR (400 MHz, $CDCl_3$): $\delta$ 8.36 (s, 1H), 7.93 (s, 1H), 7.85 (d, *J* = 8.0 Hz, 1H), 7.75 (d, *J* = 8.0 Hz, 1H), 7.44 - 7.40 (m, 1H), 7.33 - 7.29 (m, 1H), 6.62 (d, *J* = 6.8 Hz, 1H), 5.88 (d, *J* = 6.4 Hz, 1H), 5.49 - 5.48 (m, 1H), 5.29 (dd, *J* = 6.0, 4.0 Hz, 1H), 5.07 (t, *J* = 6.8 Hz, 1H), 4.70 (q, *J* = 6.0 Hz, 2H), 4.50 (td, *J* = 6.8, 1.6 Hz, 1H), 4.38 (dd, *J* = 4.4, 2.0 Hz, 1H), 4.03 (dd, *J* = 14.0, 6.0 Hz, 1H), 3.86 (dd, *J* = 14.0, 7.2 Hz, 1H), 0.92 (s, 9H), 0.75 (s, 9H), 0.12 (s, 3H), 0.11 (s, 3H), -0.12 (s, 3H), -0.38 (s, 3H).

**(2*S*,3*S*,4*R*,5*R*)-2-(1,3-Benzothiazol-2-ylsulfanylmethyl)-5- [6-(cyclopentylamino)purin-9-**

**yl]tetrahydrofuran-3,4-diol (20)**

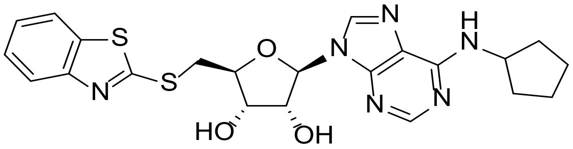

To a solution of compound **99a** (400 mg, 0.56 mmol) in MeOH (8 mL) was added potassium fluoride (325 mg, 5.61 mmol). The reaction mixture was stirred at 50 °C for 16 h, cooled to RT and concentrated under reduced pressure. The residue was purified by reverse column chromatography (0.05% $NH_3{\cdot}H_2O$ in water/MeCN gradient 0-25%) to afford compound **20** (166 mg, 61% yield) as a white solid. HRMS (Method 1) *m/z* $[M+H]^+$ calcd for $C_{22}H_{24}N_6O_3S_2$ 485.1430; found 485.1448. $^1H$ NMR (400 MHz, DMSO-$d_6$): $\delta$ 8.37 (s, 1H), 8.21 (s, 1H), 8.00 (d, *J* = 8.0 Hz, 1H), 7.86 (d, *J* = 8.0 Hz, 1H), 7.73 (d, *J* = 6.4 Hz, 1H), 7.47 (td, *J* = 8.0, 0.8 Hz, 1H), 7.37 (td, *J* = 8.0, 0.8 Hz, 1H), 5.93 (d, *J* = 6.0 Hz, 1H), 5.56 (d, *J* = 6.0 Hz, 1H), 5.47 (d, *J* = 4.8 Hz, 1H), 4.87 (q, *J* = 5.2 Hz, 1H), 4.53 (br.s, 1H), 4.30 - 4.27 (m, 1H), 4.25 - 4.23 (m, 1H), 3.87 (dd, *J* = 13.6, 5.6 Hz, 1H), 3.75 (dd, *J* = 13.6, 7.6 Hz, 1H), 2.00 - 1.92 (m, 2H), 1.78 - 1.69 (m, 2H), 1.64 - 1.54 (m, 4H).

**(2S,3S,4R,5R)-2-((benzo[d]thiazol-2-ylthio)methyl)-5-(6-(isopropylamino)-9H-purin-9-yl)tetrahydrofuran-3,4-diol (21)**

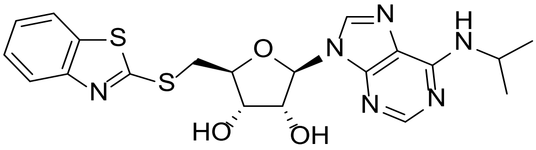

To a solution of compound **99b** (180 mg, 0.26 mmol) in MeOH (3 mL) was added potassium fluoride (123 µL, 5.24 mmol). The reaction mixture was stirred at 60 °C for 16 h, cooled to RT and then concentrated under reduced pressure. The residue was purified by reverse column chromatography (0.1% $NH_3{\cdot}H_2O$ in water/MeCN gradient 0-100%) to afford compound **21** (29.4 mg, 24% yield) as a white solid. HRMS (Method 1) *m/z* $[M+H]^+$ calcd for $C_{20}H_{22}N_6O_3S_2$ 459.1278; found 459.1292. $^1H$ NMR (400 MHz, DMSO-$d_6$): $\delta$ 8.37 (s, 1H), 8.21 - 8.18 (m, 1H), 8.00 (d, *J* = 7.6 Hz, 1H), 7.85 (d, J = 8 Hz, 1H), 7.61 - 7.59 (m, 1H), 7.47 (t, *J* = 7.6 Hz, 1H), 7.36 (t, *J* = 7.6 Hz, 1H), 5.92 (d, *J* = 5.6Hz, 1H), 5.56 (d, *J* = 6 Hz, 1H), 5.48 (d, *J* = 4.8 Hz,1H), 4.88 - 4.84 (m , 1H), 4.54 - 4.35 (m, 1H), 4.31 - 4.27 (m, 1H), 4.26 - 4.22 (m, 1H), 3.91 - 3.83 (m, 1H), 3.79 - 3.69 (m, 1H), 1.22 (d, *J* = 6.4Hz, 6H).

**(2*S*,3*S*,4*R*,5*R*)-2-(1,3-Benzothiazol-2-ylsulfanylmethyl)-5- [6-(oxetan-3-ylamino)purin-9-yl]tetrahydrofuran-3,4-diol (22)**

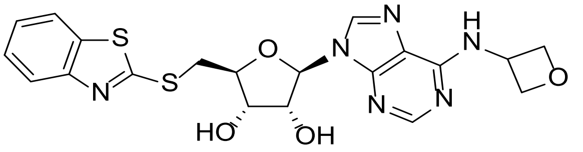

To a solution of compound **99c** (150 mg, 0.21 mmol) in methanol (4 mL) was added potassium fluoride (124 mg, 2.14 mmol). The reaction mixture was stirred at 60 °C for 16 h, cooled to RT and concentrated under reduced pressure. The residue was purified by reverse column chromatography (0.05% $NH_3 \cdot H_2O$ in water/MeCN gradient 0-25%) to afford compound **22** (29.4 mg, 28% yield) as a white solid. HRMS (Method 1) *m/z* $[M+H]^+$ calcd for $C_{20}H_{20}N_6O_4S_2$ 473.1066; found 473.1092. $^1$H NMR (400 MHz, DMSO-*d₆*): $\delta$ 8.60 (s, 1H), 8.45 (s, 1H), 8.23 (s, 1H), 8.00 (d, *J* = 7.6 Hz, 1H), 7.85 (d, *J* = 8.0 Hz, 1H), 7.47 (td, *J* = 8.0, 0.8 Hz, 1H), 7.36 (td, *J* = 8.0, 0.8 Hz, 1H), 5.94 (d, *J* = 5.6 Hz, 1H), 5.62 - 5.55 (m, 2H), 5.20 (br.s, 1H), 4.86 (t, *J* = 5.2 Hz, 1H), 4.80 (t, *J* = 6.8 Hz, 2H), 4.65 (t, *J* = 6.0 Hz, 2H), 4.29 (t, *J* = 4.8 Hz, 1H), 4.26 - 4.22 (m, 1H), 3.86 (dd, *J* = 13.6, 5.6 Hz, 1H), 3.75 (dd, *J* = 13.6, 7.6 Hz, 1H).

##### 1.7.5 Synthesis of compounds 25 and 33

**Scheme S10**

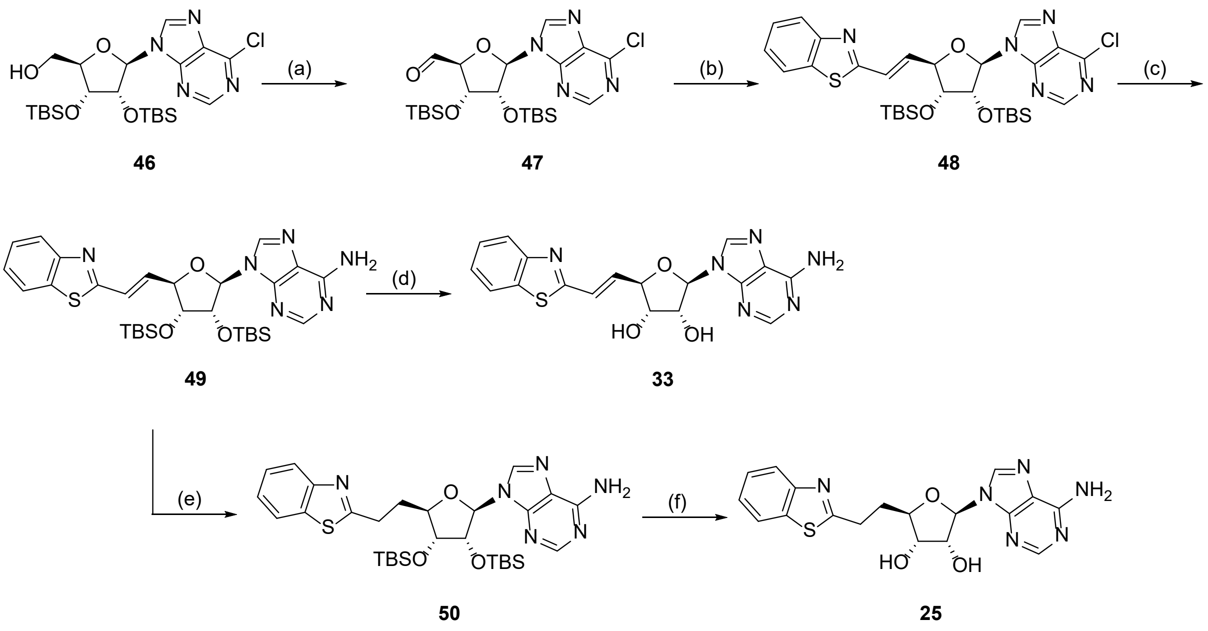

(α) (COCl)$_2$, DMSO, TEA, DCM, -60 °C, 3 h, 67%; (b) (benzo[d]thiazol-2-ylmethyl)triphenylphosphonium chloride, KHMDS, THF, -78 °C, 2 h, 87%; (c) $NH_3$, THF, -70 - 80 °C, 16 h, 51%; (d) KF, DMF, 60 °C, 16 h, 57%; (e) $H_2$, Pd/C, THF, RT, 16 h, 79%; (f) KF, DMF, 60 °C, 16 h, 35%.

**(2*S*,3*R*,4*R*,5*R*)-3,4-Bis[[tert-butyl(dimethyl)silyl]oxy]-5- (6-chloropurin-9-yl)tetrahydrofuran-2-carbaldehyde (47)**

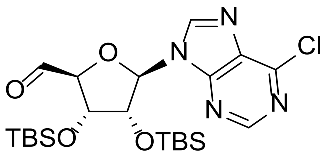

To a solution of DMSO (3.64 mL, 46.6 mmol) in DCM (30 mL) was added oxalyl chloride (3.06 mL, 34.94 mmol) at -60 °C under inert atmosphere and stirred for 30 min. A solution of compound **46** (3.00 g, 5.82 mmol) in DCM (10 mL) was then added and stirred for additional 30 min, followed by the addition of triethylamine (9.73 mL, 69.88 mmol). The reaction mixture was stirred at -60 °C for 2 h and quenched with water (100 mL) at 0 °C. The aqueous layer was extracted with DCM (3 x 50 mL). The combined organic layers were dried over sodium sulfate, filtered and the filtrate was concentrated under reduced pressure. The residue was purified by flash column chromatography (petroleum ether/ethyl acetate gradient 2-17%) to afford compound **47** (2.00 g, 67% yield) as yellow oil. LCMS (ES$^+$, Method 1) *m/z* 513.2 [M+H]$^+$.

**[(2*R*,3*R*,4*R*,5*R*)-2-[(*E*)-2-(1,3-Benzothiazol-2-yl)vinyl]-4-[tert-butyl (dimethyl)silyl]oxy-5-(6-chloropurin-9-yl)tetrahydrofuran-3-yl]oxy-tert-butyl-dimethyl-silane (48)**

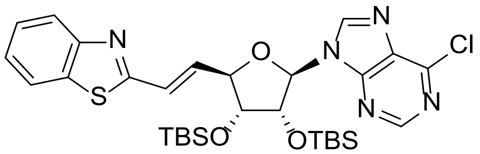

To a solution of (benzo[d]thiazol-2-ylmethyl)triphenylphosphonium chloride (86.9 mg, 194 µmol) in THF (2 mL) was added potassium bis(trimethylsilyl)amide (195 µL, 1M in THF) under inert atmosphere at -78 °C and stirred for 30 min. A solution of compound **47** (100 mg, 195 µmol) in THF (2 mL) was then added and stirred at -78 °C for 1 h. The reaction mixture was quenched with sat. aq. ammonium chloride solution (15 mL) at 0 °C and extracted with EtOAc (3 x 10 mL). The combined organic layers were washed with brine (10 mL), dried over sodium sulfate, filtered, and the filtrate was concentrated under reduced pressure. The residue was purified by preparative TLC chromatography ($SiO_2$, petroleum ether/ethyl acetate

25%) to afford compound **48** (110 mg, 87% yield) as yellow oil. LCMS ($ES^+$, Method 5) *m/z* 644.2 $[M+H]^+$; $^1H$ NMR (400 MHz, $CDCl_3$): $\delta$ 8.86 (s, 1H), 8.27 (s, 1H), 8.02 (d, *J* = 8.0 Hz, 1H), 7.90 (d, *J* = 8.0 Hz, 1H), 7.50 (t, *J* = 7.6 Hz, 1H), 7.42 (t, *J* = 7.6 Hz, 1H), 7.08 - 7.03 (m, 2H), 6.03 (d, *J* = 4.8 Hz, 1H), 5.12 (t, *J* = 4.8 Hz, 1H), 4.81 (t, *J* = 4.8 Hz, 1H), 4.36 (t, *J* = 4.0 Hz, 1H), 0.97 (s, 9H), 0.83 (s, 9H), 0.15 (s, 3H), 0.14 (s, 3H), -0.01 (s, 3H), -0.26 (s, 3H).

**9-[(2*R*,3*R*,4*R*,5*R*)-5-[(*E*)-2-(1,3-Benzothiazol-2-yl)vinyl]-3,4-bis [[tert-butyl(dimethyl)silyl] oxy]tetrahydrofuran-2-yl]purin-6-amine (49)**

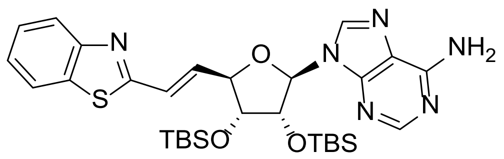

To a solution of compound **48** (30.0 mg, 0.05 µmol) in THF (10 mL) was bubbled ammonia (39.6 mg, 2.33 mmol) at -70 °C for 5 min. The solution was then transferred to a sealed tube, stirred at RT for 16 h and then heated to 80 °C for 16 h. The reaction mixture was cooled to RT and concentrated under reduced pressure. The residue was purified by preparative TLC chromatography ($SiO_2$, petroleum ether/ethyl acetate 25%) to afford compound **49** (15.0 mg, 51% yield) as yellow oil. LCMS ($ES^+$, Method 3) *m/z* 625.3 $[M+H]^+$.

**(2*R*,3*R*,4*S*,5*R*)-2-(6-Aminopurin-9-yl)-5-[(*E*)-2-(1,3-benzothiazol-2-yl) vinyl]tetrahydrofuran-3,4-diol (33)**

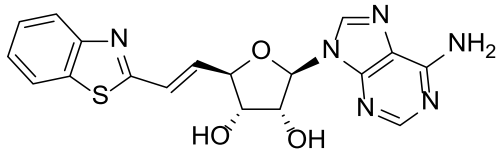

To a solution of compound **49** (15.0 mg, 24.0 µmol) in DMF (2 mL) was added potassium fluoride (13.9 mg, 240 µmol). The reaction mixture was stirred at 60 °C for 16 h, cooled to RT and concentrated under reduced pressure. The residue was purified by prep-HPLC (Method 8) to afford compound **33** (5.52 mg, 13.9 µmol) as a white solid. HRMS (Method 1) *m/z* $[M+H]^+$ calcd for $C_{18}H_{16}N_6O_3S$ 397.1083; found 397.1095; $^1H$ NMR (400 MHz, DMSO-$d_6$): $\delta$ 8.42 (s, 1H), 8.21 (s, 1H), 8.09 (d, *J* = 8.0 Hz, 1H), 7.97 (d, *J* = 7.6 Hz, 1H), 7.51 (td, *J* = 6.8, 1.2 Hz, 1H), 7.44

(td, $J$ = 8.4, 1.2 Hz, 1H), 7.33 (s, 2H), 7.13 - 6.99 (m, 2H), 6.00 (d, $J$ = 5.2 Hz, 1H), 5.63 (d, $J$ = 5.6 Hz, 1H), 5.58 (d, $J$ = 5.2 Hz, 1H), 4.85 (q, $J$ = 5.2 Hz, 1H), 4.65 (t, $J$ = 4.8 Hz, 1H), 4.33 (q, $J$ = 4.8 Hz, 1H).

**9-[(2*R*,3*R*,4*R*,5*R*)-5-[2-(1,3-Benzothiazol-2-yl)ethyl]-3,4-bis [[tert-butyl(dimethyl)silyl]oxy] tetrahydrofuran-2-yl]purin-6-amine (50)**

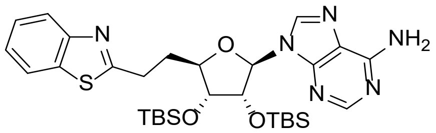

To a solution of compound **49** (100 mg, 160 µmol) in THF (10 mL) was added Pd/C (20 mg, 160 µmol, 10% wt.) under inert atmosphere. The suspension was degassed under reduced pressure and purged with $H_2$ several times. The mixture was stirred under $H_2$ at RT for 16 h, and then filtered. The filtrate was concentrated under reduced pressure and the residue was purified by preparative TLC chromatography ($SiO_2$, petroleum ether/ethyl acetate 33%) to afford compound **50** (80.0 mg, 79% yield) as yellow oil. LCMS ($ES^+$, Method 2) *m/z* 627.4 $[M+H]^+$; $^1H$ NMR (400 MHz, $CDCl_3$): $\delta$ 8.35 (s, 1H), 7.98 (d, $J$ = 8.0 Hz, 1H), 7.88 (s, 1H), 7.85 (d, $J$ = 8.0 Hz, 1H), 7.46 (t, $J$ = 8.0 Hz, 1H), 7.37 (t, $J$ = 8.0 Hz, 1H), 5.86 (d, $J$ = 6.0 Hz, 1H), 5.65 (s, 2H), 5.19 (dd, $J$ = 6.0, 4.4 Hz, 1H), 4.22 - 4.10 (m, 2H), 3.33 - 3.29 (m, 1H), 3.23 - 3.16 (m, 1H), 2.59 - 2.55 (m, 1H), 2.34 - 2.30 (m, 1H), 0.94 (s, 9H), 0.78 (s, 9H), 0.11 (s, 6H), -0.08 (s, 3H), -0.35 (s, 3H).

**(2*R*,3*R*,4*S*,5*R*)-2-(6-Aminopurin-9-yl)-5-[2-(1,3-benzothiazol-2-yl) ethyl]tetrahydrofuran-3,4-diol (25)**

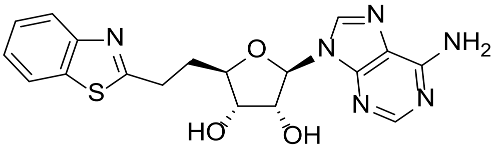

To a solution of compound **50** (80.0 mg, 0.13 mmol) in DMF (2 mL) was added potassium fluoride (74.1 mg, 1.28 mmol). The reaction mixture was stirred at 60 °C for 16 h, cooled to RT and concentrated under reduced pressure. The residue was purified by prep-HPLC (Method 9) to afford compound **25** (17.8 mg, 35% yield) as a white solid. HRMS (Method 1) *m/z* $[M+H]^+$

calcd for $C_{18}H_{18}N_6O_3S$ 399.1239; found 399.1251; $^1$H NMR (400 MHz, DMSO-$d_6$): $\delta$ 8.36 (s, 1H), 8.15 (s, 1H), 8.04 (d, $J$ = 8.0 Hz, 1H), 7.93 (d, $J$ = 8.0 Hz, 1H), 7.48 (t, $J$ = 7.2 Hz, 1H), 7.40 (d, $J$ = 8.0 Hz, 1H), 7.29 (s, 2H), 5.88 (d, $J$ = 5.2 Hz, 1H), 5.37 (s, 2H), 4.73 (t, $J$ = 5.2 Hz, 1H), 4.16 (d, $J$ = 4.8 Hz, 1H), 4.00 - 3.96 (m, 1H), 3.21 - 3.16 (m, 2H), 2.34 (q, $J$ = 7.2 Hz, 2H).

##### 1.7.6 Synthesis of compounds 26 and 27

**Scheme S11**

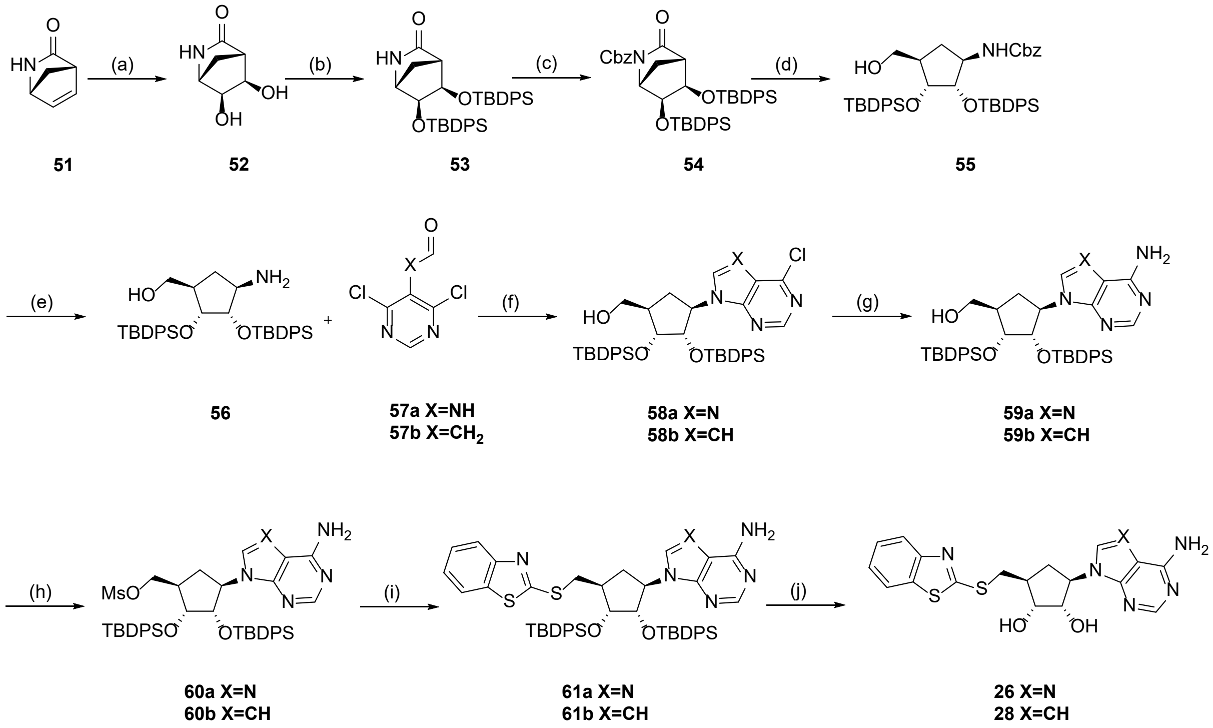

(a) $K_2OsO_4$, NMO, THF, t-BuOH, $H_2O$, RT, 16 h, 54%; (b) TBDPSCl, 1H-imidazole, DMF, RT, 24 h, 63%; (c) CbzCl, LiHMDS, THF, -70 °C, 30 min, 92%; (d) $NaBH_4$, THF, MeOH, 0 °C, 2 h, 97%; (e) $H_2$, Pd/C, THF, RT, 16 h, 82%; (f) TEA, n-BuOH, 130 °C, 24 h, 78%, 77%; (g) $NH_3$/MeOH, 90 °C, 16 h, 77%, 92%; (h) $Ms_2O$, TEA, THF, RT, 16 h, 72%, 90%; (i) sodium benzo[d]thiazole-2-thiolate, DMF, 50 °C, 16 h, 55%, 50%; (j) KF, MeOH, 50 °C, 16 h, 43%, 55%.

##### (1*R*,4*S*,5*R*,6*S*)-5,6-Dihydroxy-2-azabicyclo[2.2.1]heptan-3-one (52)

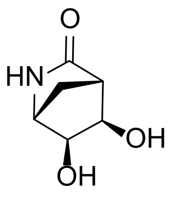

To a solution of (1*R*,4*S*)-2-azabicyclo[2.2.1]hept-5-en-3-one **51** (7.00 g, 64.2 mmol) in THF (140 mL), tert-butyl alcohol (70 mL) and water (20 mL) was added 4-methylmorpholinen-oxidemonohydrate (22.5 g, 192.44 mmol) and potassium osmate(VI) dihydrate (3.55 g, 9.62 mmol). The reaction mixture was stirred at RT for 16 h and then concentrated under reduced pressure. The residue was purified by flash column chromatography (dichloromethane/methanol 0-10%) to afford compound **52** (5.00 g, 54% yield) as a black solid. LCMS (ES$^+$, Method 4) *m/z* 144.2 [M+H]$^+$. $^1$H NMR (400 MHz, DMSO-$d_6$): $\delta$ 7.53 (s, 1H), 5.02 (d, $J$ = 4.8 Hz, 1H), 4.96 (d, $J$ = 5.2 Hz, 1H), 3.77 - 3.76 (m, 1H), 3.71 - 3.70 (m, 1H), 3.43 (s, 1H), 2.25 (s, 1H), 1.89 (d, $J$ = 10.0 Hz, 1H), 1.72 (d, $J$ = 9.6 Hz, 1H).

**(1*R*,4*S*,5*R*,6*S*)-5,6-Bis[[tert-butyl(diphenyl)silyl]oxy]-2-azabicyclo [2.2.1]heptan-3-one (53)**

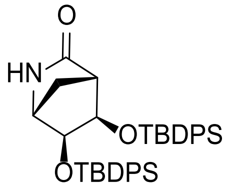

To a solution of compound **52** (4.00 g, 27.9 mmol) in DMF (20 mL) was added 1H-imidazole (19.0 g, 279 mmol) and tert-butylchlorodiphenylsilane (21.5 mL, 83.8 mmol). The reaction mixture was stirred at RT for 24 h. The reaction was then quenched with water (100 mL) and extracted with EtOAc (3 x 50 mL). The combined organic layers were washed with brine (2 x 50 mL), dried over sodium sulfate, filtered and the filtrate was concentrated under reduced pressure. The residue was purified by flash column chromatography (petroleum ether/ethyl acetate 2-50%) to afford compound **53** (11.0 g, 63% yield) as yellow oil. LCMS (ES$^+$, Method 2) *m/z* 620.2 [M+H]$^+$. $^1$H NMR (400 MHz, DMSO-$d_6$): $\delta$ 7.67 - 7.62 (m, 7H), 7.49 - 7.45 (m, 4H), 7.41 - 7.36 (m, 9H), 4.14 (d, $J$ = 5.6 Hz, 1H), 4.06 (d, $J$ = 6.0 Hz, 1H), 3.12 (s, 1H), 2.25 (d, $J$ = 9.2 Hz, 1H), 2.03 (s, 1H), 1.70 (d, $J$ = 9.6 Hz, 1H), 1.01 (s, 18H).

**Benzyl (1*R*,4*S*,5*R*,6*S*)-5,6-bis[[tert-butyl(diphenyl)silyl]oxy]-3-oxo-2-azabicyclo[2.2.1] heptane-2-carboxylate (54)**

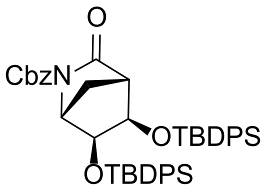

To a solution of compound **53** (8.00 g, 12.9 mmol) in THF (100 mL) under inert atmosphere at -70 °C was added dropwise lithium bis(trimethylsilyl)amide (12.9 mL , 1 M in THF) and was stirred for 10 min. Benzyl carbonochloridate (2.75 mL, 19.36 mmol) was then added and further stirred for 30 min. The reaction mixture was quenched with sat. aq. ammonium chloride solution (300 mL) and extracted with EtOAc (3 x 150 mL). The combined organic layers were washed with brine (2 x 100 mL), dried over sodium sulfate, filtered and the filtrate was concentrated under reduced pressure. The residue was purified by column chromatography by flash column chromatography (petroleum ether/ethyl acetate 2-25%) to afford compound **54** (9.00 g, 92% yield) as colorless oil. LCMS ($ES^+$, Method 3) *m/z* 754.3 $[M+H]^+$. $^1H$ NMR (400 MHz, DMSO-$d_6$): $\delta$ 7.67 - 7.57 (m, 5H), 7.46 - 7.45 (m, 5H), 7.40 - 7.34 (m, 10H), 7.29 - 7.38 (m, 3H), 7.20 - 7.17 (m, 2H), 4.94 (d, $J$ = 4.0 Hz, 2H), 4.50 (d, $J$ = 5.6 Hz, 1H), 4.35 (d, $J$ = 5.6 Hz, 1H), 4.09 (s, 1H), 2.32 (d, $J$ = 10.0 Hz, 1H), 2.22 (s, 1H), 1.93 (d, $J$ = 10.0 Hz, 1H), 0.99 (s, 9H), 0.97 (s, 9H).

**Benzyl *N*-[(1*R*,2*S*,3*R*,4*R*)-2,3-bis[[tert-butyl(diphenyl)silyl]oxy]-4 (hydroxymethyl) cyclopentyl]carbamate (55)**

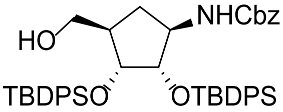

To a solution of compound **54** (9.00 g, 11.9 mmol) in THF (100 mL) and MeOH (20 mL) was added portionwise sodium borohydride (1.03 g, 27.2 mmol) at 0 °C. The resulting mixture was stirred at 0 °C for 2 h. The reaction was then quenched with water (300 mL) at 0 °C and extracted with EtOAc (3 x 100 mL). The combined organic layers were washed with brine (2 x 50 mL), dried over sodium sulfate, filtered and the filtrate was concentrated under reduced pressure. The residue was purified by flash column chromatography (petroleum ether/ethyl acetate 2-50%) to afford compound **55** (8.82 g, 97% yield) as colorless oil. LCMS ($ES^+$, Method 3) *m/z* 680.3 $[M+H]^+$. $^1H$ NMR (400 MHz, DMSO-$d_6$): $\delta$ 7.66 - 7.57 (m, 6H), 7.50 - 7.49 (m, 2H),

7.45 - 7.22 (m, 17H), 4.99 (d, $J$ = 12.4 Hz, 1H), 4.92 (d, $J$ = 12.4 Hz, 1H), 4.24 (t, $J$ = 4.8 Hz, 1H), 4.17 (t, $J$ = 8.8 Hz, 1H), 3.94 - 3.92 (m, 1H), 3.83 - 3.82 (m, 1H), 2.89 - 2.85 (m, 1H), 2.67 - 2.62 (m, 1H), 2.03 - 1.99 (m, 1H), 1.96 - 1.93 (m, 1H), 1.01 (s, 9H), 0.92 (s, 9H).

**[(1*R*,2*R*,3*S*,4*R*)-4-Amino-2,3-bis[[tert-butyl(diphenyl)silyl]oxy] cyclopentyl]methanol (56)**

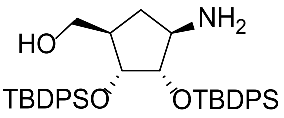

To a solution of compound **55** (8.00 g, 10.5 mmol) in THF (200 mL) was added Pd/C (10%, 100 mg, 1.06 mmol) under inert atmosphere. The suspension was degassed under reduced pressure and purged with $H_2$ several times. The mixture was stirred under $H_2$ at RT for 16 h, then filtered. The filtrate was concentrated under reduced pressure to afford compound **56** (5.40 g, 82% yield) as a brown solid. The isolated material was used in the following step without any further purification. LCMS (ES$^+$, Method 3) *m/z* 624.3 [M+H]$^+$. $^1$H NMR (400 MHz, DMSO-$d_6$): $\delta$ 7.66 - 7.58 (m, 8H), 7.45 - 7.40 (m, 8H), 7.33 - 7.24 (m, 4H), 4.00 - 3.97 (m, 1H), 3.66 - 3.63 (m, 1H), 3.62 - 3.58 (m, 1H), 3.31 (brs, 2H), 3.24 - 3.19 (m, 1H), 2.99 (dd, $J$ = 10.4, 4.8 Hz, 1H), 2.69 - 2.64 (m, 1H), 2.05 - 2.02 (m, 1H), 1.95 - 1.89 (m, 1H), 1.01 (s, 9H), 0.98 (s, 9H).

***N*-(4,6-Dichloropyrimidin-5-yl)formamide (57a)**

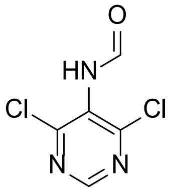

To a suspension of 4,6-dichloropyrimidin-5-amine (5.00 g, 30.5 mmol) in formic acid (62 mL) was added acetic anhydride (83.3 mL, 889 mmol) at 0 °C. The reaction mixture was warmed to RT, stirred for 1 h and then concentrated under reduced pressure to afford compound **57a** (4.00 g, 68% yield) as a white solid. The isolated material was used in the following step without any further purification. $^1$H NMR (400 MHz, DMSO-$d_6$): $\delta$ 10.62 (s, 1H), 9.41 (s, 1H), 8.36 (s, 1H).

**[(1*R*,2*R*,3*S*,4*R*)-2,3-Bis[[tert-butyl(diphenyl)silyl]oxy]-4-(6-chloropurin-9-yl)cyclopentyl] methanol (58a)**

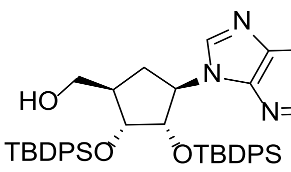

To a solution of compound **56** (5.00 g, 8.01 mmol) in n-butyl alcohol (75 mL) was added *N,N*-diisopropylethylamine (4.19 mL, 24.04 mmol) and compound **57a** (1.85 g, 9.62 mmol). The reaction mixture was heated at 130 °C for 24 h, cooled to RT and then concentrated under reduced pressure. The residue was purified by flash column chromatography (petroleum ether/ethyl acetate 2-50%) to afford compound **58a** (4.80 g, 78% yield) as a yellow solid. LCMS ($ES^+$, Method 3) *m/z* 761.4 $[M+H]^+$. $^1H$ NMR (400 MHz, DMSO-$d_6$): $\delta$ 8.48 (s, 1H), 8.43 (s, 1H), 7.78 - 7.76 (m, 2H), 7.71 - 7.69 (m, 2H), 7.50 - 7.39 (m, 6H), 7.32 - 7.23 (m, 5H), 7.11 - 7.04 (m, 5H), 5.34 (q, *J* = 9.6 Hz, 1H), 4.72 (dd, *J* = 10.0, 3.6 Hz, 1H), 4.65 (t, *J* = 5.2 Hz, 1H), 4.23 (d, *J* = 3.6 Hz, 1H), 3.14 - 3.11 (m, 1H), 3.05 - 3.02 (m, 1H), 2.21 - 2.18 (m, 1H), 2.10 - 2.07 (m, 1H), 1.93 - 1.88 (m, 1H), 1.11 (s, 9H), 0.79 (s, 9H).

**[(1*R*,2*R*,3*S*,4*R*)-2,3-Bis[[tert-butyl(diphenyl)silyl]oxy]-4-(4- chloropyrrolo[2,3-d]pyrimidin-7-yl)cyclopentyl]methanol (58b)**

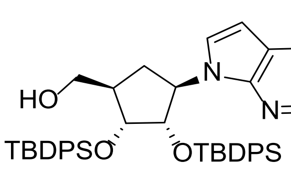

To a solution of compound **56** (0.80 g, 1.28 mmol) in n-butyl alcohol (20 mL) was added 2-(4,6-dichloropyrimidin-5-yl)acetaldehyde **57b** (293.88 mg, 1.54 mmol) and diisopropylethylamine (670 µL, 3.85 mmol). The reaction mixture was heated at 130 °C for 24 h, cooled to RT and then concentrated under reduced pressure. The residue was purified by flash column chromatography (petroleum ether/ethyl acetate 2-75%) to afford compound **58b** (0.70 g, 71% yield) as yellow oil. LCMS ($ES^+$, Method 2) *m/z* 760.3 $[M+H]^+$. $^1H$ NMR (400 MHz, $CDCl_3$): $\delta$ 7.75 - 7.71 (m, 6H), 7.65 - 7.64 (m, 2H), 7.47 - 7.40 (m, 5H), 7.39 - 7.31 (m, 6H), 7.30 - 7.28 (m, 4H), 4.97 (d, *J* = 12.4 Hz, 1H), 4.89 (d, *J* = 12.4 Hz, 1H), 4.37 (d, *J* = 7.6 Hz, 1H), 4.04 - 4.03 (m, 1H),

4.01 - 3.98 (m, 1H), 3.00 - 2.94 (m, 2H), 2.36 - 2.28 (m, 1H), 2.04 - 2.01 (m, 1H), 1.09 (s, 9H), 1.04 (s, 9H).

**[(1*R*,2*R*,3*S*,4*R*)-4-(6-Aminopurin-9-yl)-2,3-bis[[tert-butyl(diphenyl) silyl]oxy]cyclopentyl] methanol (59a)**

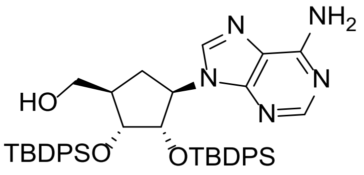

A solution of compound **58a** (4.00 g, 5.25 mmol) and ammonia (20 mL, 7 M in methanol) in a sealed tube was heated at 90 °C and stirred for 16 h. The mixture was cooled to RT and concentrated under reduced pressure. The residue was purified by flash column chromatography (petroleum ether/ethyl acetate 2-50%) to afford compound **59a** (3.00 g, 77% yield) as a yellow solid. LCMS ($ES^+$, Method 2) *m/z* 742.7 $[M+H]^+$. $^1H$ NMR (400 MHz, DMSO-$d_6$): $\delta$ 7.90 (s, 2H), 7.65 (t, $J$ = 6.0 Hz, 4H), 7.48 - 7.34 (m, 9H), 7.28 - 7.25 (m, 1H), 7.18 (t, $J$ = 7.6 Hz, 2H), 7.13 (s, 2H), 7.07 - 7.00 (m, 4H), 5.21 (q, $J$ = 9.6 Hz, 1H), 4.73 (dd, $J$ = 6.0, 4.8 Hz, 1H), 4.65 (dd, $J$ = 9.6, 3.6 Hz, 1H), 4.10 - 4.06 (m, 1H), 3.06 - 3.02 (m, 1H), 2.91 - 2.89 (m, 1H), 2.16 - 2.13 (m, 1H), 1.99 - 1.96 (m, 1H), 1.92 - 1.89 (m, 1H), 1.04 (s, 9H), 0.77 (s, 9H).

**[(1*R*,2*R*,3*S*,4*R*)-4-(4-Aminopyrrolo[2,3-d]pyrimidin-7-yl)-2,3-bis [[tert-butyl(diphenyl)silyl] oxy]cyclopentyl]methanol (59b)**

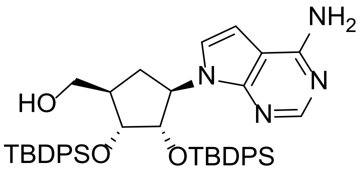

A solution of compound **58b** (100 mg, 131 µmol) and ammonia (1.14 mL, 7 M in methanol) in a sealed tube was heated at 110 °C and stirred for 16 h. The mixture was cooled to RT and concentrated under reduced pressure. The residue was purified by flash column chromatography (petroleum ether/ethyl acetate 2-50%) to afford compound **59b** (90.0 mg, 92% yield) as yellow oil. LCMS ($ES^+$, Method 1) *m/z* 741.3 $[M+H]^+$. $^1H$ NMR (400 MHz, DMSO-$d_6$): $\delta$ 7.82 (s, 1H), 7.75 - 7.73 (m, 4H), 7.44 - 7.33 (m, 9H), 7.26 - 7.24 (m, 1H), 7.21 - 7.19 (m, 3H), 7.07 (t, $J$ = 7.6 Hz, 2H), 6.99 (t, $J$ = 7.6 Hz, 2H), 6.82 (d, $J$ = 3.2 Hz, 1H), 6.12 (d, $J$ = 3.2 Hz,

1H), 5.26 (brs, 1H), 5.02 (dd, $J$ = 9.2, 3.6 Hz, 1H), 4.86 - 4.79 (m, 1H), 4.28 (d, $J$ = 3.6 Hz, 1H), 3.32 (dd, $J$ = 12.0, 2.4 Hz, 1H), 3.04 (dd, $J$ = 11.6, 2.4 Hz, 1H), 2.39 - 2.29 (m, 1H), 2.00 - 1.97(m, 2H), 1.16 (s, 9H), 0.97 (s, 9H).

**[(1*R*,2*R*,3*S*,4*R*)-4-(6-Aminopurin-9-yl)-2,3-bis[[tert-butyl(diphenyl) silyl]oxy]cyclopentyl] methyl methanesulfonate (60a)**

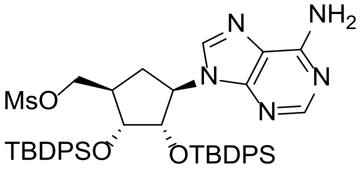

To a solution of compound **59a** (1.00 g, 1.35 mmol) and triethylamine (563 uL, 4.04 mmol) in THF (20 mL) was added a solution of methanesulfonic anhydride (281 mg, 1.62 mmol) in THF (5 mL) at 0 °C. The reaction mixture was warmed to RT and stirred for 16 h. The reaction was quenched with water (50 mL) and then extracted with EtOAc (3 x 30 mL). The combined organic layers were washed with brine (3 x 20 mL), dried over sodium sulfate, filtered and the filtrate was concentrated under reduced pressure. The residue was purified by flash column chromatography (petroleum ether/ethyl acetate 2-50%) to afford compound **60a** (800 mg, 72% yield) as yellow oil. LCMS ($ES^+$, Method 3) *m/z* 820.3 $[M+H]^+$. $^1H$ NMR (400 MHz, DMSO-$d_6$): $\delta$ 8.01 (s, 1H), 7.86 (s, 1H), 7.64 (d, $J$ = 6.4 Hz, 2H), 7.59 (d, $J$ = 6.8 Hz, 2H), 7.50 - 7.43 (m, 4H), 7.38 - 7.28 (m, 6H), 7.18 (t, $J$ = 7.6 Hz, 2H), 7.13 (s, 2H), 7.09 - 7.02 (m, 4H), 5.21 (q, $J$ = 9.2 Hz, 1H), 4.70 (dd, $J$ = 9.2, 3.6 Hz, 1H), 4.08 (d, $J$ =3.6 Hz, 1H), 3.84 (d, $J$ = 6.8 Hz, 2H), 2.93 (s, 3H), 2.28 - 2.20 (m, 2H), 2.02 - 1.99 (m, 1H), 1.02 (s, 9H), 0.78 (s, 9H).

**[(1*R*,2*R*,3*S*,4*R*)-4-(4-Aminopyrrolo[2,3-d]pyrimidin-7-yl)-2,3-bis [[tert-butyl(diphenyl) silyl]oxy]cyclopentyl]methyl methanesulfonate (60b)**

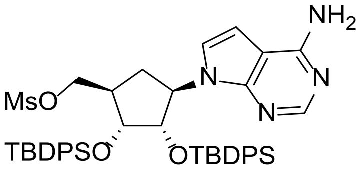

To a solution of compound **59b** (90.0 mg, 121 µmol) and triethylamine (101 µL, 728 µmol) in THF (5 mL) was added a solution of methanesulfonic anhydride (63.4 mg, 364 µmol) in THF (2 mL) at 0 °C. The reaction mixture was stirred at RT for 2 h. The reaction was quenched with water (20 mL) and then extracted with EtOAc (3 x 10 mL). The combined organic layers were washed with brine (3 x 10 mL), dried over sodium sulfate, filtered and the filtrate was concentrated under reduced pressure. The residue was purified by flash column chromatography (petroleum ether/ethyl acetate 2-50%) to afford compound **60b** (90.0 mg, 90% yield) as a white solid. $^1$H NMR (400 MHz, $CDCl_3$): $\delta$ 7.95 (d, *J* = 6.8 Hz, 2H), 7.87 - 7.81 (m, 4H), 7.75 - 7.70 (m, 2H), 7.65 - 7.57 (m, 4H), 7.54 - 7.49 (m, 4H), 7.43 - 7.34 (m, 5H), 7.15 (s, 1H), 6.93 (d, *J* = 3.6 Hz, 1H), 5.39 (d, *J* = 3.6 Hz, 1H), 4.20 (d, *J* = 4.8 Hz, 1H), 4.08 - 4.06 (m, 1H), 3.87 (d, *J* = 4.4 Hz, 1H), 3.63 (d, *J* = 13.2 Hz, 1H), 3.04 - 2.96 (m, 1H), 2.72 (s, 3H), 2.66 - 2.64 (m, 1H), 2.16 (dd, *J* = 13.6, 4.4 Hz, 1H), 1.65 (d, *J* = 13.2 Hz, 1H), 1.23 (s, 9H), 1.11 (s, 9H).

**9-[(1*R*,2*S*,3*R*,4*S*)-4-(1,3-Benzothiazol-2-ylsulfanylmethyl)-2,3-bis[[tert-butyl(diphenyl)silyl]oxy]cyclopentyl]purin-6-amine (61a)**

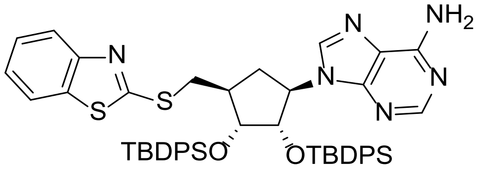

To a solution of compound **60a** (500 mg, 0.61 mmol) in DMF (5 mL) was added 1,3-benzothiazol-2-ylsulfanyl sodium (346 mg, 1.83 mmol) and heated to 50 °C for 16 h. The reaction mixture was cooled to RT, quenched with water (20 mL) and extracted with EtOAc (3 x 10 mL). The combined organic layers were washed with brine (10 mL), dried over sodium sulfate, filtered and the filtrate was concentrated under reduced pressure. The residue was purified by flash column chromatography (petroleum ether/ethyl acetate 2-100%) to afford compound **61a** (300 mg, 55% yield) as yellow oil. LCMS ($ES^+$, Method 3) *m/z* 891.3 $[M+H]^+$. $^1$H NMR (400 MHz, DMSO-$d_6$): $\delta$ 8.04 (s, 1H), 7.95 (d, *J* = 8.0 Hz, 1H), 7.86 (s, 1H), 7.62 (d, *J* = 8.0 Hz, 3H), 7.58 (d, *J* = 6.8 Hz, 2H), 7.46 - 7.41 (m, 2H), 7.38 - 7.32 (m, 4H), 7.31 - 7.22(m, 6H), 7.16 - 7.01 (m, 8H), 5.20 (q, *J* = 9.6 Hz, 1H), 4.83 (dd, *J* = 9.2, 3.2 Hz, 1H), 4.14 (d, *J* = 3.6 Hz, 1H), 3.28 - 3.25(m, 2H), 2.38 - 2.30 (m, 2H), 2.07 - 2.03 (m, 1H), 0.10 (s, 9H), 0.77 (s, 9H).

**7-[(1*R*,2*S*,3*R*,4*S*)-4-(1,3-Benzothiazol-2-ylsulfanylmethyl)-2,3-bis[[tert-butyl(diphenyl)silyl]oxy]cyclopentyl]pyrrolo[2,3-d]pyrimidin-4-amine (61b)**

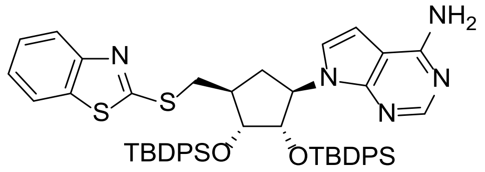

To a solution of compound **60b** (90.0 mg, 110 µmol) in DMF (2 mL) was added 1,3-benzothiazol-2-ylsulfanyl sodium (62.4 mg, 329 µmol) and heated at 50 °C for 16 h. The reaction mixture was cooled to RT, quenched with water (15 mL) and extracted with EtOAc (3 x 10 mL). The combined organic layers were washed with brine (3 x 10 mL), dried over sodium sulfate, filtered and the filtrate was concentrated under reduced pressure. The residue was purified by preparative TLC ($SiO_2$, petroleum ether/ethyl acetate 50%) to afford compound **61b** (30.0 mg, 30% yield) as yellow oil. LCMS ($ES^+$, Method 1) *m/z* 890.2 $[M+H]^+$.

**(1*R*,2*S*,3*R*,5*S*)-3-(6-aminopurin-9-yl)-5-(1,3-benzothiazol-2-ylsulfanylmethyl) cyclopentane-1,2-diol (26)**

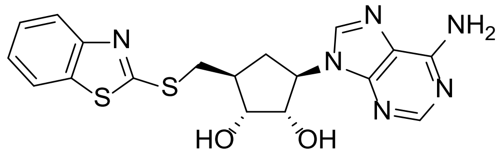

To a solution of compound **61a** (300 mg, 0.34 mmol) in MeOH (10 mL) was added potassium fluoride (196 mg, 3.37 mmol). The reaction mixture was stirred at 50 °C for 16 h, cooled to RT and then concentrated under reduced pressure. The residue was purified by reverse column chromatography (0.05% $NH_3{\cdot}H_2O$ in water/MeCN gradient 0-5%) to afford compound **26** (62.7 mg, 43% yield) as a white solid. HRMS (Method 1) *m/z* $[M+H]^+$ calcd for $C_{18}H_{18}N_6O_2S_2$ 415.1011; found 415.1022. $^1H$ NMR (400 MHz, DMSO-*d$_6$*): $\delta$ 8.22 (s, 1H), 8.11 (s, 1H), 8.01 (d, *J* = 8.0 Hz, 1H), 7.86 (d, *J* = 8.0 Hz, 1H), 7.47 (td, *J* = 8.0, 0.8 Hz, 1H), 7.37 (td, *J* = 8.4, 0.8 Hz, 1H), 7.19 (s, 2H), 5.07 (d, *J* = 6.0 Hz, 1H), 4.95 (d, *J* = 4.4 Hz, 1H), 4.72 - 4.67 (m, 1H), 4.48 - 4.43 (m, 1H), 3.96 - 3.92 (m, 1H), 3.69 (dd, *J* = 12.8, 6.4 Hz, 1H), 3.54 (dd, *J* = 13.2, 8.0 Hz, 1H), 2.43 - 2.37(m, 2H), 1.92 - 1.87(m, 1H).

**(1*R*,2*S*,3*R*,5*S*)-3-(4-Aminopyrrolo[2,3-d]pyrimidin-7-yl)-5- (1,3-benzothiazol-2 -ylsulfanylmethyl)cyclopentane-1,2-diol (28)**

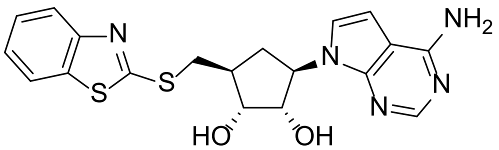

To a solution of compound **61b** (30.0 mg, 33.7 µmol) in MeOH (2 mL) was added p-toluenesulfonic acid (29.0 mg, 168 µmol). The reaction mixture was heated to 70 °C for 16 h, cooled to RT and concentrated under reduced pressure. The residue was purified by reverse column chromatography (0.05% $NH_3 \cdot H_2O$ in water/MeCN gradient 0-5%) to afford compound **28** (8.16 mg, 55% yield) as a light yellow solid. HRMS (Method 1) *m/z* $[M+H]^+$ calcd for $C_{19}H_{19}N_5O_2S_2$ 414.1058; found 414.1080. $^1H$ NMR (400 MHz, DMSO-$d_6$): $\delta$ 8.02 - 8.00 (m, 2H), 7.86 (d, *J* = 8.0 Hz, 1H), 7.47 (t, *J* = 8.0 Hz, 1H), 7.36 (t, *J* = 7.6 Hz, 1H), 7.26 (d, *J* = 3.6 Hz, 1H), 6.91 (s, 2H), 6.54 (d, *J* = 3.2 Hz, 1H), 4.96 (s, 1H), 4.85 - 4.83 (m, 1H), 4.28 (t, *J* = 7.6 Hz, 1H), 3.91 (t, *J* = 4.8 Hz, 1H), 3.67 (dd, *J* = 12.8, 6.4 Hz, 1H), 3.56 - 3.47 (m, 1H), 2.35 - 2.30 (m, 2H), 1.73 - 1.67 (m, 1H).

##### 1.7.7 Synthesis of compound 27

__Scheme S12__

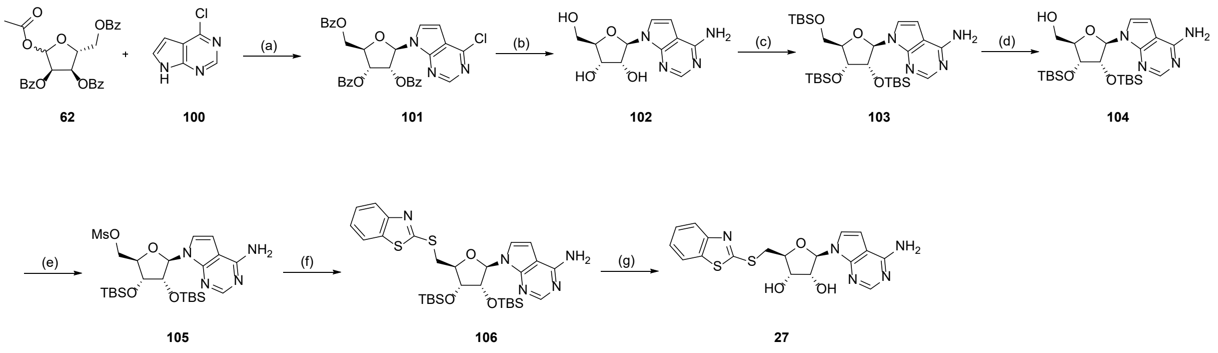

(a) BSA, TMSOTf, MeCN, 25-80 °C, 1.5 h, 27%; (b) $NH_3$/MeOH, 90 °C, 32 h, 75%; (c) TBS-Cl, 1H-imidazole, DMF, RT, 16 h, 74%; (d) $Cl_3CCO_2H$, THF/H2O, 0 °C, 16 h, 58%; (e) $Ms_2O$, TEA, THF, RT, 16 h, 73%; (f) sodium benzo[d]thiazole-2-thiolate, DMF, 50 °C, 16 h, 78%; (g) KF, MeOH, 50 °C, 16 h, 19%

**[(2*R*,3*R*,4*R*,5*R*)-3,4-Dibenzoyloxy-5-(4-chloropyrrolo[2,3-d] pyrimidin-7-yl)tetrahydrofuran-2-yl]methyl benzoate (101)**

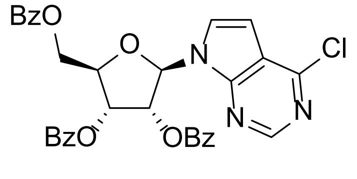

*N,O*-Bis(trimethylsilyl)acetamide (8.05 mL, 32.6 mmol) was added to a stirred suspension of 4-chloro-7*H*-pyrrolo[2,3-d]pyrimidine **100** (5.00 g, 32.6 mmol) MeCN (100 mL) at RT. After stirring for 10 min, [(2*R*,3*R*, 4*R*)-5-acetoxy-3, 4-dibenzoyloxy-tetrahydrofuran-2-yl] methyl benzoate **62** (18.1 g, 35.8 mmol) was added, followed by trimethylsilyl trifluoromethanesulfonate (5.88 mL, 32.56 mmol). The reaction mixture was stirred at RT for 15 min before being heated at 80 °C for 1 h. The reaction mixture was cooled to RT and diluted with EtOAc (75 mL). The organic phase was sequentially washed with sat. aq. sodium bicarbonate solution (2 x 200 mL) and brine (50 mL), dried over sodium sulfate, filtered and the filtrate was concentrated under reduced pressure. The residue was purified by flash column chromatography (petroleum ether/ethyl acetate gradient 0-50%) to afford compound **101** (5.60 g, 27% yield) as yellow oil. LCMS ($ES^+$, Method 1) *m/z* 598.1 $[M+H]^+$. $^1H$ NMR (400 MHz, $CDCl_3$): $\delta$ 8.61 (s, 1H), 8.23 - 8.09 (m, 2H), 8.06 - 7.98 (m, 2H), 7.96 - 7.89 (m, 2H), 7.64 - 7.53 (m, 3H), 7.51 - 7.46 (m, 2H), 7.45 - 7.34 (m, 5H), 6.69 (d, *J* = 5.6 Hz, 1H), 6.64 (d, *J* = 3.6 Hz, 1H), 6.25 (t, *J* = 5.6 Hz, 1H), 6.20 - 6.13 (m, 1H), 4.90 (dd, *J* = 3.2, 12.0 Hz, 1H), 4.83 - 4.77 (m, 1H), 4.69 (dd, *J* = 4.0, 12.0 Hz, 1H).

**(2*R*,3*R*,4*S*,5*R*)-2-(4-Aminopyrrolo[2,3-d]pyrimidin-7-yl)-5-(hydroxymethyl)tetrahydrofuran-3,4-diol (102)**

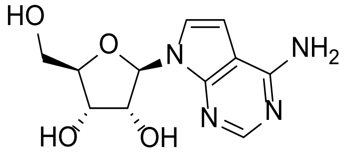

A solution of compound **101** (3.00 g, 5.02 mmol) in ammonia (36.0 mL, 7 M in MeOH) in a sealed tube was heated at 90 °C for 32 h. The mixture was cooled to RT and concentrated under reduced pressure. The residue was purified by reverse column chromatography (0.05% $NH_3 \cdot H_2O$ in water/MeCN gradient 0-5%) to afford compound **102** (1.00 g, 75% yield) as a yellow solid. LCMS ($ES^+$, Method 5) *m/z* 267.2 $[M+H]^+$.

**7-[(2*R*,3*R*,4*R*,5*R*)-3,4-Bis[[tert-butyl(dimethyl)silyl]oxy]-5-[[tert-butyl(dimethyl)silyl] oxymethyl]tetrahydrofuran-2-yl]pyrrolo[2,3-d]pyrimidin-4-amine (103)**

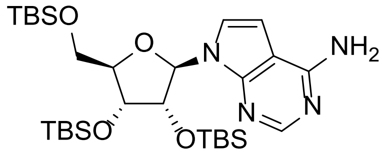

To a solution of compound **102** (1.00 g, 3.76 mmol) and imidazole (2.56 g, 37.5 mmol) in DMF (10 mL) at 0 °C was added tert-butylchlorodimethylsilane (2.83 g, 18.8 mmol). The mixture was warmed to RT and stirred for 16 h. The reaction was quenched with water (50 mL) and extracted with EtOAc (3 x 20 mL). The combined organic layers were washed with brine (3 x 20 mL), dried over sodium sulfate, filtered, and the filtrate was concentrated under reduced pressure. The residue was purified by flash column chromatography (petroleum ether/ethyl acetate gradient 0-50%) to afford compound **103** (1.70 g, 74% yield) as a white solid. LCMS ($ES^+$, Method 2) *m/z* 609.6 $[M+H]^+$. $^1H$ NMR (400 MHz, $CDCl_3$): $\delta$ 8.31 (s, 1H), 7.36 (d, *J* = 3.6 Hz, 1H), 6.37 (d, *J* = 3.6 Hz, 1H), 6.25 (d, *J* = 5.6 Hz, 1H), 5.16 (s, 2H), 4.54 (dd, *J* = 5.6, 4.4 Hz, 1H), 4.29 (dd, *J* = 4.4, 3.2 Hz, 1H), 4.10 - 4.09 (m, 1H), 3.97 (dd, *J* = 11.2, 4.0 Hz, 1H), 3.78 (dd, *J* = 11.2, 2.8 Hz, 1H), 0.97 (s, 9H), 0.94 (s, 9H), 0.78 (s, 9H), 0.13 (d, *J* = 1.6 Hz, 6H), 0.11 (s, 6H), -0.09 (s, 3H), -0.28 (s, 3H).

**[(2*R*,3*R*,4*R*,5*R*)-5-(4-Aminopyrrolo[2,3-d]pyrimidin-7-yl)-3,4- bis[[tert-butyl(dimethyl)silyl]oxy]tetrahydrofuran-2-yl]methanol (104)**

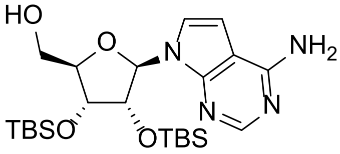

To a solution of compound **103** (1.70 g, 2.79 mmol) in THF (60 mL) at 0 °C was added dropwise a solution of trichloroacetic acid (4.56 g, 27.9 mmol) in water (20 mL). The resulting mixture was stirred at 0 °C for 16 h and then quenched with sat. aq. sodium bicarbonate (100 mL). The aqueous layer was extracted with EtOAc (3 x 50 mL). The combined organic layers were washed with brine (2 x 50 mL), dried over sodium sulfate**,** filtered, and the filtrate was concentrated under reduced pressure. The residue was purified by flash column chromatography (petroleum ether/ethyl acetate gradient 2-100%) to afford compound **104** (800 mg, 58% yield) as a white solid. LCMS ($ES^+$, Method 2) *m/z* 495.2 $[M+H]^+$. $^1H$ NMR (400 MHz, DMSO-$d_6$): $\delta$ 8.03(s, 1H), 7.37 (d, *J* = 3.6 Hz, 1H), 7.05 (s, 1H), 6.58 (d, *J* = 3.6 Hz, 1H), 5.97

(d, $J$ = 3.6 Hz, 1H), 5.69 - 5.67 (m, 1H), 4.74 (dd, $J$ = 7.2, 4.4 Hz, 1H), 4.24 (d, $J$ = 4.8 Hz, 1H), 3.92 - 3.91 (m, 1H), 3.68 - 3.64 (m, 1H), 3.58 - 3.53 (m, 1H).

**((2*R*,3*R*,4*R*,5*R*)-5-(4-Amino-7H-pyrrolo[2,3-d]pyrimidin-7-yl)-3,4- bis((tertbutyldimethyl silyl)oxy)tetrahydrofuran-2-yl)methyl methanesulfonate (105)**

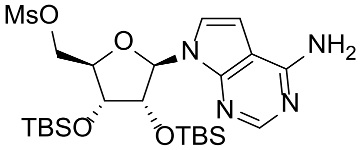

To a solution of compound **104** (0.20 g, 404 µmol) and triethylamine (169 µL, 1.21 mmol) in THF (5 mL) at 0 °C was added dropwise a solution of methanesulfonic anhydride (84.5 mg, 0.49 mmol) in THF (1 mL). The resulting mixture was warmed up to RT and stirred for 16 h. The reaction was quenched with water (20 mL) and extracted with EtOAc (3 x 10 mL). The combined organic layers were washed with brine (2 x 10 mL), dried over sodium sulfate, filtered, and the filtrate concentrated under reduced pressure. The residue was purified by flash column chromatography (petroleum ether/ethyl acetate gradient 2-100%) to afford compound **105** (170 mg, 73% yield) as a yellow oil. LCMS ($ES^+$, Method 2) *m/z* 573.1 $[M+H]^+$. $^1$H NMR (400 MHz, DMSO-$d_6$): $\delta$ 8.07 (s, 1H), 7.34 (d, $J$ = 3.6 Hz, 1H), 7.03 (s, 1H), 6.63 (d, $J$ = 3.6 Hz, 1H), 6.13 (d, $J$ = 7.2 Hz, 1H), 4.74 (dd, $J$ = 7.2, 4.8 Hz, 1H), 4.51 - 4.48 (m, 2H), 4.31 (dd, $J$ = 4.4, 1.2 Hz, 1H), 4.13 (t, $J$ = 5.2 Hz, 1H), 3.21 (s, 3H), 0.94 (s, 9H), 0.69 (s, 9H), 0.15 (s, 3H), 0.13 (s, 3H), -0.11 (s, 3H), -0.38 (s, 3H).

**7-[(2*R*,3*R*,4*R*,5*S*)-5-(1,3-Benzothiazol-2-ylsulfanylmethyl)-3,4-bis [[tert-butyl(dimethyl) silyl]oxy]tetrahydrofuran-2-yl]pyrrolo[2,3-d]pyrimidin-4-amine (106)**

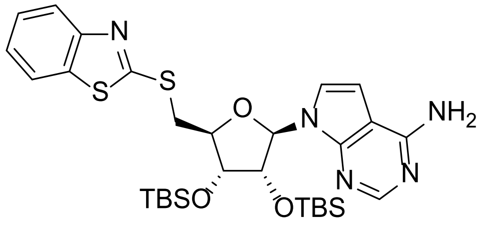

To a solution of compound **105** (170 mg, 297 µmol, 1.0) in DMF (2 mL) was added 1,3-benzothiazol-2-ylsulfanylsodium (84.2 mg, 445 µmol) and stirred at 50 °C for 16 h. The reaction was cooled to RT, quenched with water (20 mL) and extracted with EtOAc (3 x 10 mL). The combined organic layers were washed with brine (3 x 10 mL), dried over sodium sulfate, filtered, and the filtrate was concentrated under reduced pressure. The residue was purified by flash column chromatography (petroleum ether/ethyl acetate gradient 2-100%) to afford compound **106** (150 mg, 78% yield) as yellow oil. LCMS (ES$^+$, Method 3) *m/z* 644.2 [M+H]$^+$. $^1$H NMR (400 MHz, DMSO-*d*$_6$): *δ* 8.06 (s, 1H), 8.01 (d, *J* = 8.0 Hz, 1H), 7.82 (d, *J* = 8.0 Hz, 1H), 7.48 (td, *J* = 8.0, 1.2 Hz, 1H), 7.41 - 7.39 (m, 1H), 7.37 - 7.35 (m, 1H), 7.01 (s, 2H), 6.62 (d, *J* = 3.6 Hz, 1H), 6.08 (d, *J* = 7.6 Hz, 1H), 4.95 (q, *J* = 4.4 Hz, 1H), 4.37 (d, *J* = 4.4 Hz, 1H), 4.27 (t, *J* = 6.8 Hz, 1H), 3.94 (dd, *J* = 14.0, 7.2 Hz, 1H), 3.76 (dd, *J* = 14.0, 6.8 Hz, 1H), 0.88 (s, 9H), 0.65 (s, 9H), 0.08 (d, *J* = 1.2 Hz, 6H), -0.16 (s, 3H), -0.44 (s, 3H).

**(2*R*,3*R*,4*S*,5*S*)-2-(4-Aminopyrrolo[2,3-d]pyrimidin-7-yl)-5-(1,3-benzothiazol-2-ylsulfanylmethyl)tetrahydrofuran-3,4-diol (27)**

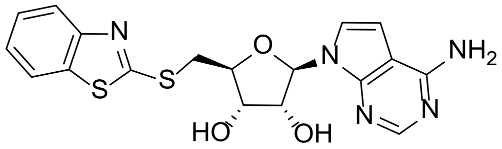

To a solution of compound **106** (150 mg, 0.23 mmol) in methanol (3 mL) was added potassium fluoride (135 mg, 2.33 mmol). The resulting mixture was stirred at 50 °C for 16 h, cooled to RT and concentrated under reduced pressure. The residue was purified by reverse column chromatography (0.05% $NH_3 \cdot H_2O$ in water/MeCN gradient 0-5%) and then triturated from acetonitrile and methanol (1:1, 4 mL) to afford compound **27** (19.2 mg, 19% yield) as a white solid. HRMS (Method 1) *m/z* [M+H]$^+$ calcd for $C_{18}H_{17}N_5O_3S_2$ 416.0851; found 416.0852. $^1$H NMR (400 MHz, DMSO-*d*$_6$): *δ* 8.06 (s, 1H), 8.00 (d, *J* = 7.2 Hz, 1H), 7.86 (d, *J* = 7.6 Hz, 1H), 7.47 (td, *J* = 8.0, 1.2 Hz, 1H), 7.39 - 7.34 (m, 2H), 7.02 (s, 2H), 6.61 (d, *J* = 3.6 Hz, 1H), 6.08 (d, *J* = 6.0 Hz, 1H), 5.42 (s, 2H), 4.57 (t, *J* = 5.6 Hz, 1H), 4.19 - 4.17 (m, 2H), 3.82 (dd, *J* = 13.6, 5.6 Hz, 1H), 3.69 (dd, *J* = 13.6, 7.2 Hz, 1H).

##### 1.7.8 Synthesis of compound 29

<u>**Scheme S13**</u>

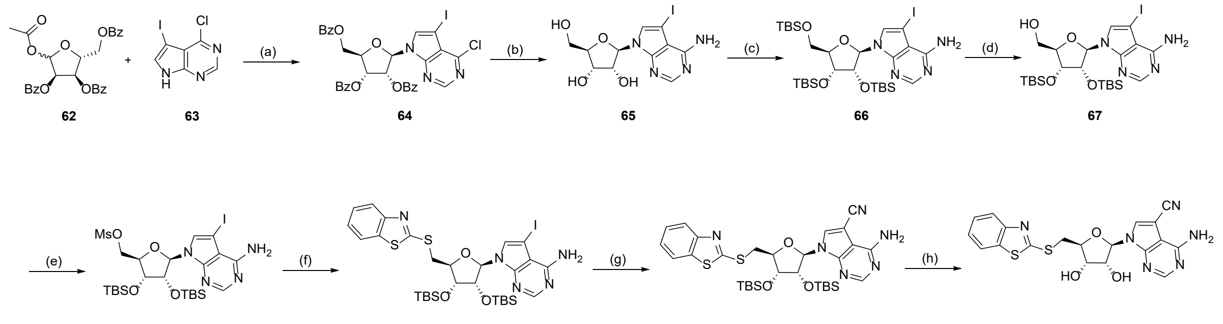

(a) BSA, TMSOTf, MeCN, 25-80 °C, 1.5 h, 49%; (b) $NH_3$/MeOH, 90 °C, 16 h, 98%; (c) TBS-Cl, 1H-imidazole, DMF, RT, 16 h, 30%; (d) $Cl_3CCO_2H$, THF/H2O, 0 °C, 1 h, 84%; (e) $Ms_2O$, TEA, THF, 0°C to RT, 2 h, 89%; (f) sodium benzo[d]thiazole-2-thiolate, DMF, 50 °C, 12 h, 95%; (g) $Pd(PPh_3)_4$, $Zn(CN)_2$, DMF, 90 °C, 16 h, 92%; (h) KF, MeOH, 60 °C, 16 h, 12%.

#### (2R,3R,4R,5R)-2-((Benzoyloxy)methyl)-5-(4-chloro-5-iodo-7H-pyrrolo[2,3-d]pyrimidin-7-yl)tetrahydrofuran-3,4-diyl dibenzoate (64)

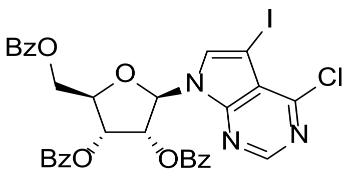

N,O-Bis(trimethylsilyl)acetamide (8.84 mL, 35.7 mmol) was added to a stirred suspension of 4-chloro-5-iodo-7H-pyrrolo[2,3-d]pyrimidine **63** (10.0 g, 35.8 mmol) in MeCN (200 mL). After stirring at RT for 10 min, [(2R,3R,4R)-5-acetoxy-3,4-dibenzoyloxy-tetrahydrofuran-2-yl]methyl benzoate **62** (19.9 g, 39.4 mmol) was added, followed by the addition of trimethylsilyl trifluoromethanesulfonate (6.47 mL, 35.78 mmol). The reaction mixture was stirred at RT for 15 min before being warmed up at 80 °C for 1 h. The reaction mixture was cooled to RT and diluted with EtOAc (75 mL). The organic phase was sequentially washed with sat. aq. sodium bicarbonate (2 x 200 mL) and brine (50 mL), dried over sodium sulfate, filtered, and the filtrate was concentrated under reduced pressure. The isolated material was recrystallized in MeCN to afford compound **64** (12.7 g, 49% yield) as a white solid. $^1$H NMR (400 MHz, $CDCl_3$): $\delta$ 8.58 (s, 1H), 8.12 (d, $J$ = 7.6 Hz, 2H), 8.00 (d, $J$ = 7.6 Hz, 2H), 7.92 (d, $J$ = 6.8 Hz, 2H), 7.56 - 7.51 (m, 6H), 7.42 - 7.37 (m, 4H), 6.67 (d, $J$ = 5.6 Hz, 1H), 6.17 - 6.14 (m, 1H), 6.12 - 6.10 (m, 1H), 4.91 (dd, $J$ = 12.4, 2.8 Hz, 1H), 4.81 (d, $J$ = 4.0Hz, 1H), 4.69 (dd, $J$ = 12.0, 3.6Hz, 1H)

**(2R,3R,4S,5R)-2-(4-Amino-5-iodo-7H-pyrrolo[2,3-d]pyrimidin-7-yl)-5-(hydroxymethyl) tetrahydrofuran-3,4-diol (65)**

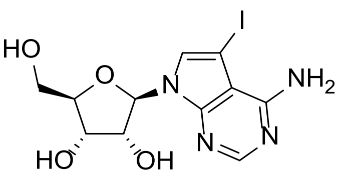

Compound **64** (3 g, 4.14 mmol) was added to a sealed tube containing ammonia (59.2 mL, 7 in MeOH) and stirred at 90 °C for 16 h. The reaction mixture was cooled to RT and concentrated under reduced pressure. The residue was triturated in EtOAc (100 mL) under ultrasonic wave for 10 min and filtered to afford compound **65** (1.60 g, 98% yield) as a black solid. The isolated material was used in the following step without any further purification. $^1$H NMR (400 MHz, DMSO-$d_6$): $\delta$ 8.09 (s, 1H), 7.67 (s, 1H), 6.67 (brs, 2H), 6.02 (d, $J$ = 6.4 Hz, 1H), 5.31 (brs, 1H), 5.16 - 5.12 (m, 2H), 4.35 (s, 1H), 4.06 (d, $J$ = 4.4 Hz, 1H), 3.87 - 3.85 (m, 1H), 3.62 - 3.59 (m, 1H), 3.53 - 3.50 (m, 1H).

**7-((2R,3R,4R,5R)-3,4-Bis((tert-butyldimethylsilyl)oxy)-5-(((tert-butyldimethylsilyl)oxy) methyl)tetrahydrofuran-2-yl)-5-iodo-7H-pyrrolo[2,3-d]pyrimidin-4-amine (66)**

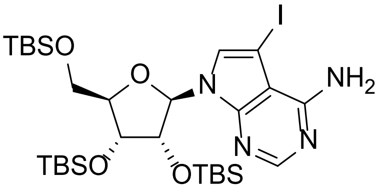

To a solution of compound **65** (2.50 g, 6.38 mmol) and 1H-imidazole (5.25 g, 77.2 mmol) in DMF (25 mL) at RT, was added *t*-butyl-chloro-dimethyl-silane (5.82 g, 38.6 mmol) and stirred for 16 h. The reaction mixture was diluted in hexane (50 mL) and washed with brine (250 mL). The separated aqueous layer was re-extracted with hexane (2 x 100 mL) and the combined organic layers dried over sodium sulfate, filtered, and the filtrate was concentrated under reduced pressure. The residue was purified by flash column chromatography (petroleum ether/ethyl acetate gradient 0-15%) to afford compound **66** (1.40 g, 30% yield) as a black solid. LCMS (ES$^+$, Method 3) *m/z* 735.3 [M+H]$^+$; $^1$H NMR (400 MHz, $CDCl_3$): $\delta$ 8.26 (s, 1H), 7.54 (s, 1H), 6.26 (d, $J$ = 5.2 Hz, 1H), 5.72 (brs, 2H), 4.40 (t, $J$ = 4.8 Hz, 1H), 4.22 - 4.12 (m, 1H), 4.10 - 4.08 (m, 1H), 3.98 - 3.97 (m, 1H), 3.94 - 3.79 (m, 1H), 1.00 (s, 9H), 0.92 (s, 9H), 0.78 (s, 9H), 0.18 (d, $J$ = 5.6 Hz, 6H), 0.11 (s, 6H), -0.071 (s, 3H), -0.27 (s, 3H).

**1-(5-Benzylsulfanyl-6-fluoro-indazol-1-yl)ethanone (67)**

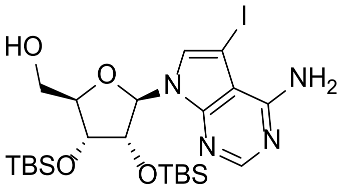

To a solution of compound **66** (1.40 g, 1.90 mmol) in THF (45 mL) cooled to 0 °C, was added dropwise a solution of 2,2,2-trichloroacetic acid (3.11 g, 19.1 mmol) in $H_2O$ (15 mL). Upon completion of the addition, the reaction mixture was stirred for 1 h at 0 °C. The reaction mixture was neutralized using sodium carbonate, diluted with water (100 mL) and extracted with EtOAc (3 x 50 mL). The combined organic layers were dried over magnesium sulphate, filtered, and the filtrate was concentrated under reduced pressure. The residue was purified by flash column chromatography (petroleum ether/ethyl acetate gradient 10-50%) to afford compound **67** (1.00 g, 84% yield) as a black solid. LCMS ($ES^+$, Method 3) *m/z* 621.3 $[M+H]^+$; $^1H$ NMR (400 MHz, $CDCl_3$): *δ* 8.2 (s, 1H), 7.14 (s, 1H), 6.57 - 6.54 (m, 1H), 5.89 (brs, 2H), 5.62 (d, *J* = 8.0 Hz, 1H), 5.04 - 5.38 (m, 1H), 4.30 (d, *J* = 4.4 Hz, 1H), 3.92 (d, *J* = 13.4 Hz, 1H), 3.72 - 3.69 (m, 1H), 0.95 (s, 9H), 0.76 (s, 9H), 0.12 (d, *J* = 5.2 Hz, 6H), -0.14 (s, 3H), -0.59 (s, 3H).

**((2R,3R,4R,5R)-5-(4-Amino-5-iodo-7H-pyrrolo[2,3-d]pyrimidin-7-yl)-3,4-bis((tert-butyldimethylsilyl)oxy)tetrahydrofuran-2-yl)methyl methanesulfonate (68)**

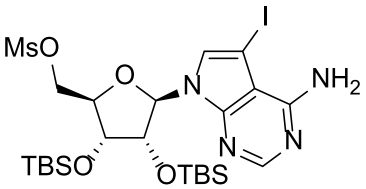

To a solution of compound **67** (1.00 g, 1.61 mmol) and triethylamine (978 mg, 9.67 mmol) in THF (10 mL) was added dropwise methanesulfonic anhydride (842 mg, 4.83 mmol) at 0 °C under inert atmosphere. The mixture was warmed to RT and stirred for 2 h. The reaction was quenched with water (20 mL) and extracted with EtOAc (3 x 20 mL). The combined organic layers were washed with brine (20 mL), dried over sodium sulfate, filtered, and the filtrate was concentrated under reduced pressure. The residue was purified by flash column chromatography (petroleum ether/ethyl acetate gradient 10-50%) to afford compound **68** (1.00 g, 89% yield) as yellow oil. LCMS (ES$^+$, Method 3) *m/z* 699.2 [M+H]$^+$; $^1$H NMR (400 MHz, MeOH-$d_4$): $\delta$ 8.27 (s, 1H), 7.28 - 7.27 (m, 1H), 6.03 (d, *J* = 4.4 Hz, 1H), 5.76 (s, 1H), 4.72 (t, *J* = 4.4 Hz, 1H), 4.58 - 4.55 (m, 1H), 4.49 - 4.45 (m, 1H), 4.35 - 4.30 (m, 2H), 3.03 (s, 1H), 0.95 (s, 9H), 0.84 (s, 9H), 0.14 (d, *J* = 4.0 Hz, 6H), 0.002 (s, 3H), -0.13 (s, 3H).

**7-((2R,3R,4R,5S)-5-((Benzo[d]thiazol-2-ylthio)methyl)-3,4-bis((tert-butyldimethylsilyl) oxy)tetrahydrofuran-2-yl)-5-iodo-7H-pyrrolo[2,3-d]pyrimidin-4-amine (69)**

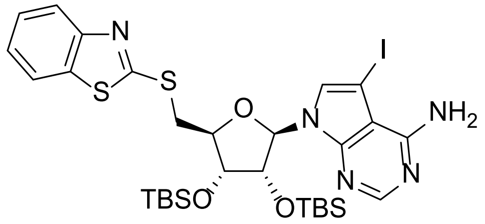

To a solution of compound **68** (1 g, 1.43 mmol) in DMF (3 mL) was added 1,3-benzothiazol-2-ylsulfanyl sodium (812 mg, 4.29 mmol). The resulting mixture was stirred at 50 °C for 12 h. The reaction mixture was cooled to RT, quenched with water (20 mL) and the resulting aqueous layer extracted with EtOAc (3 x 10 mL). The combined organic layers were washed with brine (10 mL), dried over sodium sulphate, filtered, and the filtrate was concentrated under reduced pressure. The residue was purified by flash column chromatography (petroleum ether/ethyl acetate gradient 0-15%) to afford compound **69** (1.05 g, 95% yield) as a yellow solid. LCMS (ES$^+$, Method 3) *m/z* 770.2 [M+H]$^+$; $^1$H NMR (400 MHz, $CDCl_3$): $\delta$ 8.2 (s, 1H), 7.78 (d, *J* = 8.4 Hz, 1H), 7.77 (d, *J* = 8.0 Hz, 1H), 7.45 - 7.40 (m, 1H), 7.35 - 7.33 (m, 1H), 7.24 - 7.2 2(m, 1H), 5.99 (d, *J* = 6.4 Hz, 1H), 5.66 (s, 2H), 4.99 - 4.96 (m, 1H), 4.45 (dd, *J* = 5.8, 2.4 Hz, 1H), 4.33 - 4.32(m, 1H), 3.98 - 3.93 (m, 1H), 3.83 (d, *J* = 6.4 Hz, 1H), 2.05 (s, 4H), 1.28 - 1.25 (m, 4H), 0.97 - 0.91 (m, 10H), 0.11 (d, *J* = 9.3 Hz, 6H), -0.14 (s, 3H), -0.32 (s, 1H).

**4-Amino-7-[(2*R*,3*R*,4*R*,5*S*)-5-(1,3-benzothiazol-2-ylsulfanylmethyl)- 3,4-bis[[tert-butyl (dimethyl)silyl]oxy]tetrahydrofuran-2-yl]pyrrolo[2,3-d]pyrimidine-5-carbonitrile (70)**

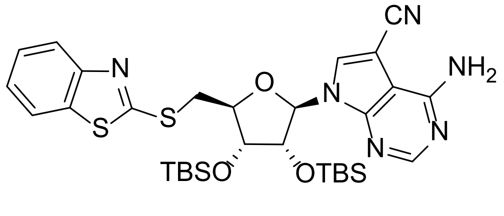

To a solution of compound **69** (100 mg, 129 µmol) in DMF (5 mL) was added $Pd(PPh_3)_4$ (30.0 mg, 25.9 µmol) and zinc cyanide (30.5 mg, 259 µmol). The reaction mixture was stirred at 90 °C under inert atmosphere for 16 h. The reaction was quenched with water (20 mL) and extracted with EtOAc (3 x 10 mL). The combined organic layers were washed with brine (10 mL), dried over sodium sulfate, filtered, and the filtrate was concentrated under reduced pressure. The residue was purified by flash column chromatography (petroleum ether/ethyl acetate gradient 2-50%) to afford compound **70** (80.0 mg, 92% yield) as yellow oil. LCMS ($ES^+$, Method 1) *m/z* 669.2 $[M+H]^+$; $^1H$ NMR (400 MHz, $CDCl_3$): $\delta$ 8.36 (s, 1H), 7.89 (d, *J* = 8.0 Hz, 1H), 7.78 - 7.77 (m, 2H), 7.44 (td, *J* = 8.4, 1.2 Hz, 1H), 7.32 (td, *J* = 8.0, 0.8 Hz, 1H), 5.97 (d, *J* = 6.0 Hz, 1H), 5.62 (s, 2H), 4.98 (dd, *J* = 5.6, 4.4 Hz, 1H), 4.50 - 4.48 (m, 1H), 4.29 (dd, *J* = 4.0, 2.8 Hz, 1H), 3.96 (dd, *J* = 14.4, 5.6 Hz, 1H), 3.88 (dd, *J* = 14.0, 5.6 Hz, 1H), 0.95 (s, 9H), 0.94 (s, 9H), 0.12 (s, 3H), 0.10 (s, 3H), -0.12 (s, 3H), -0.29 (s, 3H).

**4-Amino-7-[(2*R*,3*R*,4*S*,5*S*)-5-(1,3-benzothiazol-2-ylsulfanylmethyl) -3,4-dihydroxy-tetrahydrofuran-2-yl]pyrrolo[2,3-d]pyrimidine-5-carbonitrile (29)**

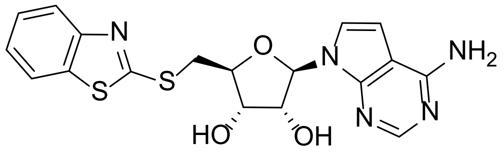

To a solution of compound **70** (80.0 mg, 0.12 mmol) in MeOH (3 mL) was added potassium fluoride (69.5 mg, 1.20 mmol). The resulting mixture was stirred at 60 °C for 16 h, cooled to RT and then concentrated under reduced pressure. The residue was purified by reverse column chromatography (0.05% $NH_3 \cdot H_2O$ in water/MeCN gradient 0-25%) and then purified by prep-HPLC (Method 10) to afford compound **29** (6.36 mg, 12% yield) as a grey solid. HRMS (Method 1) *m/z* $[M+H]^+$ calcd for $C_{19}H_{16}N_6O_3S_2$ 441.0804; found 441.0817; $^1H$ NMR (400 MHz, DMSO-$d_6$): $\delta$8.49 (s, 1H), 8.23 (s, 1H), 8.00 (d, *J* = 8.0 Hz, 1H), 7.86 (d, *J* = 8.0 Hz, 1H), 7.47 (t, *J* = 7.2 Hz, 1H), 7.37 (t, *J* = 7.6 Hz, 1H), 6.90 (brs, 2H), 6.10 (d, *J* = 5.6 Hz, 1H), 5.62 (d, *J* = 6.0Hz, 1H), 5.51 (d, *J* = 4.8 Hz, 1H), 4.59 (q, *J* = 5.2 Hz, 1H), 4.28 - 4.19 (m, 2H), 3.84 (dd, *J* = 13.6, 5.2 Hz, 1H), 3.74 (dd, *J* = 13.6, 7.6 Hz, 1H).

#### 1.7.9 Synthesis of compounds 30 and 31

<u>**Scheme S14**</u>

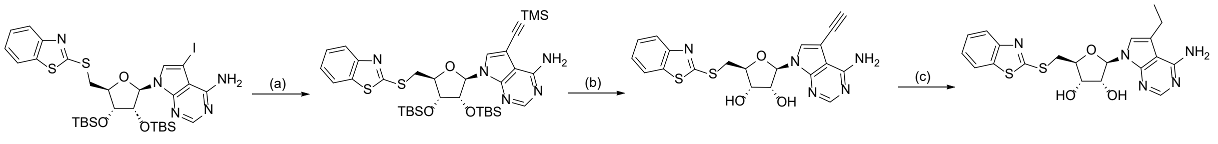

69 71 30 31

(a) TMS-acetylene, $Pd(PPh_3)_4$, $Zn(CN)_2$, DMF, 60 °C, 2 h, 62%; (b) KF, MeOH, 60 °C, 16 h, 74%; (c) $H_2$, Pd/C, RT, 2 h, 18%.

**7-[(2*R*,3*R*,4*R*,5*S*)-5-(1,3-benzothiazol-2-ylsulfanylmethyl)-3,4-bis [[tert-butyl(dimethyl) silyl]oxy]tetrahydrofuran-2-yl]-5-(2-trimethylsilylethynyl)pyrrolo[2,3-d]pyrimidin-4-amine (71)**

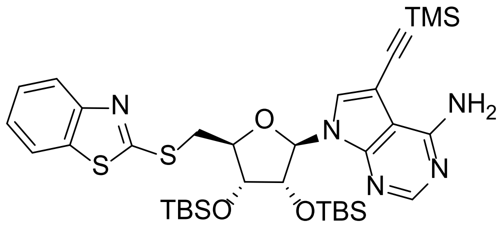

To a solution of compound **69** (200 mg, 259 µmol) in THF (10 mL) was added ethynyl (trimethyl)silane (71.9 µL, 519 µmol), triethylamine (108 µL, 779 µmol), copper iodide (14.84 mg, 77.93 µmol) and $Pd(PPh_3)_4$ (30.0 mg, 25.9 µmol). The reaction mixture was stirred at 60 °C under inert atmosphere for 2 h, cooled to RT and then concentrated under reduced pressure. The residue was purified by flash column chromatography (petroleum ether/ethyl acetate gradient 2-50%) to afford compound **71** (120 mg, 62% yield) as yellow oil. $^1$H NMR (400 MHz, $CDCl_3$): $\delta$ 8.26 (s, 1H), 7.89 - 7.87 (m, 1H), 7.77 - 7.75 (m, 1H), 7.44 - 7.40 (m, 1H), 7.36 (s, 1H), 7.33 - 7.29 (m, 1H), 5.96 (d, *J* = 6.0 Hz, 1H), 4.96 (dd, *J* = 6.4, 4.8 Hz, 1H), 4.45 - 4.43 (m, 1H), 4.31 (dd, *J* = 4.8, 2.8 Hz, 1H), 3.94 (dd, *J* = 14.0, 6.0 Hz, 1H), 3.82 (dd, *J* = 14.0, 6.8 Hz, 1H), 0.93 (s, 9H), 0.77 (s, 9H), 0.30 (s, 9H), 0.12 (s, 3H), 0.09 (s, 3H), -0.13 (s, 3H), -0.29 (s, 3H).

**(2*R*,3*R*,4*S*,5*S*)-2-(4-Amino-5-ethynyl-pyrrolo[2,3-d]pyrimidin-7-yl) -5-(1,3-benzothiazol-2-ylsulfanylmethyl)tetrahydrofuran-3,4-diol (30)**

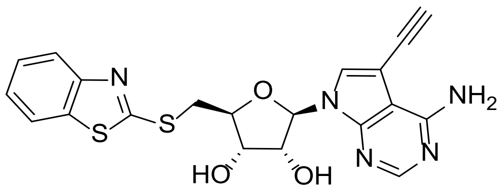

To a solution of compound **71** (100 mg, 0.14 mmol) in MeOH (10 mL) was added potassium fluoride (78.4 mg, 1.35 mmol). The reaction mixture was stirred at 70 °C for 16 h. The mixture was concentrated under reduced pressure and the residue was purified by reverse column chromatography (0.05% $NH_3{\cdot}H_2O$ in water/MeCN gradient 0-55%) to afford compound **30** (45.0 mg, 74% yield) as an off-white solid. HRMS (Method 1) *m/z* $[M+H]^+$ calcd for $C_{20}H_{17}N_5O_3S_2$ 440.0851; found 440.0851; $^1$H NMR (400 MHz, DMSO-$d_6$): $\delta$ 8.14 (s, 1H), 8.00 (d, *J* = 7.6 Hz, 1H), 7.88 - 7.86 (m, 2H), 7.47 (t, *J* = 7.6 Hz, 1H), 7.37 (t, *J* = 7.6 Hz, 1H), 6.73 (brs, 1H), 6.07 (d, *J* = 6.0 Hz, 1H), 5.58 (s, 1H), 4.58 (t, *J* = 5.6 Hz, 1H), 4.31 (s, 1H), 4.21 - 4.16 (m, 2H), 3.82 (dd, *J* = 13.2, 5.2 Hz, 1H), 3.72 (dd, *J* = 13.2, 7.2 Hz, 1H).

**(2*R*,3*R*,4*S*,5*S*)-2-(4-amino-5-ethyl-pyrrolo[2,3-d]pyrimidin-7-yl) -5-(1,3-benzothiazol-2-ylsulfanylmethyl)tetrahydrofuran-3,4-diol (31)**

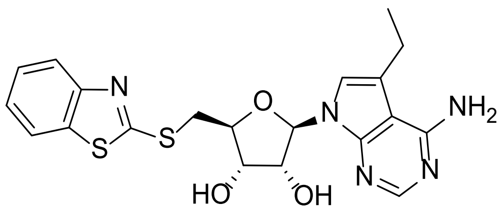

To a solution of compound **30** (20.0 mg, 45.5 µmol) in MeOH (8 mL) was added Pd/C (20.0 mg, 4.55 µmol, 10%) under inert atmosphere. The suspension was degassed under reduced pressure and purged with $H_2$ several times. The resulting mixture was stirred under hydrogen at RT for 2 h. The reaction mixture was filtered, and the filtrate was concentrated under reduced pressure. The residue was purified by reverse column chromatography (0.05% $NH_3{\cdot}H_2O$ in water/MeCN gradient 0-55%) and then by prep-HPLC (Method 11) to afford compound **31** (3.24 mg, 18% yield) as a white solid. HRMS (Method 1) *m/z* $[M+H]^+$ calcd for $C_{20}H_{21}N_5O_3S_2$ 444.1164; found 444.1124; $^1$H NMR (400 MHz, DMSO-$d_6$): $\delta$ 8.03 (s, 1H), 8.00 (d, *J* = 8.0 Hz, 1H), 7.86 (d, *J* = 8.0 Hz, 1H), 7.47 (t, *J* = 8.0 Hz, 1H), 7.36 (t, *J* = 7.6 Hz, 1H), 7.07 (s, 1H), 6.56 (s, 2H), 6.07 (d, *J* = 6.0 Hz, 1H), 5.38 - 5.35 (m, 2H), 4.52 (q, *J* = 5.2Hz, 1H), 4.16 - 4.14 (m, 2H), 3.81 (dd, *J* = 13.6, 5.2 Hz, 1H), 3.68 (dd, *J* = 13.6, 6.8 Hz, 1H), 2.75 (q, *J* = 7.2 Hz,

2H), 1.18 (t, *J* = 7.6 Hz, 3H).

##### 1.7.10 Synthesis of compound 32

**Scheme S15**

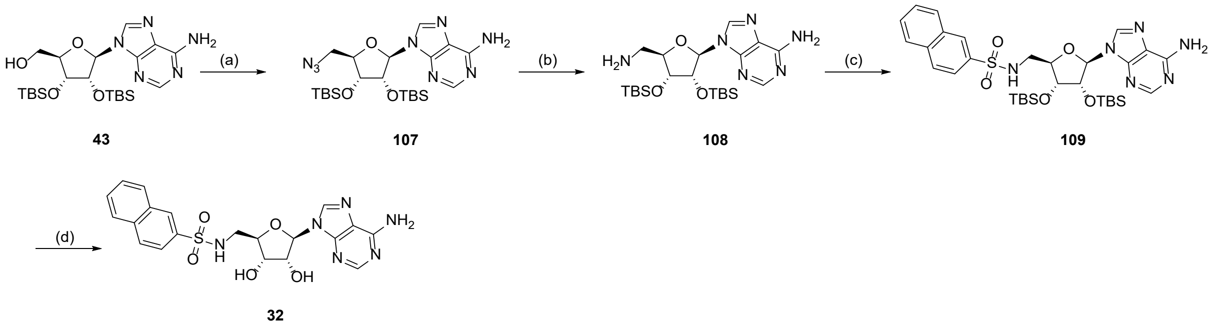

(a) DPPA, $PPh_3$, DIAD THF, 0 °C to RT, 16 h 38%; (b) $PPh_3$, THF, water, 40 °C, 16 h, 84%; (c) naphthalene-2-sulfonyl chloride, TEA, DMF, RT, 16 h, 50%; (d) KF DMF, 60 °C, 16 h, 44%.

**9-[(2*R*,3*R*,4*R*,5*R*)-5-(azidomethyl)-3,4-bis[[tert-butyl(dimethyl)silyl] oxy]tetrahydrofuran-2-yl]purin-6-amine (107)**

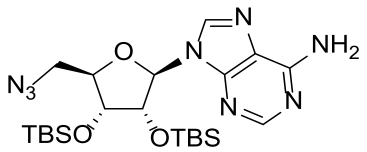

To a solution of compound **43** (1.00 g, 2.02 mmol) and triphenylphosphine (740 mg, 2.82 mmol) in THF (20 mL) was added di*iso*propyl azodicarboxylate (549 µL, 2.82 mmol), followed by a solution of diphenyl phosphoryl azide (611 µL, 2.82 mmol) in THF (10 mL) at 0 °C. The reaction mixture was warmed to RT, stirred for 16 h and then concentrated under reduced pressure. The residue was purified by flash column chromatography (petroleum ether/ethyl acetate gradient 0-25%) to afford compound **107** (400 mg, 38% yield) as a white solid. LCMS ($ES^+$, Method 1) *m/z* 521.3 $[M+H]^+$; $^1H$ NMR (400 MHz, $CDCl_3$): $\delta$ 8.35 (s, 1H), 8.01 (s, 1H), 5.90 (d, *J* = 4.4 Hz, 1H), 5.74 (s, 2H), 4.94 (t, *J* = 4.4 Hz, 1H), 4.33 (t, *J* = 4.4 Hz, 1H), 4.21 (q, *J* = 4.8 Hz, 1H), 3.73 - 3.72 (m, 2H), 0.94 (s, 9H), 0.84 (s, 9H), 0.13 (s, 3H), 0.11 (s, 3H), -0.01 (s, 3H), -0.17 (s, 3H).

**9-[(2*R*,3*R*,4*R*,5*R)*-5-(Aminomethyl)-3,4-bis[[tert-butyl(dimethyl)silyl] oxy]tetrahydrofuran-2-yl]purin-6-amine (108)**

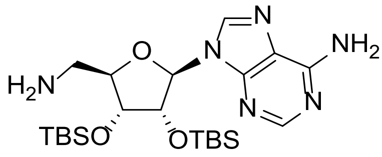

To a solution of compound **107** (400 mg, 768 µmol) in THF (8 mL) and water (0.8 mL) was added triphenylphosphine (241 mg, 921 µmol). The reaction mixture was stirred at 40 °C for 16 h and then concentrated under reduced pressure. The residue was purified by reverse column chromatography (0.05% $NH_3 \cdot H_2O$ in water/MeCN gradient 0-25%) to afford compound **108** (320 mg, 84% yield) as a yellow solid. LCMS ($ES^+$, Method 2) *m/z* 495.3 $[M+H]^+$.

**N-[[(2R,3R,4R,5R)-5-(6-Aminopurin-9-yl)-3,4-bis[[tert-butyl (dimethyl)silyl] oxy]tetrahydrofuran-2-yl]methyl]naphthalene-2-sulfonamide (109)**

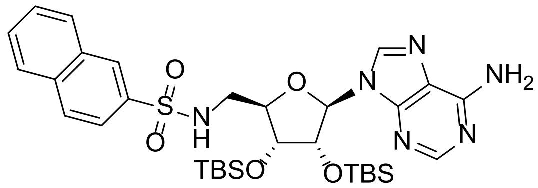

To a solution of compound **108** (50.0 mg, 101 µmol) in DMF (2 mL) was added naphthalene-2-sulfonyl chloride (22.9 mg, 101 µmol) at 0 °C and stirred at RT for 16 h. The reaction was quenched with water (10 mL) and extracted with EtOAc (3 x 10 mL). The combined organic layers were washed with brine (2 x 5 mL), dried over sodium sulfate, filtered and the filtrate was concentrated under reduced pressure. The residue was purified by preparative TLC ($SiO_2$, petroleum ether/ethyl acetate 50%) to afford compound **109** (50.0 mg, 72% yield) as a white solid. LCMS ($ES^+$, Method 2) *m/z* 669.4 $[M+H]^+$.

**N-[[(2R,3S,4R,5R)-5-(6-Aminopurin-9-yl)-3,4-dihydroxy-tetrahyd rofuran-2-yl]methyl]naphthalene-2-sulfonamide (32)**

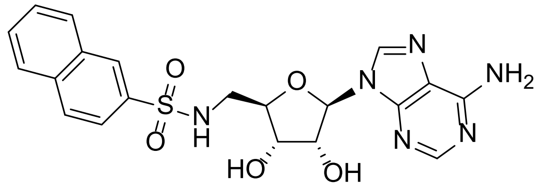

To a solution of compound **109** (50.0 mg, 72.9 µmol) in DMF (2 mL) was added potassium fluoride (42.4 mg, 729 µmol) and stirred at 60 °C for 16 h. The reaction mixture was then concentrated under reduced pressure. The residue was purified by prep-HPLC (Method 12) to afford compound **32** (14.9 mg, 44% yield) as a white solid. HRMS (Method 1) *m/z* $[M+H]^+$ calcd for $C_{20}H_{20}N_6O_5S$ 457.1294; found 457.1380; $^1H$ NMR (400 MHz, DMSO-$d_6$): $\delta$ 8.43 (s, 1H), 8.35 (s, 1H), 8.13 - 8.08 (m, 3H), 8.02 (d, *J* = 7.6 Hz, 1H), 7.80 (dd, *J* = 8.8, 2.0 Hz, 1H), 7.71 - 7.63 (m, 2H), 7.36 (s, 2H), 5.82 (d, *J* = 6.4 Hz, 1H), 5.38 (brs, 1H), 4.70 (t, *J* = 5.6 Hz, 1H), 4.09 - 4.07 (m, 1H), 4.01 - 3.99 (m, 1H), 3.10 (d, *J* = 4.8 Hz, 2H).

##### 1.7.11 Synthesis of compound 34

**Scheme S16**

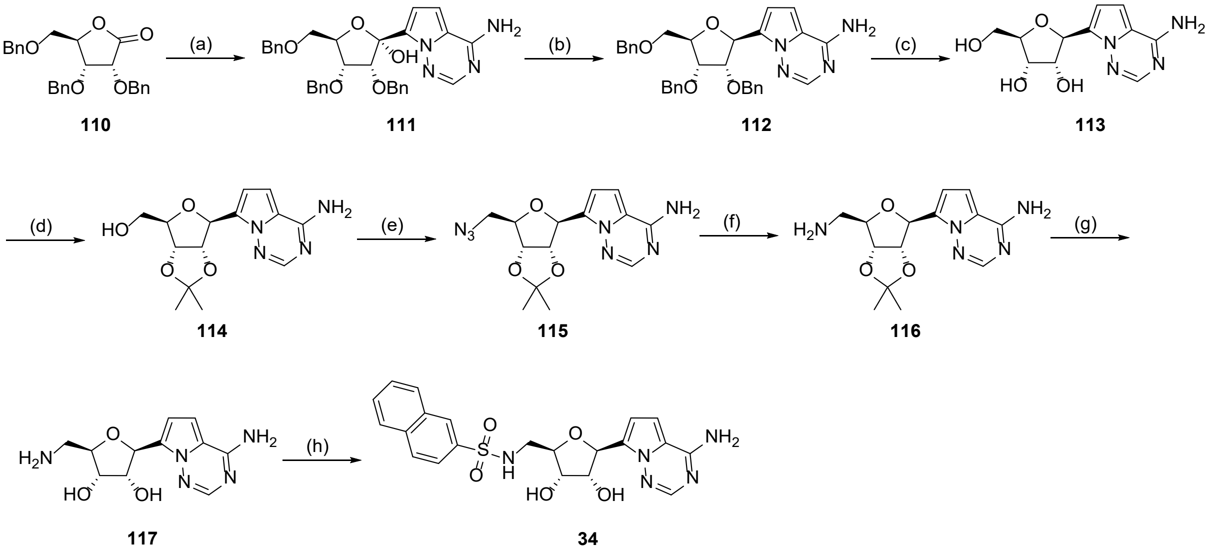

(a) 7-iodopyrrolo[2,1-f][1,2,4]triazin-4-amine 2, n-BuLi, 1,2-bis(chlorodimethylsilyl)ethane, THF, -78 °C, 26%; (b) $Et_3SiH$, $BF_3{\cdot}Et_2O$ DCM, 0 °C, RT, 97%; (c) $H_2$, Pd/C AcOH, RT, 16 h, 84%; (d) 2,2-dimethoxypropane, TsOH, DMF, Acetone, 60 °C, 16 h, 92%; (e) DPPA, DIAD, $PPh_3$, THF, 0 °C to RT, 16 h, 57%; (f) $PPh_3$, $H_2O$, THF, 40 °C, 16 h, 18% (g) HCl, MeOH, $H_2O$, RT, 2 h, 69%; (h) naphthalene-2-sulfonyl chloride TEA, DMF, 0 °C, 2 h, 17%.

**(2*S*,3*R*,4*R*,5*R*)-2-(4-Aminopyrrolo[2,1-f][1,2,4]triazin-7-yl)-3,4- dibenzyloxy-5-(benzyloxy methyl)tetrahydrofuran-2-ol (111)**

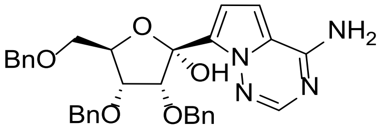

To a suspension of 7-iodopyrrolo[2,1-f][1,2,4]triazin-4-amine (4.92 g, 18.9 mmol) and chloro-[2-[chloro(dimethyl)silyl]ethyl]-dimethyl-silane (4.07 g, 18.9 mmol) in THF (70 mL) was added n-butyllithium (24.1 mL, 2.5 M in hexane) at -78 °C under inert atmosphere. The reaction mixture was stirred for 30 min prior to the addition of a solution of compound **110** (7.20 g, 17.2 mmol) in THF (20 mL) and then further stirred at -78 °C for 1 h. The reaction was quenched with sat. aq. ammonium chloride solution (150 mL) at 0 °C and then extracted with EtOAc (3 x 50 mL). The combined organic layers were washed with sat. aq. sodium bicarbonate (80 mL), brine (80 mL), dried over sodium sulfate, filtered, and the filtrate was concentrated under reduced pressure. The residue was purified by flash column chromatography (petroleum ether/ethyl acetate gradient 2-100%) to afford compound **111** (2.50 g, 26% yield) as yellow oil. LCMS (ES$^+$, Method 2) *m/z* 575.4 [M+Na]$^+$; $^1$H NMR (400 MHz, DMSO-$d_6$): $\delta$ 8.07 (s, 2H), 7.99 (s, 1H), 7.36 - 7.23 (m, 13H), 7.17 - 7.12 (m, 2H), 6.99 - 6.98 (m, 1H), 6.94 (d, *J* = 4.8 Hz, 1H), 5.38 (d, *J* = 6.0 Hz, 1H), 5.07 (d, *J* = 5.2 Hz, 1H), 7.57 (dd, *J* = 11.6, 2.4 Hz, 2H), 4.49 - 4.44 (m, 4H), 4.01 - 4.00 (m, 1H), 3.92 (t, *J* = 6.0 Hz, 1H), 3.68 (dd, *J* = 10.0, 3.2 Hz, 1H), 3.46 (dd, *J* = 10.0, 6.4 Hz, 1H).

**7-[(2*S*,3*S*,4*R*,5*R*)-3,4-Dibenzyloxy-5-(benzyloxymethyl) tetrahydrofuran-2-yl]pyrrolo[2,1-f][1,2,4]triazin-4-amine (112)**

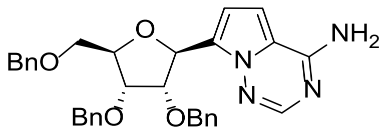

To a solution of compound **111** (2.00 g, 3.62 mmol) and triethylsilane (1.50 mL, 9.41 mmol) in DCM (20 mL) was added dropwise boron trifluoride diethyl etherate (1.43 mL, 5.43 mmol, 47% purity) at 0 °C under inert atmosphere and stirred for 2 h. The reaction mixture was quenched with sat. aq. sodium bicarbonate (50 mL) at 0 °C, and then extracted with DCM (3 x 20 mL). The combined organic layers were dried over sodium sulfate, filtered and the filtrate was concentrated under reduced pressure. The residue was purified by flash column chromatography (petroleum ether/ethyl acetate gradient 0-50%) to afford compound **112** (1.90 g, 97% yield) As yellow oil. LCMS (ES$^+$, Method 3) *m/z* 537.4 [M+H]$^+$; $^1$H NMR (400 MHz, $CDCl_3$): $\delta$ 7.92 (s, 1H), 7.36 - 7.28 (s, 15H), 6.73 (d, *J* = 4.8 Hz, 1H), 6.60 (d, *J* = 4.4 Hz, 1H), 5.68 (d, *J* = 4.0 Hz, 1H), 4.74 (s, 2H), 4.63 - 4.53 (m, 3H), 4.47 - 4.44 (m, 1H), 4.42 - 4.40 (m, 1H), 4.27 (t, *J* = 4.4 Hz, 1H), 4.16 - 4.14 (m, 1H), 3.79 (dd, *J* = 10.8, 3.6 Hz, 1H), 3.67 (dd, *J* = 10.8, 4.0 Hz, 1H).

**(2*S*,3*R*,4*S*,5*R*)-2-(4-Aminopyrrolo[2,1-f][1,2,4]triazin-7-yl) -5-(hydroxymethyl) tetra hydrofuran-3,4-diol (113)**

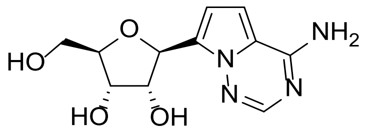

To a solution compound **112** (1.90 g, 3.54 mmol) in acetic acid (30 mL) was added Pd/C (1.00 g, 354 µmol, 10% wt.) under inert atmosphere. The reaction mixture was degassed and refilled with $H_2$ three times and stirred under a $H_2$ atmosphere (50 psi) at RT for 16 h. The mixture was filtered, and the filtrate was concentrated under reduced pressure to afford compound **113** (800 mg, 84% yield) as a white solid. The isolated product was used in the following step without any further purification. LCMS ($ES^+$, Method 4) *m/z* 267.1 $[M+H]^+$; $^1$H NMR (400 MHz, DMSO-$d_6$): $\delta$ 7.82 (s, 1H), 7.68 (s, 1H), 6.84 (d, $J$ = 4.8 Hz, 1H), 6.68 (d, $J$ = 4.8 Hz, 1H), 5.11 (d, $J$ = 6.8 Hz, 1H), 4.23 (t, $J$ = 6.0 Hz, 1H), 3.95 (t, $J$ = 5.6 Hz, 1H), 3.79 (q, $J$ = 4.8 Hz, 1H), 3.55 (dd, $J$ = 12.0, 4.0 Hz, 1H), 3.45 (dd, $J$ = 12.4, 5.2 Hz, 1H).

**[(3a*S*,4*S*,6*R*,6a*R*)-4-(4-Aminopyrrolo[2,1-f][1,2,4]triazin-7-yl)-2,2 -dimethyl-3a,4,6,6a-tetrahydrofuro[3,4-d][1,3]dioxol-6-yl]methanol (114)**

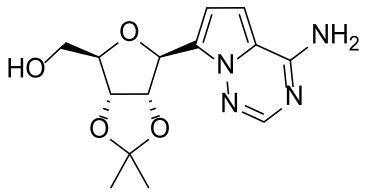

To a solution of compound **113** (800 mg, 3.00 mmol) in DMF (10 mL) and acetone (10 mL) was added 2,2-dimethoxypropane (368 µL, 3.00 mmol) and 4-methylbenzenesulfonic acid (628 mg, 3.31 mmol) and stirred at 60 °C for 16 h. The reaction mixture was cooled to RT, quenched with sat. aq. sodium bicarbonate (50 mL) and extracted with EtOAc (3 x 30 mL). The combined organic layers were washed with brine (3 x 20 mL), dried over sodium sulfate, filtered and the filtrate was concentrated under reduced pressure. The residue was purified by reverse column chromatography (0.05% TFA in water/MeCN gradient 0-55%) to afford compound **114** (850 mg, 92% yield) as a white solid. LCMS ($ES^+$, Method 3) *m/z* 307.2 $[M+H]^+$; $^1$H NMR (400 MHz,

DMSO-$d_6$): $\delta$ 7.85 (s, 1H), 7.74 (s, 2H), 6.84 (d, $J$ = 4.4 Hz, 1H), 6.74 (d, $J$ = 4.4 Hz, 1H), 5.22 (d, $J$ = 4.8 Hz, 1H), 5.03 (t, $J$ = 5.2 Hz, 1H), 4.88 (t, $J$ = 6.0 Hz, 1H), 4.72 (dd, $J$ = 6.8, 4.0 Hz, 1H), 3.97 (q, $J$ = 4.0 Hz, 1H), 3.48 (t, $J$ = 5.6 Hz, 2H), 1.50 (s, 3H),1.30 (s, 3H).

**7-[(3a*S*,4*S*,6*R*,6a*R*)-6-(Azidomethyl)-2,2-dimethyl-3a,4,6,6a-tetrahydrofuro[3,4-d][1,3] dioxol-4-yl]pyrrolo[2,1-f][1,2,4]triazin-4-amine (115)**

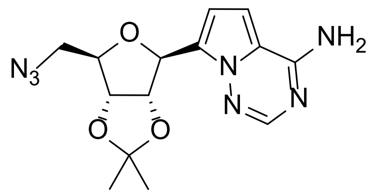

To a solution of compound **114** (480 mg, 1.57 mmol) and triphenylphosphine (575 mg, 2.19 mmol) in THF (6 mL) was added di*iso*propyl azodicarboxylate (425 µL, 2.19 mmol), followed by a solution of diphenyl phosphoryl azide (473 µL 2.19 mmol) in THF (2 mL) at 0°C under inert atmosphere. The reaction mixture was warmed to RT, stirred for 16 h and then concentrated under reduced pressure. The residue was purified by reverse column chromatography (0.05% TFA in water/MeCN gradient 0-25%) to afford compound **115** (300 mg, 57% yield) as a white solid. LCMS (ES$^+$, Method 1) *m/z* 332.2 [M+H]$^+$.

**7-[(3a*S*,4*S*,6*R*,6a*R*)-6-(Aminomethyl)-2,2-dimethyl-3a,4,6,6a- tetrahydrofuro[3,4-d][1,3] dioxol-4-yl]pyrrolo[2,1-f][1,2,4]triazin-4-amine (116)**

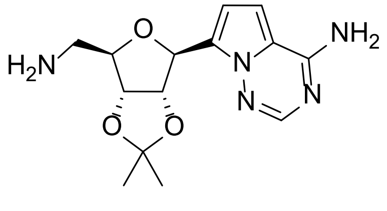

To a solution of compound **115** (300 mg, 0.91 mmol) in THF (5 mL) and water (0.5 mL) was added triphenylphosphine (284 mg, 1.09 mmol) and stirred at 40 °C for 16 h. The reaction mixture was cooled to RT and then concentrated under reduced pressure. The residue was purified by reverse column chromatography (0.05% $NH_3{\cdot}H_2O$ in water/MeCN gradient 0-25%) to afford compound **116** (50.0 mg, 18% yield) as a yellow solid. LCMS (ES$^+$, Method 2) *m/z* 306.21 [M+H]$^+$.

**(2*R*,3*S*,4*R*,5*S*)-2-(Aminomethyl)-5-(4-aminopyrrolo[2,1-f][1,2,4] triazin-7-yl)tetrahydrofuran -3,4-diol (117)**

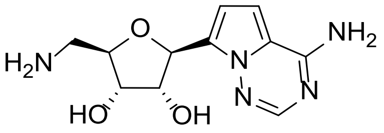

To a solution of compound **116** (50.0 mg, 163 µmol) in MeOH (3 mL) was added hydrochloric acid (3 mL, 1N in MeOH) and stirred at RT for 2 h. The reaction mixture was then concentrated under reduced pressure. The residue was purified by reverse column chromatography (0.05% $NH_3·H_2O$ in water/MeCN gradient 0-5%) to afford compound **117** (30.0 mg, 69% yield) as a yellow solid. LCMS ($ES^+$, Method 6) *m/z* 266.3 $[M+H]^+$.

***N*-[[(2*R*,3*S*,4*R*,5*S*)-5-(4-Aminopyrrolo[2,1-f][1,2,4]triazin-7-yl)-3,4- dihydroxy-tetrahydrofuran-2-yl]methyl]naphthalene-2-sulfonamide (34)**

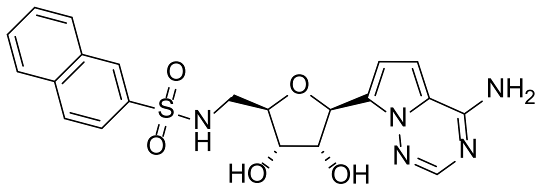

To a solution of compound **117** (30.0 mg, 113 µmol) in DMF (1 mL) was added triethylamine (47.2 µL 339 µmol) and naphthalene-2-sulfonyl chloride (25.6 mg, 113 µmol) at 0 °C and stirred for 2 h. The reaction mixture was concentrated under reduced pressure. The residue was purified by prep-HPLC (Method 13) to afford compound **34** (9.23 mg, 17% yield) as an off-white solid. HRMS (Method 1) *m/z* $[M+H]^+$ calcd for $C_{21}H_{21}N_5O_5S$ 456.1342; found 456.1329; $^1H$ NMR (400 MHz, DMSO-$d_6$): $\delta$ 8.45 (s, 1H), 8.12 (t, *J* = 8.0 Hz, 2H), 8.03 (d, *J* = 8.0 Hz, 1H), 7.97 (t, *J* = 6.0 Hz, 1H), 7.83 - 7.81 (m, 2H), 7.75 - 7.64 (m, 4H), 6.81 (d, *J* = 4.4 Hz, 1H), 6.62 (d, *J* = 4.4 Hz, 1H), 5.03(t, *J* = 6.4 Hz, 2H), 4.98 (d, *J* = 5.2 Hz, 1H), 4.29 (q, *J* = 6.0 Hz, 1H), 3.90 - 3.86 (m, 1H), 3.85 - 3.82 (m, 1H), 3.03 - 3.01 (m, 1H), 2.96 - 2.94 (m, 1H).

##### 1.7.12 Synthesis of compounds 35-38

<u>**Scheme S17**</u>

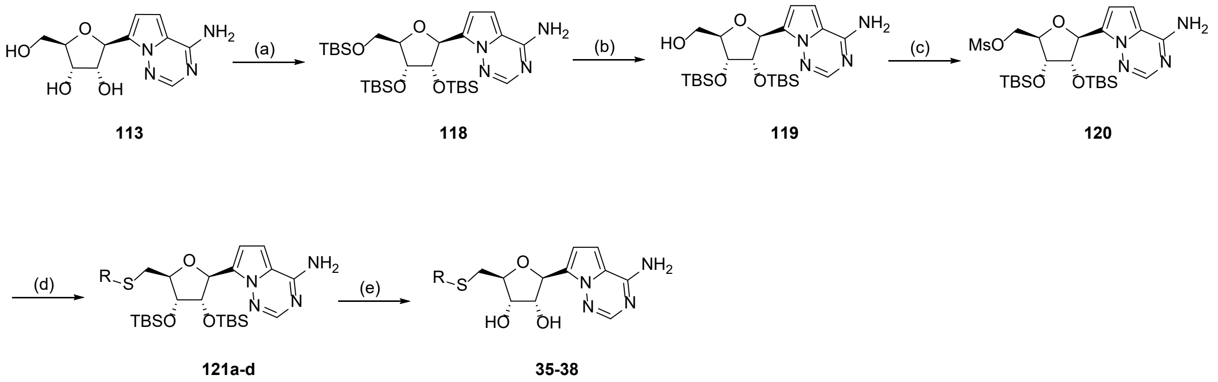

(a) TBSCl, imidazole, DMF, RT, 2 h, 94%; (b) $CCl_3COOH$, THF, 0 °C, 4 h, 98%; (c) $Ms_2O$ TEA, THF, 0 °C, 2 h, 52%; (d) RSH, NaH, THF, 0-70 °C, 16 h, 69-89%; (e) KF, DMF, 70 °C, 16 h, 13-72 %.

#### 7-[(2*S*,3*S*,4*R*,5*R*)-3,4-Bis[[tert-butyl(dimethyl)silyl]oxy]-5-[[tert-butyl(dimethyl) silyl]oxymethyl]tetrahydrofuran-2-yl]pyrrolo[2,1-f][1,2,4]triazin-4-amine (118)

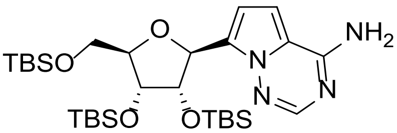

To a solution of compound **113** (3.00 g, 11.3 mmol) in DMF (30 mL) was added imidazole (8.44 g, 124 mmol) and *tert*-butylchloro dimethylsilane (7.62 mL, 61.9 mmol) and stirred at RT for 2 h. The reaction mixture was quenched with water (100 mL) and extracted with EtOAc (3 x 50 mL). The combined organic layers were washed with brine (4 x 20 mL), dried over sodium sulfate, filtered and the filtrate was concentrated under reduced pressure. The residue was purified by flash column chromatography (petroleum ether/ethyl acetate gradient 0-20%) to afford compound **118** (6.50 g, 94% yield) as yellow oil. LCMS ($ES^+$, Method 2) *m/z* 609.5 $[M+H]^+$; $^1H$ NMR (400 MHz, $CDCl_3$): $\delta$ 7.89 (s, 1H), 6.75 (d, $J$ = 4.8 Hz, 1H), 6.59 (d, $J$ = 4.4 Hz, 1H), 5.37 (d, $J$ = 5.6 Hz, 1H), 4.53 - 4.51 (m, 1H), 4.26 - 4.24 (m, 1H), 4.06 (q, $J$ = 4.0 Hz, 1H), 3.90 (dd, $J$ = 10.8, 4.8 Hz, 1H), 3.75 (dd, $J$ = 11.2, 3.6 Hz, 1H), 0.91 (s, 9H), 0.80 (s, 9H), 0.08 (s, 3H), 0.06 (s, 3H), -0.06 (s, 3H), -0.21 (s, 3H).

#### [(2*R*,3*R*,4*S*,5*S*)-5-(4-Aminopyrrolo[2,1-f][1,2,4]triazin-7-yl) -3,4-bis[[tert-butyl(dimethyl)silyl]oxy]tetrahydrofuran-2-yl]methanol (119)

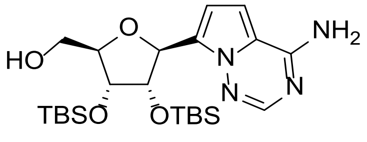

To a solution of compound **118** (5.00 g, 8.21 mmol) in THF (50 mL) was added a solution of 2,2,2-trichloroacetic acid (13.4 g, 82.2 mmol) in water (15 mL) at 0 °C and stirred for 4 h. The reaction mixture was quenched by sat. aq. sodium bicarbonate (100 mL) and extracted with EtOAc (3 x 50 mL). The combined organic layers were washed with brine (3 x 20 mL), dried over sodium sulfate, filtered and the filtrate was concentrated under reduced pressure. The residue was purified by flash column chromatography (petroleum ether/ethyl acetate gradient 0-25%) to afford compound **119** (4.00 g, 98% yield) as a white solid. LCMS (ES$^+$, Method 2) *m/z* 495.4 [M+H]$^+$; $^1$H NMR (400 MHz, $CDCl_3$): *δ* 7.92 - 7.91 (m, 1H), 6.71 - 6.70 (m, 1H), 6.61 (d, *J* = 4.4 Hz, 1H), 5.99 (brs, 2H), 5.28 (s, 1H), 5.11 (d, *J* = 9.2 Hz, 1H), 4.90 (dd, *J* = 8.8, 4.8 Hz, 1H), 4.29 (d, *J* = 4.8 Hz, 1H), 4.15 - 4.10 (m, 1H), 3.89 (dd, *J* = 12.4, 2.0 Hz, 1H), 3.70 (d, *J* = 12.4 Hz, 1H), 0.95 (s, 9H), 0.73 (s, 9H), 0.12 (s, 3H), 0.11 (s, 3H), -0.19 (s, 3H), -0.67 (s, 3H).

**[(2*R*,3*R*,4*S*,5*S*)-5-(4-Aminopyrrolo[2,1-f][1,2,4]triazin-7-yl)-3,4-bis [[tert-butyl(dimethyl)silyl]oxy]tetrahydrofuran-2-yl]methyl methanesulfonate (120)**

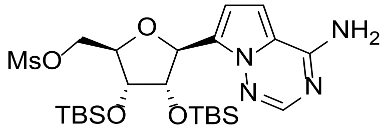

To a solution of compound **119** (1.00 g, 2.02 mmol) and triethylamine (2.81 mL, 20.2 mmol) in THF (10 mL) was added a solution of methanesulfonic anhydride (1.06 g, 6.06 mmol) in THF (5 mL) at 0 °C under inert atmosphere and stirred for 2 h. The reaction mixture was quenched with sat. aq. sodium bicarbonate (20 mL) at 0 °C, and then extracted with EtOAc (3 x 20 mL). The combined organic layers were washed with brine (3 x 10 mL), dried over sodium sulfate, filtered and the filtrate was concentrated under reduced pressure. The residue was purified by flash column chromatography (petroleum ether/ethyl acetate gradient 0-50%) to afford compound **120** (600 mg, 52% yield) as a white solid. LCMS (ES$^+$, Method 2) *m/z* 573.4 [M+H]$^+$; $^1$H NMR (400 MHz, $CDCl_3$): *δ* 7.94 (s, 1H), 6.72 (d, *J* = 4.4 Hz, 1H), 6.62 (d, *J* = 4.4 Hz, 1H), 5.67 (s, 2H), 5.39 (d, *J* = 3.6 Hz, 1H), 4.55 - 4.50 (m, 2H), 4.42 (dd, *J* = 11.2, 4.8 Hz, 1H), 4.29 - 4.21 (m, 2H), 2.96 (s, 3H), 0.93 (s, 9H), 0.87 (s, 9H), 0.11 (s, 3H), 0.09 (s, 3H), 0.02 (s, 3H), -0.06 (s, 3H).

**7-[(2*S*,3*S*,4*R*,5*S*)-5-(1,3-Benzothiazol-2-ylsulfanylmethyl) -3,4-bis[[tert-butyl(dimethyl) silyl]oxy]tetrahydrofuran-2-yl]pyrrolo[2,1-f][1,2,4]triazin-4-amine (121a)**

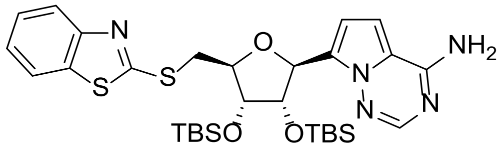

To a solution of compound **120** (100 mg, 174 µmol) in DMF (1 mL) was added 1,3-benzothiazol-2-ylsulfanyl sodium (165 mg, 872 µmol) and stirred at 50 °C for 16 h. The reaction mixture was cooled to RT, quenched with water (20 mL) and extracted with EtOAc (3 x 10 mL). The combined organic layers were washed with brine (3 x 10 mL), dried over sodium sulfate, filtered and the filtrate was concentrated under reduced pressure. The residue was purified by flash column chromatography (petroleum ether/ethyl acetate gradient 0-50%) to afford compound **121a** (100 mg, 89% yield) as a yellow solid. LCMS ($ES^+$, Method 2) *m/z* 644.4 $[M+H]^+$; $^1H$ NMR (400 MHz, $CDCl_3$): $\delta$ 7.91 (s, 1H), 7.85 (d, *J* = 8.0 Hz, 1H), 7.75 (d, *J* = 7.6 Hz, 1H), 7.42 (t, *J* = 7.2 Hz, 1H), 7.32 - 7.29 (m, 1H), 6.77 (d, *J* = 4.8 Hz, 1H), 6.64 (d, *J* = 4.8 Hz, 1H), 5.70 (s, 2H), 5.34 (d, *J* = 5.6 Hz, 1H), 4.75 (t, *J* = 4.4 Hz, 1H), 4.44 (q, *J* = 4.4 Hz, 1H), 4.26 (t, *J* = 4.0 Hz, 1H), 3.95 (dd, *J* = 13.6, 5.6 Hz, 1H), 3.70 (dd, *J* = 13.6, 6.4 Hz, 1H), 0.94 (s, 9H), 0.80 (s, 9H), 0.11 (s, 3H), 0.09 (s, 3H), -0.07 (s, 3H), -0.24 (s, 3H).

**7-[(2S,3S,4R,5S)-3,4-Bis[[tert-butyl(dimethyl)silyl]oxy]-5-[(6-methyl -1,3-benzothiazol-2-yl)sulfanylmethyl]tetrahydrofuran-2-yl]pyrrolo[2,1-f][1,2,4]triazin-4-amine (121b)**

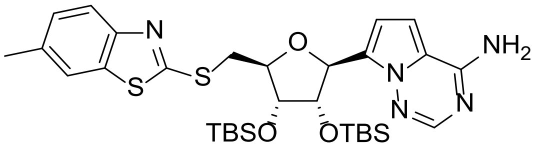

To a solution of 6-methyl-1,3-benzothiazole-2-thiol (63.3 mg, 349 µmol) in THF (2 mL) was added sodium hydride (13.9 mg, 349 µmol, 60% dispersion in mineral oil) at 0°C under inert atmosphere and stirred for 30 min. A solution of compound **120** (100 mg, 174 µmol) in THF (2 mL) was added and gradually heated to 70 °C and stirred for 16 h. The reaction mixture was cooled to RT, quenched with water (20 mL) and extracted with EtOAc (3 x 10 mL). The combined organic layers were washed with brine (10 mL), dried over sodium sulfate, filtered and the filtrate was concentrated under reduced pressure. The residue was purified by flash column chromatography (petroleum ether/ethyl acetate gradient 0-50%) to afford compound **121b** (80.0 mg, 69% yield) as a white solid. LCMS (ES$^+$, Method 2) *m/z* 658.4 [M+H]$^+$; $^1$H NMR (400 MHz, $CDCl_3$): *δ* 7.91 (s, 1H), 7.73 (d, *J* = 8.4 Hz, 1H), 7.54 (s, 1H), 7.22 (d, *J* = 8.4 Hz, 1H), 6.77 (d, *J* = 4.8 Hz, 1H), 6.64 (d, *J* = 4.8 Hz, 1H), 5.67 (s, 2H), 5.34 (d, *J* = 5.6 Hz, 1H), 4.75 (dd, *J* = 5.6, 4.8 Hz, 1H), 4.44 - 4.40 (m, 1H), 4.25 (t, *J* = 4.0 Hz, 1H), 3.92 (dd, *J* = 13.6, 5.6 Hz, 1H), 3.68 (dd, *J* = 13.6, 6.4 Hz, 1H), 0.93 (s, 9H), 0.80 (s, 9H), 0.10 (s, 3H), 0.08 (s, 9H), -0.10 (s, 3H), -0.30 (s, 3H).

**7-[(2*S*,3*S*,4*R*,5*S*)-3,4-Bis[[tert-butyl(dimethyl)silyl]oxy]-5- (2-naphthylsulfanylmethyl) tetrahydrofuran-2-yl]pyrrolo[2,1-f][1,2,4]triazin-4-amine (121c)**

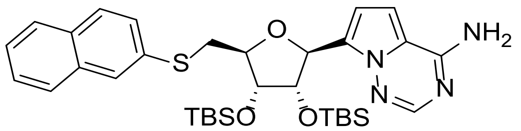

To a solution of naphthalene-2-thiol (55.9 mg, 349 µmol) in THF (2 mL) was added sodium hydride (13.9 mg, 349 µmol, 60% dispersion in mineral oil) at 0 °C under inert atmosphere and stirred for 30 min. A solution of compound **120** (100 mg, 174 µmol) in THF (2 mL) was added, gradually then heated to 70 °C and stirred for 16 h. The reaction mixture was cooled to RT, quenched with water (20 mL) and extracted with EtOAc (3 x 10 mL). The combined organic layers were washed with brine (10 mL), dried over sodium sulfate, filtered and the filtrate was concentrated under reduced pressure. The residue was purified by flash column chromatography (petroleum ether/ethyl acetate gradient 0-50%) to afford compound **121c** (80.0 mg, 72% yield) as a white solid. LCMS (ES$^+$, Method 2) *m/z* 637.5 [M+H]$^+$; $^1$H NMR (400 MHz, $CDCl_3$): *δ* 7.92 (s, 1H), 7.80 (s, 1H), 7.78 (d, *J* = 8.0 Hz, 1H), 7.73 (d, *J* = 8.8 Hz, 1H), 7.70 (d, *J* = 7.6 Hz, 1H), 7.48 - 7.41 (m, 3H), 6.74 (d, *J* = 4.4 Hz, 1H), 6.60 (d, *J* = 4.4 Hz, 1H), 5.62 (s, 2H), 5.33 (d, *J* = 5.6 Hz, 1H), 4.72 (t, *J* = 4.8 Hz, 1H), 4.32 (q, *J* = 4.8 Hz, 1H), 4.24 (t, *J* = 4.4 Hz, 1H), 3.53 (dd, *J* = 14.0, 6.4 Hz, 1H), 3.36 (dd, *J* = 14.0, 6.0 Hz, 1H), 0.91 (s, 9H), 0.79 (s, 9H), 0.08 (s, 3H), 0.07 (s, 3H), 0.01 (s, 3H), -0.24 (s, 3H).

**7-[(2*S*,3*S*,4*R*,5*S*)-3,4-Bis[[tert-butyl(dimethyl)silyl]oxy]- 5-(2-quinolylsulfanylmethyl) tetrahydrofuran-2-yl]pyrrolo[2,1-f][1,2,4]triazin-4- amine (121d)**

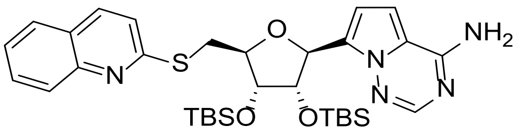

To a solution of quinoline-2-thiol (56.3 mg, 349 µmol) in THF (2 mL) was added sodium hydride (13.9 mg, 349 µmol, 60% dispersion in mineral oil) at 0°C under inert atmosphere and stirred for 30 min. A solution of compound **120** (100 mg, 174 µmol) in THF (2 mL) was then added and gradually heated to 70 °C and stirred for 16 h. The reaction mixture was cooled to RT, quenched with water (20 mL) and extracted with EtOAc (3 x 10 mL). The combined organic layers were washed with brine (10 mL), dried over sodium sulfate, filtered and the filtrate was concentrated under reduced pressure. The residue was purified by flash column chromatography (petroleum ether/ethyl acetate gradient 0-50%) to afford compound **121d** (80.0 mg, 72%) as a white solid. LCMS (ES$^+$, Method 2) *m/z* 638.5 [M+H]$^+$; $^1$H NMR (400 MHz, $CDCl_3$): *δ* 7.95 - 7.94 (m, 2H), 7.89 (d, *J* = 8.4 Hz, 1H), 7.72 (d, *J* = 8.0 Hz, 1H), 7.64 (td, *J* = 7.2, 1.6 Hz, 1H), 7.44 (t, *J* = 6.8 Hz, 1H), 7.25 (d, *J* = 8.8 Hz, 1H), 6.79 (d, *J* = 4.4 Hz, 1H), 6.63 (d, *J* = 4.4 Hz, 1H), 5.70 (s, 2H), 5.36 (d, *J* = 6.0 Hz, 1H), 4.79 (dd, *J* = 5.6, 4.4 Hz, 1H), 4.43 - 4.39 (m, 1H), 4.24 (t, *J* = 4.0 Hz, 1H), 3.99 (dd, *J* = 14.0, 6.4 Hz, 1H), 3.70 (dd, *J* = 14.0, 6.8 Hz, 1H), 0.92 (s, 9H), 0.77 (s, 9H), 0.08 (s, 3H), 0.06 (s, 3H), -0.15 (s, 3H), -0.28 (s, 3H).

**(2*S*,3*R*,4*S*,5*S*)-2-(4-Aminopyrrolo[2,1-f][1,2,4]triazin-7-yl)-5-(1,3-benzothiazol-2-ylsulfanylmethyl)tetrahydrofuran-3,4-diol (35)**

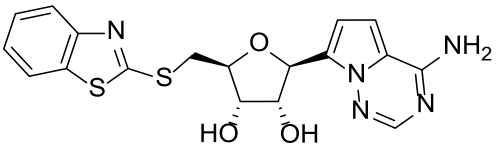

To a solution of compound **121a** (100 mg, 0.16 mmol) in DMF (1 mL) was added potassium fluoride (90.2 mg, 1.55 mmol) and stirred at 70 °C for 16 h. The reaction mixture was cooled to RT and then concentrated under reduced pressure. The residue was purified by prep-HPLC (Method 14) to afford compound **35** (59.6 mg, 91% yield) as a white solid. HRMS (Method 1) *m/z* [M+H]$^+$ calcd for $C_{18}H_{17}N_5O_3S_2$ 416.0851; found 416.0830; $^1$H NMR (400 MHz, DMSO-*d*$_6$): *δ* 8.00 (d, *J* = 7.6 Hz, 1H), 7.86 (d, *J* = 8.0 Hz, 1H), 7.84 (s, 1H), 7.70 (brs, 2H), 7.47 (td, *J* = 8.0, 0.8 Hz, 1H), 7.36 (td, *J* = 8.0, 0.8Hz, 1H), 6.85 (d, *J* = 4.4 Hz, 1H), 6.70 (d, *J* = 4.4 Hz, 1H), 5.19

(dd, *J* = 6.0, 4.0 Hz, 2H), 5.15 (d, *J* = 6.0 Hz, 1H), 4.35 (q, *J* = 5.6 Hz, 1H), 4.12 (q, *J* = 6.4Hz, 1H), 4.03 (q, *J* = 4.8 Hz, 1H), 3.77 (dd, *J* = 13.2, 4.8 Hz, 1H), 3.60 (dd, *J* = 13.2, 6.8 Hz, 1H).

**(2*S*,3*R*,4*S*,5*S*)-2-(4-Aminopyrrolo[2,1-f][1,2,4]triazin-7-yl)-5- [(6-methyl-1,3-benzothiazol-2-yl)sulfanylmethyl]tetrahydrofuran-3,4-diol (36)**

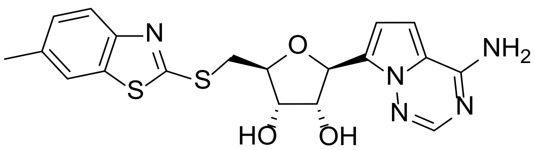

To a solution of compound **121b** (80.0 mg, 0.12 mmol) in DMF (1 mL) was added potassium fluoride (70.6 mg, 1.22 mmol) and stirred at 70 °C for 16 h. The reaction mixture was cooled to RT and then concentrated under reduced pressure. The residue was purified by prep-HPLC (Method 15) to afford compound **36** (6.85 mg, 13% yield) as a white solid. HRMS (Method 1) *m/z* $[M+H]^+$ calcd for $C_{19}H_{19}N_5O_3S_2$ 430.1008; found 430.1006; $^1H$ NMR (400 MHz, DMSO-$d_6$): $\delta$ 7.83 (s, 1H), 7.79 (s, 1H), 7.74 (d, *J* = 8.0 Hz, 1H), 7.70 (brs, 1H), 7.28 (d, *J* = 7.6 Hz, 1H), 6.85 (d, *J* = 4.4 Hz, 1H), 6.69 (d, *J* = 4.4 Hz, 1H), 5.17 (t, *J* = 6.4 Hz, 2H), 5.13 (d, *J* = 6.4 Hz, 1H), 4.34 (q, *J* = 5.2 Hz, 1H), 4.10 (d, *J* = 6.4 Hz, 1H), 4.02 (q, *J* = 4.8 Hz, 1H), 3.73 (dd, *J* = 13.6, 5.2 Hz, 1H), 3.57 (dd, *J* = 12.8, 6.4 Hz, 1H), 2.41 (s, 3H).

**(2*S*,3*R*,4*S*,5*S*)-2-(4-Aminopyrrolo[2,1-f][1,2,4]triazin-7-yl)- 5-(2-naphthylsulfanylmethyl) tetrahydrofuran-3,4-diol (37)**

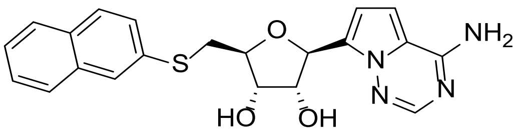

To a solution of compound **121c** (80.0 mg, 0.13 mmol) in DMF (1 mL) was added potassium fluoride (72.9 mg, 1.26 mmol) and stirred at 70 °C for 16 h. The reaction mixture was cooled to RT and then concentrated under reduced pressure. The residue was purified by prep-HPLC (Method 16) to afford compound **37** (21.6 mg, 42% yield) as a white solid. HRMS (Method 1) *m/z* $[M+H]^+$ calcd for $C_{21}H_{20}N_4O_3S$ 409.1334; found 409.1345; $^1H$ NMR (400 MHz, DMSO-$d_6$): $\delta$ 7.89 - 7.82 (m, 4H), 7.81 (d, *J* = 8.0 Hz, 1H), 7.69 (brs, 2H), 7.51 - 7.43 (m, 3H), 6.84 (d, *J* = 4.8

Hz, 1H), 6.70 (d, *J* = 4.4 Hz, 1H), 5.18 (d, *J* = 6.0 Hz, 1H), 5.10 (d, *J* = 5.6 Hz, 2H), 4.34 (q, *J* = 5.2 Hz, 1H), 4.04 - 3.97 (m, 2H), 3.42 (dd, *J* = 13.6, 4.8 Hz, 1H), 3.30 - 3.25 (m, 1H).

**(2*S*,3*R*,4*S*,5*S*)-2-(4-Aminopyrrolo[2,1-f][1,2,4]triazin-7-yl)-5- (2-quinolylsulfanylmethyl) tetrahydrofuran-3,4-diol (38)**

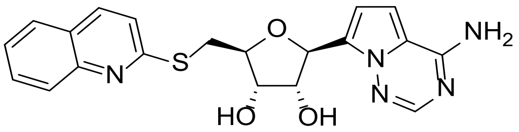

To a solution of compound **121d** (80.0 mg, 0.13 mmol) in DMF (1 mL) was added potassium fluoride (72.8 mg, 1.25 mmol) and stirred at 70 °C for 16 h. The reaction mixture was cooled to RT and then concentrated under reduced pressure. The residue was purified by prep-HPLC (Method 17) to afford compound **38** (9.52 mg, 19% yield) as a white solid. HRMS (Method 1) *m/z* $[M+H]^+$ calcd for $C_{20}H_{19}N_5O_3S$ 410.1287; found 410.1289; $^1H$ NMR (400 MHz, DMSO-$d_6$): $\delta$ 8.16 (d, *J* = 8.8 Hz, 1H), 7.89 (t, *J* = 8.4 Hz, 2H), 7.84 (s, 1H), 7.74 - 7.70 (m, 2H), 7.51 (t, *J* = 7.6 Hz, 1H), 7.40 (d, *J* = 8.4 Hz, 1H), 6.86 (d, *J* = 4.4 Hz, 1H), 6.72 (d, *J* = 4.4 Hz, 1H), 5.17 (d, *J* = 6.0 Hz, 1H), 5.08 (t, *J* = 5.6 Hz, 2H), 4.39 (q, *J* = 5.6 Hz, 1H),4.07 (q, *J* = 4.8 Hz, 1H), 4.01 (q, *J* = 4.8 Hz, 1H), 3.70 (dd, *J* = 13.6, 6.0 Hz, 1H), 3.55 (dd, *J* = 13.6, 6.4 Hz, 1H).

##### 1.7.13 Synthesis of compounds 39 and 40

**Scheme S17**

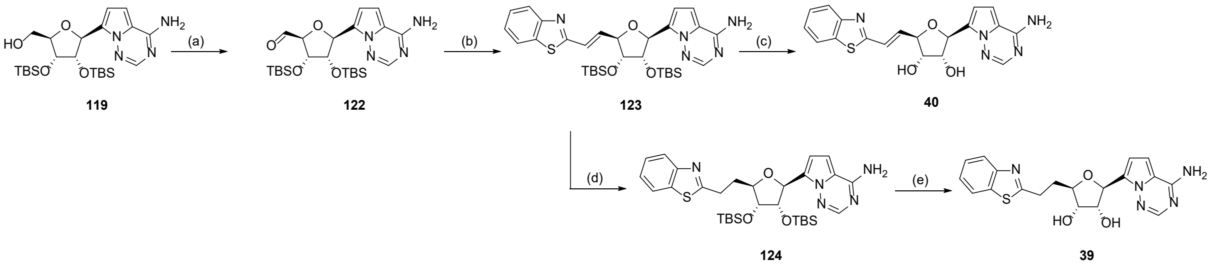

(a) IBX, MeCN, 80 °C, 2 h, 80%; (b) (benzo[d]thiazol-2-ylmethyl)triphenylphosphonium chloride, KHMDS, THF, -60 °C, 3 h, 79%; (c) KF, DMF, 70 °C, 16 h, 32%; (e) $H_2$, Pd/C, THF, RT, 2 h, 95%; (f) KF, DMF, 70 °C, 16 h, 56%

**(2S,3R,4S,5S)-5-(4-Aminopyrrolo[2,1-f][1,2,4]triazin-7-yl)-3,4-bis [[tert-butyl(dimethyl)silyl]oxy]tetrahydrofuran-2-carbaldehyde (122)**

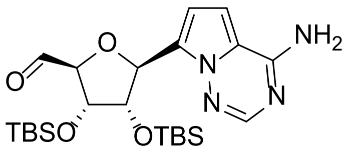

To a solution of compound **119** (300 mg, 0.61 mmol) in MeCN (10 mL) was added 1-hydroxy-1,2-benziodoxol-3(1*H*)-one 1-oxide (339 mg, 1.21 mmol) and stirred at 80 °C for 2 h. The reaction mixture was cooled to RT, quenched with sat. aq. sodium sulfite (10 mL) and extracted with EtOAc (3 x 10 mL). The combined organic layers were washed with brine (3 x 10 mL), dried over sodium sulfate, filtered and the filtrate was concentrated under reduced pressure. The residue was purified by flash column chromatography (petroleum ether/ethyl acetate gradient 0-33%) to afford compound **122** (240 mg, 80% yield) as a yellow solid. LCMS ($ES^+$, Method 2) *m/z* 493.4 $[M+H]^+$; $^1H$ NMR (400 MHz, $CDCl_3$): $\delta$ 9.98 (s, 1H), 7.96 - 7.89 (m, 1H), 7.44 (t, *J* = 7.6 Hz, 1H), 7.15 (t, *J* = 8.0 Hz, 1H), 6.81 - 6.77 (m, 1H), 5.38 (d, *J* = 6.8 Hz, 1H), 4.92 - 4.89 (m, 1H), 4.76 - 4.74 (m, 1H), 1.01 (s, 9H), 0.81 (s, 9H), 0.16 (s, 3H), 0.15 (s, 3H), 0.08 (s, 1H), 0.05 (s, 3H).

**7-[(2*S*,3*S*,4*R*,5*R*)-5-[(*E*)-2-(1,3-Benzothiazol-2-yl)vinyl]-3,4-bis [[tert-butyl(dimethyl) silyl]oxy]tetrahydrofuran-2-yl]pyrrolo[2,1-f][1,2,4]triazin-4-amine (123)**

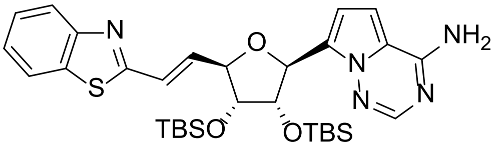

To a solution of (benzo[d]thiazol-2-ylmethyl)triphenylphosphonium chloride (108 mg, 243 µmol) in THF (3 mL) was added dropwise potassium bis(trimethylsilyl) amide (244 µL, 1.0 M in THF) at -60 °C under nitrogen and stirred for 30 min. A solution of compound **122** (80.0 mg, 162.35 µmol) in THF (2 mL) was added and stirred for 1 h. The reaction mixture was quenched with sat. aq. ammonium chloride (20 mL) and extracted with EtOAc (3 x 10 mL). The combined organic layers were washed with brine (2 x 10 mL), dried over sodium sulfate, filtered and the filtrated was concentrated under reduced pressure. The residue was purified by flash column chromatography (petroleum ether/ethyl acetate gradient 0-33%) to afford compound **123** (80.0 mg, 79% yield) as a yellow solid. LCMS ($ES^+$, Method 2) *m/z* 624.5 $[M+H]^+$; $^1H$ NMR (400

MHz, $CDCl_3$): δ 8.02 - 8.00 (m, 3H), 7.98 (s, 1H), 7.87 (d, *J* = 8.0 Hz, 1H), 7.48 (t, *J* = 7.6 Hz, 1H), 7.39 (t, *J* = 7.2 Hz, 1H), 7.17 (dd, *J* = 12.0, 1.2 Hz, 1H), 6.93 (dd, *J* = 12.0, 6.4 Hz, 1H), 6.75 (d, *J* = 4.4 Hz, 1H), 6.67 (d, *J* = 4.4 Hz, 1H), 5.64 (s, 2H), 5.45 (d, *J* = 3.6 Hz, 1H), 4.74 (t, *J* = 6.0 Hz, 1H), 4.59 (t, *J* = 4.4 Hz, 1H), 4.18 - 4.15 (m, 1H), 0.94 (s, 9H), 0.89 (s, 9H), 0.09 (s, 3H), 0.08 (s, 3H), 0.03 (s, 3H), -0.06 (s, 3H).

**(2*S*,3*R*,4*S*,5*R*)-2-(4-aminopyrrolo[2,1-f][1,2,4]triazin-7-yl)-5-[(*E*)-2-(1,3-benzothiazol-2-yl)vinyl]tetrahydrofuran-3,4-diol (40)**

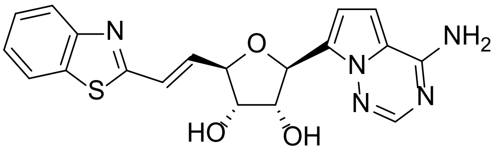

To a solution of compound **123** (50.0 mg, 80.1 µmol) in DMF (1 mL) was added potassium fluoride (46.5 mg, 801 µmol) and stirred at 70 °C for 16 h. The reaction mixture was cooled to RT and then concentrated under reduced pressure. The residue was purified by prep-HPLC (Method 18) to afford compound **40** (10.0 mg, 32% yield) as a yellow solid. HRMS (Method 1) *m/z* $[M+H]^+$ calcd for $C_{19}H_{17}N_5O_3S$ 396.1130; found 396.1139; $^1H$ NMR (400 MHz, DMSO-$d_6$): δ 8.07 (d, *J* = 7.6 Hz, 1H), 7.96 (d, *J* = 8.0 Hz, 1H), 7.87 (s, 1H), 7.74 (brs, 2H), 7.50 (td, *J* = 8.0, 0.8 Hz, 1H), 7.43 (td, *J* = 8.4, 1.2 Hz, 1H), 7.00 - 6.88 (m, 3H), 6.70 (d, *J* = 4.4 Hz, 1H), 5.33 - 5.25 (m, 3H), 4.51 (t, *J* = 5.2 Hz, 1H), 4.32 (q, *J* = 4.8 Hz, 1H), 4.04 (q, *J* = 5.2 Hz, 1H).

**7-((2*S*,3*S*,4*R*,5*R*)-5-(2-(benzo[d]thiazol-2-yl)ethyl)-3,4-bis((tert-butyl dimethylsilyl) oxy)tetrahydrofuran-2-yl)pyrrolo[2,1-f][1,2,4]triazin-4-amine (124)**

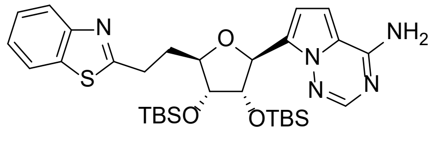

To a solution of compound **123** (100 mg, 160 µmol) in THF (10 mL) was added Pd/C (50.0 mg, 160 µmol, 10% wt.) under inert atmosphere. The suspension was degassed under vacuum and purged with $H_2$ several times. The reaction mixture was stirred under $H_2$ (50 psi) at RT for 2 h, and then filtered. The filtrate was concentrated under reduced pressure. The residue was purified by flash column chromatography (petroleum ether/ethyl acetate gradient 0-33%) to afford compound **124** (95.0 mg, 95% yield) as a yellow oil. LCMS ($ES^+$, Method 2) *m/z* 626.5

[M+H]+; $^1$H NMR (400 MHz, $CDCl_3$): $\delta$ 7.98 (d, $J$ = 8.4 Hz, 1H), 7.93 (s, 1H), 7.85 (d, $J$ = 8.0 Hz, 1H), 7.46 (t, $J$ = 7.6 Hz, 1H), 7.36 (t, $J$ = 7.6Hz, 1H), 6.68 (d, $J$ = 4.4 Hz, 1H), 6.60 (d, $J$ = 4.4 Hz, 1H), 5.70 (s, 2H), 5.32 (d, $J$ = 4.8 Hz, 1H), 4.67 (t, $J$ = 4.4 Hz, 1H), 4.17 - 4.15 (m, 1H), 4.04 (t, $J$ = 4.4 Hz, 1H), 3.46 - 3.39 (m, 1H), 3.28 - 3.21 (m, 1H), 2.34 - 2.27 (m, 2H), 0.93 (s, 9H), 0.83 (s, 9H), 0.12 - 0.09 (m, 6H), -0.06 (s, 3H), -0.21 (s, 3H).

**(2*S*,3*R*,4*S*,5*R*)-2-(4-Aminopyrrolo[2,1-f][1,2,4]triazin-7-yl)-5- [2-(1,3-benzothiazol-2-yl)ethyl]tetrahydrofuran-3,4-diol (39)**

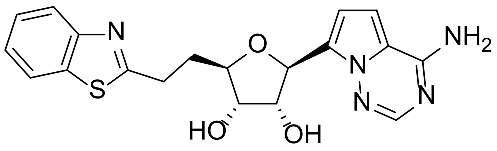

To a solution of compound **124** (90.0 mg, 0.14 mmol) in DMF (2 mL) was added potassium fluoride (83.5 mg, 1.44 mmol) and stirred at 70 °C for 16 h. The reaction mixture was cooled to RT and then concentrated under reduced pressure. The residue was purified by prep-HPLC (Method 19) to afford compound **39** (32.7 mg, 56% yield) as a white solid. HRMS (Method 1) $m/z$ [M+H]+ calcd for $C_{19}H_{19}N_5O_3S$ 398.1287; found 398.1292; $^1$H NMR (400 MHz, DMSO-$d_6$): $\delta$ 8.05 (d, $J$ = 8.0 Hz, 1H), 7.93 (d, $J$ = 8.0 Hz, 1H), 7.84 (s, 1H), 7.70 (brs, 2H), 7.48 (td, $J$ = 8.0, 0.8 Hz, 1H), 7.43 (td, $J$ = 8.4, 1.2 Hz, 1H), 7.40 (dt, $J$ = 8.4, 1.2 Hz, 1H), 6.85 (d, $J$ = 7.6 Hz, 1H), 6.65 (d, $J$ = 4.4 Hz, 1H), 5.14 (d, $J$ = 5.6 Hz, 1H), 5.05 (d, $J$ = 6.0 Hz, 1H), 4.98 (d, $J$ = 5.6 Hz, 1H), 4.26 (q, $J$ = 5.2 Hz, 1H), 3.89 - 3.83 (m, 2H), 3.23 - 3.17 (m, 2H), 2.02 - 2.18 (m, 1H), 2.08 - 2.05 (m, 1H).

### 2 Supplementary Information

#### 2.1 Crystallography table

| | SARS-CoV-2 nsp14 | | | | | |
|---|---|---|---|---|---|---|
| **Compound** | **1** | **5** | **6** | **18** | **26** | **27** |
| **PDB ID** | 9S0M | 9SAJ | 9SAK | 9SAL | 9SAM | 9SAN |
| **Wavelength** | 0.9537 | 0.9537 | 1.0000 | 1.0000 | 1.0000 | 1.0000 |
| **Resolution range** | 63.96 - 2.85 (2.90 – 2.85) | 64.04 - 2.48 (2.52 – 2.48) | 44.84 - 2.30 (2.34 – 2.30) | 41.26 - 2.29 (2.33 – 2.29) | 44.59 - 2.54 (2.58 – 2.54) | 44.79 - 2.73 (2.77 – 2.73) |
| **Space group** | P 1 21 1 | P 1 21 1 | P 1 21 1 | P 1 21 1 | P 1 21 1 | P 1 21 1 |
| **Unit cell** | 67.59 101.86 91.02 90 108.9 90 | 67.81 102.36 90.74 90 109.2 90 | 67.38 102.05 90.87 90 108.7 90 | 67.69 101.63 91.41 90 108.6 90 | 67.58 101.99 91.19 90 109.4 90 | 67.81 102.08 91.25 90 109.3 90 |
| **Total reflections** | 110868 (6040) | 174595 (9276) | 218043 (11092) | 290731 (15520) | 140935 (7106) | 134961 (6843) |
| **Unique reflections** | 26758 (1348) | 40958 (2075) | 50538 (2551) | 51355 (2638) | 36568 (1911) | 31372 (1558) |
| **Multiplicity** | 4.1 (4.5) | 4.3 (4.5) | 4.3 (4.3) | 5.7 (5.9) | 3.9 (3.7) | 4.3 (4.4) |
| **Completeness (%)** | 97.5 (100.0) | 98.6 (100.0) | 97.7 (100.0) | 97.4 (100.0) | 94.5 (99.9) | 99.9 (100.0) |
| **Mean I/sigma(I)** | 12.3 (1.4) | 13.1 (1.3) | 16.8 (1.5) | 16.3 (1.3) | 14.8 (1.3) | 10.4 (1.3) |
| **Wilson B-factor** | 72.26 | 61.48 | 66.24 | 70.21 | 75.80 | 73.51 |
| **R-merge** | 0.089 (1.031) | 0.060 (0.993) | 0.039 (0.825) | 0.044 (0.980) | 0.045 (0.926) | 0.086 (1.105) |
| **R-meas** | 0.102 (1.165) | 0.069 (1.128) | 0.044 (0.941) | 0.048 (1.077) | 0.052 (1.086) | 0.099 (1.257) |
| **CC1/2** | 0.998 (0.556) | 0.999 (0.647) | 0.999 (0.675) | 0.999 (0.653) | 0.999 (0.633) | 0.984 (0.473) |
| **Reflections used in refinement** | 25322 (1342) | 39956 (2064) | 49304 (2540) | 50028 (2609) | 35664 (1905) | 28313 (1556) |
| **Reflections used for R-free** | 625 (64) | 997 (97) | 1224 (135) | 1304 (136) | 884 (94) | 717 (82) |
| **R-work** | 0.2307 | 0.1978 | 0.1984 | 0.2012 | 0.1936 | 0.2122 |
| **R-free** | 0.2817 | 0.2378 | 0.2487 | 0.2281 | 0.2282 | 0.2647 |

| | | | | | | |
|---|---|---|---|---|---|---|
| **Overall number of atoms** | 7111 | 7225 | 7063 | 7012 | 7105 | 6898 |
| **in macromolecules** | 7019 | 6983 | 6806 | 6826 | 6920 | 6773 |
| **in ligands** | 73 | 119 | 115 | 87 | 83 | 72 |
| **in waters** | 19 | 123 | 142 | 99 | 102 | 53 |
| **Protein residues** | 897 | 884 | 876 | 881 | 886 | 882 |
| **RMS (bonds)** | 0.014 | 0.014 | 0.014 | 0.014 | 0.014 | 0.014 |
| **RMS (angles)** | 1.727 | 1.701 | 1.688 | 1.708 | 1.682 | 1.751 |
| **Ramachandran favored (%)** | 95.34 | 97.23 | 97.17 | 97.79 | 97.93 | 94.78 |
| **Ramachandran allowed (%)** | 4.66 | 2.77 | 2.83 | 2.21 | 2.07 | 5.22 |
| **Ramachandran outliers (%)** | 0.00 | 0.00 | 0.00 | 0.00 | 0.00 | 0.00 |
| **Rotamer outliers (%)** | 0.68 | 0.82 | 0.87 | 0.58 | 0.42 | 0.89 |
| **Clashscore** | 1.25 | 0.58 | 0.60 | 0.60 | 0.22 | 1.54 |
| **Average B-factor** | 74.02 | 67.05 | 75.91 | 82.16 | 81.72 | 74.65 |
| **for macromolecules** | 74.13 | 67.23 | 76.18 | 82.42 | 81.96 | 74.84 |
| **for ligands** | 65.02 | 64.78 | 71.95 | 77.27 | 72.83 | 62.25 |
| **for solvent** | 68.27 | 58.88 | 66.36 | 68.43 | 73.00 | 67.14 |

**Table S1.** X-ray crystallography data collection and refinement statistics.

#### 2.2 ADME Summary Tables

Table S2

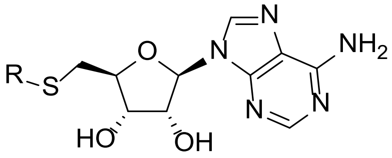

| Compound | R | nsp14 $IC_{50}$ / µM | RNMT $IC_{50}$ / µM | CHI-logD | Mouse liver Mic Clint | |
|---|---|---|---|---|---|---|
| | | | | | Ml/min/g | $T_{1/2}$ / min |
| **1** | 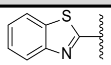 | 0.42 ± 0.3 | > 100 | 1.2 | 1.0 | > 30 |
| **2** | 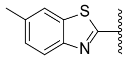 | 0.26 ± 0.07 | > 100 | 1.6 | 1.4 | > 30 |
| **3** | 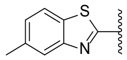 | 1.2 ± 0.15 | > 100 | 1.6 | 2.6 | 25.7 |
| **4** | 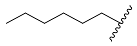 | 6.1 ± 1.1 | > 100 | 2.0 | 5.0 | 13.2 |
| **5** | 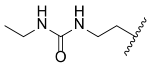 | 1.9 ± 0.30 | > 100 | -0.2 | < 0.5 | > 30 |
| **6** | 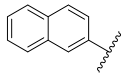 | 0.31 ± 0.12 | > 100 | 1.8 | 2.5 | 26.9 |
| **7** | 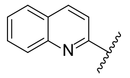 | 0.33 ± 0.09 | > 100 | 1.3 | 2.0 | > 30 |
| **8** | 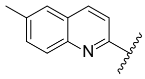 | 1.1 ± 0.07 | > 100 | 1.7 | 1.4 | > 30 |
| **9** | 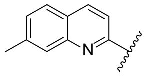 | 0.39 ± 0.06 | 70.5 | 1.7 | 2.2 | > 30 |
| **10** | 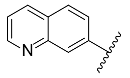 | 0.36 ± 0.11 | > 100 | 0.6 | 0.7 | > 30 |
| **11** | 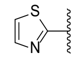 | > 100 | > 100 | 0.3 | < 0.5 | > 30 |
| **12** | 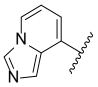 | 15 ± 12 | 9.3 | 0.3 | - | - |

| | | | | | | |
|---|---|---|---|---|---|---|
| **13** | 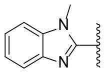 | 0.82 ± 0.13 | > 100 | 0.8 | 0.7 | > 30 |
| **14** | 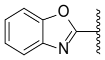 | 0.67 ± 0.02 | > 100 | 1.0 | 0.5 | > 30 |
| **15** | 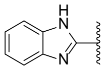 | 0.59 ± 0.11 | > 100 | 0.3 | < 0.5 | > 30 |
| **16** | 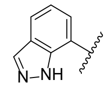 | *79 ± 18 | > 100 | 0.7 | - | - |
| **17** | 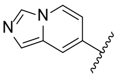 | 6.9 ± 4.6 | 31.1 | 0.1 | 6.0 | 11.1 |

Table S3

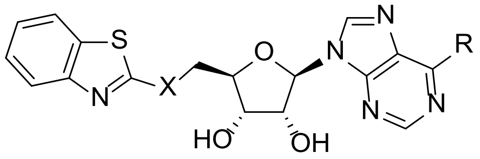

| | **X** | **Y** | **R** | **nsp14 IC$_{50}$ / µM** | **RNMT IC$_{50}$ / µM** | **CHI-logD** | **Mouse liver T$_{1/2}$ / min** |
|---|---|---|---|---|---|---|---|
| **1** | O | N | - | 0.420 ± 0.3 | > 100 | 1.2 | > 30 |
| **26** | $CH_2$ | N | - | 0.053 ± 0.03 | > 100 | 1.5 | > 30 |
| **27** | O | C | H | 0.055 ± 0.02 | > 100 | 1.6 | > 30 |
| **28** | $CH_2$ | C | H | 0.308 ± 0.34 | 22.8 | 1.8 | > 30 |
| **29** | O | C | CN | 0.170 ± 0.02 | > 100 | 1.8 | 12.1 |
| **30** | O | C | C≡CH | 0.054± 0.01 | > 100 | 2.0 | 10.1 |
| **31** | O | C | Et | 0.250 ± 0.2 | > 100 | 2.1 | 3.9 |

Table S4

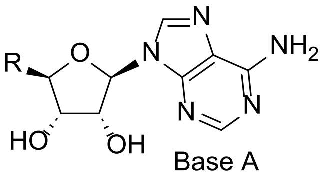

**25**, **32**-**33**

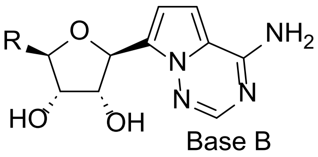

**34**-**40**

| **#** | **R** | **Base** | **CHI-logD** | **Mouse liver Mic Clint** | |
|---|---|---|---|---|---|
| | | | | **mL/min/g** | **$T_{1/2}$ / min** |
| **25** | 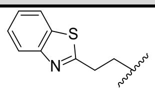 | A | 1.1 | 1.0 | > 30 |
| **32** | 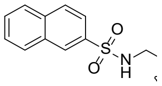 | A | 1.5 | 1.0 | > 30 |
| **33** | 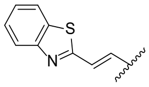 | A | 1.3 | < 0.5 | > 30 |
| **34** | 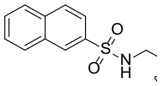 | B | 1.6 | 6.7 | 9.9 |
| **35** | 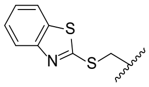 | B | 1.5 | 1.1 | > 30 |
| **36** | 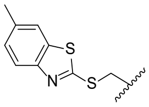 | B | 1.9 | 3.7 | 18 |
| **37** | 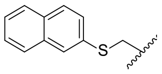 | B | 2.2 | 4.9 | 13.6 |
| **38** | 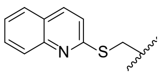 | B | 1.7 | 1.2 | > 30 |
| **39** | 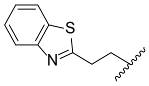 | B | 1.3 | 1.1 | > 30 |
| **40** | 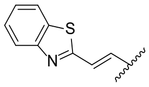 | B | 1.5 | 0.8 | > 30 |

#### 2.3 Repeats in biochemical assays

Table S5 – repeat 'n' values for different compounds in stated biochemical assays

| Compound | SARS-CoV-2 nsp14 | NL63 nsp14 | 229E nsp14 | RNMT |
|---|---|---|---|---|
| **1** | 11 | 2 | 2 | 2 |
| **2** | 4 | 2 | 2 | 2 |
| **3** | 2 | - | - | 2 |
| **4** | 4 | 2 | 2 | 2 |
| **5** | 3 | - | - | 2 |
| **6** | 12 | 2 | 2 | 2 |
| **7** | 11 | 2 | 2 | 2 |
| **8** | 4 | 2 | 2 | 2 |
| **9** | 4 | 2 | 2 | 2 |
| **10** | 8 | 2 | 2 | 2 |
| **11** | 4 | 2 | 2 | 2 |
| **12** | 9 | 2 | 2 | 2 |
| **13** | 5 | 2 | 2 | 2 |
| **14** | 5 | 2 | 2 | 2 |
| **15** | 9 | 2 | 2 | 2 |
| **16** | 3 | - | - | 2 |
| **17** | 4 | 2 | 2 | 2 |
| **18** | 2 | - | - | - |
| **19** | 6 | 2 | 2 | 2 |
| **20** | 7 | 2 | 2 | 2 |
| **21** | 7 | 2 | 2 | 2 |
| **22** | 3 | - | - | 2 |
| **23** | 2 | - | - | 2 |
| **24** | 2 | - | - | 2 |
| **25** | 10 | 2 | 2 | 2 |
| **26** | 18 | 2 | 2 | 2 |
| **27** | 9 | 2 | 2 | 2 |
| **28** | 4 | 2 | 2 | 2 |
| **29** | 4 | 2 | 2 | 2 |
| **30** | 8 | 2 | 2 | 2 |
| **31** | 4 | 2 | 2 | 2 |
| **32** | 10 | 2 | 2 | 2 |
| **33** | 6 | 2 | 2 | 2 |
| **34** | 10 | 2 | 2 | 2 |
| **35** | 8 | 2 | 2 | 2 |
| **36** | 2 | - | - | 2 |
| **37** | 2 | - | - | 2 |
| **38** | 2 | - | - | 2 |
| **39** | 8 | 2 | 2 | 2 |
| **40** | 8 | 2 | 2 | 2 |

#### 2.4 MERS AlphaFold2 model

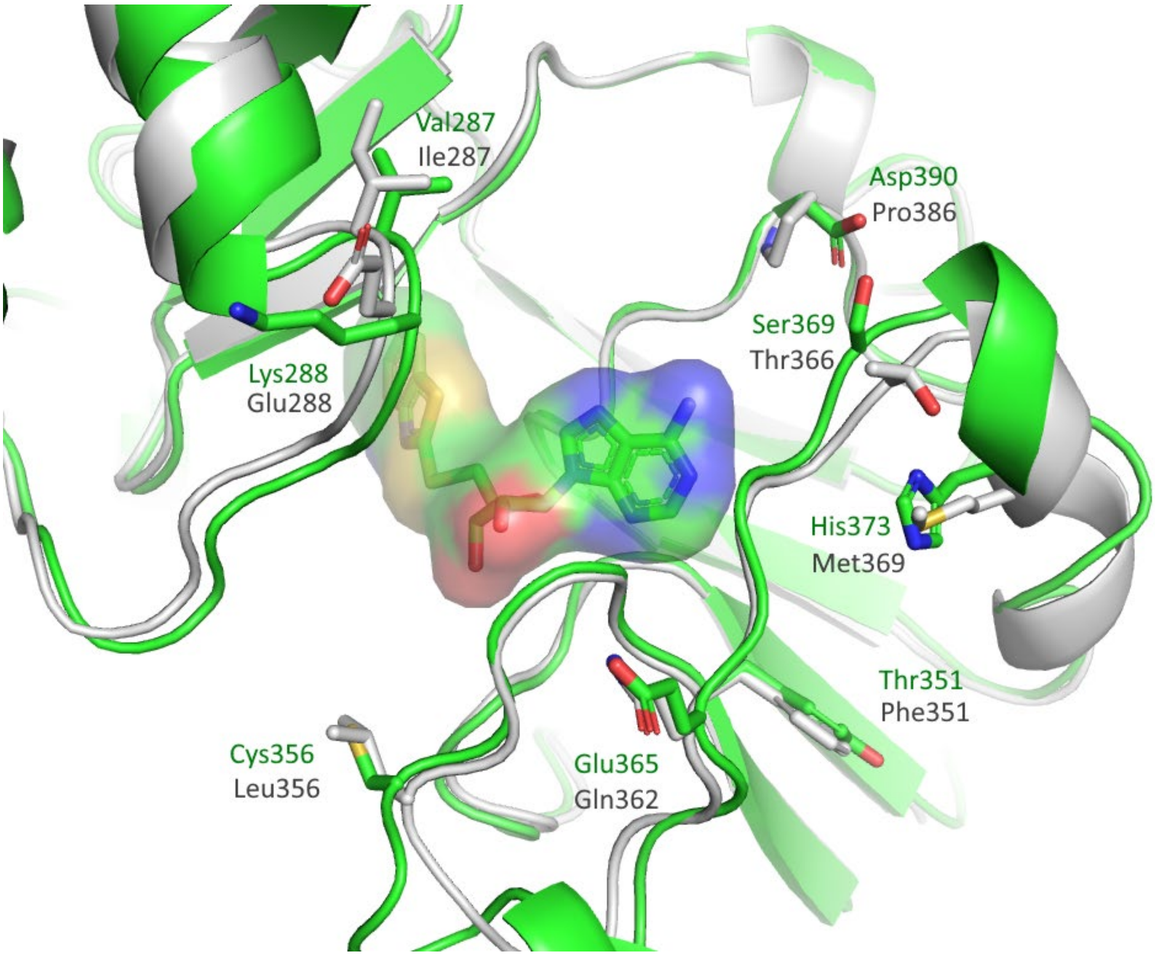

**Figure S1** Compound **1** bound to SARS-CoV-2 nsp14 (9s0m, green carbons) superimposed onto a MERS nsp14 AlphaFold2 model (grey carbons). Residues that differ between SARS-CoV-2 nsp14 (green) and MERS nsp14 (grey) in the proximity of the ligand **1** binding site are indicated.

### 3 Appendix

#### 3.1 QC Spectra for Compounds

$^{1}$**H NMR** 9-[(2*R*,3*R*,4*R*,5*R*)-3,4-Bis[[tert-butyl(dimethyl)silyl]oxy]-5-[[tert-butyl(dimethyl)silyl]oxymethyl] tetrahydrofuran-2-yl]purin-6-amine **(42)**

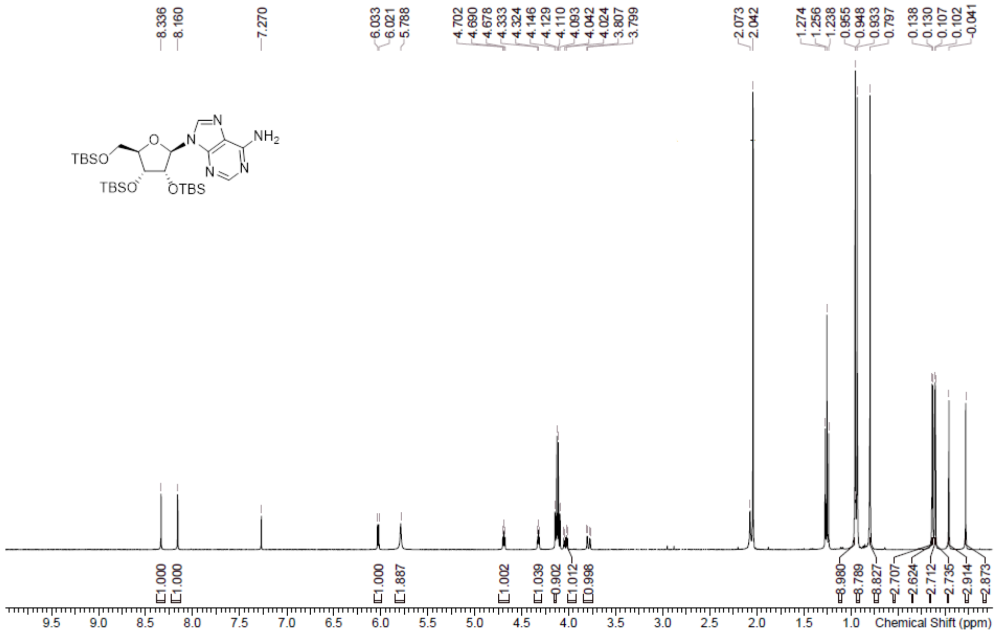

**LCMS** 9-[(2*R*,3*R*,4*R*,5*R*)-3,4-Bis[[tert-butyl(dimethyl)silyl]oxy]-5-[[tert-butyl(dimethyl)silyl]oxymethyl] tetrahydrofuran-2-yl]purin-6-amine **(42)**

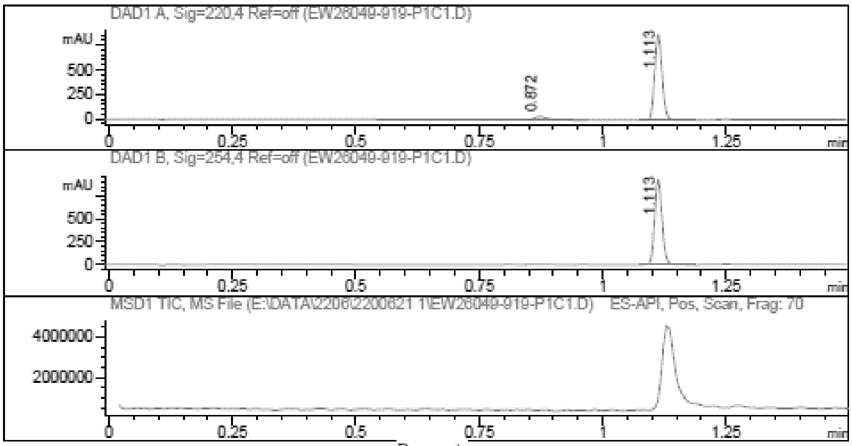

Report

```
=================================================================
   Signal 1 : DAD1 A, Sig=220,4 Ref=off
    #  Meas. Ret.   Height    Width       Area       Area %
   --------------------------------------------------------
     1      0.872      31.677   0.030          66.044   6.676
     2      1.113     871.994   0.016         923.297  93.324
   --------------------------------------------------------

   Signal 2 : DAD1 B, Sig=254,4 Ref=off
    #  Meas. Ret.   Height    Width       Area       Area %
   --------------------------------------------------------
     1      1.113     928.441   0.016         978.312 100.000
   --------------------------------------------------------
```

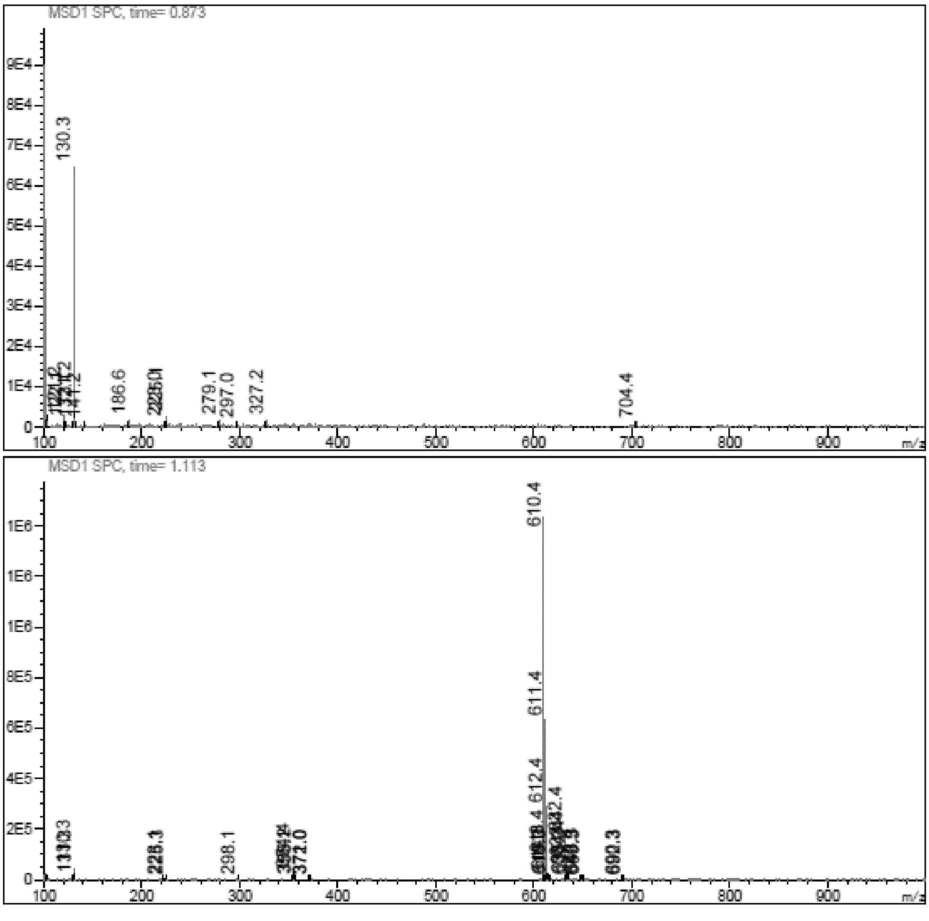

**$^{1}$H NMR** [(2*R*,3*R*,4*R*,5*R*)-5-(6-Aminopurin-9-yl)-3,4-bis[[tert-butyl(dimethyl)silyl]oxy] tetrahydrofuran-2yl] methanol **(43)**

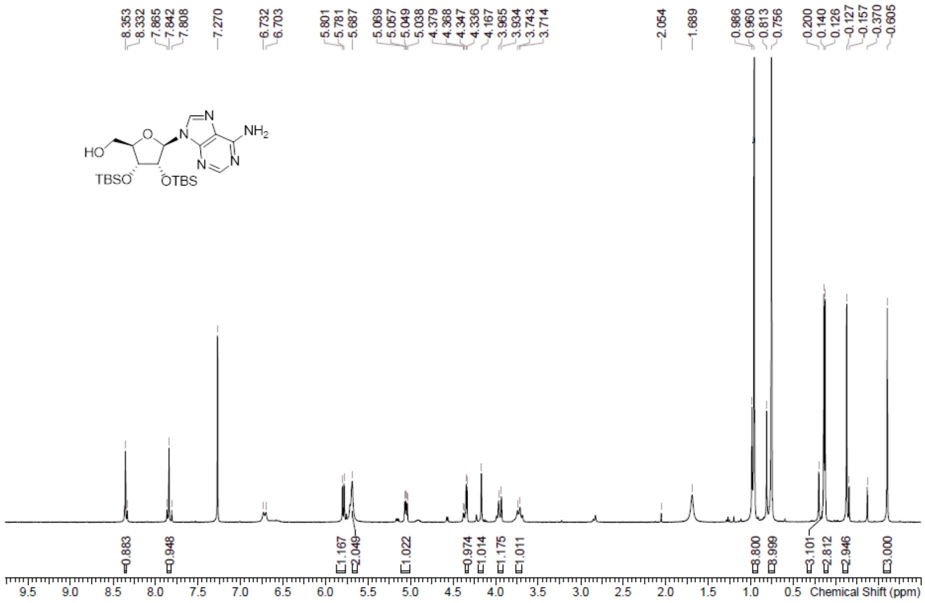

**LCMS** [(2*R*,3*R*,4*R*,5*R*)-5-(6-Aminopurin-9-yl)-3,4-bis[[tert-butyl(dimethyl)silyl]oxy] tetrahydrofuran-2yl]methanol **(43)**

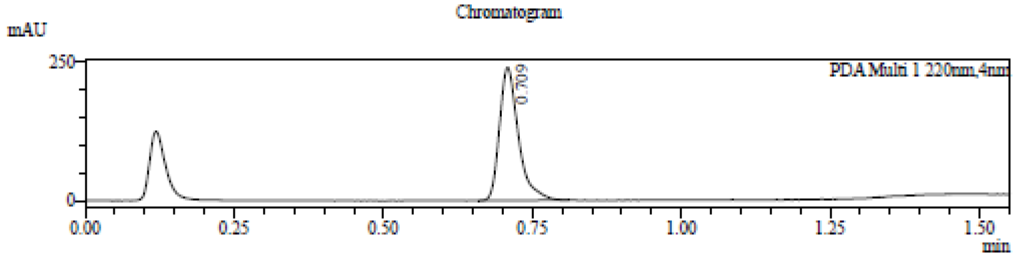

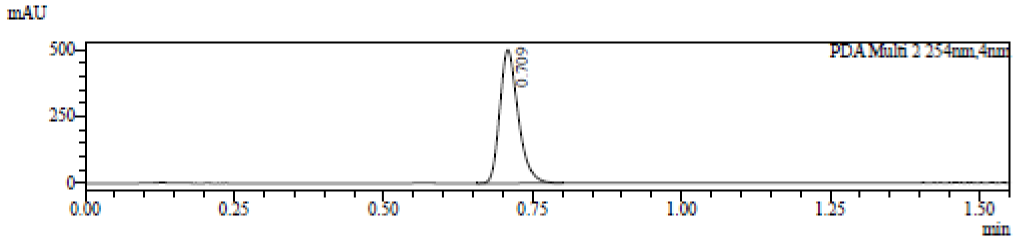

1 PDA Multi 1 / 220nm,4nm
2 PDA Multi 2 / 254nm,4nm

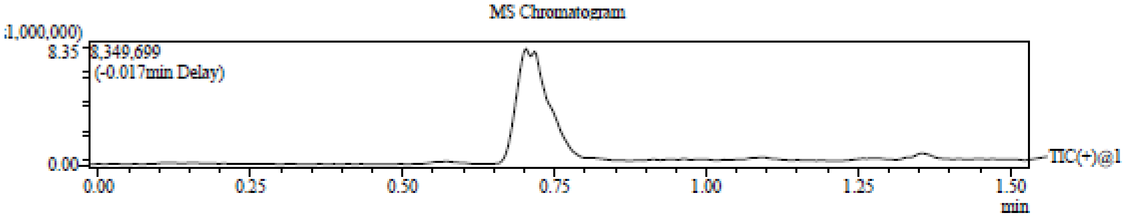

Integration Result

Peak Table

PDA Ch1 220nm

| Peak# | Ret. Time | Height | Height% | USP Width | Area | Area% |
|---|---|---|---|---|---|---|
| 1 | 0.709 | 239309 | 100.000 | 0.057 | 531891 | 100.000 |

Peak Table

PDA Ch2 254nm

| Peak# | Ret. Time | Height | Height% | USP Width | Area | Area% |
|---|---|---|---|---|---|---|
| 1 | 0.709 | 499610 | 100.000 | 0.056 | 1076670 | 100.000 |

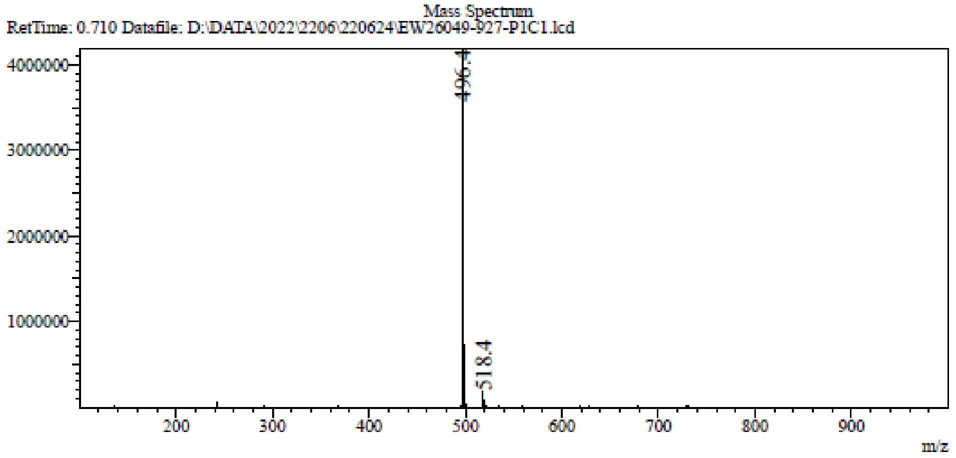

**[1]H NMR** [(2*R*,3*R*,4*R*,5*R*)-5-(6-Aminopurin-9-yl)-3,4-bis[[tert-butyl(dimethyl)silyl] oxy] tetrahydrofuran-2-yl]methyl methanesulfonate **(44)**

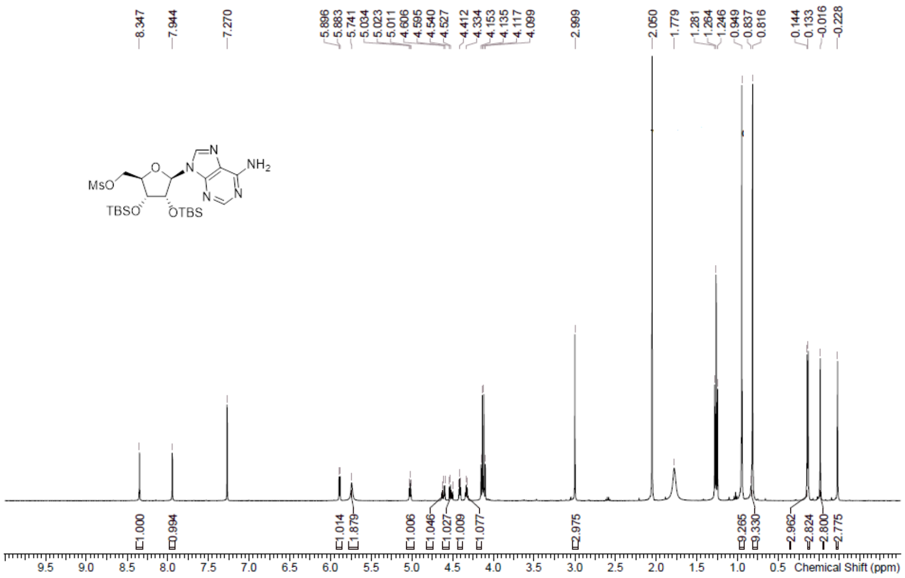

**LCMS** [(2*R*,3*R*,4*R*,5*R*)-5-(6-Aminopurin-9-yl)-3,4-bis[[tert-butyl(dimethyl)silyl] oxy] tetrahydrofuran-2-yl]methyl methanesulfonate **(44)**

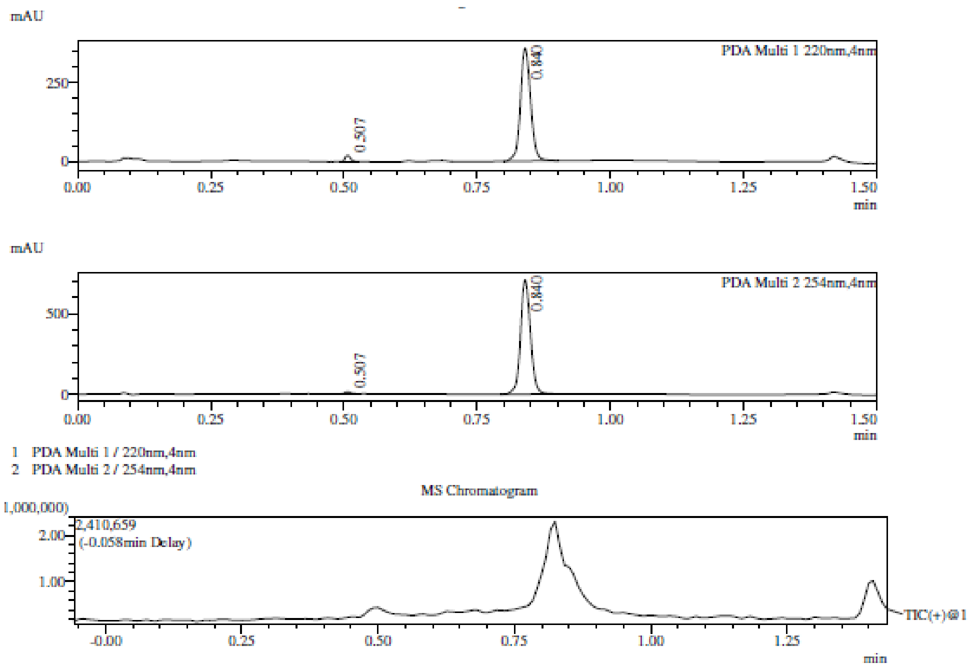

Integration Result

Peak Table

PDA Ch1 220nm

| Peak# | Ret. Time | Height | Height% | USP Width | Area | Area% |
|---|---|---|---|---|---|---|
| 1 | 0.507 | 19707 | 5.230 | 0.022 | 14520 | 2.952 |
| 2 | 0.840 | 357123 | 94.770 | 0.036 | 477346 | 97.048 |

Peak Table

PDA Ch2 254nm

| Peak# | Ret. Time | Height | Height% | USP Width | Area | Area% |
|---|---|---|---|---|---|---|
| 1 | 0.507 | 13212 | 1.825 | 0.022 | 9833 | 1.046 |
| 2 | 0.840 | 710668 | 98.175 | 0.035 | 930380 | 98.954 |

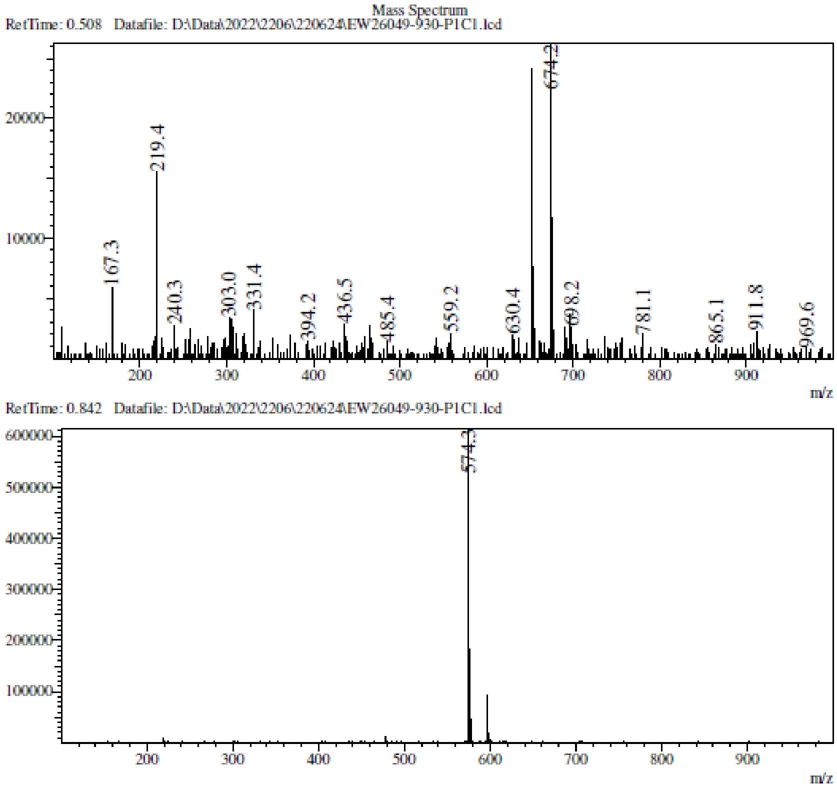

**LCMS** 9-[(2*R*,3*R*,4*R*,5*S*)-5-(1,3-Benzothiazol-2-ylsulfanylmethyl)-3,4-bis[[tert-butyl (dimethyl)silyl]oxy] tetrahydrofuran-2-yl]purin-6-amine **(45a)**

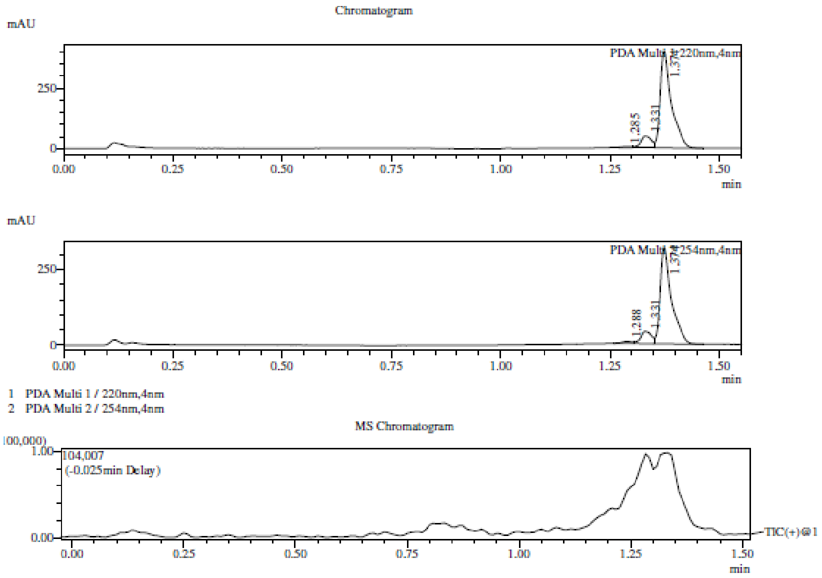

Integration Result

Peak Table

PDA Ch1 220nm

| Peak# | Ret. Time | Height | Height% | USP Width | Area | Area% |
|---|---|---|---|---|---|---|
| 1 | 1.285 | 4147 | 0.911 | 0.405 | 6479 | 0.800 |
| 2 | 1.331 | 49389 | 10.849 | 0.047 | 81089 | 10.013 |
| 3 | 1.374 | 401722 | 88.241 | 0.043 | 722269 | 89.187 |

Peak Table

PDA Ch2 254nm

| Peak# | Ret. Time | Height | Height% | USP Width | Area | Area% |
|---|---|---|---|---|---|---|
| 1 | 1.288 | 6516 | 1.781 | 0.068 | 10327 | 1.585 |
| 2 | 1.331 | 39941 | 10.919 | 0.048 | 66355 | 10.182 |
| 3 | 1.374 | 319349 | 87.300 | 0.043 | 575034 | 88.234 |

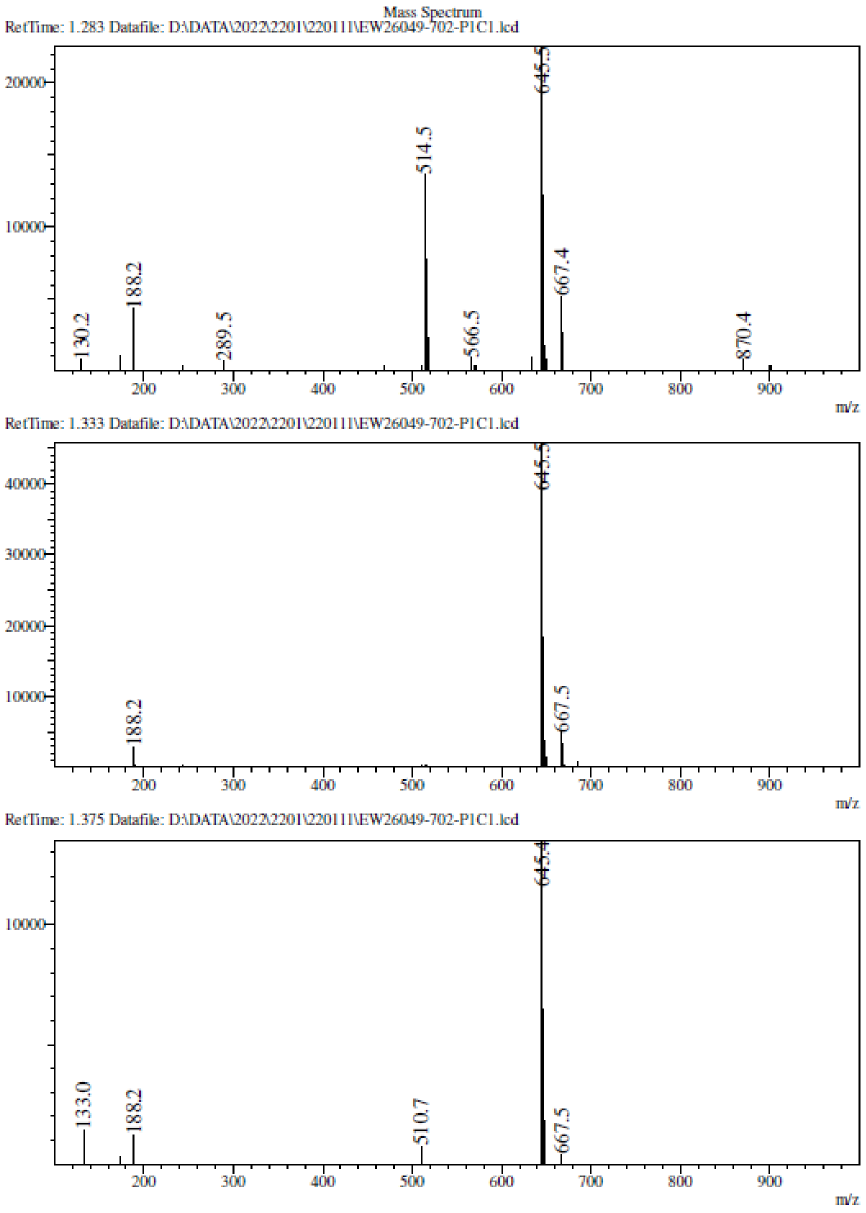
Mass Spectrum
RetTime: 1.283 Datafile: D:\DATA\2022\2201\220111\EW26049-702-P1C1.lcd
20000
10000
130.2
188.2
289.5
514.5
566.5
645.5
667.4
870.4
200
300
400
500
600
700
800
900
m/z
RetTime: 1.333 Datafile: D:\DATA\2022\2201\220111\EW26049-702-P1C1.lcd
40000
30000
20000
10000
188.2
645.5
667.5
200
300
400
500
600
700
800
900
m/z
RetTime: 1.375 Datafile: D:\DATA\2022\2201\220111\EW26049-702-P1C1.lcd
10000
133.0
188.2
510.7
645.4
667.5
200
300
400
500
600
700
800
900
m/z

**$^{1}$H NMR** 9-[(2*R*,3*R*,4*R*,5*S*)-3,4-Bis[[tert-butyl(dimethyl)silyl]oxy]-5-[(6- methyl-1,3-benzothiazol-2-yl)sulfanylmethyl] tetrahydrofuran-2-yl]purin-6-amine **(45b)**

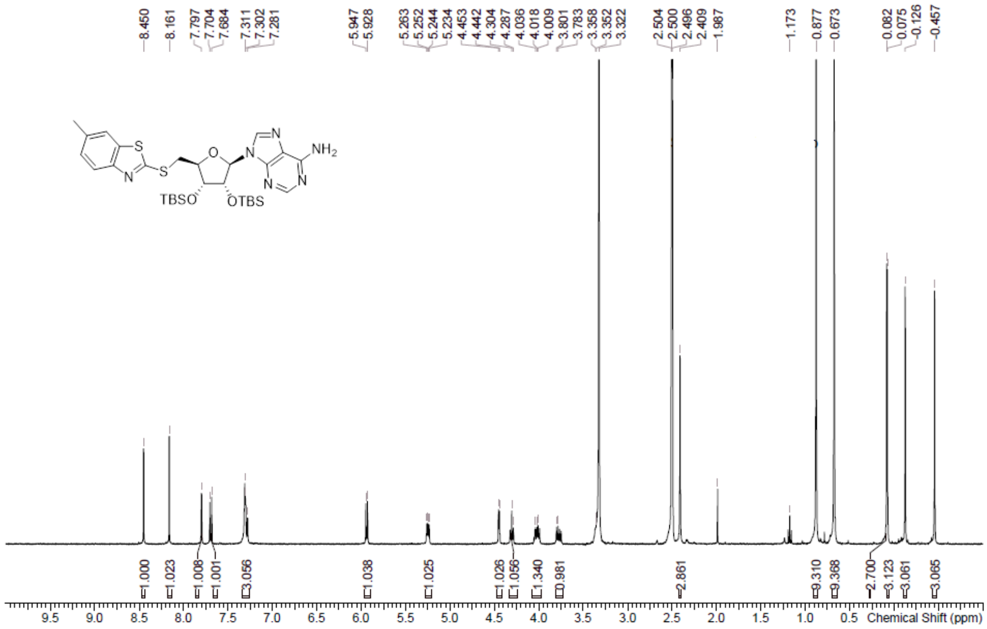

**LCMS** 9-[(2*R*,3*R*,4*R*,5*S*)-3,4-Bis[[tert-butyl(dimethyl)silyl]oxy]-5-[(6- methyl-1,3-benzothiazol-2-yl)sulfanylmethyl] tetrahydrofuran-2-yl]purin-6-amine **(45b)**

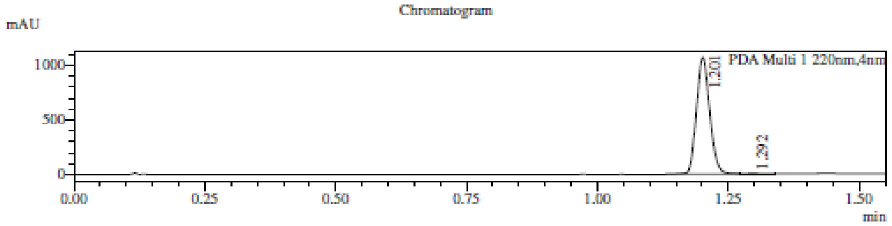

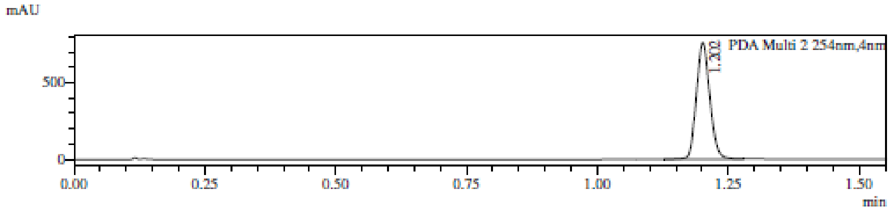

1 PDA Multi 1 / 220nm,4nm
2 PDA Multi 2 / 254nm,4nm

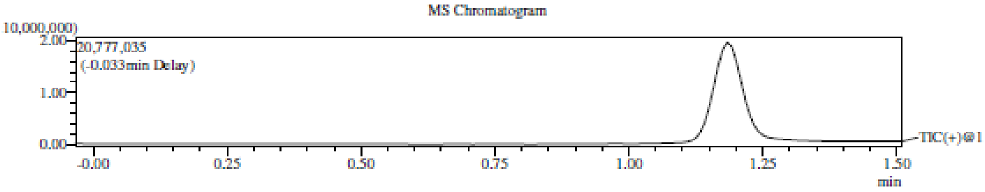

Integration Result

Peak Table

PDA Ch1 220nm

| Peak# | Ret. Time | Height | Height% | USP Width | Area | Area% |
|---|---|---|---|---|---|---|
| 1 | 1.201 | 1071362 | 99.630 | 0.049 | 1905861 | 99.448 |
| 2 | 1.292 | 3975 | 0.370 | 0.080 | 10573 | 0.552 |

Peak Table

PDA Ch2 254nm

| Peak# | Ret. Time | Height | Height% | USP Width | Area | Area% |
|---|---|---|---|---|---|---|
| 1 | 1.202 | 760491 | 100.000 | 0.049 | 1363581 | 100.000 |

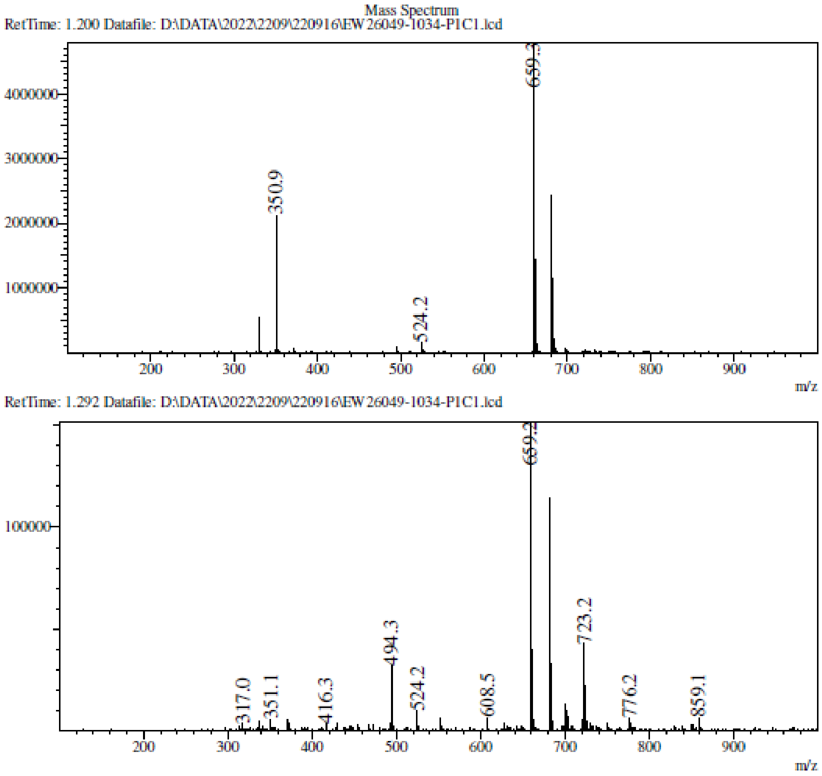

**LCMS** 9-[(2*R*,3*R*,4*R*,5*S*)-3,4-bis[[tert-butyl(dimethyl)silyl]oxy]-5- [(5-methyl-1,3-benzothiazol-2yl)sulfanylmethyl] tetrahydrofuran-2-yl]purin-6-amine **(45c)**

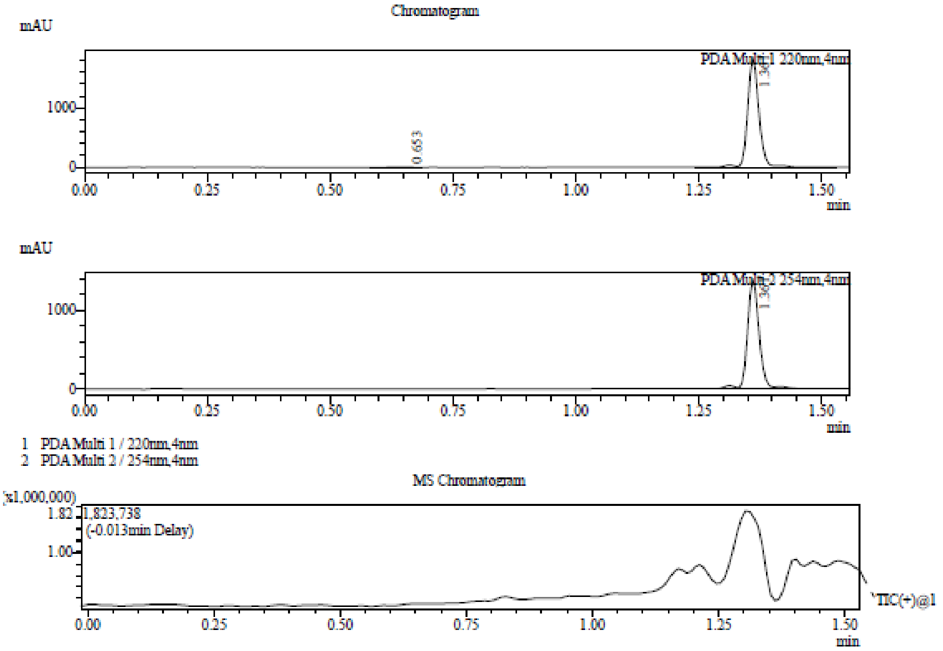

Integration Result

Peak Table

PDA Ch1 220nm

| Peak# | Ret. Time | Height | Height% | USP Width | Area | Area% |
|---|---|---|---|---|---|---|
| 1 | 0.653 | 2093 | 0.113 | 0.096 | 5952 | 0.203 |
| 2 | 1.361 | 1853171 | 99.887 | 0.048 | 2928174 | 99.797 |

Peak Table

PDA Ch2 254nm

| Peak# | Ret. Time | Height | Height% | USP Width | Area | Area% |
|---|---|---|---|---|---|---|
| 1 | 1.361 | 1401706 | 100.000 | 0.049 | 2307954 | 100.000 |

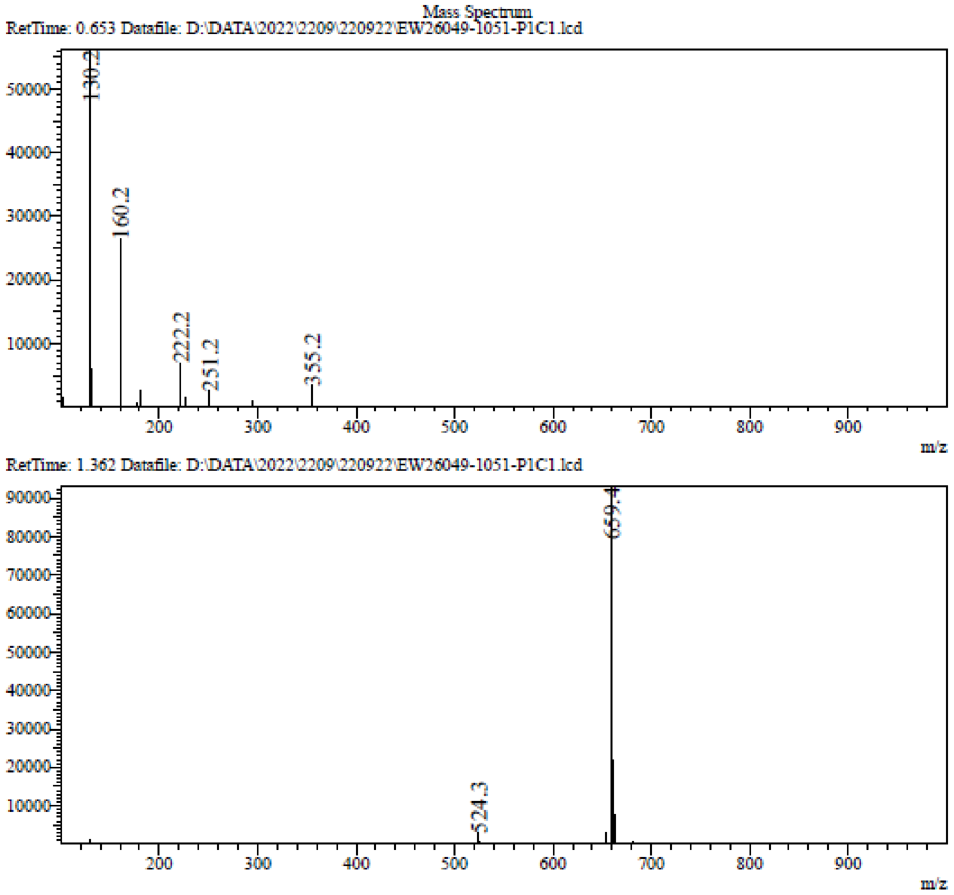

**¹H NMR** 9-[(2*R*,3*R*,4*R*,5*S*)-3,4-bis[[tert-butyl(dimethyl)silyl]oxy]-5-(2-quinolylsulfanylmethyl) tetrahydrofuran-2-yl]purin-6-amine **(45d)**

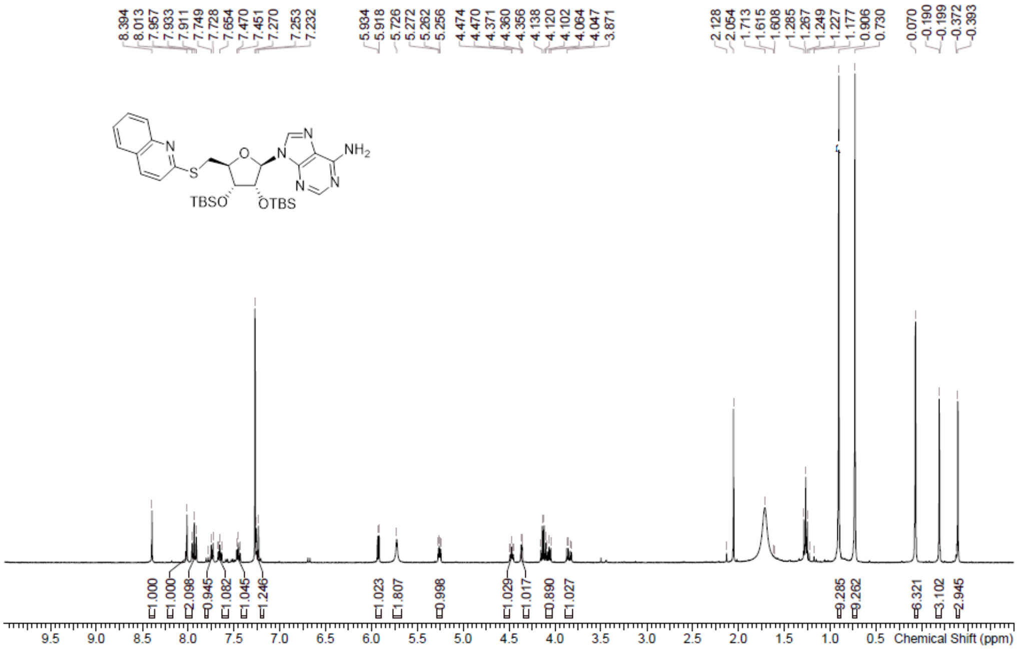

**LCMS** 9-[(2R,3R,4R,5S)-3,4-Bis[[tert-butyl(dimethyl)silyl]oxy]-5- [(6-methyl-2-quinolyl) sulfanylmethyl]

tetrahydrofuran-2-yl]purin-6-amine **(45e)**

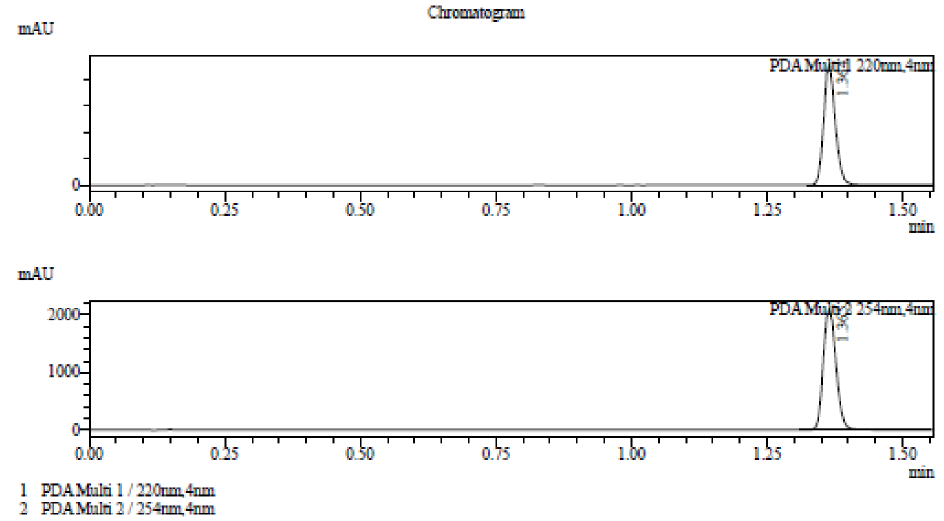

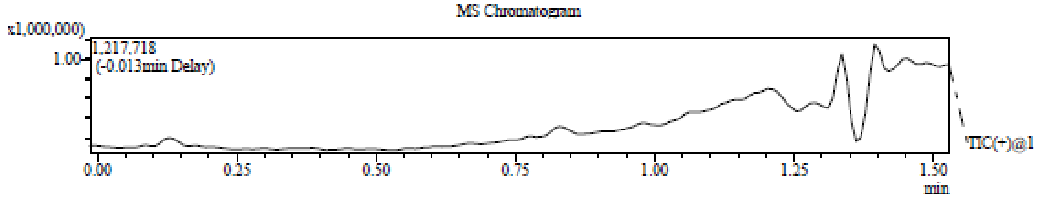

Integration Result

Peak Table

PDA Ch1 220nm

| Peak# | Ret. Time | Height | Height% | USP Width | Area | Area% |
|---|---|---|---|---|---|---|
| 1 | 1.364 | 2289479 | 100.000 | 0.045 | 3372238 | 100.000 |

Peak Table

PDA Ch2 254nm

| Peak# | Ret. Time | Height | Height% | USP Width | Area | Area% |
|---|---|---|---|---|---|---|
| 1 | 1.365 | 2112825 | 100.000 | 0.050 | 3448768 | 100.000 |

Mass Spectrum

RetTime: 1.362 Datafile: D:\DATA\2022\2209\220922\EW26049-1050-P1C1.lcd

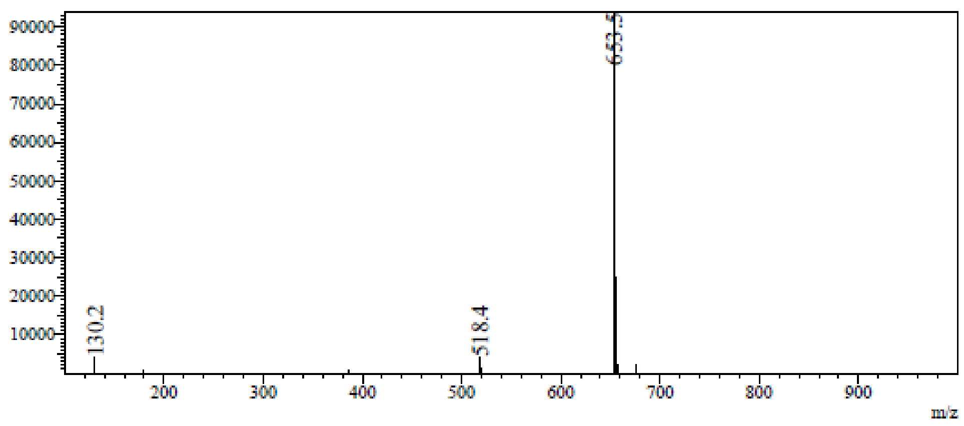

**LCMS** 9-[(2*R*,3*R*,4*R*,5*S*)-3,4-Bis[[tert-butyl(dimethyl)silyl]oxy]-5-[(7 -methyl-2-quinolyl)sulfanyl methyl]tetrahydrofuran-2-yl]purin-6-amine **(45f)**

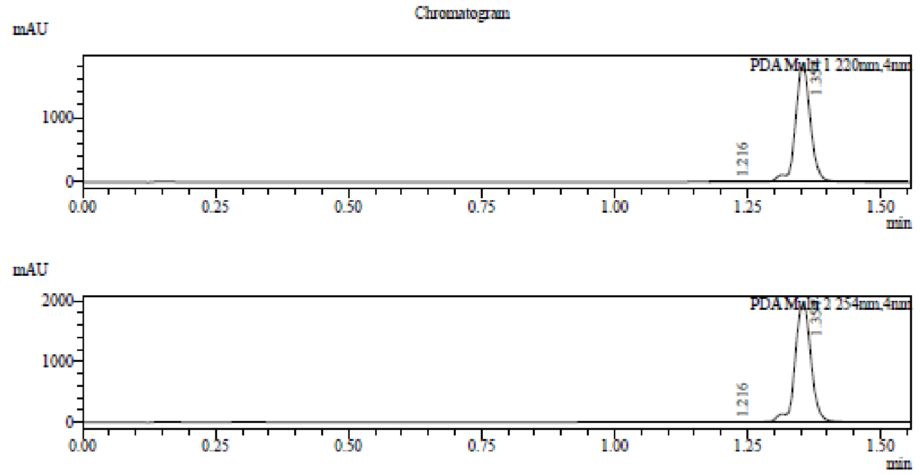

1 PDA Multi 1 / 220nm,4nm
2 PDA Multi 2 / 254nm,4nm

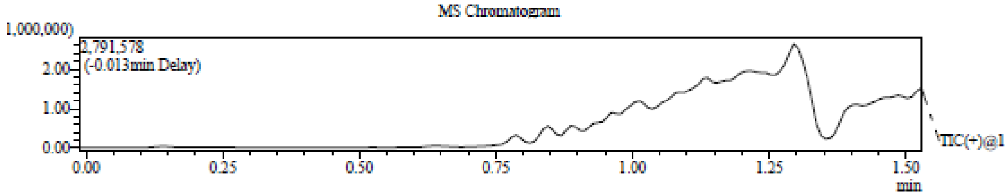

Integration Result

Peak Table

PDA Ch1 220nm

| Peak# | Ret. Time | Height | Height% | USP Width | Area | Area% |
|---|---|---|---|---|---|---|
| 1 | 1.216 | 4330 | 0.234 | 0.087 | 11400 | 0.318 |
| 2 | 1.354 | 1843472 | 99.766 | 0.058 | 3568278 | 99.682 |

Peak Table

PDA Ch2 254nm

| Peak# | Ret. Time | Height | Height% | USP Width | Area | Area% |
|---|---|---|---|---|---|---|
| 1 | 1.216 | 2462 | 0.125 | 0.165 | 8051 | 0.196 |
| 2 | 1.354 | 1964935 | 99.875 | 0.061 | 4098328 | 99.804 |

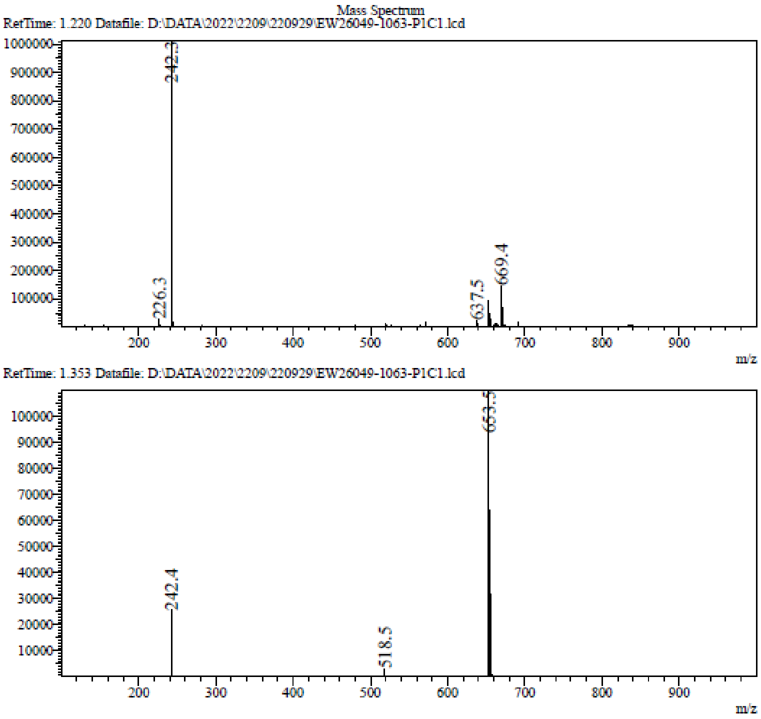

**[1]H NMR** 9-((2R,3R,4R,5S)-3,4-Bis((tert-butyldimethylsilyl)oxy)-5-((quinolin-7-ylthio)methyl) tetrahydrofuran-2-yl)-9H-purin-6-amine **(45g)**

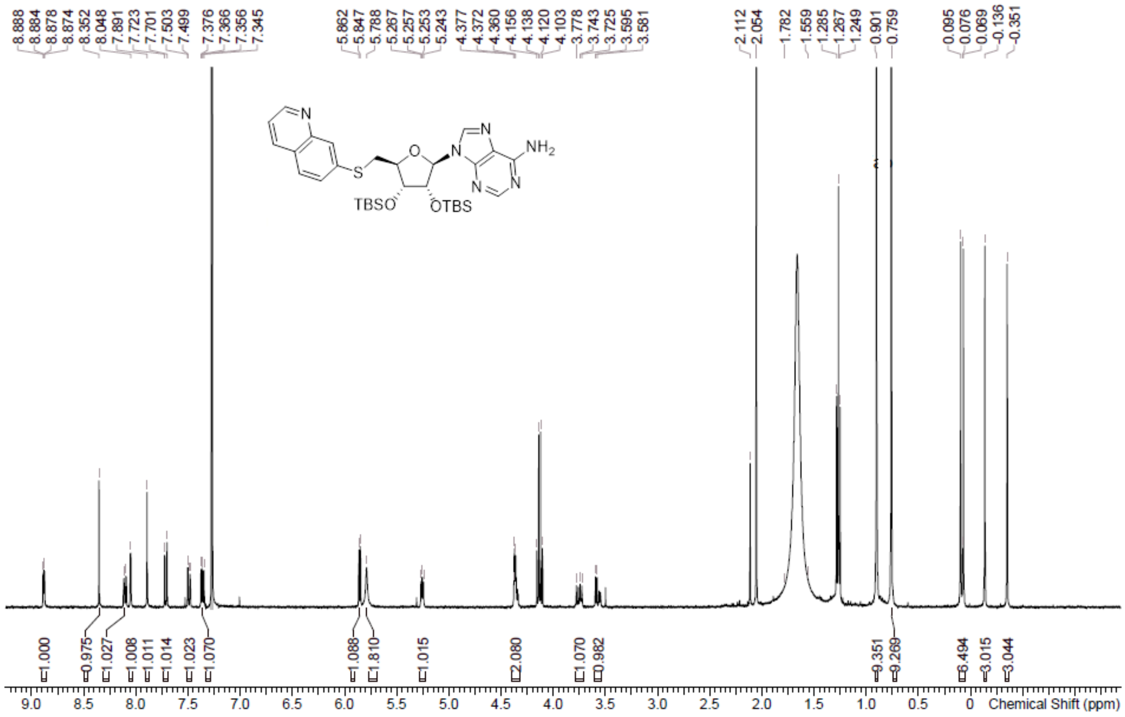

**LCMS** 9-((2R,3R,4R,5S)-3,4-Bis((tert-butyldimethylsilyl)oxy)-5-((quinolin-7-ylthio)methyl) tetrahydrofuran-2-yl)-9H-purin-6-amine **(45g)**

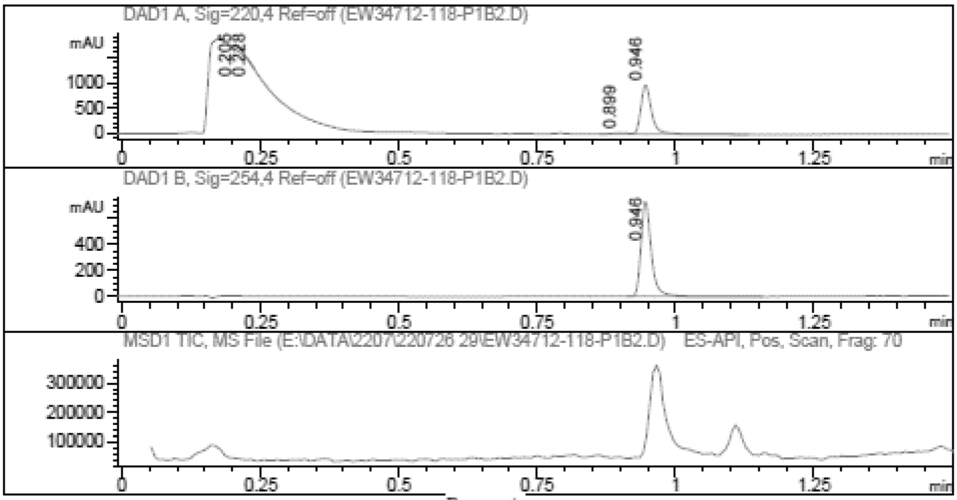

Report

Signal 1 : DAD1 A, Sig=220,4 Ref=off

| # | Meas. Ret. | Height | Width | Area | Area % |
|---|---|---|---|---|---|
| 1 | 0.205 | 56.491 | 0.020 | 71.408 | 5.091 |
| 2 | 0.228 | 21.645 | 0.007 | 8.642 | 0.616 |
| 3 | 0.899 | 18.134 | 0.028 | 33.545 | 2.392 |
| 4 | 0.946 | 964.954 | 0.021 | 1289.056 | 91.901 |

Signal 2 : DAD1 B, Sig=254,4 Ref=off

| # | Meas. Ret. | Height | Width | Area | Area % |
|---|---|---|---|---|---|
| 1 | 0.946 | 728.104 | 0.020 | 940.752 | 100.000 |

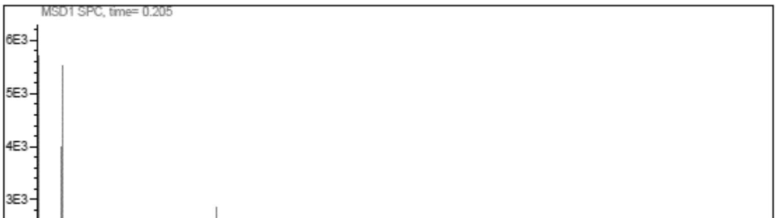

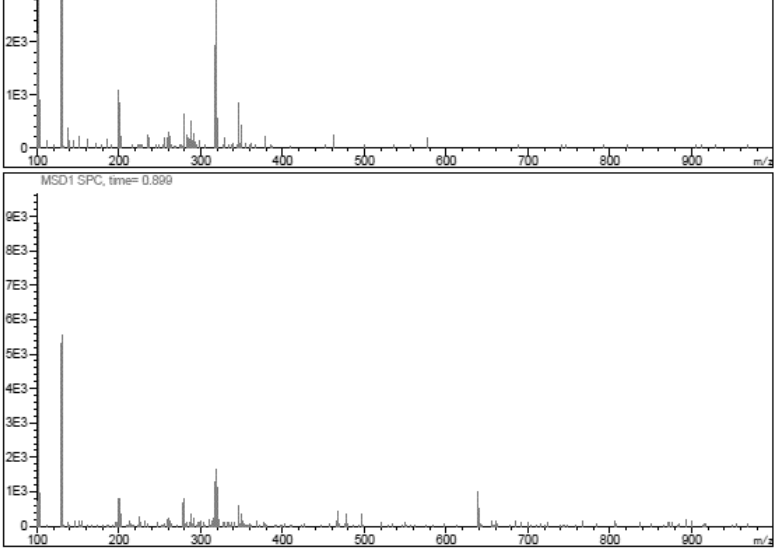

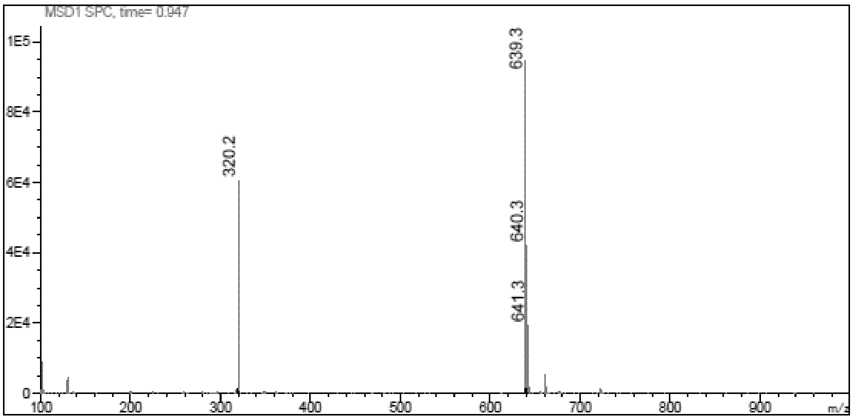

**$^{1}$H NMR** 9-[(2*R*,3*R*,4*R*,5*S*)-3,4-Bis[[tert-butyl(dimethyl)silyl]oxy]-5- (thiazol-2-ylsulfanylmethyl) tetrahydrofuran-2-yl]purin-6-amine **(45h)**

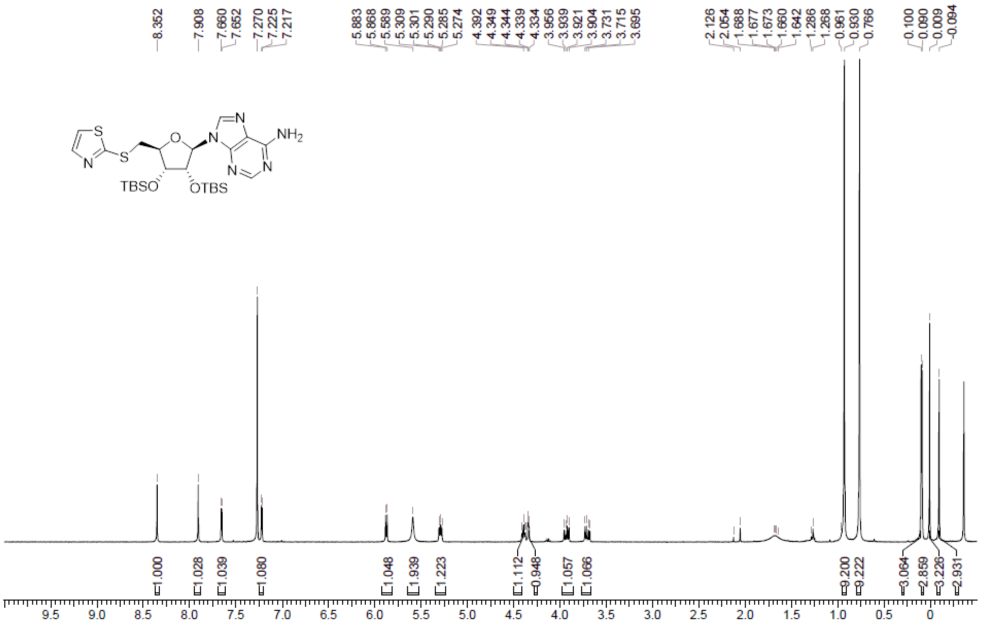

**LCMS** 9-[(2*R*,3*R*,4*R*,5*S*)-3,4-Bis[[tert-butyl(dimethyl)silyl]oxy]-5- (thiazol-2-ylsulfanylmethyl) tetrahydrofuran-2-yl]purin-6-amine **(45h)**

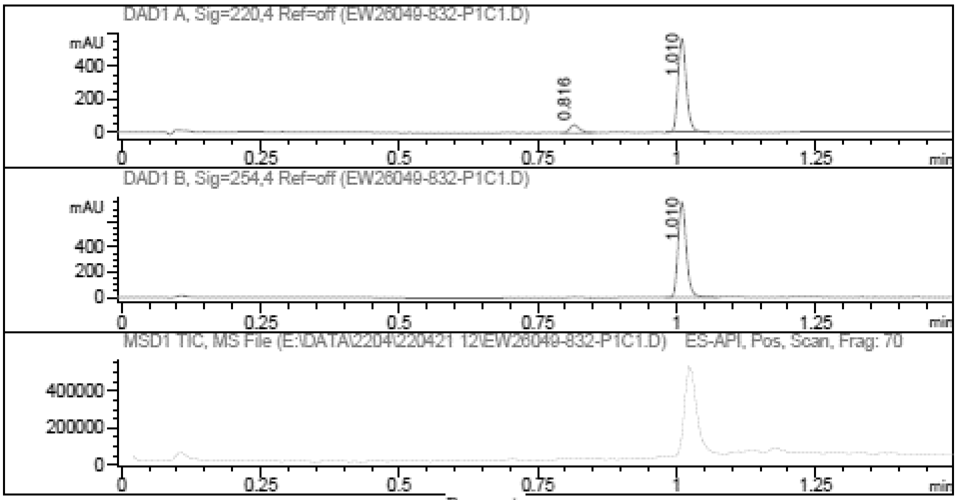

Report

Signal 1 : DAD1 A, Sig=220,4 Ref=off

| # | Meas. Ret. | Height | Width | Area | Area % |
|---|---|---|---|---|---|
| 1 | 0.816 | 48.042 | 0.019 | 60.401 | 9.113 |
| 2 | 1.010 | 576.954 | 0.016 | 602.418 | 90.887 |

Signal 2 : DAD1 B, Sig=254,4 Ref=off

| # | Meas. Ret. | Height | Width | Area | Area % |
|---|---|---|---|---|---|
| 1 | 1.010 | 759.583 | 0.016 | 784.489 | 100.000 |

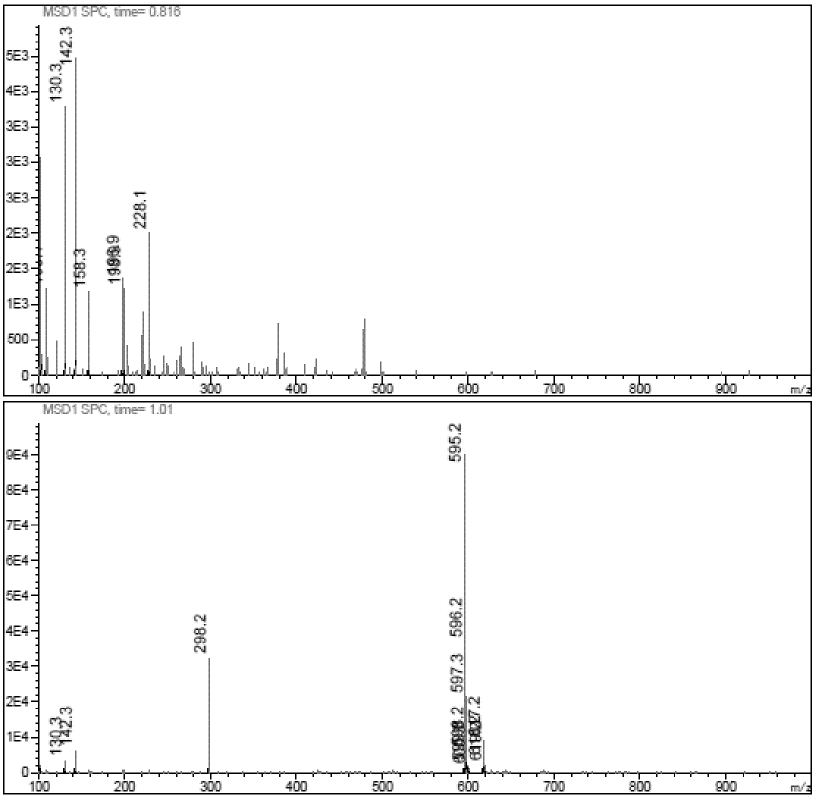

**$^{1}$H NMR** 9-[(2*R*,3*R*,4*R*,5*S*)-3,4-Bis[[tert-butyl(dimethyl)silyl]oxy]-5- (imidazo[1,5-a]pyridin-8-ylsulfanylmethyl)tetrahydrofuran-2-yl]purin-6-amine **(45i)**

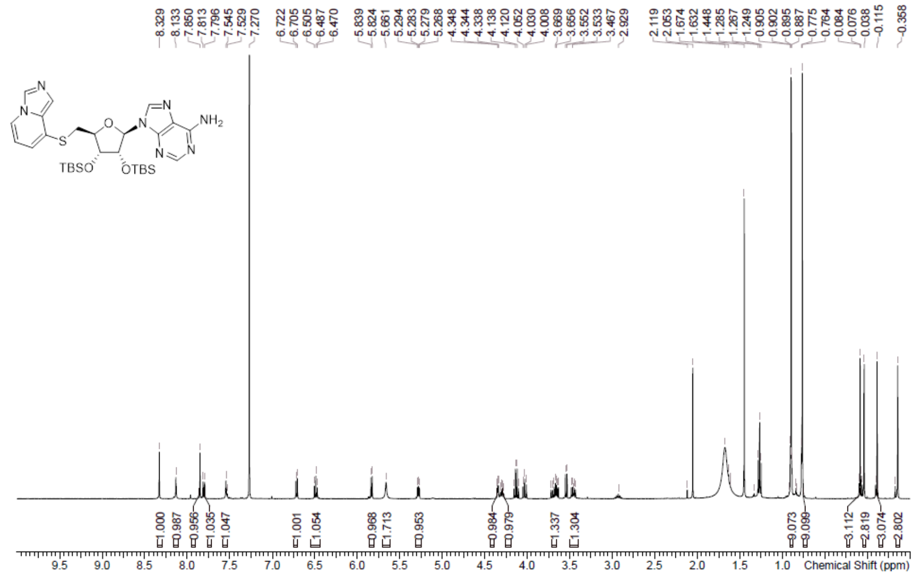

**LCMS** 9-[(2*R*,3*R*,4*R*,5*S*)-3,4-Bis[[tert-butyl(dimethyl)silyl]oxy]-5- (imidazo[1,5-a]pyridin-8-ylsulfanylmethyl)tetrahydrofuran-2-yl]purin-6-amine **(45i)**

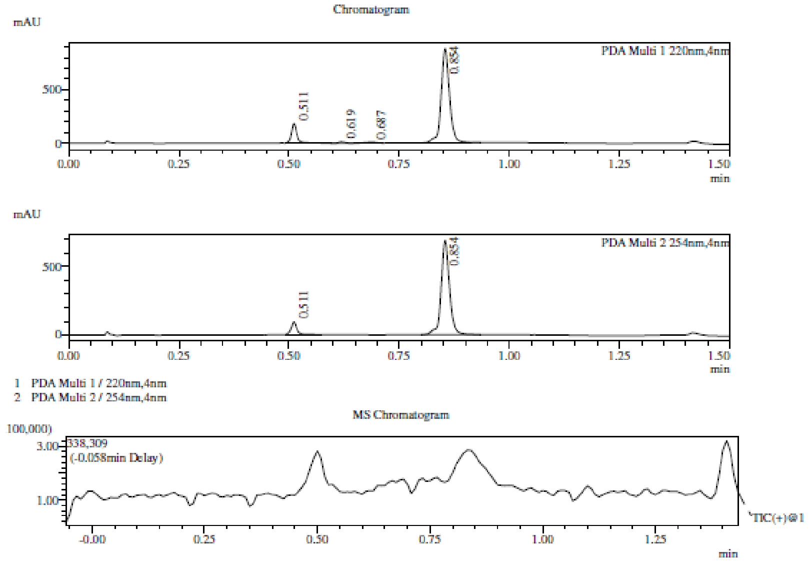

Integration Result

Peak Table

PDA Ch1 220nm

| Peak# | Ret. Time | Height | Height% | USP Width | Area | Area% |
|---|---|---|---|---|---|---|
| 1 | 0.511 | 183504 | 16.951 | 0.024 | 167988 | 12.251 |
| 2 | 0.619 | 12630 | 1.167 | 0.030 | 13184 | 0.961 |
| 3 | 0.687 | 9897 | 0.914 | 0.050 | 17364 | 1.266 |
| 4 | 0.854 | 876552 | 80.969 | 0.035 | 1172720 | 85.522 |

Peak Table

PDA Ch2 254nm

| Peak# | Ret. Time | Height | Height% | USP Width | Area | Area% |
|---|---|---|---|---|---|---|
| 1 | 0.511 | 92065 | 11.781 | 0.025 | 86936 | 8.655 |
| 2 | 0.854 | 689417 | 88.219 | 0.034 | 917488 | 91.345 |

Mass Spectrum

RetTime: 0.508 Datafile: D:\Data\2022\2206\220621\EW26049-920-P1C1.lcd

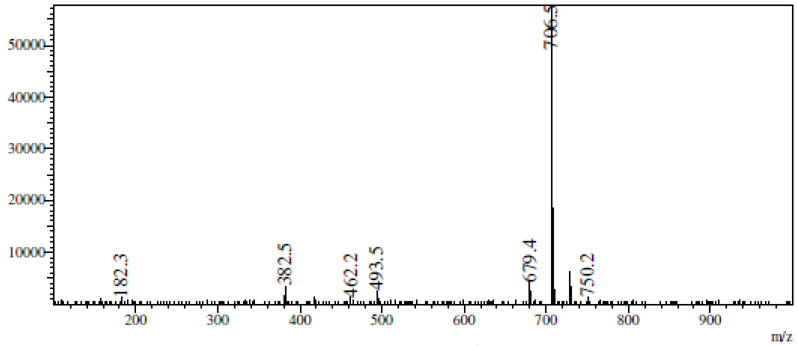

RetTime: 0.617 Datafile: D:\Data\2022\2206\220621\EW26049-920-P1C1.lcd

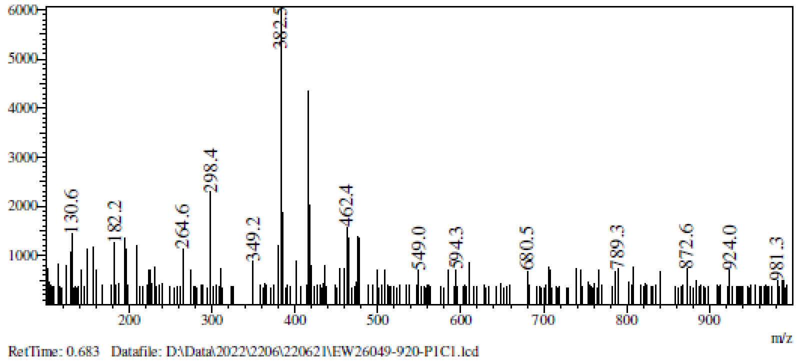

RetTime: 0.683 Datafile: D:\Data\2022\2206\220621\EW26049-920-P1C1.lcd

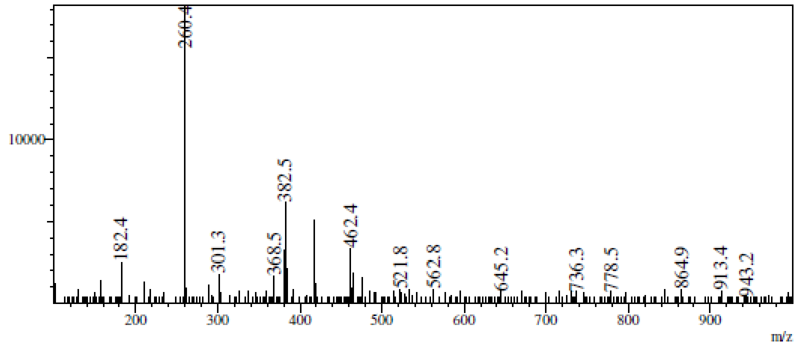

RetTime: 0.858 Datafile: D:\Data\2022\2206\220621\EW26049-920-P1C1.lcd

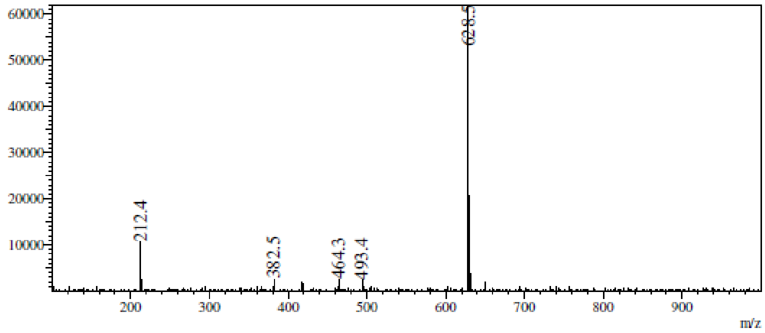

**¹H NMR** 9-((2R,3R,4R,5S)-3,4-Bis((tert-butyldimethylsilyl)oxy)-5-(((1-methyl-1H-benzo[d]imidazol-2-yl)thio)methyl)tetrahydrofuran-2-yl)-9H-purin-6-amine **(45j)**

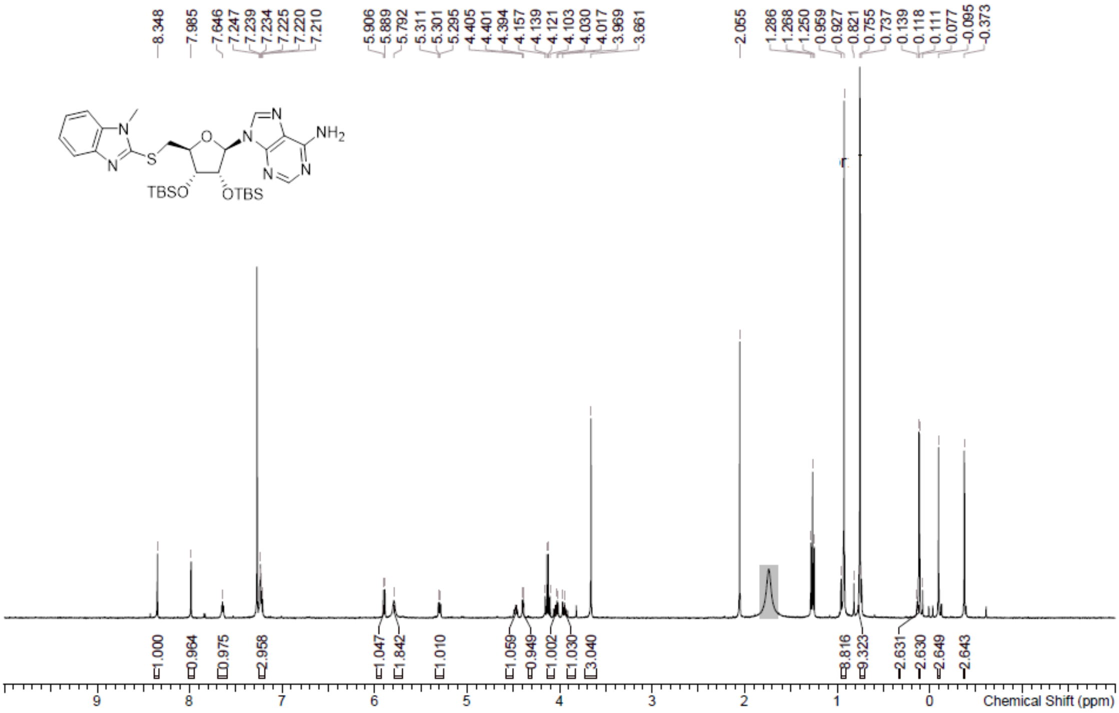

**LCMS** 9-((2R,3R,4R,5S)-3,4-Bis((tert-butyldimethylsilyl)oxy)-5-(((1-methyl-1H-benzo[d]imidazol-2-yl)thio)methyl)tetrahydrofuran-2-yl)-9H-purin-6-amine **(45j)**

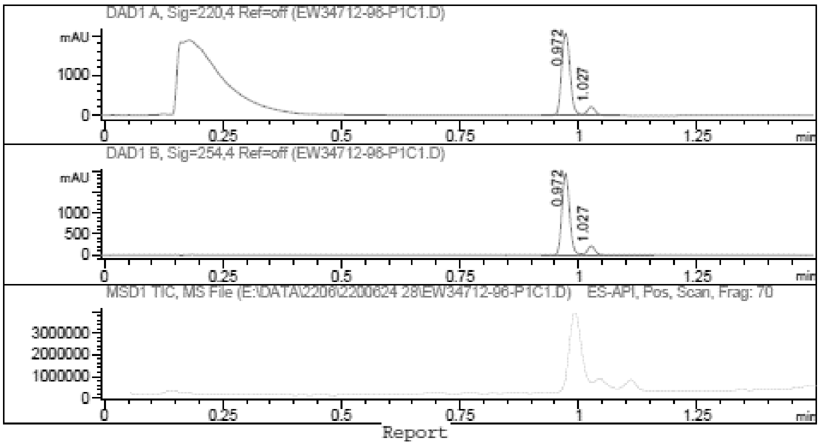

Report

Signal 1 : DAD1 A, Sig=220,4 Ref=off

| # | Meas. Ret. | Height | Width | Area | Area % |
|---|---|---|---|---|---|
| 1 | 0.972 | 2073.677 | 0.020 | 2568.364 | 90.738 |
| 2 | 1.027 | 220.629 | 0.018 | 262.159 | 9.262 |

Signal 2 : DAD1 B, Sig=254,4 Ref=off

| # | Meas. Ret. | Height | Width | Area | Area % |
|---|---|---|---|---|---|
| 1 | 0.972 | 1939.357 | 0.018 | 2162.180 | 89.326 |
| 2 | 1.027 | 214.098 | 0.018 | 258.357 | 10.674 |

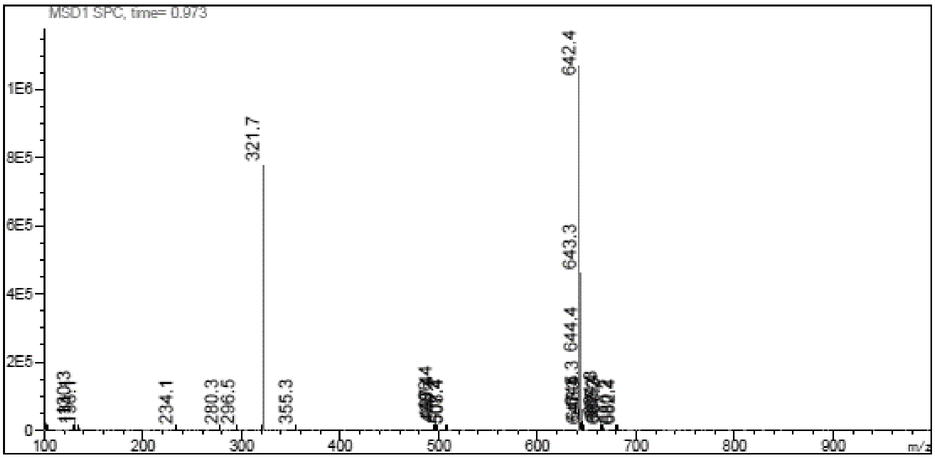
MSD1 SPC, time= 0.973
642.4
321.7
643.3
644.4
234.1
280.3
296.5
355.3
m/z
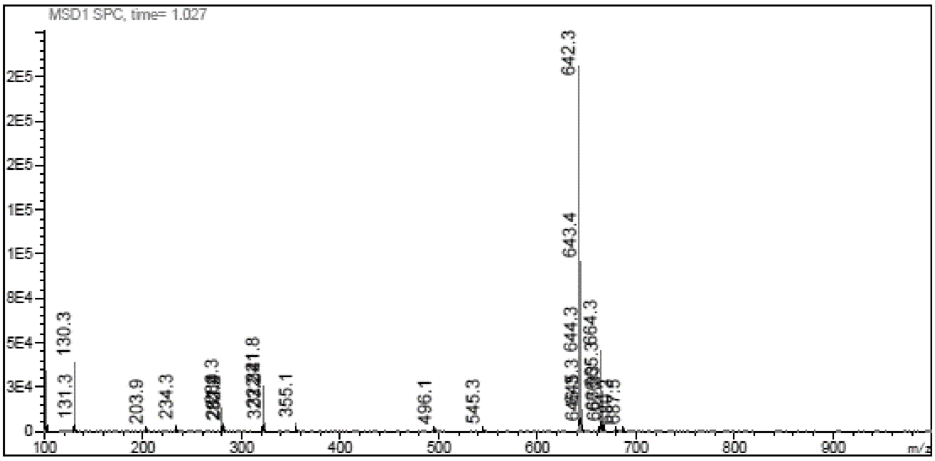
MSD1 SPC, time= 1.027
642.3
643.4
130.3
131.3
203.9
234.3
355.1
496.1
545.3
m/z

**LCMS** 9-((2R,3R,4R,5S)-5-((Benzo[d]oxazol-2-ylthio)methyl)-3,4-bis((tertbutyldimethylsilyl) oxy)tetrahydrofuran-2-yl)-9H-purin-6-amine **(45k)**

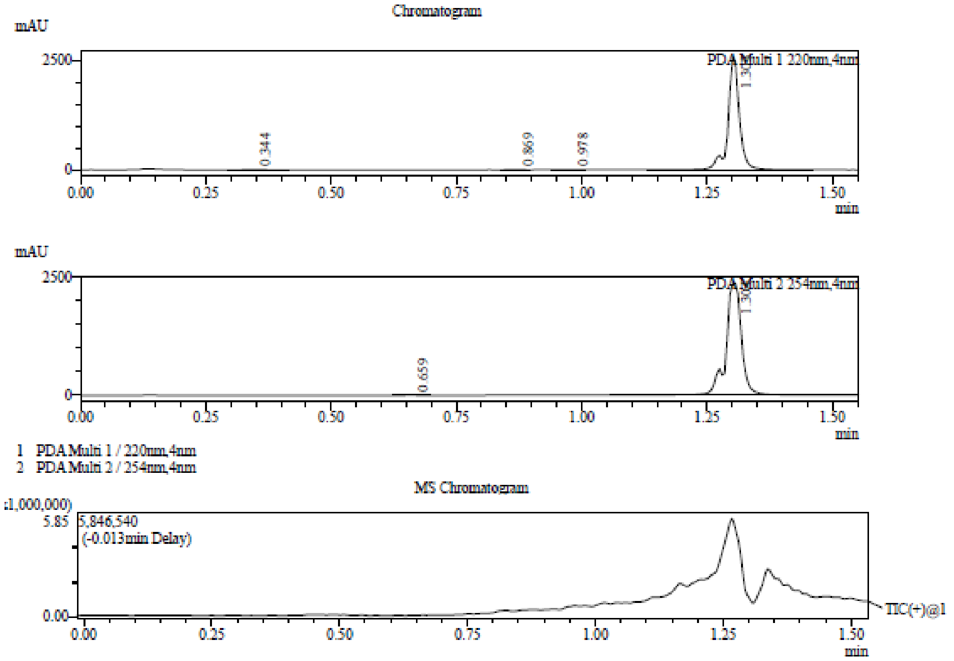

Integration Result

Peak Table

PDA Ch1 220nm

| Peak# | Ret. Time | Height | Height% | USP Width | Area | Area% |
|---|---|---|---|---|---|---|
| 1 | 0.344 | 1963 | 0.077 | 0.100 | 6715 | 0.164 |
| 2 | 0.869 | 2955 | 0.115 | 0.045 | 5135 | 0.125 |
| 3 | 0.978 | 4117 | 0.160 | 0.064 | 8736 | 0.213 |
| 4 | 1.303 | 2556725 | 99.648 | 0.043 | 4076287 | 99.498 |

Peak Table

PDA Ch2 254nm

| Peak# | Ret. Time | Height | Height% | USP Width | Area | Area% |
|---|---|---|---|---|---|---|
| 1 | 0.659 | 4142 | 0.174 | 0.044 | 6144 | 0.119 |
| 2 | 1.304 | 2375088 | 99.826 | 0.059 | 5163276 | 99.881 |

Mass Spectrum

RetTime: 0.343 Datafile: D:\DATA\2022\2207\220701\EW34712-102-P1C1.lcd

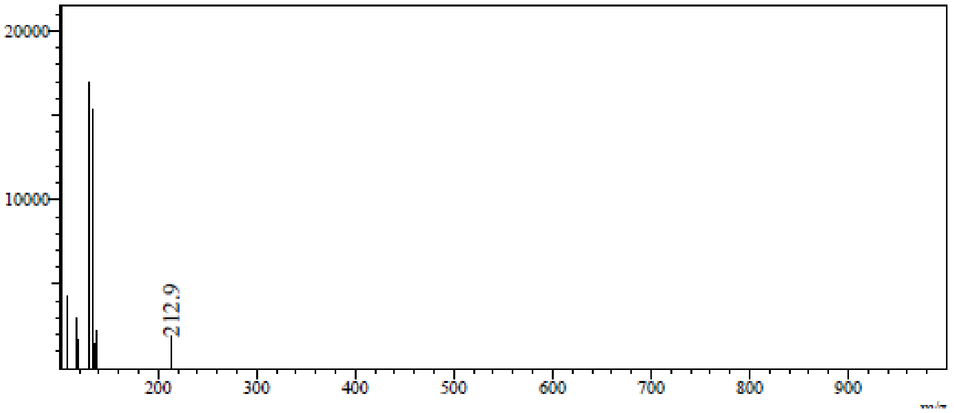

RetTime: 0.870 Datafile: D:\DATA\2022\2207\220701\EW34712-102-P1C1.lcd

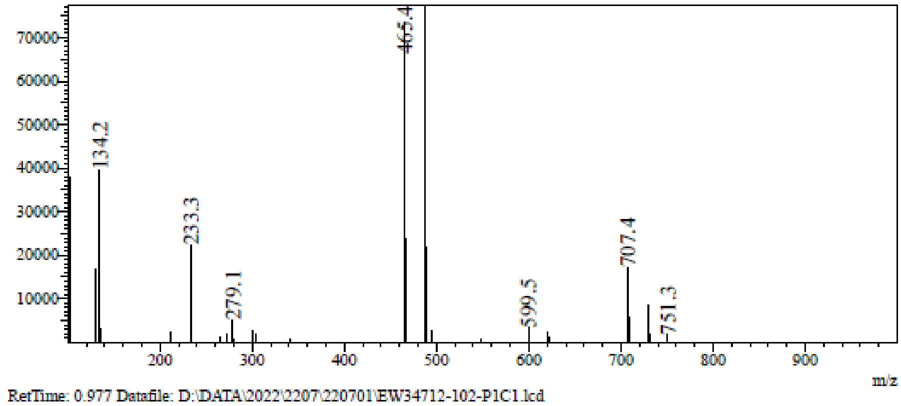

RetTime: 0.977 Datafile: D:\DATA\2022\2207\220701\EW34712-102-P1C1.lcd

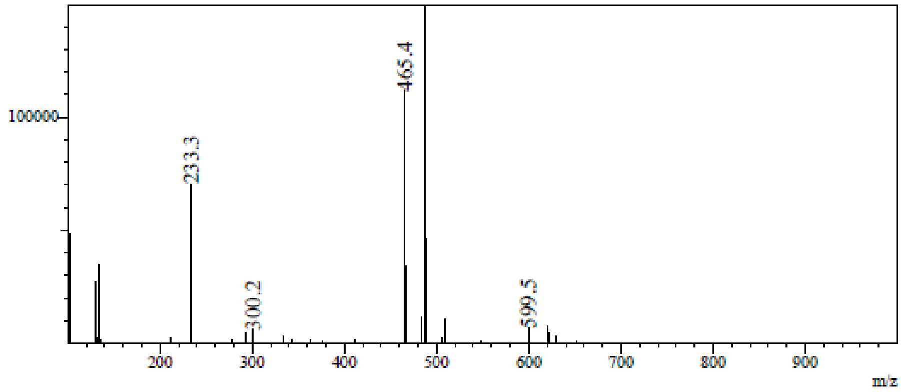

RetTime: 1.303 Datafile: D:\DATA\2022\2207\220701\EW34712-102-P1C1.lcd

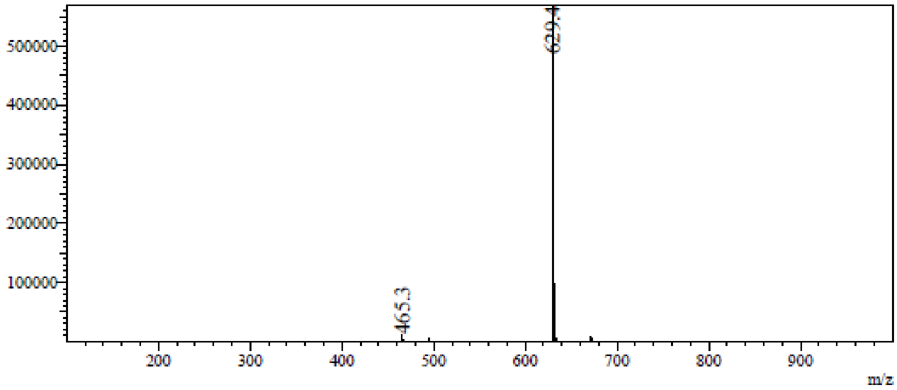

**LCMS** 9-((2R,3R,4R,5S)-5-(((1H-Benzo[d]imidazol-2-yl)thio)methyl)-3,4-bis((tertbutyldimethyl silyl)oxy)tetrahydrofuran-2-yl)-9H-purin-6-amine **(45l)**

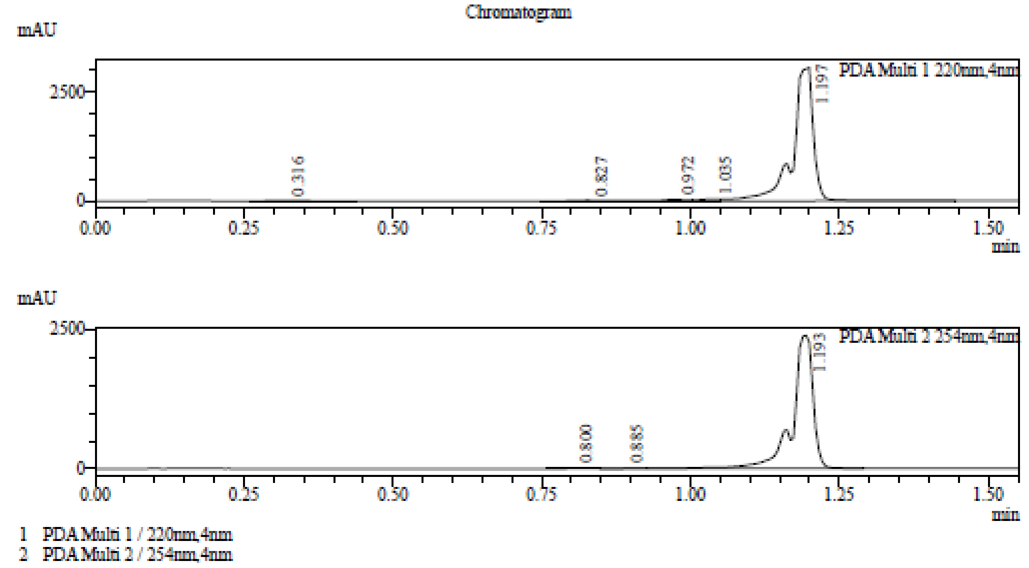

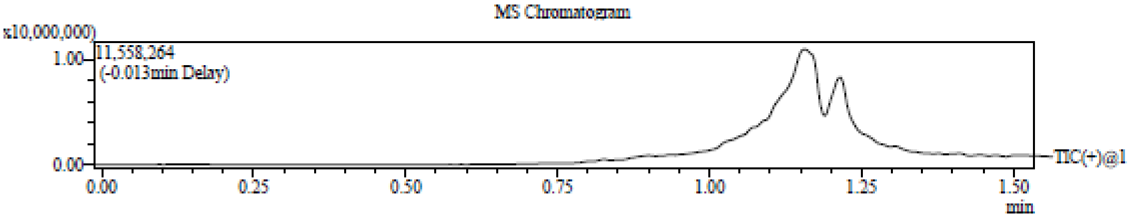

Integration Result

Peak Table

PDA Ch1 220nm

| Peak# | Ret. Time | Height | Height% | USP Width | Area | Area% |
|---|---|---|---|---|---|---|
| 1 | 0.316 | 5662 | 0.179 | 0.109 | 21695 | 0.281 |
| 2 | 0.827 | 17028 | 0.540 | 0.045 | 37134 | 0.481 |
| 3 | 0.972 | 35702 | 1.132 | 0.087 | 138947 | 1.798 |
| 4 | 1.035 | 41918 | 1.329 | 0.230 | 95377 | 1.235 |
| 5 | 1.197 | 3054270 | 96.820 | 0.224 | 7432714 | 96.206 |

Peak Table

PDA Ch2 254nm

| Peak# | Ret. Time | Height | Height% | USP Width | Area | Area% |
|---|---|---|---|---|---|---|
| 1 | 0.800 | 9089 | 0.379 | 0.091 | 17746 | 0.303 |
| 2 | 0.885 | 5920 | 0.247 | 0.258 | 23826 | 0.407 |
| 3 | 1.193 | 2384951 | 99.375 | 0.059 | 5815172 | 99.290 |

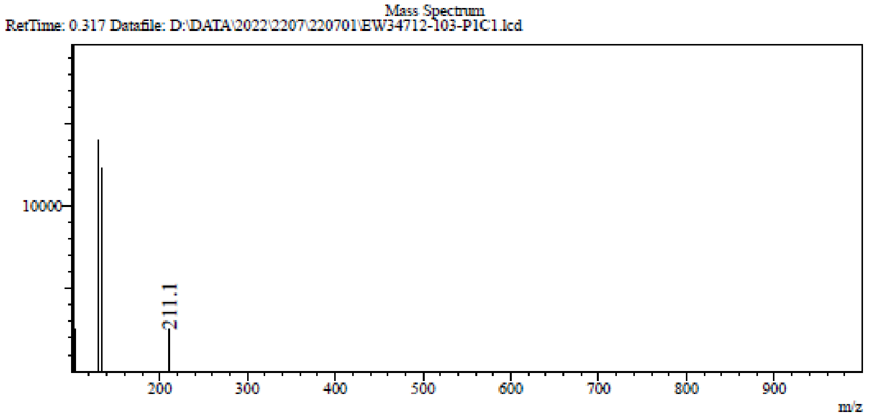

RetTime: 0.827 Datafile: D:\DATA\2022\2207\220701\EW34712-103-P1C1.lcd

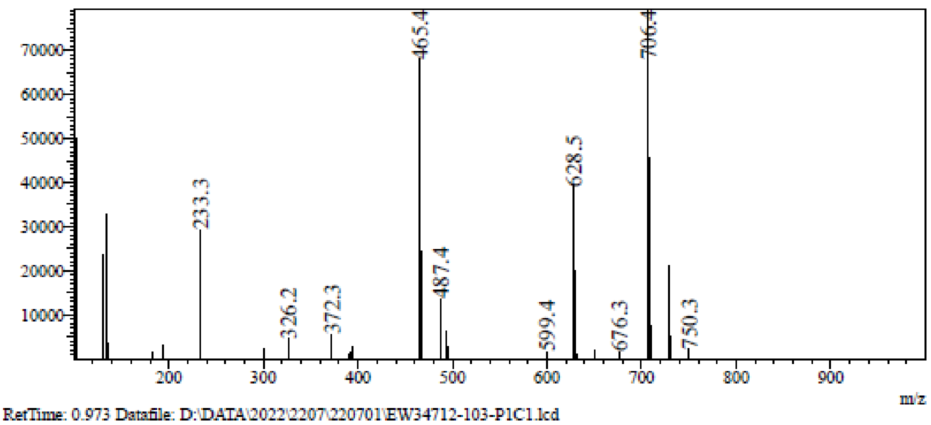

RetTime: 0.973 Datafile: D:\DATA\2022\2207\220701\EW34712-103-P1C1.lcd

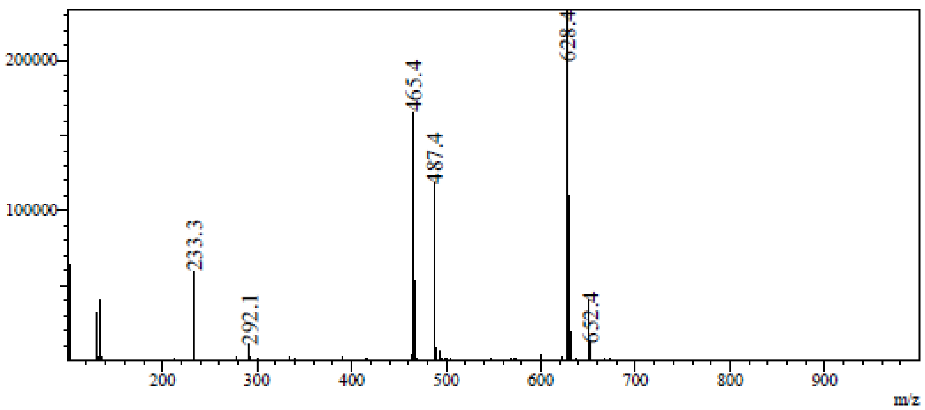

RetTime: 1.037 Datafile: D:\DATA\2022\2207\220701\EW34712-103-P1C1.lcd

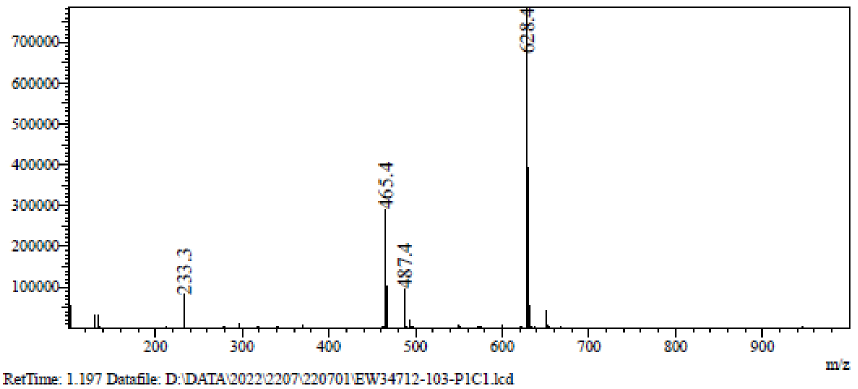

RetTime: 1.197 Datafile: D:\DATA\2022\2207\220701\EW34712-103-P1C1.lcd

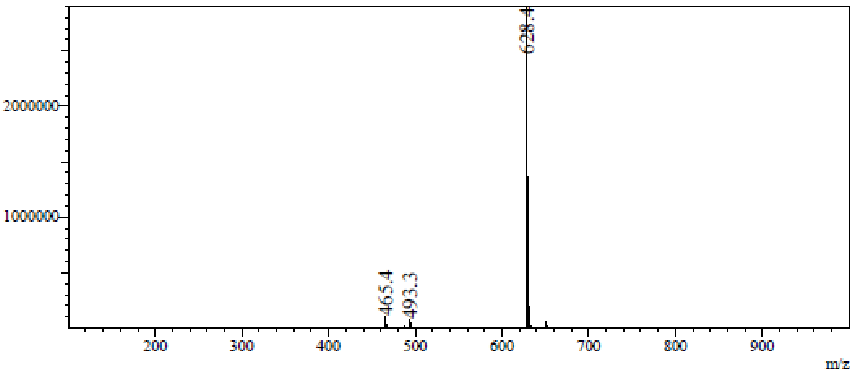

**LCMS** 9-((2R,3R,4R,5S)-5-(((1H-Indazol-7-yl)thio)methyl)-3,4-bis((tert-butyldimethylsilyl)oxy) tetrahydrofuran-2-yl)-9H-purin-6-amine **(45m)**

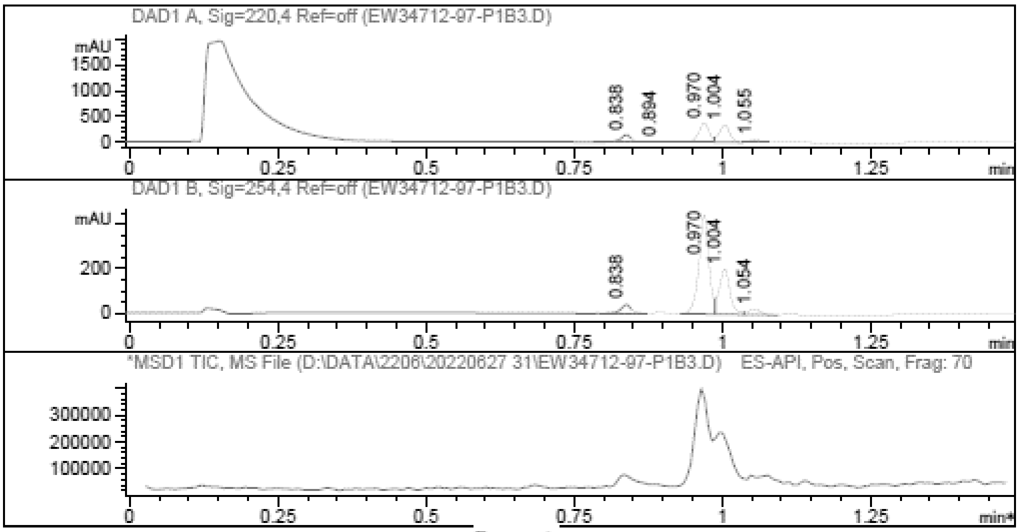

Report

=================================================================

Signal 1 : DAD1 A, Sig=220,4 Ref=off

| # | Meas. Ret. | Height | Width | Area | Area % |
|---|---|---|---|---|---|
| 1 | 0.838 | 135.808 | 0.019 | 168.799 | 14.768 |
| 2 | 0.894 | 24.034 | 0.021 | 34.284 | 2.999 |
| 3 | 0.970 | 372.911 | 0.018 | 443.779 | 38.825 |
| 4 | 1.004 | 343.672 | 0.019 | 425.769 | 37.249 |
| 5 | 1.055 | 46.438 | 0.022 | 70.403 | 6.159 |

Signal 2 : DAD1 B, Sig=254,4 Ref=off

| # | Meas. Ret. | Height | Width | Area | Area % |
|---|---|---|---|---|---|
| 1 | 0.838 | 39.049 | 0.019 | 49.638 | 5.668 |
| 2 | 0.970 | 446.271 | 0.018 | 532.231 | 60.779 |
| 3 | 1.004 | 203.776 | 0.019 | 257.435 | 29.398 |
| 4 | 1.054 | 24.235 | 0.022 | 36.381 | 4.155 |

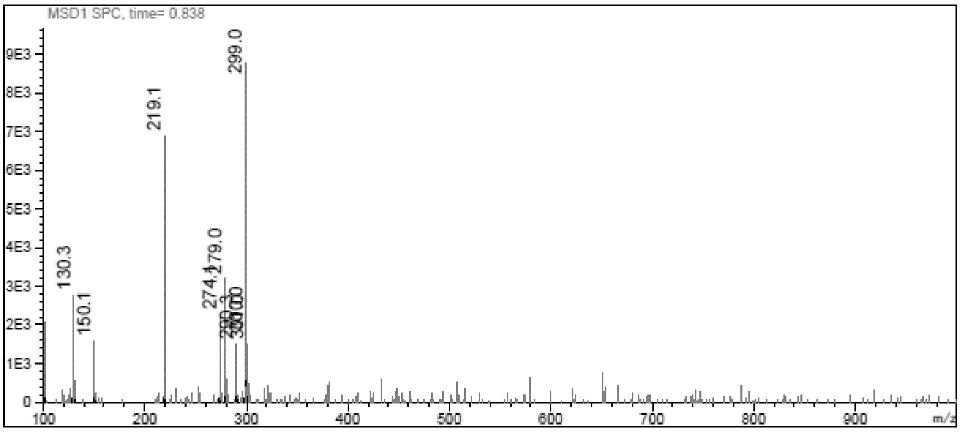

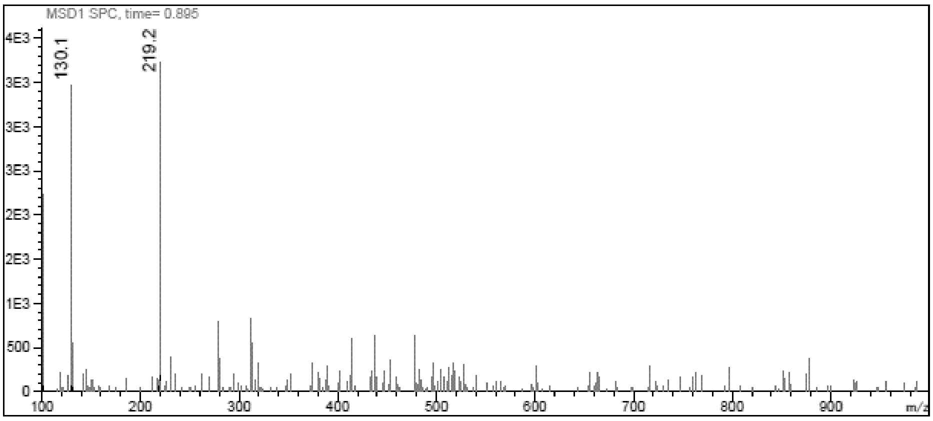
MSD1 SPC, time= 0.895
130.1
219.2
m/z

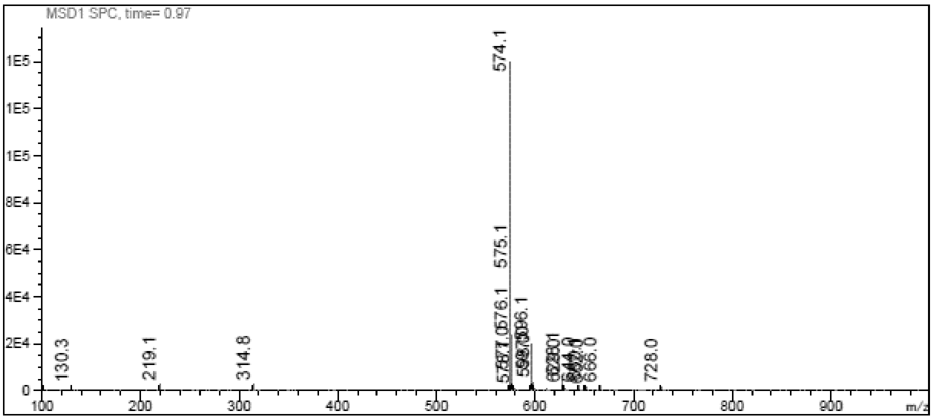
MSD1 SPC, time= 0.97
574.1
575.1
130.3
219.1
314.8
666.0
728.0
m/z

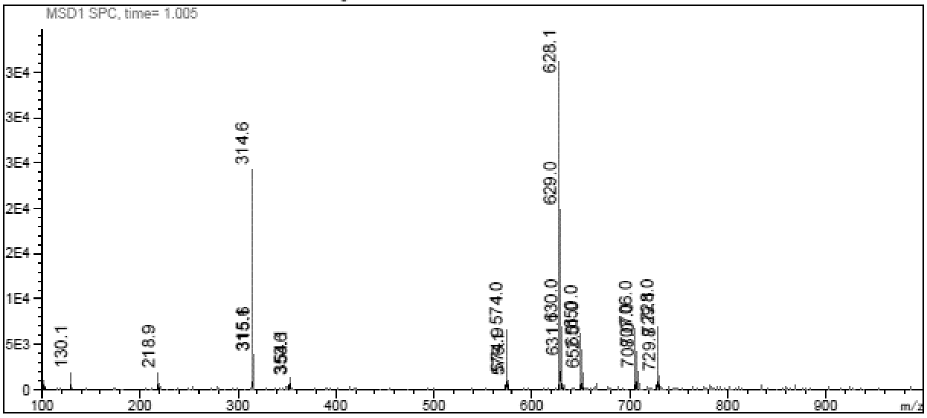
MSD1 SPC, time= 1.005
628.1
314.6
629.0
130.1
218.9
574.0
m/z

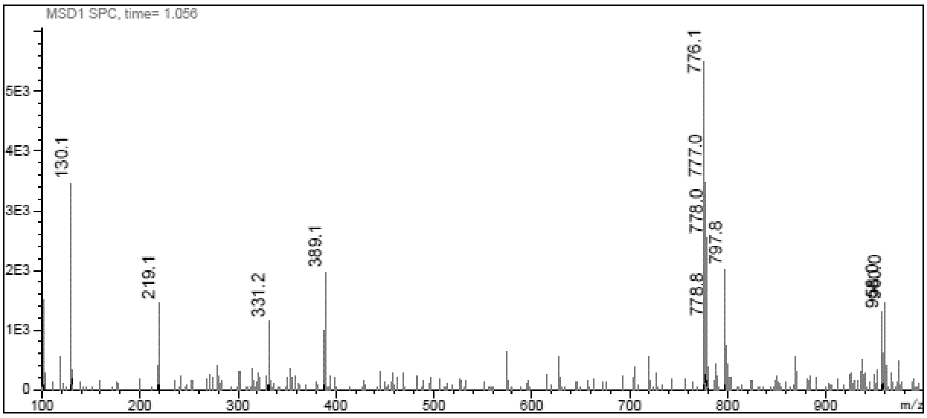
MSD1 SPC, time= 1.056
776.1
777.0
130.1
778.0
797.8
389.1
219.1
331.2
778.8
m/z

**[1]H NMR** 9-[(2*R*,3*R*,4*R*,5*S*)-3,4-bis[[tert-butyl(dimethyl)silyl]oxy]-5-(imidazo [1,5-a]pyridin-7-ylsulfanylmethyl)tetrahydrofuran-2-yl]purin-6-amine **(45n)**

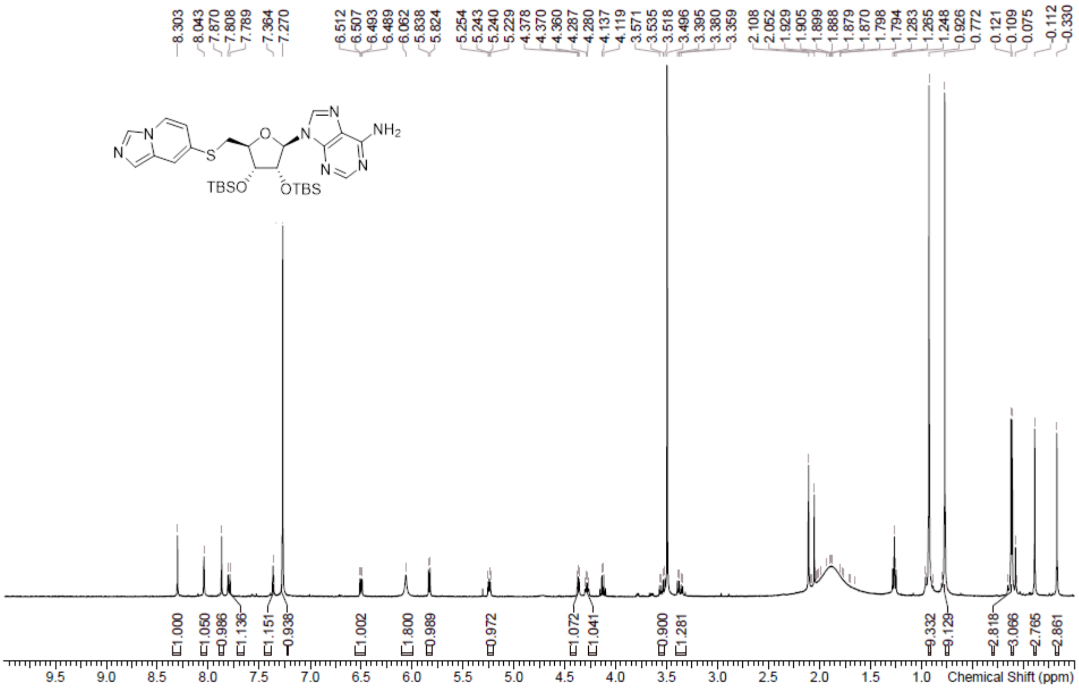

**LCMS** 9-[(2*R*,3*R*,4*R*,5*S*)-3,4-bis[[tert-butyl(dimethyl)silyl]oxy]-5-(imidazo [1,5-a]pyridin-7-ylsulfanylmethyl)tetrahydrofuran-2-yl]purin-6-amine **(45n)**

Chromatogram

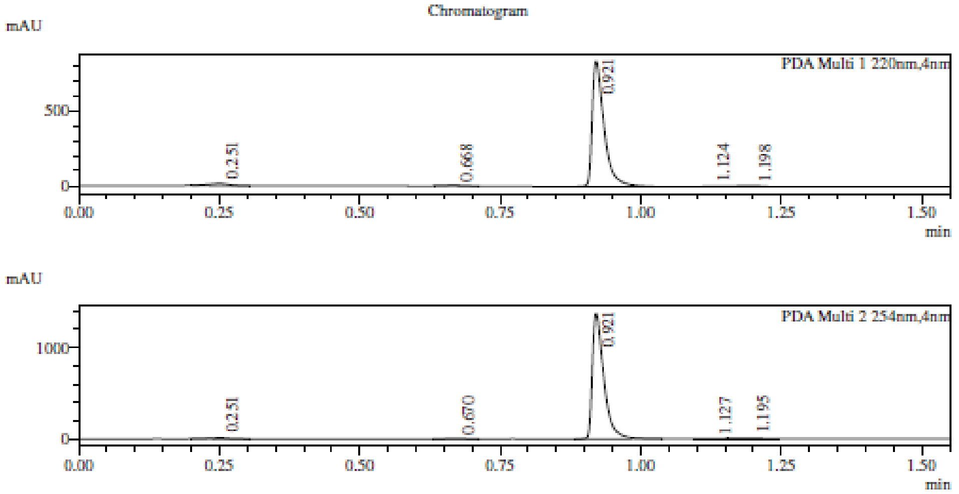

1 PDA Multi 1 / 220nm,4nm
2 PDA Multi 2 / 254nm,4nm

MS Chromatogram

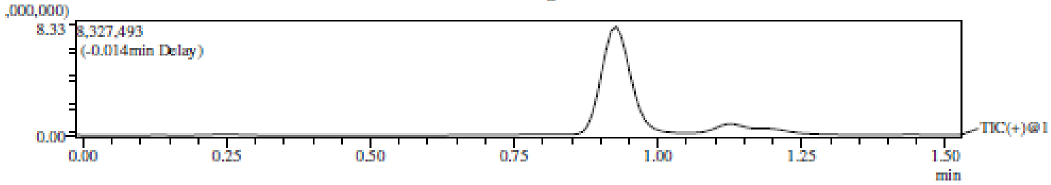

Integration Result

Peak Table

PDA Ch1 220nm

| Peak# | Ret. Time | Height | Height% | USP Width | Area | Area% |
|---|---|---|---|---|---|---|
| 1 | 0.251 | 14041 | 1.628 | 0.068 | 34180 | 2.610 |
| 2 | 0.668 | 4342 | 0.504 | 0.069 | 11039 | 0.843 |
| 3 | 0.921 | 833221 | 96.636 | 0.038 | 1232865 | 94.145 |
| 4 | 1.124 | 3272 | 0.380 | 0.286 | 15568 | 1.189 |
| 5 | 1.198 | 7355 | 0.853 | 0.096 | 15891 | 1.213 |

Peak Table

PDA Ch2 254nm

| Peak# | Ret. Time | Height | Height% | USP Width | Area | Area% |
|---|---|---|---|---|---|---|
| 1 | 0.251 | 11948 | 0.858 | 0.068 | 29469 | 1.451 |
| 2 | 0.670 | 3903 | 0.280 | 0.068 | 10322 | 0.508 |
| 3 | 0.921 | 1367846 | 98.179 | 0.037 | 1960629 | 96.540 |
| 4 | 1.127 | 2830 | 0.203 | 0.000 | 8108 | 0.399 |
| 5 | 1.195 | 6695 | 0.481 | 0.109 | 22376 | 1.102 |

Mass Spectrum

RetTime: 0.253 Datafile: D:\DATA\2022\2207\220726\EW26049-970-P1S1.lcd

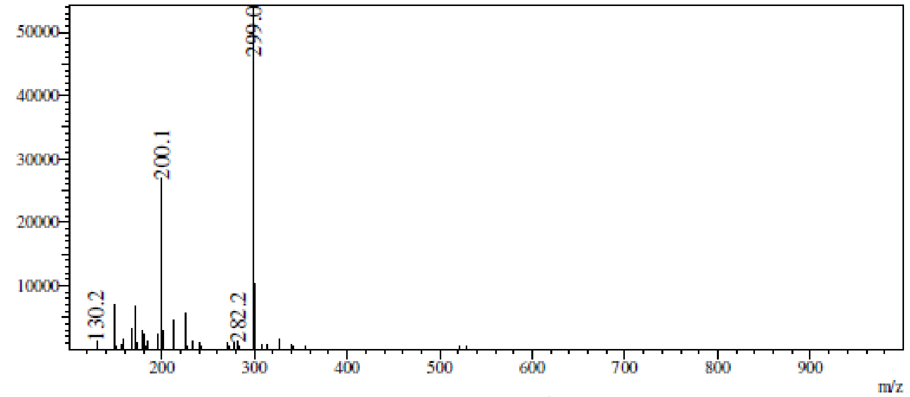

m/z

RetTime: 0.669 Datafile: D:\DATA\2022\2207\220726\EW26049-970-P1S1.lcd

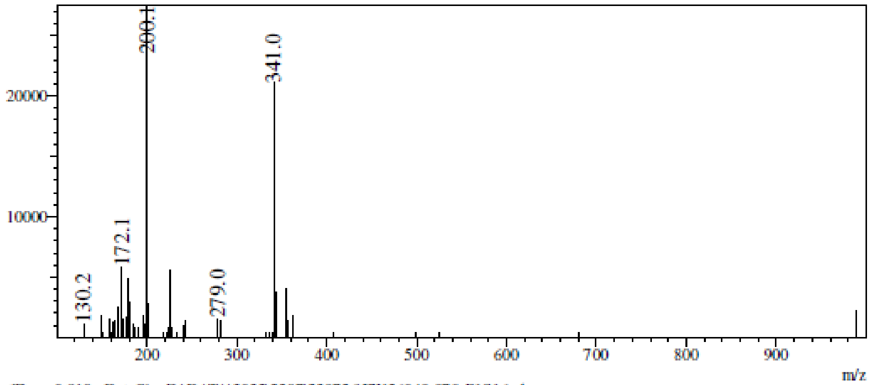

RetTime: 0.919 Datafile: D:\DATA\2022\2207\220726\EW26049-970-P1S1.lcd

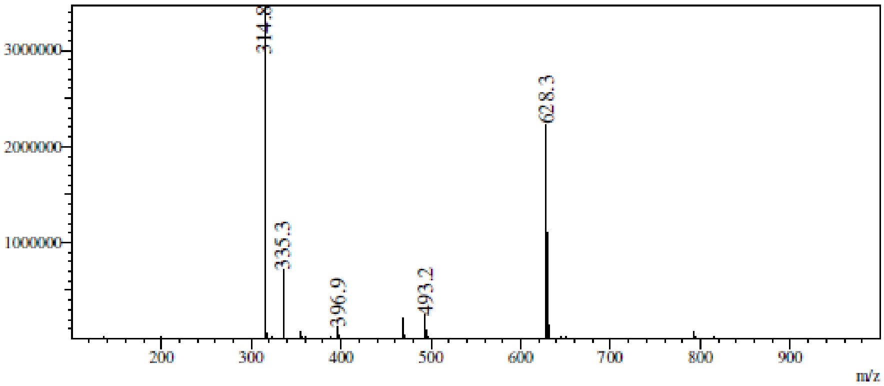

RetTime: 1.128 Datafile: D:\DATA\2022\2207\220726\EW26049-970-P1S1.lcd

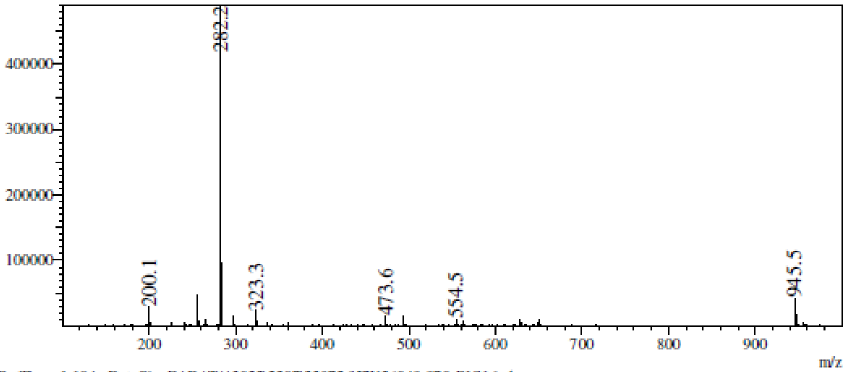

RetTime: 1.194 Datafile: D:\DATA\2022\2207\220726\EW26049-970-P1S1.lcd

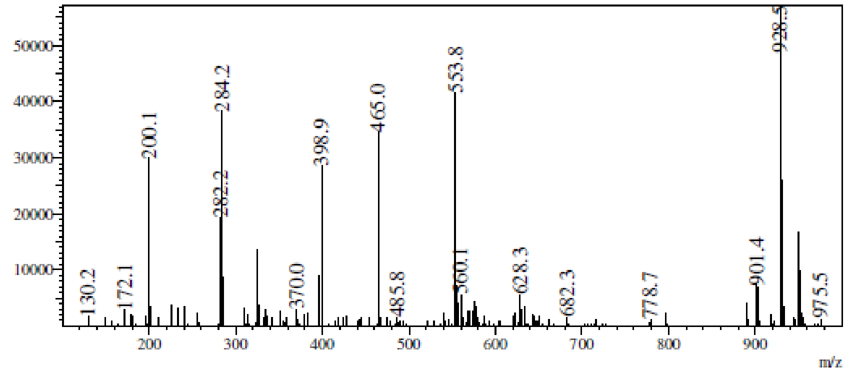

**$^1$H NMR** 9-[(2*R*,3*R*,4*R*,5*R*)-5-(1,3-Benzothiazol-2-yloxymethyl)-3,4-bis[[tert-butyl(dimethyl)silyl]oxy]tetrahydrofuran-2-yl]purin-6-amine **(45o)**

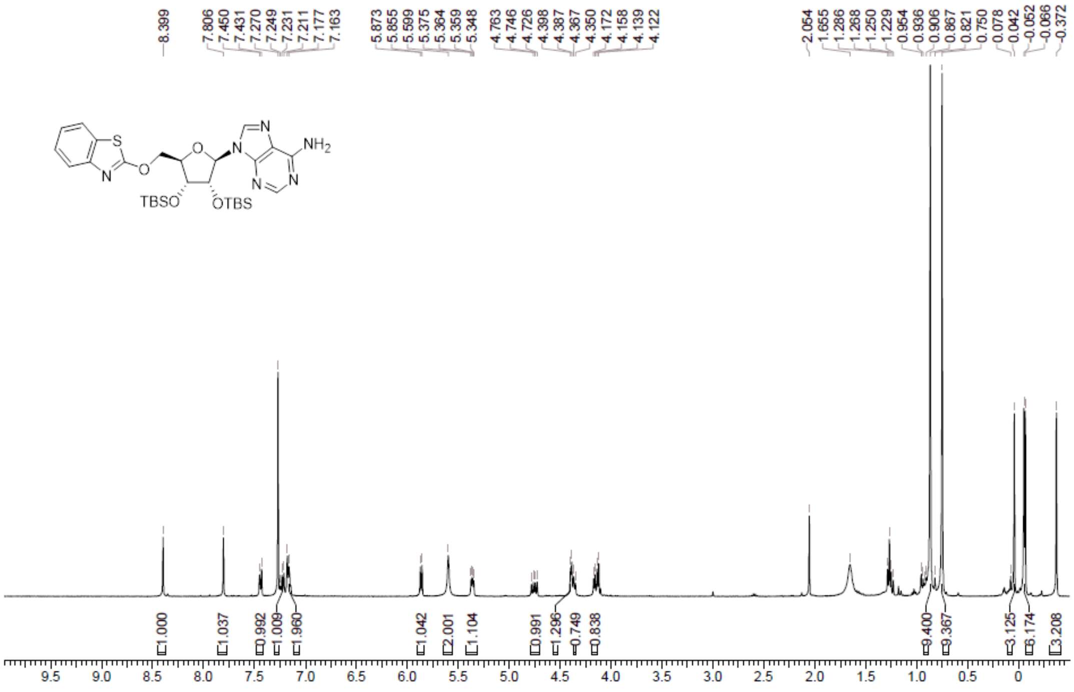

**LCMS** 9-[(2*R*,3*R*,4*R*,5*R*)-5-(1,3-Benzothiazol-2-yloxymethyl)-3,4-bis[[tert-butyl(dimethyl)silyl]oxy]tetrahydrofuran-2-yl]purin-6-amine **(45o)**

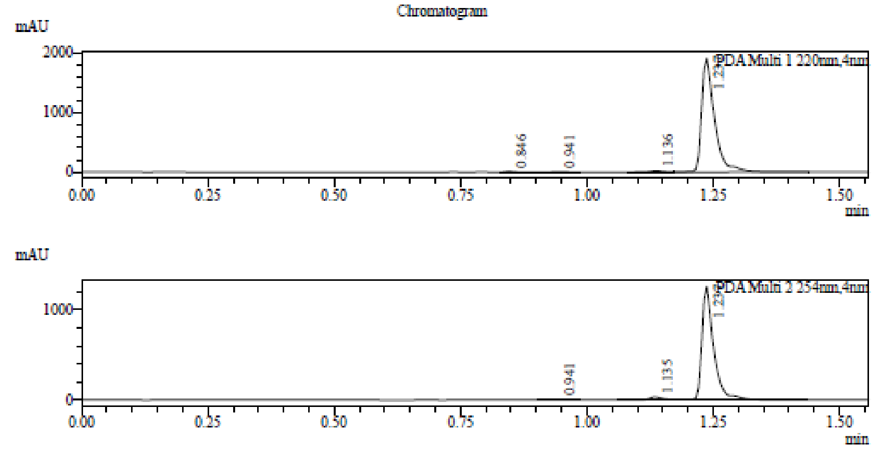

1 PDA Multi 1 / 220nm,4nm
2 PDA Multi 2 / 254nm,4nm

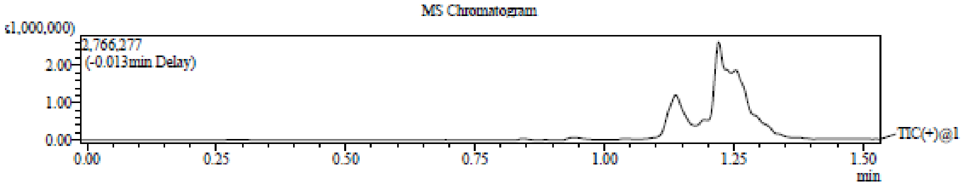

Integration Result

Peak Table

PDA Ch1 220nm

| Peak# | Ret. Time | Height | Height% | USP Width | Area | Area% |
|---|---|---|---|---|---|---|
| 1 | 0.846 | 10568 | 0.545 | 0.029 | 12036 | 0.353 |
| 2 | 0.941 | 6503 | 0.336 | 0.042 | 9611 | 0.282 |
| 3 | 1.136 | 19149 | 0.988 | 0.055 | 36774 | 1.077 |
| 4 | 1.237 | 1901544 | 98.131 | 0.049 | 3355166 | 98.289 |

Peak Table

PDA Ch2 254nm

| Peak# | Ret. Time | Height | Height% | USP Width | Area | Area% |
|---|---|---|---|---|---|---|
| 1 | 0.941 | 3443 | 0.266 | 0.042 | 5125 | 0.244 |
| 2 | 1.135 | 34723 | 2.680 | 0.050 | 62322 | 2.970 |
| 3 | 1.237 | 1257587 | 97.055 | 0.044 | 2030859 | 96.786 |

Mass Spectrum
RetTime: 0.847 Datafile: D:\DATA\2022\2201\220120\EW26049-718-P1S2.lcd

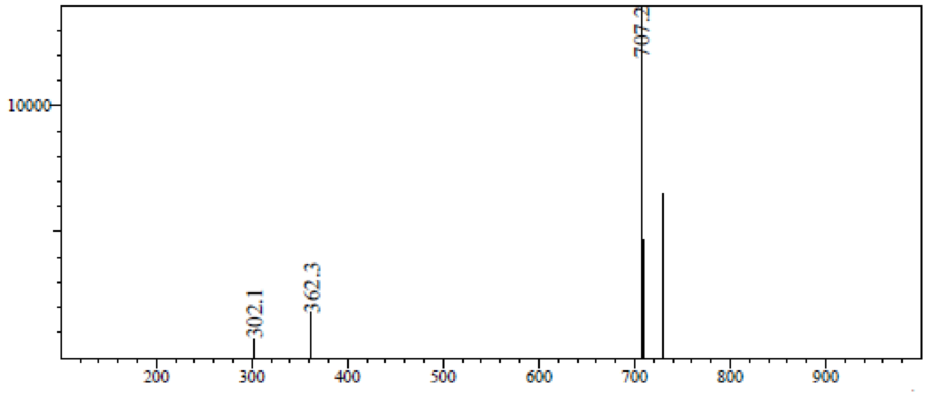

RetTime: 0.940 Datafile: D:\DATA\2022\2201\220120\EW26049-718-P1S2.lcd

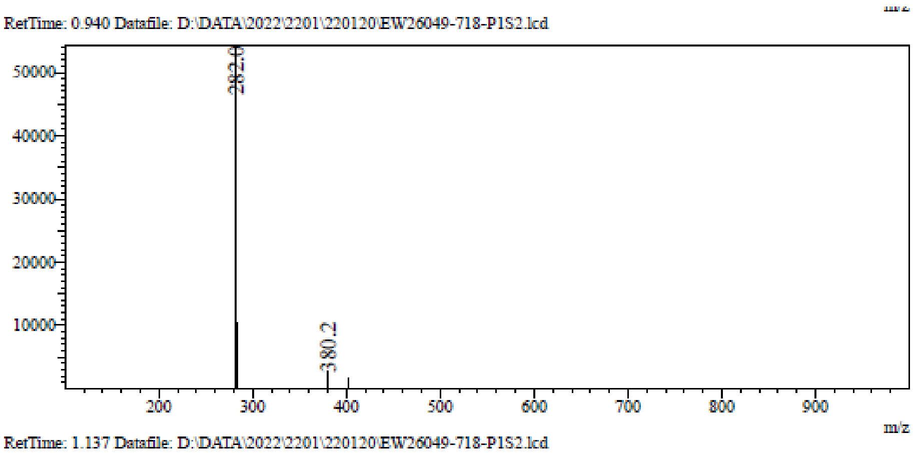

RetTime: 1.137 Datafile: D:\DATA\2022\2201\220120\EW26049-718-P1S2.lcd

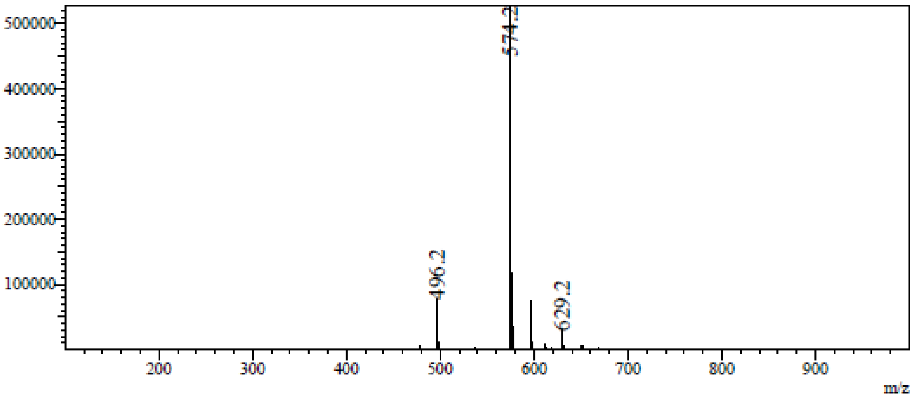

RetTime: 1.237 Datafile: D:\DATA\2022\2201\220120\EW26049-718-P1S2.lcd

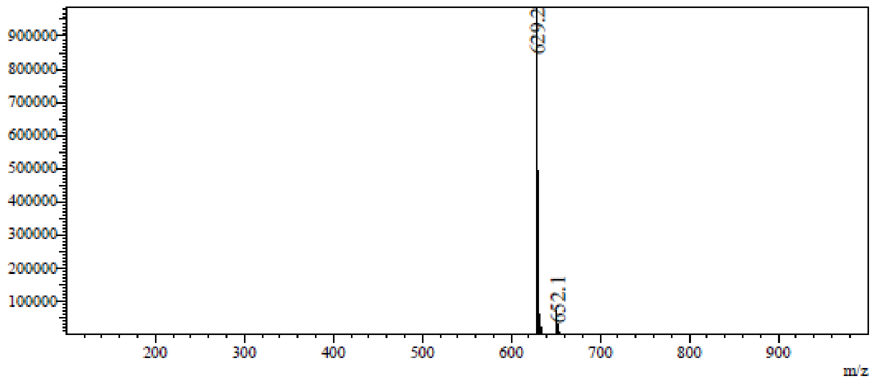

**LCMS** *N*-[[(2*R*,3*R*,4*R*,5*R*)-5-(6-Aminopurin-9-yl)-3,4-bis[[tert-butyl(dimethyl)silyl]oxy] tetrahydrofuran-2-yl]methyl]-1,3-benzothiazol-2-amine **(45p)**

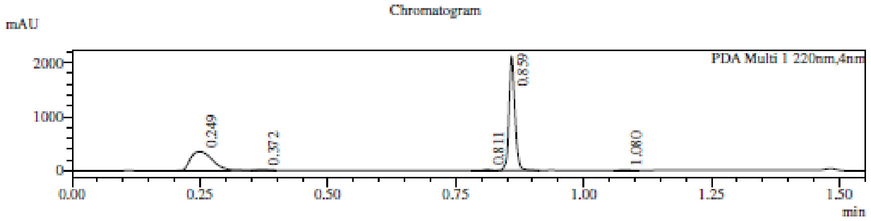

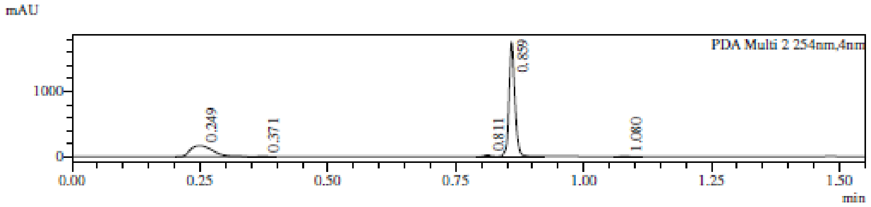

1 PDA Multi 1 / 220nm,4nm
2 PDA Multi 2 / 254nm,4nm

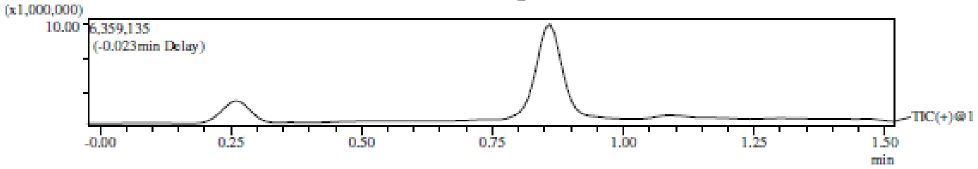

Integration Result

Peak Table

PDA Ch1 220nm

| Peak# | Ret. Time | Height | Height% | USP Width | Area | Area% |
|---|---|---|---|---|---|---|
| 1 | 0.249 | 351253 | 14.042 | 0.077 | 1063423 | 37.939 |
| 2 | 0.372 | 9852 | 0.394 | 0.029 | 10096 | 0.360 |
| 3 | 0.811 | 18509 | 0.740 | 0.027 | 18464 | 0.659 |
| 4 | 0.859 | 2115149 | 84.559 | 0.023 | 1705112 | 60.832 |
| 5 | 1.080 | 6626 | 0.265 | 0.024 | 5869 | 0.209 |

Peak Table

PDA Ch2 254nm

| Peak# | Ret. Time | Height | Height% | USP Width | Area | Area% |
|---|---|---|---|---|---|---|
| 1 | 0.249 | 167094 | 8.561 | 0.077 | 501190 | 25.976 |
| 2 | 0.371 | 4869 | 0.249 | 0.035 | 7309 | 0.379 |
| 3 | 0.811 | 20137 | 1.032 | 0.026 | 18101 | 0.938 |
| 4 | 0.859 | 1753802 | 89.852 | 0.023 | 1397283 | 72.419 |
| 5 | 1.080 | 5984 | 0.307 | 0.025 | 5547 | 0.288 |

Mass Spectrum
RetTime: 0.252 Datafile: D:\DATA\2022\2201\220117\EW26049-712-P1C2.lcd

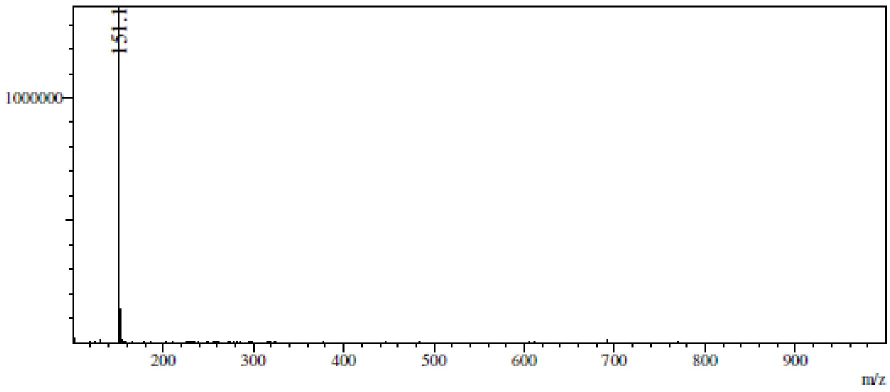

RetTime: 0.368 Datafile: D:\DATA\2022\2201\220117\EW26049-712-P1C2.lcd

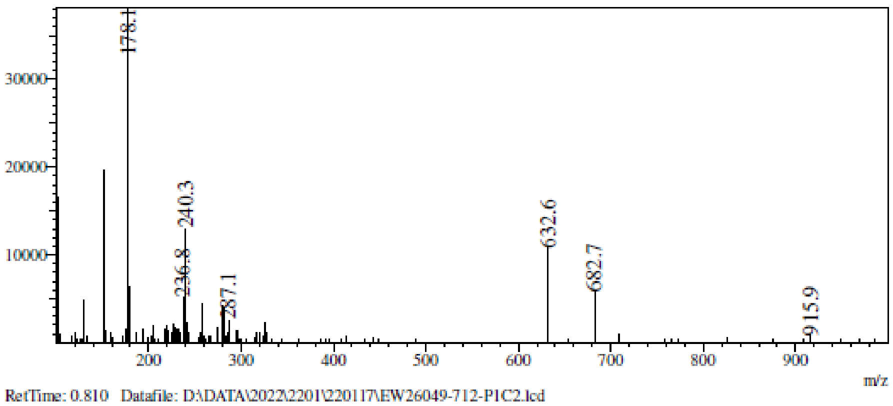

RetTime: 0.810 Datafile: D:\DATA\2022\2201\220117\EW26049-712-P1C2.lcd

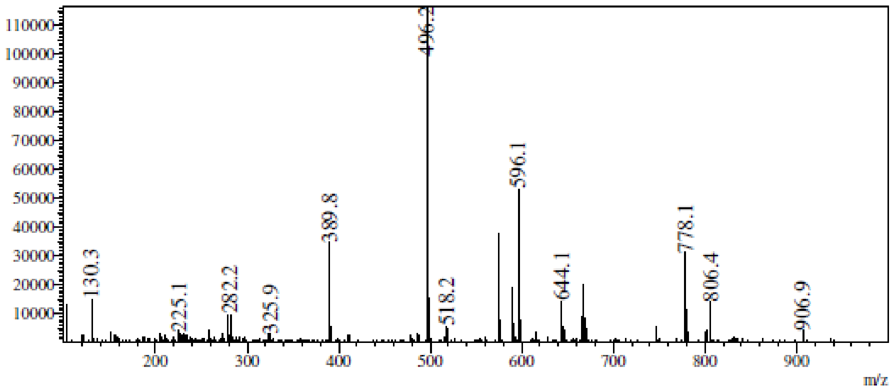

RetTime: 0.860 Datafile: D:\DATA\2022\2201\220117\EW26049-712-P1C2.lcd

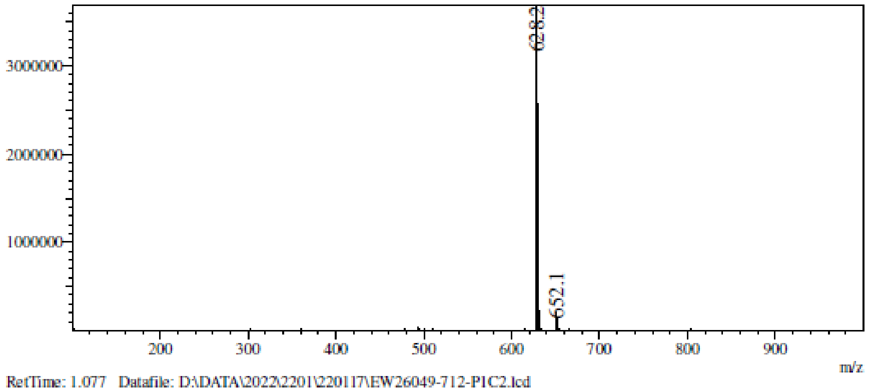

RetTime: 1.077 Datafile: D:\DATA\2022\2201\220117\EW26049-712-P1C2.lcd

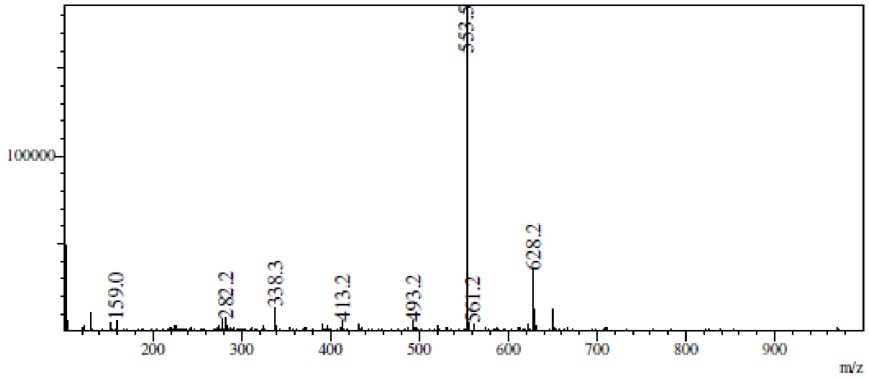

**[1]H NMR** (2*R*,3*R*,4*S*,5*S*)-2-(6-aminopurin-9-yl)-5-(1,3-benzothiazol-2-ylsulfanylmethyl) tetrahydrofuran-3,4-diol **(1)**

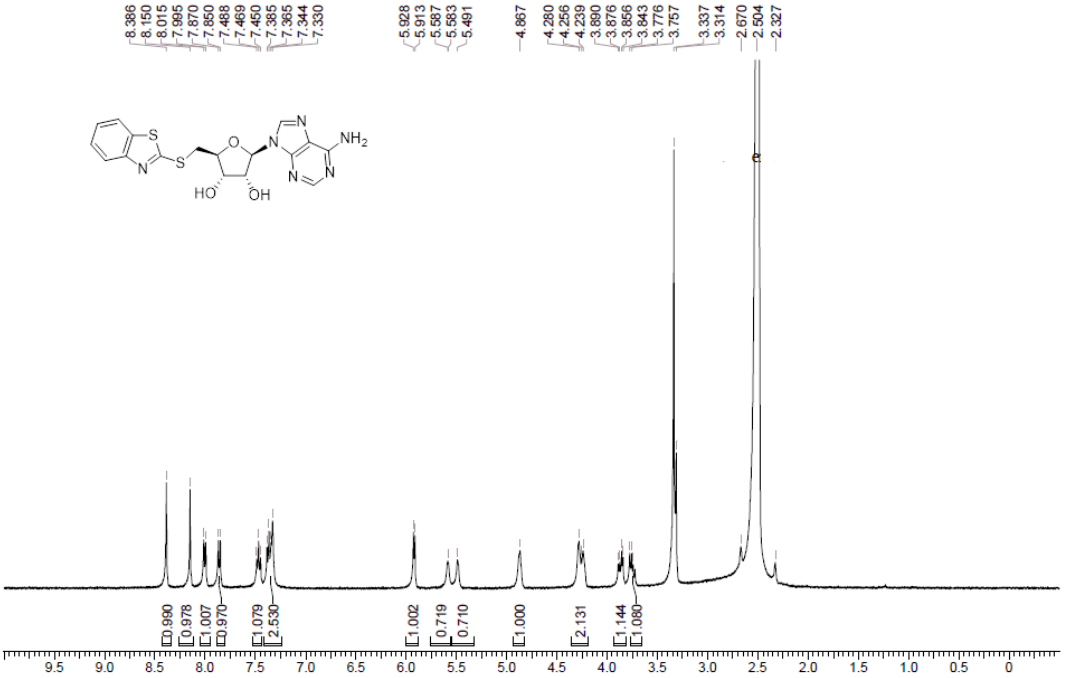

**HPLC** (2*R*,3*R*,4*S*,5*S*)-2-(6-aminopurin-9-yl)-5-(1,3-benzothiazol-2-ylsulfanylmethyl) tetrahydrofuran-3,4-diol **(1)**

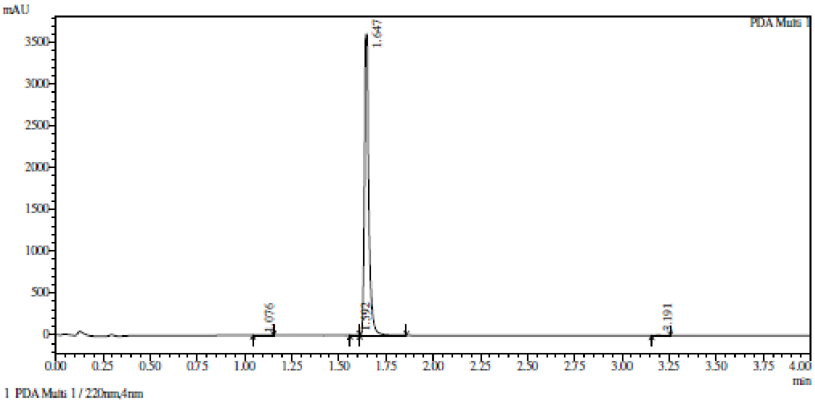

Integration result

PeakTable

PDA Ch1 220nm

| Peak# | Ret. Time | USP Width | Resolution | Height | Area | Area % |
|---|---|---|---|---|---|---|
| 1 | 1.076 | 0.070 | 0.000 | 1021 | 2906 | 0.051 |
| 2 | 1.592 | 0.090 | 6.453 | 601 | 1368 | 0.024 |
| 3 | 1.647 | 0.036 | 0.869 | 3609456 | 5666061 | 99.744 |
| 4 | 3.191 | 0.050 | 35.904 | 5442 | 10272 | 0.181 |
| Total | | | | 3616520 | 5680607 | 100.000 |

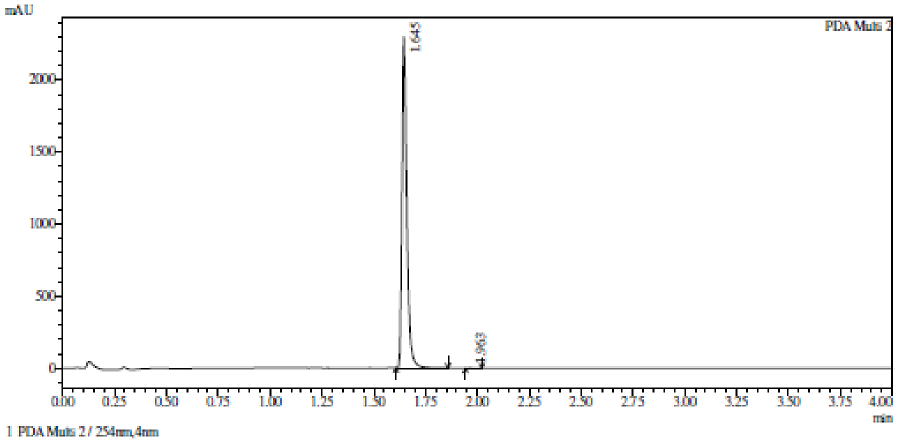

Integration result

PeakTable

PDA Ch2 254nm

| Peak# | Ret. Time | USP Width | Resolution | Height | Area | Area % |
|---|---|---|---|---|---|---|
| 1 | 1.645 | 0.044 | 0.000 | 2297104 | 3741542 | 99.968 |
| 2 | 1.963 | 0.051 | 6.742 | 596 | 1183 | 0.032 |
| Total | | | | 2297700 | 3742726 | 100.000 |

**HRMS** (2*R*,3*R*,4*S*,5*S*)-2-(6-aminopurin-9-yl)-5-(1,3-benzothiazol-2-ylsulfanylmethyl) tetrahydrofuran-3,4-diol **(1)**

Compound Table

| Label | Tgt Score | Mass Error (ppm) | Tgt Formula | Obs. RT | Ref. Mass | Obs. Mass |
|---|---|---|---|---|---|---|
| Cpd 1: C17 H16 N6 O3 S2; 1.687 | 97.92 | 1.73 | C17 H16 N6 O3 S2 | 1.687 | 416.0725 | 416.0733 |

| Obs. *m/z* | Obs. RT | Obs. Mass | Tgt Formula | Tgt Mass | Tgt Mass Error (ppm) | Find Cpds Algorithm |
|---|---|---|---|---|---|---|
| 417.0803 | 1.687 | 416.0733 | C17 H16 N6 O3 S2 | 416.0725 | 1.73 | Find by Formula |

Compound Chromatograms

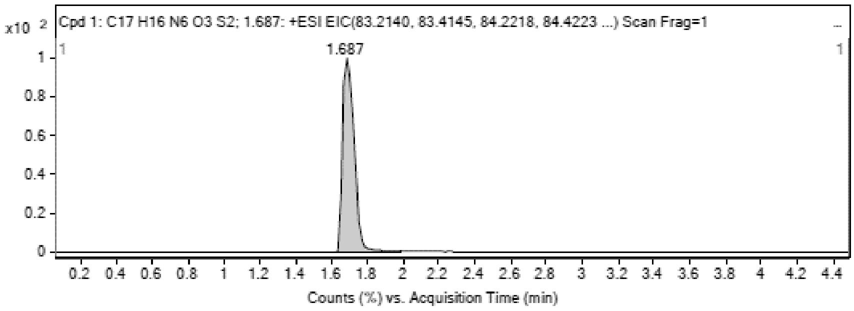

MS Zoomed Spectrum

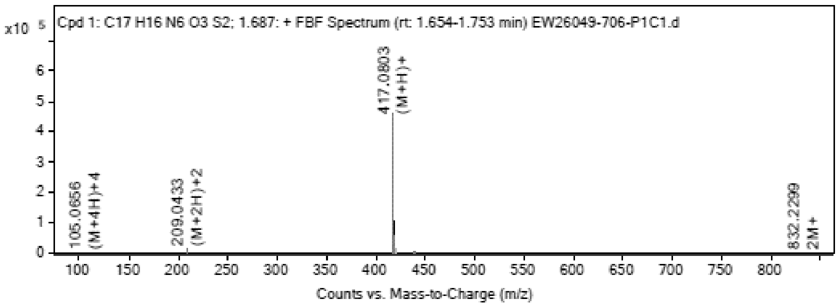

MS Spectrum Peak List

| Obs. *m/z* | Charge | Abund | Ion/Isotope |
|---|---|---|---|
| 105.0656 | 4 | 462.41 | (M+4H)+4 |
| 208.1331 | 2 | 393.25 | M+2 |

| | | | |
|---|---|---|---|
| 209.0433 | 2 | 11833.25 | (M+2H)+2 |
| 209.5443 | 2 | 2876.12 | (M+2H)+2 |
| 210.0417 | 2 | 1317.21 | (M+2H)+2 |
| 417.0803 | 1 | 460341.03 | (M+H)+ |
| 418.0823 | 1 | 102473.61 | (M+H)+ |
| 419.0779 | 1 | 51885.25 | (M+H)+ |
| 439.0625 | 1 | 2859.47 | (M+Na)+ |
| 440.0637 | 1 | 622.13 | (M+Na)+ |

MS Zoomed Spectrum

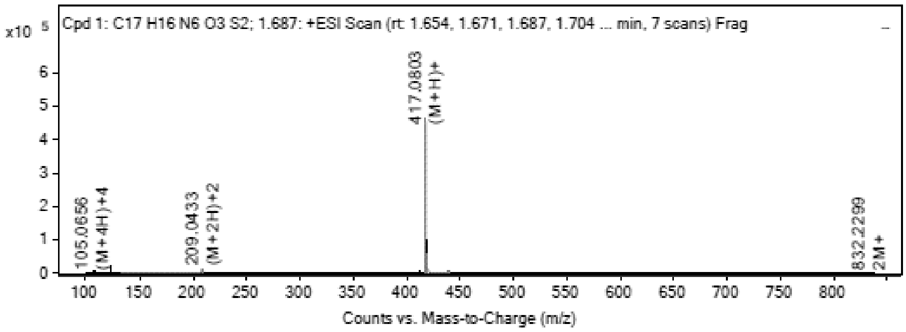

MS Spectrum Peak List

| Obs. m/z | Charge | Abund | Ion/Isotope | Tgt Mass Error (ppm) |
|---|---|---|---|---|
| 105.0656 | 4 | 462.41 | (M+4H)+4 | -382.52 |
| 208.1331 | 2 | 393.25 | M+2 | -468.11 |
| 209.0433 | 2 | 11833.25 | (M+2H)+2 | 1.16 |
| 209.5443 | 2 | 2876.12 | (M+2H)+2 | 2.16 |
| 210.0417 | 2 | 1317.21 | (M+2H)+2 | 4.41 |
| 417.0803 | 1 | 460341.03 | (M+H)+ | -1.14 |
| 418.0823 | 1 | 102473.61 | (M+H)+ | 0.02 |
| 419.0779 | 1 | 51885.25 | (M+H)+ | 0.07 |
| 439.0625 | 1 | 2859.47 | (M+Na)+ | -1.59 |
| 440.0637 | 1 | 622.13 | (M+Na)+ | 1.32 |

--- End Of Report ---

**$^{1}$H NMR** (2*R*,3*R*,4*S*,5*S*)-2-(6-aminopurin-9-yl)-5-[(6-methyl-1,3-benzothiazol -2-yl)sulfanylmethyl] tetrahydrofuran-3,4-diol **(2)**

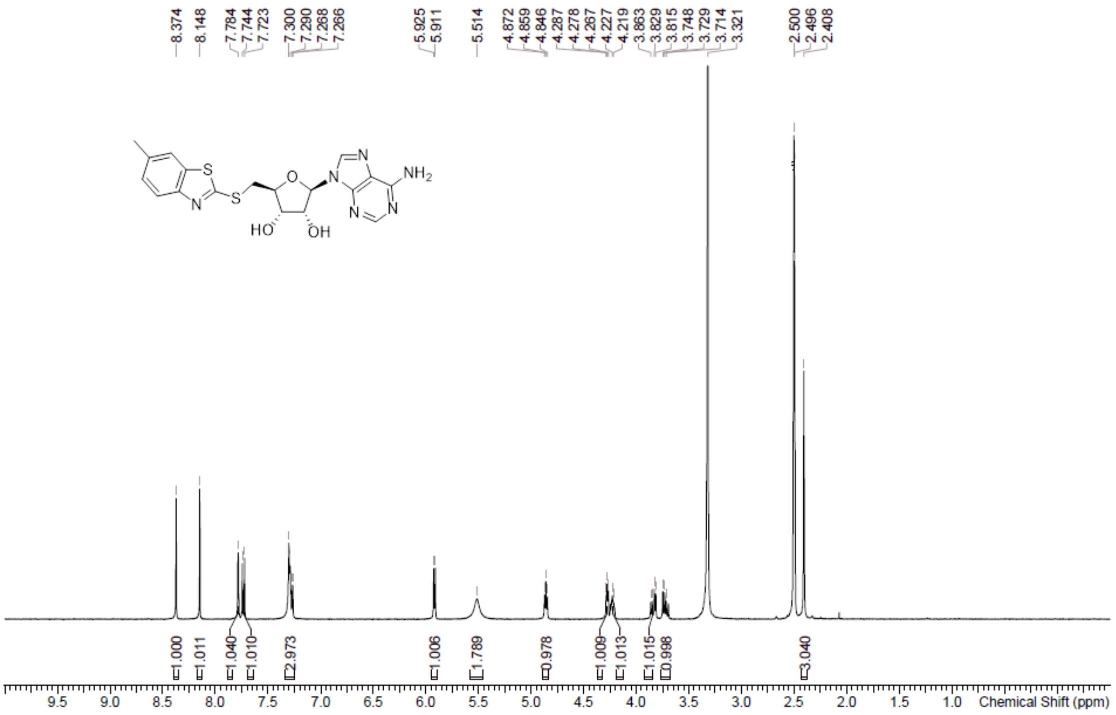

**HPLC** (2*R*,3*R*,4*S*,5*S*)-2-(6-aminopurin-9-yl)-5-[(6-methyl-1,3-benzothiazol -2-yl)sulfanylmethyl] tetrahydrofuran-3,4-diol **(2)**

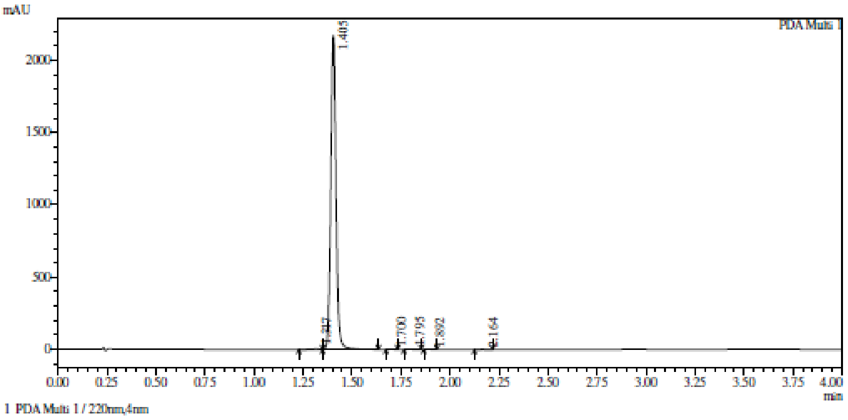

Integration result

PeakTable

PDA Ch1 220nm

| Peak# | Ret. Time | USP Width | Resolution | Height | Area | Area % |
|---|---|---|---|---|---|---|
| 1 | 1.317 | 0.126 | 0.000 | 4624 | 17873 | 0.438 |
| 2 | 1.405 | 0.049 | 0.999 | 2174873 | 4050027 | 99.268 |
| 3 | 1.700 | 0.045 | 6.315 | 1309 | 2157 | 0.053 |
| 4 | 1.795 | 0.052 | 1.960 | 1584 | 3148 | 0.077 |
| 5 | 1.892 | 0.041 | 2.099 | 1749 | 2600 | 0.064 |
| 6 | 2.164 | 0.069 | 4.957 | 1614 | 4085 | 0.100 |
| Total | | | | 2185753 | 4079890 | 100.000 |

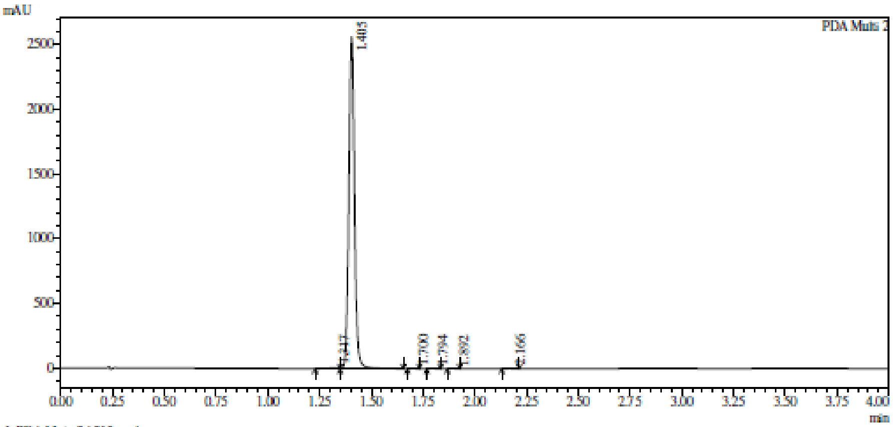

Integration result

PeakTable

PDA Ch2 215nm

| Peak# | Ret. Time | USP Width | Resolution | Height | Area | Area % |
|---|---|---|---|---|---|---|
| 1 | 1.317 | 0.117 | 0.000 | 5693 | 21767 | 0.451 |
| 2 | 1.405 | 0.049 | 1.062 | 2563316 | 4790990 | 99.355 |
| 3 | 1.700 | 0.045 | 6.302 | 988 | 1627 | 0.034 |
| 4 | 1.794 | 0.050 | 1.977 | 1090 | 1938 | 0.040 |
| 5 | 1.892 | 0.040 | 2.172 | 1766 | 2558 | 0.053 |
| 6 | 2.166 | 0.065 | 5.216 | 1329 | 3189 | 0.066 |
| Total | | | | 2574182 | 4822069 | 100.000 |

**HRMS** (2R,3R,4S,5S)-2-(6-aminopurin-9-yl)-5-[(6-methyl-1,3-benzothiazol -2-yl)sulfanylmethyl] tetrahydrofuran-3,4-diol **(2)**

**Compound Table**

| Label | Tgt Score | Mass Error (ppm) | Tgt Formula | Obs. RT | Ref. Mass | Obs. Mass |
|---|---|---|---|---|---|---|
| Cpd 1: C18 H18 N6 O3 S2; 1.452 | 97.44 | 1.66 | C18 H18 N6 O3 S2 | 1.452 | 430.0882 | 430.0889 |

| Obs. *m/z* | Obs. RT | Obs. Mass | Tgt Formula | Tgt Mass | Tgt Mass Error (ppm) | Find Cpds Algorithm |
|---|---|---|---|---|---|---|
| 216.0518 | 1.452 | 430.0889 | C18 H18 N6 O3 S2 | 430.0882 | 1.66 | Find by Formula |

**Compound Chromatograms**

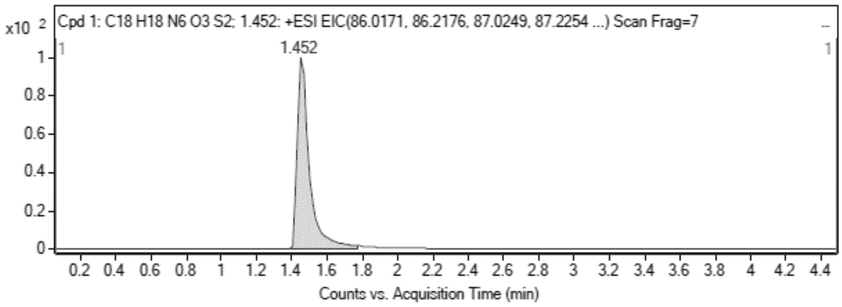

MS Zoomed Spectrum

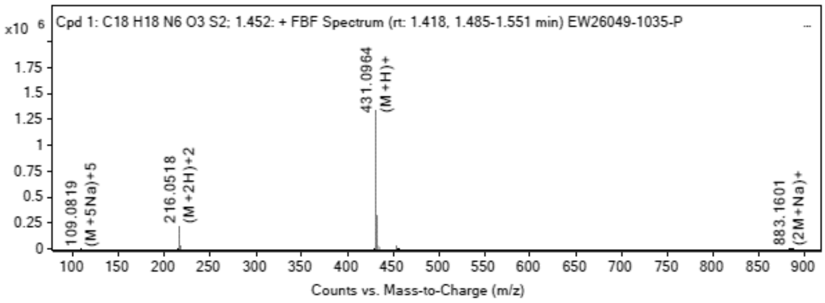

**MS Spectrum Peak List**

| Obs. *m/z* | Charge | Abund | Ion/Isotope |
|---|---|---|---|
| 216.0518 | 2 | 220369.45 | (M+2H)+2 |
| 216.553 | 2 | 53893.43 | (M+2H)+2 |
| 217.0509 | 2 | 26690.65 | (M+2H)+2 |
| 431.0964 | 1 | 1321847.88 | (M+H)+ |
| 432.0981 | 1 | 325651.41 | (M+H)+ |
| 433.0936 | 1 | 158972.36 | (M+H)+ |
| 453.0764 | 1 | 25026.08 | (M+Na)+ |
| 454.0789 | 1 | 6208.18 | (M+Na)+ |
| 455.0758 | 1 | 3294.22 | (M+Na)+ |
| 883.1601 | 1 | 1169.77 | (2M+Na)+ |

MS Zoomed Spectrum

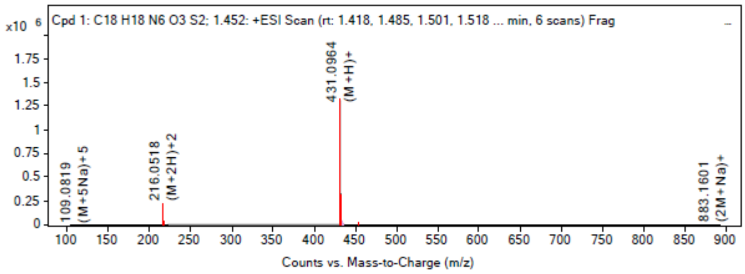

**MS Spectrum Peak List**

| Obs. *m/z* | Charge | Abund | Ion/Isotope | Tgt Mass Error (ppm) |
|---|---|---|---|---|
| 216.0518 | 2 | 220369.45 | (M+2H)+2 | -1.9 |
| 216.553 | 2 | 53893.43 | (M+2H)+2 | -1.66 |
| 217.0509 | 2 | 26690.65 | (M+2H)+2 | -2.08 |
| 431.0964 | 1 | 1321847.88 | (M+H)+ | -2.29 |
| 432.0981 | 1 | 325651.41 | (M+H)+ | -0.16 |
| 433.0936 | 1 | 158972.36 | (M+H)+ | 0.18 |
| 453.0764 | 1 | 25026.08 | (M+Na)+ | 2.22 |
| 454.0789 | 1 | 6208.18 | (M+Na)+ | 2.23 |
| 455.0758 | 1 | 3294.22 | (M+Na)+ | -0.31 |
| 883.1601 | 1 | 1169.77 | (2M+Na)+ | 6.17 |

--- End Of Report ---

**[1]H NMR** 2R,3R,4S,5S)-2-(6-aminopurin-9-yl)-5-[(5-methyl-1,3-benzothiazol -2-yl)sulfanylmethyl] tetrahydrofuran-3,4-diol **(3)**

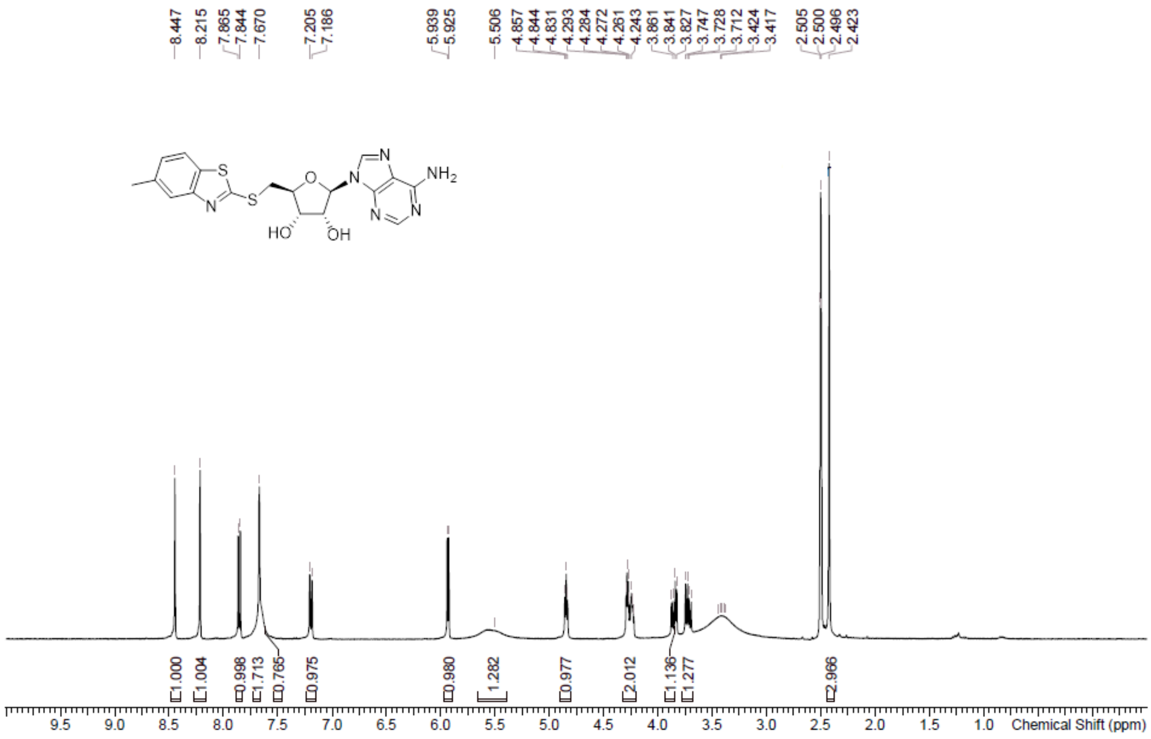

**HPLC** 2*R*,3*R*,4*S*,5*S*)-2-(6-aminopurin-9-yl)-5-[(5-methyl-1,3-benzothiazol -2-yl)sulfanylmethyl] tetrahydrofuran-3,4-diol **(3)**

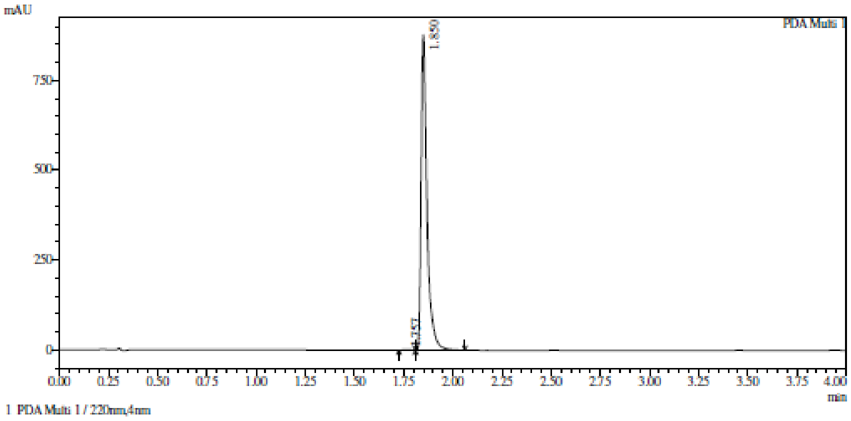

Integration result

PeakTable

PDA Ch1 220nm

| Peak# | Ret. Time | USP Width | Resolution | Height | Area | Area % |
|---|---|---|---|---|---|---|
| 1 | 1.757 | 0.077 | 0.000 | 1117 | 3310 | 0.180 |
| 2 | 1.850 | 0.051 | 1.447 | 877291 | 1840677 | 99.820 |
| Total | | | | 878408 | 1843987 | 100.000 |

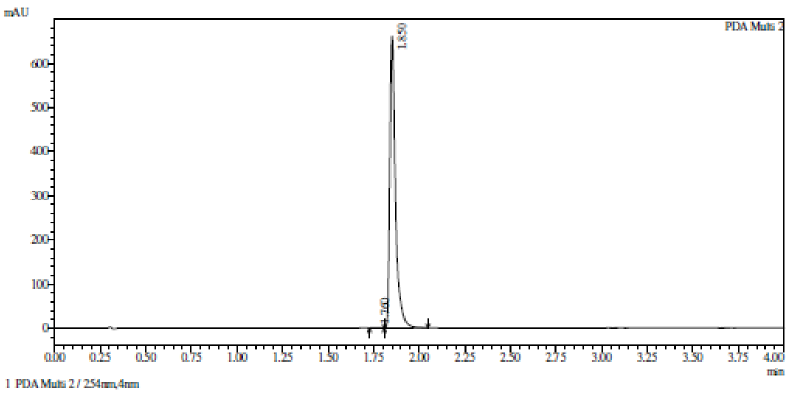

Integration result

PeakTable

PDA Ch2 254nm

| Peak# | Ret. Time | USP Width | Resolution | Height | Area | Area % |
|---|---|---|---|---|---|---|
| 1 | 1.760 | 0.073 | 0.000 | 830 | 2335 | 0.167 |
| 2 | 1.850 | 0.051 | 1.451 | 662072 | 1395294 | 99.833 |
| Total | | | | 662902 | 1397629 | 100.000 |

**HPLC** 2*R*,3*R*,4*S*,5*S*)-2-(6-aminopurin-9-yl)-5-[(5-methyl-1,3-benzothiazol -2-yl)sulfanylmethyl] tetrahydrofuran-3,4-diol **(3)**

**Compound Table**

| Label | Tgt Score | Mass Error (ppm) | Tgt Formula | Obs. RT | Ref. Mass | Obs. Mass |
|---|---|---|---|---|---|---|
| Cpd 1: C18 H18 N6 O3 S2; 1.455 | 97.84 | 4.46 | C18 H18 N6 O3 S2 | 1.455 | 430.0882 | 430.0901 |

| Obs. *m/z* | Obs. RT | Obs. Mass | Tgt Formula | Tgt Mass | Tgt Mass Error (ppm) | Find Cpds Algorithm |
|---|---|---|---|---|---|---|
| 453.0779 | 1.455 | 430.0901 | C18 H18 N6 O3 S2 | 430.0882 | 4.46 | Find by Formula |

**Compound Chromatograms**

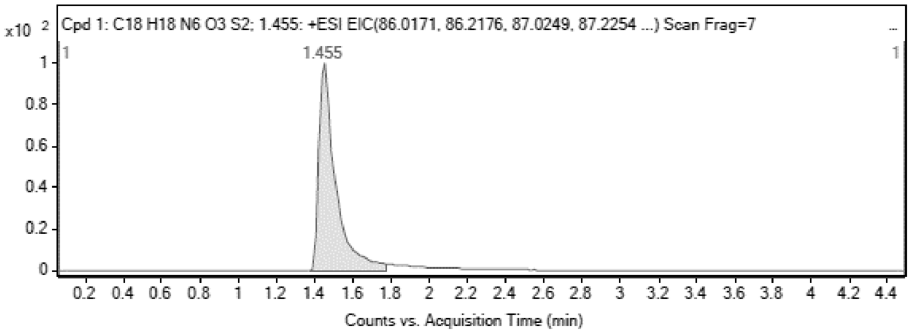

**MS Zoomed Spectrum**

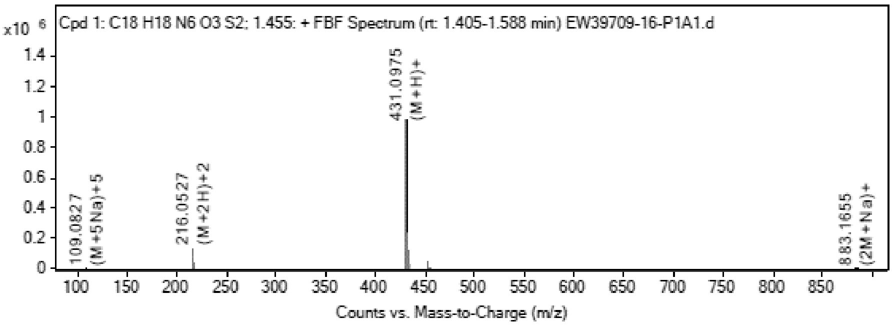

**MS Spectrum Peak List**

| Obs. *m/z* | Charge | Abund | Ion/Isotope |
|---|---|---|---|
| 216.0527 | 2 | 131618.92 | (M+2H)+2 |
| 216.5539 | 2 | 31740.88 | (M+2H)+2 |

| | | | |
|---|---|---|---|
| 217.0517 | 2 | 16412.5 | (M+2H)+2 |
| 431.0975 | 1 | 981331.69 | (M+H)+ |
| 432.0993 | 1 | 237460.64 | (M+H)+ |
| 433.0948 | 1 | 115323.02 | (M+H)+ |
| 453.0779 | 1 | 46647.75 | (M+Na)+ |
| 454.0804 | 1 | 11184.8 | (M+Na)+ |
| 455.0761 | 1 | 5599.68 | (M+Na)+ |
| 883.1655 | 1 | 4335.82 | (2M+Na)+ |

MS Zoomed Spectrum

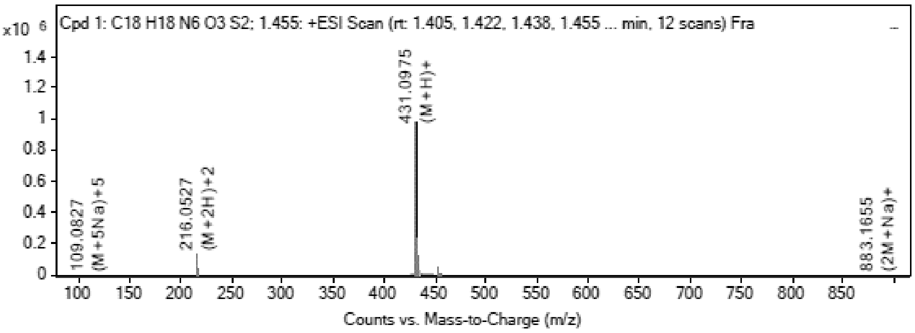

MS Spectrum Peak List

| Obs. m/z | Charge | Abund | Ion/Isotope | Tgt Mass Error (ppm) |
|---|---|---|---|---|
| 216.0527 | 2 | 131618.92 | (M+2H)+2 | -6.37 |
| 216.5539 | 2 | 31740.88 | (M+2H)+2 | -5.74 |
| 217.0517 | 2 | 16412.5 | (M+2H)+2 | -5.75 |
| 431.0975 | 1 | 981331.69 | (M+H)+ | -4.85 |
| 432.0993 | 1 | 237460.64 | (M+H)+ | -2.93 |
| 433.0948 | 1 | 115323.02 | (M+H)+ | -2.45 |
| 453.0779 | 1 | 46647.75 | (M+Na)+ | -1.08 |
| 454.0804 | 1 | 11184.8 | (M+Na)+ | -1.02 |
| 455.0761 | 1 | 5599.68 | (M+Na)+ | -0.94 |
| 883.1655 | 1 | 4335.82 | (2M+Na)+ | 0.07 |

--- End Of Report ---

**$^1$H NMR** (2*R*,3*R*,4*S*,5*S*)-2-(6-aminopurin-9-yl)-5-(2-quinolylsulfanylmethyl) tetrahydrofuran-3,4-diol **(7)**

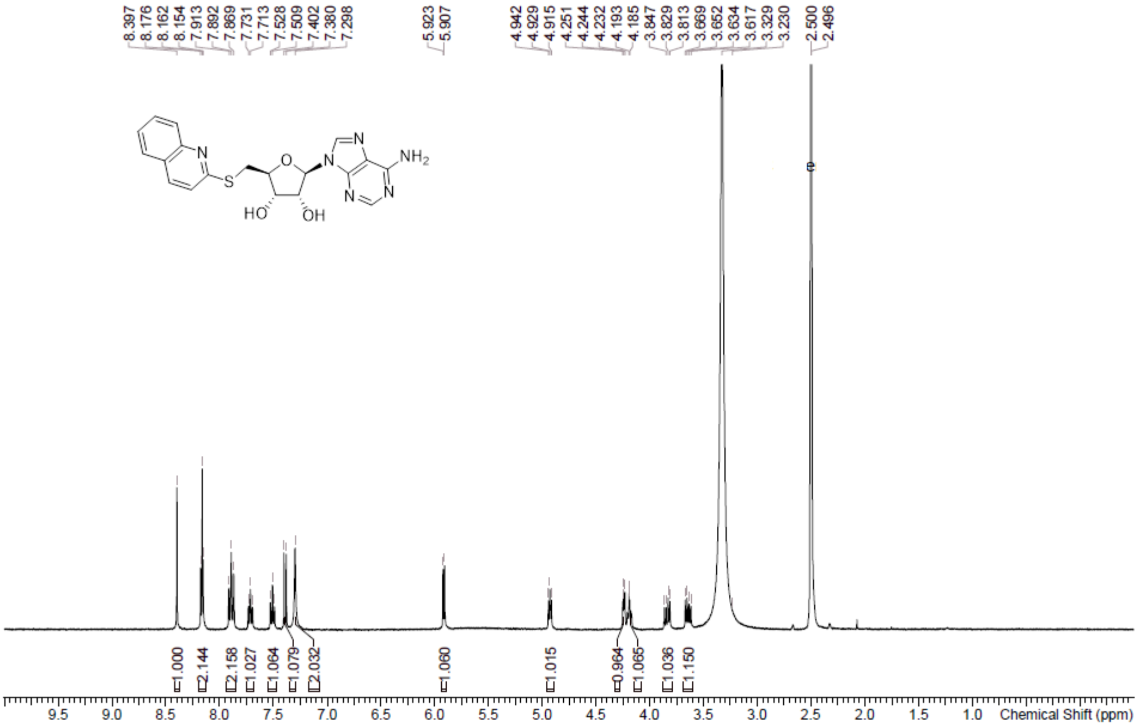

**HPLC** (2*R*,3*R*,4*S*,5*S*)-2-(6-aminopurin-9-yl)-5-(2-quinolylsulfanylmethyl) tetrahydrofuran-3,4-diol **(7)**

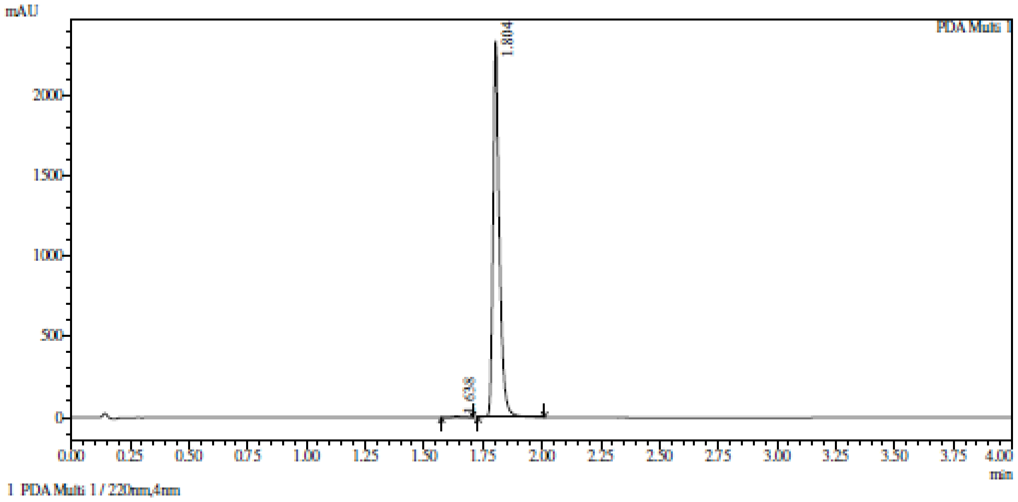

Integration result

PeakTable

PDA Ch1 220nm

| Peak# | Ret. Time | USP Width | Resolution | Height | Area | Area % |
|---|---|---|---|---|---|---|
| 1 | 1.638 | 0.073 | 0.000 | 5837 | 16239 | 0.370 |
| 2 | 1.804 | 0.047 | 2.782 | 2342534 | 4378211 | 99.630 |
| Total | | | | 2348370 | 4394450 | 100.000 |

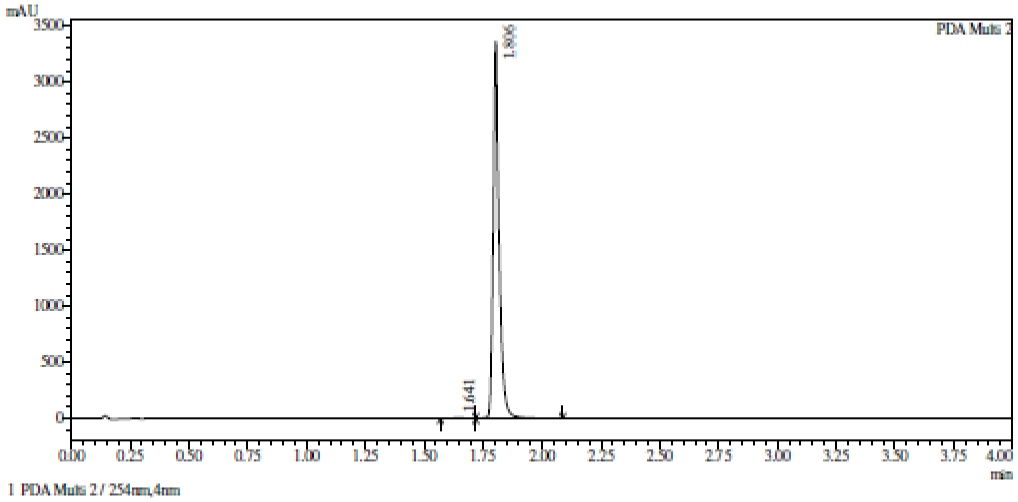

Integration result

PeakTable

PDA Ch2 254nm

| Peak# | Ret. Time | USP Width | Resolution | Height | Area | Area % |
|---|---|---|---|---|---|---|
| 1 | 1.641 | 0.139 | 0.000 | 5120 | 23930 | 0.406 |
| 2 | 1.806 | 0.042 | 1.820 | 3360897 | 5864740 | 99.594 |
| Total | | | | 3366017 | 5888670 | 100.000 |

**HRMS** (2*R*,3*R*,4*S*,5*S*)-2-(6-aminopurin-9-yl)-5-(2-quinolylsulfanylmethyl) tetrahydrofuran-3,4-diol **(7)**

**Compound Table**

| Label | Tgt Score | Mass Error (ppm) | Tgt Formula | Obs. RT | Ref. Mass | Obs. Mass |
|---|---|---|---|---|---|---|
| Cpd 1: C19 H18 N6 O3 S; 1.659 | 99.01 | 2.99 | C19 H18 N6 O3 S | 1.659 | 410.1161 | 410.1173 |

| Obs. *m/z* | Obs. RT | Obs. Mass | Tgt Formula | Tgt Mass | Tgt Mass Error (ppm) | Find Cpds Algorithm |
|---|---|---|---|---|---|---|
| 433.1057 | 1.659 | 410.1173 | C19 H18 N6 O3 S | 410.1161 | 2.99 | Find by Formula |

**Compound Chromatograms**

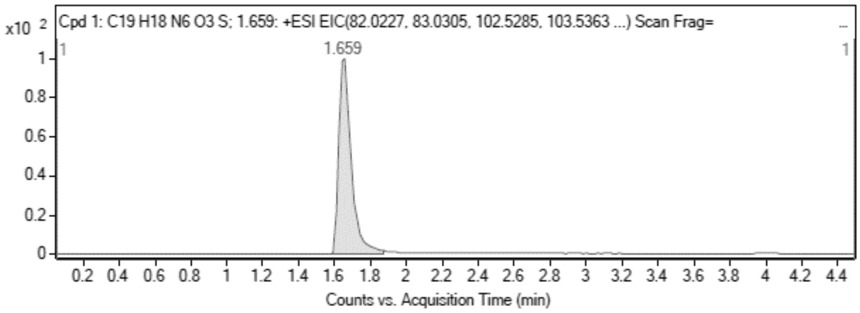

MS Zoomed Spectrum

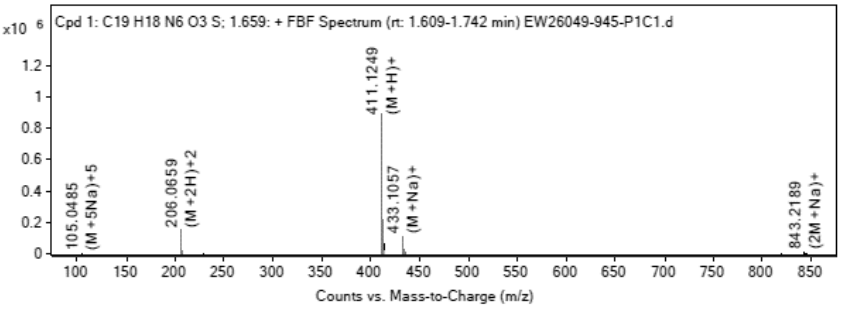

**MS Spectrum Peak List**

| Obs. *m/z* | Charge | Abund | Ion/Isotope |
|---|---|---|---|
| 206.0659 | 2 | 152511.06 | (M+2H)+2 |
| 206.5671 | 2 | 37882.84 | (M+2H)+2 |
| 207.0661 | 2 | 12751.92 | (M+2H)+2 |
| 411.1249 | 1 | 894294.31 | (M+H)+ |
| 412.1269 | 1 | 214500.81 | (M+H)+ |
| 413.1242 | 1 | 65846.22 | (M+H)+ |
| 433.1057 | 1 | 111117.71 | (M+Na)+ |
| 434.1083 | 1 | 25837.89 | (M+Na)+ |
| 435.1059 | 1 | 8540.43 | (M+Na)+ |
| 843.2189 | 1 | 7718.13 | (2M+Na)+ |

MS Zoomed Spectrum

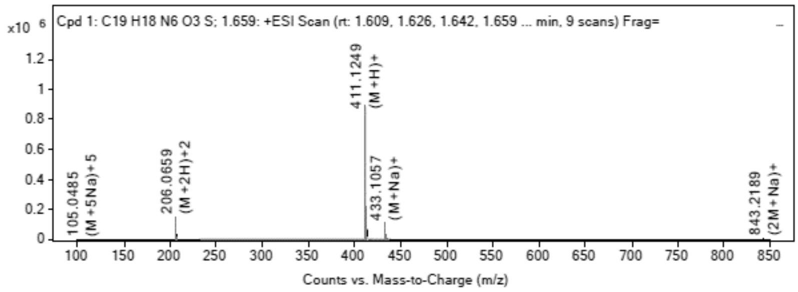

**MS Spectrum Peak List**

| Obs. *m/z* | Charge | Abund | Ion/Isotope | Tgt Mass Error (ppm) |
|---|---|---|---|---|
| 206.0659 | 2 | 152511.06 | (M+2H)+2 | -2.54 |
| 206.5671 | 2 | 37882.84 | (M+2H)+2 | -2.14 |
| 207.0661 | 2 | 12751.92 | (M+2H)+2 | -4.07 |
| 411.1249 | 1 | 894294.31 | (M+H)+ | -3.68 |
| 412.1269 | 1 | 214500.81 | (M+H)+ | -2.04 |
| 413.1242 | 1 | 65846.22 | (M+H)+ | -2.58 |
| 433.1057 | 1 | 111117.71 | (M+Na)+ | -0.9 |
| 434.1083 | 1 | 25837.89 | (M+Na)+ | -0.65 |
| 435.1059 | 1 | 8540.43 | (M+Na)+ | -1.88 |
| 843.2189 | 1 | 7718.13 | (2M+Na)+ | 3.03 |

--- End Of Report ---

**$^{1}$H NMR** (2*R*,3*R*,4*S*,5*S*)-2-(6-Aminopurin-9-yl)-5-[(6-methyl-2-quinolyl) sulfanylmethyl] tetrahydrofuran-3,4-diol **(8)**

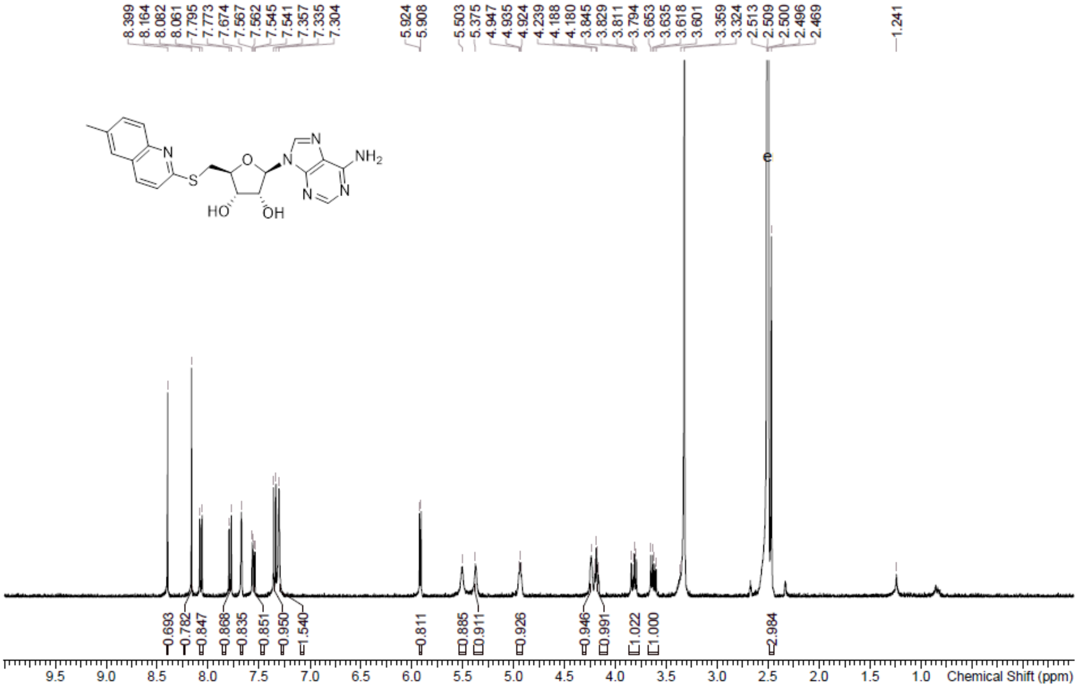

**HPLC** (2*R*,3*R*,4*S*,5*S*)-2-(6-Aminopurin-9-yl)-5-[(6-methyl-2-quinolyl) sulfanylmethyl] tetrahydrofuran-3,4-diol **(8)**

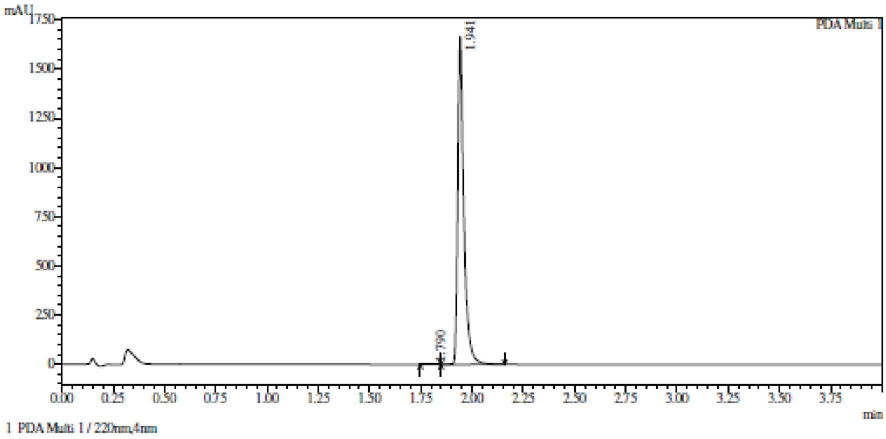

1 PDA Multi 1 / 220nm,4nm

Integration result

PeakTable

PDA Ch1 220nm

| Peak# | Ret. Time | USP Width | Resolution | Height | Area | Area % |
|---|---|---|---|---|---|---|
| 1 | 1.790 | 0.087 | 0.000 | 1481 | 4733 | 0.137 |
| 2 | 1.941 | 0.052 | 2.190 | 1668135 | 3439618 | 99.863 |
| Total | | | | 1669616 | 3444350 | 100.000 |

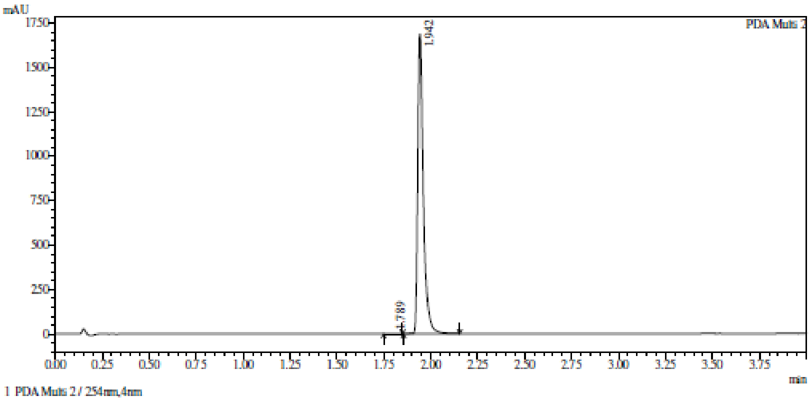

Integration result

PeakTable

PDA Ch2 254nm

| Peak# | Ret. Time | USP Width | Resolution | Height | Area | Area % |
|---|---|---|---|---|---|---|
| 1 | 1.789 | 0.087 | 0.000 | 1574 | 4811 | 0.140 |
| 2 | 1.942 | 0.051 | 2.211 | 1688326 | 3438317 | 99.860 |
| Total | | | | 1689900 | 3443128 | 100.000 |

**HRMS** (2*R*,3*R*,4*S*,5*S*)-2-(6-Aminopurin-9-yl)-5-[(6-methyl-2-quinolyl) sulfanylmethyl] tetrahydrofuran-3,4-diol **(8)**

**Compound Table**

| Label | Tgt Score | Mass Error (ppm) | Tgt Formula | Obs. RT | Ref. Mass | Obs. Mass |
|---|---|---|---|---|---|---|
| Cpd 1: C20 H20 N6 O3 S; 1.299 | 96.21 | 5.97 | C20 H20 N6 O3 S | 1.299 | 424.1318 | 424.1343 |

| Obs. *m/z* | Obs. RT | Obs. Mass | Tgt Formula | Tgt Mass | Tgt Mass Error (ppm) | Find Cpds Algorithm |
|---|---|---|---|---|---|---|
| 447.1219 | 1.299 | 424.1343 | C20 H20 N6 O3 S | 424.1318 | 5.97 | Find by Formula |

**Compound Chromatograms**

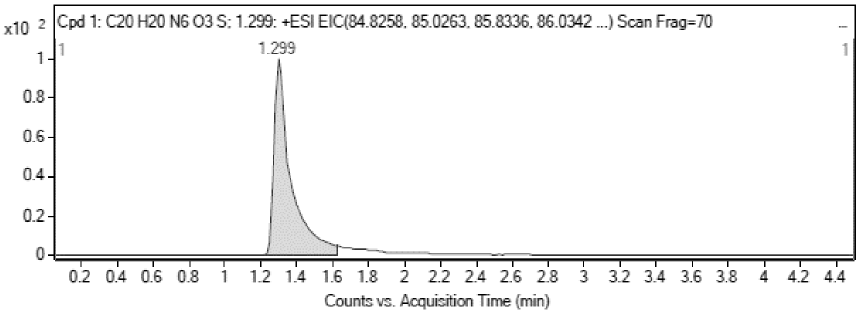

**MS Zoomed Spectrum**

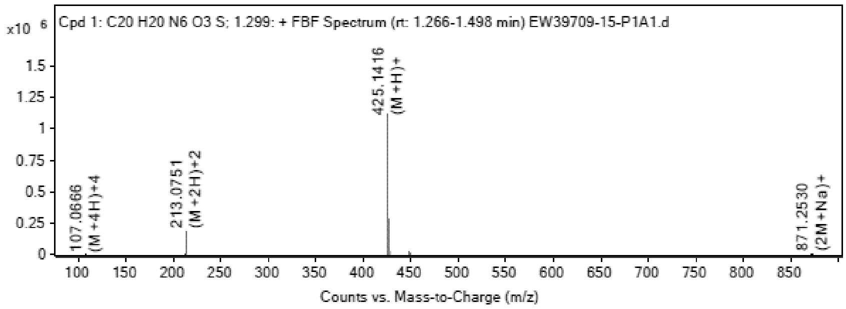

**MS Spectrum Peak List**

| Obs. *m/z* | Charge | Abund | Ion/Isotope |
|---|---|---|---|
| 213.0751 | 2 | 186816.16 | (M+2H)+2 |
| 213.5762 | 2 | 47819.52 | (M+2H)+2 |

| | | | |
|---|---|---|---|
| 214.0748 | 2 | 15425.82 | (M+2H)+2 |
| 425.1416 | 1 | 1121128.88 | (M+H)+ |
| 426.1435 | 1 | 287162.13 | (M+H)+ |
| 427.1407 | 1 | 88244.41 | (M+H)+ |
| 447.1219 | 1 | 20542.54 | (M+Na)+ |
| 448.1245 | 1 | 5157.79 | (M+Na)+ |
| 449.1234 | 1 | 1794.27 | (M+Na)+ |
| 871.253 | 1 | 1756.87 | (2M+Na)+ |

MS Zoomed Spectrum

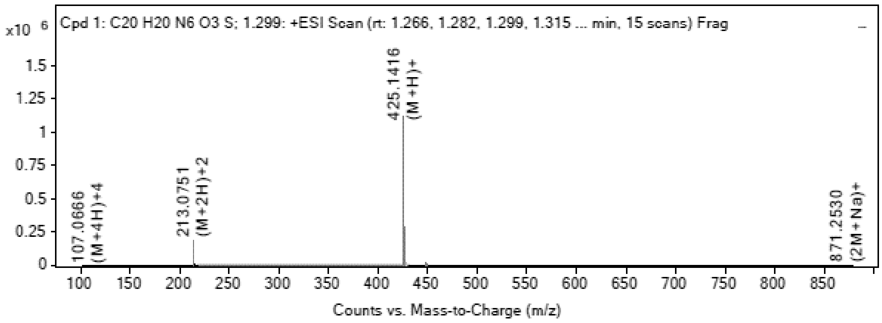

MS Spectrum Peak List

| Obs. m/z | Charge | Abund | Ion/Isotope | Tgt Mass Error (ppm) |
|---|---|---|---|---|
| 213.0751 | 2 | 186816.16 | (M+2H)+2 | -8.98 |
| 213.5762 | 2 | 47819.52 | (M+2H)+2 | -7.74 |
| 214.0748 | 2 | 15425.82 | (M+2H)+2 | -7.85 |
| 425.1416 | 1 | 1121128.88 | (M+H)+ | -6.06 |
| 426.1435 | 1 | 287162.13 | (M+H)+ | -4.13 |
| 427.1407 | 1 | 88244.41 | (M+H)+ | -3.86 |
| 447.1219 | 1 | 20542.54 | (M+Na)+ | -2.12 |
| 448.1245 | 1 | 5157.79 | (M+Na)+ | -1.78 |
| 449.1234 | 1 | 1794.27 | (M+Na)+ | -5.35 |
| 871.253 | 1 | 1756.87 | (2M+Na)+ | -0.26 |

--- End Of Report ---

**$^1$H NMR** (2*R*,3*R*,4*S*,5*S*)-2-(6-Aminopurin-9-yl)-5-[(7-methyl-2-quinolyl) sulfanylmethyl] tetrahydrofuran-3,4-diol **(9)**

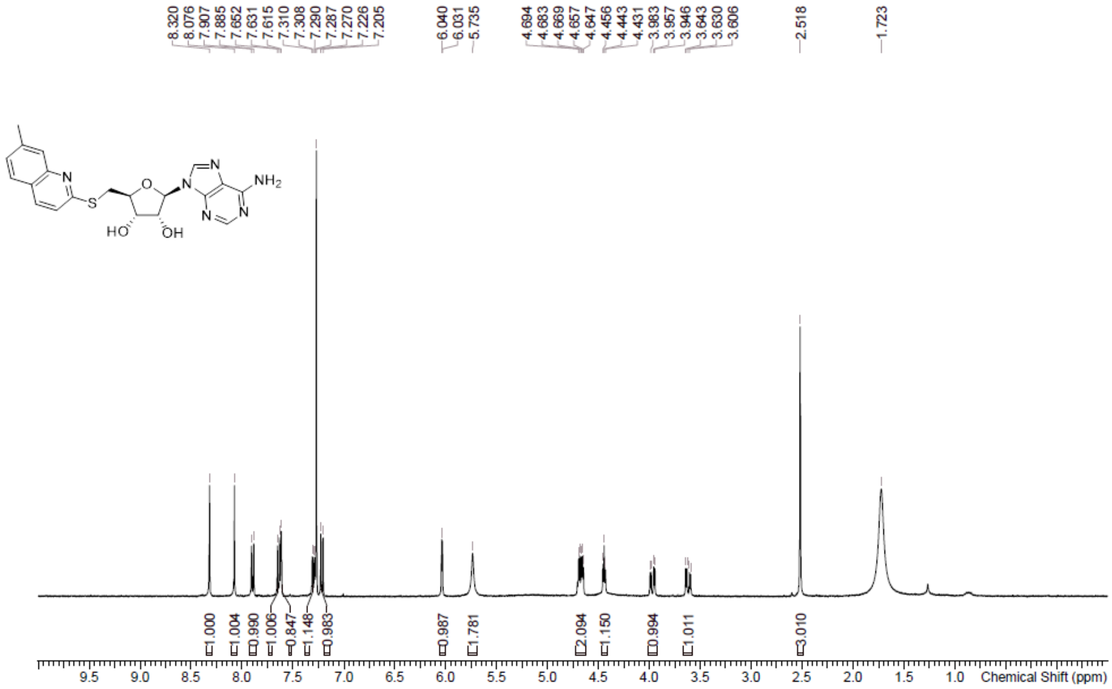

**HPLC** (2*R*,3*R*,4*S*,5*S*)-2-(6-Aminopurin-9-yl)-5-[(7-methyl-2-quinolyl) sulfanylmethyl] tetrahydrofuran-3,4-diol **(9)**

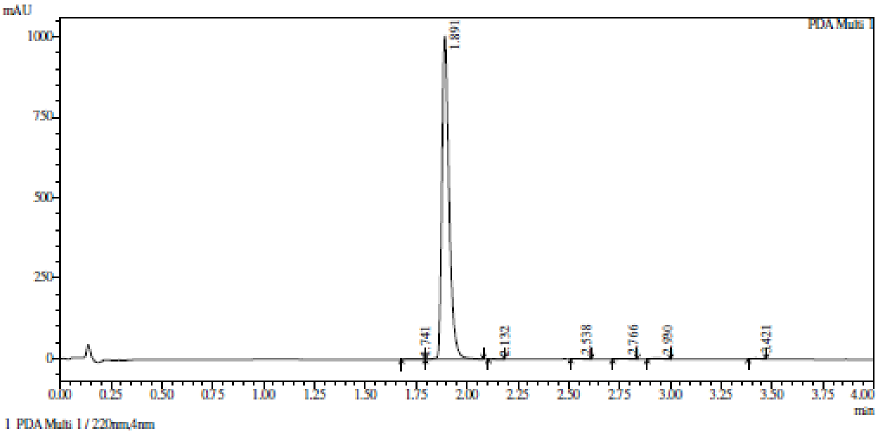

Integration result

PeakTable

PDA Ch1 220nm

| Peak# | Ret. Time | USP Width | Resolution | Height | Area | Area % |
|---|---|---|---|---|---|---|
| 1 | 1.741 | 0.117 | 0.000 | 1219 | 5083 | 0.206 |
| 2 | 1.891 | 0.063 | 1.672 | 1003302 | 2440310 | 99.011 |
| 3 | 2.132 | 0.055 | 4.099 | 803 | 1649 | 0.067 |
| 4 | 2.538 | 0.076 | 6.182 | 688 | 1950 | 0.079 |
| 5 | 2.766 | 0.107 | 2.495 | 1213 | 4380 | 0.178 |
| 6 | 2.930 | 0.080 | 1.764 | 2718 | 7986 | 0.324 |
| 7 | 3.421 | 0.064 | 6.814 | 1339 | 3335 | 0.135 |
| Total | | | | 1011282 | 2464694 | 100.000 |

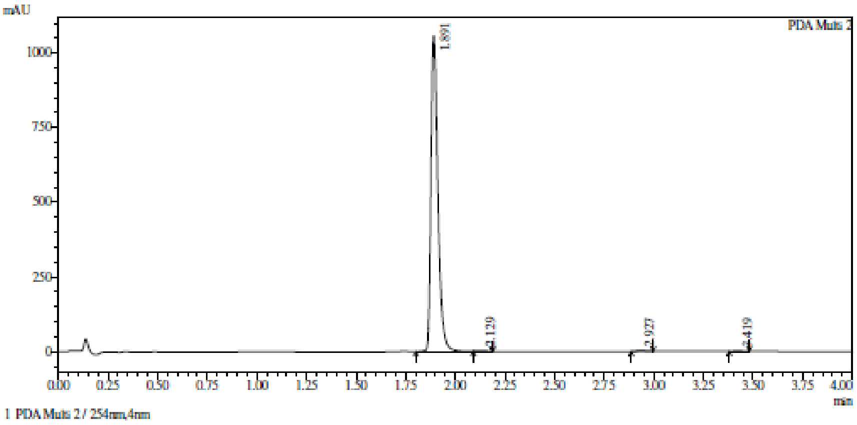

Integration result

PeakTable

PDA Ch2 254nm

| Peak# | Ret. Time | USP Width | Resolution | Height | Area | Area % |
|---|---|---|---|---|---|---|
| 1 | 1.891 | 0.062 | 0.000 | 1057931 | 2545001 | 99.417 |
| 2 | 2.129 | 0.058 | 3.965 | 893 | 1942 | 0.076 |
| 3 | 2.927 | 0.079 | 11.690 | 2620 | 7715 | 0.301 |
| 4 | 3.419 | 0.070 | 6.602 | 1987 | 5268 | 0.206 |
| Total | | | | 1063431 | 2559927 | 100.000 |

**HRMS** (2*R*,3*R*,4*S*,5*S*)-2-(6-Aminopurin-9-yl)-5-[(7-methyl-2-quinolyl) sulfanylmethyl] tetrahydrofuran-3,4-diol **(9)**

**Compound Table**

| Label | Tgt Score | Mass Error (ppm) | Tgt Formula | Obs. RT | Ref. Mass | Obs. Mass |
|---|---|---|---|---|---|---|
| Cpd 1: C20 H20 N6 O3 S; 4.174 | 97.42 | -11.9 | C20 H20 N6 O3 S | 4.174 | 424.1318 | 424.1267 |

| Obs. *m/z* | Obs. RT | Obs. Mass | Tgt Formula | Tgt Mass | Tgt Mass Error (ppm) | Find Cpds Algorithm |
|---|---|---|---|---|---|---|
| 425.1391 | 4.174 | 424.1267 | C20 H20 N6 O3 S | 424.1318 | -11.9 | Find by Formula |

**Compound Chromatograms**

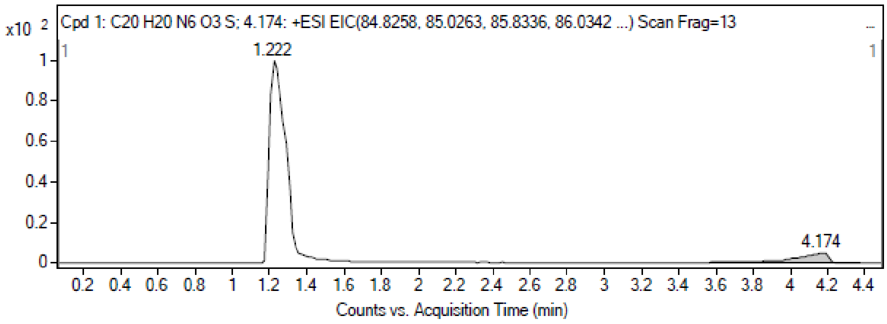

MS Zoomed Spectrum

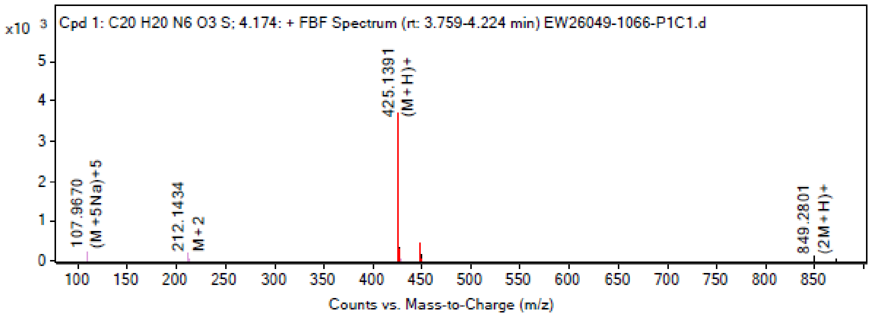

**MS Spectrum Peak List**

| Obs. *m/z* | Charge | Abund | Ion/Isotope |
|---|---|---|---|
| 107.967 | 5 | 240.43 | (M+5Na)+5 |
| 212.1434 | 2 | 163.47 | M+2 |
| 425.1391 | 1 | 3628.15 | (M+H)+ |
| 426.1412 | 1 | 989.4 | (M+H)+ |
| 427.14 | 1 | 342.28 | (M+H)+ |
| 447.1164 | 1 | 408.88 | (M+Na)+ |
| 448.1233 | 1 | 158.47 | (M+Na)+ |
| 848.2738 | 1 | 102.45 | 2M+ |
| 849.2801 | 1 | 114.34 | (2M+H)+ |
| 871.2689 | 1 | 67.29 | (2M+Na)+ |

MS Zoomed Spectrum

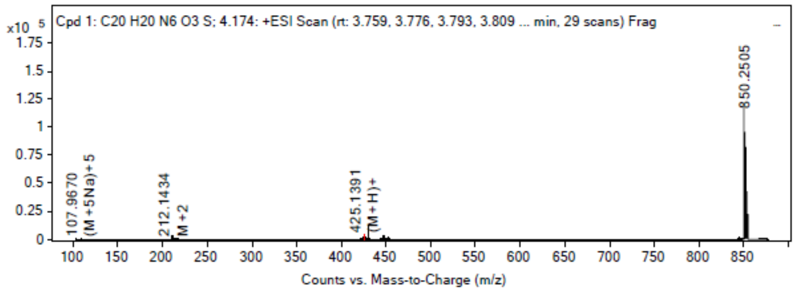

**MS Spectrum Peak List**

| Obs. *m/z* | Charge | Abund | Ion/Isotope | Tgt Mass Error (ppm) |
|---|---|---|---|---|
| 107.967 | 5 | 240.43 | (M+5Na)+5 | 454.94 |
| 212.1434 | 2 | 163.47 | M+2 | -368.17 |
| 425.1391 | 1 | 3628.15 | (M+H)+ | -0.11 |
| 426.1412 | 1 | 989.4 | (M+H)+ | 1.22 |
| 427.14 | 1 | 342.28 | (M+H)+ | -2.29 |
| 447.1164 | 1 | 408.88 | (M+Na)+ | 10.15 |
| 448.1233 | 1 | 158.47 | (M+Na)+ | 0.95 |
| 848.2738 | 1 | 102.45 | 2M+ | -12.73 |
| 849.2801 | 1 | 114.34 | (2M+H)+ | -10.95 |
| 871.2689 | 1 | 67.29 | (2M+Na)+ | -18.5 |

--- End Of Report ---

**[1]H NMR** (2*R*,3*R*,4*S*,5*S*)-2-(6-Aminopurin-9-yl)-5-(7-quinolylsulfanylmethyl) tetrahydrofuran-3,4-diol **(10)**

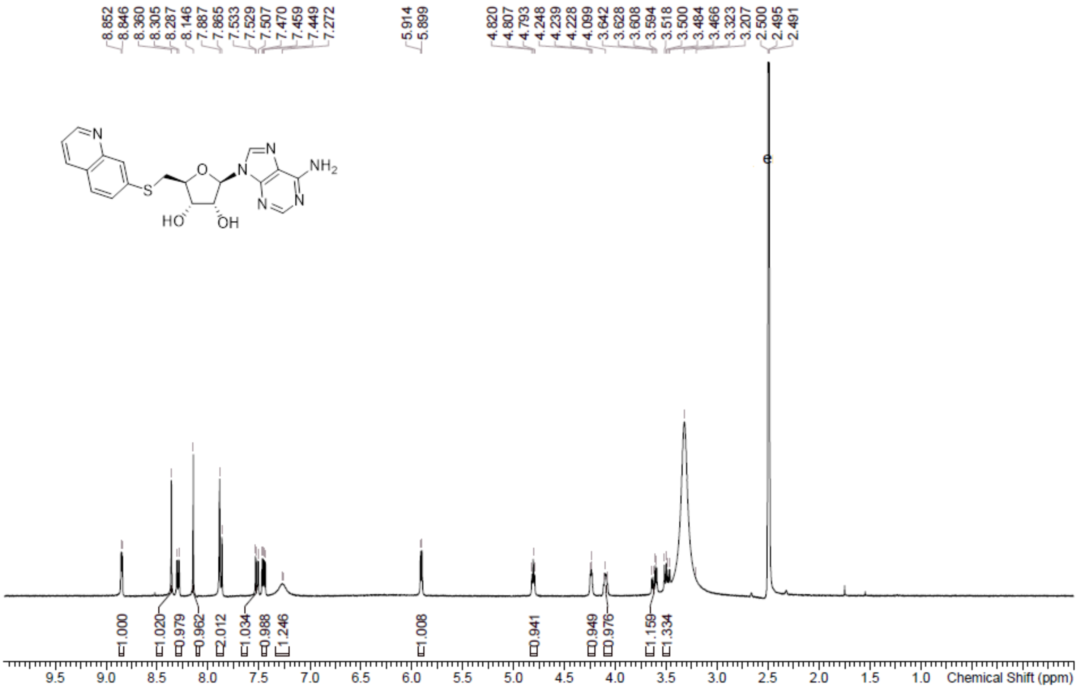

**HPLC** (2*R*,3*R*,4*S*,5*S*)-2-(6-Aminopurin-9-yl)-5-(7-quinolylsulfanylmethyl) tetrahydrofuran-3,4-diol **(10)**

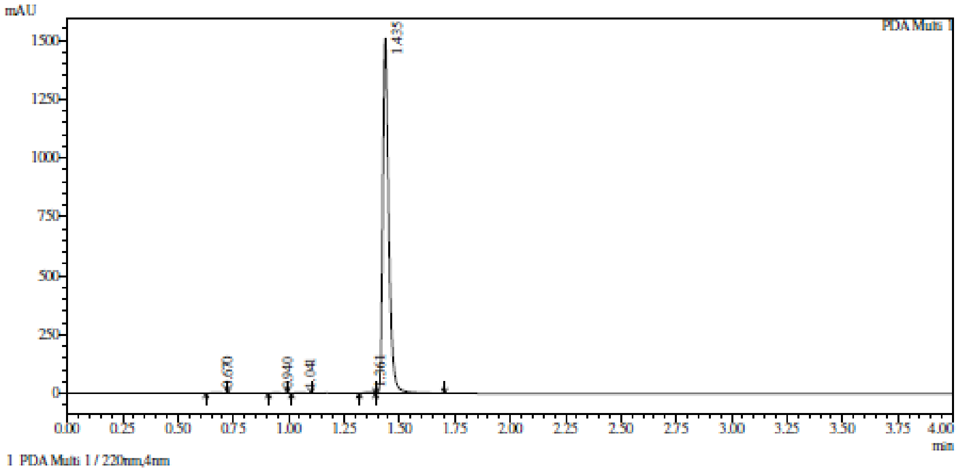

1 PDA Multi 1 / 220nm,4nm

Integration result

PeakTable

PDA Ch1 220nm

| Peak# | Ret. Time | USP Width | Resolution | Height | Area | Area % |
|---|---|---|---|---|---|---|
| 1 | 0.670 | 0.072 | 0.000 | 2617 | 6767 | 0.237 |
| 2 | 0.940 | 0.067 | 3.905 | 1108 | 2773 | 0.097 |
| 3 | 1.041 | 0.059 | 1.598 | 2104 | 4841 | 0.169 |
| 4 | 1.361 | 0.093 | 4.222 | 4233 | 12102 | 0.424 |
| 5 | 1.435 | 0.049 | 1.049 | 1506546 | 2830566 | 99.073 |
| Total | | | | 1516608 | 2857049 | 100.000 |

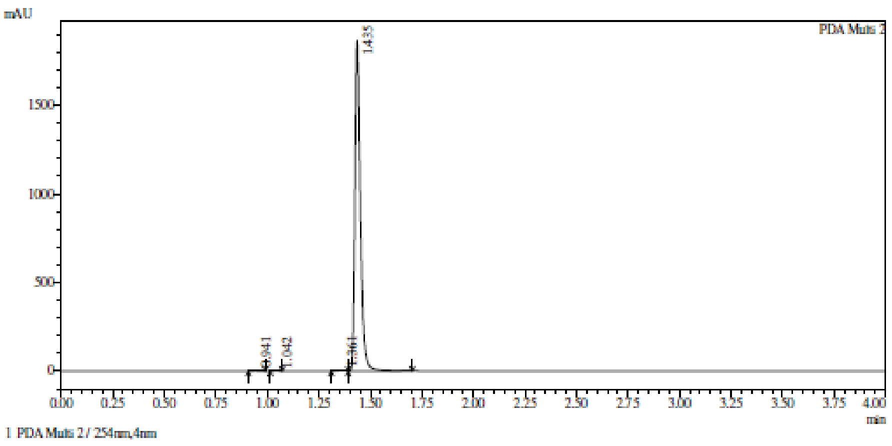

Integration result

PeakTable

PDA Ch2 254nm

| Peak# | Ret. Time | USP Width | Resolution | Height | Area | Area % |
|---|---|---|---|---|---|---|
| 1 | 0.941 | 0.068 | 0.000 | 664 | 1662 | 0.047 |
| 2 | 1.042 | 0.050 | 1.698 | 875 | 1539 | 0.044 |
| 3 | 1.361 | 0.093 | 4.481 | 4539 | 12849 | 0.367 |
| 4 | 1.435 | 0.049 | 1.049 | 1867227 | 3483963 | 99.541 |
| Total | | | | 1873304 | 3500014 | 100.000 |

**HRMS** (2*R*,3*R*,4*S*,5*S*)-2-(6-Aminopurin-9-yl)-5-(7-quinolylsulfanylmethyl) tetrahydrofuran-3,4-diol **(10)**

Compound Table

| Label | Tgt Score | Mass Error (ppm) | Tgt Formula | Obs. RT | Ref. Mass | Obs. Mass |
|---|---|---|---|---|---|---|
| Cpd 1: C19 H18 N6 O3 S; 1.315 | 98.62 | 1.32 | C19 H18 N6 O3 S | 1.315 | 410.1161 | 410.1167 |

| Obs. m/z | Obs. RT | Obs. Mass | Tgt Formula | Tgt Mass | Tgt Mass Error (ppm) | Find Cpds Algorithm |
|---|---|---|---|---|---|---|
| 433.1046 | 1.315 | 410.1167 | C19 H18 N6 O3 S | 410.1161 | 1.32 | Find by Formula |

Compound Chromatograms

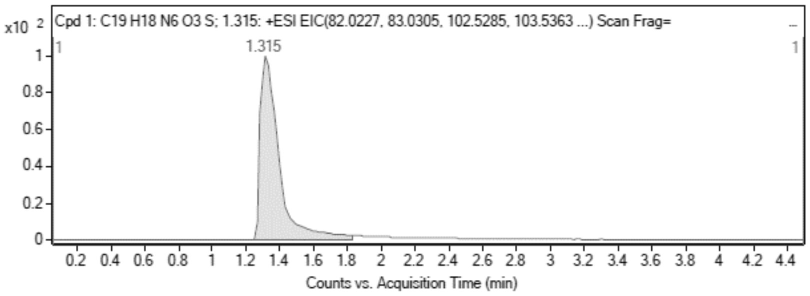

MS Zoomed Spectrum

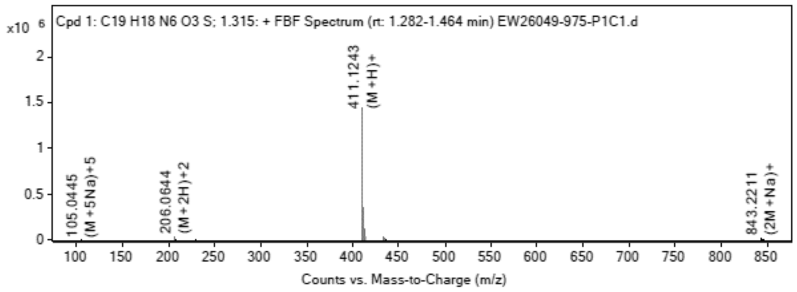

MS Spectrum Peak List

| Obs. m/z | Charge | Abund | Ion/Isotope |
|---|---|---|---|
| 206.0644 | 2 | 34937 | (M+2H)+2 |
| 206.5657 | 2 | 8542.68 | (M+2H)+2 |

| | | | |
|---|---|---|---|
| 411.1243 | 1 | 1438607.63 | (M+H)+ |
| 412.1261 | 1 | 356487.84 | (M+H)+ |
| 413.1232 | 1 | 108992.29 | (M+H)+ |
| 433.1046 | 1 | 34449.09 | (M+Na)+ |
| 434.1074 | 1 | 8316.24 | (M+Na)+ |
| 843.2211 | 1 | 18047.82 | (2M+Na)+ |
| 844.2238 | 1 | 8722.7 | (2M+Na)+ |
| 845.2147 | 1 | 5844.32 | (2M+Na)+ |

MS Zoomed Spectrum

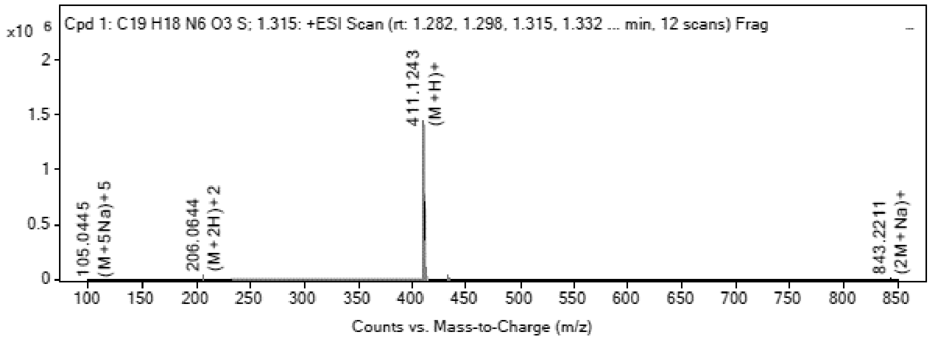

**MS Spectrum Peak List**

| Obs. *m/z* | Charge | Abund | Ion/Isotope | Tgt Mass Error (ppm) |
|---|---|---|---|---|
| 206.0644 | 2 | 34937 | (M+2H)+2 | 4.35 |
| 206.5657 | 2 | 8542.68 | (M+2H)+2 | 4.79 |
| 411.1243 | 1 | 1438607.63 | (M+H)+ | -2.11 |
| 412.1261 | 1 | 356487.84 | (M+H)+ | -0.17 |
| 413.1232 | 1 | 108992.29 | (M+H)+ | -0.01 |
| 433.1046 | 1 | 34449.09 | (M+Na)+ | 1.59 |
| 434.1074 | 1 | 8316.24 | (M+Na)+ | 1.33 |
| 843.2211 | 1 | 18047.82 | (2M+Na)+ | 0.36 |
| 844.2238 | 1 | 8722.7 | (2M+Na)+ | 0.35 |
| 845.2147 | 1 | 5844.32 | (2M+Na)+ | 9.43 |

--- End Of Report ---

$^{1}$**H NMR** (2*R*,3*R*,4*S*,5*S*)-2-(6-Aminopurin-9-yl)-5-(thiazol-2-ylsulfanylmethyl) tetrahydrofuran-3,4-diol **(11)**

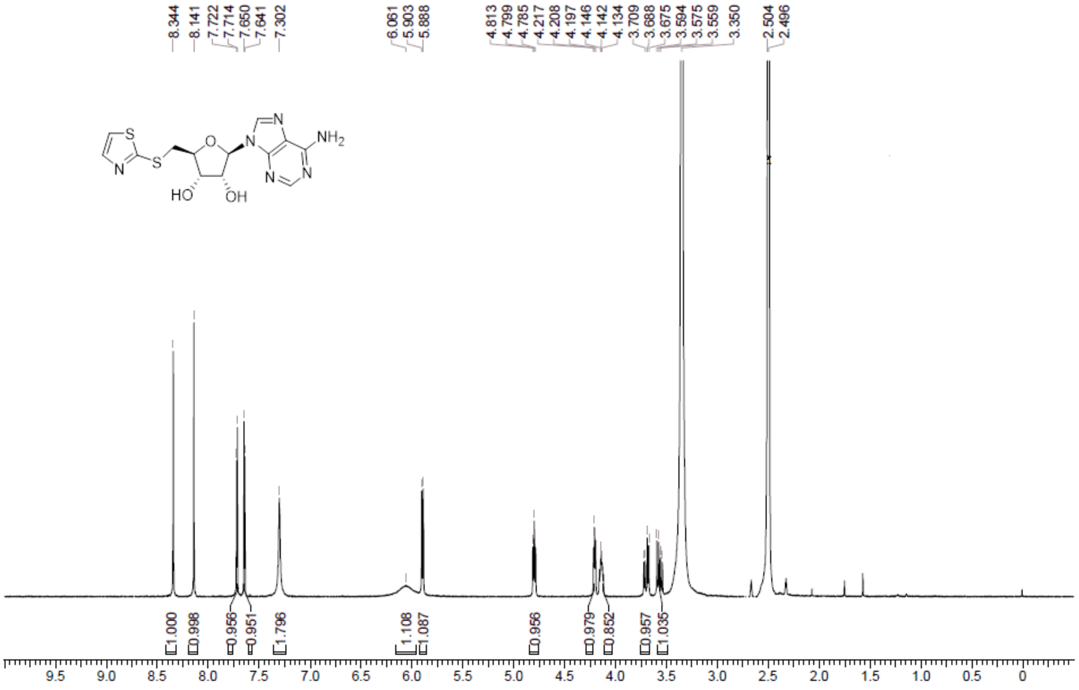

**HPLC** (2*R*,3*R*,4*S*,5*S*)-2-(6-Aminopurin-9-yl)-5-(thiazol-2-ylsulfanylmethyl) tetrahydrofuran-3,4-diol **(11)**

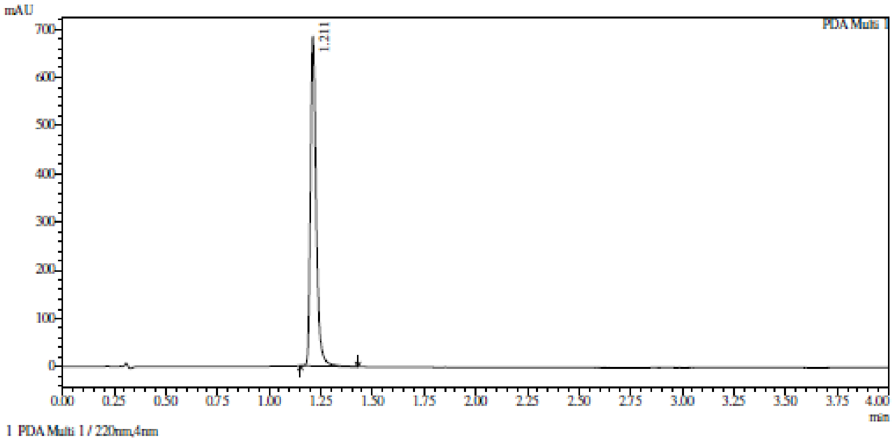

Integration result

PeakTable

PDA Ch1 220nm

| Peak# | Ret. Time | USP Width | Resolution | Height | Area | Area % |
|---|---|---|---|---|---|---|
| 1 | 1.211 | 0.050 | 0.000 | 683586 | 1329817 | 100.000 |
| Total | | | | 683586 | 1329817 | 100.000 |

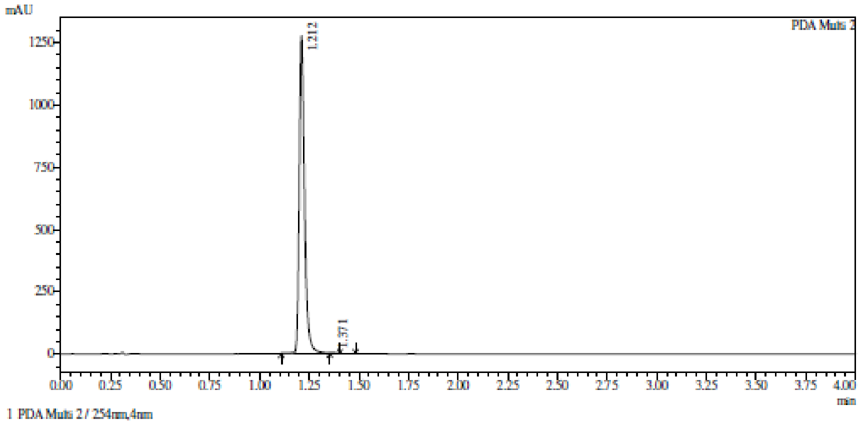

Integration result

PeakTable

PDA Ch2 254nm

| Peak# | Ret. Time | USP Width | Resolution | Height | Area | Area % |
|---|---|---|---|---|---|---|
| 1 | 1.212 | 0.049 | 0.000 | 1275983 | 2410010 | 99.930 |
| 2 | 1.371 | 0.040 | 3.611 | 1169 | 1685 | 0.070 |
| Total | | | | 1277152 | 2411695 | 100.000 |

**HRMS** (2*R*,3*R*,4*S*,5*S*)-2-(6-Aminopurin-9-yl)-5-(thiazol-2-ylsulfanylmethyl) tetrahydrofuran-3,4-diol **(11)**

**Compound Table**

| Label | Tgt Score | Mass Error (ppm) | Tgt Formula | Obs. RT | Ref. Mass | Obs. Mass |
|---|---|---|---|---|---|---|
| Cpd 1: C13 H14 N6 O3 S2; 1.425 | 99.58 | 3.92 | C13 H14 N6 O3 S2 | 1.425 | 366.0569 | 366.0583 |

| Obs. *m/z* | Obs. RT | Obs. Mass | Tgt Formula | Tgt Mass | Tgt Mass Error (ppm) | Find Cpds Algorithm |
|---|---|---|---|---|---|---|
| 389.0464 | 1.425 | 366.0583 | C13 H14 N6 O3 S2 | 366.0569 | 3.92 | Find by Formula |

**Compound Chromatograms**

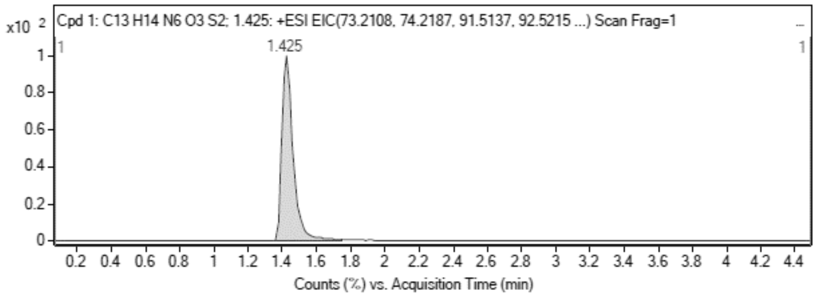

**MS Zoomed Spectrum**

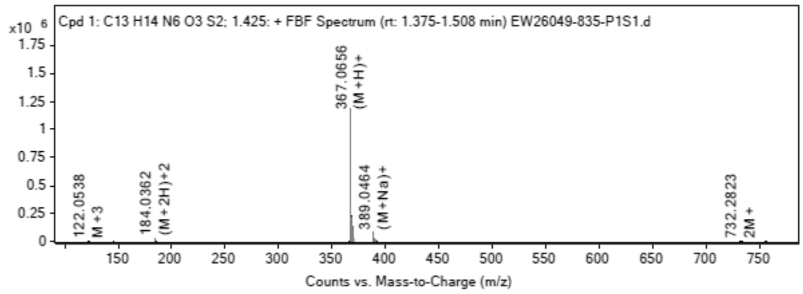

**MS Spectrum Peak List**

| Obs. *m/z* | Charge | Abund | Ion/Isotope |
|---|---|---|---|
| 184.0362 | 2 | 21635.29 | (M+2H)+2 |
| 184.5373 | 2 | 4148.75 | (M+2H)+2 |
| 185.0347 | 2 | 2450.13 | (M+2H)+2 |
| 367.0656 | 1 | 1184609 | (M+H)+ |
| 368.0671 | 1 | 219750.53 | (M+H)+ |
| 369.0624 | 1 | 127705.57 | (M+H)+ |
| 389.0464 | 1 | 82741.53 | (M+Na)+ |
| 390.0484 | 1 | 15333.03 | (M+Na)+ |
| 391.0439 | 1 | 8879.48 | (M+Na)+ |
| 392.0455 | 1 | 1312.33 | (M+Na)+ |

**MS Zoomed Spectrum**

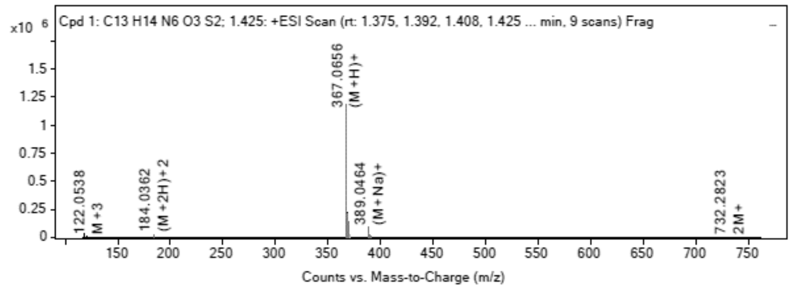

**MS Spectrum Peak List**

| Obs. *m/z* | Charge | Abund | Ion/Isotope | Tgt Mass Error (ppm) |
|---|---|---|---|---|
| 184.0362 | 2 | 21635.29 | (M+2H)+2 | -2.65 |
| 184.5373 | 2 | 4148.75 | (M+2H)+2 | -2.41 |
| 185.0347 | 2 | 2450.13 | (M+2H)+2 | -1.07 |
| 367.0656 | 1 | 1184609 | (M+H)+ | -3.99 |
| 368.0671 | 1 | 219750.53 | (M+H)+ | -1.93 |
| 369.0624 | 1 | 127705.57 | (M+H)+ | -1.93 |
| 389.0464 | 1 | 82741.53 | (M+Na)+ | -0.72 |
| 390.0484 | 1 | 15333.03 | (M+Na)+ | -0.02 |
| 391.0439 | 1 | 8879.48 | (M+Na)+ | -0.86 |
| 392.0455 | 1 | 1312.33 | (M+Na)+ | -0.66 |

--- End Of Report ---

**$^1$H NMR** (2*R*,3*R*,4*S*,5*S*)-2-(6-Aminopurin-9-yl)-5-(imidazo[1,5-a]pyridin-8- ylsulfanylmethyl) tetrahydrofuran-3,4-diol **(12)**

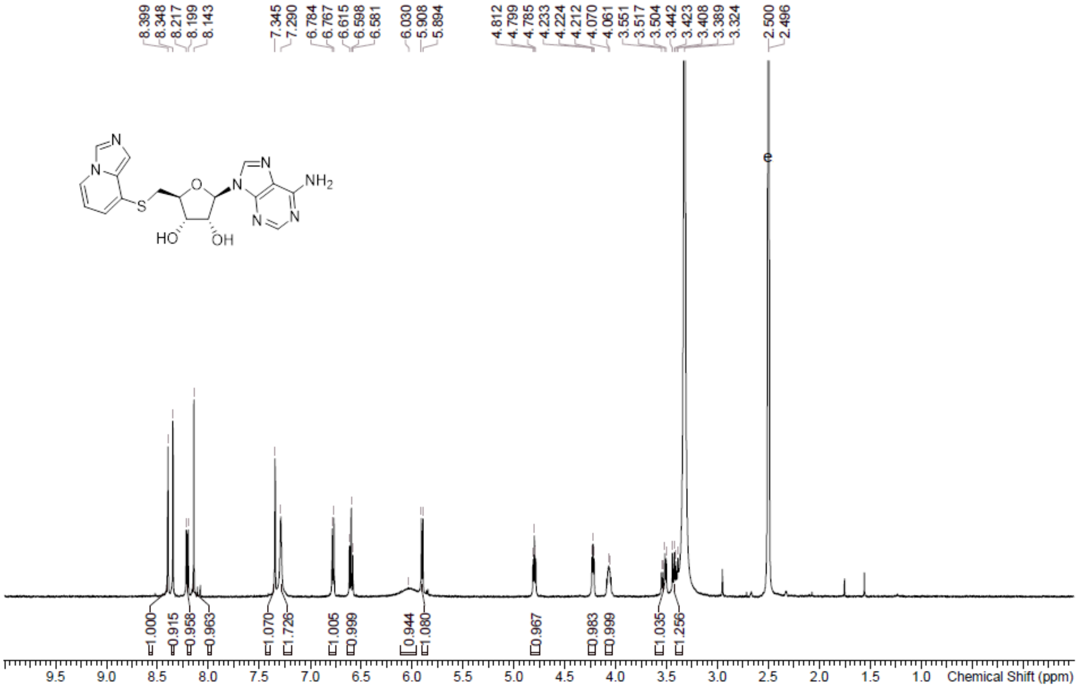

**HPLC** (2*R*,3*R*,4*S*,5*S*)-2-(6-Aminopurin-9-yl)-5-(imidazo[1,5-a]pyridin-8- ylsulfanylmethyl) tetrahydrofuran-3,4-diol **(12)**

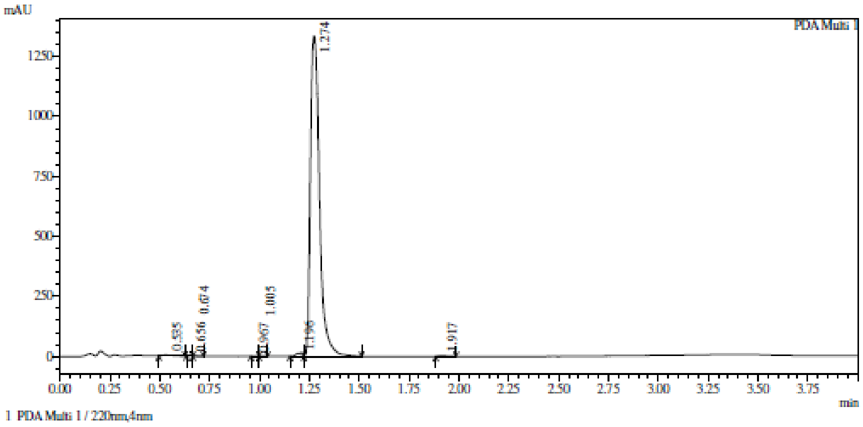

Integration result

PeakTable

PDA Ch1 220nm

| Peak# | Ret. Time | USP Width | Resolution | Height | Area | Area % |
|---|---|---|---|---|---|---|
| 1 | 0.535 | 0.109 | 0.000 | 4051 | 14907 | 0.346 |
| 2 | 0.656 | 0.044 | 1.582 | 1156 | 1387 | 0.032 |
| 3 | 0.674 | 0.052 | 0.365 | 1362 | 2175 | 0.050 |
| 4 | 0.967 | 0.085 | 4.255 | 428 | 1002 | 0.023 |
| 5 | 1.005 | 0.048 | 0.569 | 571 | 1063 | 0.025 |
| 6 | 1.196 | 0.065 | 3.382 | 12166 | 26572 | 0.616 |
| 7 | 1.274 | 0.076 | 1.101 | 1333985 | 4254352 | 98.614 |
| 8 | 1.917 | 0.084 | 8.073 | 4127 | 12668 | 0.294 |
| Total | | | | 1357846 | 4314127 | 100.000 |

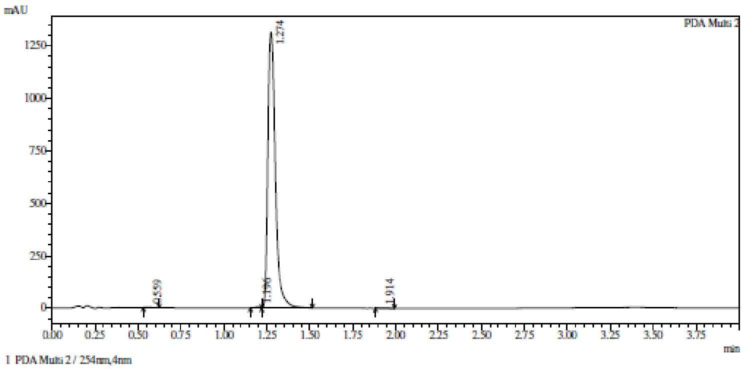

Integration result

PeakTable

PDA Ch2 254nm

| Peak# | Ret. Time | USP Width | Resolution | Height | Area | Area % |
|---|---|---|---|---|---|---|
| 1 | 0.559 | 0.098 | 0.000 | 1360 | 4519 | 0.114 |
| 2 | 1.196 | 0.070 | 7.620 | 4743 | 10395 | 0.263 |
| 3 | 1.274 | 0.073 | 1.085 | 1314521 | 3939312 | 99.509 |
| 4 | 1.914 | 0.072 | 8.819 | 1563 | 4516 | 0.114 |
| Total | | | | 1322187 | 3958742 | 100.000 |

**HRMS** (2*R*,3*R*,4*S*,5*S*)-2-(6-Aminopurin-9-yl)-5-(imidazo[1,5-a]pyridin-8- ylsulfanylmethyl) tetrahydrofuran-3,4-diol **(12)**

**Compound Table**

| Label | Tgt Score | Mass Error (ppm) | Tgt Formula | Obs. RT | Ref. Mass | Obs. Mass |
|---|---|---|---|---|---|---|
| Cpd 1: C17 H17 N7 O3 S; 1.258 | 94.79 | 3.11 | C17 H17 N7 O3 S | 1.258 | 399.1114 | 399.1126 |

| Obs. *m/z* | Obs. RT | Obs. Mass | Tgt Formula | Tgt Mass | Tgt Mass Error (ppm) | Find Cpds Algorithm |
|---|---|---|---|---|---|---|
| 400.1201 | 1.258 | 399.1126 | C17 H17 N7 O3 S | 399.1114 | 3.11 | Find by Formula |

**Compound Chromatograms**

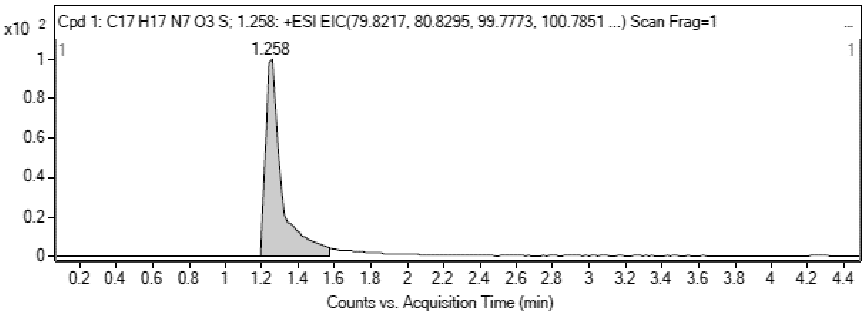

**MS Zoomed Spectrum**

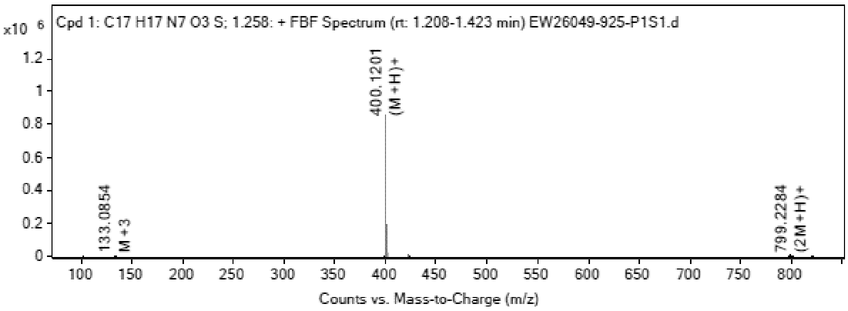

**MS Spectrum Peak List**

| Obs. *m/z* | Charge | Abund | Ion/Isotope |
|---|---|---|---|
| 399.1109 | 1 | 3222.74 | M+ |
| 400.1201 | 1 | 858207.69 | (M+H)+ |

| | | | |
|---|---|---|---|
| 401.1218 | 1 | 189835.05 | (M+H)+ |
| 402.1186 | 1 | 59762.9 | (M+H)+ |
| 422.1006 | 1 | 3575.11 | (M+Na)+ |
| 799.2284 | 1 | 5936.94 | (2M+H)+ |
| 800.2299 | 1 | 2762.15 | (2M+H)+ |
| 801.2293 | 1 | 1206.66 | (2M+H)+ |
| 821.2113 | 1 | 3012.22 | (2M+Na)+ |
| 822.2126 | 1 | 1454.71 | (2M+Na)+ |

MS Zoomed Spectrum

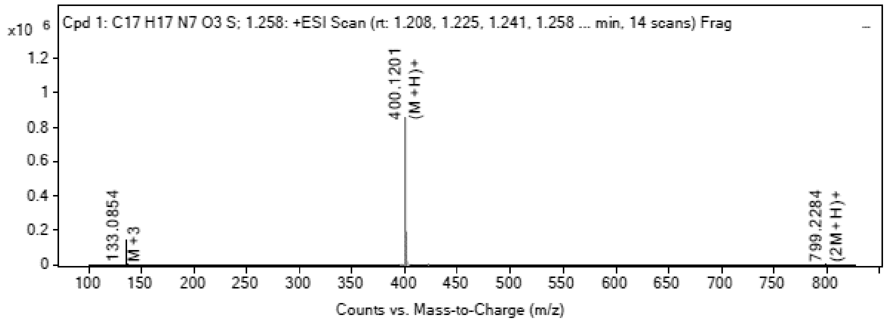

MS Spectrum Peak List

| Obs. m/z | Charge | Abund | Ion/Isotope | Tgt Mass Error (ppm) |
|---|---|---|---|---|
| 399.1109 | 1 | 3222.74 | M+ | -0.26 |
| 400.1201 | 1 | 858207.69 | (M+H)+ | -3.55 |
| 401.1218 | 1 | 189835.05 | (M+H)+ | -1.66 |
| 402.1186 | 1 | 59762.9 | (M+H)+ | -1.45 |
| 422.1006 | 1 | 3575.11 | (M+Na)+ | 0.02 |
| 799.2284 | 1 | 5936.94 | (2M+H)+ | 2.01 |
| 800.2299 | 1 | 2762.15 | (2M+H)+ | 3.26 |
| 801.2293 | 1 | 1206.66 | (2M+H)+ | 1.77 |
| 821.2113 | 1 | 3012.22 | (2M+Na)+ | 0.8 |
| 822.2126 | 1 | 1454.71 | (2M+Na)+ | 2.23 |

--- End Of Report ---

**$^1$H NMR** (2R,3R,4S,5S)-2-(6-Amino-9H-purin-9-yl)-5-(((1-methyl-1H-benzo[d]imidazol-2-yl)thio) methyl)tetrahydrofuran-3,4-diol **(13)**

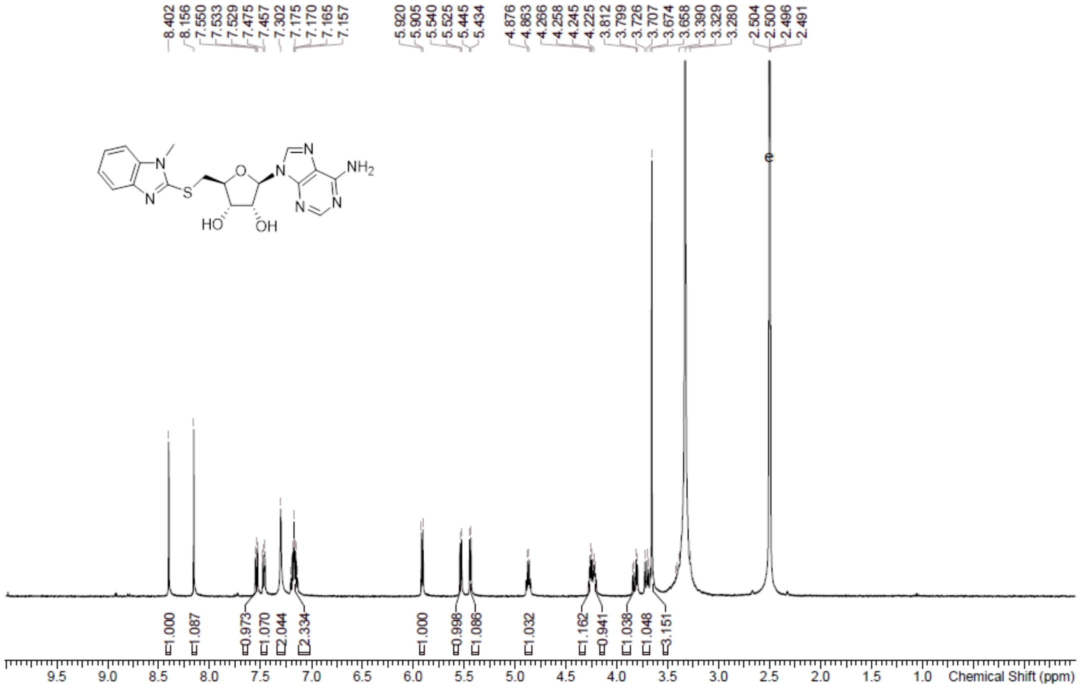

**HPLC** (2R,3R,4S,5S)-2-(6-Amino-9H-purin-9-yl)-5-(((1-methyl-1H-benzo[d]imidazol-2-yl)thio)methyl)tetrahydrofuran-3,4-diol **(13)**

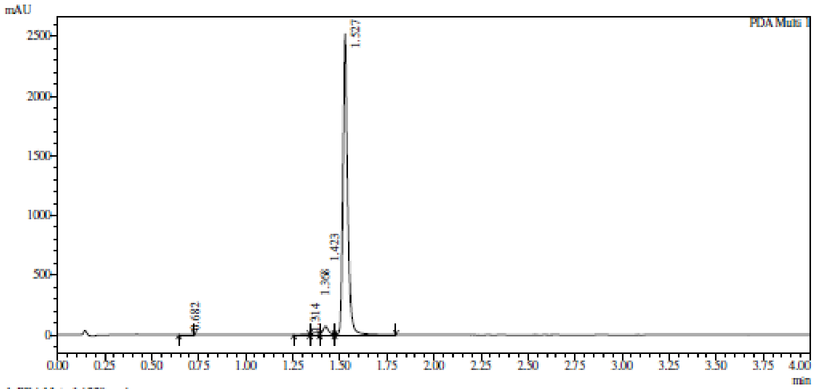

1 PDA Multi 1 / 220nm,4nm

Integration result

PeakTable

PDA Ch1 220nm

| Peak# | Ret. Time | USP Width | Resolution | Height | Area | Area % |
|---|---|---|---|---|---|---|
| 1 | 0.682 | 0.067 | 0.000 | 939 | 2273 | 0.050 |
| 2 | 1.314 | 0.078 | 8.709 | 4803 | 12506 | 0.275 |
| 3 | 1.368 | 0.071 | 0.718 | 19126 | 40212 | 0.886 |
| 4 | 1.423 | 0.051 | 0.896 | 70026 | 141471 | 3.115 |
| 5 | 1.527 | 0.045 | 2.160 | 2521338 | 4344684 | 95.674 |
| Total | | | | 2616232 | 4541146 | 100.000 |

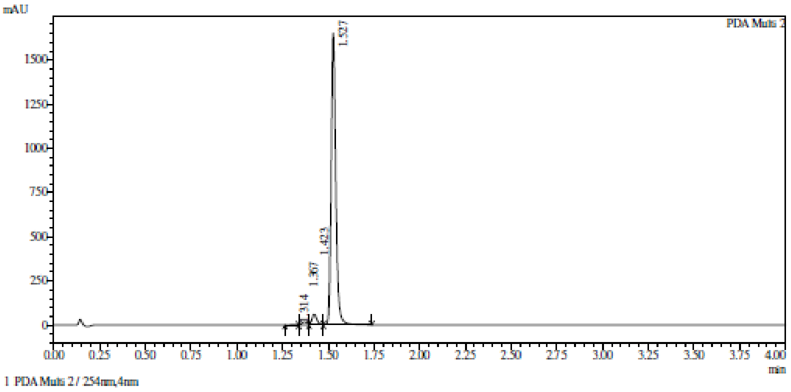

1 PDA Multi 2 / 254nm,4nm

Integration result

PeakTable

PDA Ch2 254nm

| Peak# | Ret. Time | USP Width | Resolution | Height | Area | Area % |
|---|---|---|---|---|---|---|
| 1 | 1.314 | 0.076 | 0.000 | 3112 | 7581 | 0.244 |
| 2 | 1.367 | 0.062 | 0.775 | 18220 | 37337 | 1.200 |
| 3 | 1.423 | 0.051 | 0.978 | 58517 | 116609 | 3.747 |
| 4 | 1.527 | 0.047 | 2.118 | 1650937 | 2950694 | 94.810 |
| Total | | | | 1730786 | 3112221 | 100.000 |

**HRMC** (2R,3R,4S,5S)-2-(6-Amino-9H-purin-9-yl)-5-(((1-methyl-1H-benzo[d]imidazol-2-yl)thio) methyl)tetrahydrofuran-3,4-diol **(13)**

**Compound Table**

| Label | Tgt Score | Mass Error (ppm) | Tgt Formula | Obs. RT | Ref. Mass | Obs. Mass |
|---|---|---|---|---|---|---|
| Cpd 1: C18 H19 N7 O3 S; 1.233 | 99.59 | 2.51 | C18 H19 N7 O3 S | 1.233 | 413.127 | 413.128 |

| Obs. m/z | Obs. RT | Obs. Mass | Tgt Formula | Tgt Mass | Tgt Mass Error (ppm) | Find Cpds Algorithm |
|---|---|---|---|---|---|---|
| 207.5707 | 1.233 | 413.128 | C18 H19 N7 O3 S | 413.127 | 2.51 | Find by Formula |

**Compound Chromatograms**

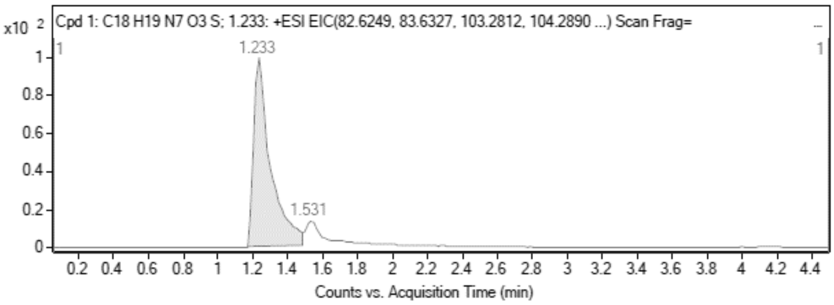

**MS Zoomed Spectrum**

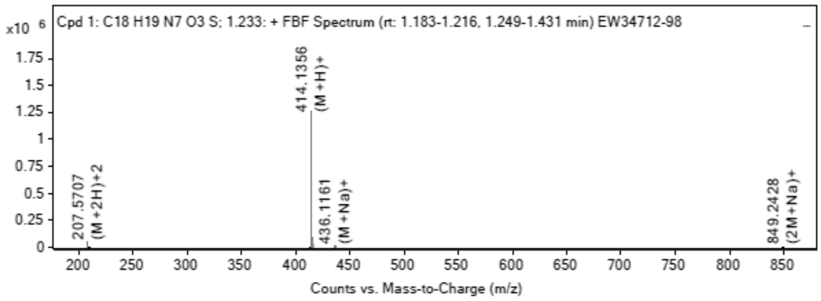

**MS Spectrum Peak List**

| Obs. m/z | Charge | Abund | Ion/Isotope |
|---|---|---|---|
| 207.5707 | 2 | 47508.82 | (M+2H)+2 |
| 208.0721 | 2 | 11656.19 | (M+2H)+2 |
| 208.5706 | 2 | 3979.88 | (M+2H)+2 |
| 413.1211 | 1 | 776.37 | M+ |
| 414.1356 | 1 | 1260667.75 | (M+H)+ |
| 415.1374 | 1 | 296669.78 | (M+H)+ |
| 416.1342 | 1 | 92065.24 | (M+H)+ |
| 436.1161 | 1 | 7755.8 | (M+Na)+ |
| 437.119 | 1 | 1760.48 | (M+Na)+ |
| 849.2428 | 1 | 1189.16 | (2M+Na)+ |

**MS Zoomed Spectrum**

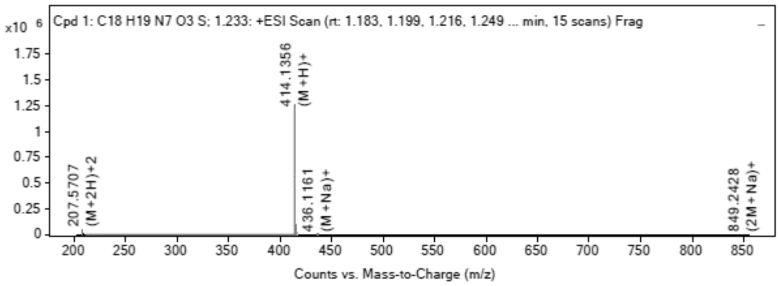

**MS Spectrum Peak List**

| Obs. m/z | Charge | Abund | Ion/Isotope | Tgt Mass Error (ppm) |
|---|---|---|---|---|
| 207.5707 | 2 | 47508.82 | (M+2H)+2 | 0.23 |
| 208.0721 | 2 | 11656.19 | (M+2H)+2 | -0.21 |
| 208.5706 | 2 | 3979.88 | (M+2H)+2 | -0.14 |
| 413.1211 | 1 | 776.37 | M+ | 12.98 |
| 414.1356 | 1 | 1260667.75 | (M+H)+ | -3.07 |
| 415.1374 | 1 | 296669.78 | (M+H)+ | -1.32 |
| 416.1342 | 1 | 92065.24 | (M+H)+ | -0.76 |
| 436.1161 | 1 | 7755.8 | (M+Na)+ | 0.32 |
| 437.119 | 1 | 1760.48 | (M+Na)+ | -0.41 |
| 849.2428 | 1 | 1189.16 | (2M+Na)+ | 0.54 |

--- End Of Report ---

**$^{1}$H NMR** (2R,3R,4S,5S)-2-(6-Amino-9H-purin-9-yl)-5-((benzo[d]oxazol-2-ylthio)methyl) tetrahydrofuran-3,4-diol **(14)**

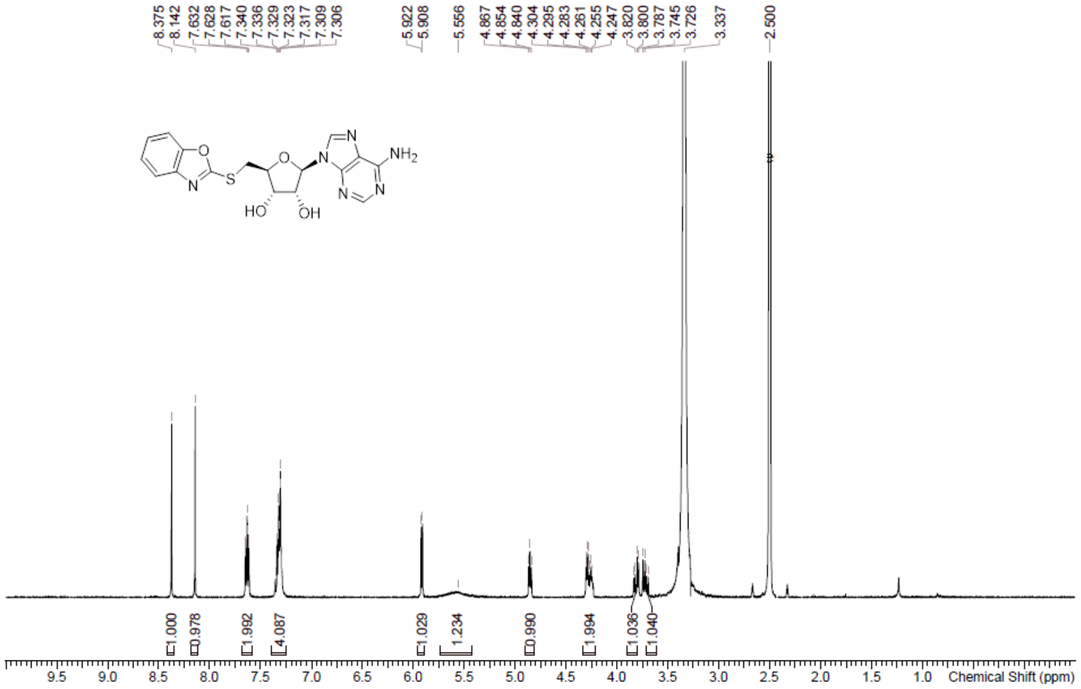

**HPLC** (2R,3R,4S,5S)-2-(6-Amino-9H-purin-9-yl)-5-((benzo[d]oxazol-2-ylthio)methyl) tetrahydrofuran-3,4-diol **(14)**

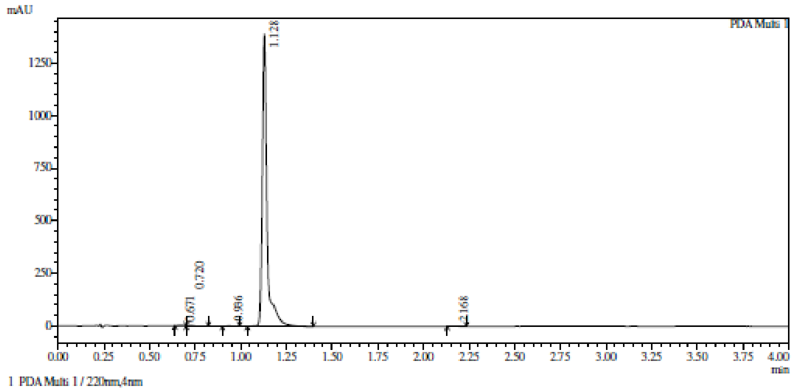

Integration result

PeakTable

PDA Ch1 220nm

| Peak# | Ret. Time | USP Width | Resolution | Height | Area | Area % |
|---|---|---|---|---|---|---|
| 1 | 0.671 | 0.000 | 0.000 | 1265 | 3992 | 0.155 |
| 2 | 0.720 | 0.067 | 0.000 | 3067 | 6431 | 0.250 |
| 3 | 0.936 | 0.054 | 3.568 | 2280 | 4941 | 0.192 |
| 4 | 1.128 | 0.043 | 3.958 | 1390751 | 2548165 | 99.230 |
| 5 | 2.168 | 0.067 | 19.003 | 1747 | 4407 | 0.172 |
| Total | | | | 1399110 | 2567936 | 100.000 |

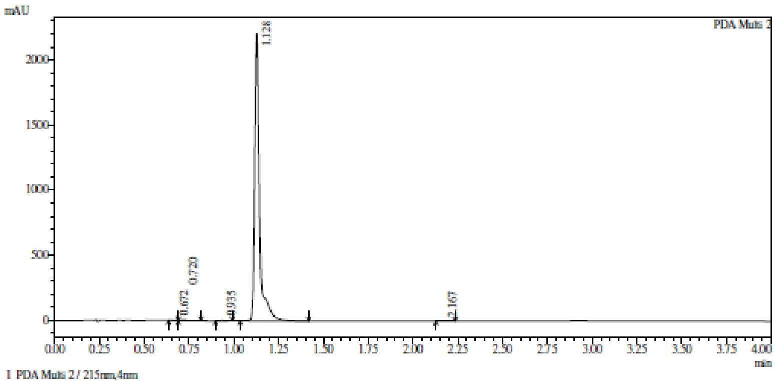

1 PDA Multi 2 / 215nm,4nm

Integration result

PeakTable

PDA Ch2 215nm

| Peak# | Ret. Time | USP Width | Resolution | Height | Area | Area % |
|---|---|---|---|---|---|---|
| 1 | 0.672 | 0.000 | 0.000 | 2448 | 5476 | 0.134 |
| 2 | 0.720 | 0.112 | 0.000 | 5950 | 14812 | 0.362 |
| 3 | 0.935 | 0.052 | 2.631 | 3477 | 6898 | 0.169 |
| 4 | 1.128 | 0.043 | 4.079 | 2202921 | 4060572 | 99.237 |
| 5 | 2.167 | 0.067 | 18.840 | 1561 | 4054 | 0.099 |
| Total | | | | 2216357 | 4091810 | 100.000 |

**HRMS** (2R,3R,4S,5S)-2-(6-Amino-9H-purin-9-yl)-5-((benzo[d]oxazol-2-ylthio)methyl) tetrahydrofuran-3,4-diol **(14)**

**Compound Table**

| Label | Tgt Score | Mass Error (ppm) | Tgt Formula | Obs. RT | Ref. Mass | Obs. Mass |
|---|---|---|---|---|---|---|
| Cpd 1: C17 H16 N6 O4 S; 1.644 | 98.19 | 3.31 | C17 H16 N6 O4 S | 1.644 | 400.0954 | 400.0967 |

| Obs. m/z | Obs. RT | Obs. Mass | Tgt Formula | Tgt Mass | Tgt Mass Error (ppm) | Find Cpds Algorithm |
|---|---|---|---|---|---|---|
| 423.0844 | 1.644 | 400.0967 | C17 H16 N6 O4 S | 400.0954 | 3.31 | Find by Formula |

**Compound Chromatograms**

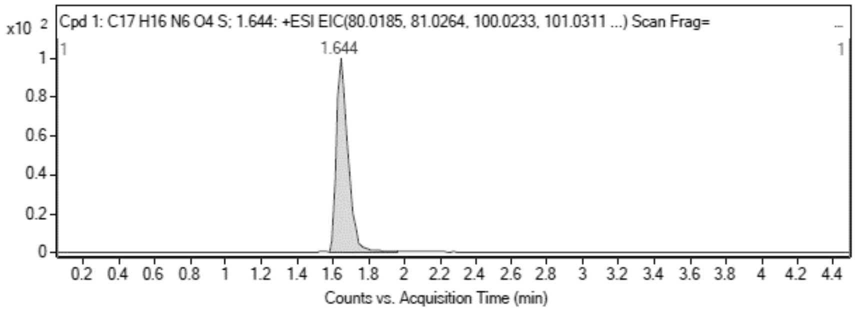

**MS Zoomed Spectrum**

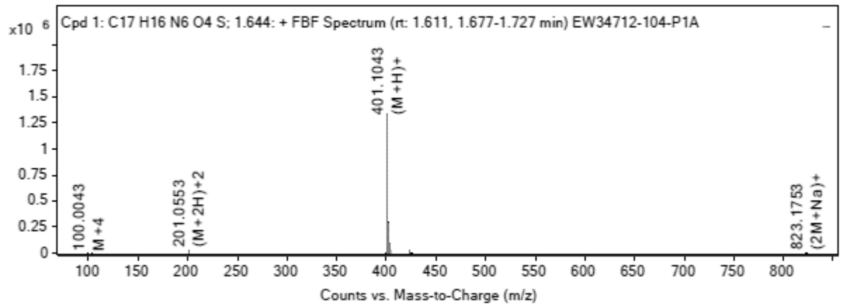

**MS Spectrum Peak List**

| Obs. m/z | Charge | Abund | Ion/Isotope |
|---|---|---|---|
| 201.0553 | 2 | 29501.22 | (M+2H)+2 |
| 201.5566 | 2 | 7101.29 | (M+2H)+2 |

| | | | |
|---|---|---|---|
| 202.055 | 2 | 2179.13 | (M+2H)+2 |
| 401.1043 | 1 | 1335291.75 | (M+H)+ |
| 402.106 | 1 | 292246.5 | (M+H)+ |
| 403.1028 | 1 | 95815.08 | (M+H)+ |
| 423.0844 | 1 | 24024.24 | (M+Na)+ |
| 424.0863 | 1 | 5721.87 | (M+Na)+ |
| 425.0841 | 1 | 2038.85 | (M+Na)+ |
| 823.1753 | 1 | 392.42 | (2M+Na)+ |

MS Zoomed Spectrum

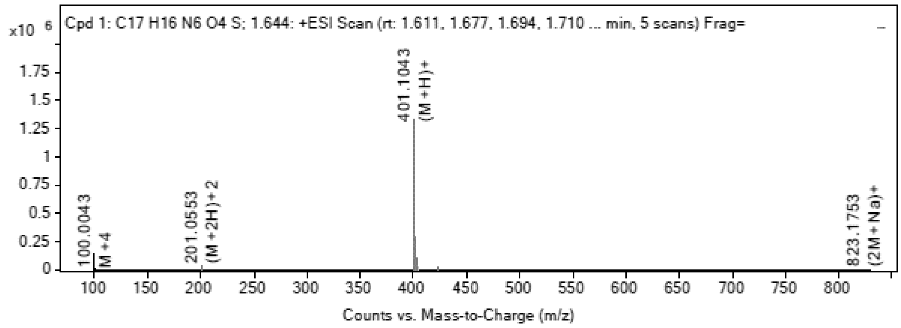

MS Spectrum Peak List

| Obs. m/z | Charge | Abund | Ion/Isotope | Tgt Mass Error (ppm) |
|---|---|---|---|---|
| 201.0553 | 2 | 29501.22 | (M+2H)+2 | -1.44 |
| 201.5566 | 2 | 7101.29 | (M+2H)+2 | -1.69 |
| 202.055 | 2 | 2179.13 | (M+2H)+2 | -1.23 |
| 401.1043 | 1 | 1335291.75 | (M+H)+ | -4 |
| 402.106 | 1 | 292246.5 | (M+H)+ | -1.96 |
| 403.1028 | 1 | 95815.08 | (M+H)+ | -1.54 |
| 423.0844 | 1 | 24024.24 | (M+Na)+ | 0.46 |
| 424.0863 | 1 | 5721.87 | (M+Na)+ | 2.22 |
| 425.0841 | 1 | 2038.85 | (M+Na)+ | 0.04 |
| 823.1753 | 1 | 392.42 | (2M+Na)+ | 5.64 |

--- End Of Report ---

**$^1$H NMR** (2S,3S,4R,5R)-2-(((1H-Benzo[d]imidazol-2-yl)thio)methyl)-5-(6-amino-9H-purin-9-yl) tetrahydrofuran-3,4-diol **(15)**

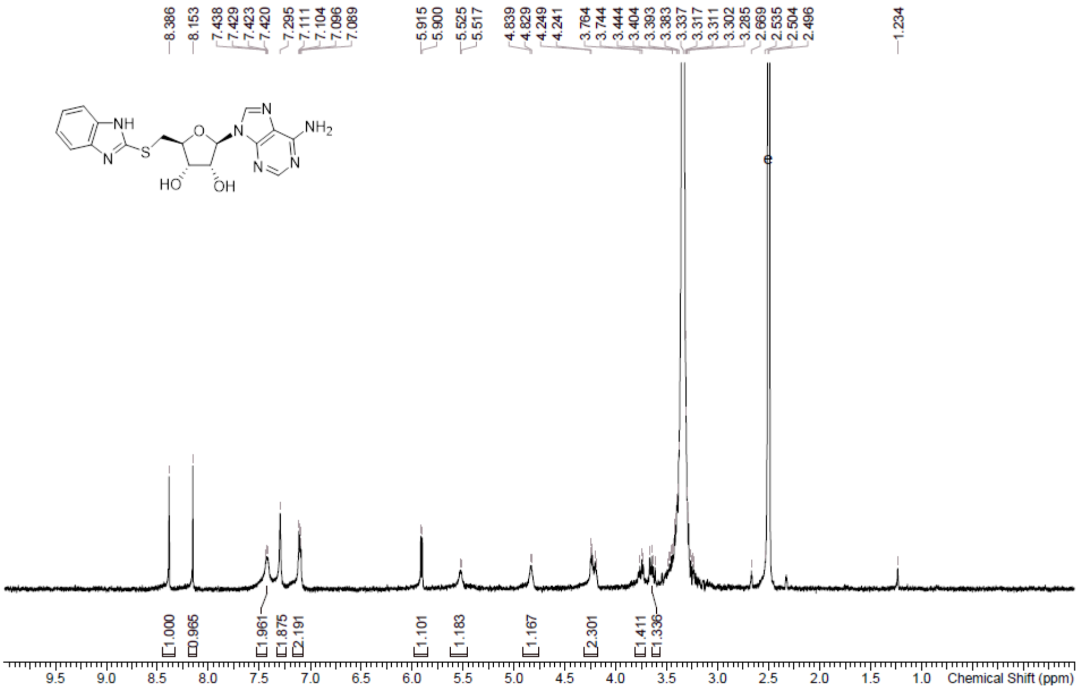

**HPLC** (2S,3S,4R,5R)-2-(((1H-Benzo[d]imidazol-2-yl)thio)methyl)-5-(6-amino-9H-purin-9-yl) tetrahydrofuran-3,4-diol **(15)**

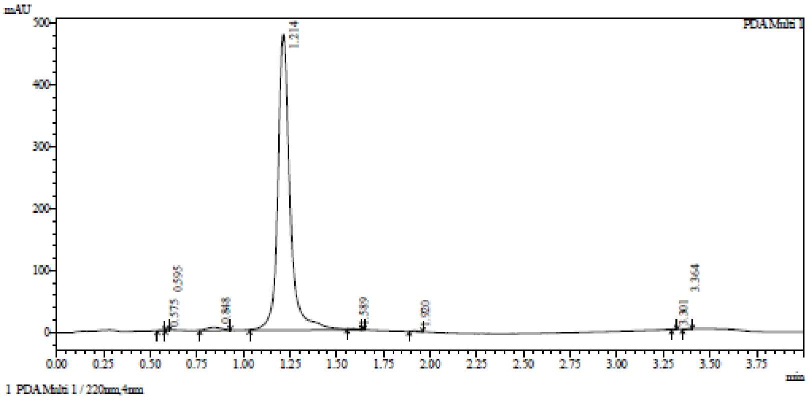

Integration result

PeakTable

PDA Ch1 220nm

| Peak# | Ret. Time | USP Width | Resolution | Height | Area | Area % |
|---|---|---|---|---|---|---|
| 1 | 0.575 | 0.057 | 0.000 | 886 | 1572 | 0.071 |
| 2 | 0.595 | 0.064 | 0.332 | 951 | 1130 | 0.051 |
| 3 | 0.848 | 0.143 | 2.449 | 4925 | 27014 | 1.212 |
| 4 | 1.214 | 0.106 | 2.932 | 479310 | 2190445 | 98.284 |
| 5 | 1.589 | 0.047 | 4.891 | 1010 | 1949 | 0.087 |
| 6 | 1.920 | 0.054 | 6.520 | 1639 | 3246 | 0.146 |
| 7 | 3.301 | 0.087 | 19.565 | 890 | 1420 | 0.064 |
| 8 | 3.364 | 0.119 | 0.610 | 673 | 1915 | 0.086 |
| Total | | | | 490283 | 2228692 | 100.000 |

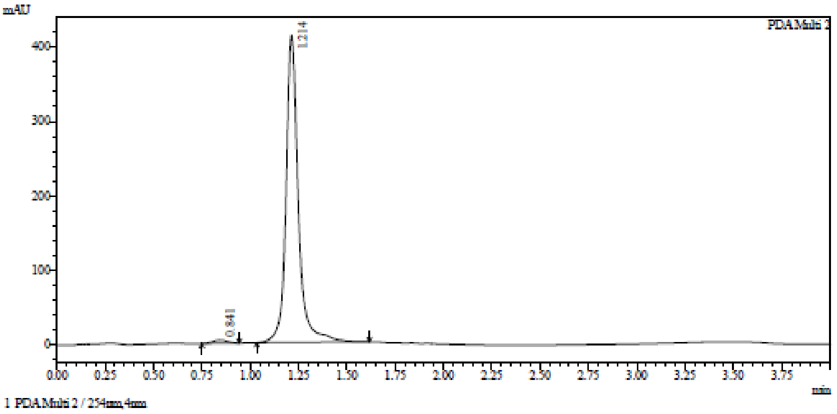

Integration result

PeakTable

PDA Ch2 254nm

| Peak# | Ret. Time | USP Width | Resolution | Height | Area | Area % |
|---|---|---|---|---|---|---|
| 1 | 0.841 | 0.151 | 0.000 | 4303 | 23862 | 1.292 |
| 2 | 1.214 | 0.105 | 2.923 | 412225 | 1822897 | 98.708 |
| Total | | | | 416528 | 1846759 | 100.000 |

**HRMC** (2S,3S,4R,5R)-2-(((1H-Benzo[d]imidazol-2-yl)thio)methyl)-5-(6-amino-9H-purin-9-yl) tetrahydrofuran-3,4-diol **(15)**

**Compound Table**

| Label | Tgt Score | Mass Error (ppm) | Tgt Formula | Obs. RT | Ref. Mass | Obs. Mass |
|---|---|---|---|---|---|---|
| Cpd 1: C17 H17 N7 O3 S; 1.155 | 95.02 | 3.16 | C17 H17 N7 O3 S | 1.155 | 399.1114 | 399.1126 |

| Obs. m/z | Obs. RT | Obs. Mass | Tgt Formula | Tgt Mass | Tgt Mass Error (ppm) | Find Cpds Algorithm |
|---|---|---|---|---|---|---|
| 400.1201 | 1.155 | 399.1126 | C17 H17 N7 O3 S | 399.1114 | 3.16 | Find by Formula |

**Compound Chromatograms**

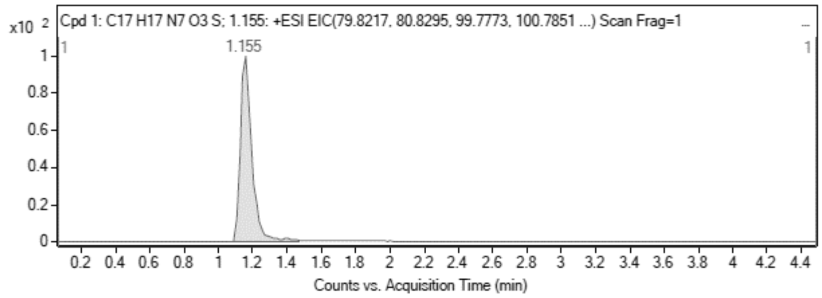

MS Zoomed Spectrum

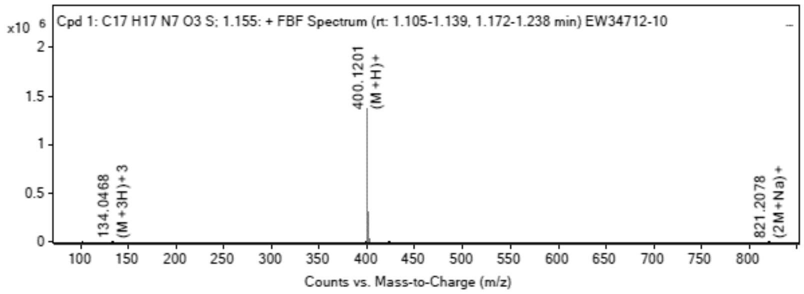

**MS Spectrum Peak List**

| Obs. m/z | Charge | Abund | Ion/Isotope |
|---|---|---|---|
| 133.073 | 3 | 289.63 | M+3 |
| 134.0468 | 3 | 4387.62 | (M+3H)+3 |
| 399.1109 | 1 | 298.95 | M+ |
| 400.1201 | 1 | 1368297 | (M+H)+ |
| 401.1218 | 1 | 306489.69 | (M+H)+ |
| 402.1186 | 1 | 94816.95 | (M+H)+ |
| 403.119 | 1 | 15879.96 | (M+H)+ |
| 422.1008 | 1 | 3490.26 | (M+Na)+ |
| 423.1031 | 1 | 772.61 | (M+Na)+ |
| 821.2078 | 1 | 325.2 | (2M+Na)+ |

MS Zoomed Spectrum

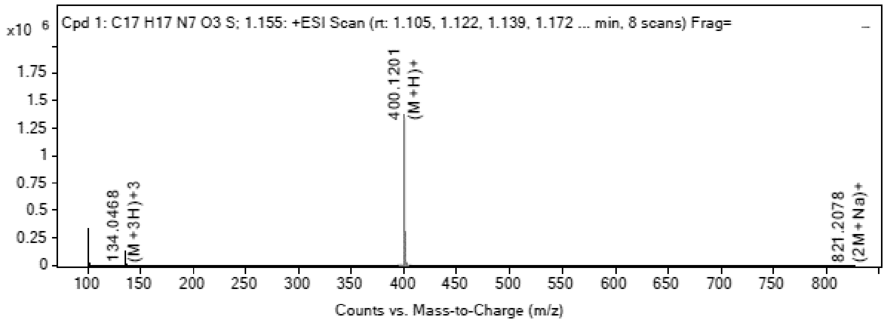

**MS Spectrum Peak List**

| Obs. m/z | Charge | Abund | Ion/Isotope | Tgt Mass Error (ppm) |
|---|---|---|---|---|
| 133.073 | 3 | 289.63 | M+3 | -274.01 |
| 134.0468 | 3 | 4387.62 | (M+3H)+3 | -17.76 |
| 399.1109 | 1 | 298.95 | M+ | -0.34 |
| 400.1201 | 1 | 1368297 | (M+H)+ | -3.63 |
| 401.1218 | 1 | 306489.69 | (M+H)+ | -1.52 |
| 402.1186 | 1 | 94816.95 | (M+H)+ | -1.44 |
| 403.119 | 1 | 15879.96 | (M+H)+ | 0.23 |
| 422.1008 | 1 | 3490.26 | (M+Na)+ | -0.5 |
| 423.1031 | 1 | 772.61 | (M+Na)+ | -0.09 |
| 821.2078 | 1 | 325.2 | (2M+Na)+ | 5 |

--- End Of Report ---

**$^{1}$H NMR** (2S,3S,4R,5R)-2-(((1H-Indazol-7-yl)thio)methyl)-5-(6-amino-9H-purin-9-yl)tetrahydrofuran-3,4-diol **(16)**

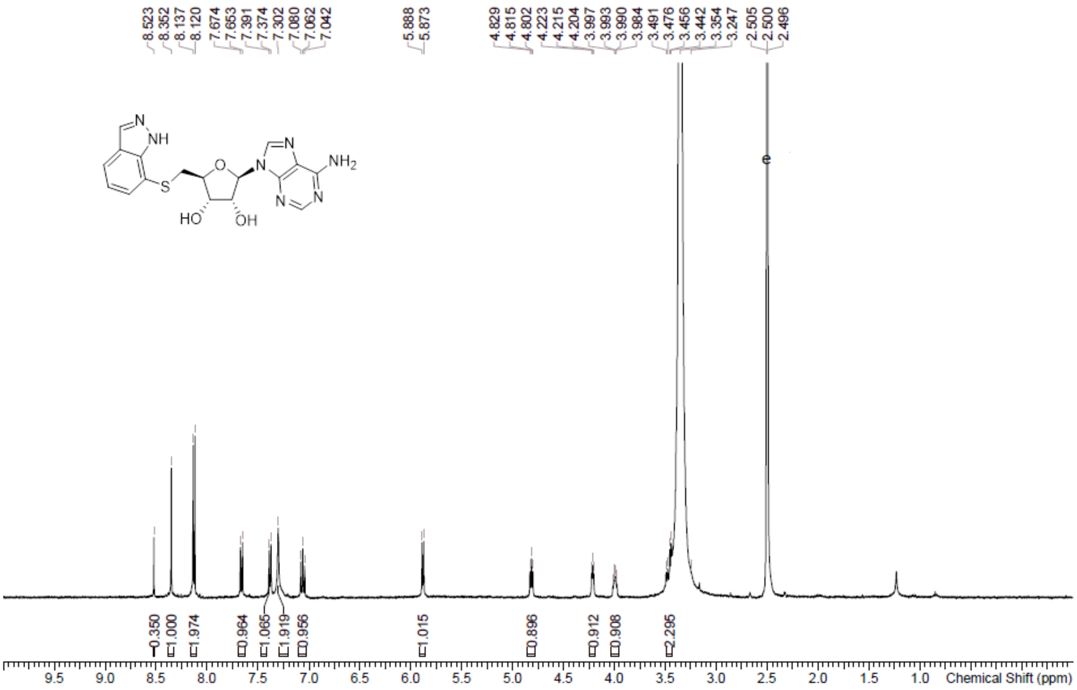

**HPLC** (2S,3S,4R,5R)-2-(((1H-Indazol-7-yl)thio)methyl)-5-(6-amino-9H-purin-9-yl)tetrahydrofuran-3,4-diol **(16)**

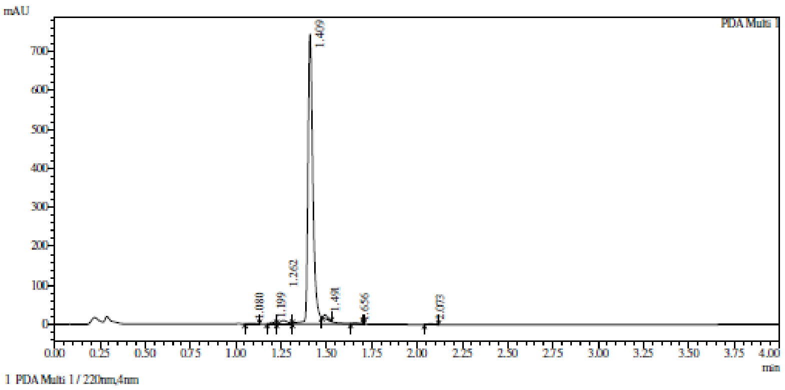

Integration result

PeakTable

PDA Ch1 220nm

| Peak# | Ret. Time | USP Width | Resolution | Height | Area | Area % |
|---|---|---|---|---|---|---|
| 1 | 1.080 | 0.061 | 0.000 | 1226 | 2823 | 0.182 |
| 2 | 1.199 | 0.045 | 2.235 | 3697 | 6006 | 0.387 |
| 3 | 1.262 | 0.073 | 1.066 | 8161 | 23077 | 1.487 |
| 4 | 1.409 | 0.050 | 2.389 | 744300 | 1498579 | 96.580 |
| 5 | 1.491 | 0.038 | 1.874 | 10411 | 14230 | 0.917 |
| 6 | 1.656 | 0.047 | 3.855 | 1963 | 3486 | 0.225 |
| 7 | 2.073 | 0.052 | 8.426 | 1868 | 3448 | 0.222 |
| Total | | | | 771627 | 1551649 | 100.000 |

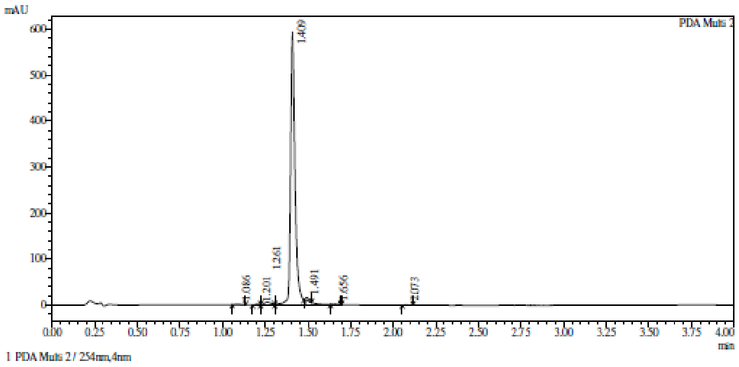

1 PDA Multi 2 / 254nm,4nm

Integration result

PeakTable

PDA Ch2 254nm

| Peak# | Ret. Time | USP Width | Resolution | Height | Area | Area % |
|---|---|---|---|---|---|---|
| 1 | 1.086 | 0.065 | 0.000 | 776 | 1792 | 0.151 |
| 2 | 1.201 | 0.051 | 1.988 | 1362 | 2224 | 0.187 |
| 3 | 1.261 | 0.072 | 0.973 | 5489 | 15103 | 1.270 |
| 4 | 1.409 | 0.048 | 2.451 | 593115 | 1158941 | 97.489 |
| 5 | 1.491 | 0.037 | 1.918 | 5790 | 7794 | 0.656 |
| 6 | 1.656 | 0.047 | 3.933 | 1109 | 1904 | 0.160 |
| 7 | 2.073 | 0.046 | 9.012 | 592 | 1041 | 0.088 |
| Total | | | | 608233 | 1188798 | 100.000 |

**HRMC** (2S,3S,4R,5R)-2-(((1H-Indazol-7-yl)thio)methyl)-5-(6-amino-9H-purin-9-yl)tetrahydrofuran-3,4-diol **(16)**

**Compound Table**

| Label | Tgt Score | Mass Error (ppm) | Tgt Formula | Obs. RT | Ref. Mass | Obs. Mass |
|---|---|---|---|---|---|---|
| Cpd 1: C17 H17 N7 O3 S; 1.547 | 99.21 | -0.76 | C17 H17 N7 O3 S | 1.547 | 399.1114 | 399.1111 |

| Obs. m/z | Obs. RT | Obs. Mass | Tgt Formula | Tgt Mass | Tgt Mass Error (ppm) | Find Cpds Algorithm |
|---|---|---|---|---|---|---|
| 400.1187 | 1.547 | 399.1111 | C17 H17 N7 O3 S | 399.1114 | -0.76 | Find by Formula |

**Compound Chromatograms**

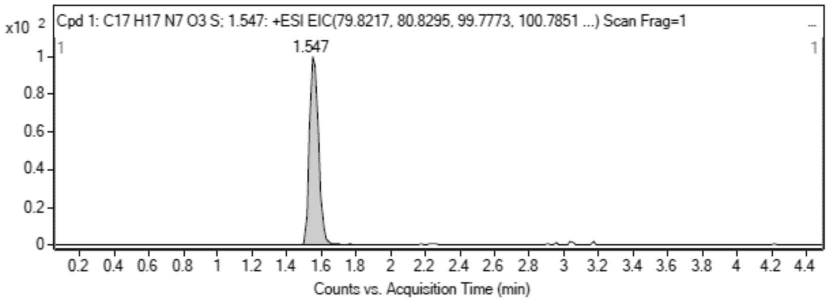

**MS Zoomed Spectrum**

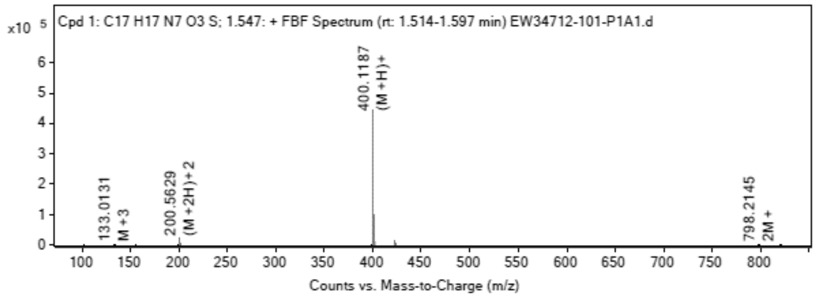

**MS Spectrum Peak List**

| Obs. m/z | Charge | Abund | Ion/Isotope |
|---|---|---|---|
| 133.0131 | 3 | 1616.92 | M+3 |
| 200.5629 | 2 | 20456.9 | (M+2H)+2 |

| | | | |
|---|---|---|---|
| 201.0662 | 2 | 5054.11 | (M+2H)+2 |
| 400.1187 | 1 | 444134.88 | (M+H)+ |
| 401.1209 | 1 | 94815.27 | (M+H)+ |
| 402.1177 | 1 | 29685.77 | (M+H)+ |
| 422.0997 | 1 | 12629.72 | (M+Na)+ |
| 423.1028 | 1 | 2998.7 | (M+Na)+ |
| 424.0982 | 1 | 1082.64 | (M+Na)+ |
| 798.2145 | 1 | 1554.26 | 2M+ |

MS Zoomed Spectrum

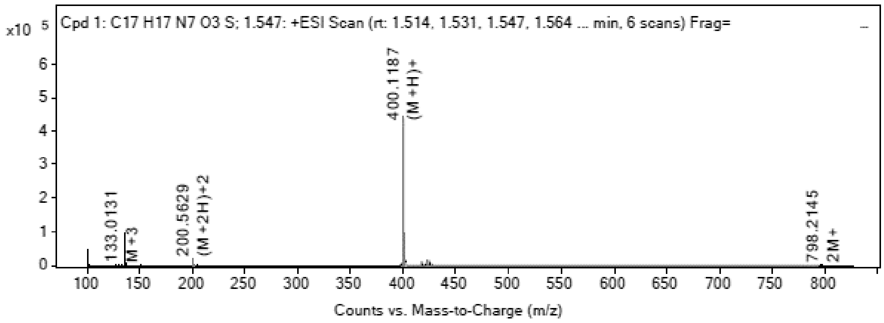

MS Spectrum Peak List

| Obs. m/z | Charge | Abund | Ion/Isotope | Tgt Mass Error (ppm) |
|---|---|---|---|---|
| 133.0131 | 3 | 1616.92 | M+3 | 176.69 |
| 200.5629 | 2 | 20456.9 | (M+2H)+2 | 0.42 |
| 201.0662 | 2 | 5054.11 | (M+2H)+2 | -9.98 |
| 400.1187 | 1 | 444134.88 | (M+H)+ | -0.28 |
| 401.1209 | 1 | 94815.27 | (M+H)+ | 0.71 |
| 402.1177 | 1 | 29685.77 | (M+H)+ | 0.86 |
| 422.0997 | 1 | 12629.72 | (M+Na)+ | 2.06 |
| 423.1028 | 1 | 2998.7 | (M+Na)+ | 0.66 |
| 424.0982 | 1 | 1082.64 | (M+Na)+ | 4.25 |
| 798.2145 | 1 | 1554.26 | 2M+ | 9.57 |

--- End Of Report ---

**$^1$H NMR** (2*R*,3*R*,4*S*,5*S*)-2-(6-Aminopurin-9-yl)-5-(imidazo[1,5-a]pyridine-7-ylsulfanylmethyl) tetrahydrofuran-3,4-diol **(17)**

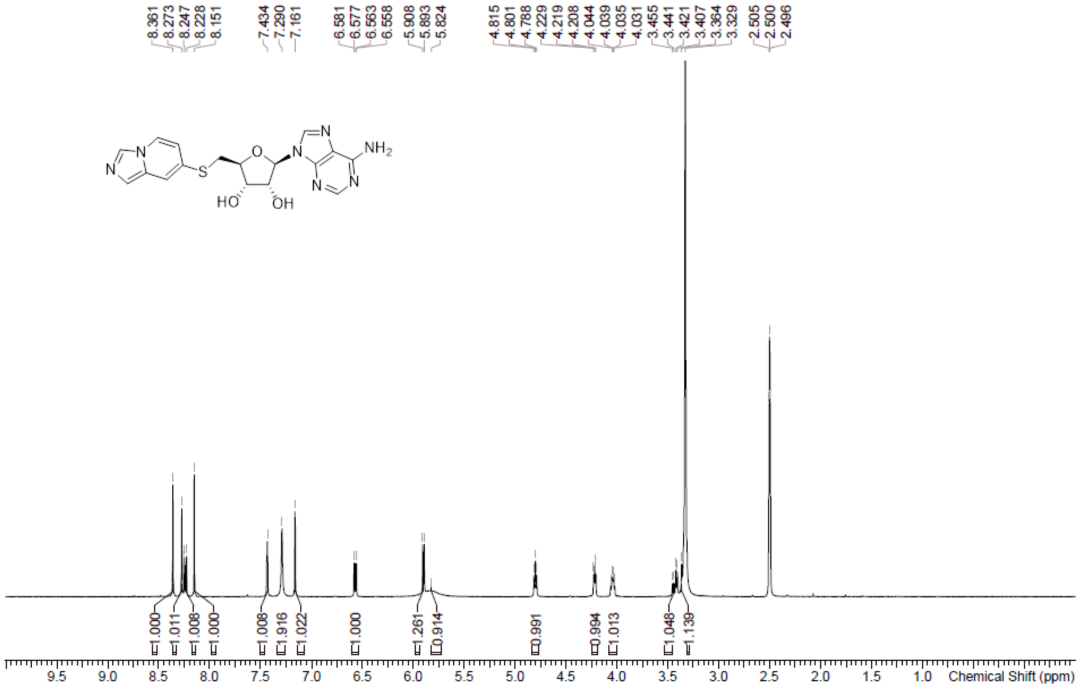

**HPLC** (2*R*,3*R*,4*S*,5*S*)-2-(6-Aminopurin-9-yl)-5-(imidazo[1,5-a]pyridine-7-ylsulfanylmethyl) tetrahydrofuran-3,4-diol **(17)**

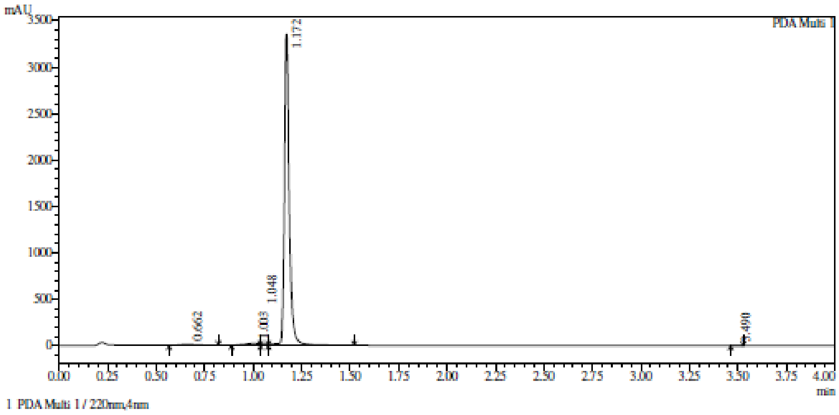

Integration result

PeakTable

PDA Ch1 220nm

| Peak# | Ret. Time | USP Width | Resolution | Height | Area | Area % |
|---|---|---|---|---|---|---|
| 1 | 0.662 | 0.163 | 0.000 | 10342 | 62083 | 1.037 |
| 2 | 1.003 | 0.153 | 2.158 | 22221 | 109086 | 1.822 |
| 3 | 1.048 | 0.364 | 0.174 | 15616 | 33664 | 0.562 |
| 4 | 1.172 | 0.040 | 0.613 | 3348987 | 5781359 | 96.557 |
| 5 | 3.490 | 0.057 | 47.911 | 647 | 1296 | 0.022 |
| Total | | | | 3397813 | 5987488 | 100.000 |

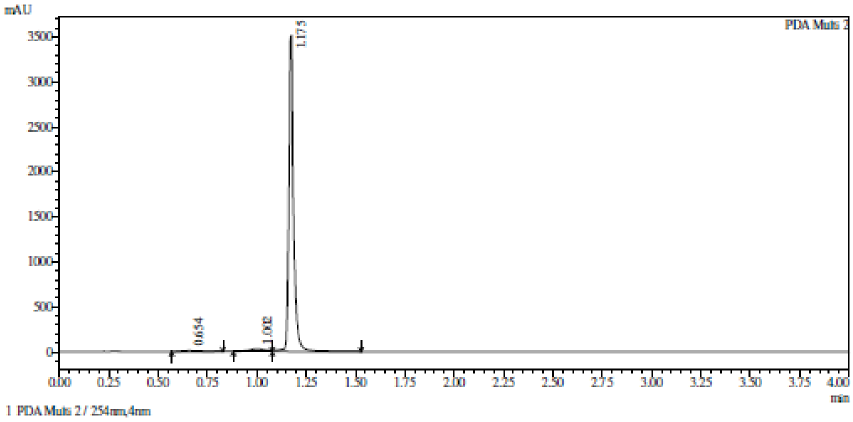

Integration result

PeakTable

PDA Ch2 254nm

| Peak# | Ret. Time | USP Width | Resolution | Height | Area | Area % |
|---|---|---|---|---|---|---|
| 1 | 0.654 | 0.180 | 0.000 | 10105 | 68263 | 1.112 |
| 2 | 1.002 | 0.143 | 2.149 | 31580 | 184359 | 3.004 |
| 3 | 1.175 | 0.036 | 1.929 | 3516010 | 5885093 | 95.884 |
| Total | | | | 3557695 | 6137715 | 100.000 |

**HRMC** (2*R*,3*R*,4*S*,5*S*)-2-(6-Aminopurin-9-yl)-5-(imidazo[1,5-a]pyridine-7-ylsulfanylmethyl) tetrahydrofuran-3,4-diol **(17)**

**Compound Table**

| Label | Tgt Score | Mass Error (ppm) | Tgt Formula | Obs. RT | Ref. Mass | Obs. Mass |
|---|---|---|---|---|---|---|
| Cpd 1: C17 H17 N7 O3 S; 1.195 | 94.54 | -3.76 | C17 H17 N7 O3 S | 1.195 | 399.1114 | 399.1099 |

| Obs. *m/z* | Obs. RT | Obs. Mass | Tgt Formula | Tgt Mass | Tgt Mass Error (ppm) | Find Cpds Algorithm |
|---|---|---|---|---|---|---|
| 400.1176 | 1.195 | 399.1099 | C17 H17 N7 O3 S | 399.1114 | -3.76 | Find by Formula |

**Compound Chromatograms**

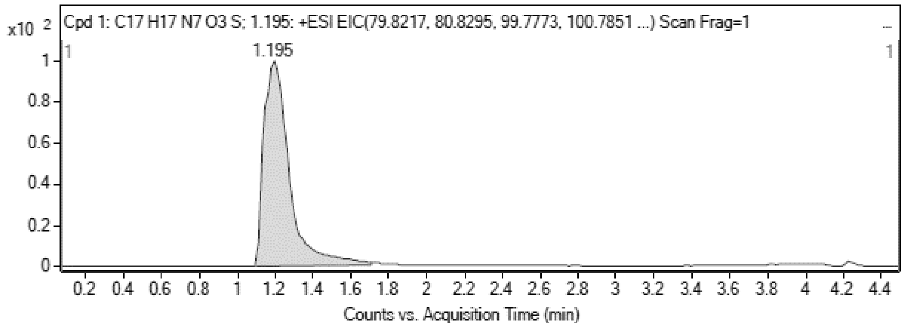

**MS Zoomed Spectrum**

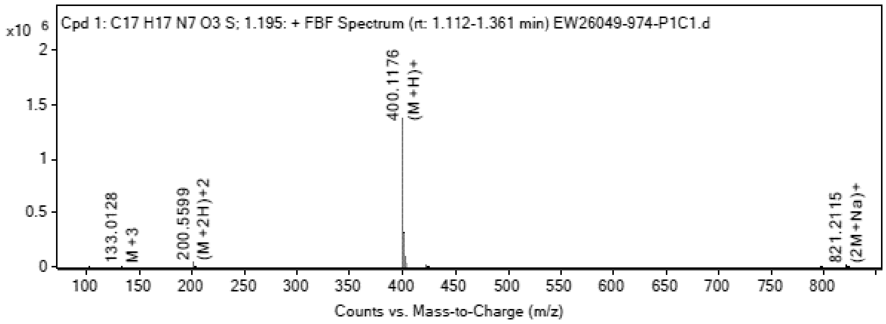

**MS Spectrum Peak List**

| Obs. *m/z* | Charge | Abund | Ion/Isotope |
|---|---|---|---|
| 200.5599 | 2 | 44219.67 | (M+2H)+2 |
| 201.0621 | 2 | 9302.93 | (M+2H)+2 |
| 399.1079 | 1 | 11667.74 | M+ |
| 400.1176 | 1 | 1360646.63 | (M+H)+ |
| 401.1193 | 1 | 313413.34 | (M+H)+ |
| 402.1162 | 1 | 98309.98 | (M+H)+ |
| 422.0981 | 1 | 11562.54 | (M+Na)+ |
| 798.2137 | 1 | 4018.8 | 2M+ |
| 821.2115 | 1 | 11299.91 | (2M+Na)+ |
| 822.2134 | 1 | 4923.16 | (2M+Na)+ |

**MS Zoomed Spectrum**

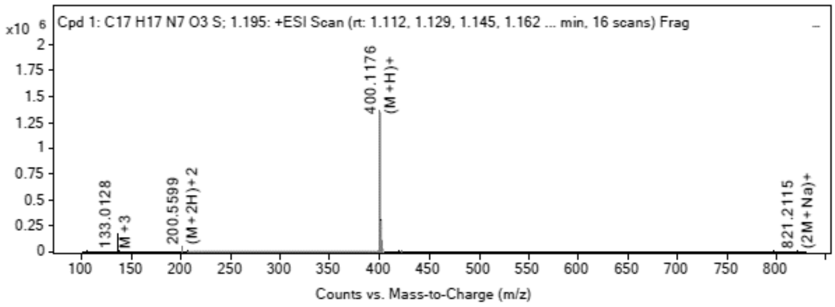

**MS Spectrum Peak List**

| Obs. *m/z* | Charge | Abund | Ion/Isotope | Tgt Mass Error (ppm) |
|---|---|---|---|---|
| 200.5599 | 2 | 44219.67 | (M+2H)+2 | 15.39 |
| 201.0621 | 2 | 9302.93 | (M+2H)+2 | 10.72 |
| 399.1079 | 1 | 11667.74 | M+ | 7.3 |
| 400.1176 | 1 | 1360646.63 | (M+H)+ | 2.64 |
| 401.1193 | 1 | 313413.34 | (M+H)+ | 4.58 |
| 402.1162 | 1 | 98309.98 | (M+H)+ | 4.55 |
| 422.0981 | 1 | 11562.54 | (M+Na)+ | 5.94 |
| 798.2137 | 1 | 4018.8 | 2M+ | 10.64 |
| 821.2115 | 1 | 11299.91 | (2M+Na)+ | 0.54 |
| 822.2134 | 1 | 4923.16 | (2M+Na)+ | 1.25 |

--- End Of Report ---

**[1]H NMR** (2*R*,3*R*,4*S*,5*R*)-2-(6-aminopurin-9-yl)-5-(1,3-benzothiazol-2-yloxymethyl)tetrahydrofuran-3,4-diol **(23)**

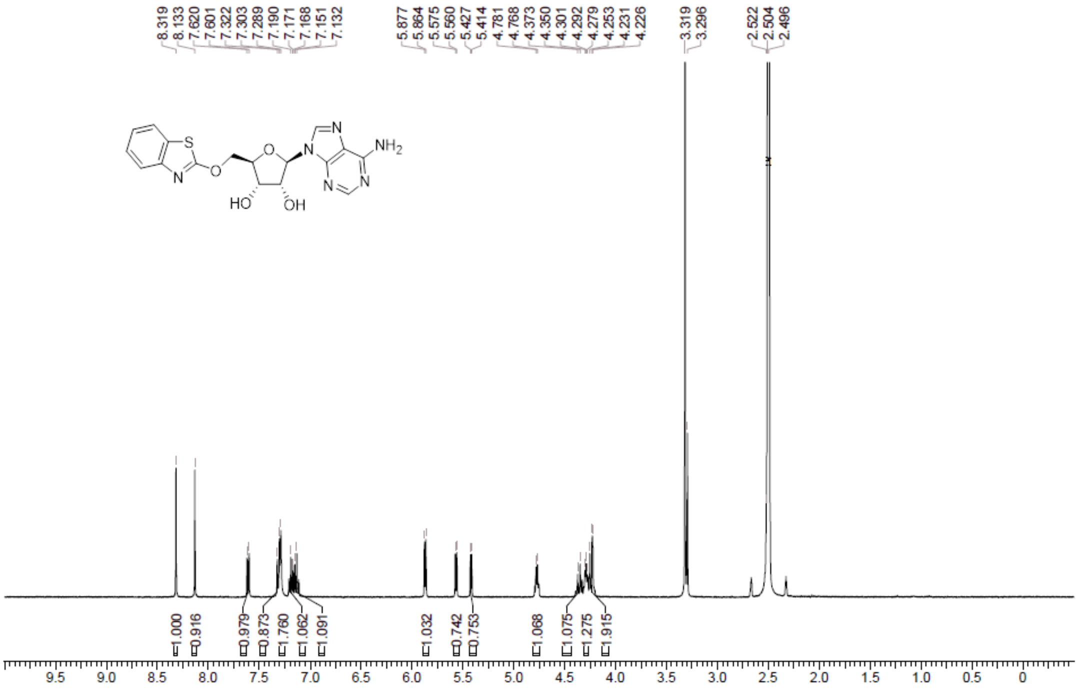

**HPLC** (2*R*,3*R*,4*S*,5*R*)-2-(6-aminopurin-9-yl)-5-(1,3-benzothiazol-2-yloxymethyl)tetrahydrofuran-3,4-diol **(23)**

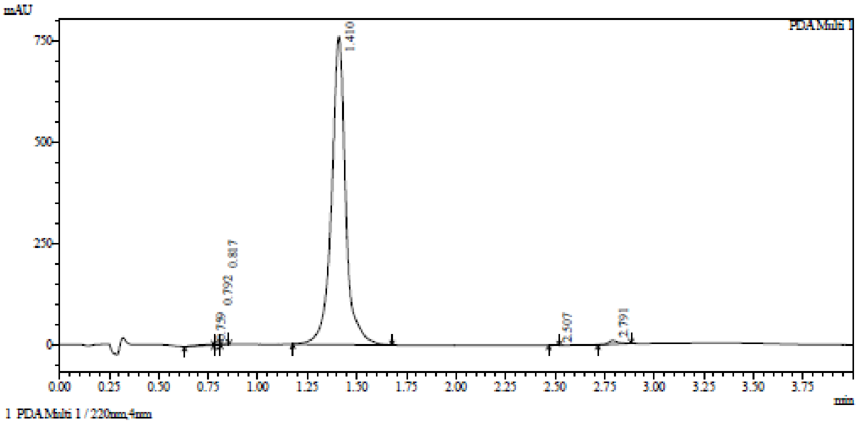

Integration result

PeakTable

PDA Ch1 220nm

| Peak# | Ret. Time | USP Width | Resolution | Height | Area | Area % |
|---|---|---|---|---|---|---|
| 1 | 0.759 | 0.152 | 0.000 | 1900 | 11739 | 0.305 |
| 2 | 0.792 | 0.081 | 0.287 | 1500 | 2633 | 0.069 |
| 3 | 0.817 | 0.094 | 0.290 | 952 | 1554 | 0.040 |
| 4 | 1.410 | 0.119 | 5.568 | 763684 | 3795218 | 98.755 |
| 5 | 2.507 | 0.042 | 13.607 | 531 | 1082 | 0.028 |
| 6 | 2.791 | 0.087 | 4.386 | 9091 | 30844 | 0.803 |
| Total | | | | 777658 | 3843069 | 100.000 |

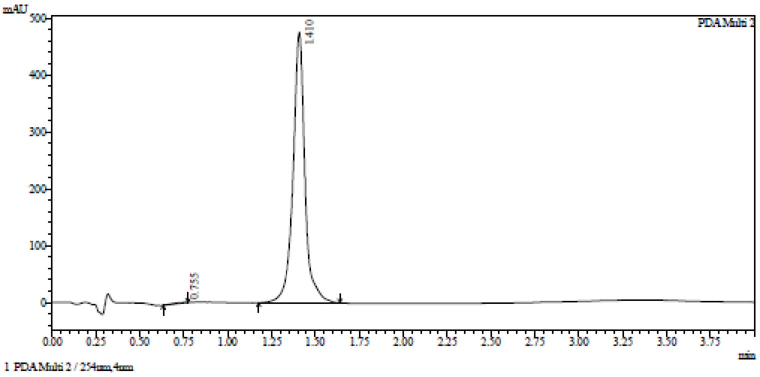

Integration result

PeakTable

PDA Ch2 254nm

| Peak# | Ret. Time | USP Width | Resolution | Height | Area | Area % |
|---|---|---|---|---|---|---|
| 1 | 0.755 | 0.156 | 0.000 | 771 | 6723 | 0.299 |
| 2 | 1.410 | 0.114 | 4.858 | 476220 | 2243465 | 99.701 |
| Total | | | | 476991 | 2250188 | 100.000 |

**HRMC** (2*R*,3*R*,4*S*,5*R*)-2-(6-aminopurin-9-yl)-5-(1,3-benzothiazol-2-yloxymethyl)tetrahydrofuran-3,4-diol **(23)**

Compound Table

| Label | Tgt Score | Mass Error (ppm) | Tgt Formula | Obs. RT | Ref. Mass | Obs. Mass |
|---|---|---|---|---|---|---|
| Cpd 1: C17 H16 N6 O4 S; 1.465 | 98.84 | 1.31 | C17 H16 N6 O4 S | 1.465 | 400.0954 | 400.0959 |

| Obs. *m/z* | Obs. RT | Obs. Mass | Tgt Formula | Tgt Mass | Tgt Mass Error (ppm) | Find Cpds Algorithm |
|---|---|---|---|---|---|---|
| 401.1031 | 1.465 | 400.0959 | C17 H16 N6 O4 S | 400.0954 | 1.31 | Find by Formula |

Compound Chromatograms

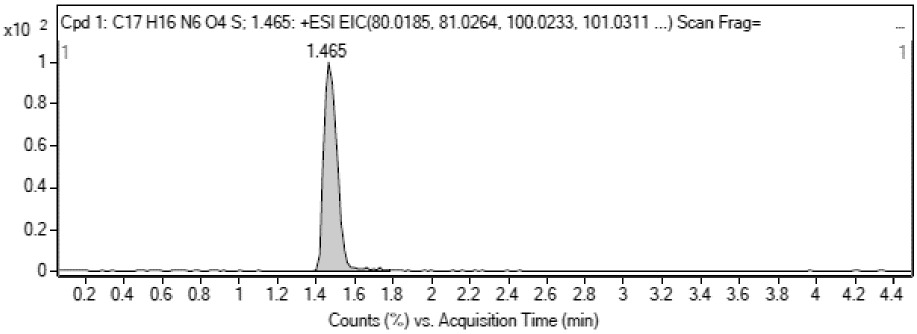

MS Zoomed Spectrum

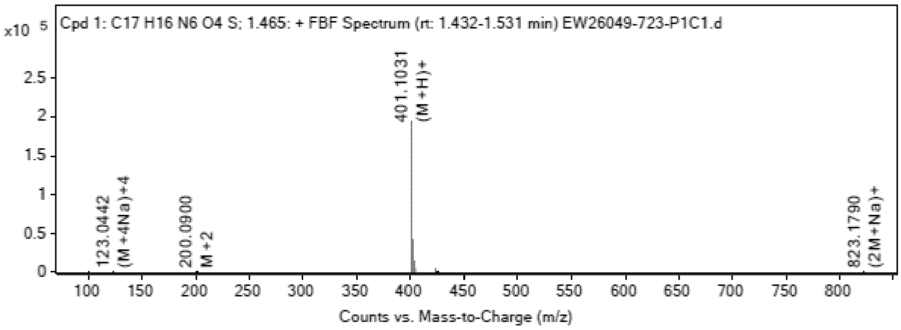

MS Spectrum Peak List

| Obs. *m/z* | Charge | Abund | Ion/Isotope |
|---|---|---|---|
| 123.0442 | 4 | 88.43 | (M+4Na)+4 |
| 200.09 | 2 | 86.06 | M+2 |

| | | | |
|---|---|---|---|
| 201.1037 | 2 | 70.64 | (M+2H)+2 |
| 401.1031 | 1 | 193524 | (M+H)+ |
| 402.1054 | 1 | 41576.28 | (M+H)+ |
| 403.1031 | 1 | 14133.26 | (M+H)+ |
| 423.0849 | 1 | 4287.03 | (M+Na)+ |
| 424.0883 | 1 | 1036.19 | (M+Na)+ |
| 425.0861 | 1 | 340.45 | (M+Na)+ |
| 823.179 | 1 | 99.71 | (2M+Na)+ |

**MS Zoomed Spectrum**

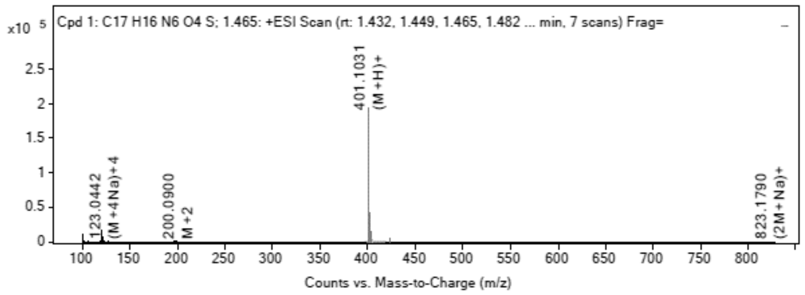

**MS Spectrum Peak List**

| Obs. *m/z* | Charge | Abund | Ion/Isotope | Tgt Mass Error (ppm) |
|---|---|---|---|---|
| 123.0442 | 4 | 88.43 | (M+4Na)+4 | -253.41 |
| 200.09 | 2 | 86.06 | M+2 | -214.23 |
| 201.1037 | 2 | 70.64 | (M+2H)+2 | -242.4 |
| 401.1031 | 1 | 193524 | (M+H)+ | -1.17 |
| 402.1054 | 1 | 41576.28 | (M+H)+ | -0.27 |
| 403.1031 | 1 | 14133.26 | (M+H)+ | -2.29 |
| 423.0849 | 1 | 4287.03 | (M+Na)+ | -0.78 |
| 424.0883 | 1 | 1036.19 | (M+Na)+ | -2.6 |
| 425.0861 | 1 | 340.45 | (M+Na)+ | -4.68 |
| 823.179 | 1 | 99.71 | (2M+Na)+ | 1.15 |

--- End Of Report ---

**$^{1}$H NMR** 2*R*,3*R*,4*S*,5*R*)-2-(6-Aminopurin-9-yl)-5-[(1,3-benzothiazol-2-ylamino)methyl] tetrahydrofuran-3,4-diol **(24)**

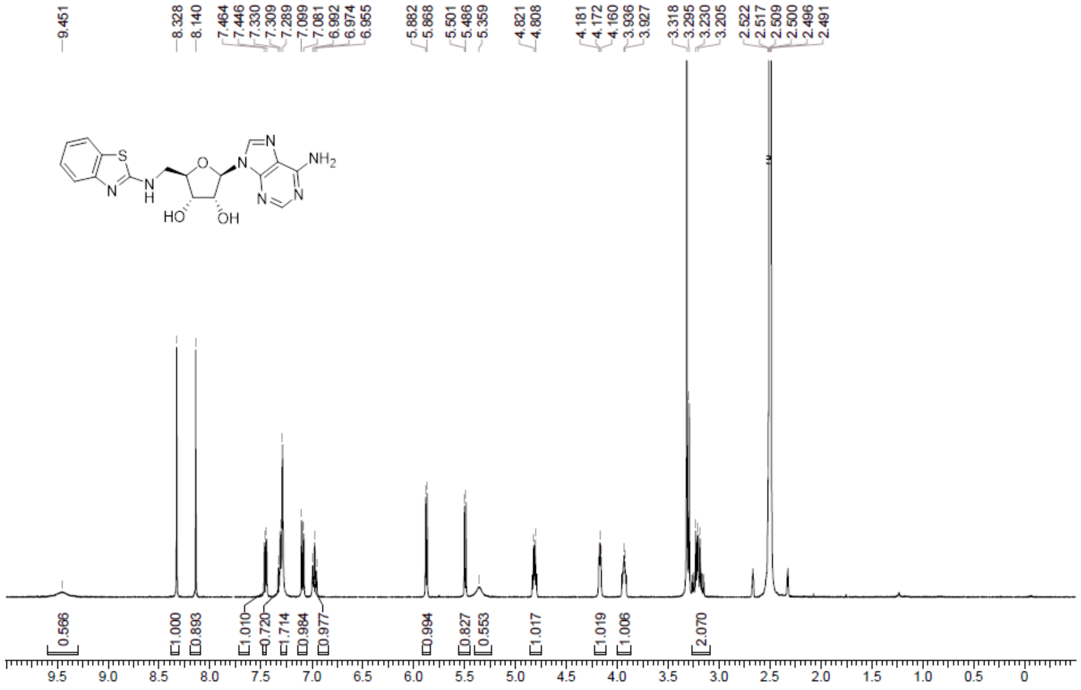

**HPLC** 2*R*,3*R*,4*S*,5*R*)-2-(6-Aminopurin-9-yl)-5-[(1,3-benzothiazol-2-ylamino)methyl] tetrahydrofuran-3,4-diol **(24)**

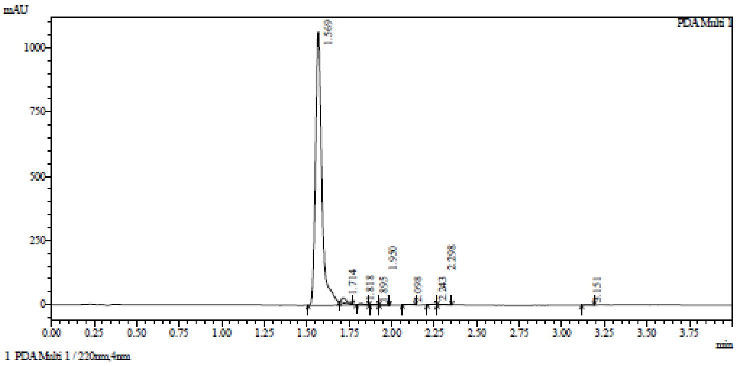

Integration result

PeakTable

PDA Ch1 220nm

| Peak# | Ret. Time | USP Width | Resolution | Height | Area | Area % |
|---|---|---|---|---|---|---|
| 1 | 1.569 | 0.066 | 0.000 | 1063984 | 2852502 | 97.489 |
| 2 | 1.714 | 0.053 | 2.433 | 18519 | 35180 | 1.202 |
| 3 | 1.818 | 0.055 | 1.902 | 5814 | 11667 | 0.399 |
| 4 | 1.895 | 0.051 | 1.450 | 1468 | 2728 | 0.093 |
| 5 | 1.950 | 0.049 | 1.098 | 663 | 1069 | 0.037 |
| 6 | 2.098 | 0.066 | 2.587 | 2040 | 4651 | 0.159 |
| 7 | 2.243 | 0.059 | 2.323 | 3383 | 6906 | 0.236 |
| 8 | 2.298 | 0.101 | 0.686 | 2277 | 7191 | 0.246 |
| 9 | 3.151 | 0.063 | 10.388 | 1743 | 4079 | 0.139 |
| Total | | | | 1099891 | 2925974 | 100.000 |

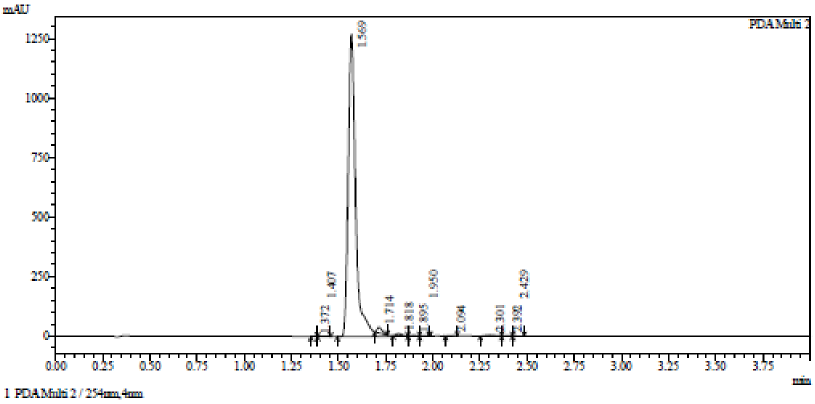

Integration result

PeakTable

PDA Ch2 254nm

| Peak# | Ret. Time | USP Width | Resolution | Height | Area | Area % |
|---|---|---|---|---|---|---|
| 1 | 1.372 | 0.052 | 0.000 | 705 | 1086 | 0.030 |
| 2 | 1.407 | 0.076 | 0.549 | 795 | 1746 | 0.048 |
| 3 | 1.569 | 0.068 | 2.267 | 1269004 | 3549920 | 97.422 |
| 4 | 1.714 | 0.051 | 2.442 | 25467 | 45291 | 1.243 |
| 5 | 1.818 | 0.057 | 1.920 | 8014 | 16255 | 0.446 |
| 6 | 1.895 | 0.050 | 1.445 | 2257 | 4291 | 0.118 |
| 7 | 1.950 | 0.044 | 1.165 | 747 | 1247 | 0.034 |
| 8 | 2.094 | 0.054 | 2.954 | 1593 | 3019 | 0.083 |
| 9 | 2.301 | 0.101 | 2.674 | 3897 | 14466 | 0.397 |
| 10 | 2.392 | 0.205 | 0.592 | 1664 | 4689 | 0.129 |
| 11 | 2.429 | 0.824 | 0.073 | 1111 | 1831 | 0.050 |
| Total | | | | 1315256 | 3643841 | 100.000 |

**HRMC** 2*R*,3*R*,4*S*,5*R*)-2-(6-Aminopurin-9-yl)-5-[(1,3-benzothiazol-2-ylamino)methyl] tetrahydrofuran-3,4-diol **(24)**

**Compound Table**

| Label | Tgt Score | Mass Error (ppm) | Tgt Formula | Obs. RT | Ref. Mass | Obs. Mass |
|---|---|---|---|---|---|---|
| Cpd 1: C17 H17 N7 O3 S; 1.595 | 98.29 | 2.05 | C17 H17 N7 O3 S | 1.595 | 399.1114 | 399.1122 |

| Obs. m/z | Obs. RT | Obs. Mass | Tgt Formula | Tgt Mass | Tgt Mass Error (ppm) | Find Cpds Algorithm |
|---|---|---|---|---|---|---|
| 422.1002 | 1.595 | 399.1122 | C17 H17 N7 O3 S | 399.1114 | 2.05 | Find by Formula |

**Compound Chromatograms**

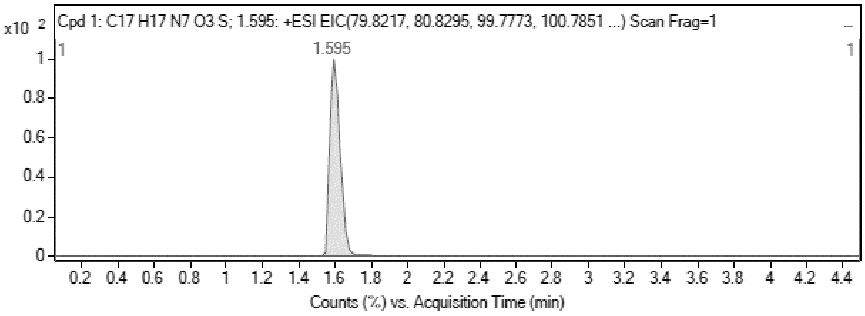

MS Zoomed Spectrum

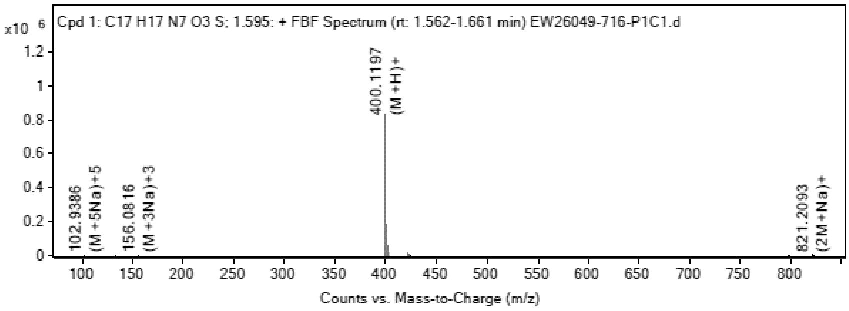

**MS Spectrum Peak List**

| Obs. m/z | Charge | Abund | Ion/Isotope |
|---|---|---|---|
| 399.1116 | 1 | 7979.15 | M+ |
| 400.1197 | 1 | 831777.56 | (M+H)+ |
| 401.1214 | 1 | 183561.86 | (M+H)+ |
| 402.1183 | 1 | 57424.29 | (M+H)+ |
| 422.1002 | 1 | 22912.8 | (M+Na)+ |
| 423.1037 | 1 | 5037.99 | (M+Na)+ |
| 424.1011 | 1 | 1867.23 | (M+Na)+ |
| 798.216 | 1 | 1916.53 | 2M+ |
| 821.2093 | 1 | 5269.97 | (2M+Na)+ |
| 822.2124 | 1 | 2410.98 | (2M+Na)+ |

MS Zoomed Spectrum

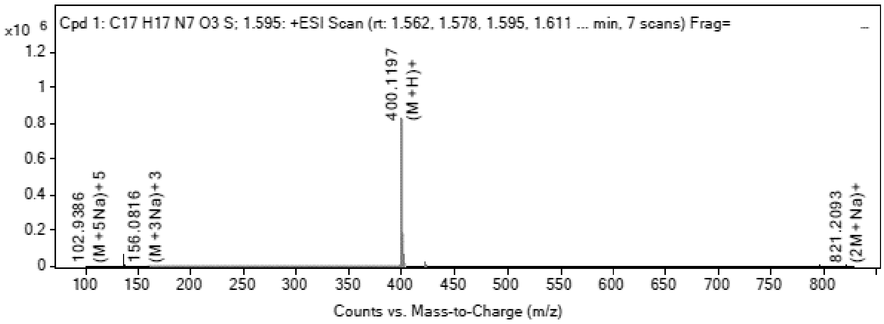

**MS Spectrum Peak List**

| Obs. m/z | Charge | Abund | Ion/Isotope | Tgt Mass Error (ppm) |
|---|---|---|---|---|
| 399.1116 | 1 | 7979.15 | M+ | -2.06 |
| 400.1197 | 1 | 831777.56 | (M+H)+ | -2.54 |
| 401.1214 | 1 | 183561.86 | (M+H)+ | -0.65 |
| 402.1183 | 1 | 57424.29 | (M+H)+ | -0.79 |
| 422.1002 | 1 | 22912.8 | (M+Na)+ | 0.84 |
| 423.1037 | 1 | 5037.99 | (M+Na)+ | -1.34 |
| 424.1011 | 1 | 1867.23 | (M+Na)+ | -2.79 |
| 798.216 | 1 | 1916.53 | 2M+ | 7.71 |
| 821.2093 | 1 | 5269.97 | (2M+Na)+ | 3.19 |
| 822.2124 | 1 | 2410.98 | (2M+Na)+ | 2.44 |

--- End Of Report ---

**[1]H NMR** 6-Methylquinoline-2-thiol **(73)**

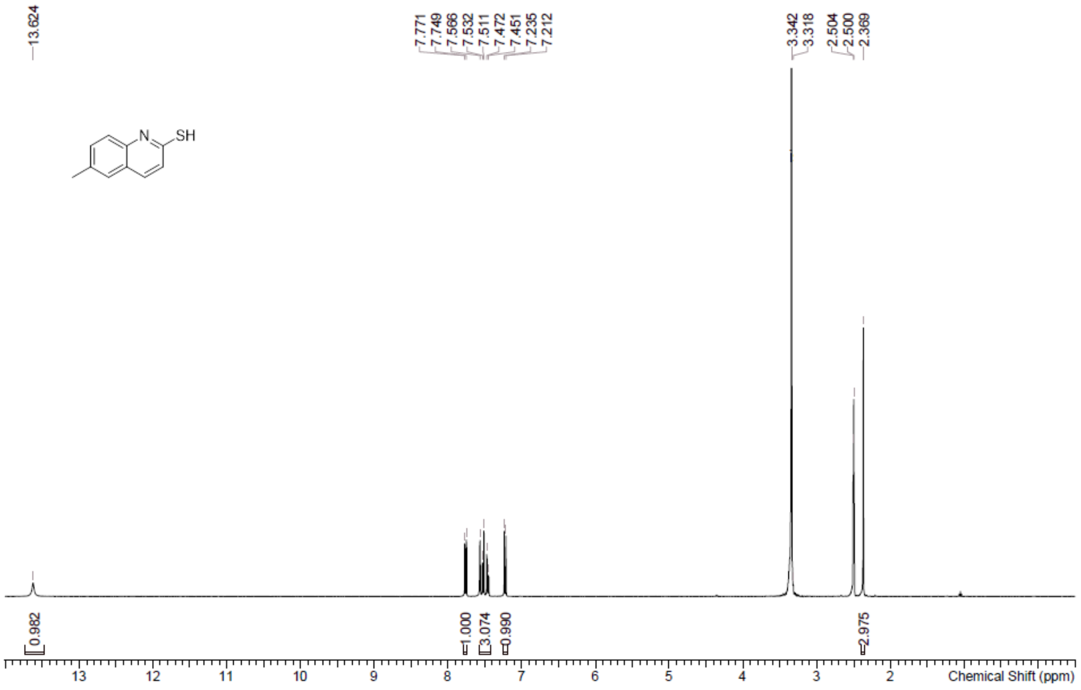

**LCMS** 6-Methylquinoline-2-thiol **(73)**

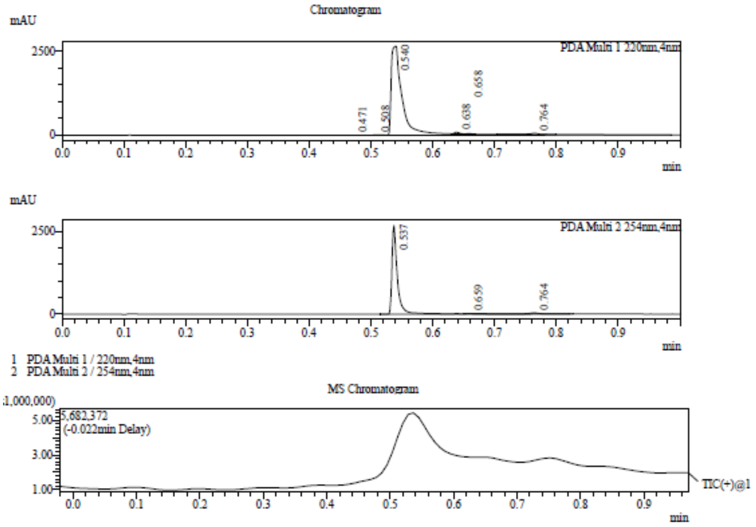

Integration Result

Peak Table

PDA Ch1 220nm

| Peak# | Ret. Time | Height | Height% | USP Width | Area | Area% |
|---|---|---|---|---|---|---|
| 1 | 0.471 | 3720 | 0.134 | 0.069 | 10372 | 0.280 |
| 2 | 0.508 | 14380 | 0.517 | 0.021 | 11374 | 0.307 |
| 3 | 0.540 | 2661842 | 95.668 | 0.027 | 3610215 | 97.363 |
| 4 | 0.638 | 49637 | 1.784 | 0.016 | 25549 | 0.689 |
| 5 | 0.658 | 14118 | 0.507 | 0.015 | 6852 | 0.185 |
| 6 | 0.764 | 38669 | 1.390 | 0.026 | 43637 | 1.177 |

Peak Table

PDA Ch2 254nm

| Peak# | Ret. Time | Height | Height% | USP Width | Area | Area% |
|---|---|---|---|---|---|---|
| 1 | 0.537 | 2687593 | 98.323 | 0.016 | 1699781 | 97.208 |
| 2 | 0.659 | 9647 | 0.353 | 0.016 | 5595 | 0.320 |
| 3 | 0.764 | 36192 | 1.324 | 0.026 | 43228 | 2.472 |

Mass Spectrum

RetTime: 0.470 Datafile: D:\DATA\2022\2209\220921\EW26049-1048-P1D1.lcd

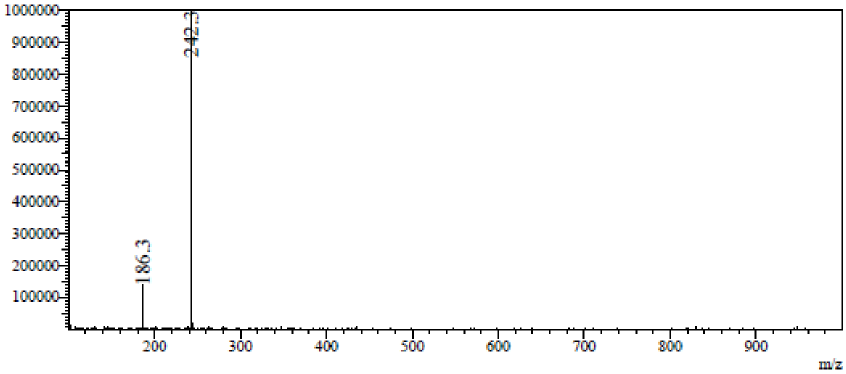

RetTime: 0.512 Datafile: D:\DATA\2022\2209\220921\EW26049-1048-P1D1.lcd

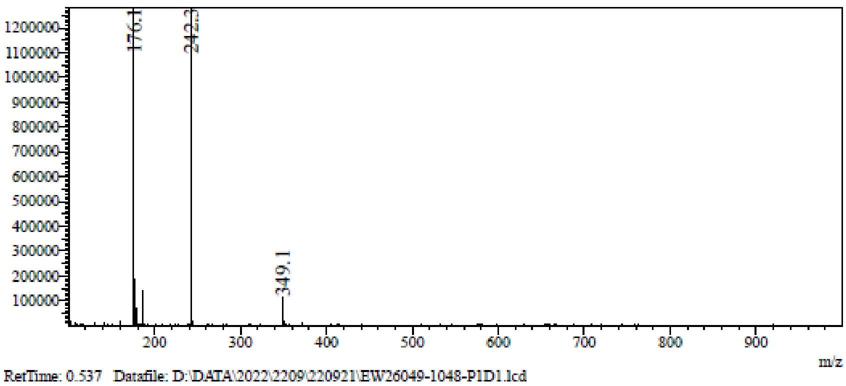

RetTime: 0.537 Datafile: D:\DATA\2022\2209\220921\EW26049-1048-P1D1.lcd

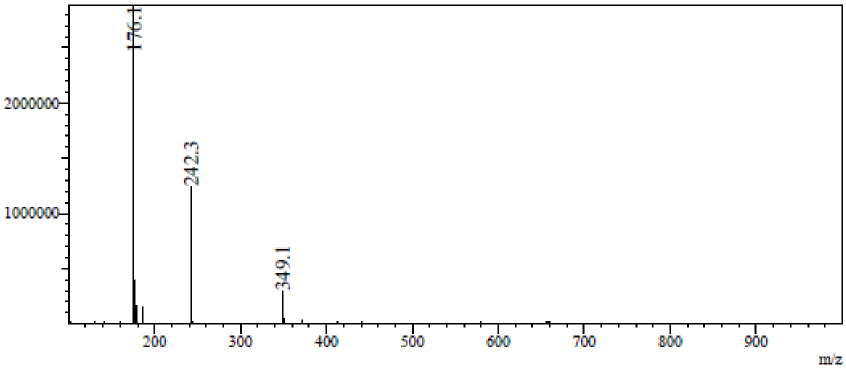

RetTime: 0.637 Datafile: D:\DATA\2022\2209\220921\EW26049-1048-P1D1.lcd

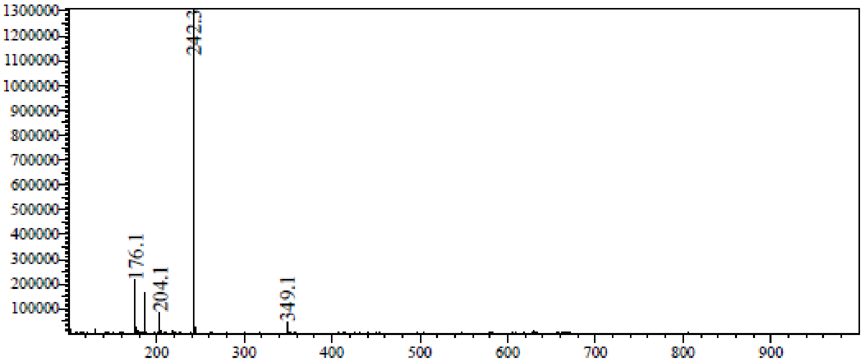

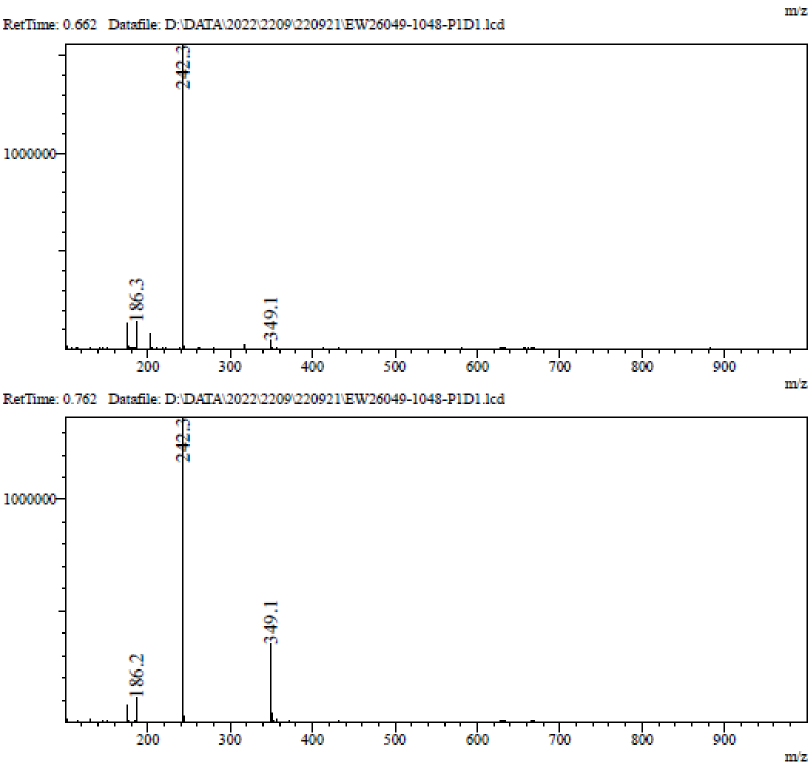

**$^1$H NMR** 7-Methyl-1-oxido-quinolin-1-ium **(75)**

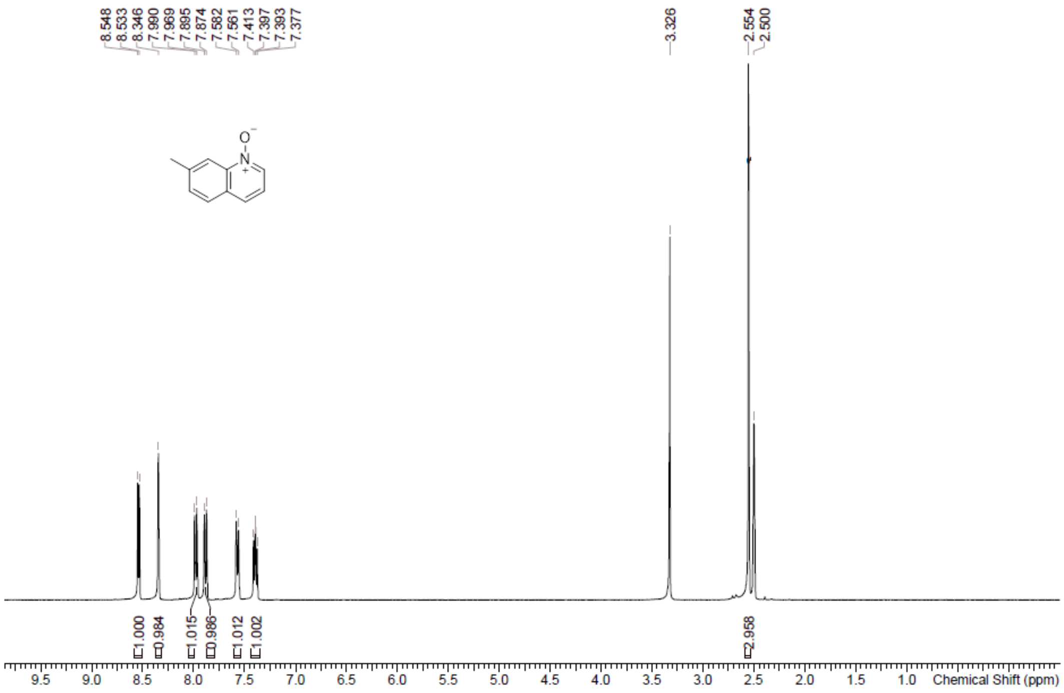

**LCMS** 7-Methyl-1-oxido-quinolin-1-ium **(75)**

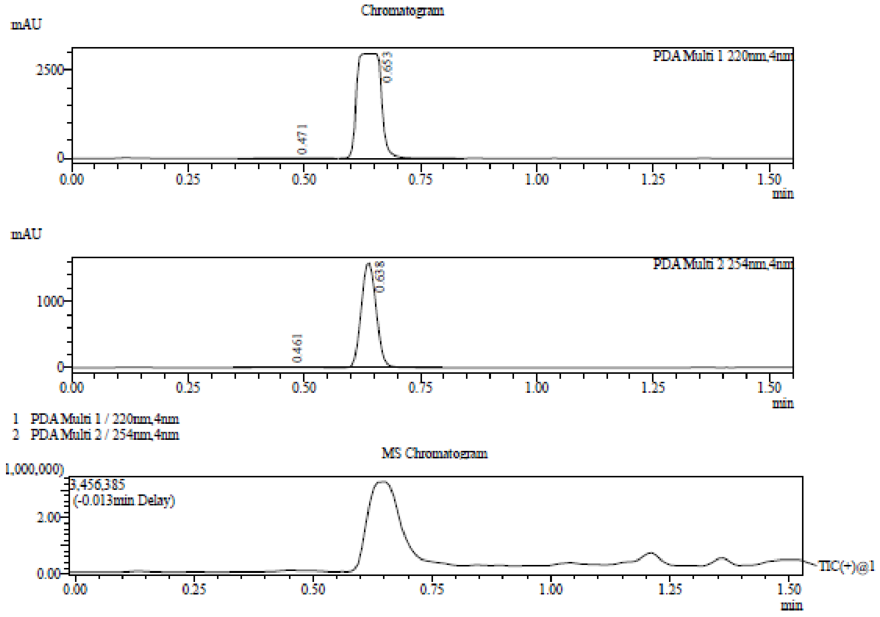

Integration Result

Peak Table

PDA Ch1 220nm

| Peak# | Ret. Time | Height | Height% | USP Width | Area | Area% |
|---|---|---|---|---|---|---|
| 1 | 0.471 | 11394 | 0.384 | 0.181 | 73212 | 0.688 |
| 2 | 0.653 | 2959336 | 99.616 | 0.080 | 10567329 | 99.312 |

Peak Table

PDA Ch2 254nm

| Peak# | Ret. Time | Height | Height% | USP Width | Area | Area% |
|---|---|---|---|---|---|---|
| 1 | 0.461 | 4803 | 0.304 | 0.181 | 30554 | 0.859 |
| 2 | 0.638 | 1573636 | 99.696 | 0.068 | 3524642 | 99.141 |

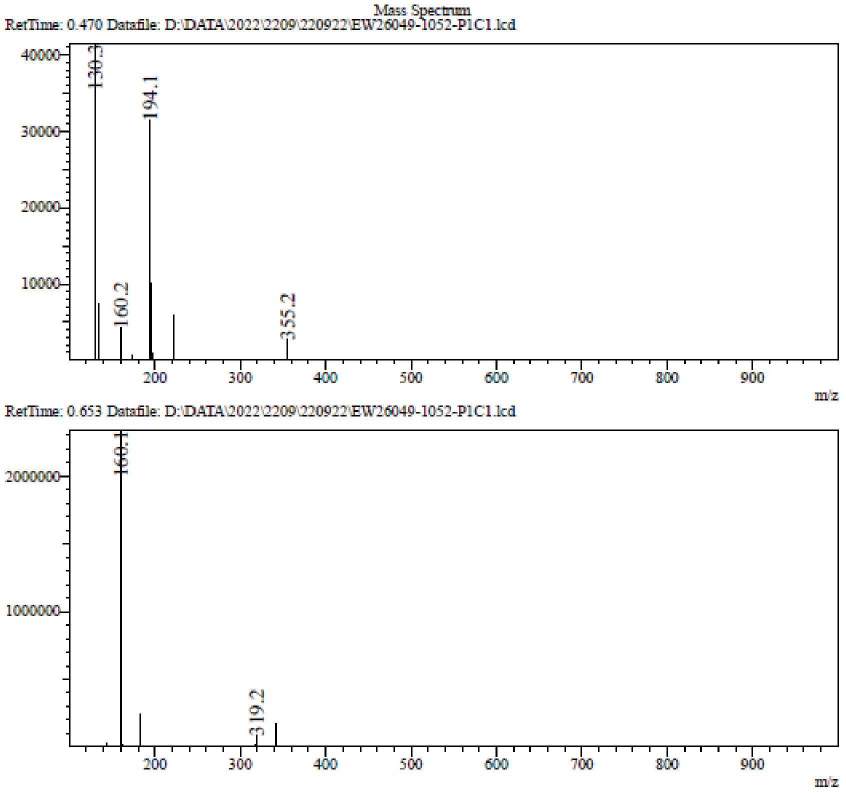

**$^1$H NMR** 2-Chloro-7-methyl-quinoline **(76)**

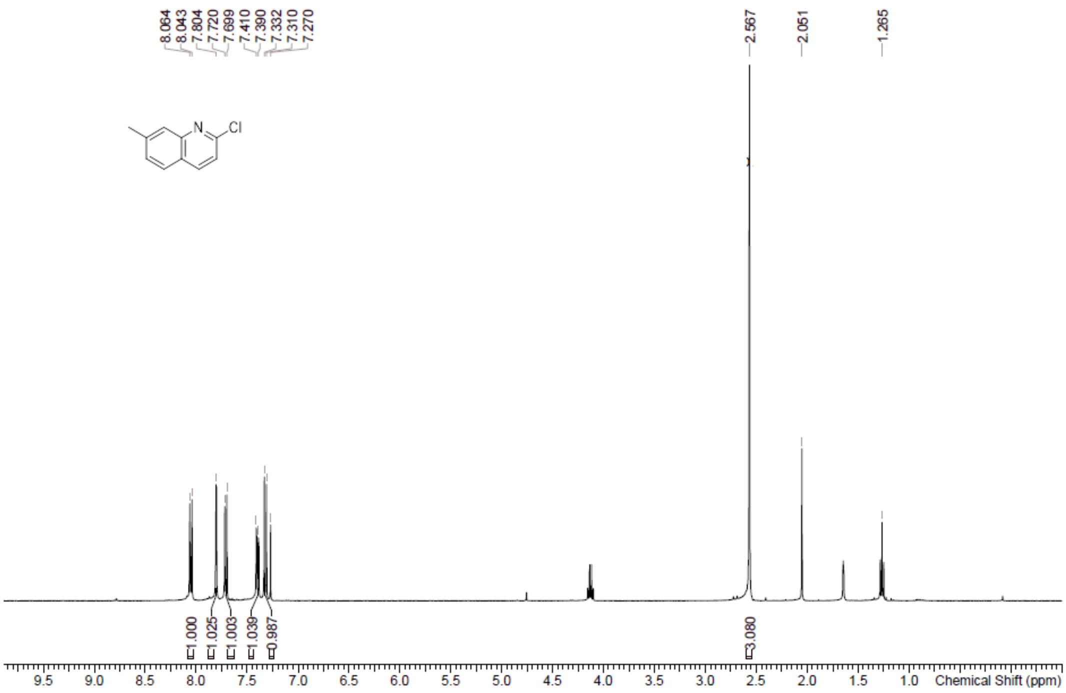

**LCMS** 2-Chloro-7-methyl-quinoline **(76)**

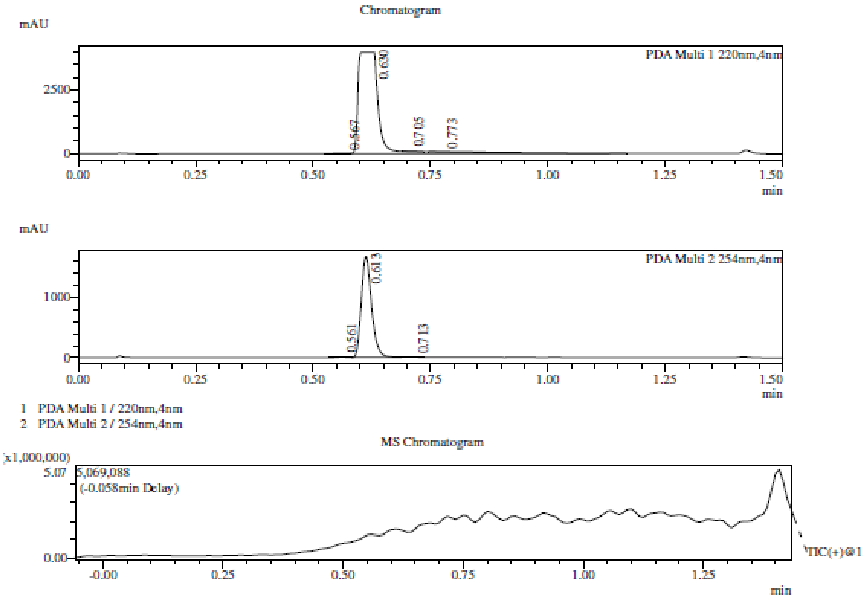

Integration Result

Peak Table

PDA Ch1 220nm

| Peak# | Ret. Time | Height | Height% | USP Width | Area | Area% |
|---|---|---|---|---|---|---|
| 1 | 0.567 | 13962 | 0.347 | 0.057 | 22856 | 0.187 |
| 2 | 0.630 | 3986957 | 99.198 | 0.056 | 12179344 | 99.510 |
| 3 | 0.705 | 14429 | 0.359 | 0.037 | 18133 | 0.148 |
| 4 | 0.773 | 3833 | 0.095 | 0.117 | 18990 | 0.155 |

Peak Table

PDA Ch2 254nm

| Peak# | Ret. Time | Height | Height% | USP Width | Area | Area% |
|---|---|---|---|---|---|---|
| 1 | 0.561 | 6277 | 0.373 | 0.037 | 8222 | 0.323 |
| 2 | 0.613 | 1675305 | 99.438 | 0.041 | 2528900 | 99.403 |
| 3 | 0.713 | 3188 | 0.189 | 0.084 | 6962 | 0.274 |

Mass Spectrum

RetTime: 0.567   Datafile: D:\Data\2022\2209\220927\EW26049-1054-P1C1.lcd

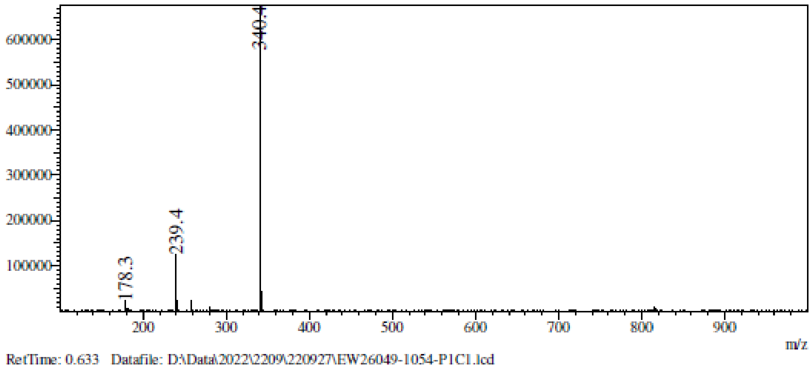

RetTime: 0.633   Datafile: D:\Data\2022\2209\220927\EW26049-1054-P1C1.lcd

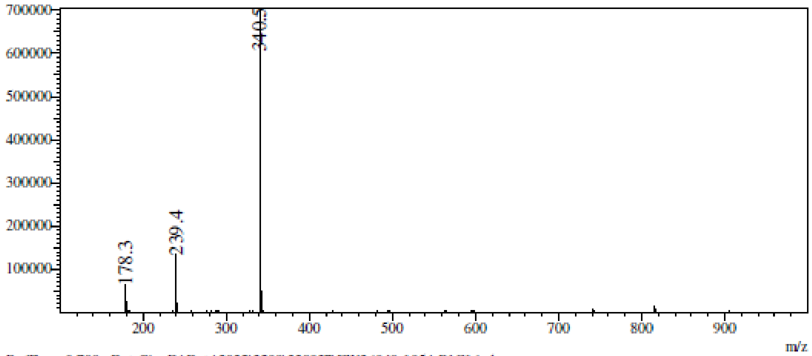

RetTime: 0.708   Datafile: D:\Data\2022\2209\220927\EW26049-1054-P1C1.lcd

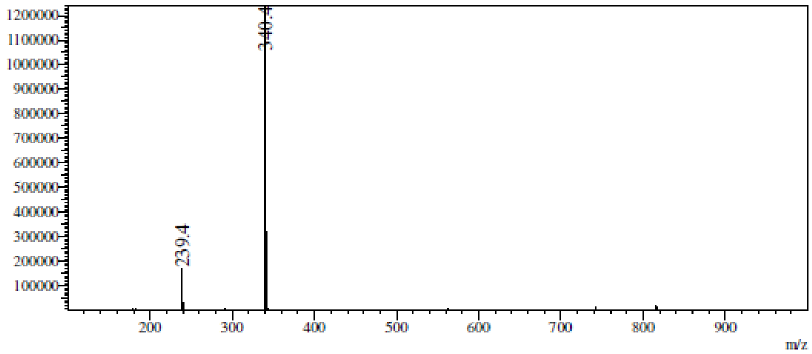

RetTime: 0.775   Datafile: D:\Data\2022\2209\220927\EW26049-1054-P1C1.lcd

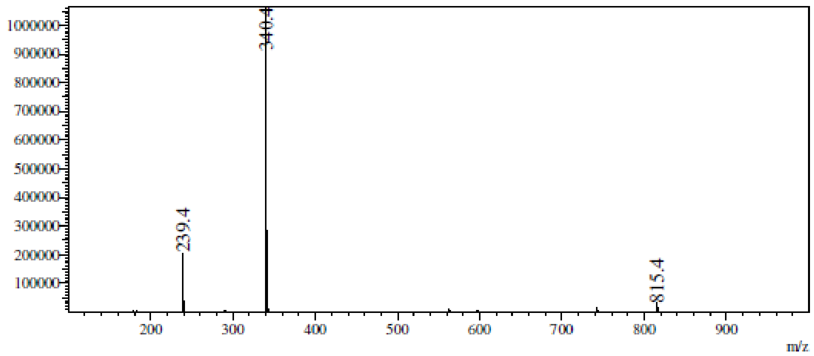

**[1]H NMR** 7-Methylquinoline-2-thiol **(77)**

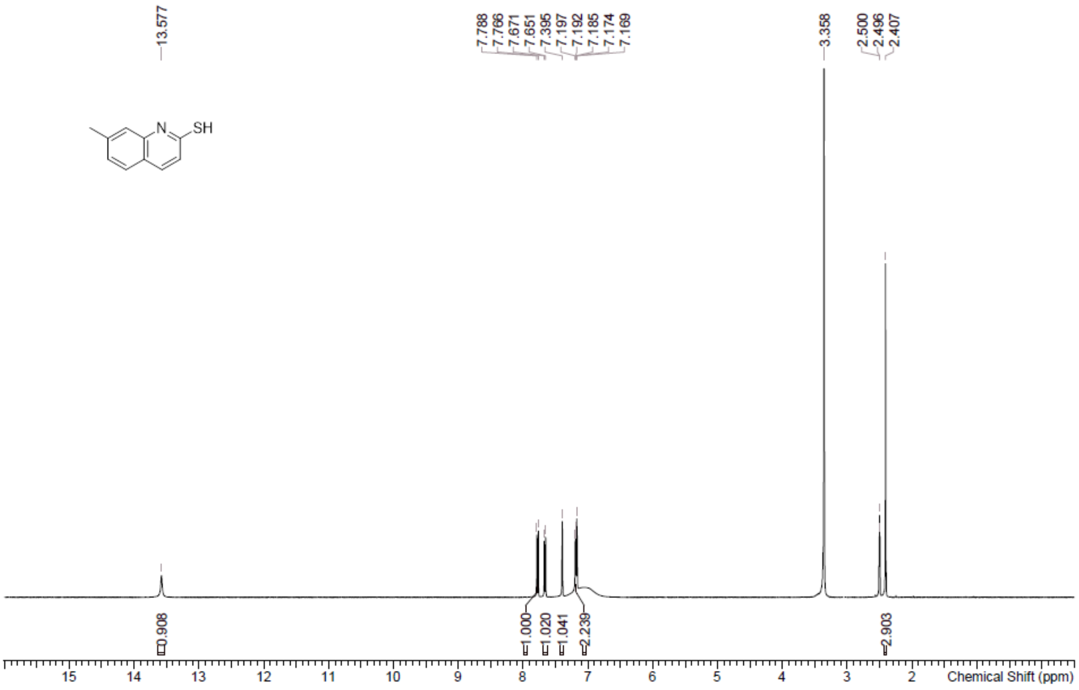

#### **LCMS** 7-Methylquinoline-2-thiol **(77)**

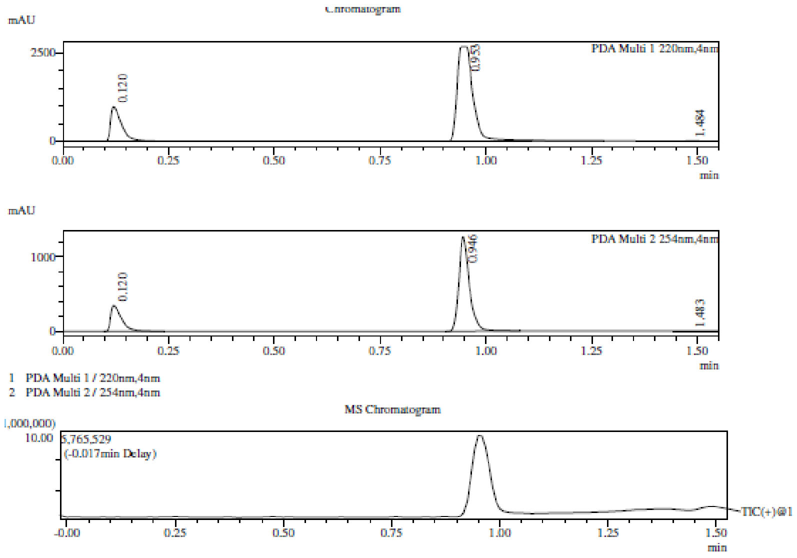

Integration Result

Peak Table

PDA Ch1 220nm

| Peak# | Ret. Time | Height | Height% | USP Width | Area | Area% |
|---|---|---|---|---|---|---|
| 1 | 0.120 | 981276 | 26.708 | 0.047 | 1764832 | 20.097 |
| 2 | 0.953 | 2689178 | 73.192 | 0.056 | 7008323 | 79.806 |
| 3 | 1.484 | 3677 | 0.100 | 0.067 | 8559 | 0.097 |

Peak Table

PDA Ch2 254nm

| Peak# | Ret. Time | Height | Height% | USP Width | Area | Area% |
|---|---|---|---|---|---|---|
| 1 | 0.120 | 346312 | 21.394 | 0.047 | 616222 | 21.838 |
| 2 | 0.946 | 1268178 | 78.343 | 0.045 | 2191839 | 77.676 |
| 3 | 1.483 | 4260 | 0.263 | 0.082 | 13710 | 0.486 |

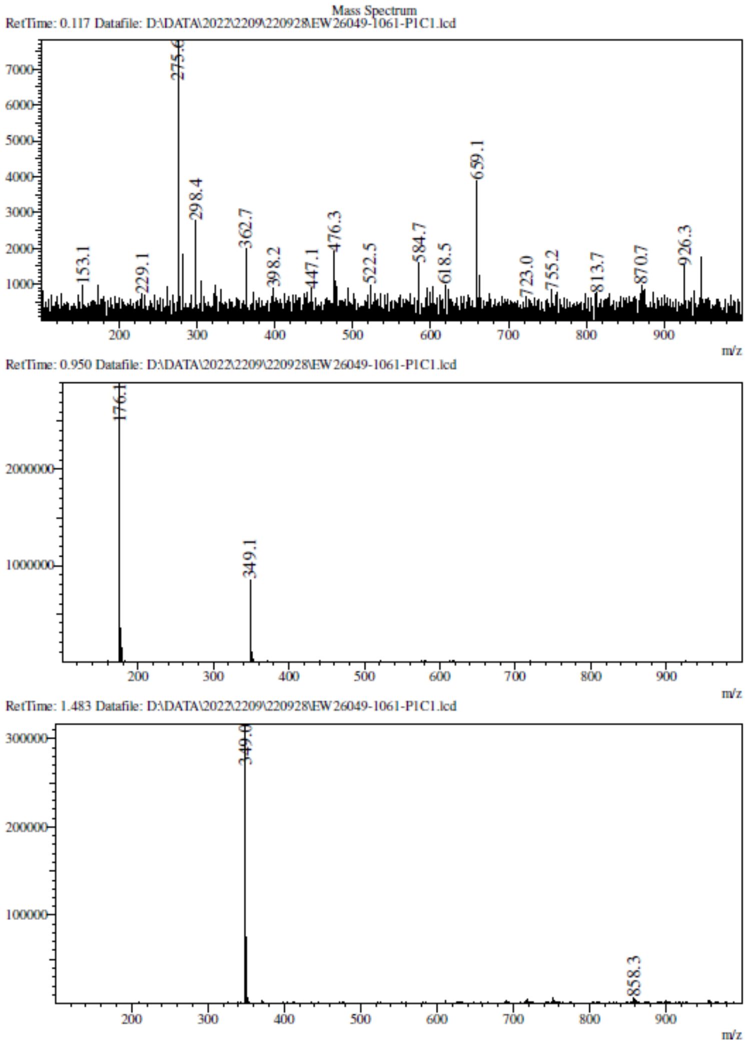
Mass Spectrum
RetTime: 0.117 Datafile: D:\DATA\2022\2209\220928\EW26049-1061-P1C1.lcd
275.6
659.1
298.4
362.7
476.3
584.7
153.1
229.1
398.2
447.1
522.5
618.5
723.0
755.2
813.7
870.7
926.3
m/z
RetTime: 0.950 Datafile: D:\DATA\2022\2209\220928\EW26049-1061-P1C1.lcd
176.1
349.1
m/z
RetTime: 1.483 Datafile: D:\DATA\2022\2209\220928\EW26049-1061-P1C1.lcd
349.0
858.3
m/z

**¹H NMR** 8-[(4-methoxyphenyl)methylsulfanyl]imidazo[1,5-a]pyridine **(79)**

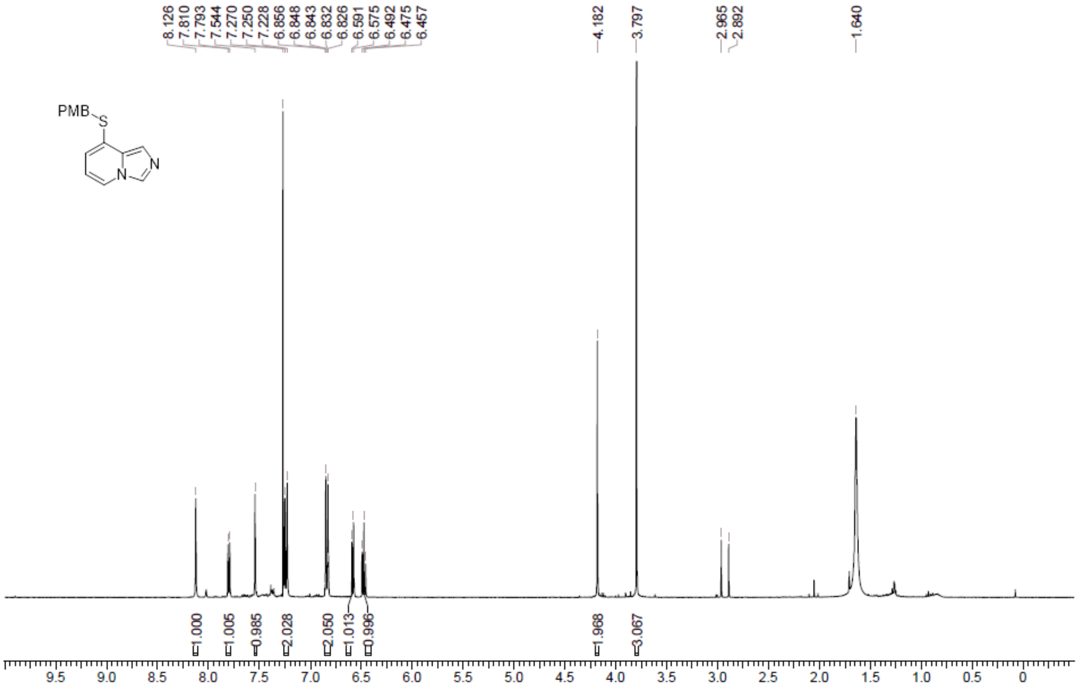

**LCMS** 8-[(4-methoxyphenyl)methylsulfanyl]imidazo[1,5-a]pyridine **(79)**

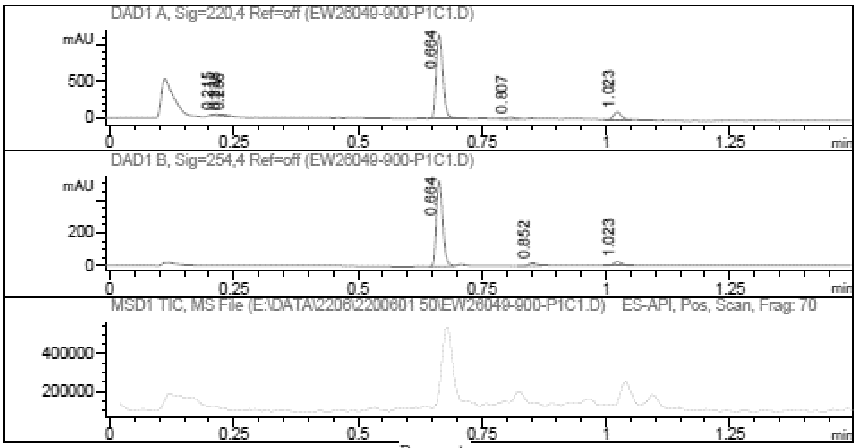

Report

Signal 1 : DAD1 A, Sig=220,4 Ref=off

| # | Meas. Ret. | Height | Width | Area | Area % |
|---|---|---|---|---|---|
| 1 | 0.215 | 20.078 | 0.016 | 22.199 | 1.759 |
| 2 | 0.228 | 19.674 | 0.008 | 9.549 | 0.757 |
| 3 | 0.236 | 15.687 | 0.008 | 7.158 | 0.567 |
| 4 | 0.664 | 1152.937 | 0.014 | 1074.950 | 85.184 |
| 5 | 0.807 | 22.970 | 0.015 | 22.046 | 1.747 |
| 6 | 1.023 | 111.341 | 0.017 | 126.014 | 9.986 |

Signal 2 : DAD1 B, Sig=254,4 Ref=off

| # | Meas. Ret. | Height | Width | Area | Area % |
|---|---|---|---|---|---|
| 1 | 0.664 | 521.269 | 0.014 | 465.063 | 91.003 |
| 2 | 0.852 | 17.640 | 0.017 | 19.793 | 3.873 |
| 3 | 1.023 | 24.565 | 0.016 | 26.185 | 5.124 |

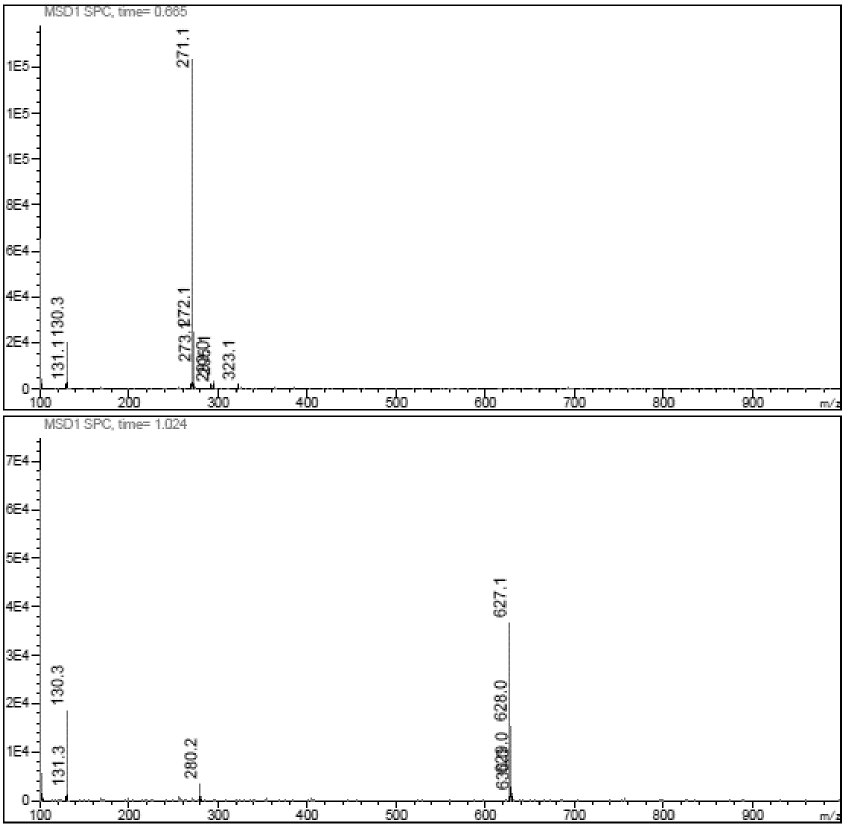

**$^1$H NMR** Imidazo[1,5-a]pyridine-8-thiol **(80)**

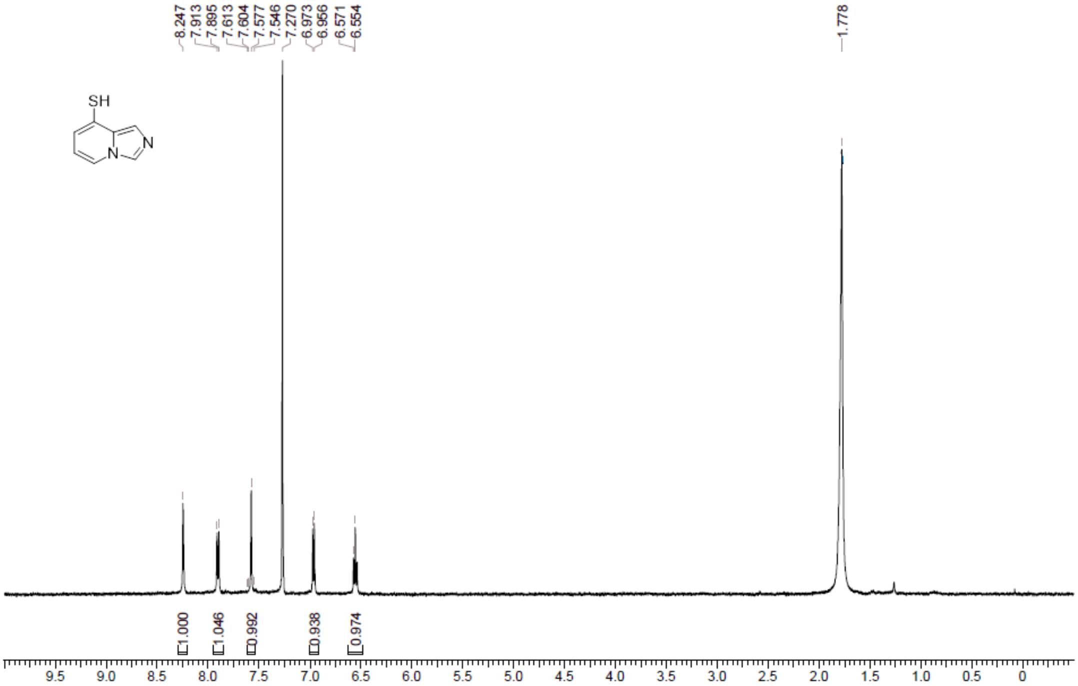

**[1]H NMR** 7-Bromo-1-((2-(trimethylsilyl)ethoxy)methyl)-1H-indazole **(82)**

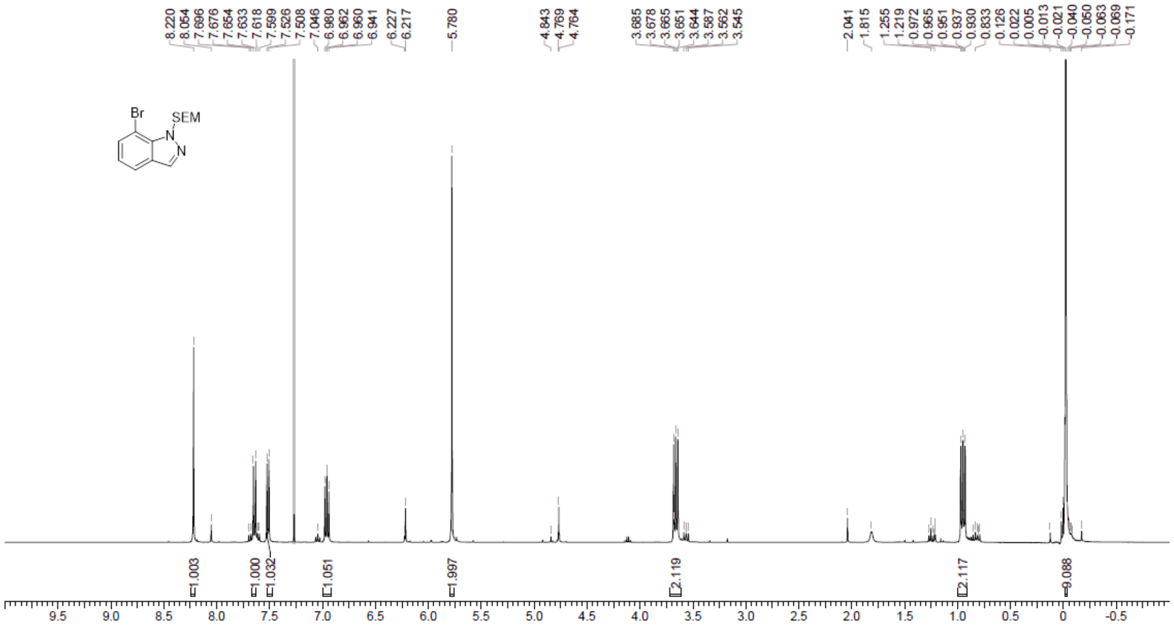

**LCMS** 7-Bromo-1-((2-(trimethylsilyl)ethoxy)methyl)-1H-indazole **(82)**

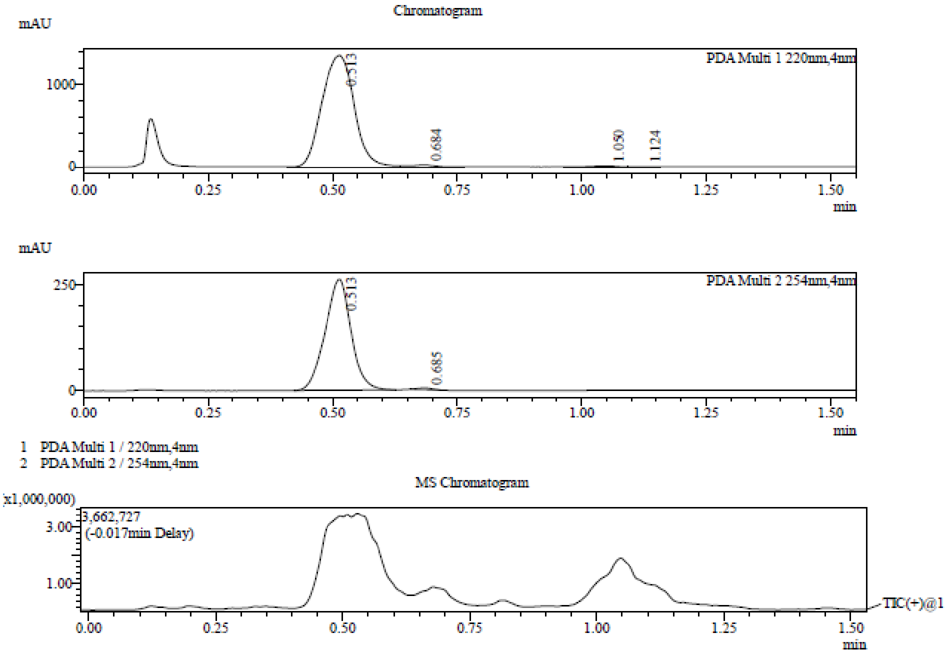

Integration Result

Peak Table

PDA Ch1 220nm

| Peak# | Ret. Time | Height | Height% | USP Width | Area | Area% |
|---|---|---|---|---|---|---|
| 1 | 0.513 | 1348759 | 97.208 | 0.114 | 6057321 | 97.814 |
| 2 | 0.684 | 22531 | 1.624 | 0.243 | 81431 | 1.315 |
| 3 | 1.050 | 12340 | 0.889 | 0.094 | 44505 | 0.719 |
| 4 | 1.124 | 3866 | 0.279 | 0.106 | 9442 | 0.152 |

Peak Table

PDA Ch2 254nm

| Peak# | Ret. Time | Height | Height% | USP Width | Area | Area% |
|---|---|---|---|---|---|---|
| 1 | 0.513 | 260850 | 98.559 | 0.097 | 949349 | 99.188 |
| 2 | 0.685 | 3814 | 1.441 | 0.059 | 7770 | 0.812 |

Mass Spectrum

RetTime: 0.513 Datafile: D:\DATA\2022\2206\220610\EW34712-79-P1A2.lcd

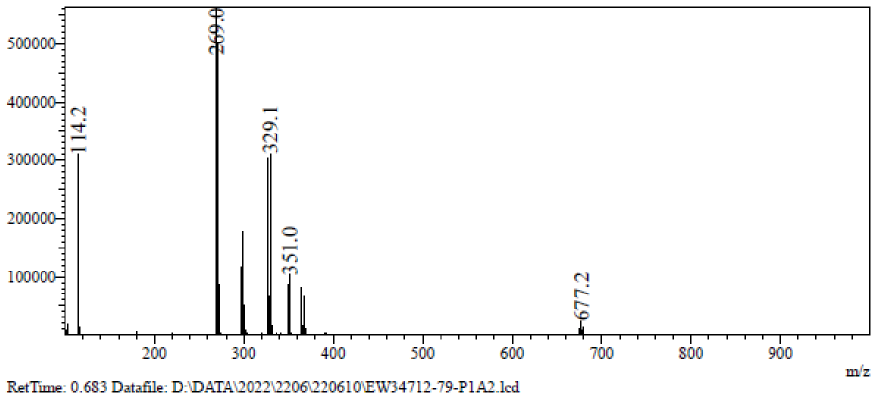

RetTime: 0.683 Datafile: D:\DATA\2022\2206\220610\EW34712-79-P1A2.lcd

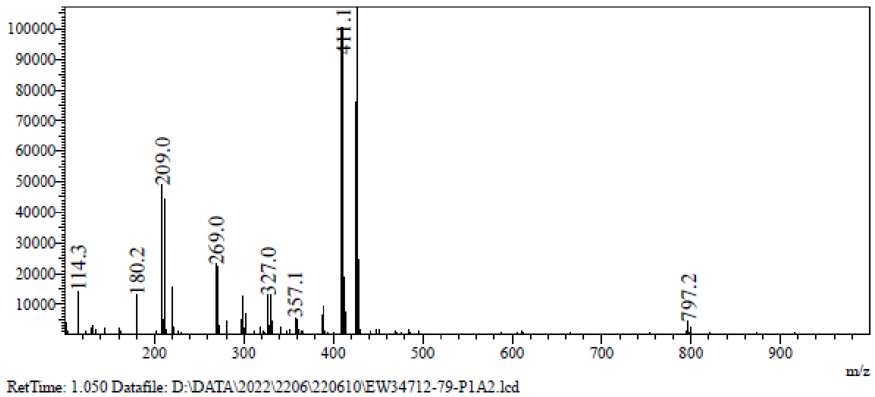

RetTime: 1.050 Datafile: D:\DATA\2022\2206\220610\EW34712-79-P1A2.lcd

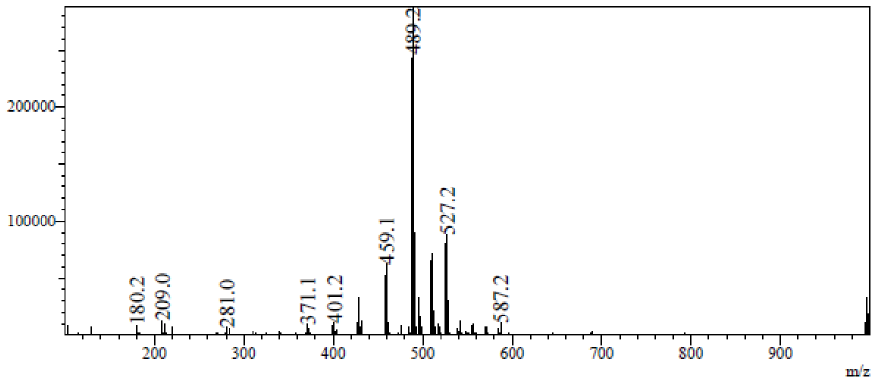

RetTime: 1.123 Datafile: D:\DATA\2022\2206\220610\EW34712-79-P1A2.lcd

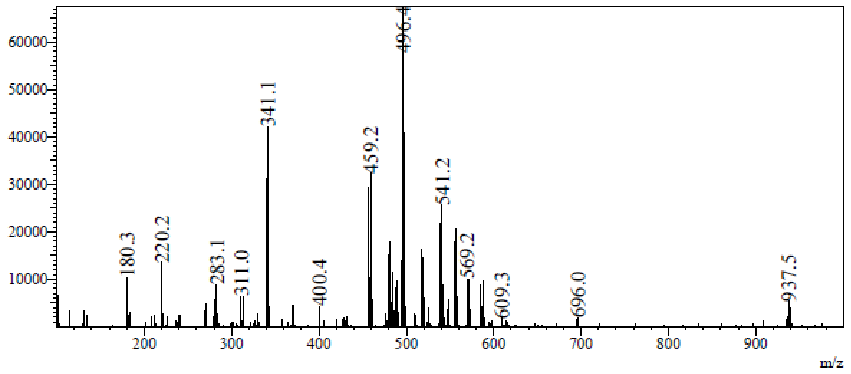

**LCMS** S7-((4-Methoxybenzyl)thio)-1H-indazole **(84)**

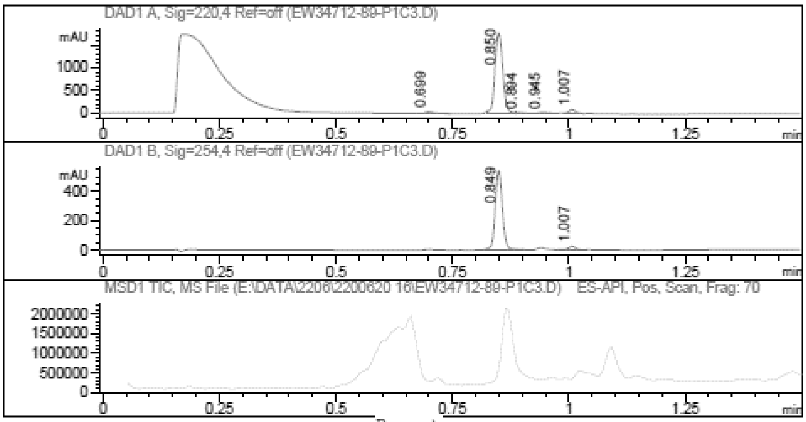

Report

Signal 1 : DAD1 A, Sig=220,4 Ref=off

| # | Meas. Ret. | Height | Width | Area | Area % |
|---|---|---|---|---|---|
| 1 | 0.699 | 35.876 | 0.018 | 44.023 | 1.989 |
| 2 | 0.850 | 1777.898 | 0.018 | 2030.185 | 91.741 |
| 3 | 0.894 | 20.543 | 0.016 | 23.058 | 1.042 |
| 4 | 0.945 | 23.152 | 0.021 | 31.548 | 1.426 |
| 5 | 1.007 | 83.924 | 0.015 | 84.138 | 3.802 |

Signal 2 : DAD1 B, Sig=254,4 Ref=off

| # | Meas. Ret. | Height | Width | Area | Area % |
|---|---|---|---|---|---|
| 1 | 0.849 | 543.465 | 0.016 | 572.278 | 95.748 |
| 2 | 1.007 | 24.332 | 0.016 | 25.413 | 4.252 |

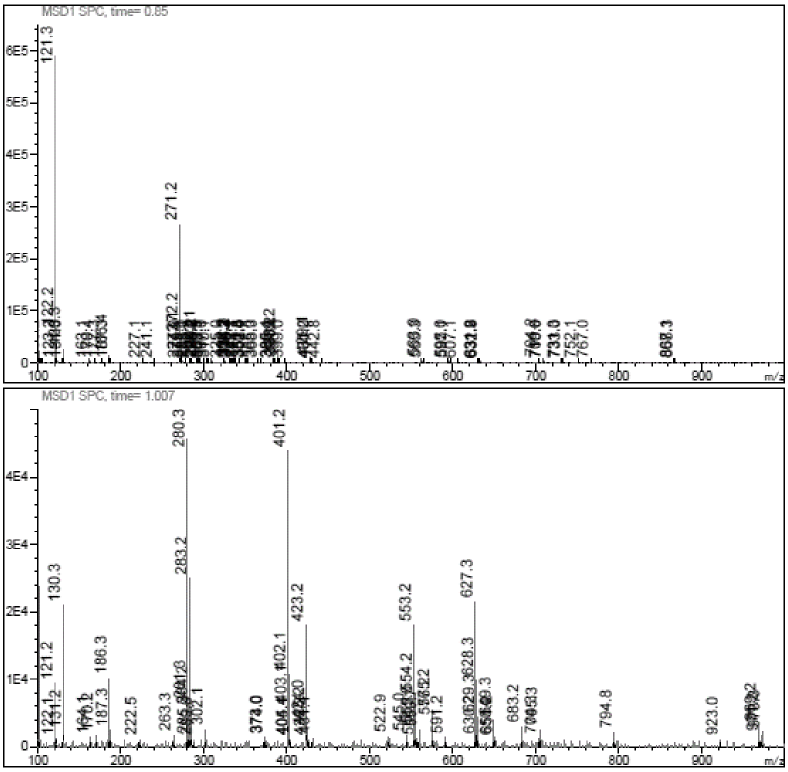

[1]**H NMR** 1H-Indazole-7-thiol **(85)**

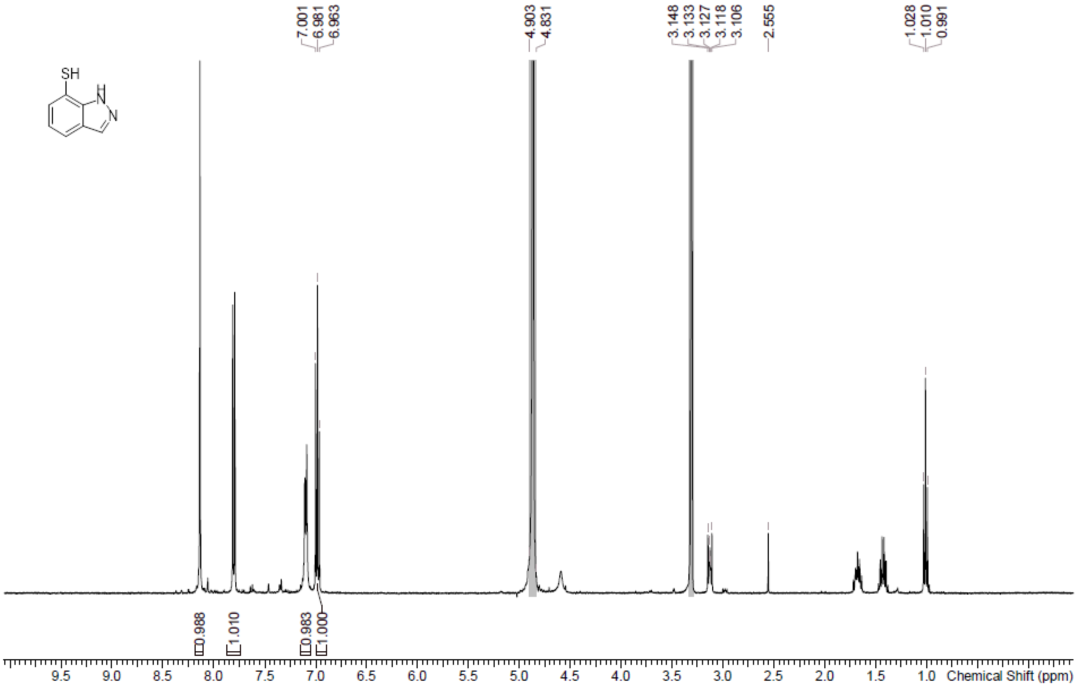

**LCMS** 1H-Indazole-7-thiol **(85)**

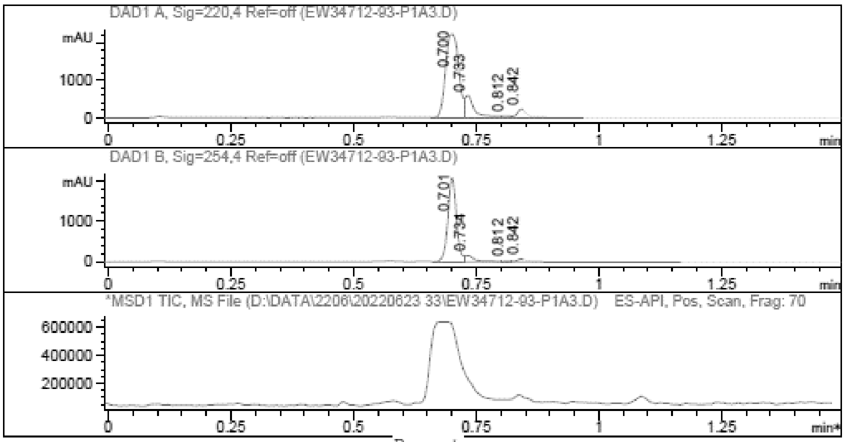

Report

Signal 1 : DAD1 A, Sig=220,4 Ref=off

| # | Meas. Ret. | Height | Width | Area | Area % |
|---|---|---|---|---|---|
| 1 | 0.700 | 2242.561 | 0.030 | 4175.400 | 76.974 |
| 2 | 0.733 | 594.393 | 0.020 | 825.402 | 15.216 |
| 3 | 0.812 | 51.604 | 0.016 | 57.294 | 1.056 |
| 4 | 0.842 | 230.986 | 0.022 | 366.334 | 6.753 |

Signal 2 : DAD1 B, Sig=254,4 Ref=off

| # | Meas. Ret. | Height | Width | Area | Area % |
|---|---|---|---|---|---|
| 1 | 0.701 | 2089.253 | 0.019 | 2522.999 | 83.574 |
| 2 | 0.734 | 157.495 | 0.021 | 230.486 | 7.635 |
| 3 | 0.812 | 20.125 | 0.017 | 23.690 | 0.785 |
| 4 | 0.842 | 74.143 | 0.042 | 241.720 | 8.007 |

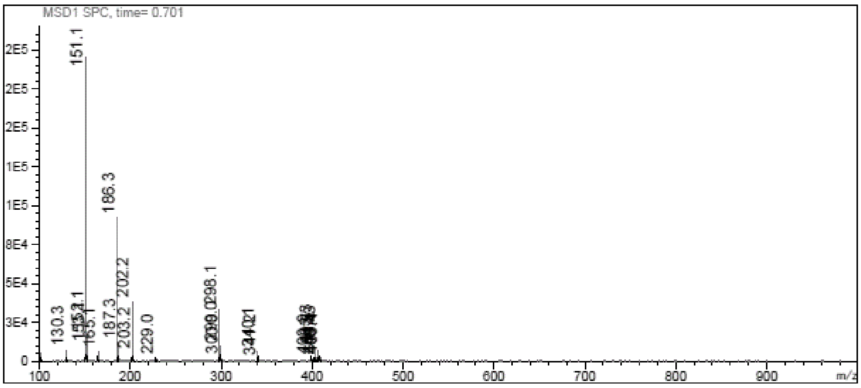
MSD1 SPC, time= 0.701
151.1
186.3
202.2
298.1
130.3
187.3
203.2
229.0
m/z

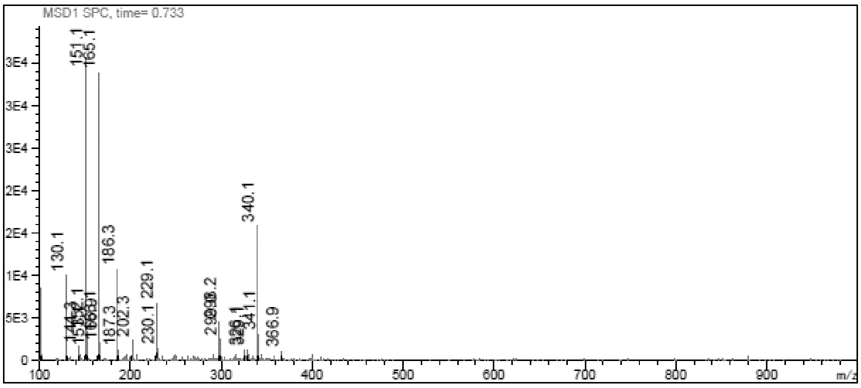
MSD1 SPC, time= 0.733
151.1
165.1
340.1
130.1
186.3
229.1
187.3
202.3
230.1
341.1
366.9
m/z

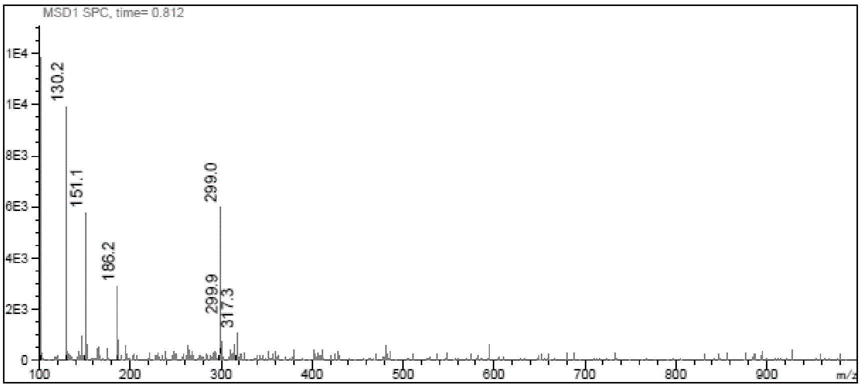
MSD1 SPC, time= 0.812
130.2
151.1
299.0
186.2
299.9
317.3
m/z

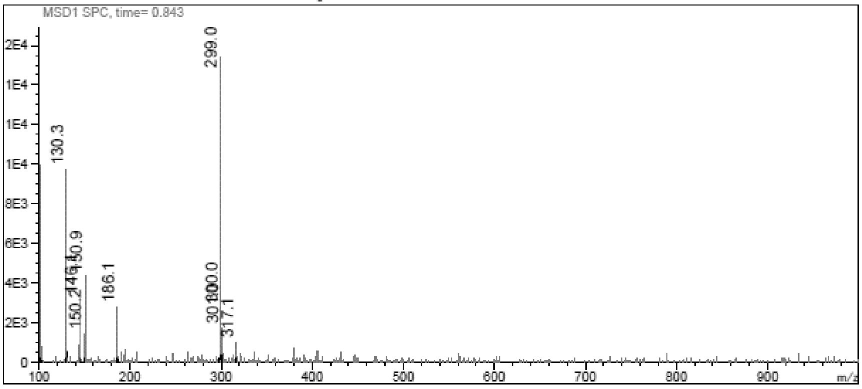
MSD1 SPC, time= 0.843
299.0
130.3
150.9
186.1
150.2
317.1
m/z

**$^{1}$H NMR** (4-Bromo-2-pyridyl)methanamine **(87)**

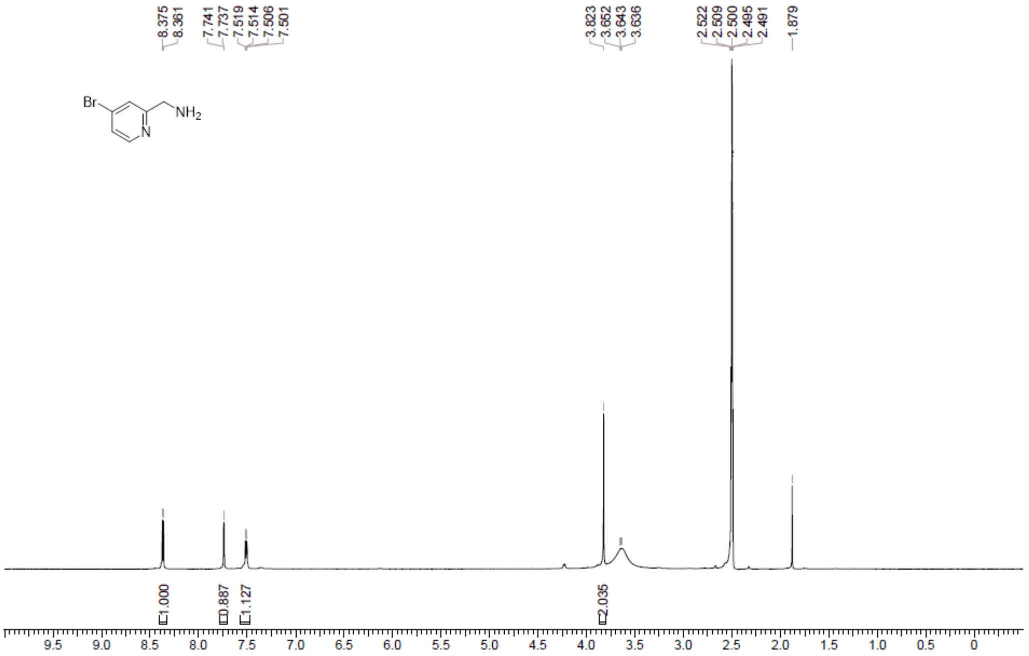

**LCMS** (4-Bromo-2-pyridyl)methanamine **(87)**

Compound ID : 1
Sample ID : EW26049-913-P1C1
Injection Vol : 5ul
Location : vial64
Acq Method : D:\method\0-60CD_R_220&254_POS.lcm
Org DataFile : D:\DATA\2022\2206\220614\EW26049-913-P1C1.lc
Injection Date : 2022-06-14 18:01:25
Instrument :LCMS-O 15-105

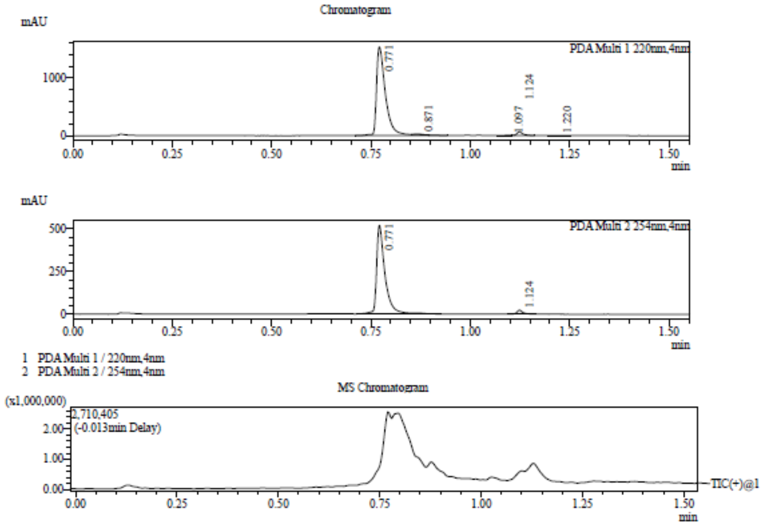

Integration Result

Peak Table

PDA Ch1 220nm

| Peak# | Ret. Time | Height | Height% | USP Width | Area | Area% |
|---|---|---|---|---|---|---|
| 1 | 0.771 | 1532626 | 94.715 | 0.040 | 2414874 | 96.401 |
| 2 | 0.871 | 9480 | 0.586 | 0.035 | 11159 | 0.445 |
| 3 | 1.097 | 5905 | 0.365 | 0.067 | 7198 | 0.287 |
| 4 | 1.124 | 66608 | 4.116 | 0.026 | 66713 | 2.663 |
| 5 | 1.220 | 3524 | 0.218 | 0.040 | 5076 | 0.203 |

Peak Table

PDA Ch2 254nm

| Peak# | Ret. Time | Height | Height% | USP Width | Area | Area% |
|---|---|---|---|---|---|---|
| 1 | 0.771 | 518067 | 95.740 | 0.037 | 764959 | 97.058 |
| 2 | 1.124 | 23054 | 4.260 | 0.026 | 23189 | 2.942 |

Mass Spectrum

RetTime: 0.770 Datafile: D:\DATA\2022\2206\220614\EW26049-913-P1C1.lcd

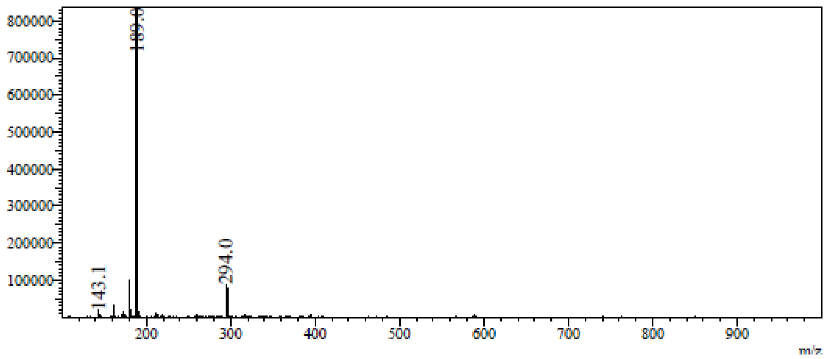

m/z

RetTime: 0.870 Datafile: D:\DATA\2022\2206\220614\EW26049-913-P1C1.lcd

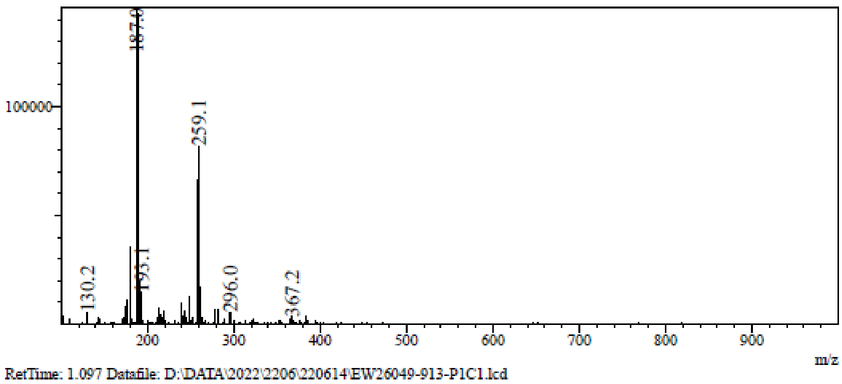

RetTime: 1.097 Datafile: D:\DATA\2022\2206\220614\EW26049-913-P1C1.lcd

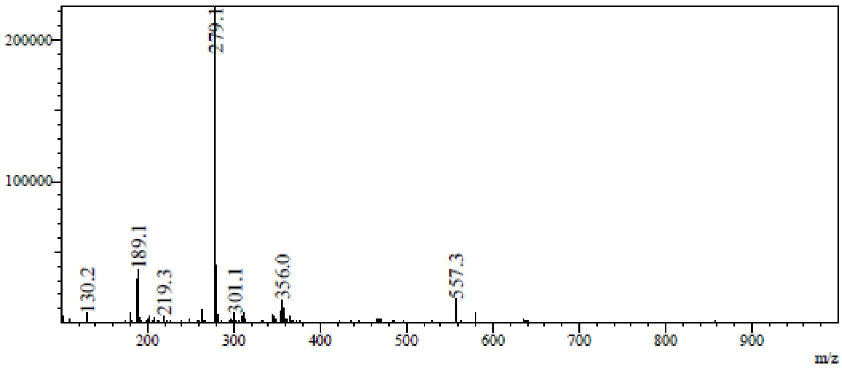

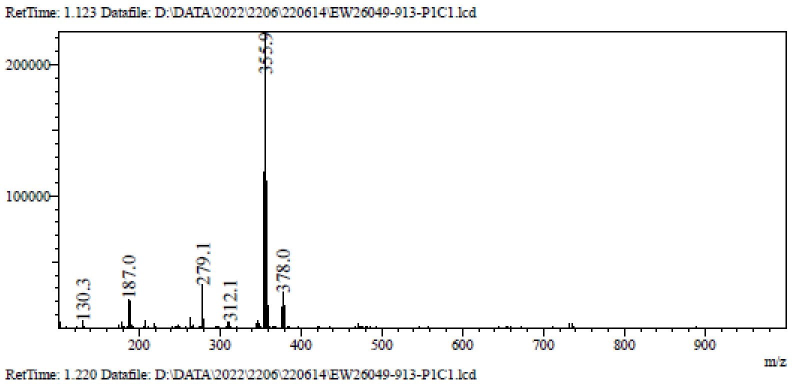

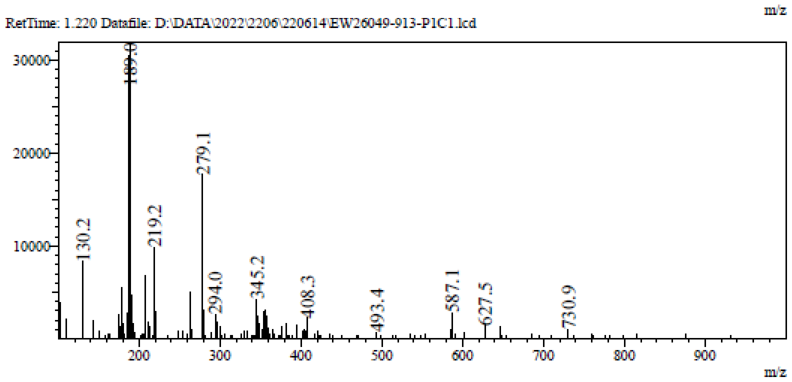

**[1]H NMR** N-[(4-Bromo-2-pyridyl)methyl]formamide **(88)**

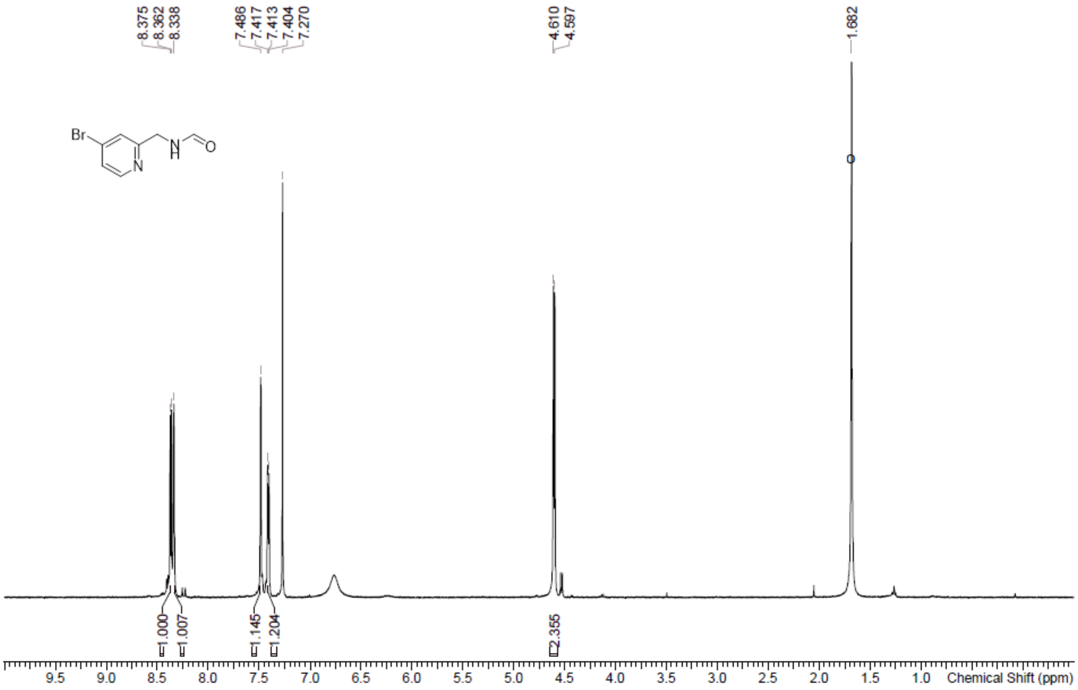

**[1]H NMR** 7-Bromoimidazo[1,5-a]pyridine **(89)**

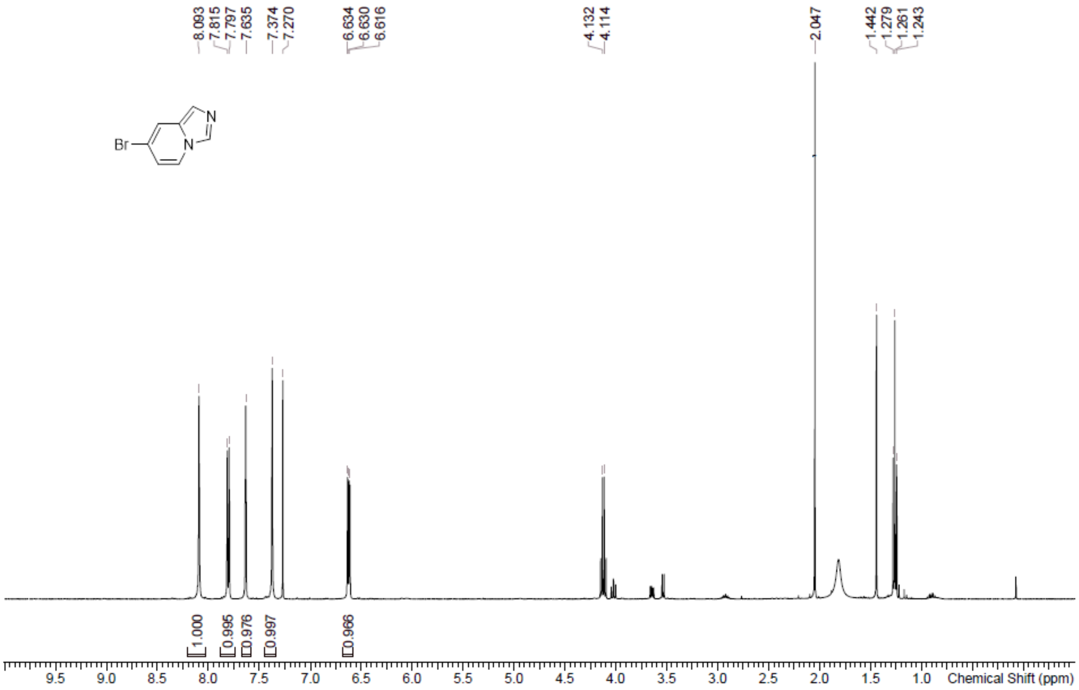

**[1]H NMR** 7-[(4-Methoxyphenyl)methylsulfanyl]imidazo[1,5-a]pyridine **(90)**

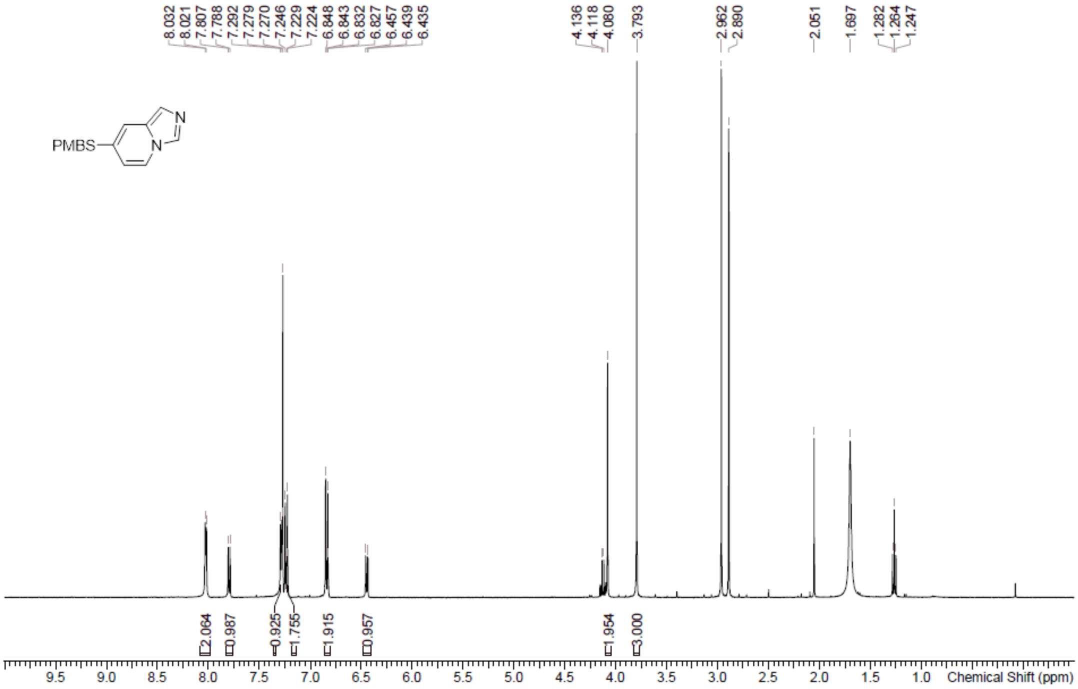

**LCMS** 7-[(4-Methoxyphenyl)methylsulfanyl]imidazo[1,5-a]pyridine **(90)**

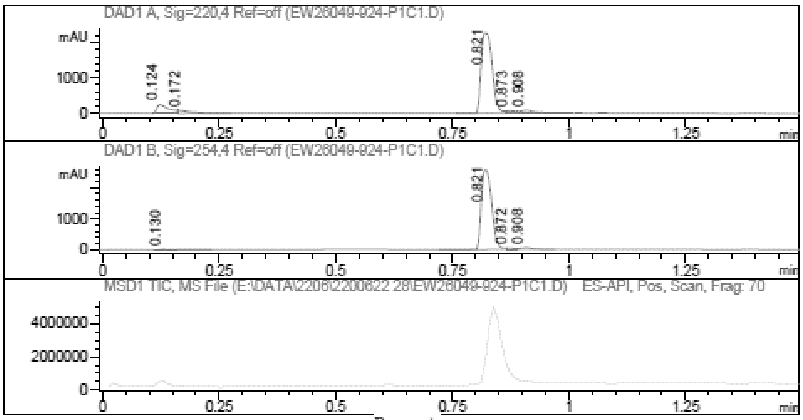

Report

Signal 1 : DAD1 A, Sig=220,4 Ref=off

| # | Meas. Ret. | Height | Width | Area | Area % |
|---|---|---|---|---|---|
| 1 | 0.124 | 253.178 | 0.025 | 447.109 | 9.825 |
| 2 | 0.172 | 69.786 | 0.032 | 161.664 | 3.552 |
| 3 | 0.821 | 2268.042 | 0.027 | 3732.992 | 82.027 |
| 4 | 0.873 | 57.333 | 0.019 | 75.903 | 1.668 |
| 5 | 0.908 | 78.055 | 0.025 | 133.251 | 2.928 |

Signal 2 : DAD1 B, Sig=254,4 Ref=off

| # | Meas. Ret. | Height | Width | Area | Area % |
|---|---|---|---|---|---|
| 1 | 0.130 | 16.372 | 0.045 | 56.535 | 1.344 |
| 2 | 0.821 | 2578.774 | 0.025 | 3990.766 | 94.887 |
| 3 | 0.872 | 44.519 | 0.018 | 53.897 | 1.281 |
| 4 | 0.908 | 55.745 | 0.027 | 104.626 | 2.488 |

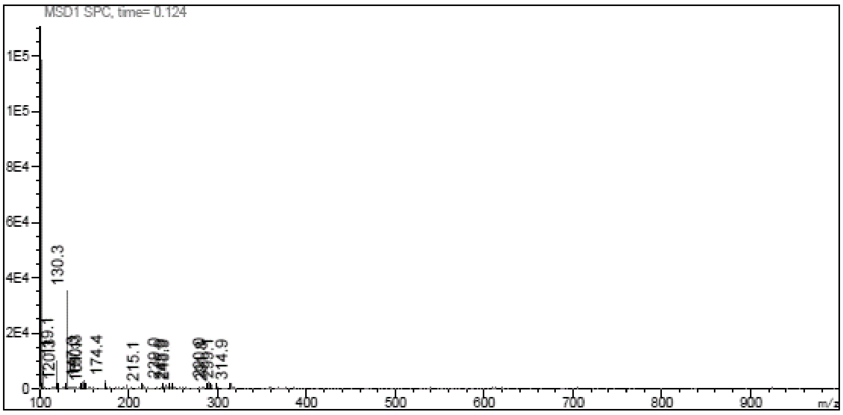

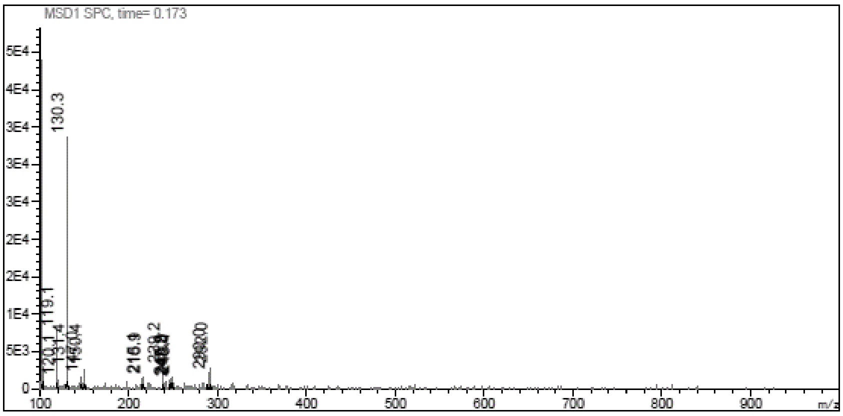

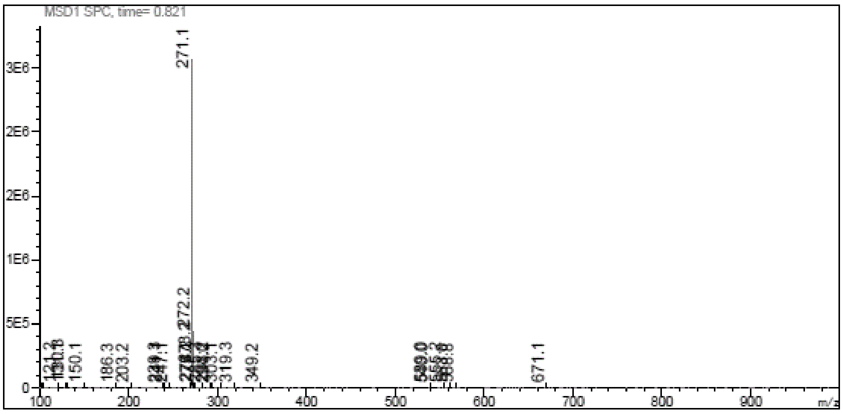

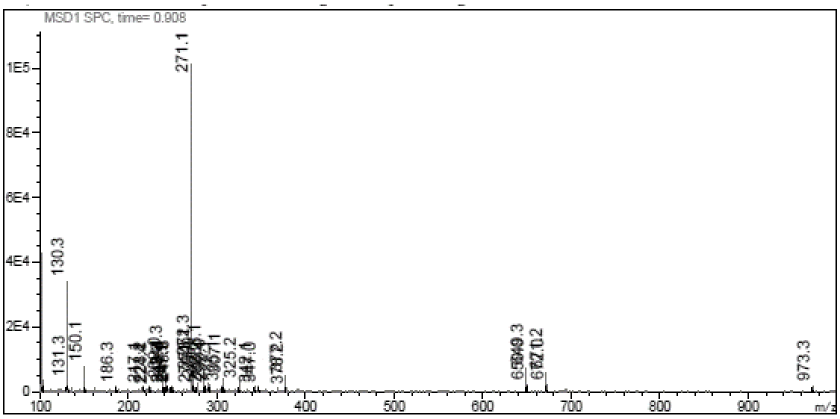

**[1]H NMR** Imidazo[1,5-a]pyridine-7-thiol **(91)**

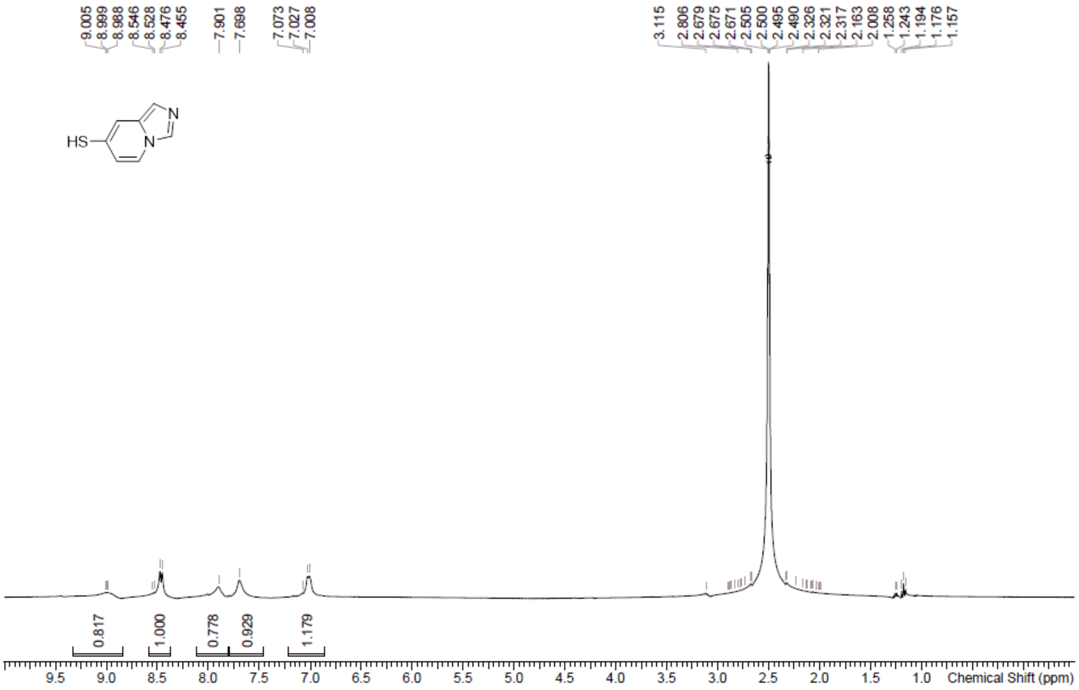

**$^{1}$H NMR** 9-[(2*R*,3*R*,4*R*,5*S*)-5-(1,3-Benzothiazol-2-ylsulfanylmethyl)-3,4-bis[[tert-butyl(dimethyl) silyl]oxy]tetrahydrofuran-2-yl]-*N*-methyl-purin-6-amine **(93)**

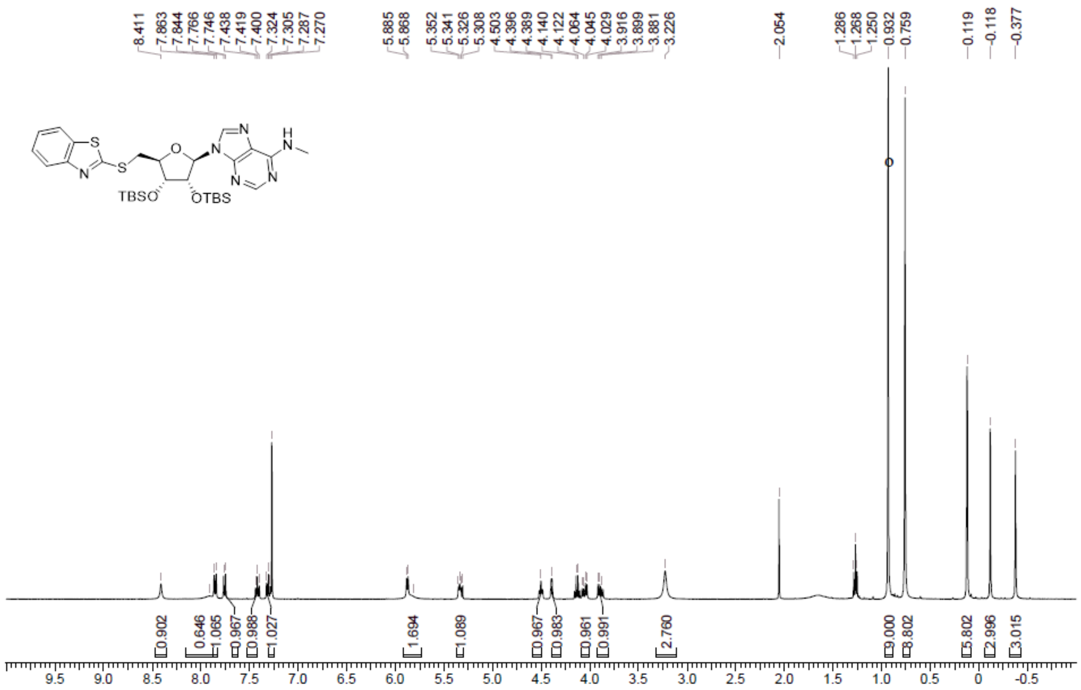

**LCMS** 9-[(2*R*,3*R*,4*R*,5*S*)-5-(1,3-Benzothiazol-2-ylsulfanylmethyl)-3,4-bis[[tert-butyl(dimethyl) silyl]oxy]tetrahydrofuran-2-yl]-*N*-methyl-purin-6-amine **(93)**

Compound ID : 1
Sample ID : EW26049-722-P1S1
Injection Vol : 5ul
Location : vial84
Acq Method : D:\method\50-100CD_R_220&254_POS.lcm
Org DataFile : D:\DATA\2022\2201\220121\EW26049-722-P1S1.lc
Injection Date : 2022-01-21 19:30:41
Instrument :LCMS-W

Chromatogram

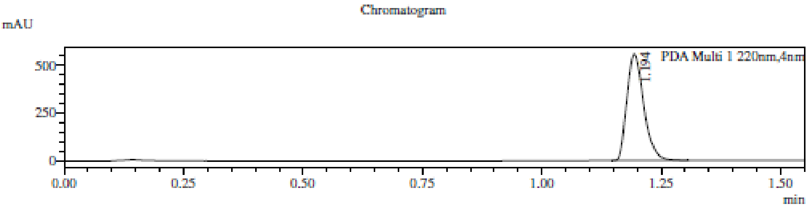

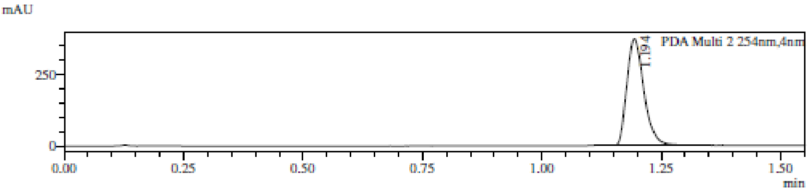

1 PDA Multi 1 / 220nm,4nm
2 PDA Multi 2 / 254nm,4nm

MS Chromatogram

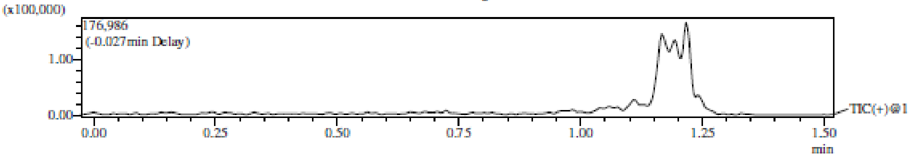

Integration Result

Peak Table

PDA Ch1 220nm

| Peak# | Ret. Time | Height | Height% | USP Width | Area | Area% |
|---|---|---|---|---|---|---|
| 1 | 1.194 | 555080 | 100.000 | 0.065 | 1369531 | 100.000 |

Peak Table

PDA Ch2 254nm

| Peak# | Ret. Time | Height | Height% | USP Width | Area | Area% |
|---|---|---|---|---|---|---|
| 1 | 1.194 | 372151 | 100.000 | 0.065 | 919959 | 100.000 |

Mass Spectrum
RetTime: 1.193 Datafile: D:\DATA\2022\2201\220121\EW26049-722-P1S1.lcd

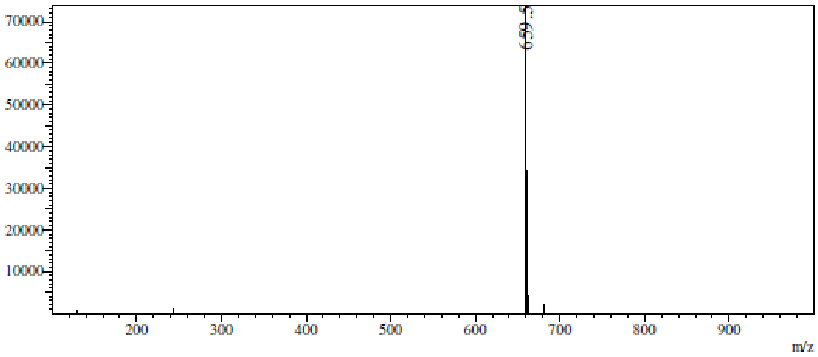

**[1]H NMR** 9-[(2*R*,3*R*,4*R*,5*S*)-5-(1,3-Benzothiazol-2-ylsulfanylmethyl)-3,4-bis[[tert-butyl(dimethyl)

silyl]oxy]tetrahydrofuran-2-yl]-*N*,*N*-dimethyl-purin-6-amine **(94)**

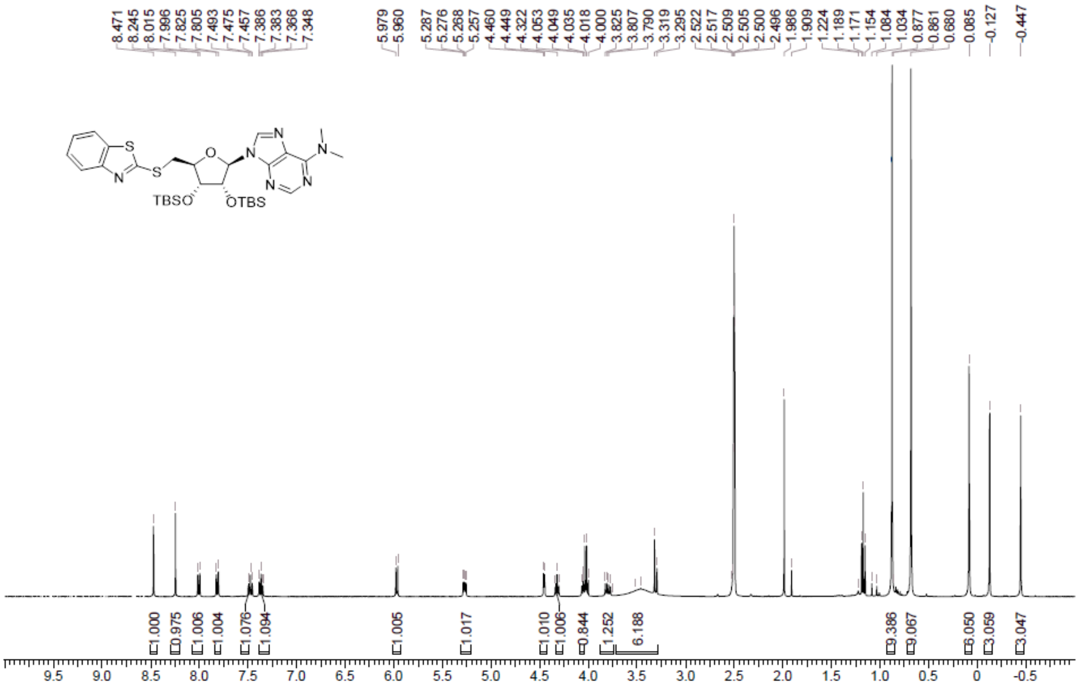

**LCMS** 9-[(2*R*,3*R*,4*R*,5*S*)-5-(1,3-Benzothiazol-2-ylsulfanylmethyl)-3,4-bis[[tert-butyl(dimethyl) silyl]oxy]tetrahydrofuran-2-yl]-*N*,*N*-dimethyl-purin-6-amine **(94)**

Compound ID : 1
Sample ID : EW26049-754-P1C1
Injection Vol : 5ul
Location : vial61
Acq Method : D:\method\50-100AB_R_220&254.lcm
Org DataFile : D:\DATA\2022\2202\220222\EW26049-754-P1C1.lc
Injection Date : 2022-02-22 19:29:53
Instrument : LCMS-AA 15-105

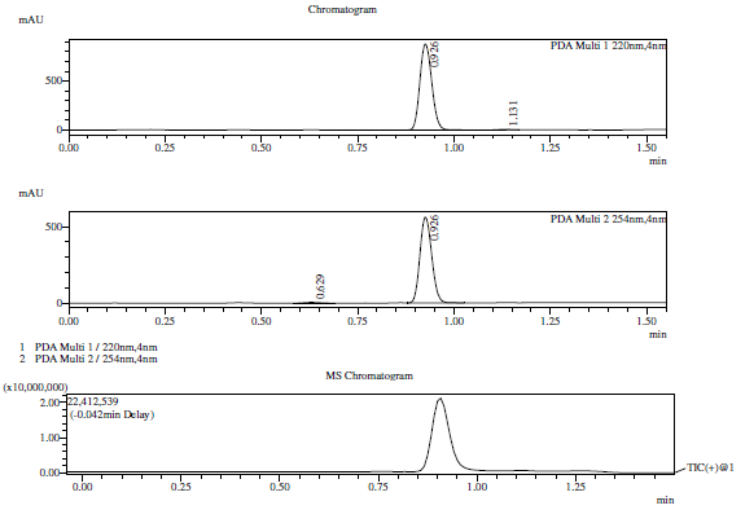

Integration Result

Peak Table

PDA Ch1 220nm

| Peak# | Ret. Time | Height | Height% | USP Width | Area | Area% |
|---|---|---|---|---|---|---|
| 1 | 0.926 | 879108 | 99.321 | 0.058 | 1890220 | 99.348 |
| 2 | 1.131 | 6012 | 0.679 | 0.057 | 12402 | 0.652 |

Peak Table

PDA Ch2 254nm

| Peak# | Ret. Time | Height | Height% | USP Width | Area | Area% |
|---|---|---|---|---|---|---|
| 1 | 0.629 | 5654 | 0.998 | 0.085 | 17247 | 1.421 |
| 2 | 0.926 | 560908 | 99.002 | 0.057 | 1196730 | 98.579 |

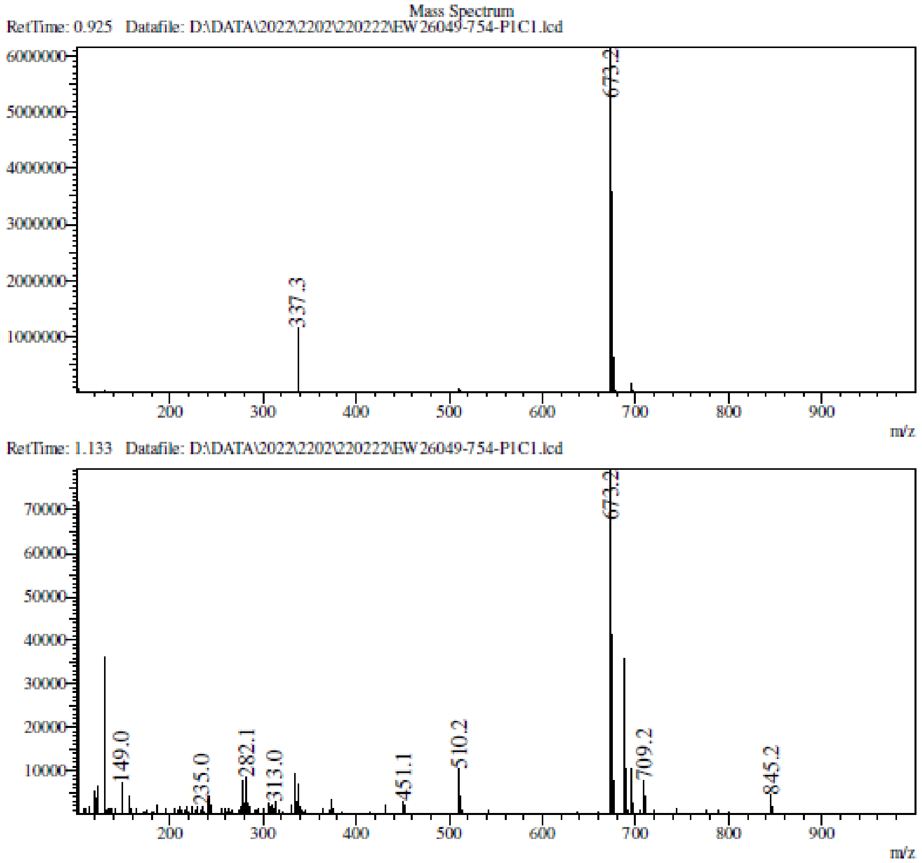

**[1]H NMR** (2*S*,3*S*,4*R*,5*R*)-2-(1,3-Benzothiazol-2-ylsulfanylmethyl)-5-[6-(methylamino)purin-9-yl]tetrahydrofuran-3,4-diol **(18)**

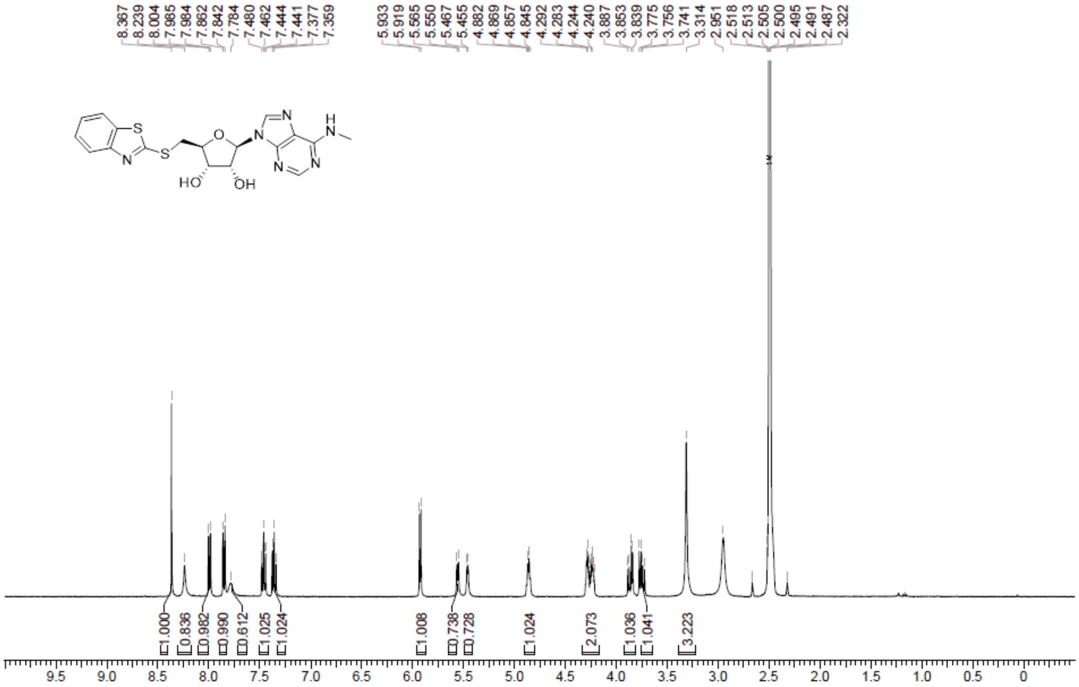

**HPLC** (2*S*,3*S*,4*R*,5*R*)-2-(1,3-Benzothiazol-2-ylsulfanylmethyl)-5-[6-(methylamino)purin-9-yl]tetrahydrofuran-3,4-diol **(18)**

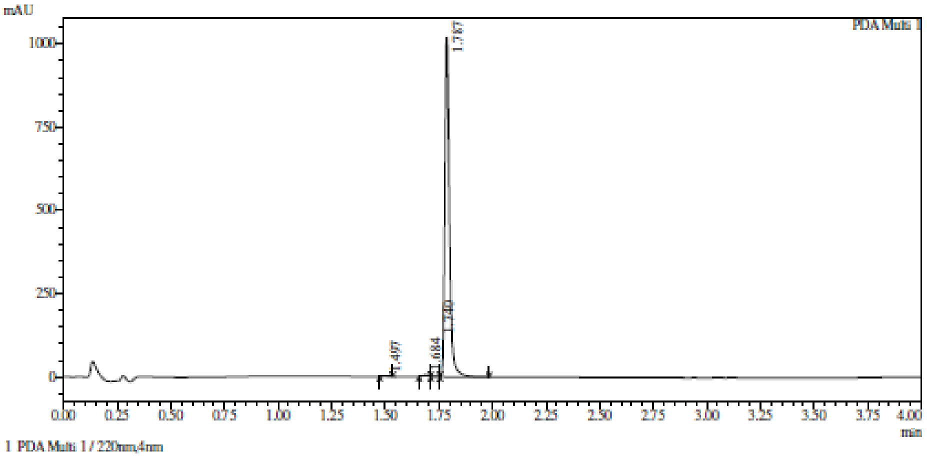

Integration result

PeakTable

PDA Ch1 220nm

| Peak# | Ret. Time | USP Width | Resolution | Height | Area | Area % |
|---|---|---|---|---|---|---|
| 1 | 1.497 | 0.043 | 0.000 | 926 | 1468 | 0.095 |
| 2 | 1.684 | 0.039 | 4.558 | 5830 | 8404 | 0.541 |
| 3 | 1.740 | 0.104 | 0.779 | 2140 | 4374 | 0.282 |
| 4 | 1.787 | 0.039 | 0.651 | 1020359 | 1538690 | 99.083 |
| Total | | | | 1029255 | 1552936 | 100.000 |

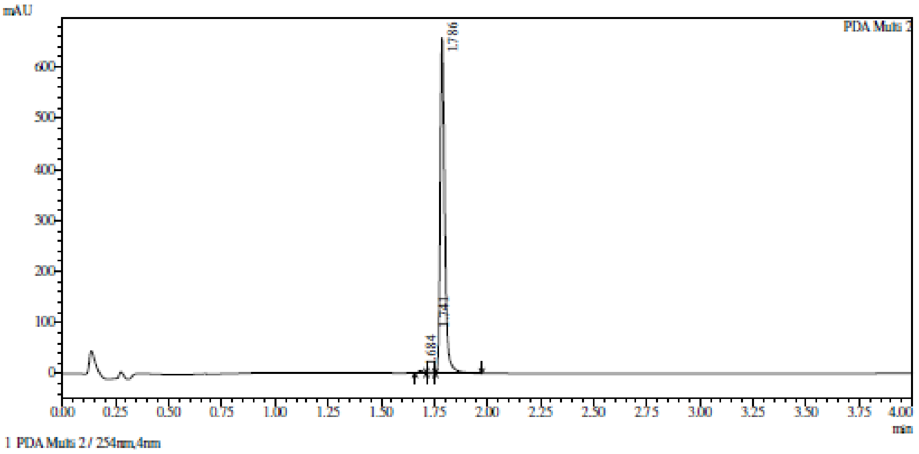

Integration result

PeakTable

PDA Ch2 254nm

| Peak# | Ret. Time | USP Width | Resolution | Height | Area | Area % |
|---|---|---|---|---|---|---|
| 1 | 1.684 | 0.039 | 0.000 | 4448 | 6454 | 0.647 |
| 2 | 1.741 | 0.109 | 0.764 | 1359 | 2765 | 0.277 |
| 3 | 1.786 | 0.039 | 0.617 | 656410 | 988544 | 99.076 |
| Total | | | | 662217 | 997762 | 100.000 |

**HRMS** (2*S*,3*S*,4*R*,5*R*)-2-(1,3-Benzothiazol-2-ylsulfanylmethyl)-5-[6-(methylamino)purin-9-yl]tetrahydrofuran-3,4-diol **(18)**

**Compound Table**

| Label | Tgt Score | Mass Error (ppm) | Tgt Formula | Obs. RT | Ref. Mass | Obs. Mass |
|---|---|---|---|---|---|---|
| Cpd 1: C18 H18 N6 O3 S2; 1.710 | 98.27 | 2.08 | C18 H18 N6 O3 S2 | 1.71 | 430.0882 | 430.0891 |

| Obs. m/z | Obs. RT | Obs. Mass | Tgt Formula | Tgt Mass | Tgt Mass Error (ppm) | Find Cpds Algorithm |
|---|---|---|---|---|---|---|
| 216.0514 | 1.71 | 430.0891 | C18 H18 N6 O3 S2 | 430.0882 | 2.08 | Find by Formula |

**Compound Chromatograms**

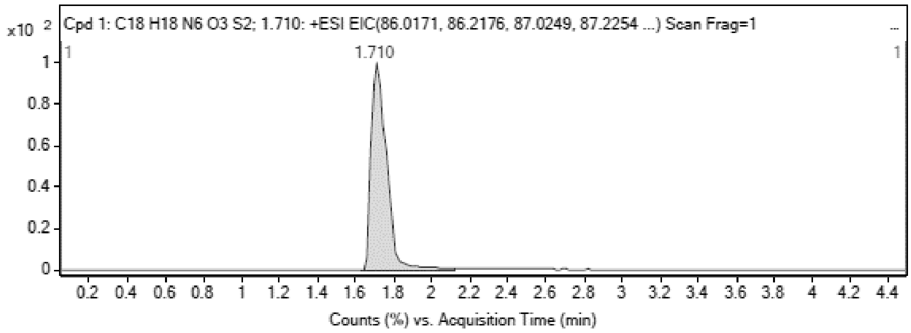

MS Zoomed Spectrum

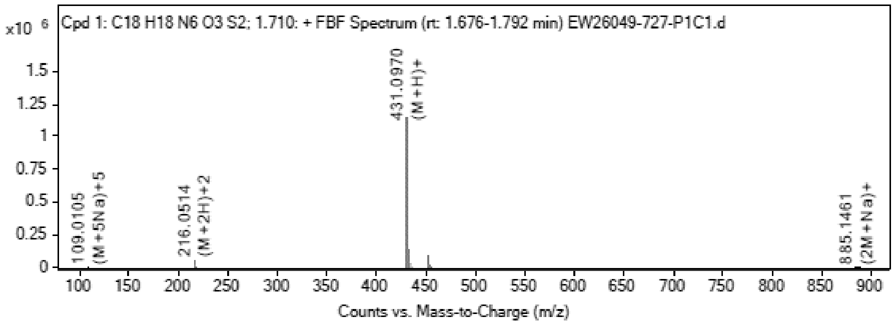

**MS Spectrum Peak List**

| Obs. m/z | Charge | Abund | Ion/Isotope |
|---|---|---|---|
| 216.0514 | 2 | 57524.63 | (M+2H)+2 |
| 216.5526 | 2 | 13759.6 | (M+2H)+2 |
| 217.0505 | 2 | 7395.06 | (M+2H)+2 |
| 431.097 | 1 | 1132331.75 | (M+H)+ |
| 432.0982 | 1 | 283200.53 | (M+H)+ |
| 433.0938 | 1 | 136242.98 | (M+H)+ |
| 453.0772 | 1 | 94407.89 | (M+Na)+ |
| 454.0789 | 1 | 21746.32 | (M+Na)+ |
| 455.0751 | 1 | 11364.44 | (M+Na)+ |
| 885.1461 | 1 | 3790.71 | (2M+Na)+ |

MS Zoomed Spectrum

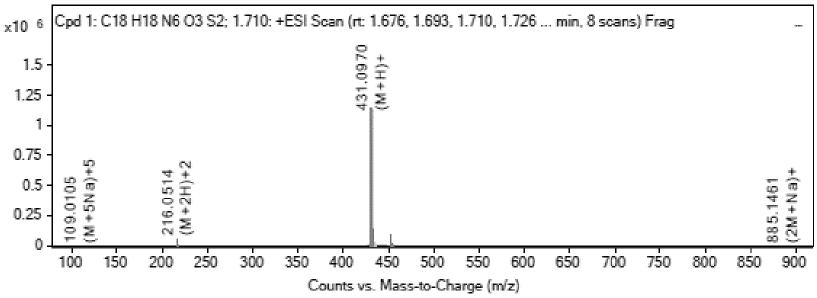

**MS Spectrum Peak List**

| Obs. m/z | Charge | Abund | Ion/Isotope | Tgt Mass Error (ppm) |
|---|---|---|---|---|
| 216.0514 | 2 | 57524.63 | (M+2H)+2 | -0.26 |
| 216.5526 | 2 | 13759.6 | (M+2H)+2 | 0.28 |
| 217.0505 | 2 | 7395.06 | (M+2H)+2 | -0.19 |
| 431.097 | 1 | 1132331.75 | (M+H)+ | -3.48 |
| 432.0982 | 1 | 283200.53 | (M+H)+ | -0.41 |
| 433.0938 | 1 | 136242.98 | (M+H)+ | -0.35 |
| 453.0772 | 1 | 94407.89 | (M+Na)+ | 0.51 |
| 454.0789 | 1 | 21746.32 | (M+Na)+ | 2.23 |
| 455.0751 | 1 | 11364.44 | (M+Na)+ | 1.23 |
| 885.1461 | 1 | 3790.71 | (2M+Na)+ | 21.47 |

--- End Of Report ---

**$^1$H NMR** (2*R*,3*R*,4*S*,5*S*)-2-(6-Aminopurin-9-yl)-5-(1,3-benzothiazol-2-ylsulfanylmethyl) tetrahydrofuran-3,4-diol **(19)**

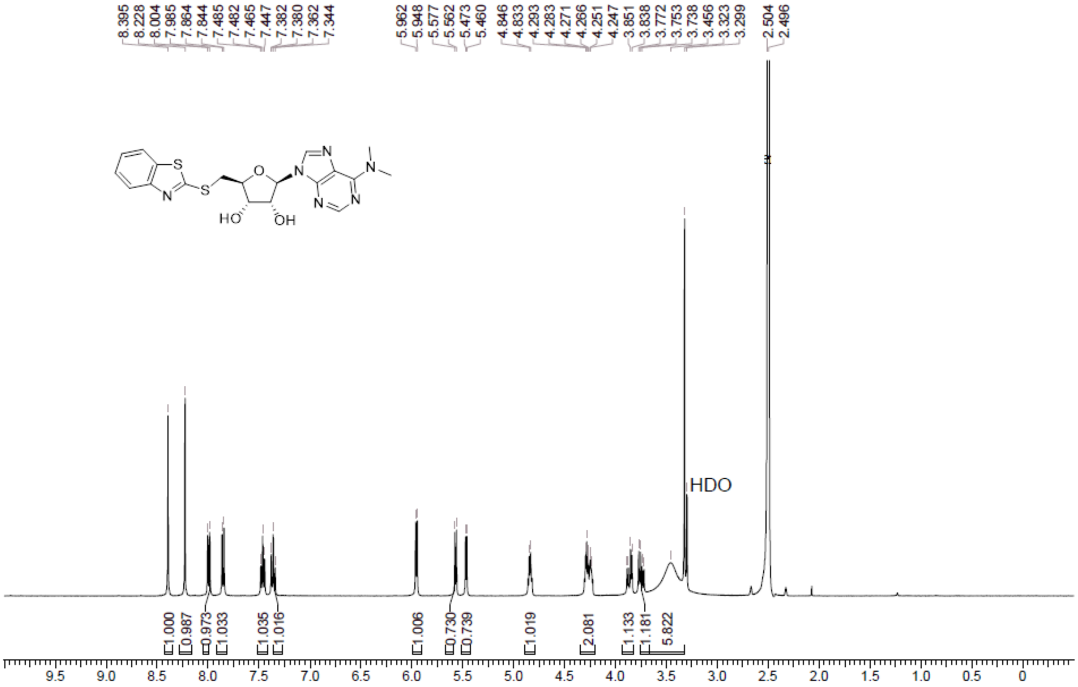

**HPLC** (2*R*,3*R*,4*S*,5*S*)-2-(6-Aminopurin-9-yl)-5-(1,3-benzothiazol-2-ylsulfanylmethyl) tetrahydrofuran-3,4-diol **(19)**

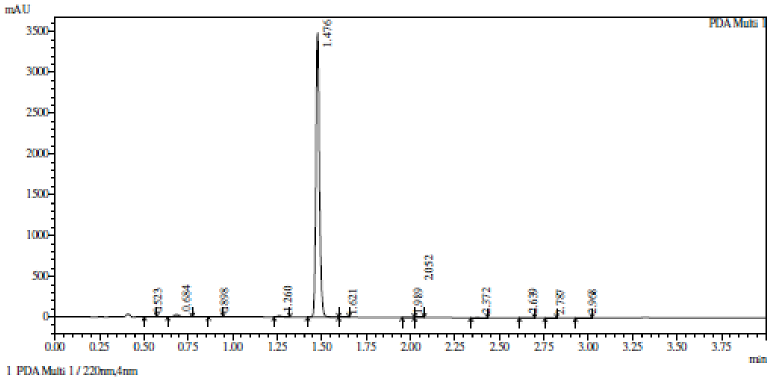

Integration result

PeakTable

PDA Ch1 220nm

| Peak# | Ret. Time | USP Width | Resolution | Height | Area | Area % |
|---|---|---|---|---|---|---|
| 1 | 0.523 | 0.045 | 0.000 | 2666 | 4485 | 0.090 |
| 2 | 0.684 | 0.056 | 3.175 | 26727 | 56430 | 1.133 |
| 3 | 0.898 | 0.067 | 3.500 | 1261 | 3054 | 0.061 |
| 4 | 1.260 | 0.042 | 6.707 | 14542 | 22571 | 0.453 |
| 5 | 1.476 | 0.035 | 5.636 | 3490065 | 4874839 | 97.901 |
| 6 | 1.621 | 0.043 | 3.685 | 1262 | 1941 | 0.039 |
| 7 | 1.989 | 0.048 | 8.087 | 1557 | 2687 | 0.054 |
| 8 | 2.052 | 0.046 | 1.353 | 880 | 1408 | 0.028 |
| 9 | 2.372 | 0.080 | 5.084 | 1178 | 3370 | 0.068 |
| 10 | 2.639 | 0.069 | 3.604 | 653 | 1759 | 0.035 |
| 11 | 2.787 | 0.051 | 2.479 | 942 | 1788 | 0.036 |
| 12 | 2.968 | 0.063 | 3.170 | 2024 | 5009 | 0.101 |
| Total | | | | 3543758 | 4979342 | 100.000 |

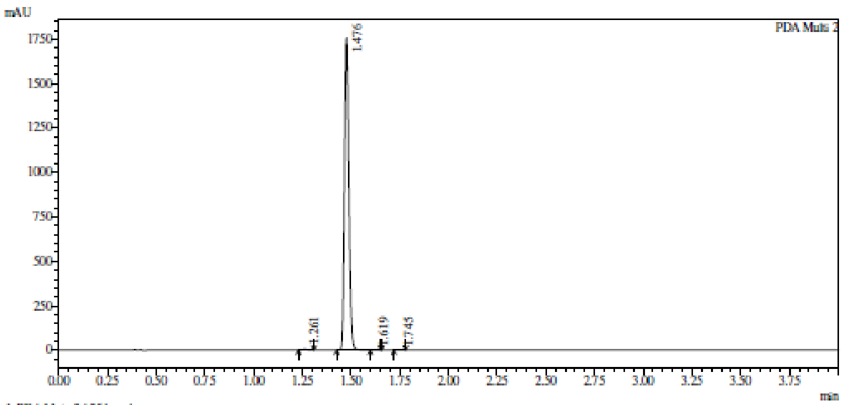

1 PDA Multi 2 / 254nm,4nm

Integration result

PeakTable

PDA Ch2 254nm

| Peak# | Ret. Time | USP Width | Resolution | Height | Area | Area % |
|---|---|---|---|---|---|---|
| 1 | 1.261 | 0.042 | 0.000 | 6344 | 9836 | 0.358 |
| 2 | 1.476 | 0.042 | 5.087 | 1758970 | 2734198 | 99.560 |
| 3 | 1.619 | 0.042 | 3.388 | 746 | 1125 | 0.041 |
| 4 | 1.745 | 0.045 | 2.918 | 674 | 1111 | 0.040 |
| Total | | | | 1766734 | 2746270 | 100.000 |

**HRMS** (2*R*,3*R*,4*S*,5*S*)-2-(6-Aminopurin-9-yl)-5-(1,3-benzothiazol-2-ylsulfanylmethyl) tetrahydrofuran-3,4-diol **(19)**

Compound Table

| Label | Tgt Score | Mass Error (ppm) | Tgt Formula | Obs. RT | Ref. Mass | Obs. Mass |
|---|---|---|---|---|---|---|
| Cpd 1: C19 H20 N6 O3 S2; 1.810 | 96.37 | 6.14 | C19 H20 N6 O3 S2 | 1.81 | 444.1038 | 444.1066 |

| Obs. *m/z* | Obs. RT | Obs. Mass | Tgt Formula | Tgt Mass | Tgt Mass Error (ppm) | Find Cpds Algorithm |
|---|---|---|---|---|---|---|
| 223.0598 | 1.81 | 444.1066 | C19 H20 N6 O3 S2 | 444.1038 | 6.14 | Find by Formula |

Compound Chromatograms

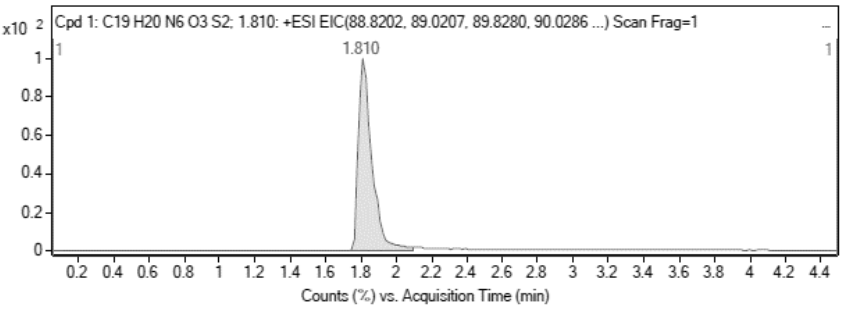

MS Zoomed Spectrum

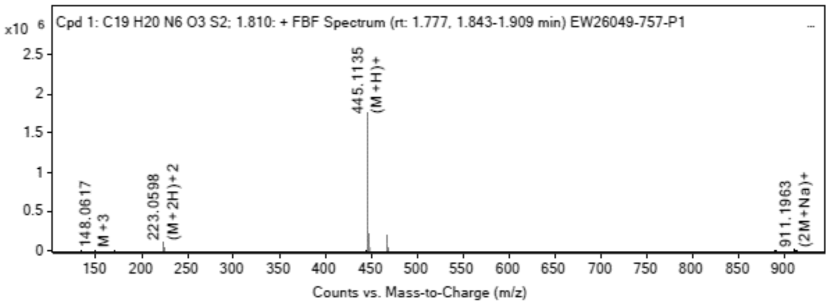

MS Spectrum Peak List

| Obs. *m/z* | Charge | Abund | Ion/Isotope |
|---|---|---|---|
| 223.0598 | 2 | 97239.34 | (M+2H)+2 |
| 223.5612 | 2 | 24231.24 | (M+2H)+2 |

| | | | |
|---|---|---|---|
| 224.0591 | 2 | 12233.51 | (M+2H)+2 |
| 445.1135 | 1 | 1760436.13 | (M+H)+ |
| 446.1153 | 1 | 444699.91 | (M+H)+ |
| 447.1109 | 1 | 214511.19 | (M+H)+ |
| 467.0945 | 1 | 200039.48 | (M+Na)+ |
| 468.0965 | 1 | 47714.09 | (M+Na)+ |
| 469.0922 | 1 | 23735.76 | (M+Na)+ |
| 911.1963 | 1 | 11867.88 | (2M+Na)+ |

MS Zoomed Spectrum

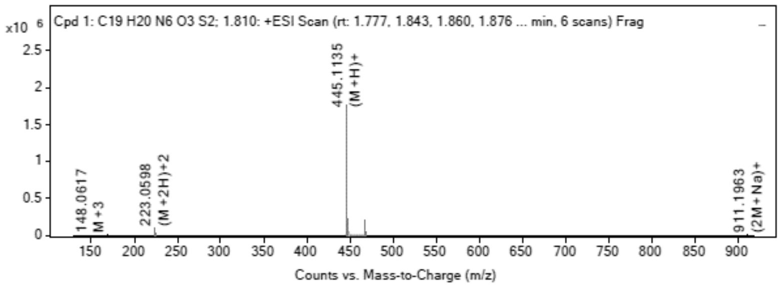

MS Spectrum Peak List

| Obs. m/z | Charge | Abund | Ion/Isotope | Tgt Mass Error (ppm) |
|---|---|---|---|---|
| 223.0598 | 2 | 97239.34 | (M+2H)+2 | -2.8 |
| 223.5612 | 2 | 24231.24 | (M+2H)+2 | -3.37 |
| 224.0591 | 2 | 12233.51 | (M+2H)+2 | -3.03 |
| 445.1135 | 1 | 1760436.13 | (M+H)+ | -5.43 |
| 446.1153 | 1 | 444699.91 | (M+H)+ | -3.55 |
| 447.1109 | 1 | 214511.19 | (M+H)+ | -3.17 |
| 467.0945 | 1 | 200039.48 | (M+Na)+ | -3.19 |
| 468.0965 | 1 | 47714.09 | (M+Na)+ | -1.8 |
| 469.0922 | 1 | 23735.76 | (M+Na)+ | -1.55 |
| 911.1963 | 1 | 11867.88 | (2M+Na)+ | 0.64 |

--- End Of Report ---

**$^1$H NMR** [(2*R*,3*R*,4*R*,5*R)*-3,4-Bis[[tert-butyl(dimethyl)silyl]oxy]-5-(6- chloropurin-9-yl) tetrahydrofuran-2-yl]methoxy-tert-butyl-dimethyl-silane **(96)**

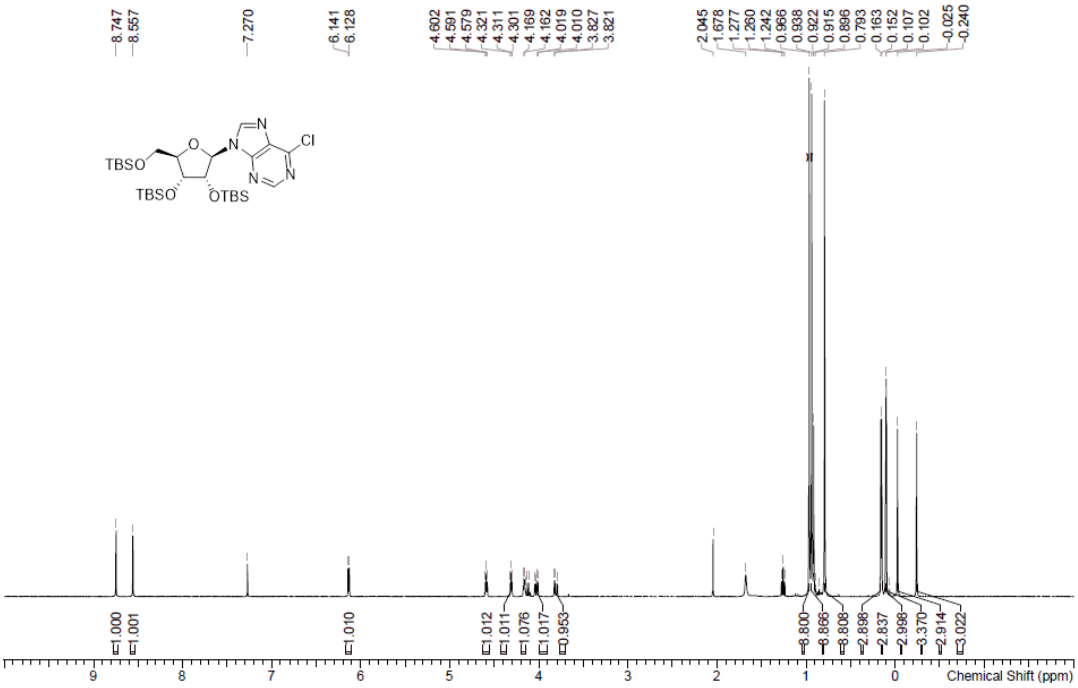

**[1]H NMR** [(2*R*,3*R*,4*R*,5*R*)-3,4-Bis[[tert-butyl(dimethyl)silyl]oxy]-5- (6-chloropurin-9-yl)tetrahydrofuran-2-yl]methanol **(46)**

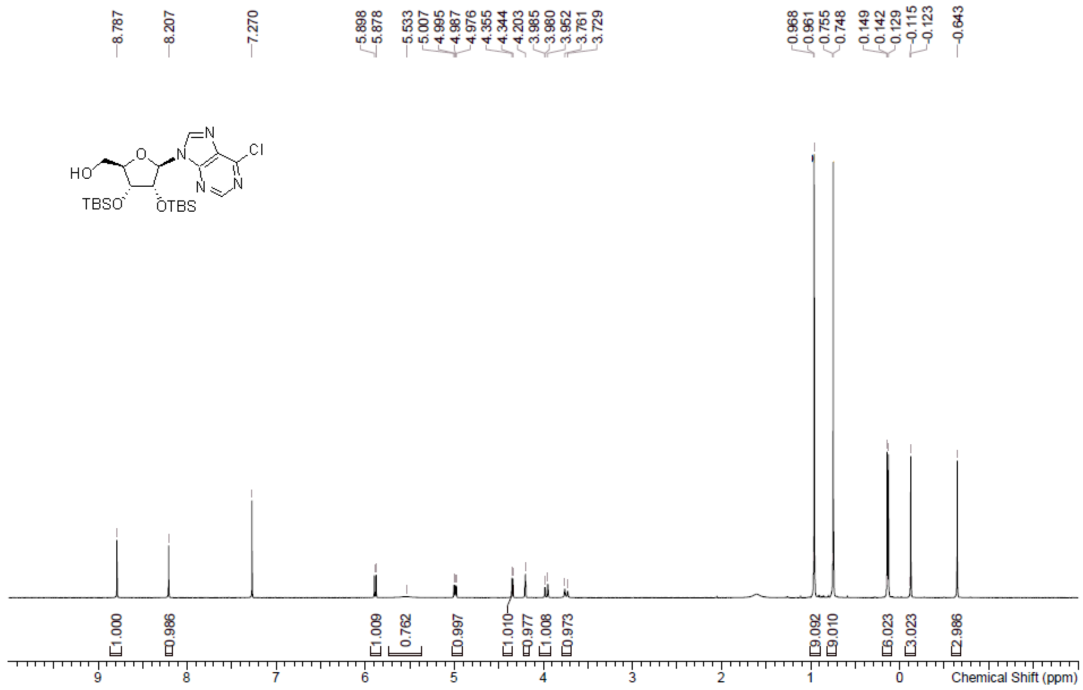

**LCMS** [(2*R*,3*R*,4*R*,5*R*)-3,4-Bis[[tert-butyl(dimethyl)silyl]oxy]-5- (6-chloropurin-9-yl)tetrahydrofuran-2-yl]methanol **(46)**

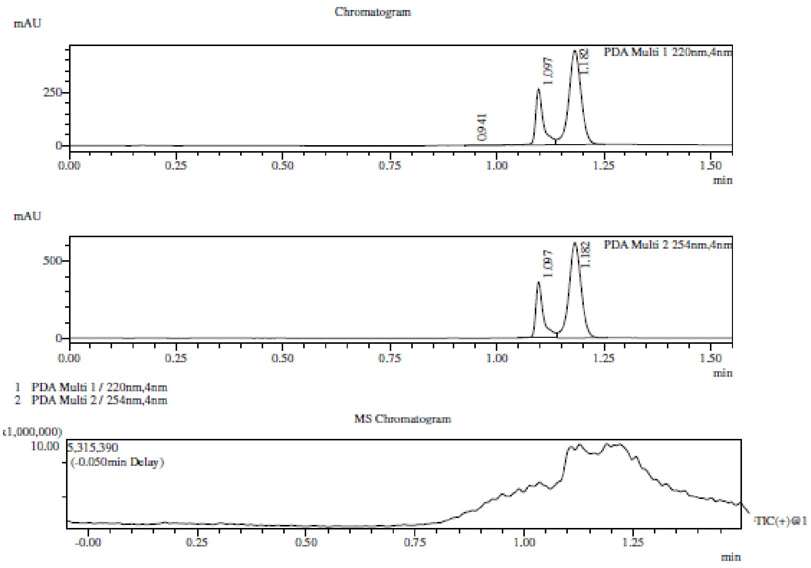

Integration Result

Peak Table

PDA Ch1 220nm

| Peak# | Ret. Time | Height | Height% | USP Width | Area | Area% |
|---|---|---|---|---|---|---|
| 1 | 0.941 | 2690 | 0.382 | 0.084 | 14131 | 1.160 |
| 2 | 1.097 | 261386 | 37.121 | 0.031 | 330533 | 27.132 |
| 3 | 1.182 | 440076 | 62.497 | 0.052 | 873570 | 71.708 |

Peak Table

PDA Ch2 254nm

| Peak# | Ret. Time | Height | Height% | USP Width | Area | Area% |
|---|---|---|---|---|---|---|
| 1 | 1.097 | 361984 | 37.089 | 0.031 | 452505 | 27.211 |
| 2 | 1.182 | 614003 | 62.911 | 0.052 | 1210427 | 72.789 |

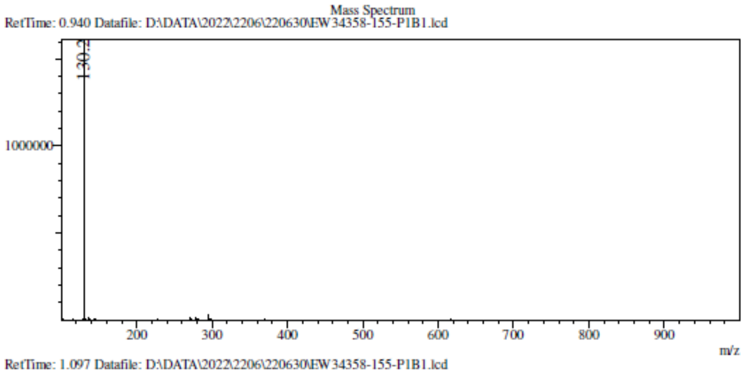
Mass Spectrum
RetTime: 0.940 Datafile: D:\DATA\2022\2206\220630\EW34358-155-P1B1.lcd
130.2
1000000
200
300
400
500
600
700
800
900
m/z

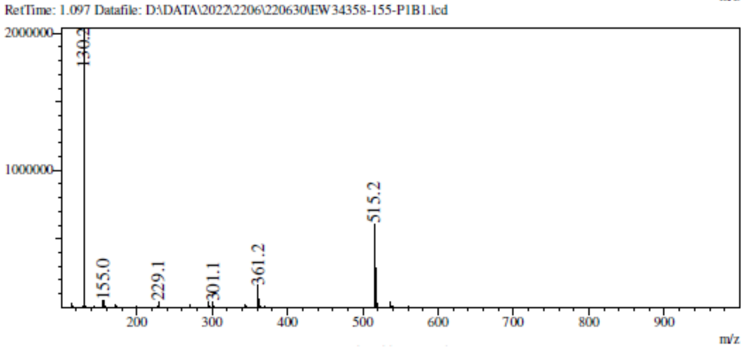
RetTime: 1.097 Datafile: D:\DATA\2022\2206\220630\EW34358-155-P1B1.lcd
2000000
1000000
130.2
155.0
229.1
301.1
361.2
515.2
200
300
400
500
600
700
800
900
m/z

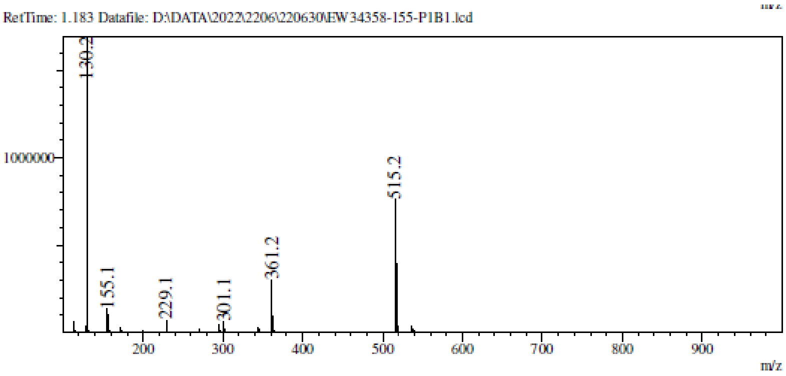
RetTime: 1.183 Datafile: D:\DATA\2022\2206\220630\EW34358-155-P1B1.lcd
1000000
130.2
155.1
229.1
301.1
361.2
515.2
200
300
400
500
600
700
800
900
m/z

**¹H NMR** [(2R,3R,4R,5R)-3,4-Bis[[tert-butyl(dimethyl)silyl]oxy]-5- (6-chloropurin-9-yl)tetrahydrofuran-2-yl]methyl methanesulfonate **(97)**

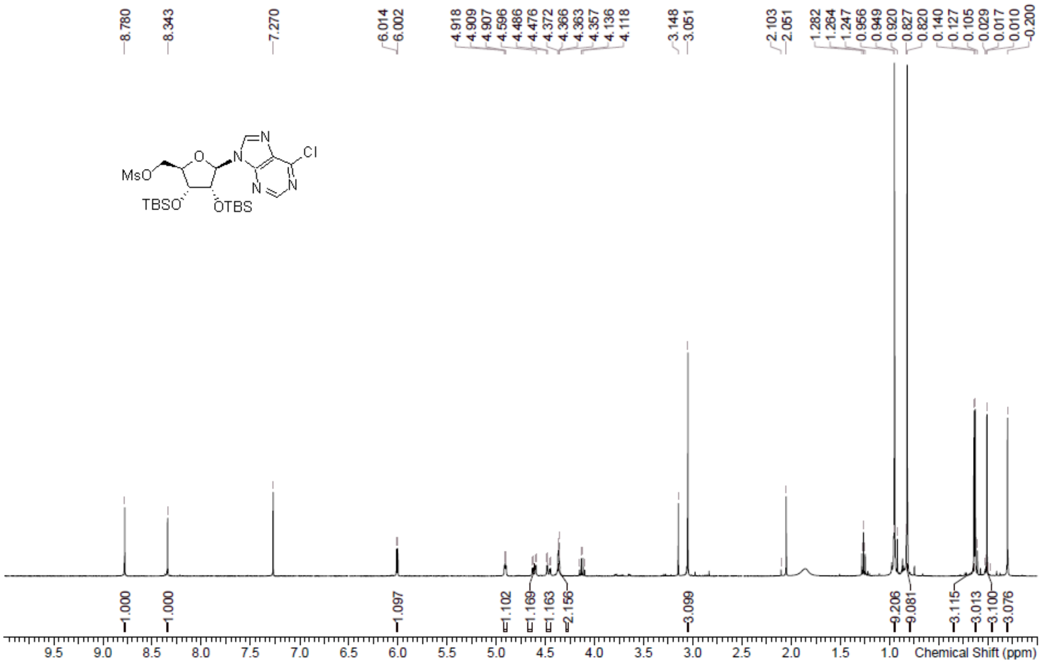

**LCMS** [(2R,3R,4R,5R)-3,4-Bis[[tert-butyl(dimethyl)silyl]oxy]-5- (6-chloropurin-9-yl)tetrahydrofuran-2-yl]methyl methanesulfonate **(97)**

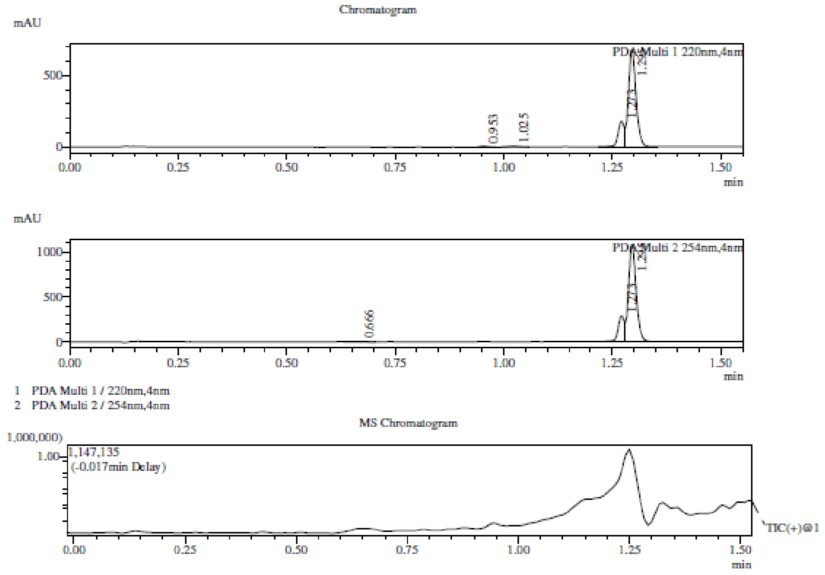

Integration Result

Peak Table

PDA Ch1 220nm

| Peak# | Ret. Time | Height | Height% | USP Width | Area | Area% |
|---|---|---|---|---|---|---|
| 1 | 0.953 | 5887 | 0.681 | 0.035 | 7556 | 0.763 |
| 2 | 1.025 | 4975 | 0.576 | 0.043 | 7642 | 0.771 |
| 3 | 1.273 | 176648 | 20.435 | 0.053 | 183282 | 18.499 |
| 4 | 1.295 | 676946 | 78.309 | 0.032 | 792270 | 79.967 |

Peak Table

PDA Ch2 254nm

| Peak# | Ret. Time | Height | Height% | USP Width | Area | Area% |
|---|---|---|---|---|---|---|
| 1 | 0.666 | 3744 | 0.275 | 0.071 | 8982 | 0.570 |
| 2 | 1.273 | 287287 | 21.079 | 0.039 | 294613 | 18.706 |
| 3 | 1.295 | 1071906 | 78.647 | 0.032 | 1271330 | 80.723 |

Mass Spectrum
RetTime: 0.950 Datafile: D:\DATA\2022\2207\220701\EW34358-160-P1B2.lcd

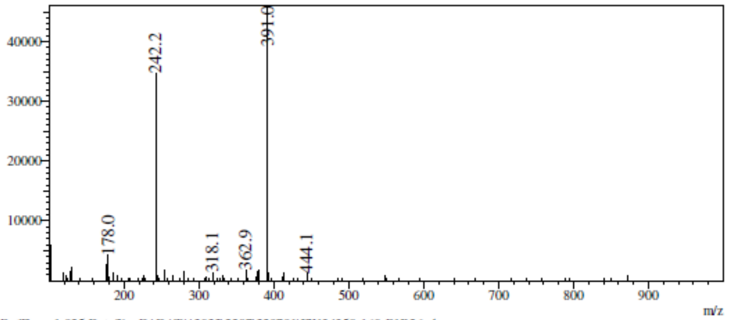

RetTime: 1.025 Datafile: D:\DATA\2022\2207\220701\EW34358-160-P1B2.lcd

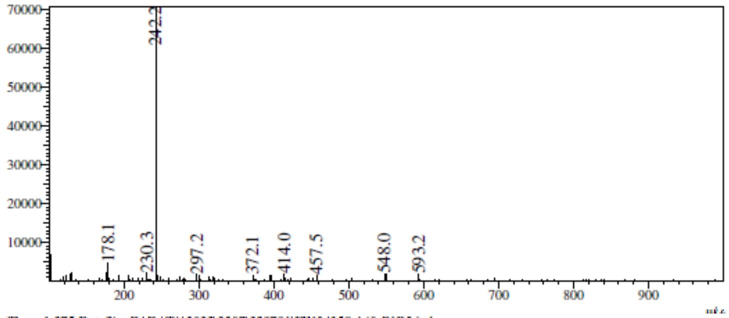

RetTime: 1.275 Datafile: D:\DATA\2022\2207\220701\EW34358-160-P1B2.lcd

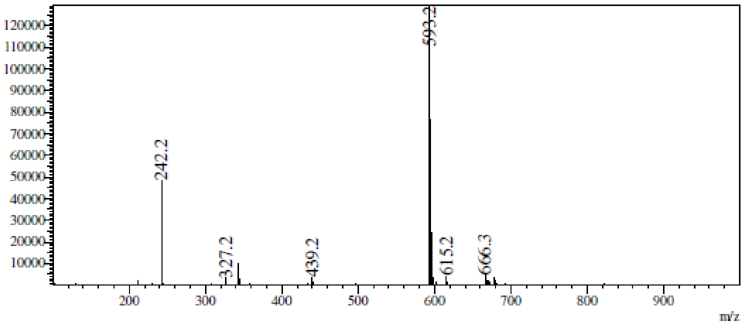

RetTime: 1.292 Datafile: D:\DATA\2022\2207\220701\EW34358-160-P1B2.lcd

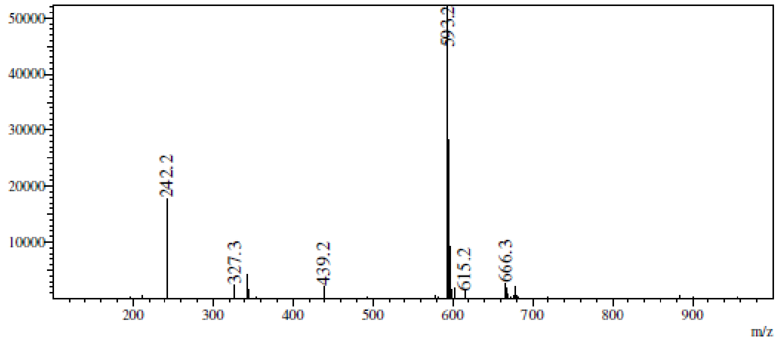

**$^1$H NMR** [(2*R*,3*R*,4*R*,5*R*)-3,4-Bis[[tert-butyl(dimethyl)silyl]oxy]-5-[6- (cyclopentylamino)purin-9-yl]tetrahydrofuran-2-yl]methyl methanesulfonate **(98a)**

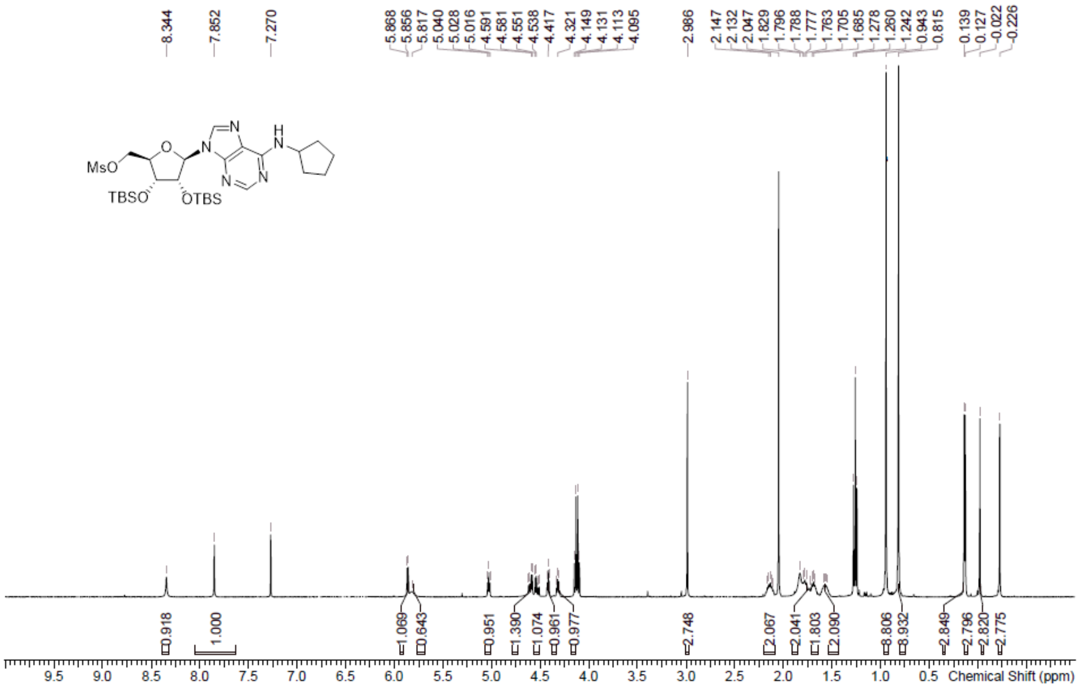

**LCMS** [(2*R*,3*R*,4*R*,5*R*)-3,4-Bis[[tert-butyl(dimethyl)silyl]oxy]-5-[6- (cyclopentylamino)purin-9-yl]tetrahydrofuran-2-yl]methyl methanesulfonate **(98a)**

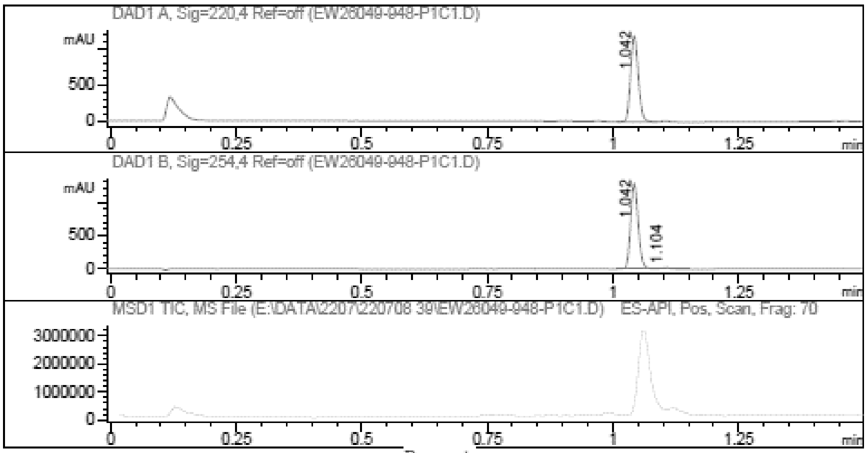

Report

Signal 1 : DAD1 A, Sig=220,4 Ref=off

| # | Meas. Ret. | Height | Width | Area | Area % |
|---|---|---|---|---|---|
| 1 | 1.042 | 1193.730 | 0.017 | 1272.045 | 100.000 |

Signal 2 : DAD1 B, Sig=254,4 Ref=off

| # | Meas. Ret. | Height | Width | Area | Area % |
|---|---|---|---|---|---|
| 1 | 1.042 | 1282.981 | 0.017 | 1355.217 | 98.286 |
| 2 | 1.104 | 19.359 | 0.018 | 23.638 | 1.714 |

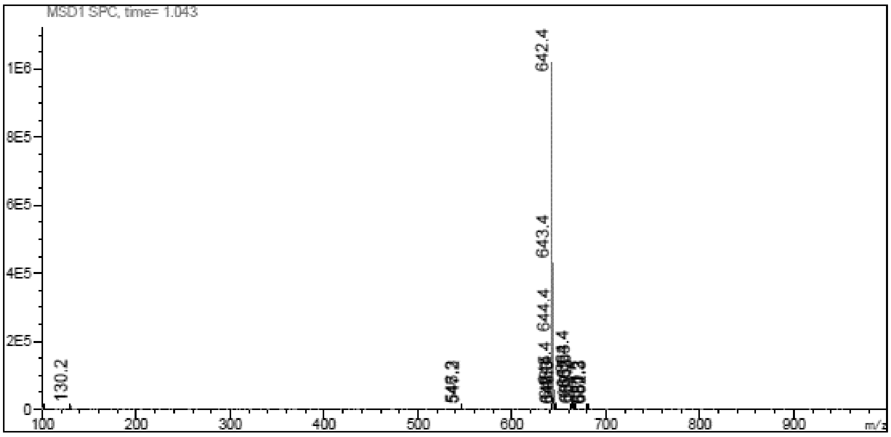

**$^{1}$H NMR** ((2R,3R,4R,5R)-3,4-bis((tert-butyldimethylsilyl)oxy)-5-(6-(isopropylamino)-9H-purin-9-yl)tetrahydrofuran-2-yl)methyl methanesulfonate **(98b)**

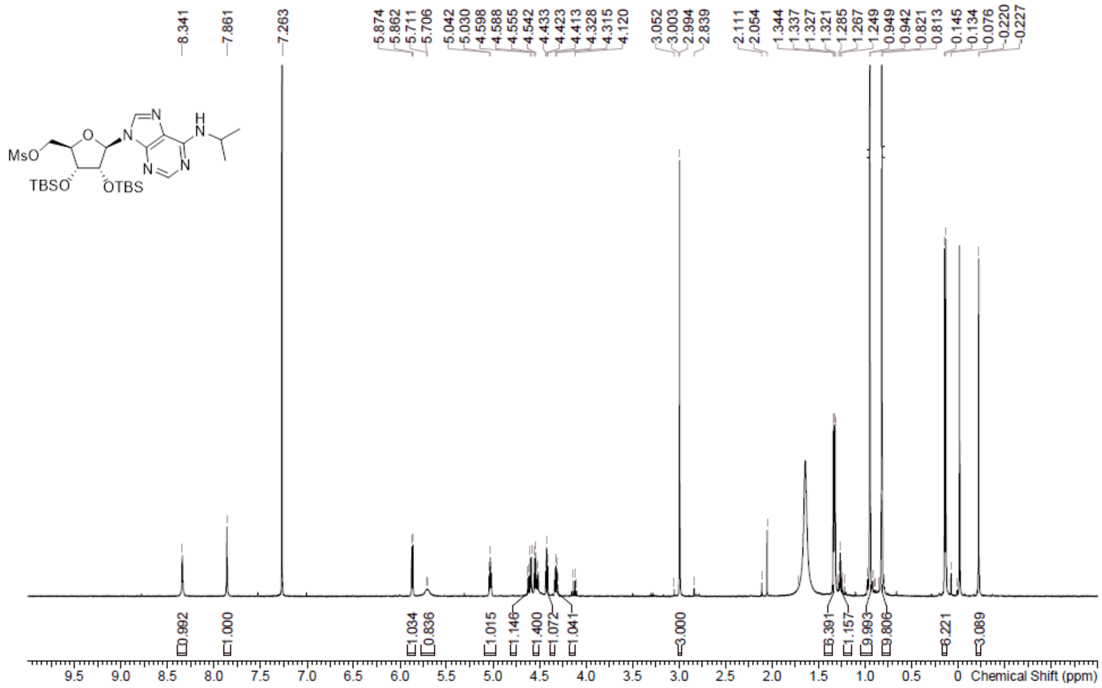

**LCMS** ((2R,3R,4R,5R)-3,4-bis((tert-butyldimethylsilyl)oxy)-5-(6-(isopropylamino)-9H-purin-9-yl)tetrahydrofuran-2-yl)methyl methanesulfonate **(98b)**

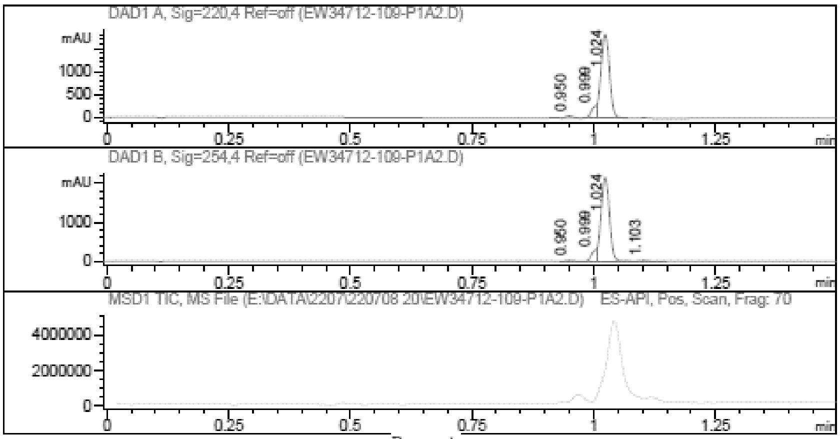

Report

Signal 1 : DAD1 A, Sig=220,4 Ref=off

| # | Meas. Ret. | Height | Width | Area | Area % |
|---|---|---|---|---|---|
| 1 | 0.950 | 45.902 | 0.015 | 46.440 | 1.855 |
| 2 | 0.999 | 229.513 | 0.012 | 181.405 | 7.246 |
| 3 | 1.024 | 1824.052 | 0.020 | 2275.589 | 90.899 |

Signal 2 : DAD1 B, Sig=254,4 Ref=off

| # | Meas. Ret. | Height | Width | Area | Area % |
|---|---|---|---|---|---|
| 1 | 0.950 | 19.879 | 0.016 | 20.378 | 0.707 |
| 2 | 0.999 | 251.896 | 0.012 | 197.907 | 6.865 |
| 3 | 1.024 | 2149.022 | 0.019 | 2637.716 | 91.503 |
| 4 | 1.103 | 21.182 | 0.019 | 26.654 | 0.925 |

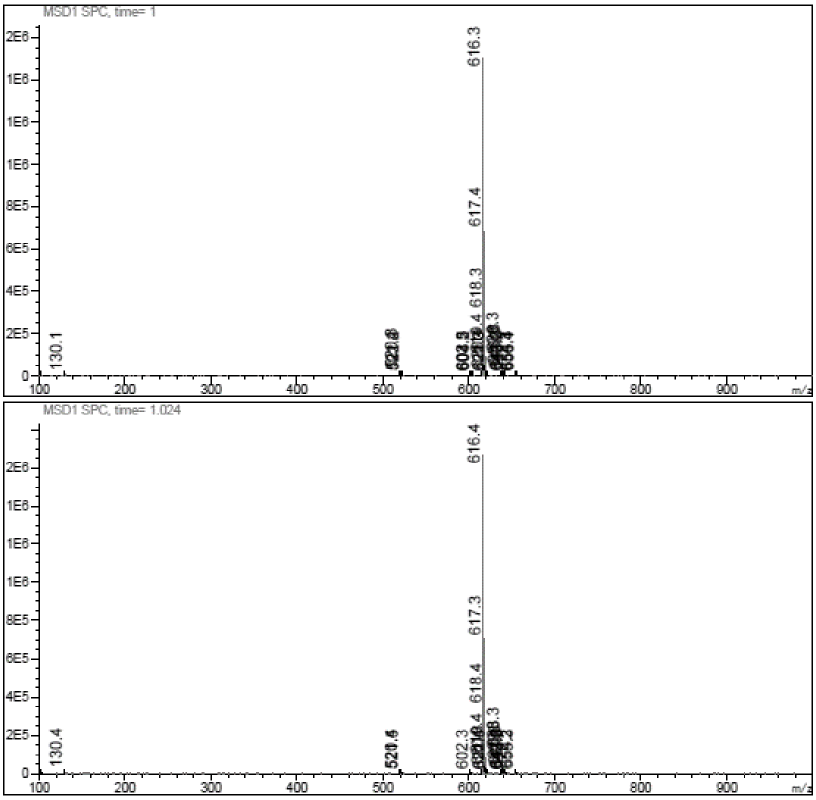

**$^1$H NMR** [(2*R*,3*R*,4*R*,5*R*)-3,4-Bis[[tert-butyl(dimethyl)silyl]oxy]-5-[6-(oxetan -3-ylamino)purin-9-yl]tetrahydrofuran-2-yl]methyl methanesulfonate **(98c)**

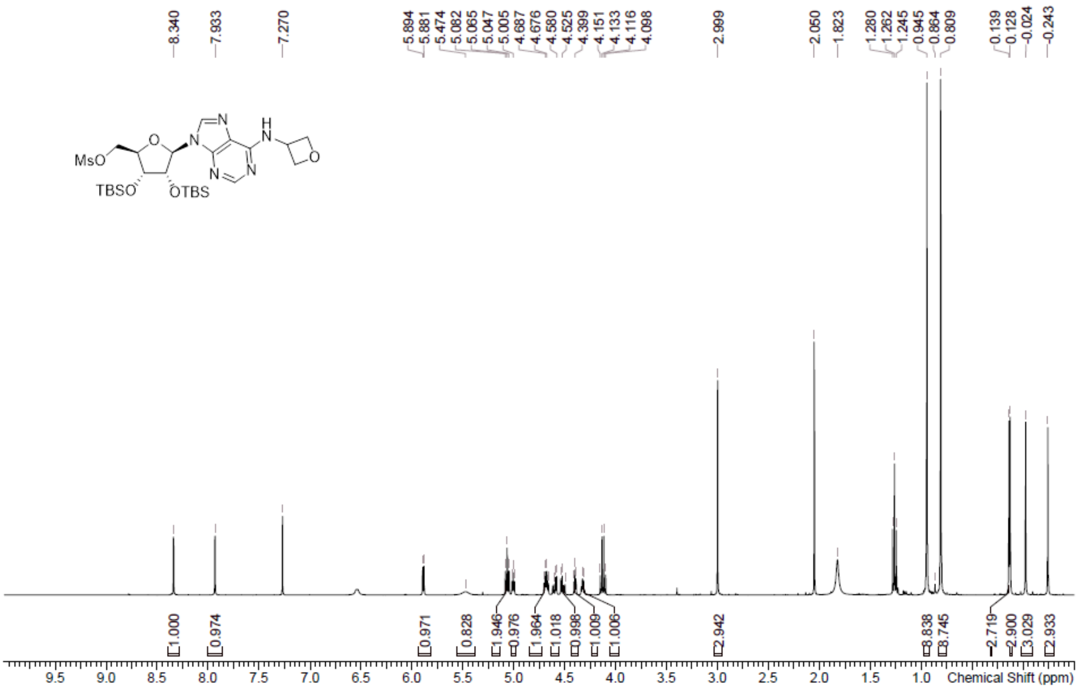

**LCMS** [(2*R*,3*R*,4*R*,5*R*)-3,4-Bis[[tert-butyl(dimethyl)silyl]oxy]-5-[6-(oxetan -3-ylamino)purin-9-yl]tetrahydrofuran-2-yl]methyl methanesulfonate **(98c)**

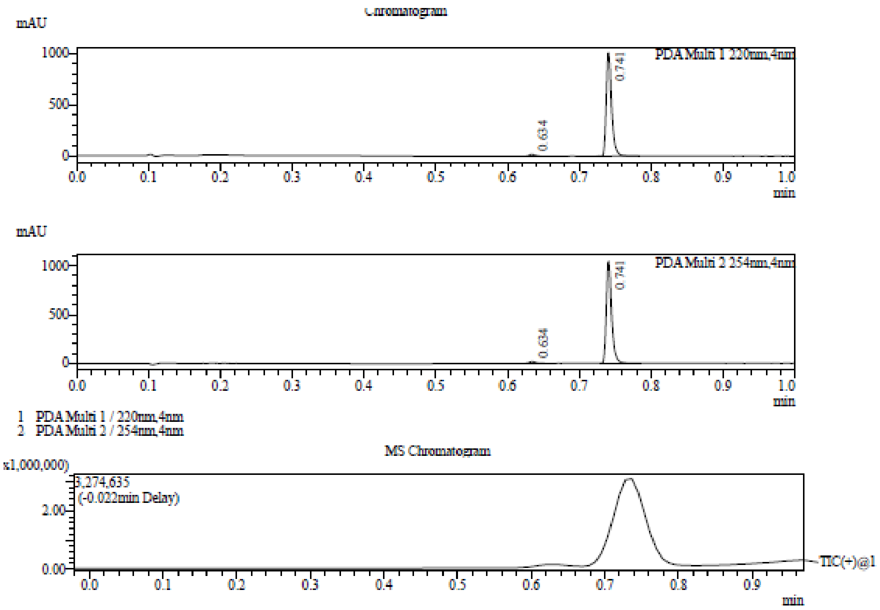

Integration Result

Peak Table

PDA Ch1 220nm

| Peak# | Ret. Time | Height | Height% | USP Width | Area | Area% |
|---|---|---|---|---|---|---|
| 1 | 0.634 | 21291 | 2.064 | 0.018 | 12874 | 2.408 |
| 2 | 0.741 | 1010192 | 97.936 | 0.015 | 521661 | 97.592 |

Peak Table

PDA Ch2 254nm

| Peak# | Ret. Time | Height | Height% | USP Width | Area | Area% |
|---|---|---|---|---|---|---|
| 1 | 0.634 | 19927 | 1.871 | 0.019 | 13277 | 2.449 |
| 2 | 0.741 | 1045336 | 98.129 | 0.014 | 528851 | 97.551 |

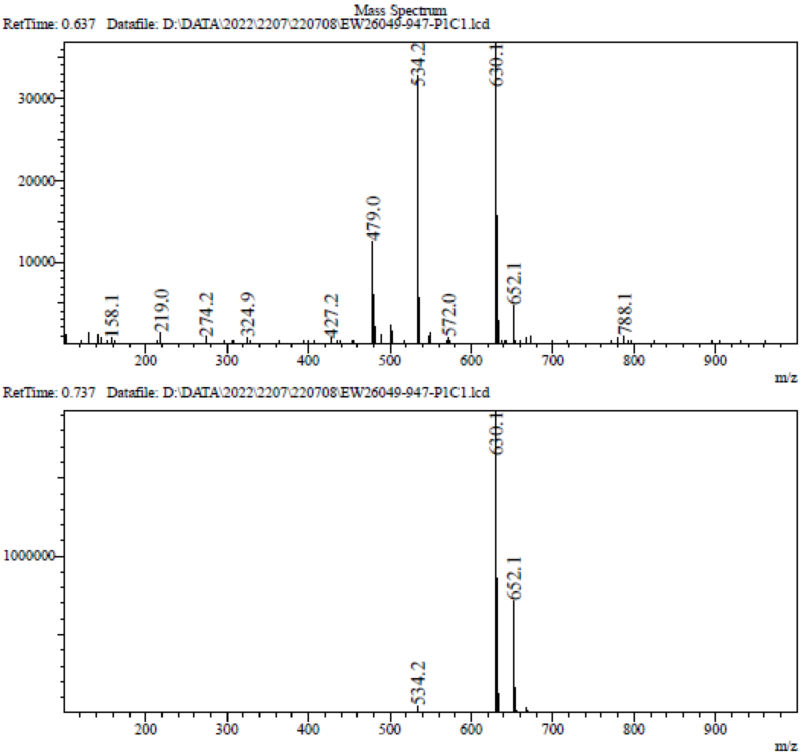

**$^1$H NMR** 9-[(2*R*,3*R*,4*R*,5*S*)-5-(1,3-Benzothiazol-2-ylsulfanylmethyl)-3,4-bis [[tert-butyl(dimethyl) silyl]oxy]tetrahydrofuran-2-yl]-*N*-cyclopentyl-purin-6-amine **(99a)**

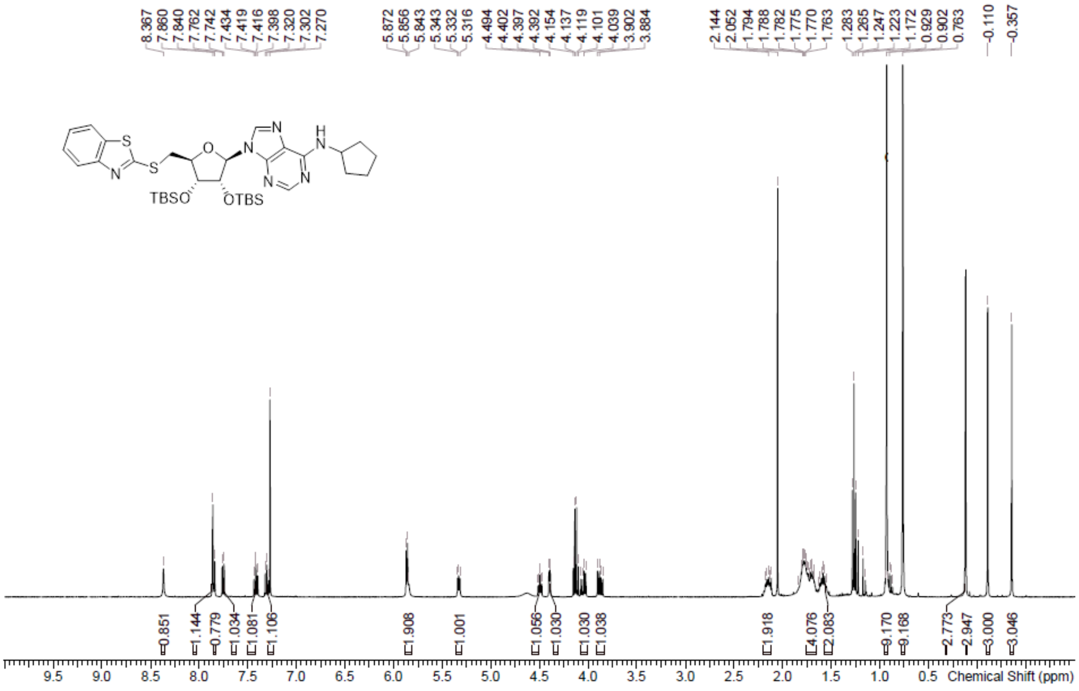

**LCMS** 9-[(2*R*,3*R*,4*R*,5*S*)-5-(1,3-Benzothiazol-2-ylsulfanylmethyl)-3,4-bis [[tert-butyl(dimethyl) silyl]oxy]tetrahydrofuran-2-yl]-*N*-cyclopentyl-purin-6-amine **(99a)**

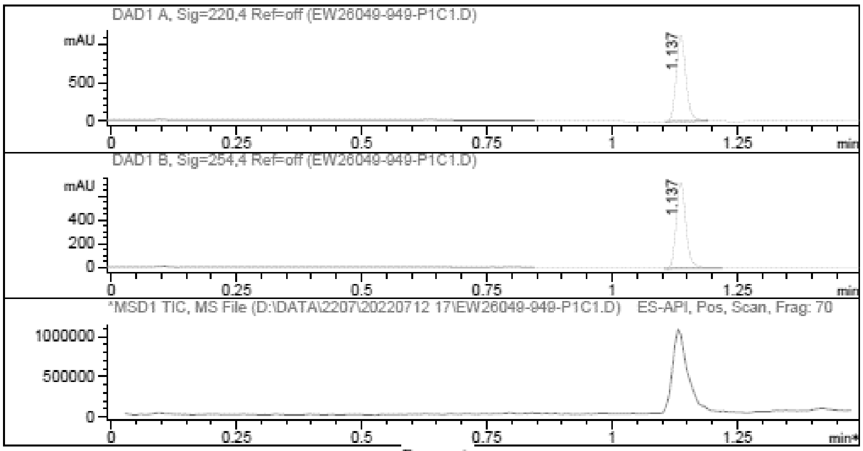

Report

Signal 1 : DAD1 A, Sig=220,4 Ref=off

| # | Meas. Ret. | Height | Width | Area | Area % |
|---|---|---|---|---|---|
| 1 | 1.137 | 1146.083 | 0.021 | 1555.363 | 100.000 |

Signal 2 : DAD1 B, Sig=254,4 Ref=off

| # | Meas. Ret. | Height | Width | Area | Area % |
|---|---|---|---|---|---|
| 1 | 1.137 | 733.089 | 0.021 | 991.781 | 100.000 |

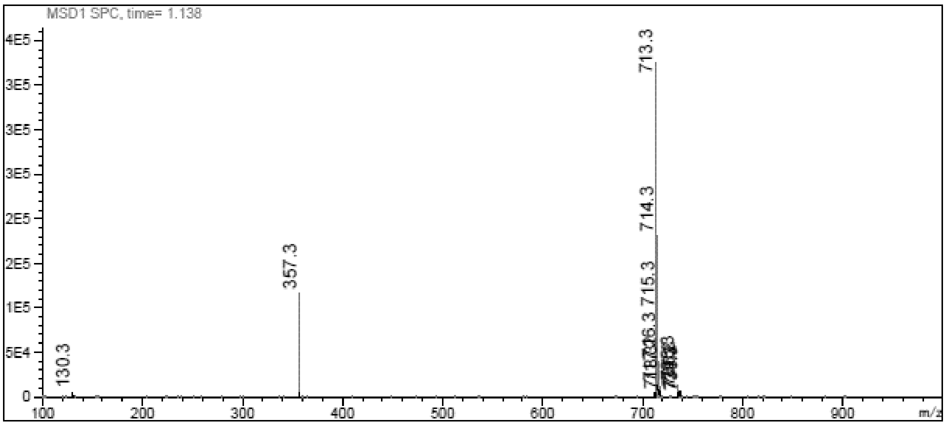

**$^1$H NMR** 9-((2R,3R,4R,5S)-5-((Benzo[d]thiazol-2-ylthio)methyl)-3,4-bis((tert-butyldimethylsilyl) oxy)tetrahydrofuran-2-yl)-N-isopropyl-9H-purin-6-amine **(99b)**

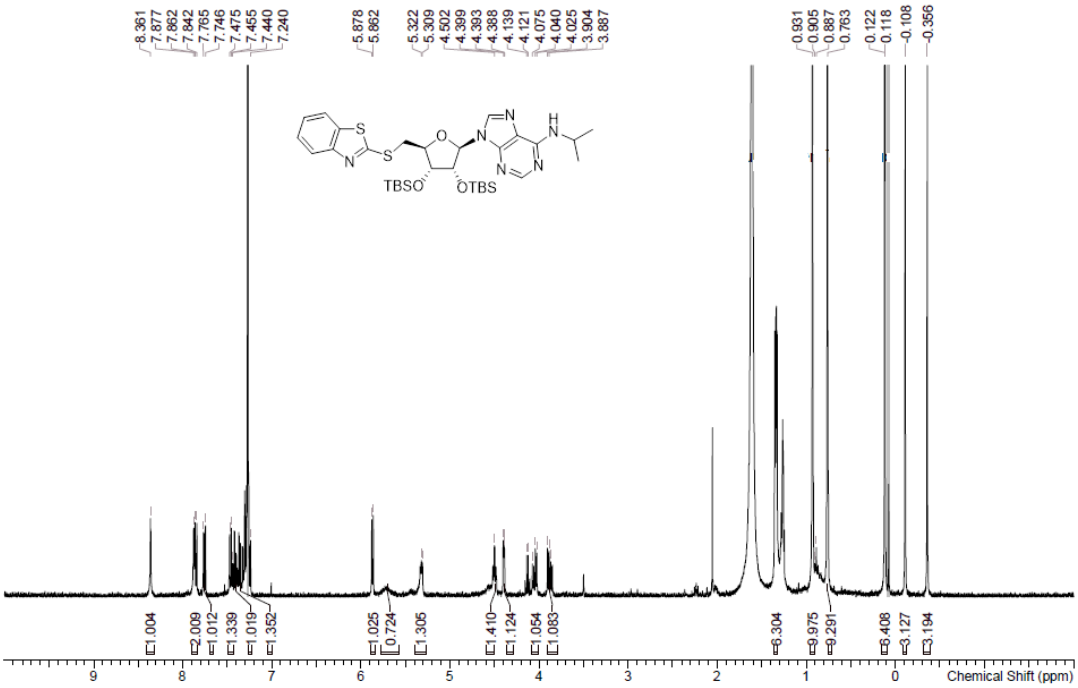

**LCMS** 9-((2R,3R,4R,5S)-5-((Benzo[d]thiazol-2-ylthio)methyl)-3,4-bis((tert-butyldimethylsilyl) oxy)tetrahydrofuran-2-yl)-N-isopropyl-9H-purin-6-amine **(99b)**

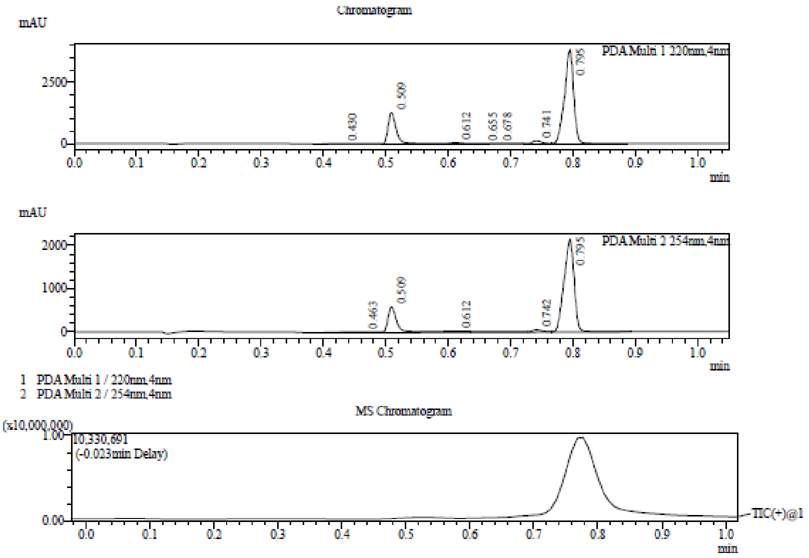

Integration Result

Peak Table

PDA Ch1 220nm

| Peak# | Ret. Time | Height | Height% | USP Width | Area | Area% |
|---|---|---|---|---|---|---|
| 1 | 0.430 | 4972 | 0.096 | 0.168 | 21712 | 0.411 |
| 2 | 0.509 | 1251280 | 24.182 | 0.022 | 1155805 | 21.852 |
| 3 | 0.612 | 37196 | 0.719 | 0.040 | 65123 | 1.231 |
| 4 | 0.655 | 8850 | 0.171 | 0.701 | 11215 | 0.212 |
| 5 | 0.678 | 7442 | 0.144 | 0.152 | 21279 | 0.402 |
| 6 | 0.741 | 111391 | 2.153 | 0.025 | 114920 | 2.173 |
| 7 | 0.795 | 3753388 | 72.536 | 0.025 | 3899128 | 73.719 |

Peak Table

PDA Ch2 254nm

| Peak# | Ret. Time | Height | Height% | USP Width | Area | Area% |
|---|---|---|---|---|---|---|
| 1 | 0.463 | 14669 | 0.530 | 0.322 | 73035 | 2.229 |
| 2 | 0.509 | 586325 | 21.186 | 0.023 | 649717 | 19.830 |

Mass Spectrum

RetTime: 0.427 Datafile: D:\DATA\2022\2207\220712\EW34712-112-P1A3.lcd

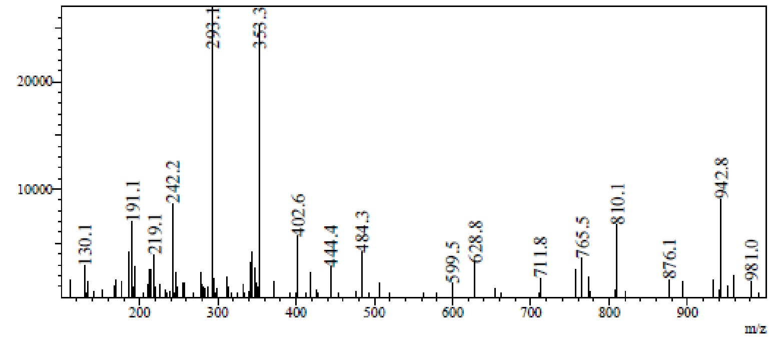

RetTime: 0.510 Datafile: D:\DATA\2022\2207\220712\EW34712-112-P1A3.lcd

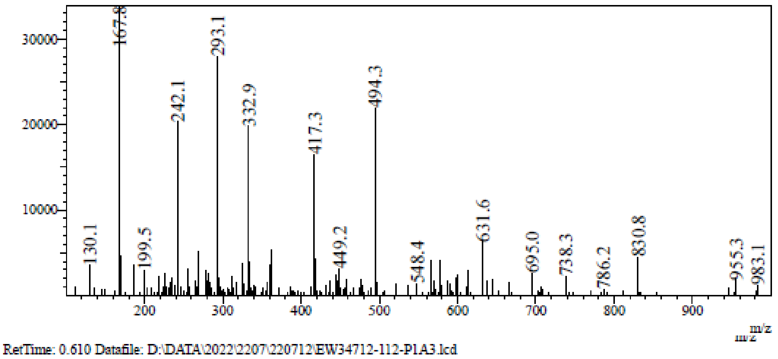

RetTime: 0.610 Datafile: D:\DATA\2022\2207\220712\EW34712-112-P1A3.lcd

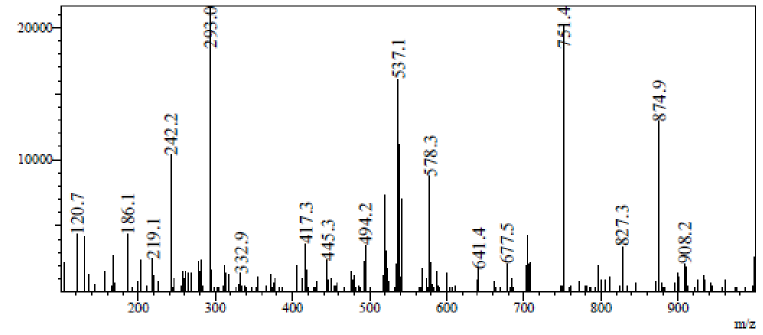

RetTime: 0.677 Datafile: D:\DATA\2022\2207\220712\EW34712-112-P1A3.lcd

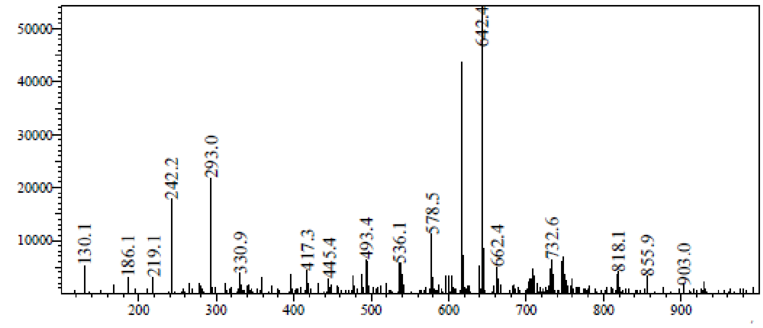

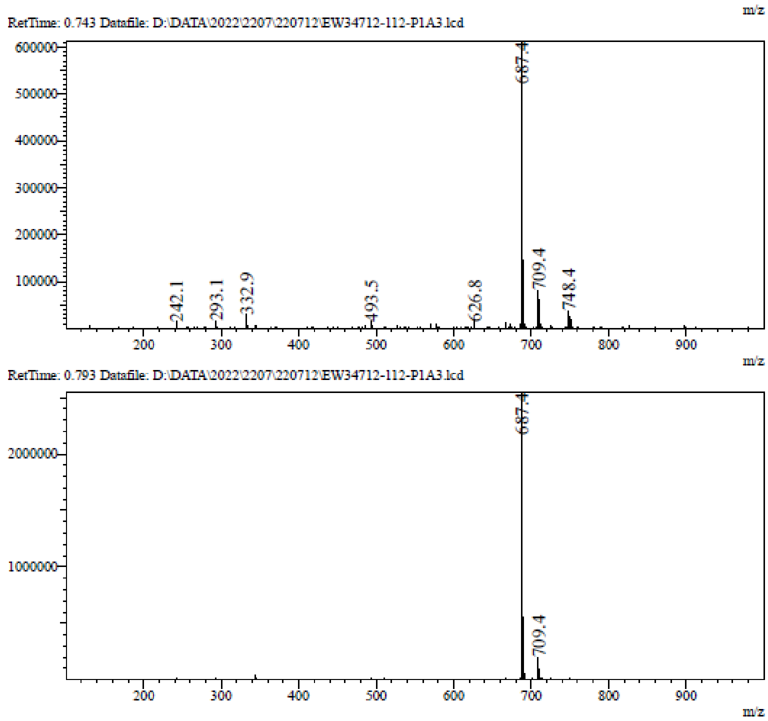

**$^1$H NMR** 9-[(2*R*,3*R*,4*R*,5*S*)-5-(1,3-Benzothiazol-2-ylsulfanylmethyl)-3,4-bis [[tert-butyl(dimethyl) silyl]oxy]tetrahydrofuran-2-yl]-*N*-(oxetan-3-yl)purin-6-amine **(99c)**

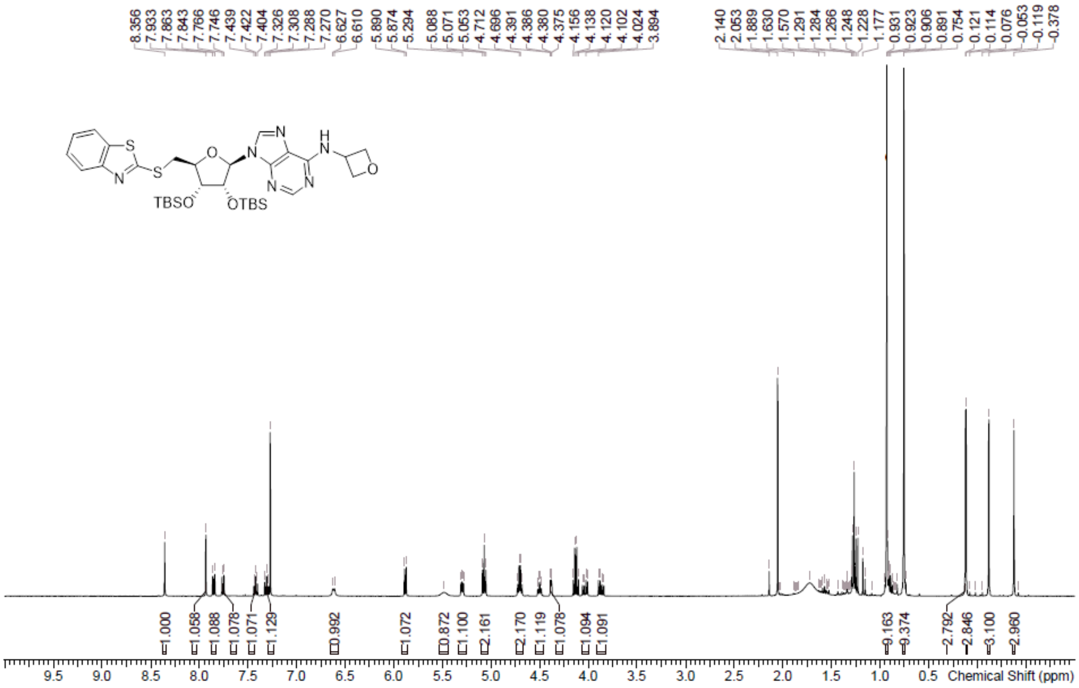

**LCMS** 9-[(2*R*,3*R*,4*R*,5*S*)-5-(1,3-Benzothiazol-2-ylsulfanylmethyl)-3,4-bis [[tert-butyl(dimethyl) silyl]oxy]tetrahydrofuran-2-yl]-*N*-(oxetan-3-yl)purin-6-amine **(99c)**

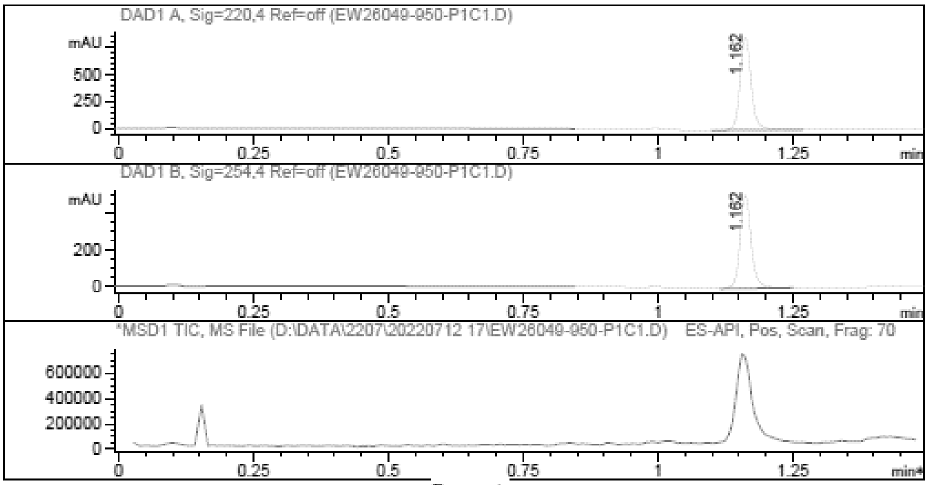

Report

Signal 1 : DAD1 A, Sig=220,4 Ref=off

| # | Meas. Ret. | Height | Width | Area | Area % |
|---|---|---|---|---|---|
| 1 | 1.162 | 861.984 | 0.022 | 1249.093 | 100.000 |

Signal 2 : DAD1 B, Sig=254,4 Ref=off

| # | Meas. Ret. | Height | Width | Area | Area % |
|---|---|---|---|---|---|
| 1 | 1.162 | 504.668 | 0.022 | 724.241 | 100.000 |

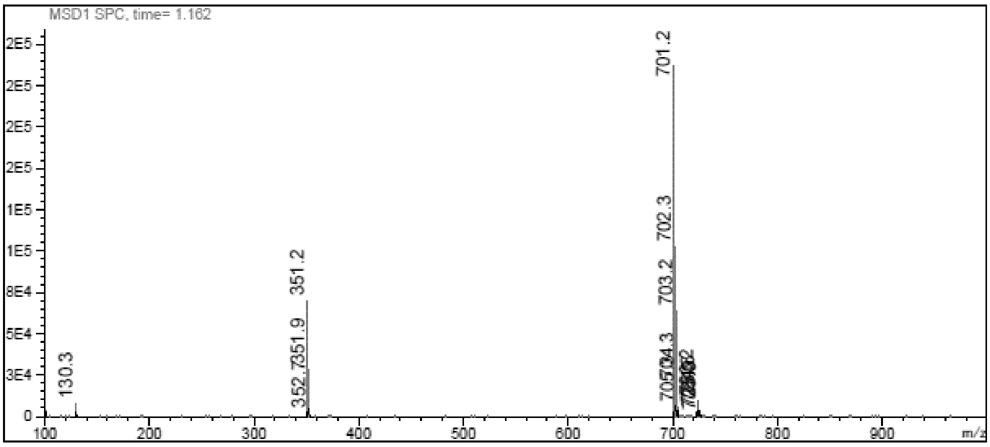

**$^1$H NMR** (2*S*,3*S*,4*R*,5*R*)-2-(1,3-Benzothiazol-2-ylsulfanylmethyl)-5- [6-(cyclopentylamino)purin-9-yl]tetrahydrofuran-3,4-diol **(20)**

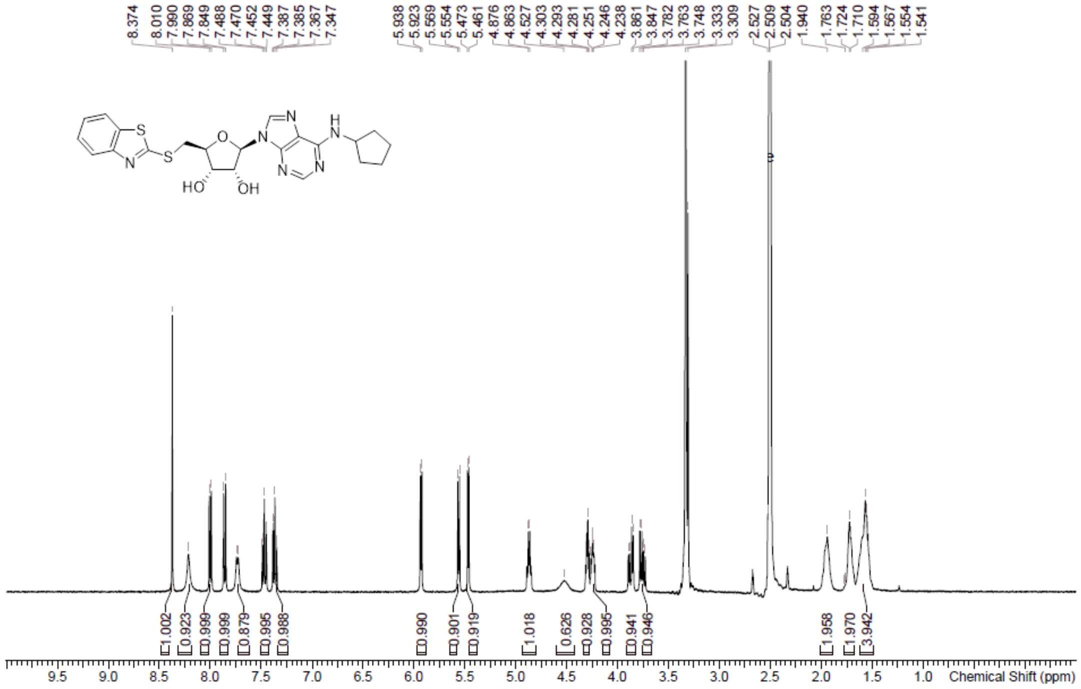

**HPLC** (2*S*,3*S*,4*R*,5*R*)-2-(1,3-Benzothiazol-2-ylsulfanylmethyl)-5- [6-(cyclopentylamino)purin-9-yl]tetrahydrofuran-3,4-diol **(20)**

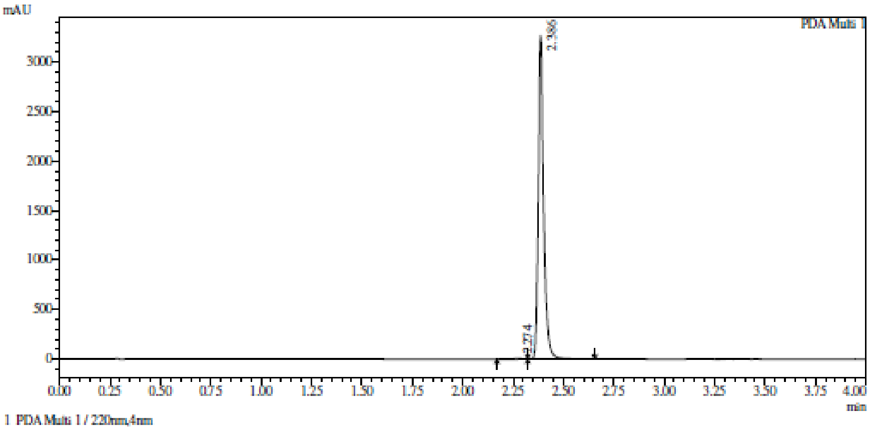

Integration result

PeakTable

PDA Ch1 220nm

| Peak# | Ret. Time | USP Width | Resolution | Height | Area | Area % |
|---|---|---|---|---|---|---|
| 1 | 2.274 | 0.306 | 0.000 | 5003 | 26154 | 0.427 |
| 2 | 2.386 | 0.044 | 0.643 | 3267417 | 6099750 | 99.573 |
| Total | | | | 3272420 | 6125904 | 100.000 |

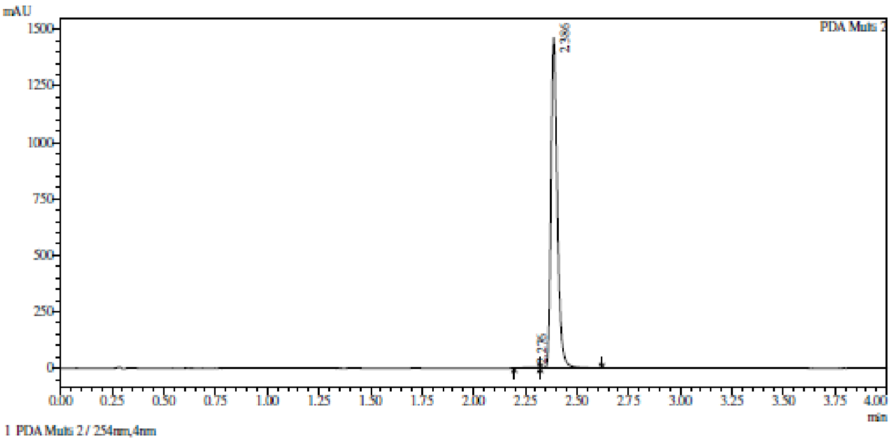

Integration result

PeakTable

PDA Ch2 254nm

| Peak# | Ret. Time | USP Width | Resolution | Height | Area | Area % |
|---|---|---|---|---|---|---|
| 1 | 2.276 | 0.156 | 0.000 | 2919 | 13658 | 0.434 |
| 2 | 2.386 | 0.056 | 1.040 | 1463094 | 3132168 | 99.566 |
| Total | | | | 1466014 | 3145826 | 100.000 |

**HRMS** (2*S*,3*S*,4*R*,5*R*)-2-(1,3-Benzothiazol-2-ylsulfanylmethyl)-5- [6-(cyclopentylamino)purin-9-yl]tetrahydrofuran-3,4-diol **(20)**

**Compound Table**

| Label | Tgt Score | Mass Error (ppm) | Tgt Formula | Obs. RT | Ref. Mass | Obs. Mass |
|---|---|---|---|---|---|---|
| Cpd 1: C22 H24 N6 O3 S2; 2.106 | 98.42 | 4.38 | C22 H24 N6 O3 S2 | 2.106 | 484.1351 | 484.1373 |

| Obs. *m/z* | Obs. RT | Obs. Mass | Tgt Formula | Tgt Mass | Tgt Mass Error (ppm) | Find Cpds Algorithm |
|---|---|---|---|---|---|---|
| 507.1251 | 2.106 | 484.1373 | C22 H24 N6 O3 S2 | 484.1351 | 4.38 | Find by Formula |

**Compound Chromatograms**

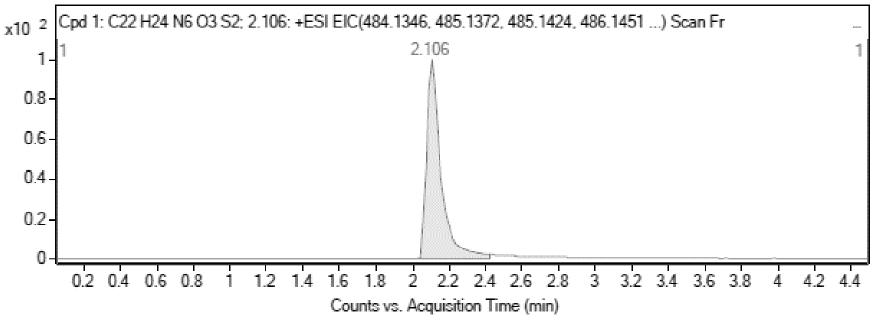

**MS Zoomed Spectrum**

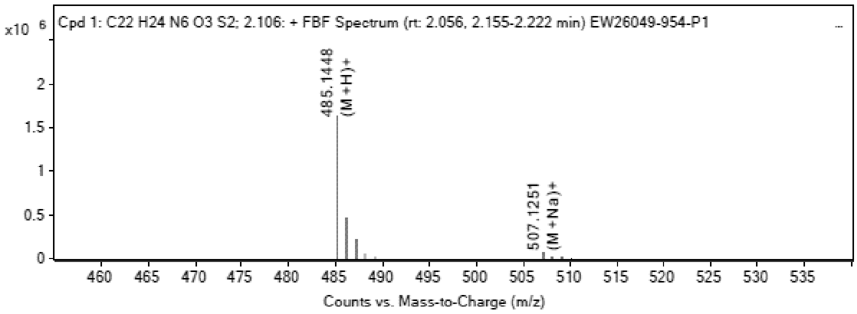

**MS Spectrum Peak List**

| Obs. *m/z* | Charge | Abund | Ion/Isotope |
|---|---|---|---|
| 485.1448 | 1 | 1631015 | (M+H)+ |
| 486.1466 | 1 | 455562.72 | (M+H)+ |

| | | | |
|---|---|---|---|
| 487.1428 | 1 | 205090.11 | (M+H)+ |
| 507.1251 | 1 | 62494.4 | (M+Na)+ |
| 508.1276 | 1 | 17014.72 | (M+Na)+ |
| 509.1246 | 1 | 8084.87 | (M+Na)+ |
| 510.1248 | 1 | 1715.72 | (M+Na)+ |

**MS Zoomed Spectrum**

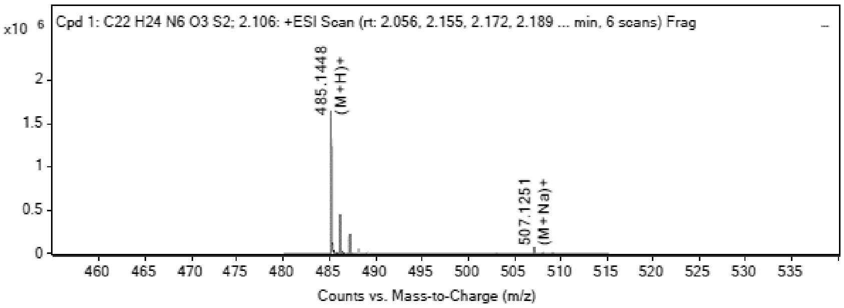

**MS Spectrum Peak List**

| Obs. m/z | Charge | Abund | Ion/Isotope | Tgt Mass Error (ppm) |
|---|---|---|---|---|
| 485.1448 | | 1631001.66 | | -4.98 |
| 485.1448 | 1 | 1631015 | (M+H)+ | -5 |
| 486.1466 | 1 | 455562.72 | (M+H)+ | -3.23 |
| 487.1428 | 1 | 205090.11 | (M+H)+ | -3.13 |
| 507.1251 | 1 | 62494.4 | (M+Na)+ | -1.57 |
| 508.1276 | 1 | 17014.72 | (M+Na)+ | -1.18 |
| 509.1246 | 1 | 8084.87 | (M+Na)+ | -2.67 |
| 510.1248 | 1 | 1715.72 | (M+Na)+ | -0.75 |

--- End Of Report ---

**$^{1}$H NMR** (2S,3S,4R,5R)-2-((benzo[d]thiazol-2-ylthio)methyl)-5-(6-(isopropylamino)-9H-purin-9-yl)tetrahydrofuran-3,4-diol **(21)**

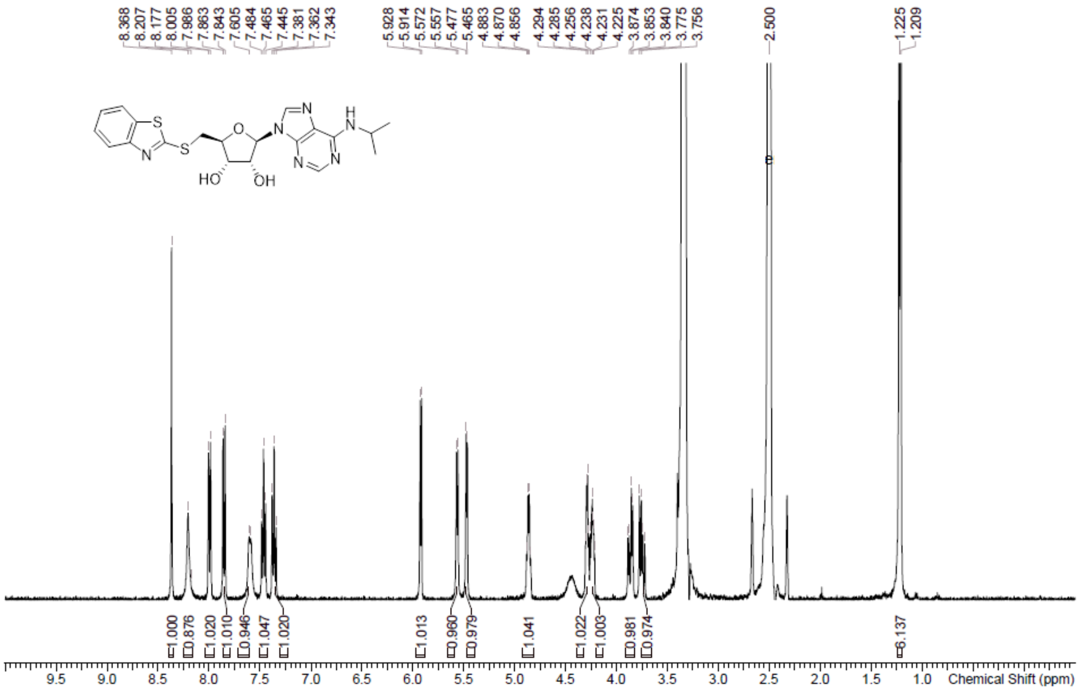

**HPLC** (2S,3S,4R,5R)-2-((benzo[d]thiazol-2-ylthio)methyl)-5-(6-(isopropylamino)-9H-purin-9-yl)tetrahydrofuran-3,4-diol **(21)**

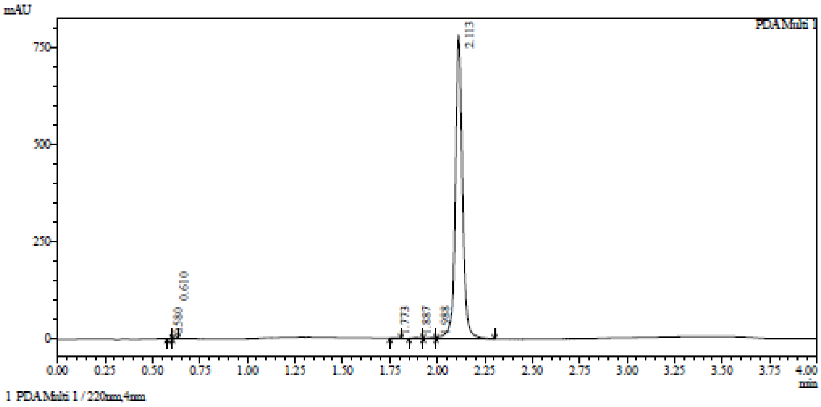

Integration result

PeakTable

PDA Ch1 220nm

| Peak# | Ret. Time | USP Width | Resolution | Height | Area | Area % |
|---|---|---|---|---|---|---|
| 1 | 0.580 | 0.105 | 0.000 | 1047 | 1749 | 0.084 |
| 2 | 0.610 | 0.083 | 0.322 | 1237 | 1869 | 0.090 |
| 3 | 1.773 | 0.070 | 15.234 | 1507 | 3648 | 0.175 |
| 4 | 1.887 | 0.071 | 1.620 | 2577 | 6517 | 0.312 |
| 5 | 1.988 | 0.112 | 1.106 | 3452 | 8442 | 0.405 |
| 6 | 2.113 | 0.066 | 1.399 | 781817 | 2063561 | 98.934 |
| Total | | | | 791636 | 2085786 | 100.000 |

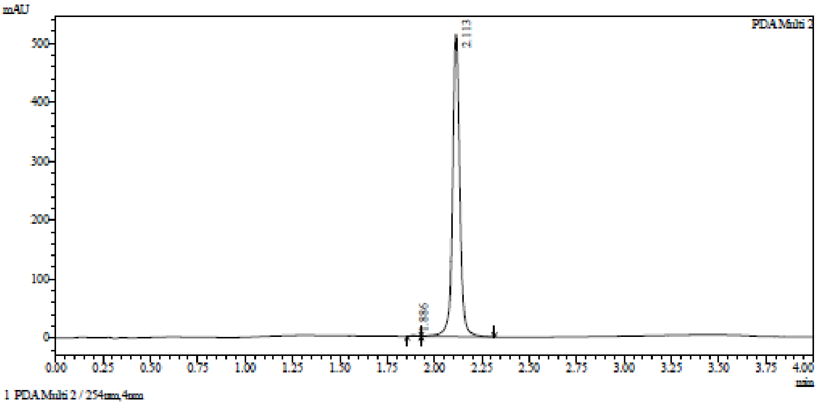

Integration result

PeakTable

PDA Ch2 254nm

| Peak# | Ret. Time | USP Width | Resolution | Height | Area | Area % |
|---|---|---|---|---|---|---|
| 1 | 1.886 | 0.065 | 0.000 | 1655 | 3797 | 0.278 |
| 2 | 2.113 | 0.066 | 3.452 | 513140 | 1359565 | 99.722 |
| Total | | | | 514794 | 1363362 | 100.000 |

**HRMS** (2S,3S,4R,5R)-2-((benzo[d]thiazol-2-ylthio)methyl)-5-(6-(isopropylamino)-9H-purin-9-yl)tetrahydrofuran-3,4-diol **(21)**

Compound Table

| Label | Tgt Score | Mass Error (ppm) | Tgt Formula | Obs. RT | Ref. Mass | Obs. Mass |
|---|---|---|---|---|---|---|
| Cpd 1: C20 H22 N6 O3 S2; 1.966 | 96.29 | 4.56 | C20 H22 N6 O3 S2 | 1.966 | 458.1195 | 458.1216 |

| Obs. *m/z* | Obs. RT | Obs. Mass | Tgt Formula | Tgt Mass | Tgt Mass Error (ppm) | Find Cpds Algorithm |
|---|---|---|---|---|---|---|
| 481.1097 | 1.966 | 458.1216 | C20 H22 N6 O3 S2 | 458.1195 | 4.56 | Find by Formula |

Compound Chromatograms

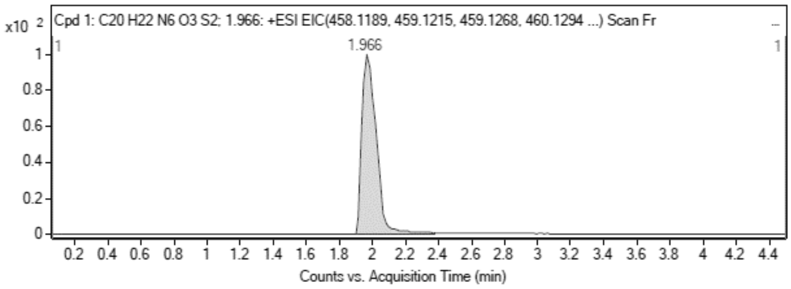

MS Zoomed Spectrum

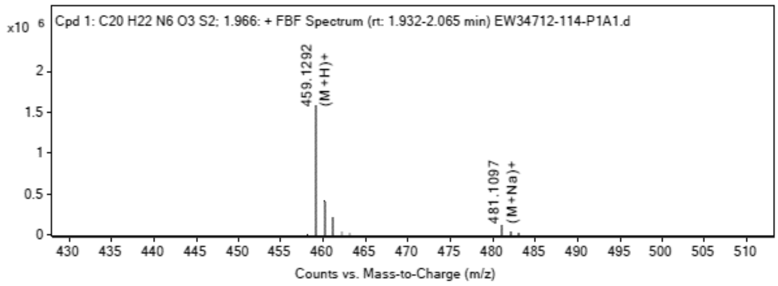

MS Spectrum Peak List

| Obs. *m/z* | Charge | Abund | Ion/Isotope |
|---|---|---|---|
| 458.1155 | 1 | 571.2 | M+ |
| 459.1292 | 1 | 1586056 | (M+H)+ |
| 460.131 | 1 | 420726.25 | (M+H)+ |
| 461.1268 | 1 | 196635.44 | (M+H)+ |
| 481.1097 | 1 | 117045.18 | (M+Na)+ |
| 482.1119 | 1 | 29380.34 | (M+Na)+ |
| 483.1077 | 1 | 13929.76 | (M+Na)+ |

MS Zoomed Spectrum

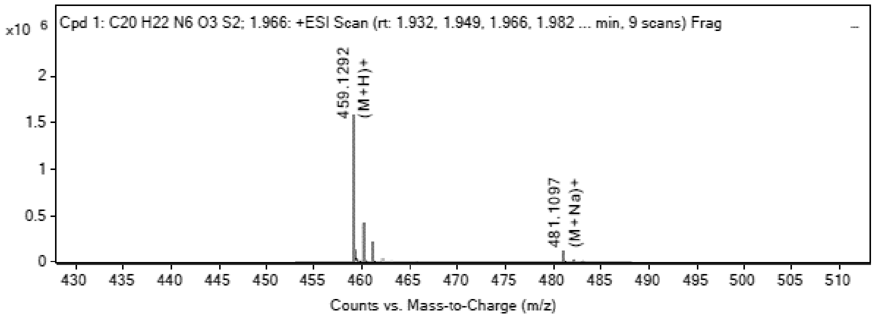

MS Spectrum Peak List

| Obs. *m/z* | Charge | Abund | Ion/Isotope | Tgt Mass Error (ppm) |
|---|---|---|---|---|
| 458.1155 | 1 | 571.2 | M+ | 7.41 |
| 459.1292 | | 1585992.68 | | -5.27 |
| 459.1292 | 1 | 1586056 | (M+H)+ | -5.28 |
| 460.131 | 1 | 420726.25 | (M+H)+ | -3.45 |
| 461.1268 | 1 | 196635.44 | (M+H)+ | -3.17 |
| 481.1097 | 1 | 117045.18 | (M+Na)+ | -2.09 |
| 482.1119 | 1 | 29380.34 | (M+Na)+ | -1.17 |
| 483.1077 | 1 | 13929.76 | (M+Na)+ | -0.94 |

--- End Of Report ---

**[1]H NMR** (2*S*,3*S*,4*R*,5*R*)-2-(1,3-Benzothiazol-2-ylsulfanylmethyl)-5- [6-(oxetan-3-ylamino)purin-9-yl]tetrahydrofuran-3,4-diol **(22)**

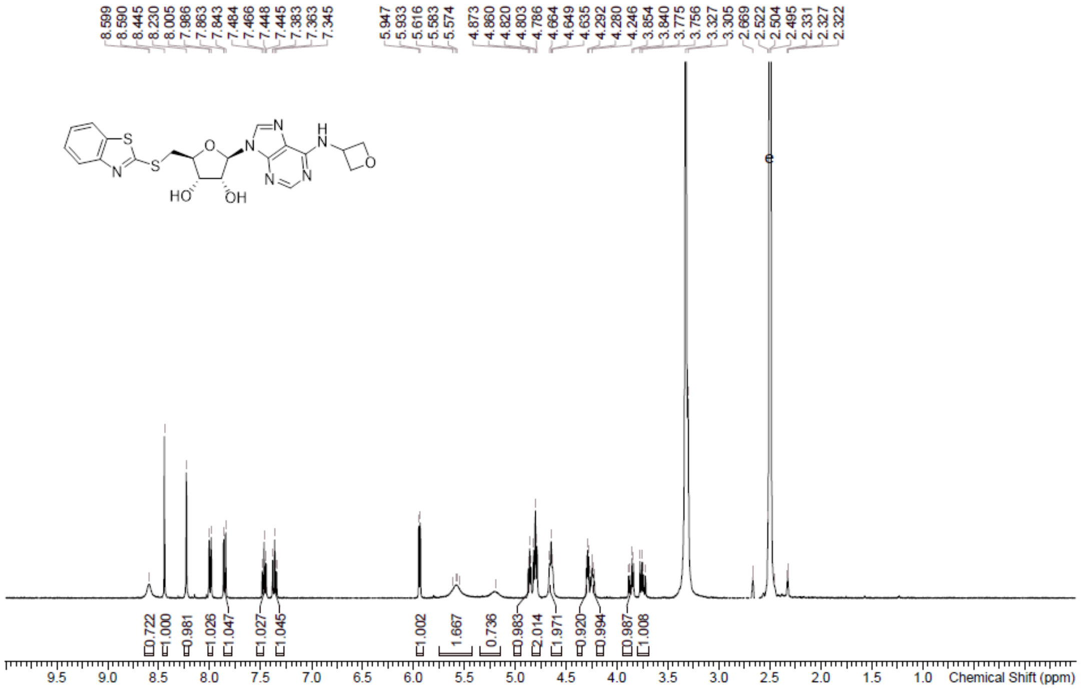

**HPLC** (2*S*,3*S*,4*R*,5*R*)-2-(1,3-Benzothiazol-2-ylsulfanylmethyl)-5- [6-(oxetan-3-ylamino)purin-9-yl]tetrahydrofuran-3,4-diol **(22)**

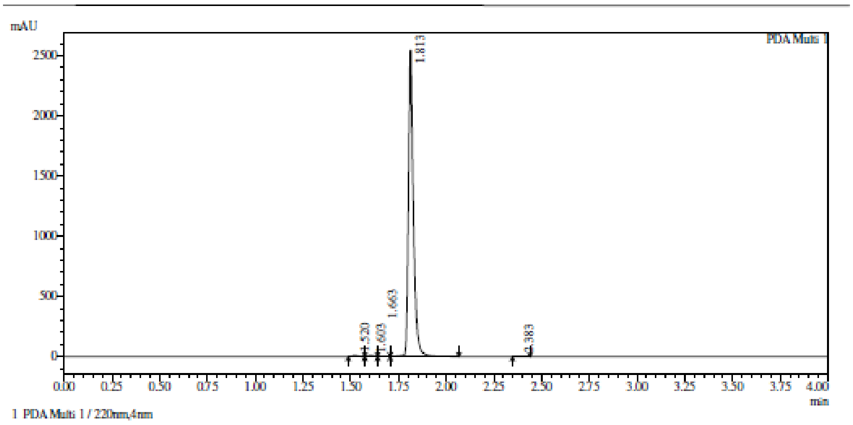

Integration result

PeakTable

PDA Ch1 220nm

| Peak# | Ret. Time | USP Width | Resolution | Height | Area | Area % |
|---|---|---|---|---|---|---|
| 1 | 1.520 | 0.052 | 0.000 | 11404 | 22626 | 0.494 |
| 2 | 1.603 | 0.057 | 1.524 | 11345 | 24466 | 0.534 |
| 3 | 1.663 | 0.107 | 0.719 | 4431 | 11797 | 0.258 |
| 4 | 1.813 | 0.046 | 1.966 | 2546174 | 4515952 | 98.617 |
| 5 | 2.383 | 0.065 | 10.284 | 1856 | 4454 | 0.097 |
| Total | | | | 2575210 | 4579296 | 100.000 |

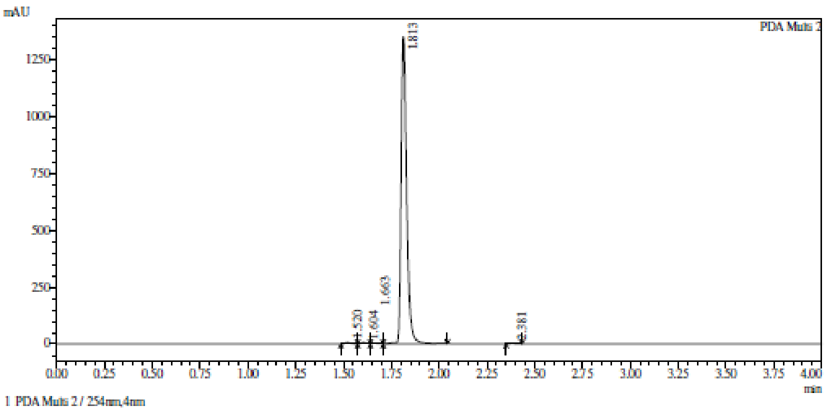

Integration result

PeakTable

PDA Ch2 254nm

| Peak# | Ret. Time | USP Width | Resolution | Height | Area | Area % |
|---|---|---|---|---|---|---|
| 1 | 1.520 | 0.052 | 0.000 | 6786 | 13329 | 0.487 |
| 2 | 1.604 | 0.057 | 1.530 | 6548 | 13936 | 0.509 |
| 3 | 1.663 | 0.097 | 0.776 | 2985 | 7865 | 0.287 |
| 4 | 1.813 | 0.052 | 2.010 | 1352967 | 2699705 | 98.628 |
| 5 | 2.381 | 0.062 | 9.987 | 1058 | 2433 | 0.089 |
| Total | | | | 1370343 | 2737267 | 100.000 |

**HRMC** (2*S*,3*S*,4*R*,5*R*)-2-(1,3-Benzothiazol-2-ylsulfanylmethyl)-5- [6-(oxetan-3-ylamino)purin-9-yl]tetrahydrofuran-3,4-diol **(22)**

Compound Table

| Label | Tgt Score | Mass Error (ppm) | Tgt Formula | Obs. RT | Ref. Mass | Obs. Mass |
|---|---|---|---|---|---|---|
| Cpd 1: C20 H20 N6 O4 S2; 1.901 | 93.11 | 6.11 | C20 H20 N6 O4 S2 | 1.901 | 472.0987 | 472.1016 |

| Obs. *m/z* | Obs. RT | Obs. Mass | Tgt Formula | Tgt Mass | Tgt Mass Error (ppm) | Find Cpds Algorithm |
|---|---|---|---|---|---|---|
| 495.0896 | 1.901 | 472.1016 | C20 H20 N6 O4 S2 | 472.0987 | 6.11 | Find by Formula |

Compound Chromatograms

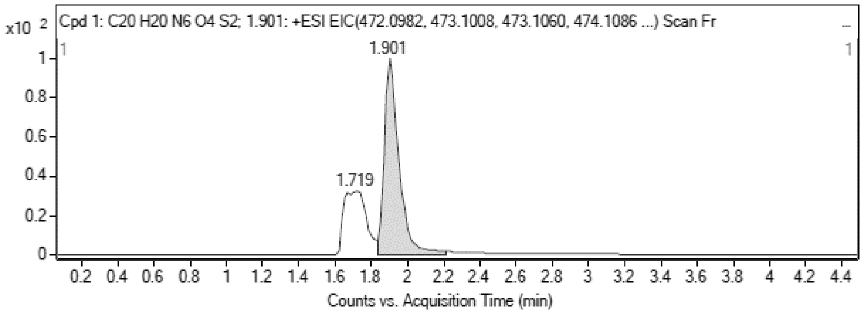

MS Zoomed Spectrum

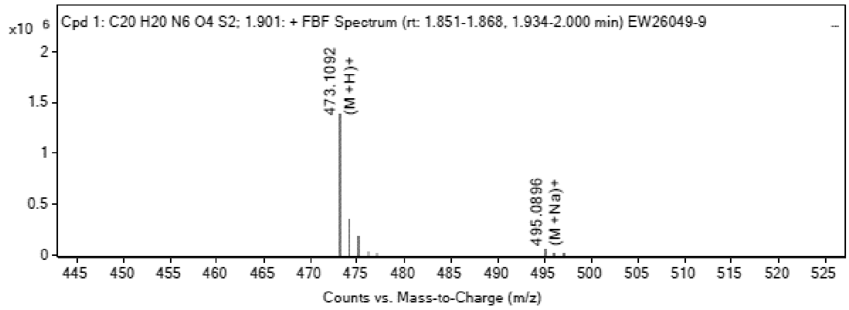

MS Spectrum Peak List

| Obs. *m/z* | Charge | Abund | Ion/Isotope |
|---|---|---|---|
| 473.1092 | 1 | 1384917.88 | (M+H)+ |
| 474.111 | 1 | 352973.34 | (M+H)+ |

| | | | |
|---|---|---|---|
| 475.1069 | 1 | 170793.89 | (M+H)+ |
| 495.0896 | 1 | 61714.53 | (M+Na)+ |
| 496.0916 | 1 | 15825.19 | (M+Na)+ |
| 497.0886 | 1 | 7490.06 | (M+Na)+ |

MS Zoomed Spectrum

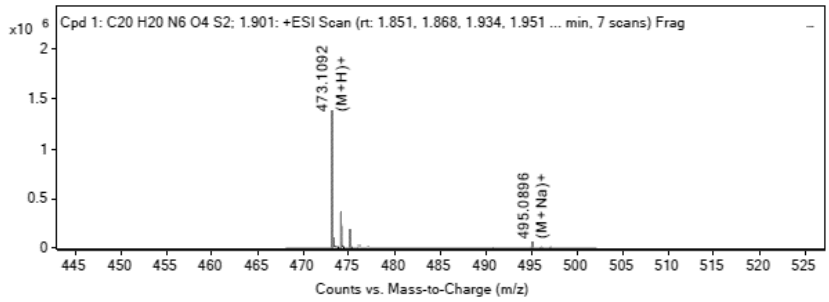

**MS Spectrum Peak List**

| Obs. *m/z* | Charge | Abund | Ion/Isotope | Tgt Mass Error (ppm) |
|---|---|---|---|---|
| 473.1092 | | 1384897.09 | | -6.7 |
| 473.1092 | 1 | 1384917.88 | (M+H)+ | -6.71 |
| 474.111 | 1 | 352973.34 | (M+H)+ | -5.04 |
| 475.1069 | 1 | 170793.89 | (M+H)+ | -4.72 |
| 495.0896 | 1 | 61714.53 | (M+Na)+ | -3.24 |
| 496.0916 | 1 | 15825.19 | (M+Na)+ | -2.05 |
| 497.0886 | 1 | 7490.06 | (M+Na)+ | -3.94 |

--- End Of Report ---

**LCMS** (2*S*,3*R*,4*R*,5*R*)-3,4-Bis[[tert-butyl(dimethyl)silyl]oxy]-5- (6-chloropurin-9-yl)tetrahydrofuran-2-carbaldehyde **(47)**

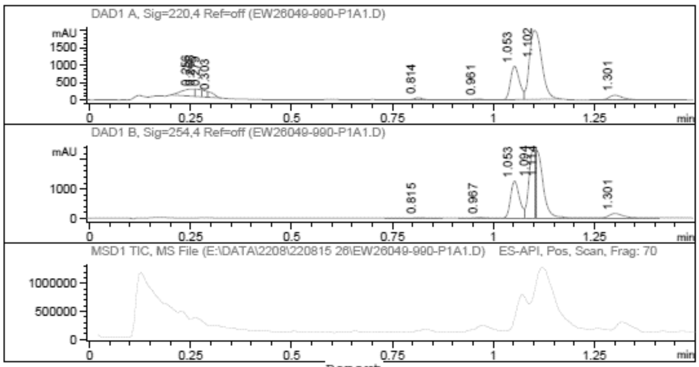

```
Report
=================================================================
Signal 1 : DAD1 A, Sig=220,4 Ref=off
 #  Meas. Ret.   Height    Width       Area       Area %
----------------------------------------------------------
 1      0.256    210.044   0.037        459.425    6.435
 2      0.266    211.443   0.015        206.461    2.892
 3      0.279    191.610   0.014        159.638    2.236
 4      0.303    130.858   0.018        160.967    2.255
 5      0.814     55.720   0.022         79.615    1.115
 6      0.961     23.491   0.025         37.717    0.528
 7      1.053    961.657   0.023       1471.560   20.611
 8      1.102   1981.939   0.033       4216.002   59.051
 9      1.301    136.826   0.038        348.233    4.877
----------------------------------------------------------
Signal 2 : DAD1 B, Sig=254,4 Ref=off
 #  Meas. Ret.   Height    Width       Area       Area %
----------------------------------------------------------
 1      0.815     17.101   0.033         39.699    0.519
 2      0.967     23.487   0.026         42.230    0.552
 3      1.053   1259.189   0.023       1903.633   24.898
 4      1.094   2148.858   0.017       2478.554   32.418
 5      1.114   2063.032   0.022       2772.413   36.261
 6      1.301    153.644   0.039        409.161    5.352
----------------------------------------------------------
```

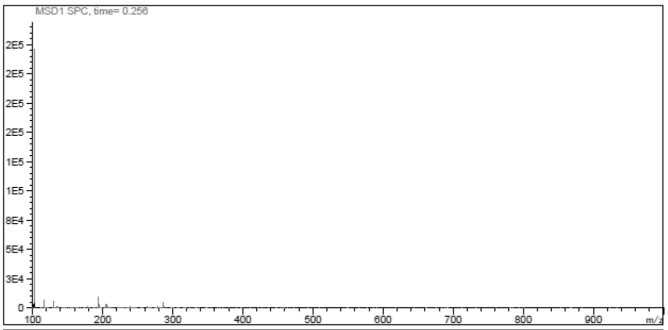
MSD1 SPC, time= 0.256
2E5
2E5
2E5
2E5
1E5
1E5
8E4
5E4
3E4
0
100 200 300 400 500 600 700 800 900 m/z

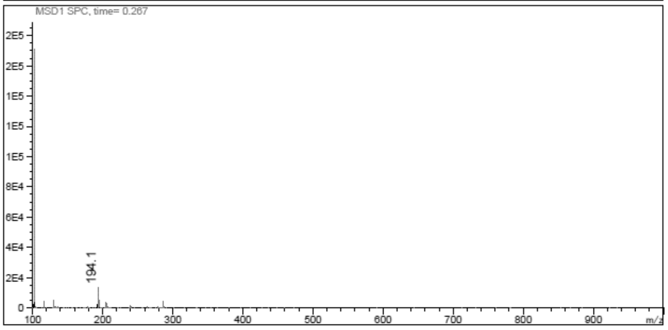
MSD1 SPC, time= 0.267
2E5
2E5
1E5
1E5
1E5
8E4
6E4
4E4
2E4
0
194.1
100 200 300 400 500 600 700 800 900 m/z

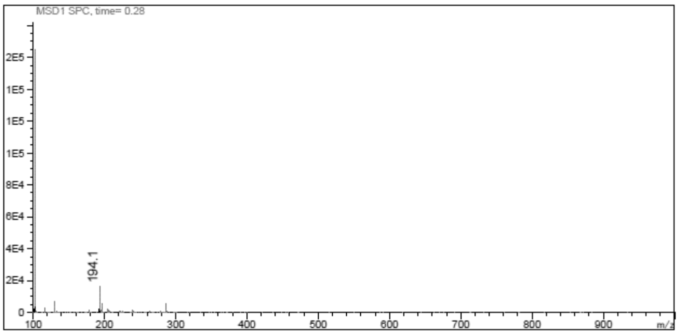
MSD1 SPC, time= 0.28
2E5
1E5
1E5
1E5
8E4
6E4
4E4
2E4
0
194.1
100 200 300 400 500 600 700 800 900 m/z

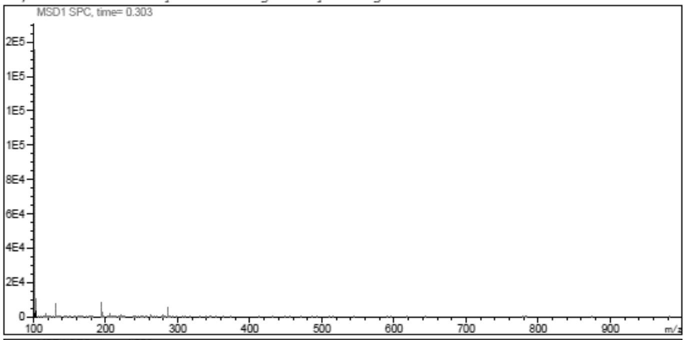
MSD1 SPC, time= 0.303
2E5
1E5
1E5
1E5
8E4
6E4
4E4
2E4
0
100 200 300 400 500 600 700 800 900 m/z

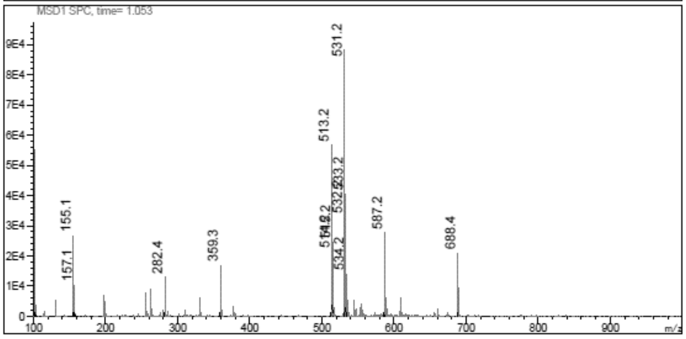
MSD1 SPC, time= 1.053
9E4
8E4
7E4
6E4
5E4
4E4
3E4
2E4
1E4
0
531.2
513.2
533.2
587.2
688.4
155.1
157.1
282.4
359.3
534.2
100 200 300 400 500 600 700 800 900 m/z

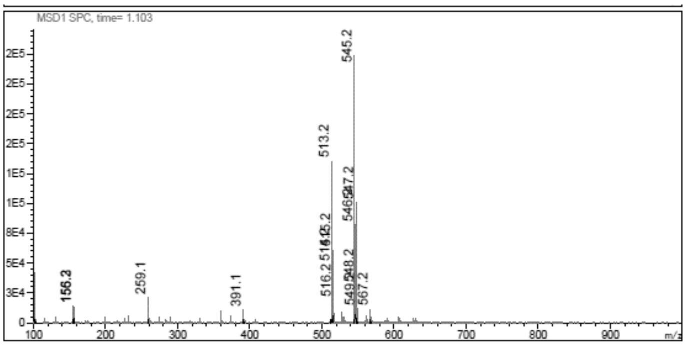

**$^{1}$H NMR** [(2*R*,3*R*,4*R*,5*R*)-2-[(*E*)-2-(1,3-Benzothiazol-2-yl)vinyl]-4-[tert-butyl (dimethyl)silyl]oxy-5-(6-chloropurin-9-yl)tetrahydrofuran-3-yl]oxy-tert-butyl-dimethyl-silane **(48)**

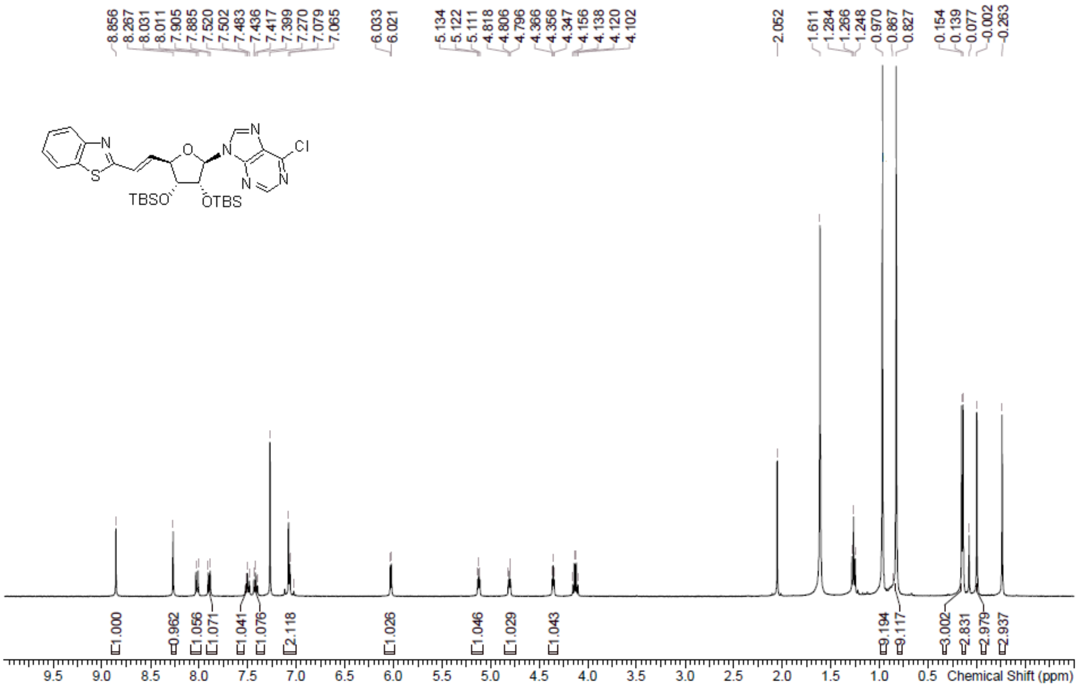

**LCMS** [(2*R*,3*R*,4*R*,5*R*)-2-[(*E*)-2-(1,3-Benzothiazol-2-yl)vinyl]-4-[tert-butyl (dimethyl)silyl]oxy-5-(6-chloropurin-9-yl)tetrahydrofuran-3-yl]oxy-tert-butyl-dimethyl-silane **(48)**

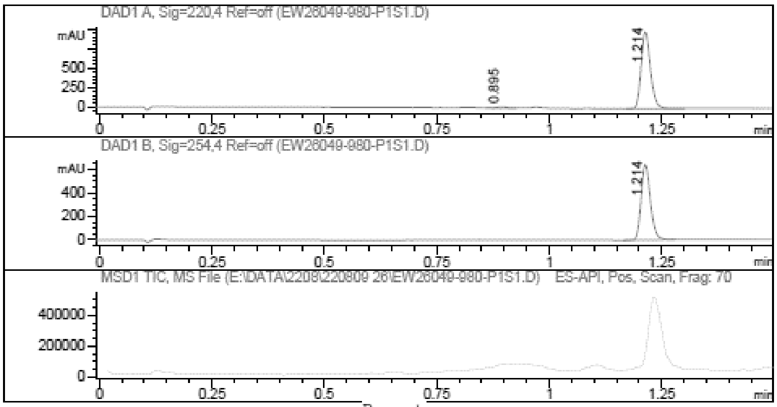

```
Report
=================================================================
   Signal 1 : DAD1 A, Sig=220,4 Ref=off
    #  Meas. Ret.   Height    Width       Area       Area %
   ----------------------------------------------------------
     1      0.895     15.259   0.028         29.316   2.000
     2      1.214    987.105   0.022       1436.830  98.000
   ----------------------------------------------------------

   Signal 2 : DAD1 B, Sig=254,4 Ref=off
    #  Meas. Ret.   Height    Width       Area       Area %
   ----------------------------------------------------------
     1      1.214    641.980   0.022        923.254 100.000
   ----------------------------------------------------------
```

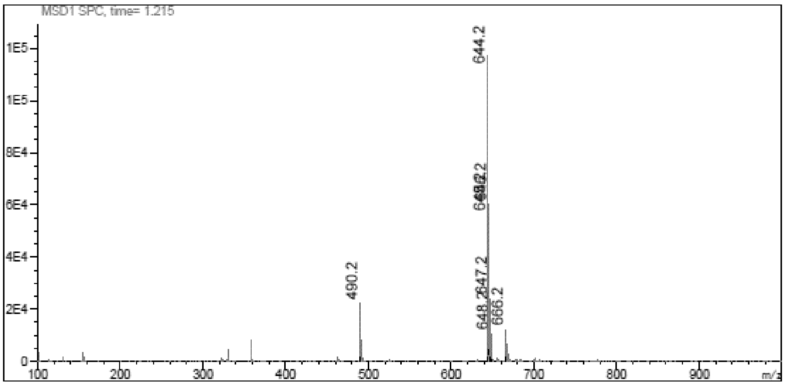

**LCMS** 9-[(2*R*,3*R*,4*R*,5*R*)-5-[(*E*)-2-(1,3-Benzothiazol-2-yl)vinyl]-3,4-bis [[tert-butyl(dimethyl)silyl]oxy]tetrahydrofuran-2-yl]purin-6-amine **(49)**

Chromatogram

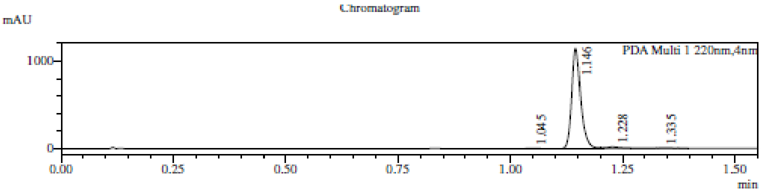

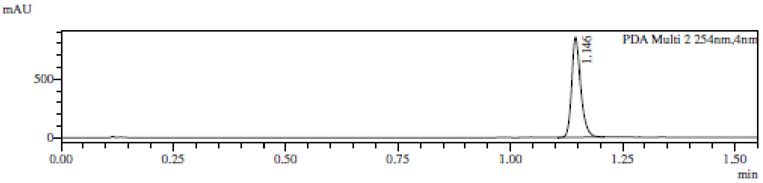

1 PDA Multi 1 / 220nm,4nm
2 PDA Multi 2 / 254nm,4nm

MS Chromatogram

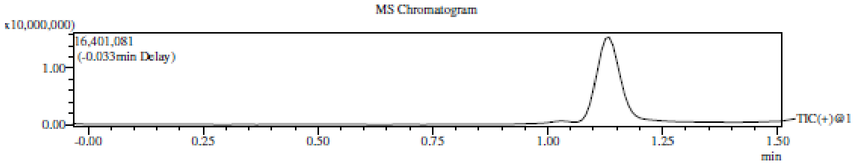

Integration Result

Peak Table

PDA Ch1 220nm

| Peak# | Ret. Time | Height | Height% | USP Width | Area | Area% |
|---|---|---|---|---|---|---|
| 1 | 1.045 | 5825 | 0.497 | 0.038 | 9068 | 0.560 |
| 2 | 1.146 | 1146419 | 97.862 | 0.036 | 1558756 | 96.202 |
| 3 | 1.228 | 16314 | 1.393 | 0.060 | 36825 | 2.273 |
| 4 | 1.335 | 2909 | 0.248 | 0.553 | 15652 | 0.966 |

Peak Table

PDA Ch2 254nm

| Peak# | Ret. Time | Height | Height% | USP Width | Area | Area% |
|---|---|---|---|---|---|---|
| 1 | 1.146 | 851413 | 100.000 | 0.036 | 1160600 | 100.000 |

Mass Spectrum

RetTime: 1.042 Datafile: D:\DATA\2022\2208\220818\EW26049-988-P1C2.lcd

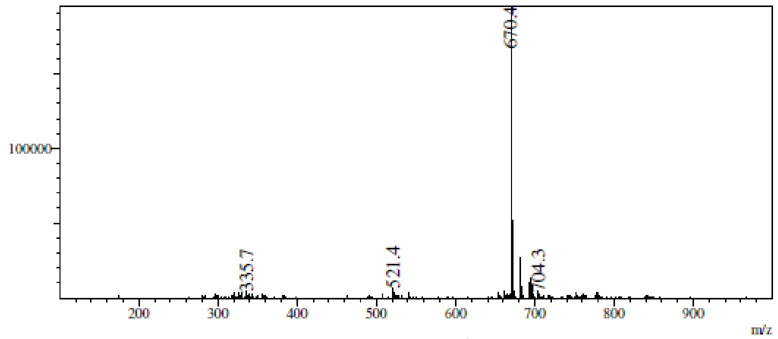

RetTime: 1.142 Datafile: D:\DATA\2022\2208\220818\EW26049-988-P1C2.lcd

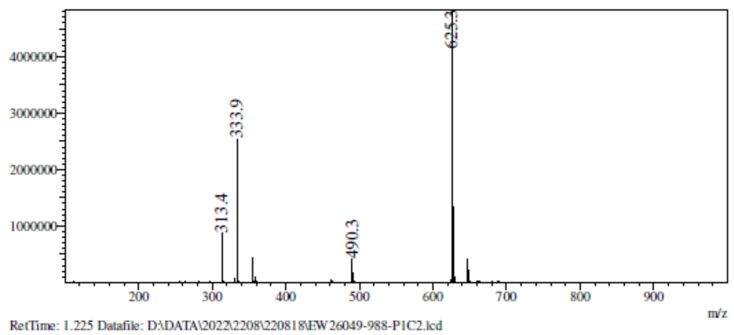

RetTime: 1.225 Datafile: D:\DATA\2022\2208\220818\EW26049-988-P1C2.lcd

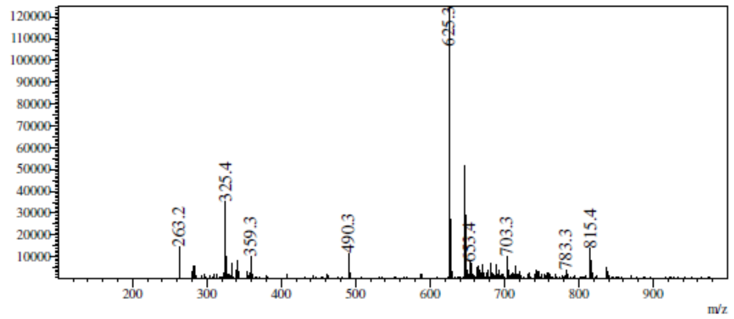

RetTime: 1.333 Datafile: D:\DATA\2022\2208\220818\EW26049-988-P1C2.lcd

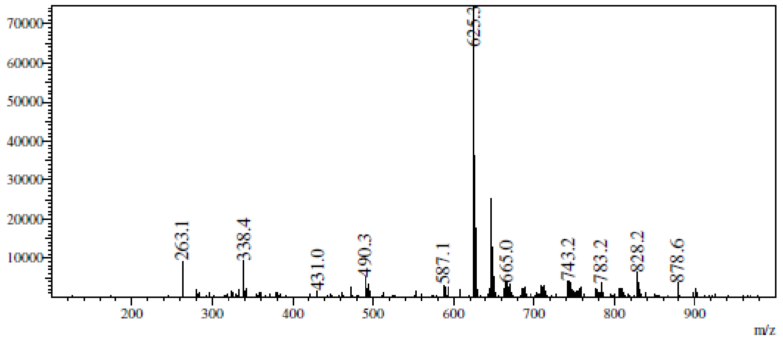

**$^1$H NMR** (2*R*,3*R*,4*S*,5*R*)-2-(6-Aminopurin-9-yl)-5-[(*E*)-2-(1,3-benzothiazol-2-yl) vinyl]tetrahydrofuran-3,4-diol **(33)**

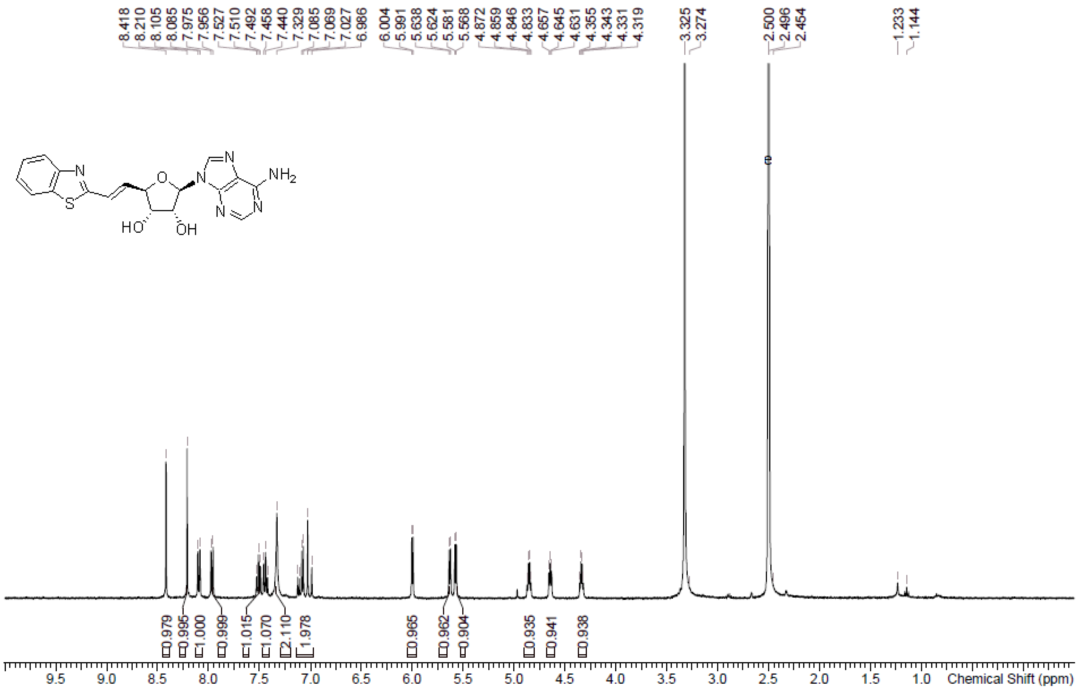

**HPLC** (2*R*,3*R*,4*S*,5*R*)-2-(6-Aminopurin-9-yl)-5-[(*E*)-2-(1,3-benzothiazol-2-yl) vinyl]tetrahydrofuran-3,4-diol **(33)**

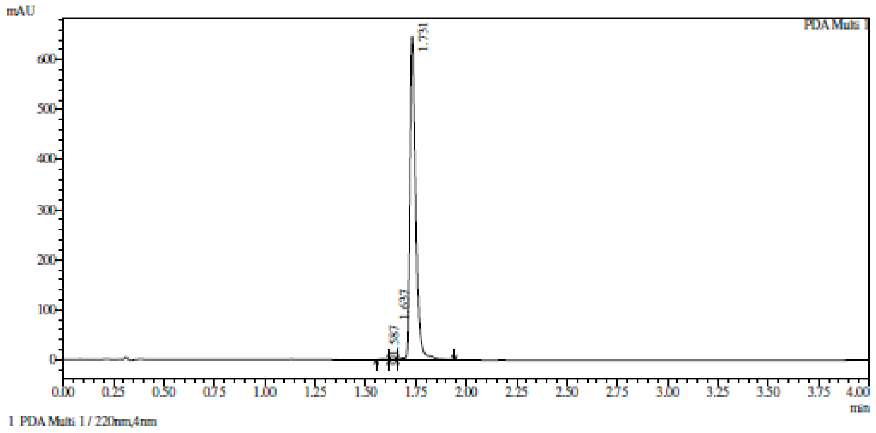

Integration result

PeakTable

PDA Ch1 220nm

| Peak# | Ret. Time | USP Width | Resolution | Height | Area | Area % |
|---|---|---|---|---|---|---|
| 1 | 1.587 | 0.053 | 0.000 | 1563 | 3091 | 0.237 |
| 2 | 1.637 | 0.100 | 0.662 | 1055 | 1736 | 0.133 |
| 3 | 1.731 | 0.051 | 1.238 | 647135 | 1300375 | 99.630 |
| Total | | | | 649752 | 1305202 | 100.000 |

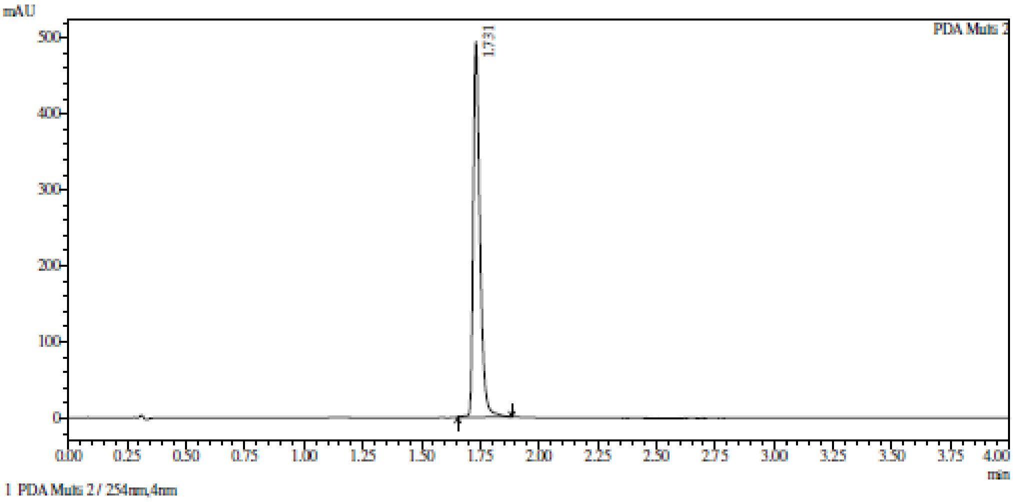

Integration result

PeakTable

PDA Ch2 254nm

| Peak# | Ret. Time | USP Width | Resolution | Height | Area | Area % |
|---|---|---|---|---|---|---|
| 1 | 1.731 | 0.051 | 0.000 | 494603 | 984589 | 100.000 |
| Total | | | | 494603 | 984589 | 100.000 |

**HRMC** (2*R*,3*R*,4*S*,5*R*)-2-(6-Aminopurin-9-yl)-5-[(*E*)-2-(1,3-benzothiazol-2-yl) vinyl]tetrahydrofuran-3,4-diol **(33)**

Compound Table

| Label | Tgt Score | Mass Error (ppm) | Tgt Formula | Obs. RT | Ref. Mass | Obs. Mass |
|---|---|---|---|---|---|---|
| Cpd 1: C18 H16 N6 O3 S; 1.768 | 99.29 | 3.28 | C18 H16 N6 O3 S | 1.768 | 396.1005 | 396.1018 |

| Obs. *m/z* | Obs. RT | Obs. Mass | Tgt Formula | Tgt Mass | Tgt Mass Error (ppm) | Find Cpds Algorithm |
|---|---|---|---|---|---|---|
| 419.0899 | 1.768 | 396.1018 | C18 H16 N6 O3 S | 396.1005 | 3.28 | Find by Formula |

Compound Chromatograms

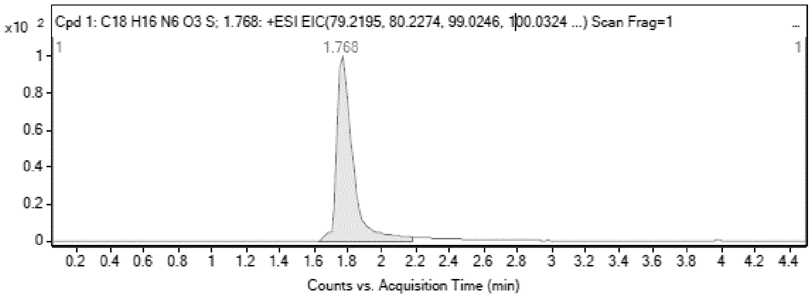

MS Zoomed Spectrum

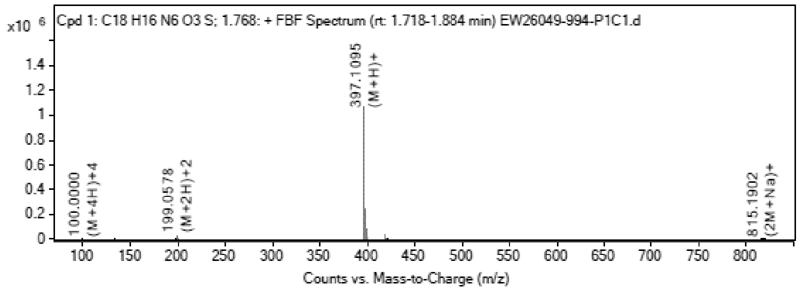

MS Spectrum Peak List

| Obs. *m/z* | Charge | Abund | Ion/Isotope |
|---|---|---|---|
| 199.0578 | 2 | 29187.99 | (M+2H)+2 |
| 199.559 | 2 | 7087.75 | (M+2H)+2 |

| | | | |
|---|---|---|---|
| 397.1095 | 1 | 1066689 | (M+H)+ |
| 398.1113 | 1 | 248296.41 | (M+H)+ |
| 399.108 | 1 | 76921.66 | (M+H)+ |
| 419.0899 | 1 | 38582.68 | (M+Na)+ |
| 420.0924 | 1 | 8908.7 | (M+Na)+ |
| 815.1902 | 1 | 8208.4 | (2M+Na)+ |
| 816.1932 | 1 | 3695.78 | (2M+Na)+ |
| 817.1763 | 1 | 4415.6 | (2M+Na)+ |

MS Zoomed Spectrum

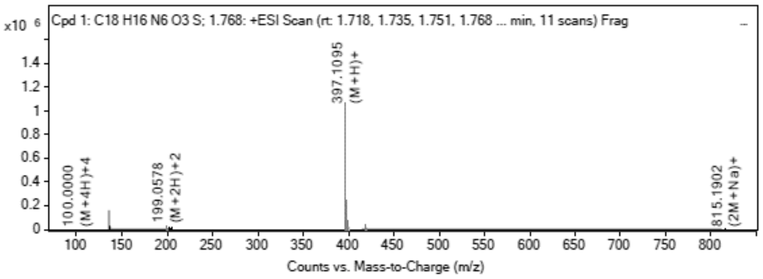

**MS Spectrum Peak List**

| Obs. *m/z* | Charge | Abund | Ion/Isotope | Tgt Mass Error (ppm) |
|---|---|---|---|---|
| 199.0578 | 2 | 29187.99 | (M+2H)+2 | -1.63 |
| 199.559 | 2 | 7087.75 | (M+2H)+2 | -0.81 |
| 397.1095 | 1 | 1066689 | (M+H)+ | -4.46 |
| 398.1113 | 1 | 248296.41 | (M+H)+ | -2.35 |
| 399.108 | 1 | 76921.66 | (M+H)+ | -1.72 |
| 419.0899 | 1 | 38582.68 | (M+Na)+ | -0.45 |
| 420.0924 | 1 | 8908.7 | (M+Na)+ | -0.26 |
| 815.1902 | 1 | 8208.4 | (2M+Na)+ | -0.06 |
| 816.1932 | 1 | 3695.78 | (2M+Na)+ | -0.5 |
| 817.1763 | 1 | 4415.6 | (2M+Na)+ | 18.15 |

--- [illegible] ---

**$^1$H NMR** 9-[(2*R*,3*R*,4*R*,5*R*)-5-[2-(1,3-Benzothiazol-2-yl)ethyl]-3,4-bis [[tert-butyl(dimethyl)silyl]oxy] tetrahydrofuran-2-yl]purin-6-amine **(50)**

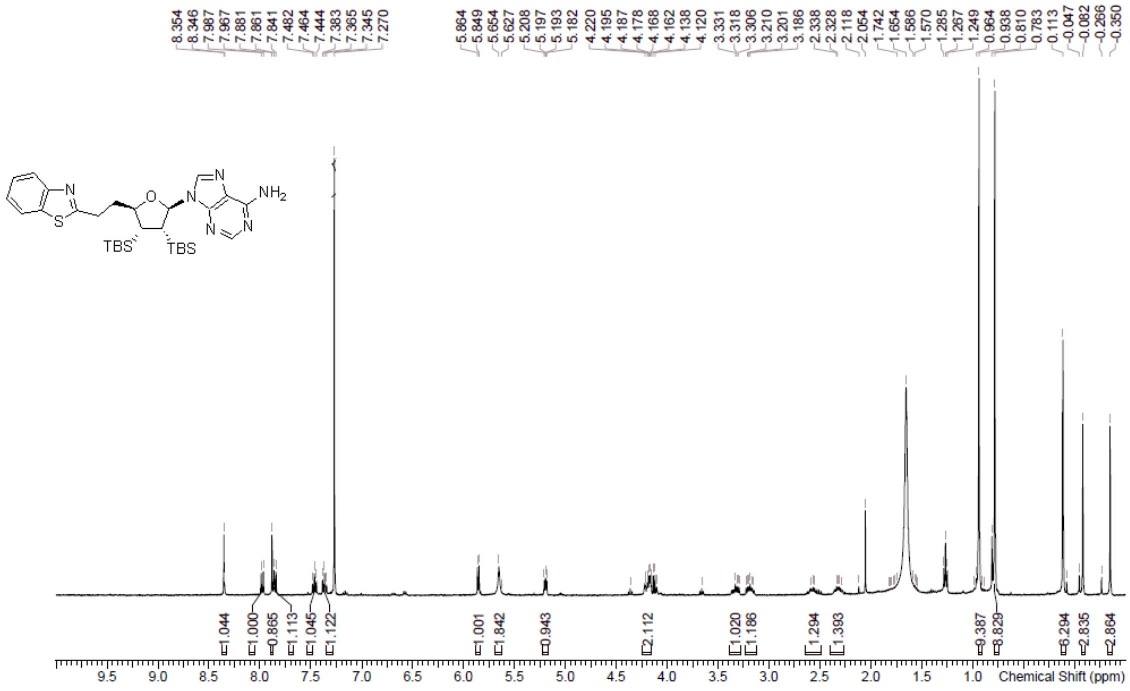

**LCMS** 9-[(2*R*,3*R*,4*R*,5*R*)-5-[2-(1,3-Benzothiazol-2-yl)ethyl]-3,4-bis [[tert-butyl(dimethyl)silyl]oxy]tetrahydrofuran-2-yl]purin-6-amine **(50)**

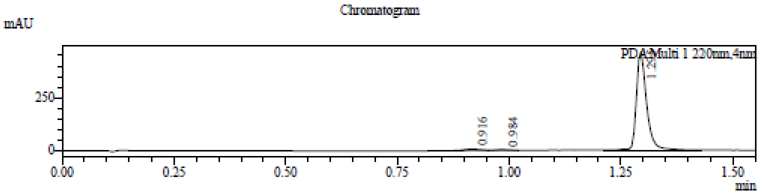

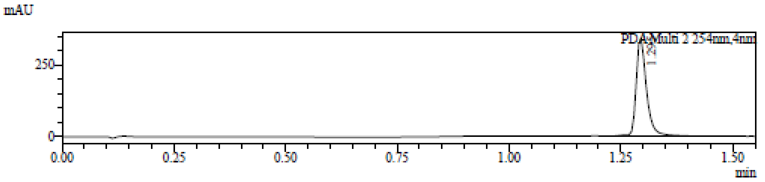

1 PDA Multi 1 / 220nm,4nm
2 PDA Multi 2 / 254nm,4nm

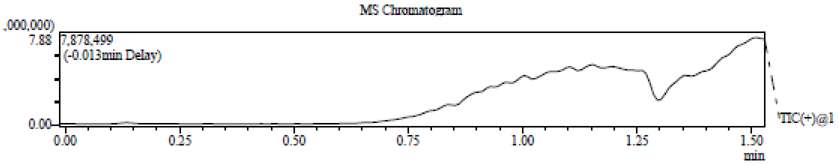

Integration Result

Peak Table

PDA Ch1 220nm

| Peak# | Ret. Time | Height | Height% | USP Width | Area | Area% |
|---|---|---|---|---|---|---|
| 1 | 0.916 | 6607 | 1.397 | 0.081 | 18435 | 2.442 |
| 2 | 0.984 | 3107 | 0.657 | 0.055 | 5752 | 0.762 |
| 3 | 1.295 | 463329 | 97.947 | 0.046 | 730841 | 96.797 |

Peak Table

PDA Ch2 254nm

| Peak# | Ret. Time | Height | Height% | USP Width | Area | Area% |
|---|---|---|---|---|---|---|
| 1 | 1.295 | 342660 | 100.000 | 0.046 | 547623 | 100.000 |

Mass Spectrum

RetTime: 0.920 Datafile: D:\DATA\2022\2208\220829\EW26049-1008-P1C1.lcd

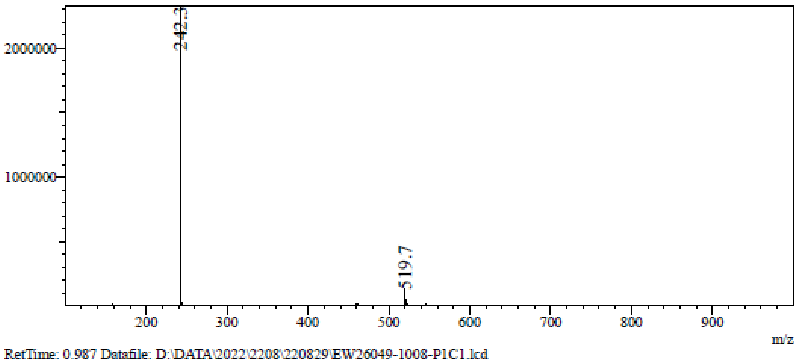

RetTime: 0.987 Datafile: D:\DATA\2022\2208\220829\EW26049-1008-P1C1.lcd

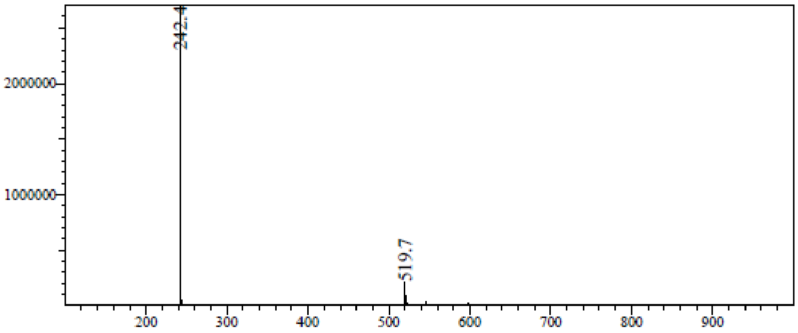

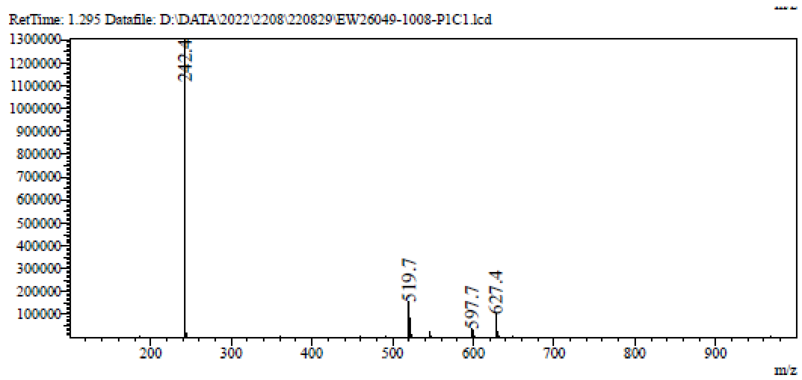

**$^{1}$H NMR** (2*R*,3*R*,4*S*,5*R*)-2-(6-Aminopurin-9-yl)-5-[2-(1,3-benzothiazol-2-yl) ethyl]tetrahydrofuran-3,4-diol **(25)**

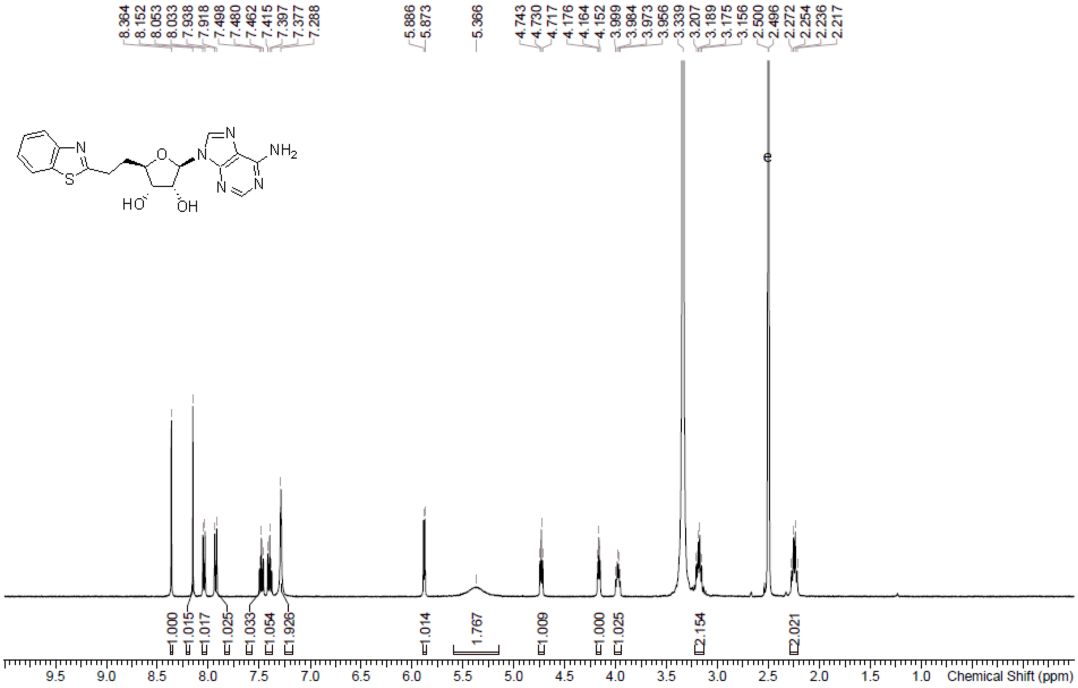

**HPLC** (2*R*,3*R*,4*S*,5*R*)-2-(6-Aminopurin-9-yl)-5-[2-(1,3-benzothiazol-2-yl) ethyl]tetrahydrofuran-3,4-diol **(25)**

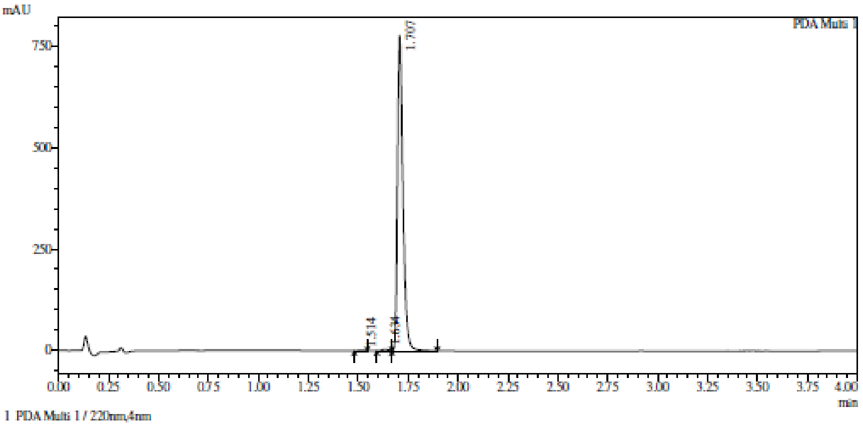

1 PDA Multi 1 / 220nm,4nm

Integration result

PeakTable

PDA Ch1 220nm

| Peak# | Ret. Time | USP Width | Resolution | Height | Area | Area % |
|---|---|---|---|---|---|---|
| 1 | 1.514 | 0.103 | 0.000 | 1264 | 3218 | 0.211 |
| 2 | 1.634 | 0.077 | 1.345 | 3528 | 9421 | 0.618 |
| 3 | 1.707 | 0.051 | 1.137 | 778638 | 1511865 | 99.171 |
| Total | | | | 783431 | 1524505 | 100.000 |

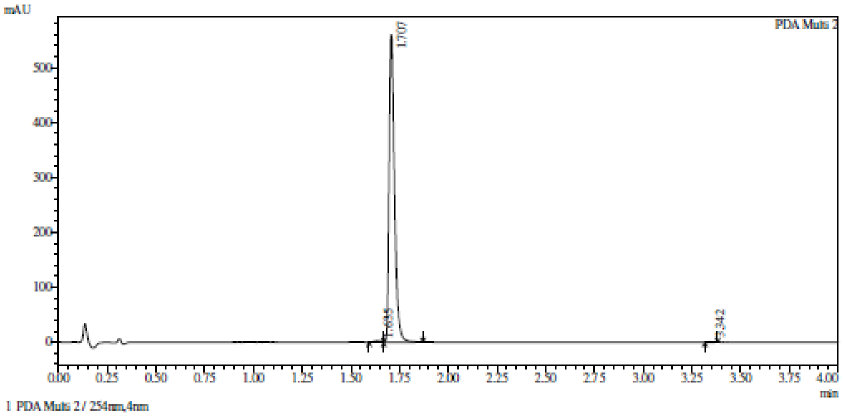

1 PDA Multi 2 / 254nm,4nm

Integration result

PeakTable

PDA Ch2 254nm

| Peak# | Ret. Time | USP Width | Resolution | Height | Area | Area % |
|---|---|---|---|---|---|---|
| 1 | 1.635 | 0.077 | 0.000 | 2550 | 6858 | 0.629 |
| 2 | 1.707 | 0.051 | 1.122 | 559197 | 1081077 | 99.229 |
| 3 | 3.342 | 0.052 | 31.750 | 822 | 1547 | 0.142 |
| Total | | | | 562570 | 1089482 | 100.000 |

**HRMC** (2*R*,3*R*,4*S*,5*R*)-2-(6-Aminopurin-9-yl)-5-[2-(1,3-benzothiazol-2-yl) ethyl]tetrahydrofuran-3,4-diol **(25)**

**Compound Table**

| Label | Tgt Score | Mass Error (ppm) | Tgt Formula | Obs. RT | Ref. Mass | Obs. Mass |
|---|---|---|---|---|---|---|
| Cpd 1: C18 H18 N6 O3 S; 1.666 | 99.11 | 3.22 | C18 H18 N6 O3 S | 1.666 | 398.1161 | 398.1174 |

| Obs. *m/z* | Obs. RT | Obs. Mass | Tgt Formula | Tgt Mass | Tgt Mass Error (ppm) | Find Cpds Algorithm |
|---|---|---|---|---|---|---|
| 421.1054 | 1.666 | 398.1174 | C18 H18 N6 O3 S | 398.1161 | 3.22 | Find by Formula |

**Compound Chromatograms**

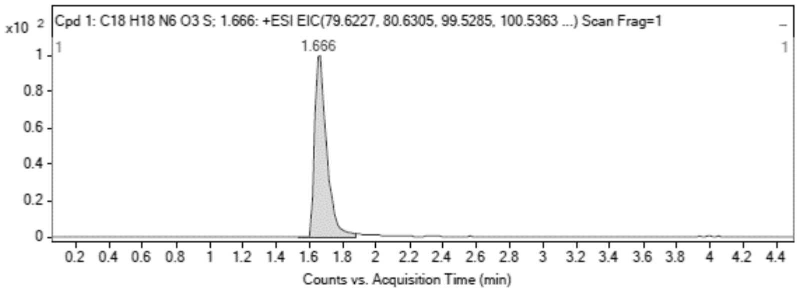

**MS Zoomed Spectrum**

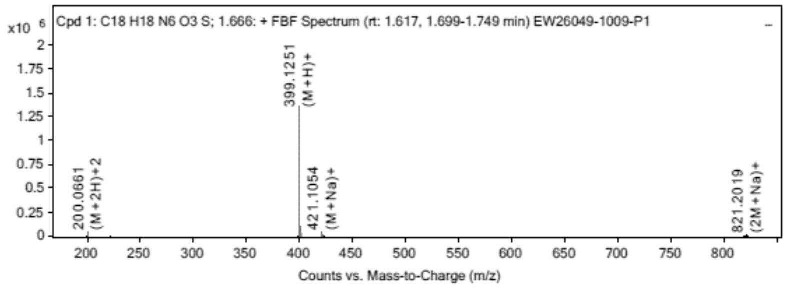

**MS Spectrum Peak List**

| Obs. *m/z* | Charge | Abund | Ion/Isotope |
|---|---|---|---|
| 200.0661 | 2 | 38145.5 | (M+2H)+2 |
| 200.5672 | 2 | 8738.28 | (M+2H)+2 |
| 201.0661 | 2 | 3324.03 | (M+2H)+2 |
| 399.1251 | 1 | 1358144.75 | (M+H)+ |
| 400.1268 | 1 | 316680.78 | (M+H)+ |
| 401.1235 | 1 | 98099.39 | (M+H)+ |
| 421.1054 | 1 | 43094.38 | (M+Na)+ |
| 422.1076 | 1 | 9586.86 | (M+Na)+ |
| 819.2206 | 1 | 5366.74 | (2M+Na)+ |
| 821.2019 | 1 | 6874.68 | (2M+Na)+ |

MS Zoomed Spectrum

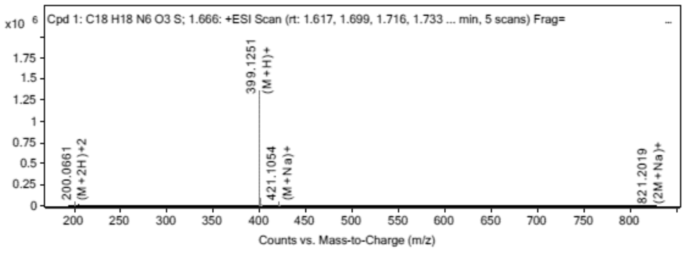

**MS Spectrum Peak List**

| Obs. *m/z* | Charge | Abund | Ion/Isotope | Tgt Mass Error (ppm) |
|---|---|---|---|---|
| 200.0661 | 2 | 38145.5 | (M+2H)+2 | -3.6 |
| 200.5672 | 2 | 8738.28 | (M+2H)+2 | -2.95 |
| 201.0661 | 2 | 3324.03 | (M+2H)+2 | -5.06 |
| 399.1251 | 1 | 1358144.75 | (M+H)+ | -4.29 |
| 400.1268 | 1 | 316680.78 | (M+H)+ | -1.97 |
| 401.1235 | 1 | 98099.39 | (M+H)+ | -1.39 |
| 421.1054 | 1 | 43094.38 | (M+Na)+ | -0.12 |
| 422.1076 | 1 | 9586.86 | (M+Na)+ | 0.96 |
| 819.2206 | 1 | 5366.74 | (2M+Na)+ | 1.06 |
| 821.2019 | 1 | 6874.68 | (2M+Na)+ | 25 |

--- End Of Report ---

**[1]H NMR** (1*R*,4*S*,5*R*,6*S*)-5,6-Dihydroxy-2-azabicyclo[2.2.1]heptan-3-one **(52)**

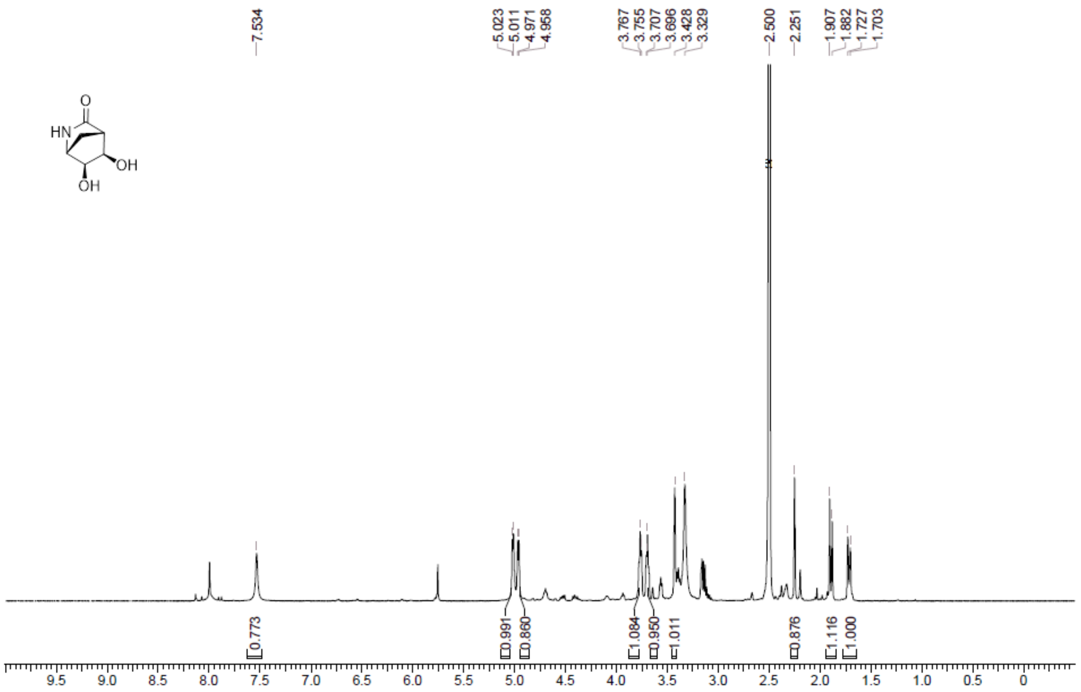

**LCMS** (1*R*,4*S*,5*R*,6*S*)-5,6-Dihydroxy-2-azabicyclo[2.2.1]heptan-3-one **(52)**

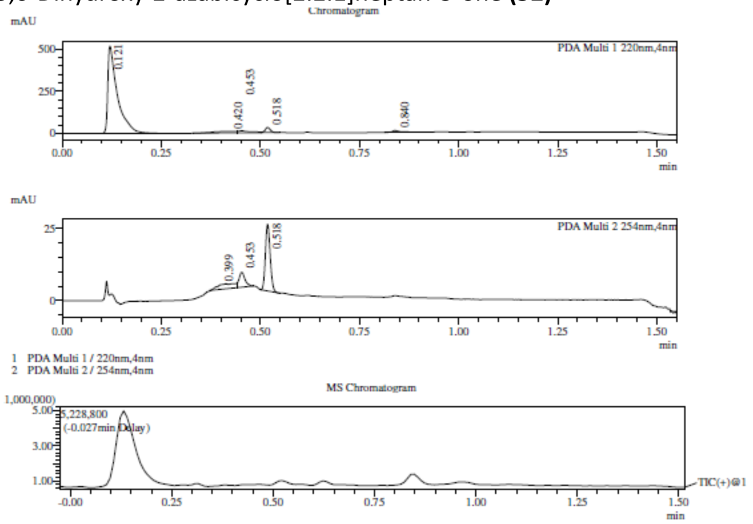

Integration Result

Peak Table

PDA Ch1 220nm

| Peak# | Ret. Time | Height | Height% | USP Width | Area | Area% |
|---|---|---|---|---|---|---|
| 1 | 0.121 | 516265 | 89.294 | 0.041 | 872473 | 89.391 |
| 2 | 0.420 | 9172 | 1.586 | 0.193 | 37956 | 3.889 |
| 3 | 0.453 | 11901 | 2.058 | 0.060 | 24618 | 2.522 |
| 4 | 0.518 | 30231 | 5.229 | 0.026 | 27963 | 2.865 |
| 5 | 0.840 | 10592 | 1.832 | 0.030 | 13011 | 1.333 |

Peak Table

PDA Ch2 254nm

| Peak# | Ret. Time | Height | Height% | USP Width | Area | Area% |
|---|---|---|---|---|---|---|
| 1 | 0.399 | 1629 | 5.432 | 0.241 | 5179 | 17.148 |
| 2 | 0.453 | 5198 | 17.334 | 0.031 | 5330 | 17.648 |
| 3 | 0.518 | 23161 | 77.234 | 0.025 | 19693 | 65.203 |

Mass Spectrum

RetTime: 0.123 Datafile: D:\DATA\2022\2201\220124\EW26049-724-P1C2.lcd

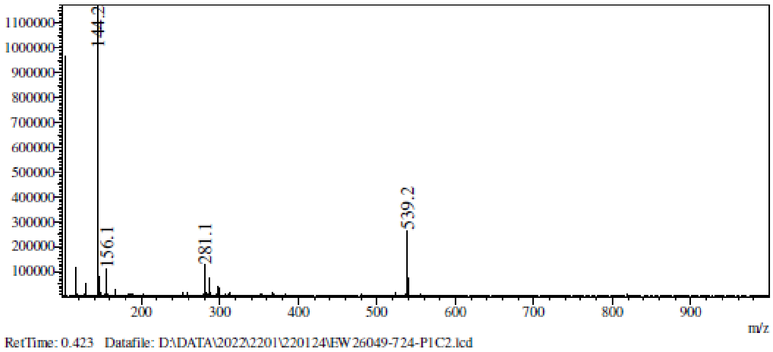

RetTime: 0.423 Datafile: D:\DATA\2022\2201\220124\EW26049-724-P1C2.lcd

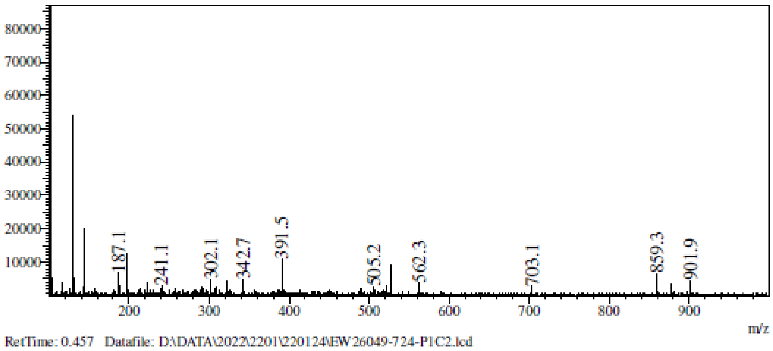

RetTime: 0.457 Datafile: D:\DATA\2022\2201\220124\EW26049-724-P1C2.lcd

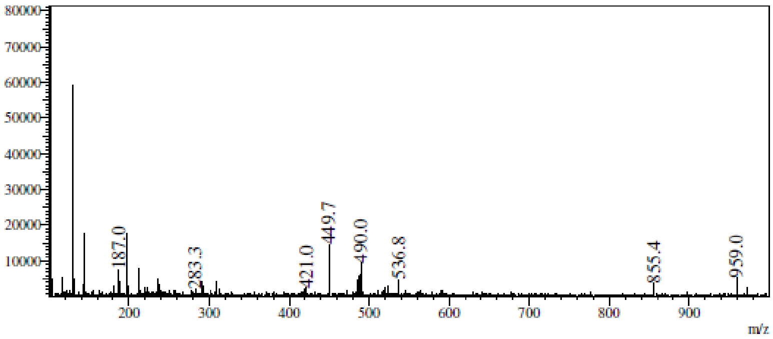

RetTime: 0.515 Datafile: D:\DATA\2022\2201\220124\EW26049-724-P1C2.lcd

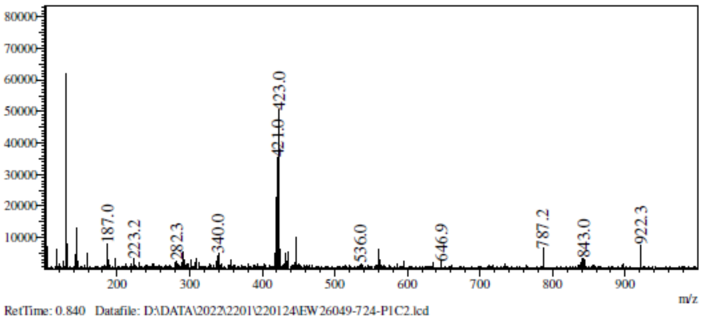

RetTime: 0.840 Datafile: D:\DATA\2022\2201\220124\EW26049-724-P1C2.lcd

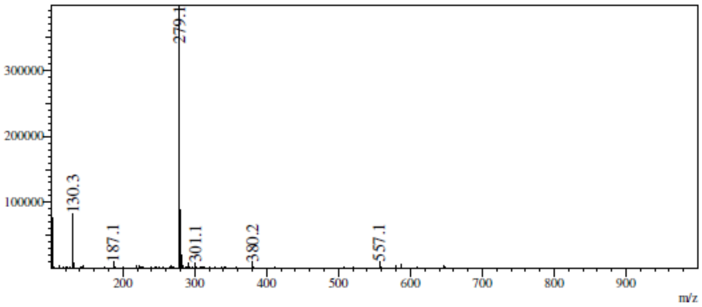

**$^{1}$H NMR** (1*R*,4*S*,5*R*,6*S*)-5,6-Bis[[tert-butyl(diphenyl)silyl]oxy]-2-azabicyclo [2.2.1]heptan-3-one **(53)**

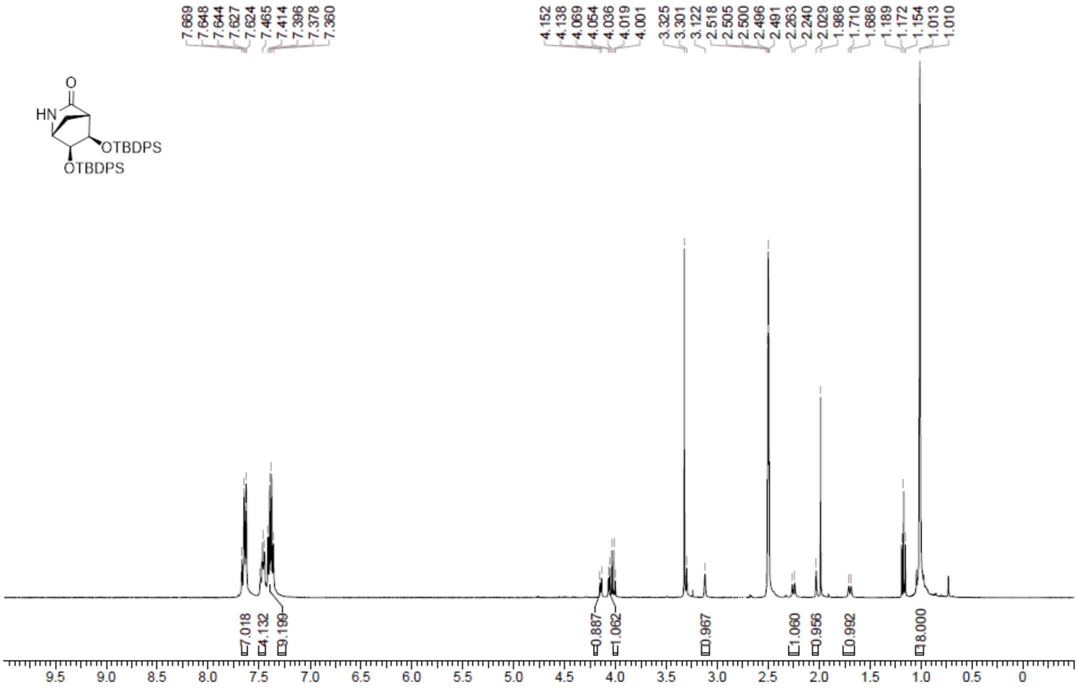

**LCMS** (1*R*,4*S*,5*R*,6*S*)-5,6-Bis[[tert-butyl(diphenyl)silyl]oxy]-2-azabicyclo [2.2.1]heptan-3-one **(53)**

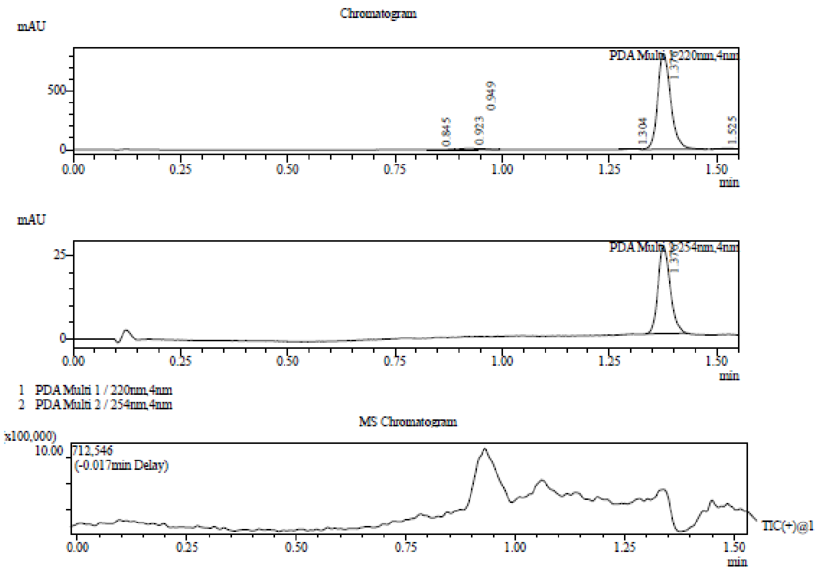

Integration Result

Peak Table

PDA Ch1 220nm

| Peak# | Ret. Time | Height | Height% | USP Width | Area | Area% |
|---|---|---|---|---|---|---|
| 1 | 0.845 | 1696 | 0.201 | 0.056 | 6157 | 0.346 |
| 2 | 0.923 | 10582 | 1.254 | 0.084 | 24960 | 1.403 |
| 3 | 0.949 | 7345 | 0.870 | 0.113 | 9285 | 0.522 |
| 4 | 1.304 | 5284 | 0.626 | 0.058 | 9768 | 0.549 |
| 5 | 1.376 | 814867 | 96.569 | 0.055 | 1720930 | 96.738 |
| 6 | 1.525 | 4041 | 0.479 | 0.051 | 7855 | 0.442 |

Peak Table

PDA Ch2 254nm

| Peak# | Ret. Time | Height | Height% | USP Width | Area | Area% |
|---|---|---|---|---|---|---|
| 1 | 1.376 | 26015 | 100.000 | 0.052 | 51285 | 100.000 |

Mass Spectrum

RetTime: 0.847 Datafile: D:\DATA\2022\2201\220128\EW26049-730-P1C1.lcd

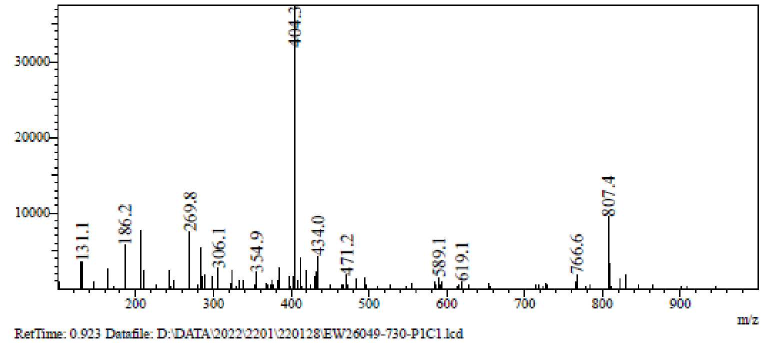

RetTime: 0.923 Datafile: D:\DATA\2022\2201\220128\EW26049-730-P1C1.lcd

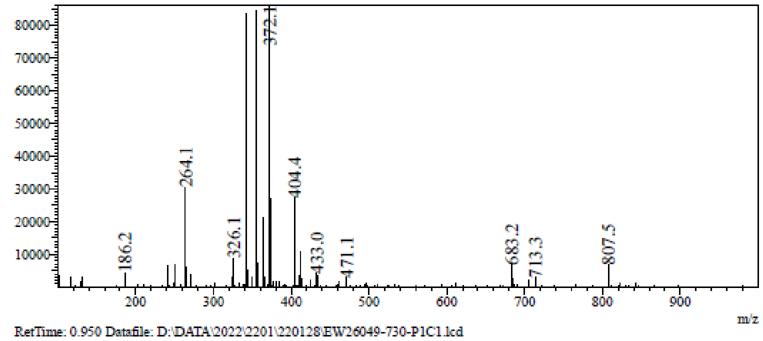

RetTime: 0.950 Datafile: D:\DATA\2022\2201\220128\EW26049-730-P1C1.lcd

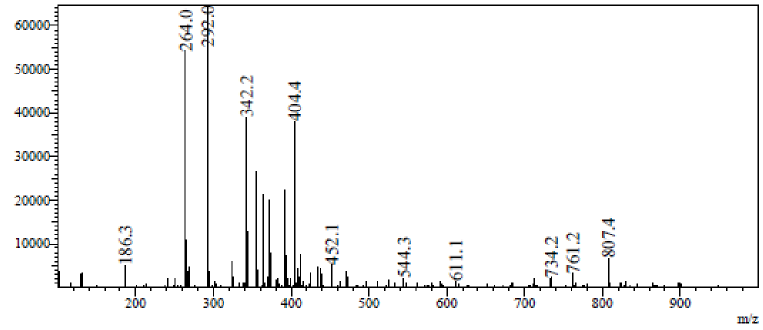

RetTime: 1.303 Datafile: D:\DATA\2022\2201\220128\EW26049-730-P1C1.lcd

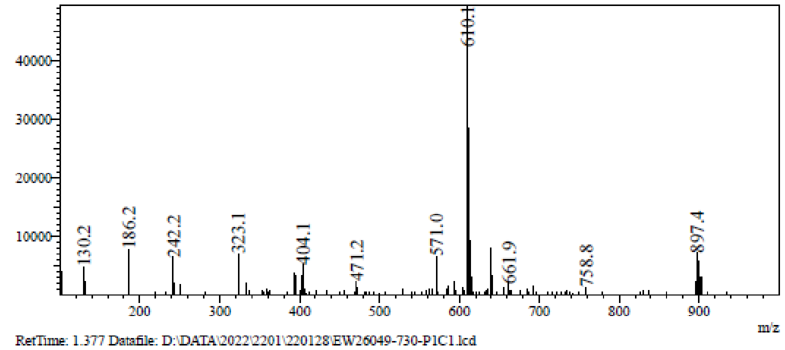

RetTime: 1.377 Datafile: D:\DATA\2022\2201\220128\EW26049-730-P1C1.lcd

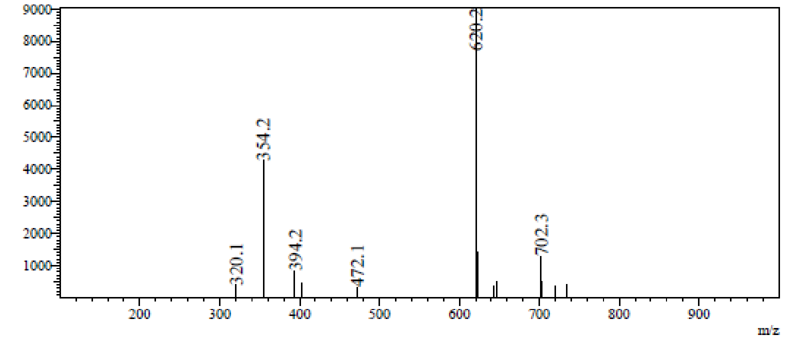

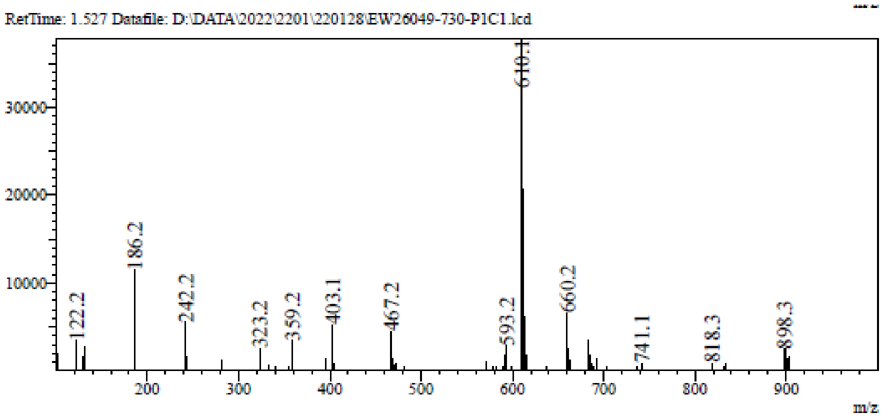

**[1]H NMR** Benzyl (1*R*,4*S*,5*R*,6*S*)-5,6-bis[[tert-butyl(diphenyl)silyl]oxy]-3-oxo-2-azabicyclo[2.2.1] heptane-2-carboxylate **(54)**

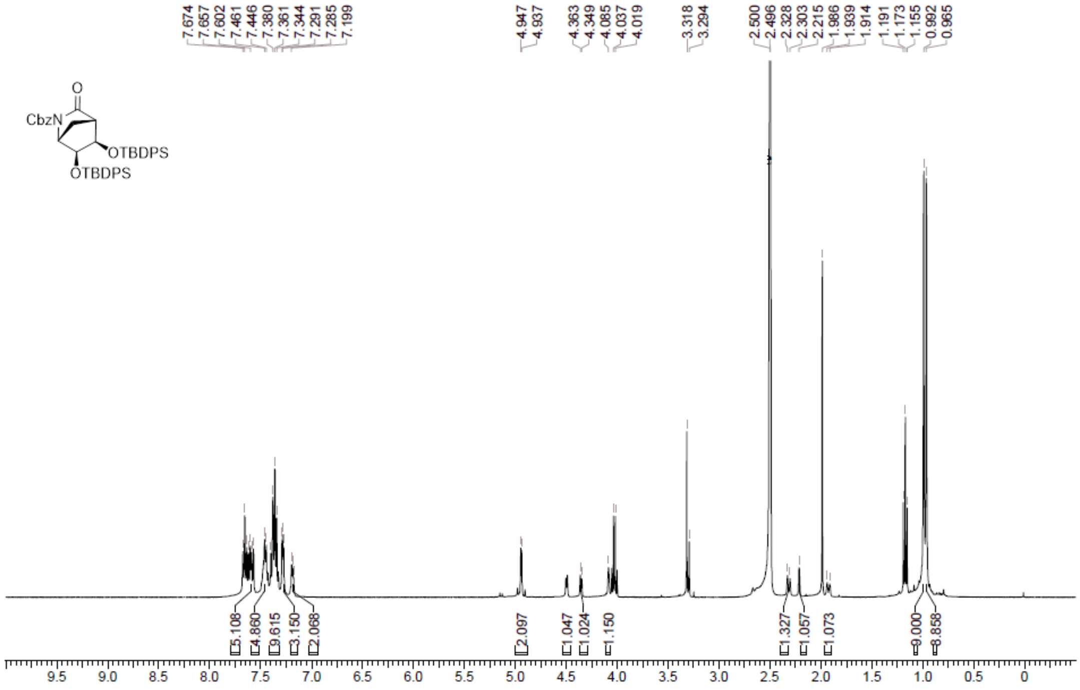

**LCMS** Benzyl (1*R*,4*S*,5*R*,6*S*)-5,6-bis[[tert-butyl(diphenyl)silyl]oxy]-3-oxo-2-azabicyclo[2.2.1] heptane-2-carboxylate **(54)**

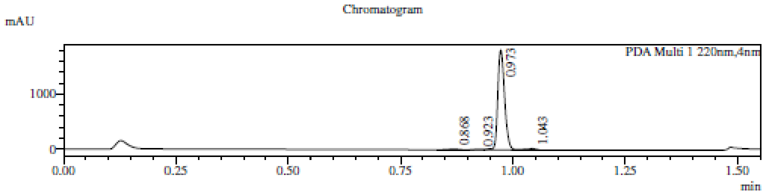

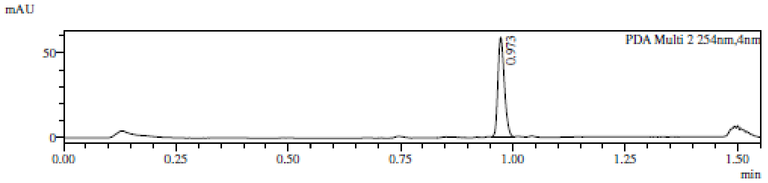

1 PDA Multi 1 / 220nm,4nm
2 PDA Multi 2 / 254nm,4nm

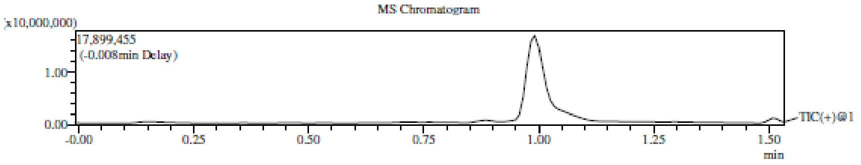

Integration Result

Peak Table

PDA Ch1 220nm

| Peak# | Ret. Time | Height | Height% | USP Width | Area | Area% |
|---|---|---|---|---|---|---|
| 1 | 0.868 | 14991 | 0.798 | 0.041 | 22814 | 1.121 |
| 2 | 0.923 | 9179 | 0.489 | 0.039 | 9826 | 0.483 |
| 3 | 0.973 | 1828083 | 97.370 | 0.030 | 1979253 | 97.248 |
| 4 | 1.043 | 25209 | 1.343 | 0.027 | 23377 | 1.149 |

Peak Table

PDA Ch2 254nm

| Peak# | Ret. Time | Height | Height% | USP Width | Area | Area% |
|---|---|---|---|---|---|---|
| 1 | 0.973 | 58471 | 100.000 | 0.029 | 59369 | 100.000 |

Mass Spectrum

RetTime: 0.867 Datafile: D:\DATA\2022\2202\220209\EW26049-732-P1C2.lcd

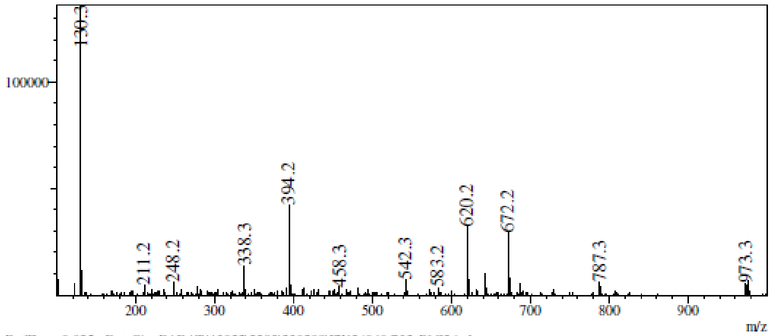

RetTime: 0.925 Datafile: D:\DATA\2022\2202\220209\EW26049-732-P1C2.lcd

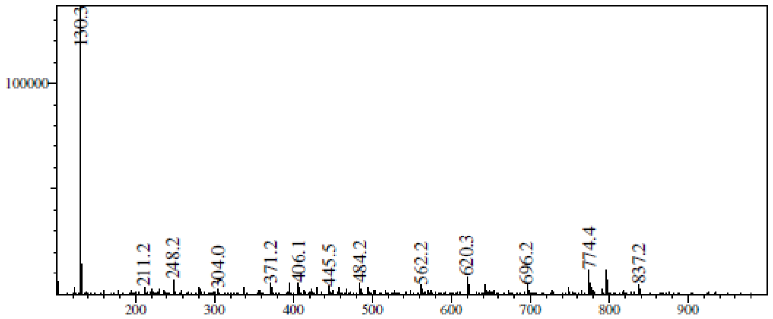

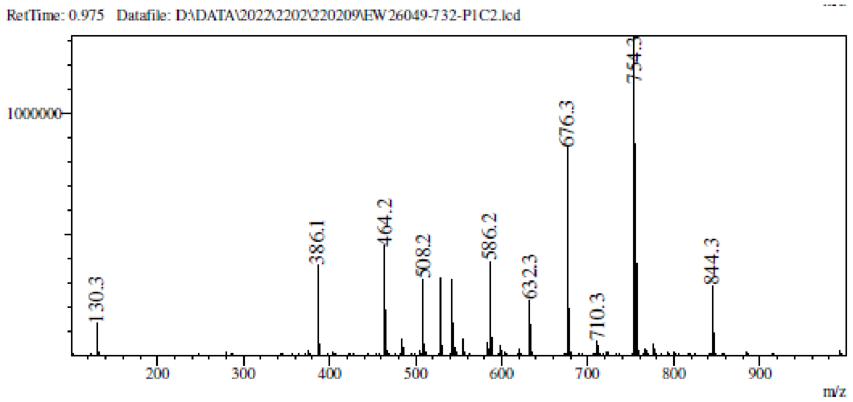

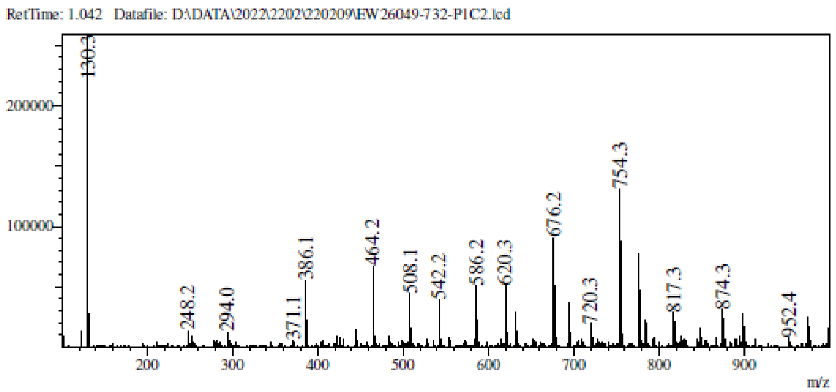

**[1]H NMR** Benzyl *N*-[(1*R*,2*S*,3*R*,4*R*)-2,3-bis[[tert-butyl(diphenyl)silyl]oxy]-4 (hydroxymethyl) cyclopentyl]carbamate **(55)**

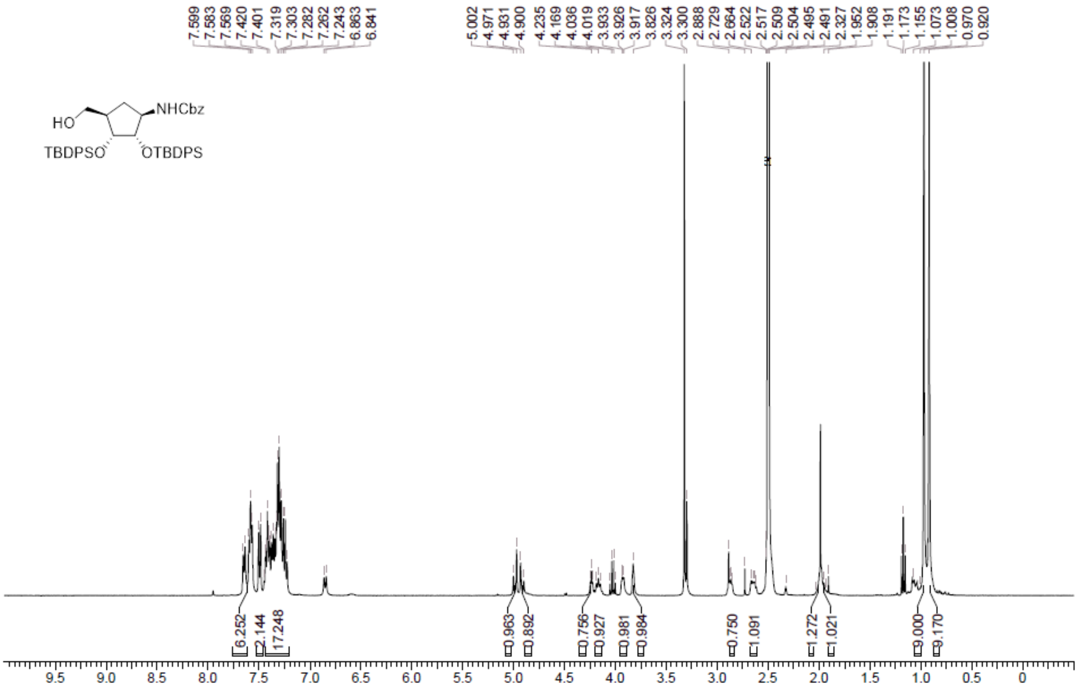

**LCMS** Benzyl *N*-[(1*R*,2*S*,3*R*,4*R*)-2,3-bis[[tert-butyl(diphenyl)silyl]oxy]-4 (hydroxymethyl) cyclopentyl]carbamate **(55)**

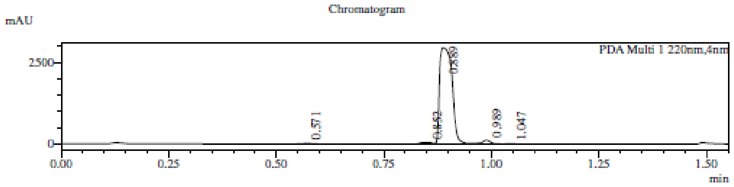

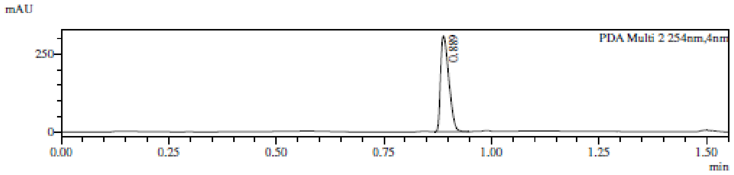

1 PDA Multi 1 / 220nm,4nm
2 PDA Multi 2 / 254nm,4nm

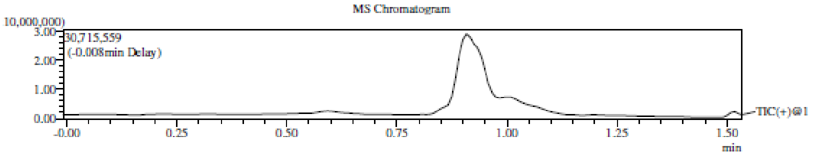

Integration Result

Peak Table

PDA Ch1 220nm

| Peak# | Ret. Time | Height | Height% | USP Width | Area | Area% |
|---|---|---|---|---|---|---|
| 1 | 0.571 | 18871 | 0.596 | 0.046 | 35343 | 0.557 |
| 2 | 0.852 | 50866 | 1.608 | 0.051 | 98446 | 1.550 |
| 3 | 0.889 | 2955632 | 93.424 | 0.047 | 6004051 | 94.549 |
| 4 | 0.989 | 124214 | 3.926 | 0.034 | 188584 | 2.970 |
| 5 | 1.047 | 14108 | 0.446 | 0.052 | 23776 | 0.374 |

Peak Table

PDA Ch2 254nm

| Peak# | Ret. Time | Height | Height% | USP Width | Area | Area% |
|---|---|---|---|---|---|---|
| 1 | 0.889 | 308139 | 100.000 | 0.037 | 414289 | 100.000 |

Mass Spectrum

RetTime: 0.575 Datafile: D:\DATA\2022\2202\220209\EW26049-734-P1C1.lcd

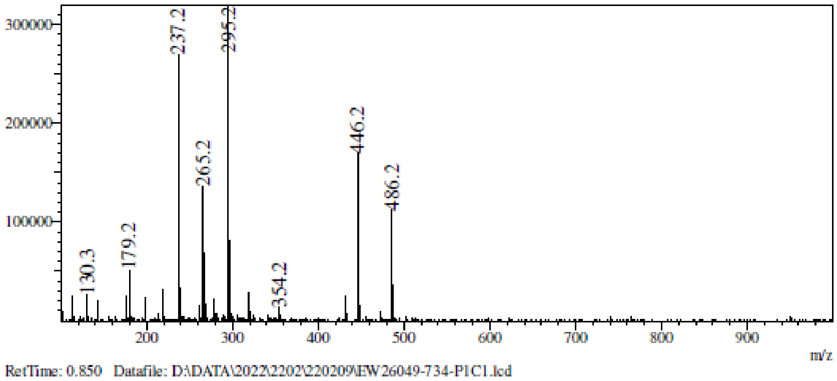

RetTime: 0.850 Datafile: D:\DATA\2022\2202\220209\EW26049-734-P1C1.lcd

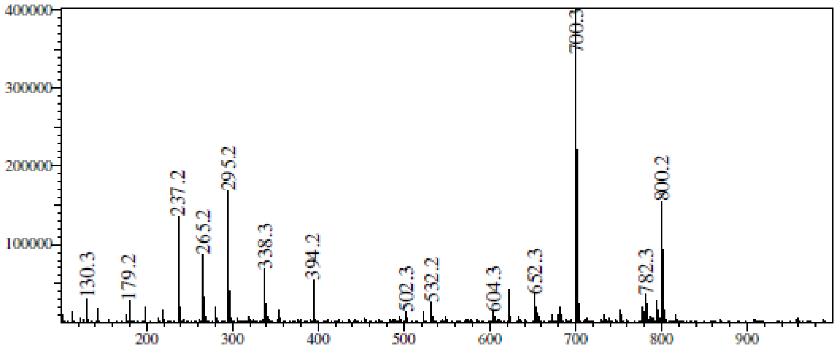

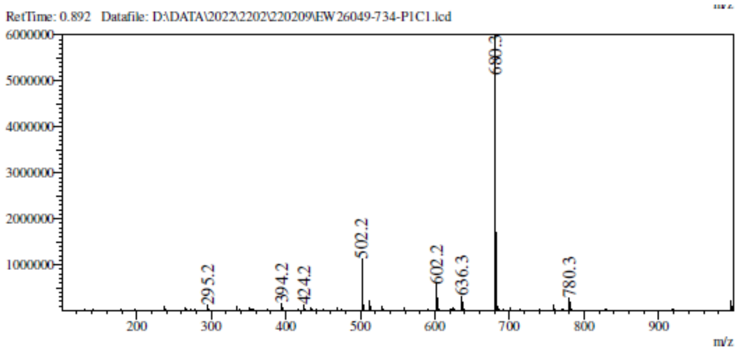

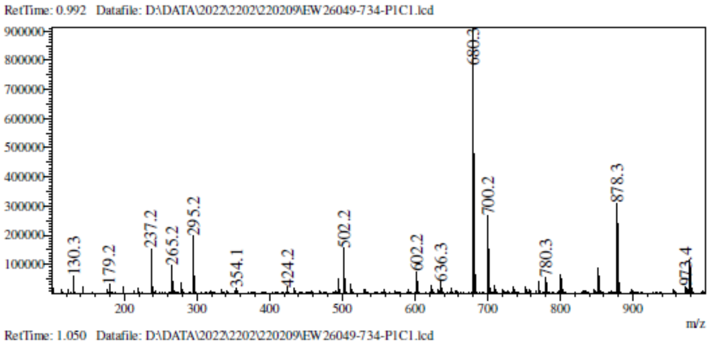

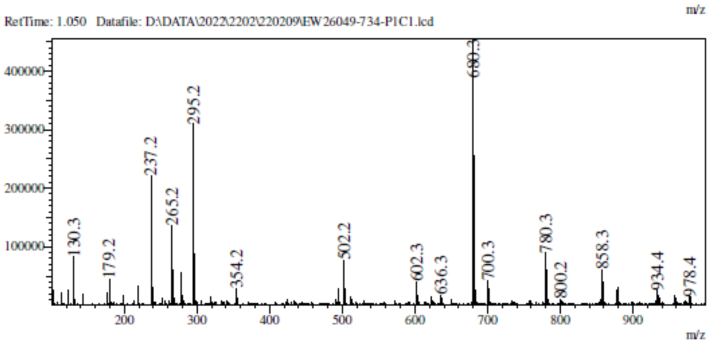

**$^1$H NMR** [(1*R*,2*R*,3*S*,4*R*)-4-Amino-2,3-bis[[tert-butyl(diphenyl)silyl]oxy] cyclopentyl]methanol **(56)**

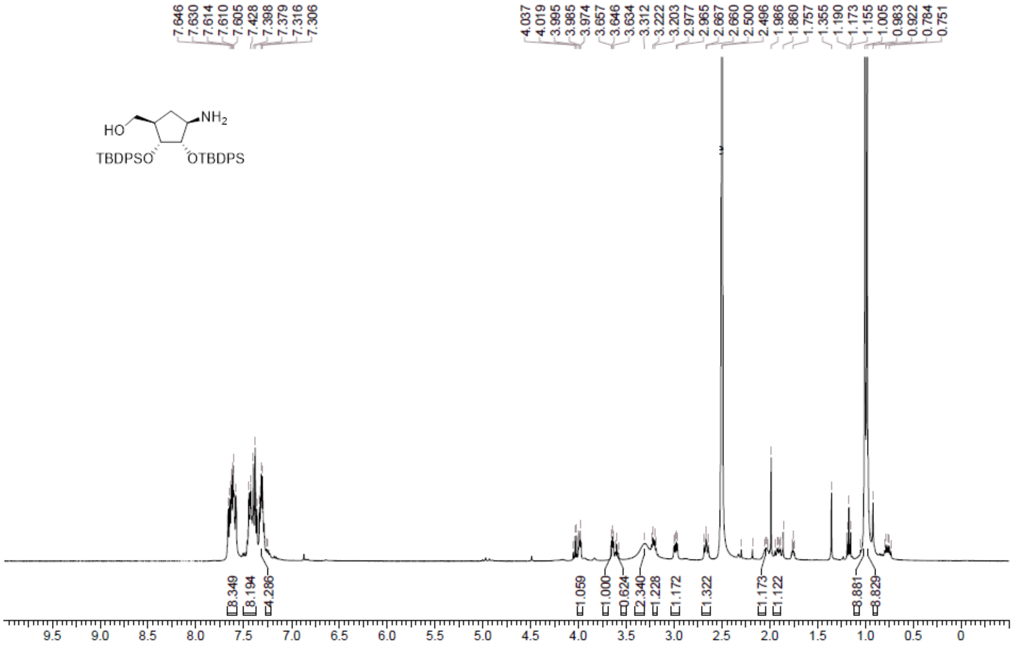

**LCMS** [(1*R*,2*R*,3*S*,4*R*)-4-Amino-2,3-bis[[tert-butyl(diphenyl)silyl]oxy] cyclopentyl]methanol **(56)**

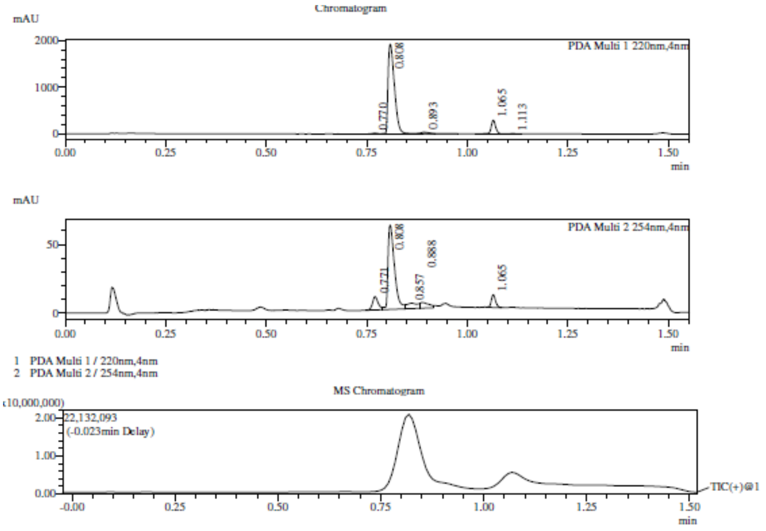

Integration Result

Peak Table

PDA Ch1 220nm

| Peak# | Ret. Time | Height | Height% | USP Width | Area | Area% |
|---|---|---|---|---|---|---|
| 1 | 0.770 | 19528 | 0.859 | 0.030 | 20112 | 0.815 |
| 2 | 0.808 | 1921466 | 84.510 | 0.030 | 2165763 | 87.722 |
| 3 | 0.893 | 30257 | 1.331 | 0.035 | 38798 | 1.571 |
| 4 | 1.065 | 293255 | 12.898 | 0.022 | 233744 | 9.468 |
| 5 | 1.113 | 9141 | 0.402 | 0.030 | 10484 | 0.425 |

Peak Table

PDA Ch2 254nm

| Peak# | Ret. Time | Height | Height% | USP Width | Area | Area% |
|---|---|---|---|---|---|---|
| 1 | 0.771 | 9727 | 11.014 | 0.028 | 9671 | 9.771 |
| 2 | 0.808 | 61503 | 69.644 | 0.030 | 68799 | 69.509 |
| 3 | 0.857 | 3740 | 4.235 | 0.103 | 7591 | 7.670 |
| 4 | 0.888 | 4137 | 4.685 | 0.067 | 6218 | 6.282 |
| 5 | 1.065 | 9204 | 10.422 | 0.022 | 6699 | 6.768 |

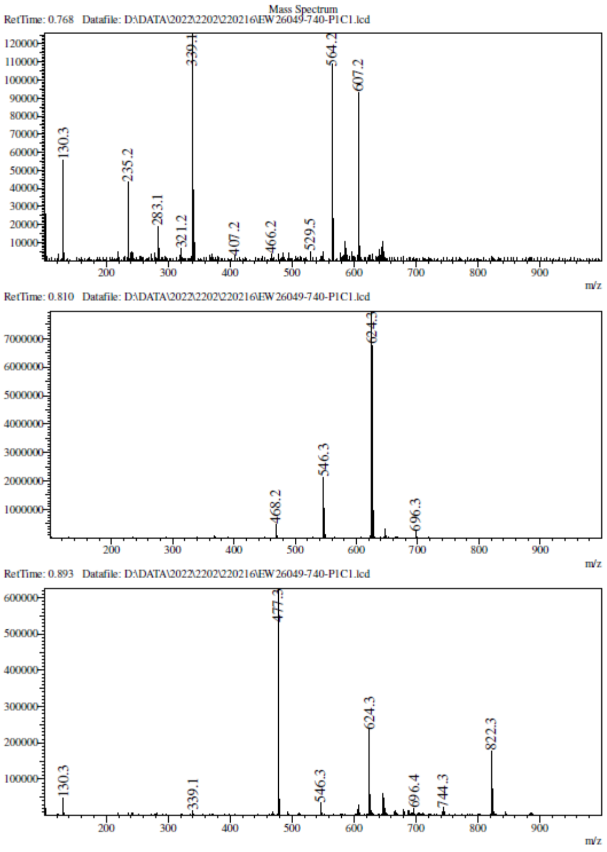

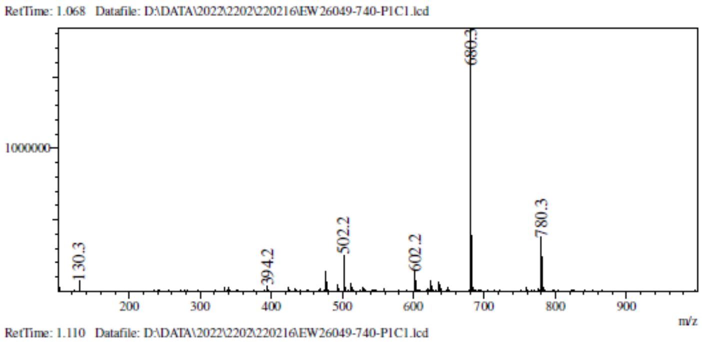

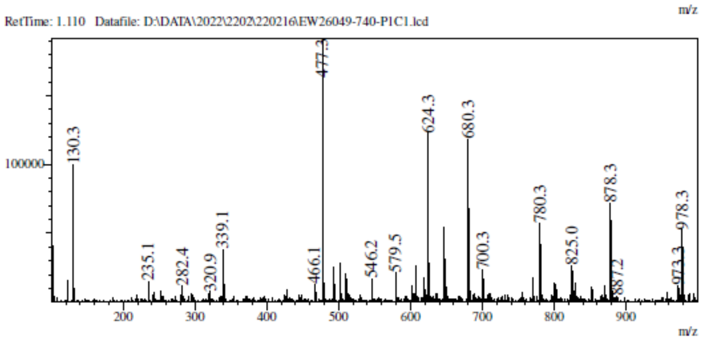

**$^{1}$H NMR** *N*-(4,6-dichloropyrimidin-5-yl)formamide **(57a)**

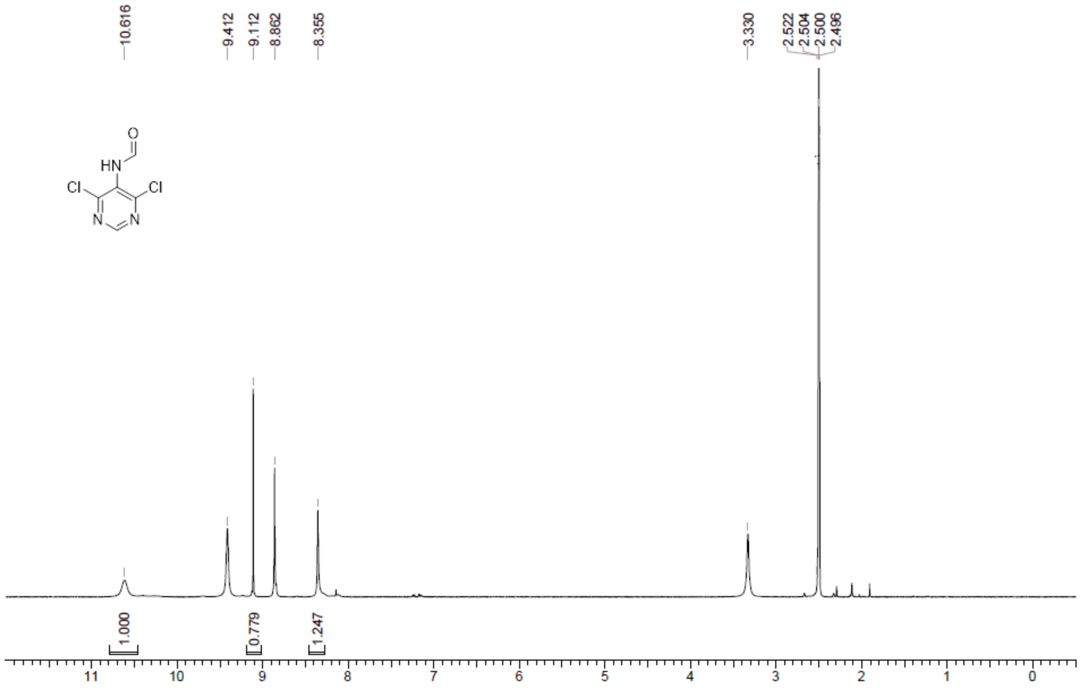

**¹H NMR** [(1*R*,2*R*,3*S*,4*R*)-2,3-Bis[[tert-butyl(diphenyl)silyl]oxy]-4-(6-chloropurin-9-yl)cyclopentyl] methanol **(58a)**

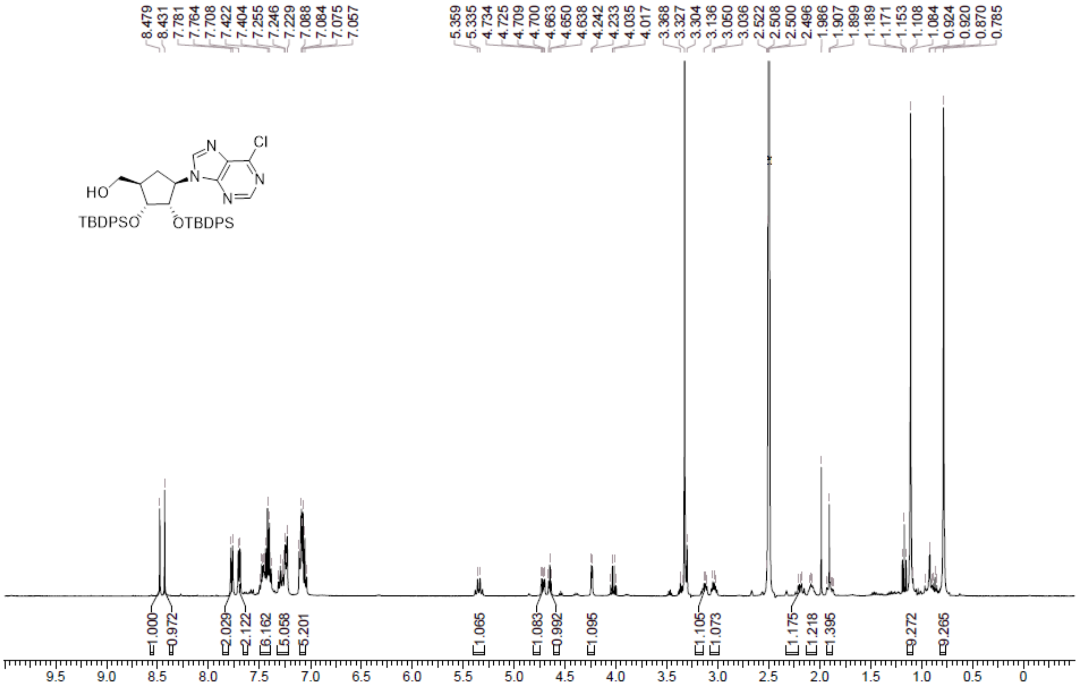

**LCMS** [(1*R*,2*R*,3*S*,4*R*)-2,3-Bis[[tert-butyl(diphenyl)silyl]oxy]-4-(6-chloropurin-9-yl)cyclopentyl] methanol **(58a)**

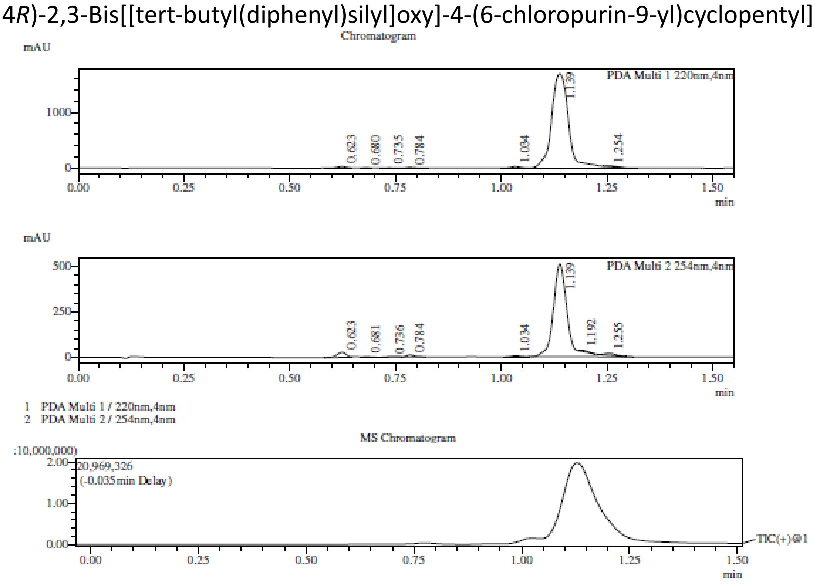

Integration Result

Peak Table

PDA Ch1 220nm

| Peak# | Ret. Time | Height | Height% | USP Width | Area | Area% |
|---|---|---|---|---|---|---|
| 1 | 0.623 | 29126 | 1.628 | 0.042 | 45911 | 0.938 |
| 2 | 0.680 | 11007 | 0.615 | 0.029 | 10774 | 0.220 |
| 3 | 0.735 | 5426 | 0.303 | 0.035 | 7038 | 0.144 |
| 4 | 0.784 | 13591 | 0.760 | 0.035 | 17550 | 0.358 |
| 5 | 1.034 | 27123 | 1.516 | 0.044 | 43629 | 0.891 |
| 6 | 1.139 | 1691265 | 94.520 | 0.064 | 4752648 | 97.067 |
| 7 | 1.254 | 11778 | 0.658 | 0.041 | 18707 | 0.382 |

Peak Table

PDA Ch2 254nm

| Peak# | Ret. Time | Height | Height% | USP Width | Area | Area% |
|---|---|---|---|---|---|---|
| 1 | 0.623 | 28614 | 4.927 | 0.041 | 43739 | 3.358 |
| 2 | 0.681 | 4593 | 0.791 | 0.036 | 5660 | 0.435 |
| 3 | 0.736 | 2859 | 0.492 | 0.052 | 5677 | 0.436 |

Mass Spectrum

RetTime: 0.622 Datafile: D:\DATA\2022\2202\220218\EW26049-749-P1C1.lcd

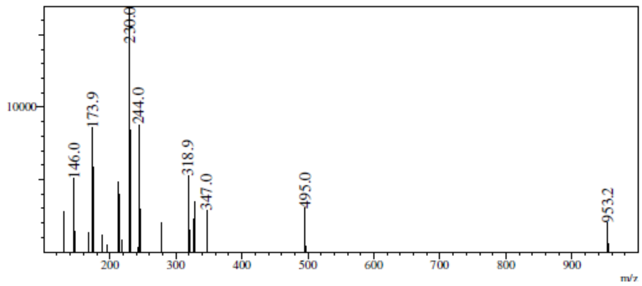

RetTime: 0.678 Datafile: D:\DATA\2022\2202\220218\EW26049-749-P1C1.lcd

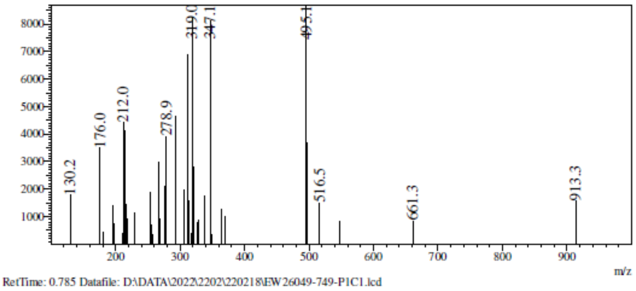

RetTime: 0.785 Datafile: D:\DATA\2022\2202\220218\EW26049-749-P1C1.lcd

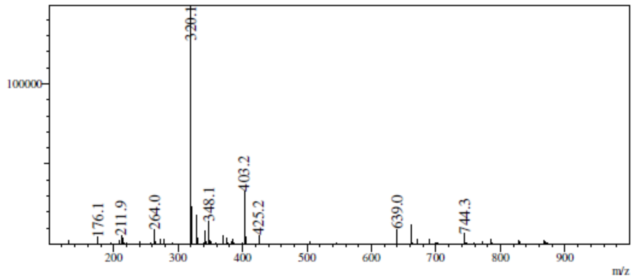

RetTime: 1.035 Datafile: D:\DATA\2022\2202\220218\EW26049-749-P1C1.lcd

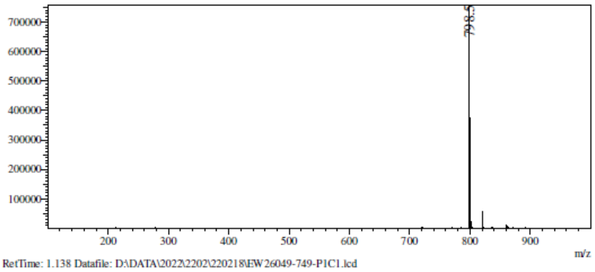

RetTime: 1.138 Datafile: D:\DATA\2022\2202\220218\EW26049-749-P1C1.lcd

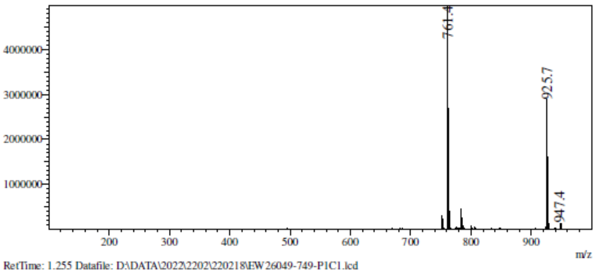

RetTime: 1.255 Datafile: D:\DATA\2022\2202\220218\EW26049-749-P1C1.lcd

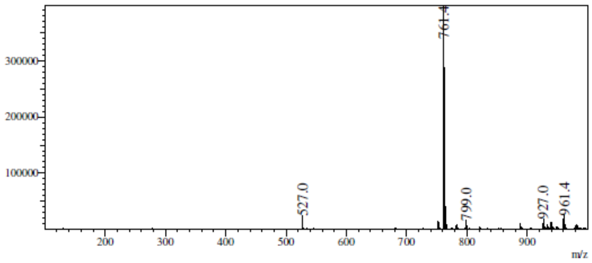

**$^1$H NMR** [(1*R*,2*R*,3*S*,4*R*)-2,3-Bis[[tert-butyl(diphenyl)silyl]oxy]-4-(4-chloropyrrolo[2,3-d]pyrimidin-7-yl)cyclopentyl]methanol **(58b)**

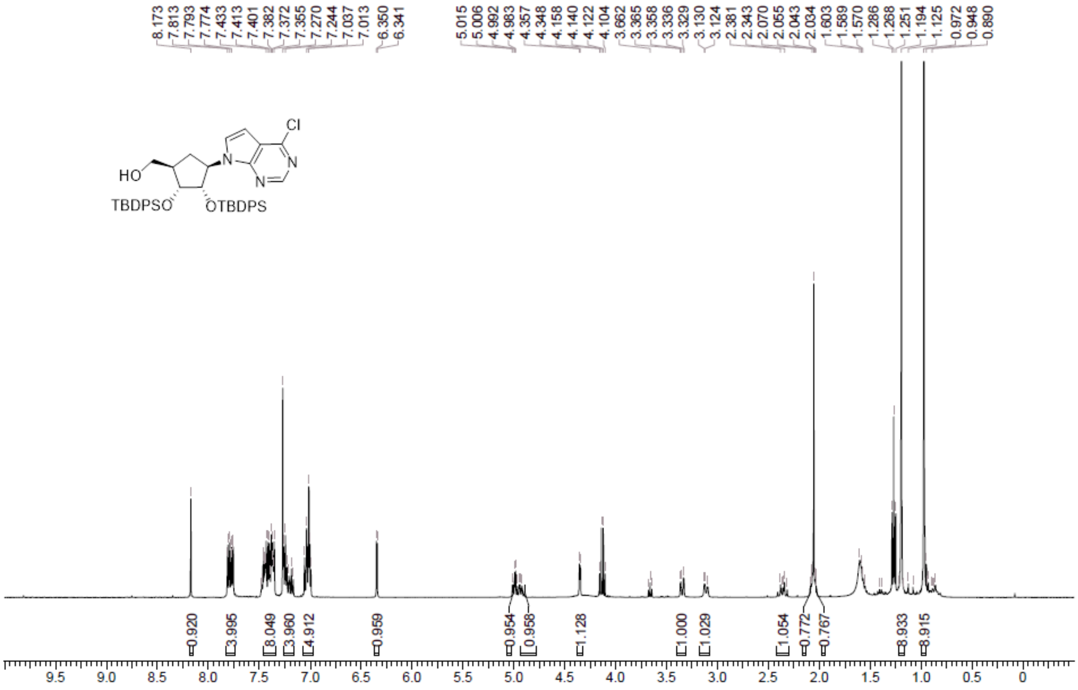

**LCMS** [(1*R*,2*R*,3*S*,4*R*)-2,3-Bis[[tert-butyl(diphenyl)silyl]oxy]-4-(4-chloropyrrolo[2,3-d]pyrimidin-7-yl)cyclopentyl]methanol **(58b)**

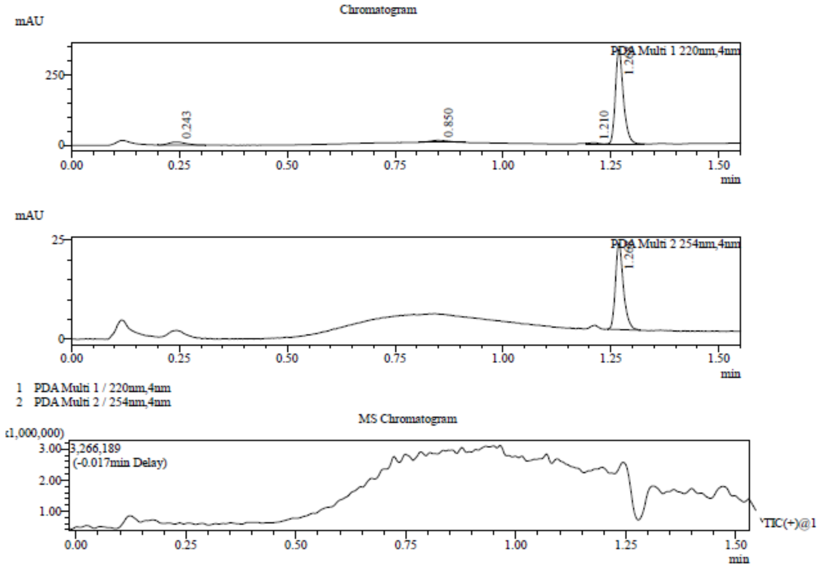

Integration Result

Peak Table

PDA Ch1 220nm

| Peak# | Ret. Time | Height | Height% | USP Width | Area | Area% |
|---|---|---|---|---|---|---|
| 1 | 0.243 | 10742 | 2.959 | 0.073 | 28396 | 5.729 |
| 2 | 0.850 | 7754 | 2.136 | 0.084 | 23277 | 4.696 |
| 3 | 1.210 | 4749 | 1.308 | 0.042 | 6334 | 1.278 |
| 4 | 1.268 | 339802 | 93.597 | 0.035 | 437637 | 88.297 |

Peak Table

PDA Ch2 254nm

| Peak# | Ret. Time | Height | Height% | USP Width | Area | Area% |
|---|---|---|---|---|---|---|
| 1 | 1.268 | 21948 | 100.000 | 0.035 | 28302 | 100.000 |

Mass Spectrum

RetTime: 0.243 Datafile: D:\DATA\2022\2205\220530\EW26049-892-P1C1.lcd

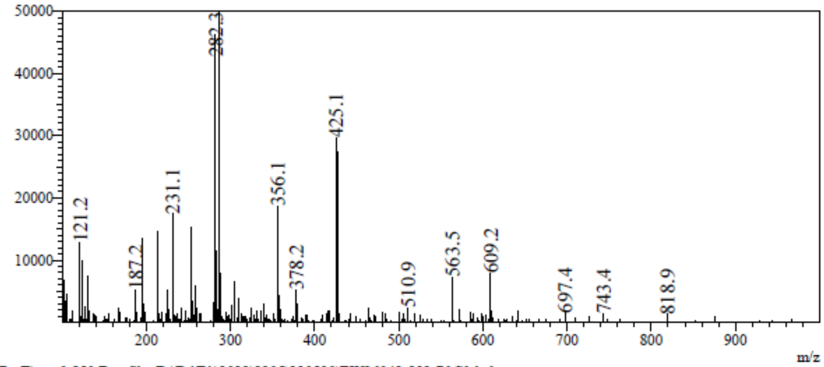

RetTime: 0.850 Datafile: D:\DATA\2022\2205\220530\EW26049-892-P1C1.lcd

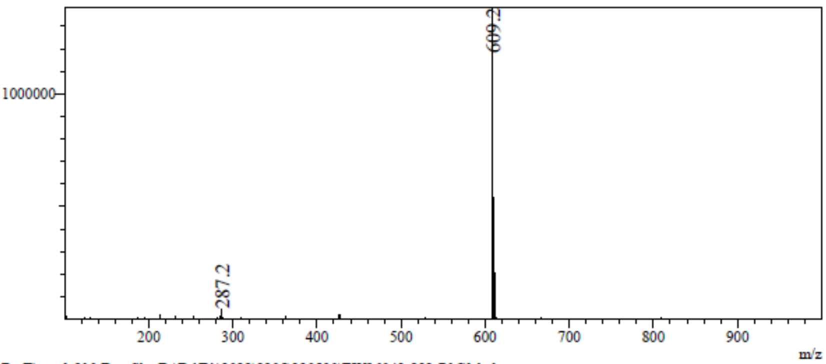

RetTime: 1.210 Datafile: D:\DATA\2022\2205\220530\EW26049-892-P1C1.lcd

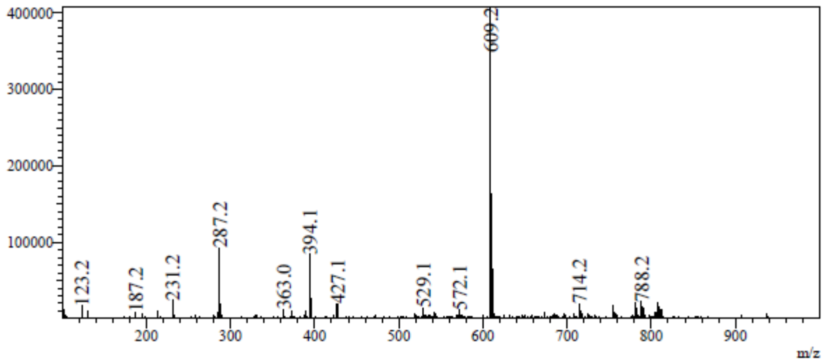

RetTime: 1.270 Datafile: D:\DATA\2022\2205\220530\EW26049-892-P1C1.lcd

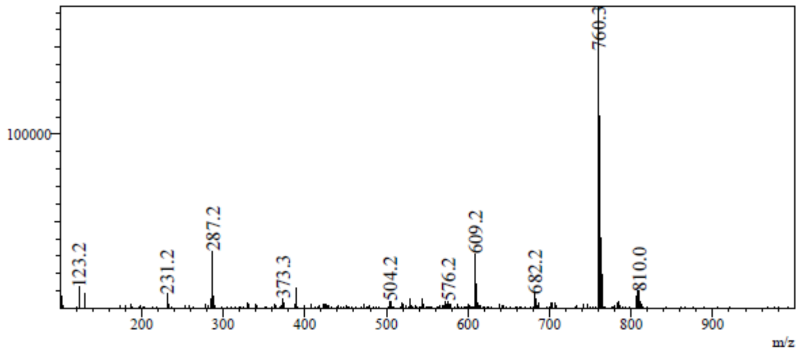

**$^1$H NMR** [(1*R*,2*R*,3*S*,4*R*)-4-(6-Aminopurin-9-yl)-2,3-bis[[tert-butyl(diphenyl) silyl]oxy]cyclopentyl] methanol **(59a)**

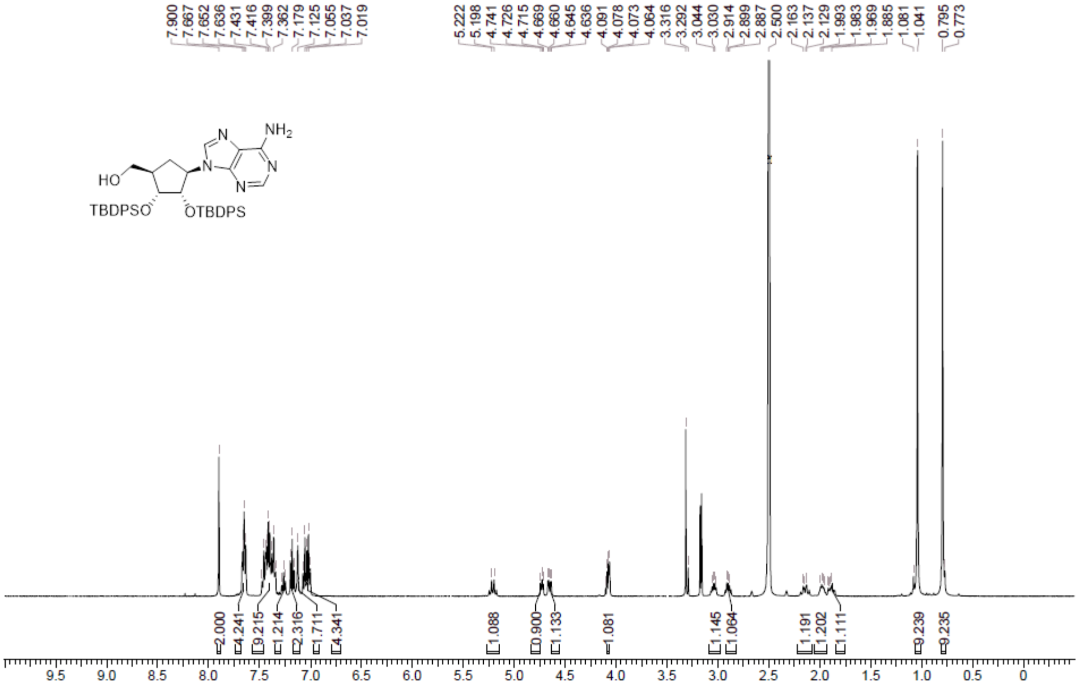

**LCMS** [(1*R*,2*R*,3*S*,4*R*)-4-(6-Aminopurin-9-yl)-2,3-bis[[tert-butyl(diphenyl) silyl]oxy]cyclopentyl] methanol **(59a)**

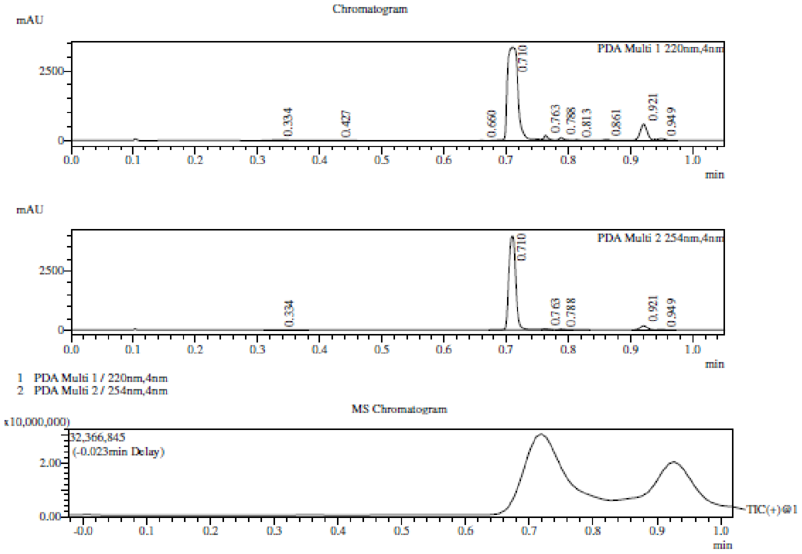

Integration Result

Peak Table

PDA Ch1 220nm

| Peak# | Ret. Time | Height | Height% | USP Width | Area | Area% |
|---|---|---|---|---|---|---|
| 1 | 0.334 | 21694 | 0.498 | 0.070 | 62427 | 1.288 |
| 2 | 0.427 | 4895 | 0.112 | 0.176 | 17195 | 0.355 |
| 3 | 0.660 | 7537 | 0.173 | 0.024 | 5899 | 0.122 |
| 4 | 0.710 | 3345641 | 76.855 | 0.025 | 4058045 | 83.726 |
| 5 | 0.763 | 149749 | 3.440 | 0.012 | 65744 | 1.356 |
| 6 | 0.788 | 101841 | 2.339 | 0.013 | 52753 | 1.088 |
| 7 | 0.813 | 23496 | 0.540 | 0.014 | 11959 | 0.247 |
| 8 | 0.861 | 37883 | 0.870 | 0.017 | 25472 | 0.526 |
| 9 | 0.921 | 590027 | 13.554 | 0.021 | 485661 | 10.020 |
| 10 | 0.949 | 70444 | 1.618 | 0.024 | 61639 | 1.272 |

Peak Table

PDA Ch2 254nm

Mass Spectrum

RetTime: 0.335 Datafile: D:\DATA\2022\2202\220221\EW26049-753-P1C1.lcd

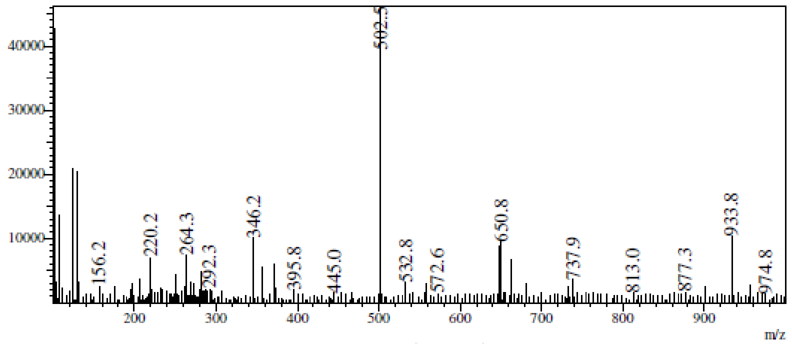

RetTime: 0.710 Datafile: D:\DATA\2022\2202\220221\EW26049-753-P1C1.lcd

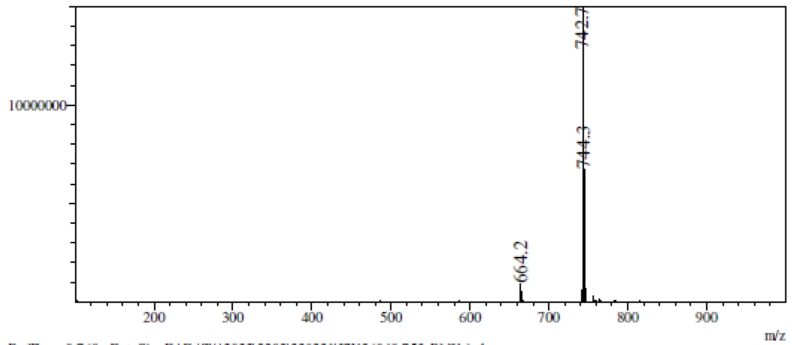

RetTime: 0.760 Datafile: D:\DATA\2022\2202\220221\EW26049-753-P1C1.lcd

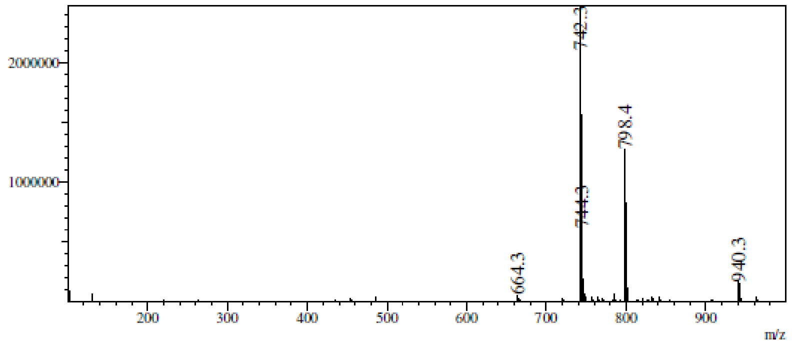

RetTime: 0.785 Datafile: D:\DATA\2022\2202\220221\EW26049-753-P1C1.lcd

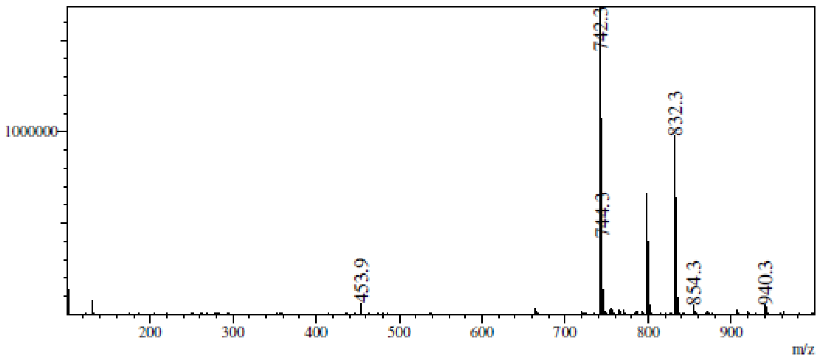

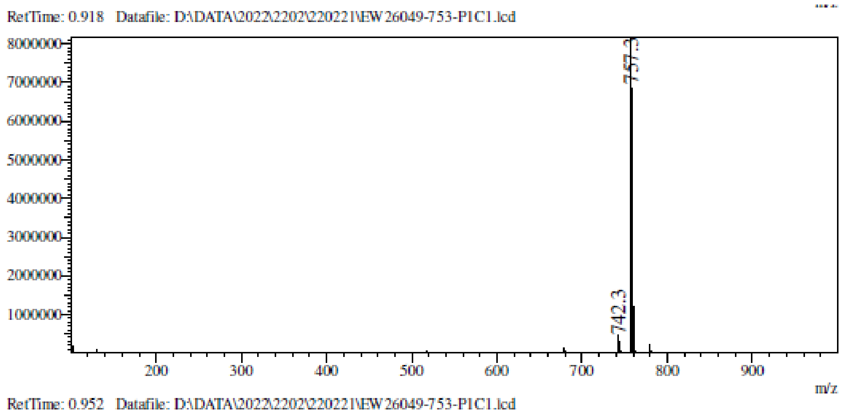

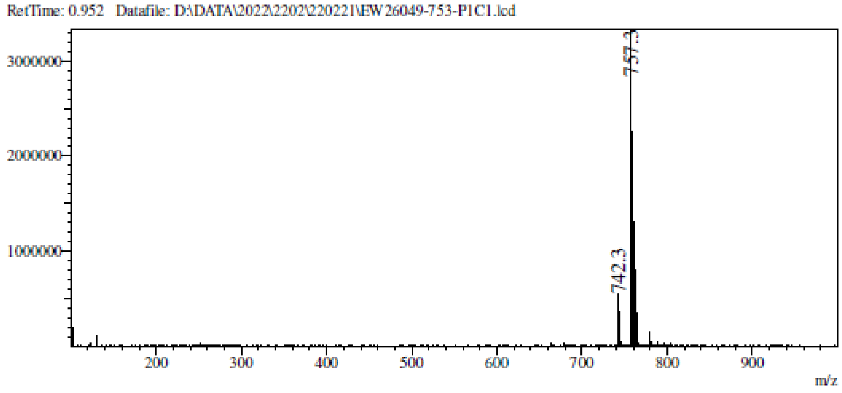

**$^1$H NMR** [(1*R*,2*R*,3*S*,4*R*)-4-(4-Aminopyrrolo[2,3-d]pyrimidin-7-yl)-2,3-bis [[tert-butyl(diphenyl)silyl] oxy]cyclopentyl]methanol **(59b)**

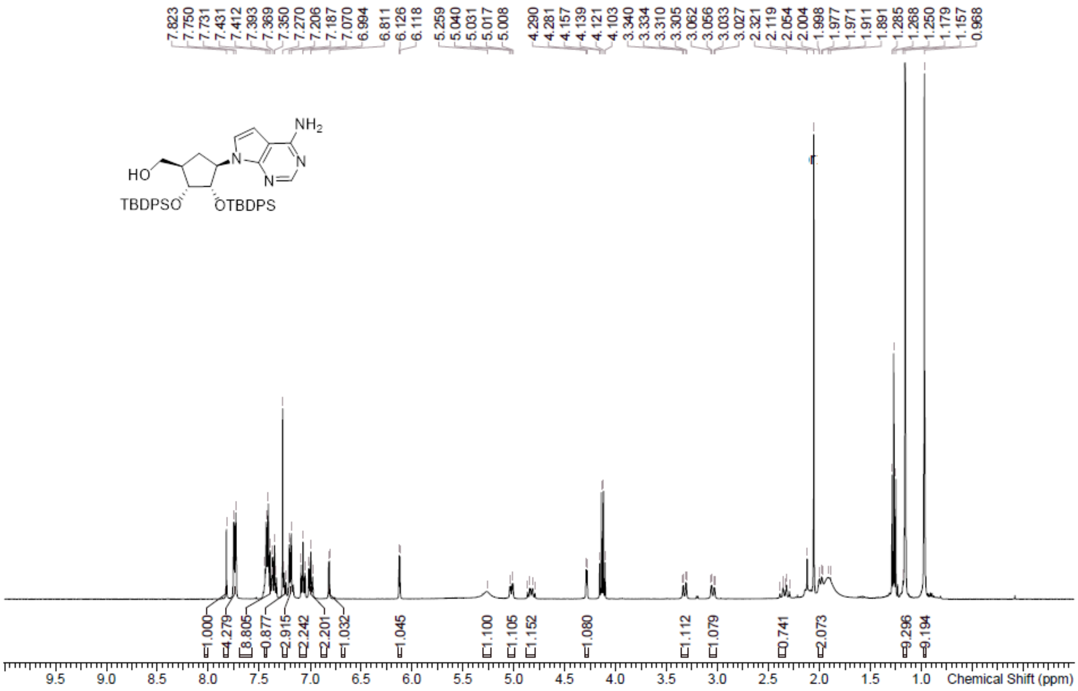

**LCMS** [(1*R*,2*R*,3*S*,4*R*)-4-(4-Aminopyrrolo[2,3-d]pyrimidin-7-yl)-2,3-bis [[tert-butyl(diphenyl)silyl] oxy]cyclopentyl]methanol **(59b)**

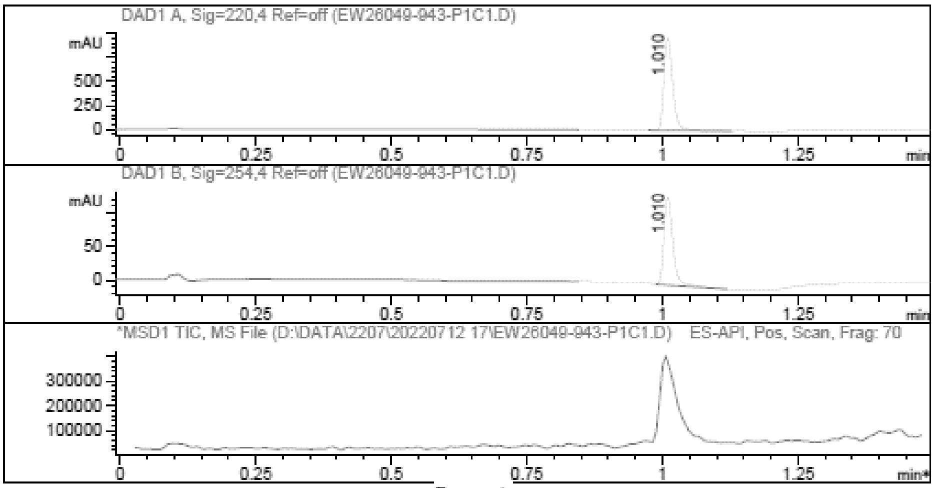

Report

Signal 1 : DAD1 A, Sig=220,4 Ref=off

| # | Meas. Ret. | Height | Width | Area | Area % |
|---|---|---|---|---|---|
| 1 | 1.010 | 964.532 | 0.018 | 1091.277 | 100.000 |

Signal 2 : DAD1 B, Sig=254,4 Ref=off

| # | Meas. Ret. | Height | Width | Area | Area % |
|---|---|---|---|---|---|
| 1 | 1.010 | 130.812 | 0.018 | 151.883 | 100.000 |

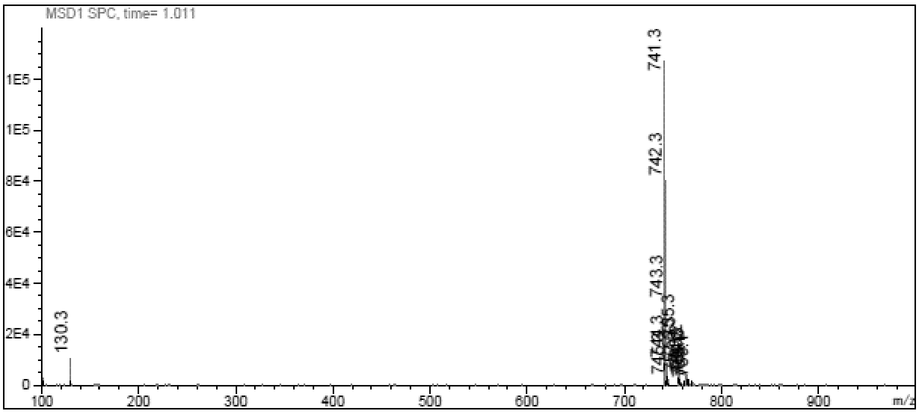

[1]H NMR [(1*R*,2*R*,3*S*,4*R*)-4-(6-Aminopurin-9-yl)-2,3-bis[[tert-butyl(diphenyl) silyl]oxy]cyclopentyl] methyl methanesulfonate **(60a)**

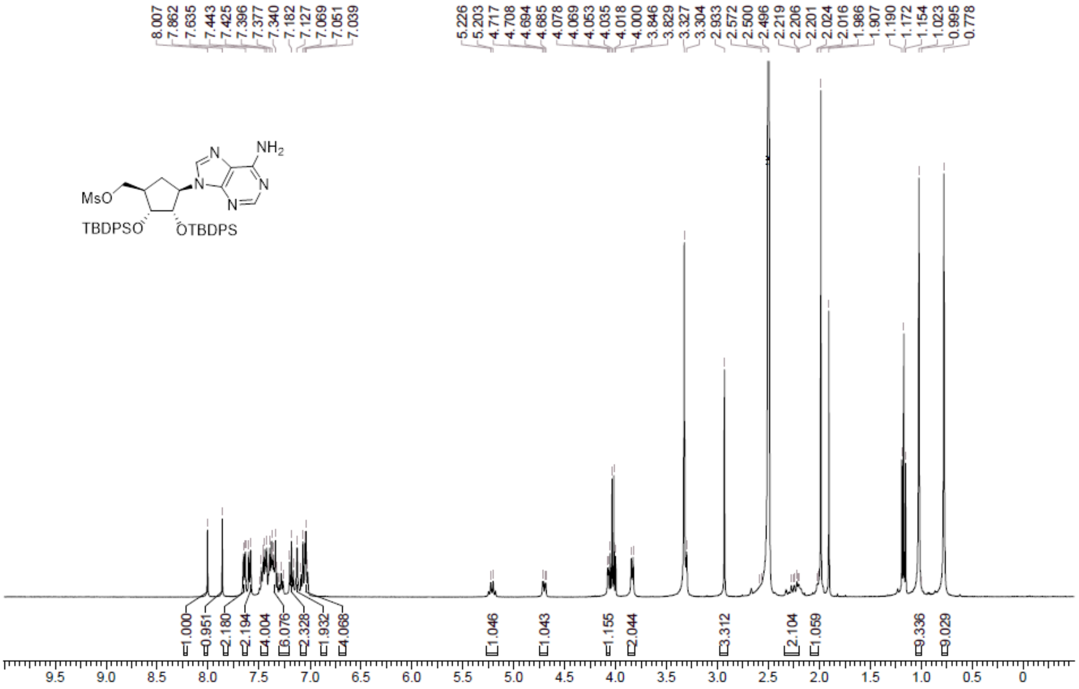

**LCMS** [(1*R*,2*R*,3*S*,4*R*)-4-(6-Aminopurin-9-yl)-2,3-bis[[tert-butyl(diphenyl) silyl]oxy]cyclopentyl] methyl methanesulfonate **(60a)**

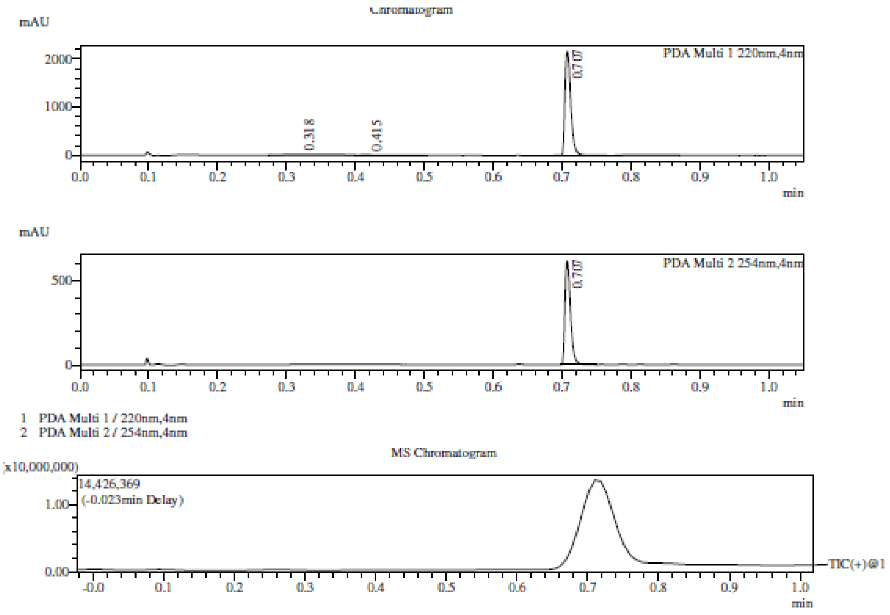

Integration Result

Peak Table

PDA Ch1 220nm

| Peak# | Ret. Time | Height | Height% | USP Width | Area | Area% |
|---|---|---|---|---|---|---|
| 1 | 0.318 | 11191 | 0.519 | 0.134 | 47540 | 3.601 |
| 2 | 0.415 | 7737 | 0.359 | 0.036 | 16666 | 1.262 |
| 3 | 0.707 | 2135871 | 99.122 | 0.015 | 1255957 | 95.136 |

Peak Table

PDA Ch2 254nm

| Peak# | Ret. Time | Height | Height% | USP Width | Area | Area% |
|---|---|---|---|---|---|---|
| 1 | 0.707 | 599679 | 100.000 | 0.014 | 338533 | 100.000 |

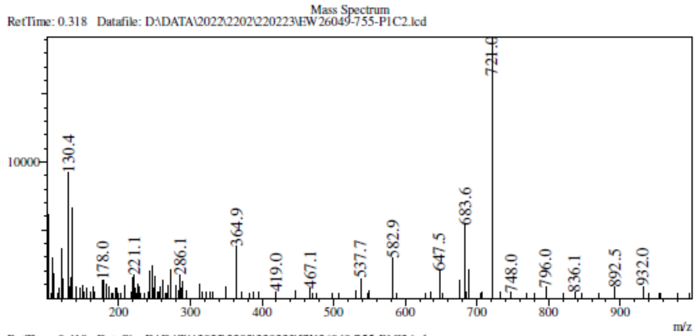

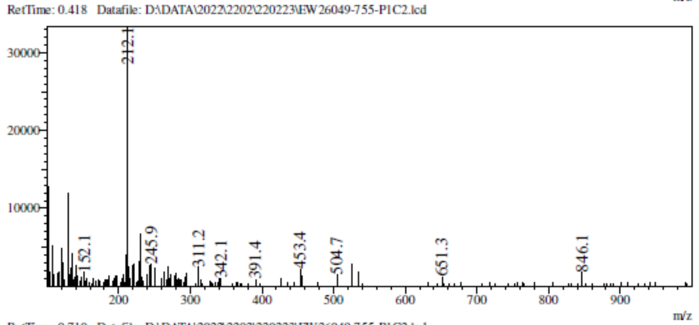

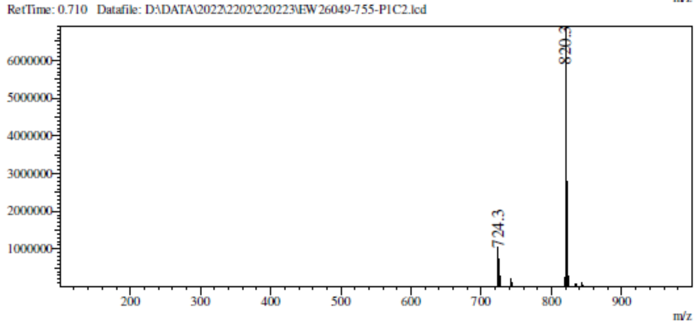

**[1]H NMR** [(1*R*,2*R*,3*S*,4*R*)-4-(4-Aminopyrrolo[2,3-d]pyrimidin-7-yl)-2,3-bis [[tert-butyl(diphenyl) silyl]oxy]cyclopentyl]methyl methanesulfonate **(60b)**

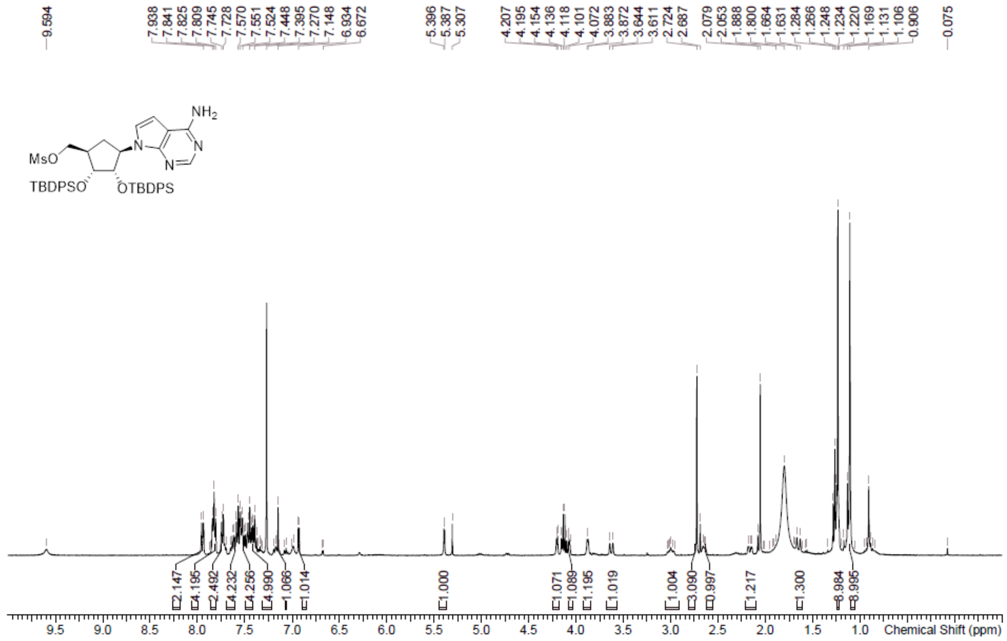

**[1]H NMR** 9-[(1*R*,2*S*,3*R*,4*S*)-4-(1,3-benzothiazol-2-ylsulfanylmethyl)-2,3- bis[[tert-butyl(diphenyl) silyl]oxy]cyclopentyl]purin-6-amine **(61a)**

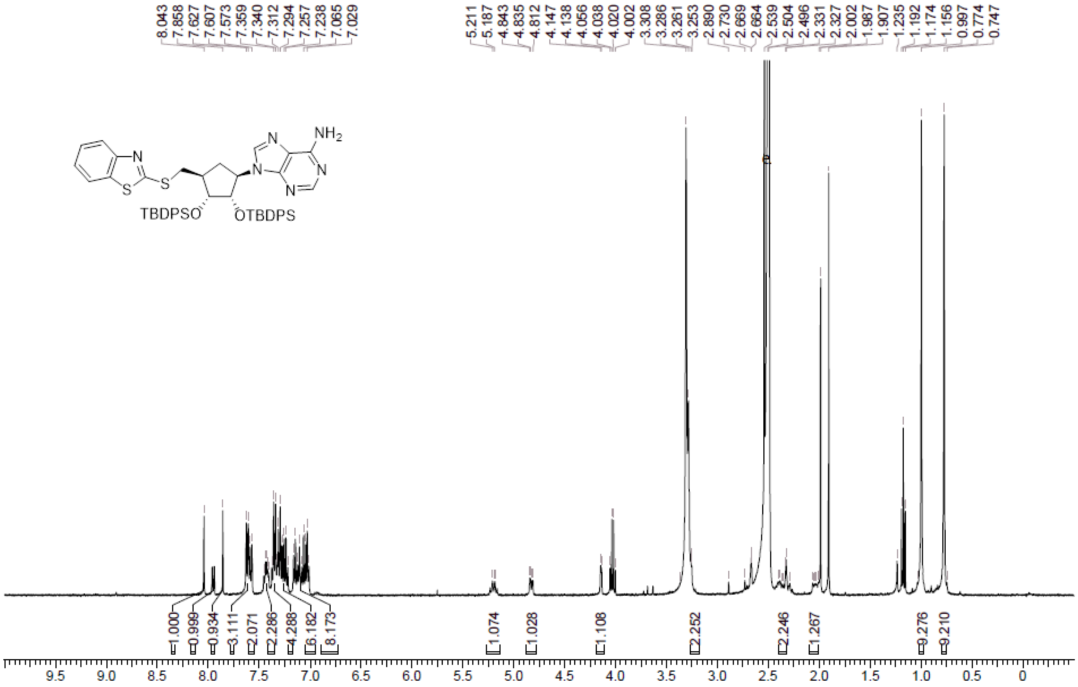

**LCMS** 9-[(1*R*,2*S*,3*R*,4*S*)-4-(1,3-benzothiazol-2-ylsulfanylmethyl)-2,3- bis[[tert-butyl(diphenyl) silyl]oxy]cyclopentyl]purin-6-amine **(61a)**

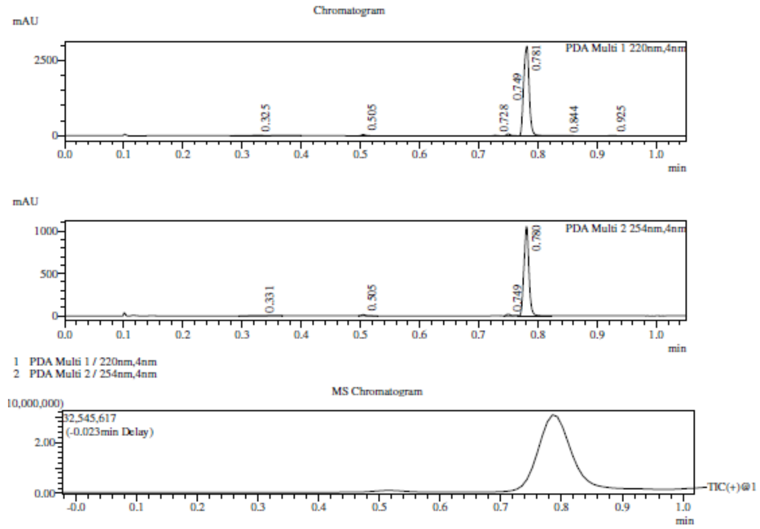

Integration Result

Peak Table

PDA Ch1 220nm

| Peak# | Ret. Time | Height | Height% | USP Width | Area | Area% |
|---|---|---|---|---|---|---|
| 1 | 0.325 | 12737 | 0.411 | 0.071 | 39615 | 1.932 |
| 2 | 0.505 | 41596 | 1.343 | 0.013 | 20613 | 1.005 |
| 3 | 0.728 | 17178 | 0.555 | 0.013 | 9817 | 0.479 |
| 4 | 0.749 | 67089 | 2.166 | 0.012 | 32922 | 1.606 |
| 5 | 0.781 | 2947073 | 95.132 | 0.017 | 1928779 | 94.078 |
| 6 | 0.844 | 9266 | 0.299 | 0.044 | 13018 | 0.635 |
| 7 | 0.925 | 2952 | 0.095 | 0.026 | 5419 | 0.264 |

Peak Table

PDA Ch2 254nm

| Peak# | Ret. Time | Height | Height% | USP Width | Area | Area% |
|---|---|---|---|---|---|---|
| 1 | 0.331 | 3717 | 0.349 | 0.050 | 7580 | 1.199 |
| 2 | 0.505 | 13047 | 1.225 | 0.013 | 6492 | 1.026 |
| 3 | 0.749 | 20319 | 1.907 | 0.012 | 9260 | 1.464 |

Mass Spectrum

RetTime: 0.327 Datafile: D:\DATA\2022\2202\220225\EW26049-759-P1S1.lcd

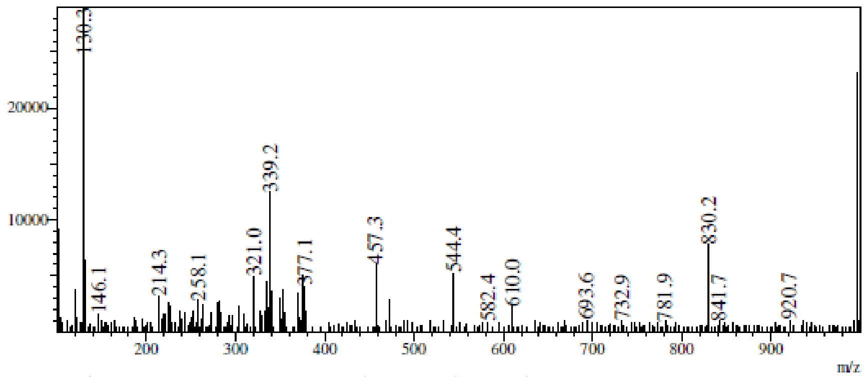

RetTime: 0.502 Datafile: D:\DATA\2022\2202\220225\EW26049-759-P1S1.lcd

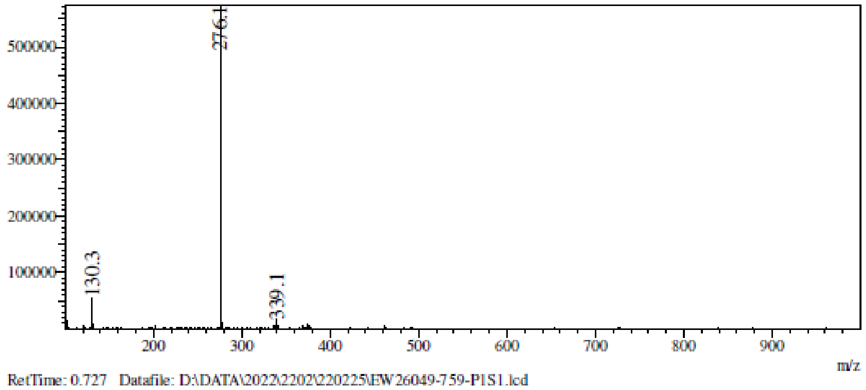

RetTime: 0.727 Datafile: D:\DATA\2022\2202\220225\EW26049-759-P1S1.lcd

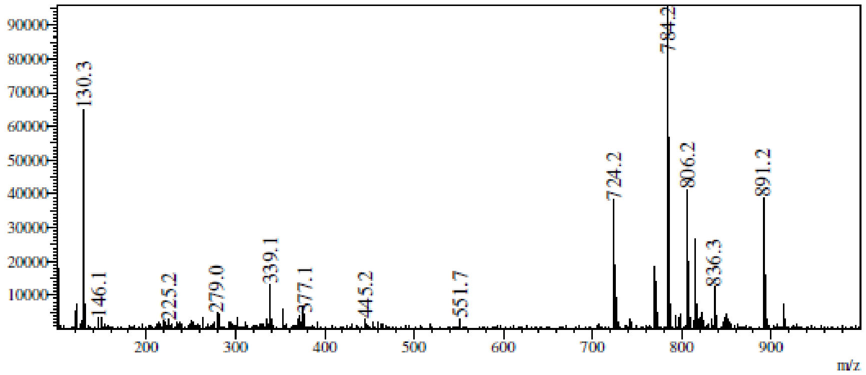

RetTime: 0.752 Datafile: D:\DATA\2022\2202\220225\EW26049-759-P1S1.lcd

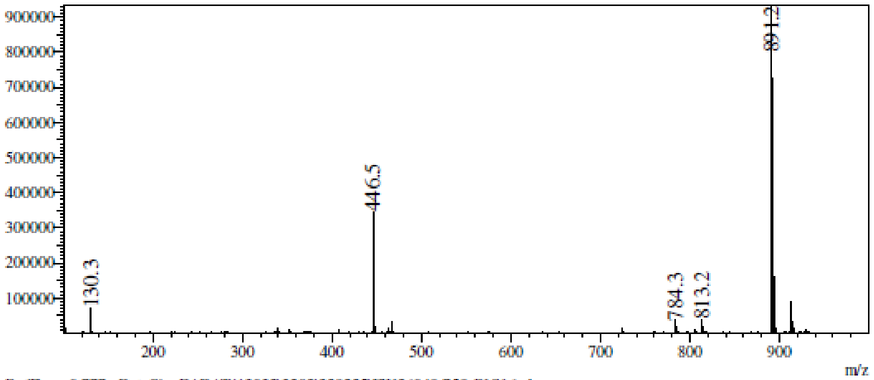

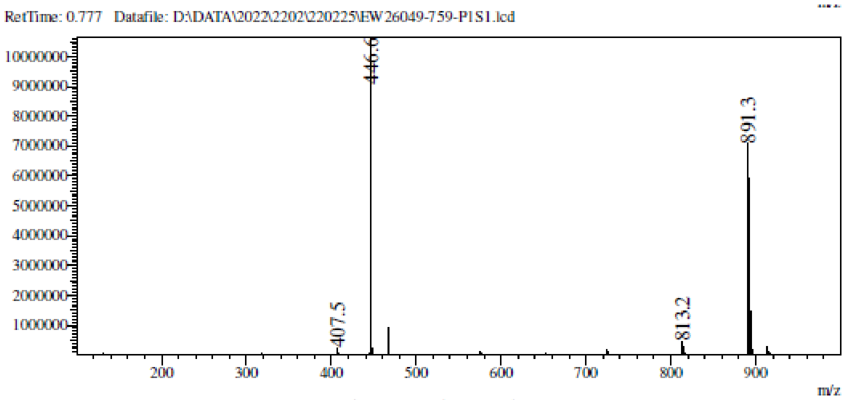

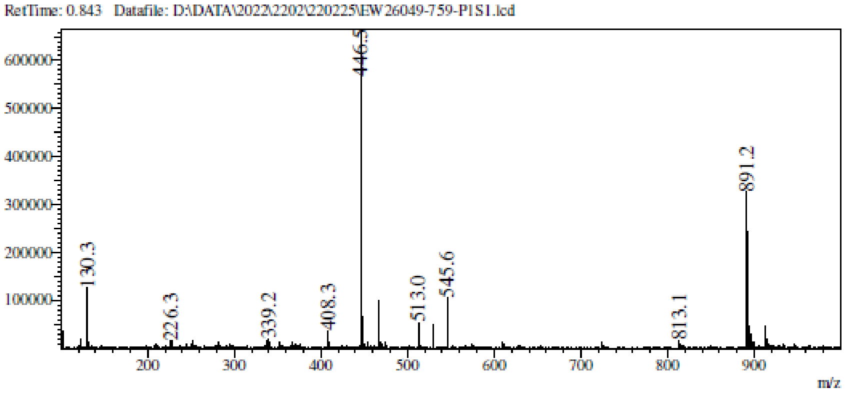

**LCMS** 7-[(1*R*,2*S*,3*R*,4*S*)-4-(1,3-benzothiazol-2-ylsulfanylmethyl)-2,3-bis [[tert-butyl(diphenyl) silyl]oxy]cyclopentyl]pyrrolo[2,3-d]pyrimidin-4-amine **(61b)**

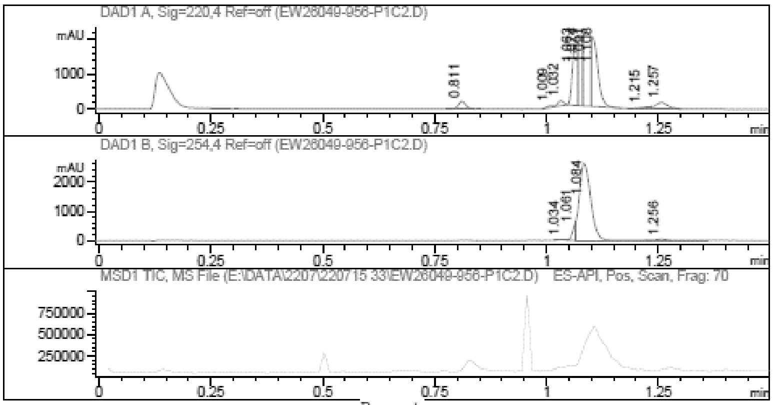

Report

Signal 1 : DAD1 A, Sig=220,4 Ref=off

| # | Meas. Ret. | Height | Width | Area | Area % |
|---|---|---|---|---|---|
| 1 | 0.811 | 220.230 | 0.018 | 251.464 | 3.125 |
| 2 | 1.009 | 51.364 | 0.013 | 43.223 | 0.537 |
| 3 | 1.032 | 138.683 | 0.016 | 135.898 | 1.689 |
| 4 | 1.063 | 1823.765 | 0.013 | 1461.010 | 18.156 |
| 5 | 1.074 | 2049.824 | 0.009 | 1394.962 | 17.335 |
| 6 | 1.091 | 2155.928 | 0.016 | 2324.598 | 28.888 |
| 7 | 1.108 | 1817.417 | 0.017 | 2049.738 | 25.472 |
| 8 | 1.215 | 35.014 | 0.026 | 66.348 | 0.825 |
| 9 | 1.257 | 178.943 | 0.027 | 319.715 | 3.973 |

Signal 2 : DAD1 B, Sig=254,4 Ref=off

| # | Meas. Ret. | Height | Width | Area | Area % |
|---|---|---|---|---|---|
| 1 | 1.034 | 21.738 | 0.017 | 23.470 | 0.467 |
| 2 | 1.061 | 459.130 | 0.008 | 245.248 | 4.878 |
| 3 | 1.084 | 2601.377 | 0.028 | 4643.146 | 92.361 |
| 4 | 1.256 | 44.536 | 0.036 | 115.307 | 2.294 |

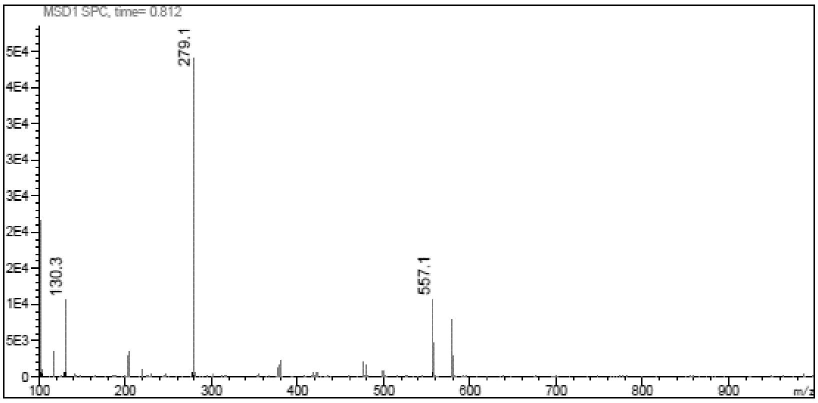
MSD1 SPC, time= 0.812
279.1
130.3
557.1
m/z

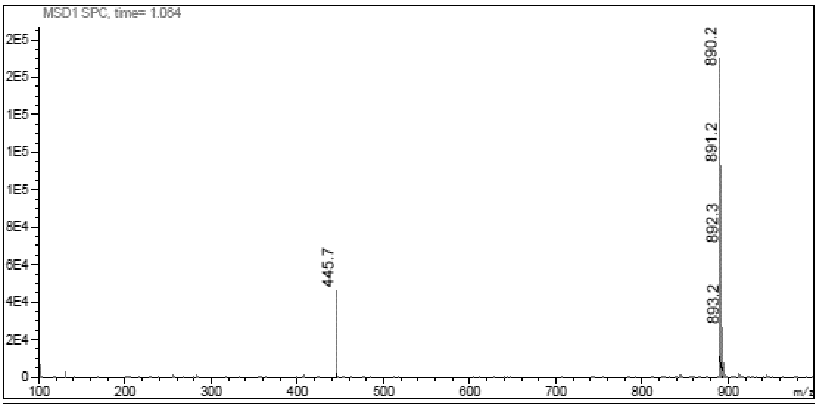
MSD1 SPC, time= 1.064
890.2
891.2
892.3
893.2
445.7
m/z

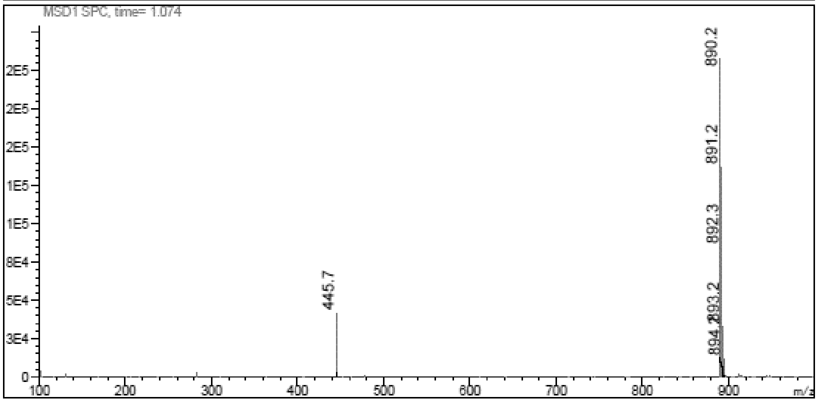
MSD1 SPC, time= 1.074
890.2
891.2
892.3
893.2
445.7
m/z

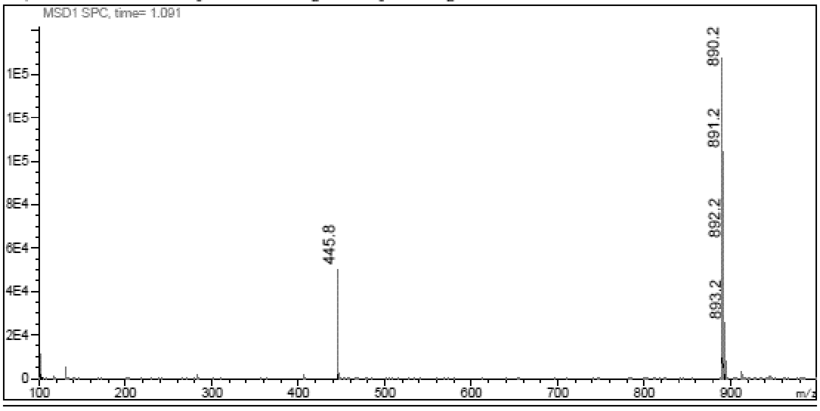
MSD1 SPC, time= 1.091
890.2
891.2
892.2
893.2
445.8
m/z

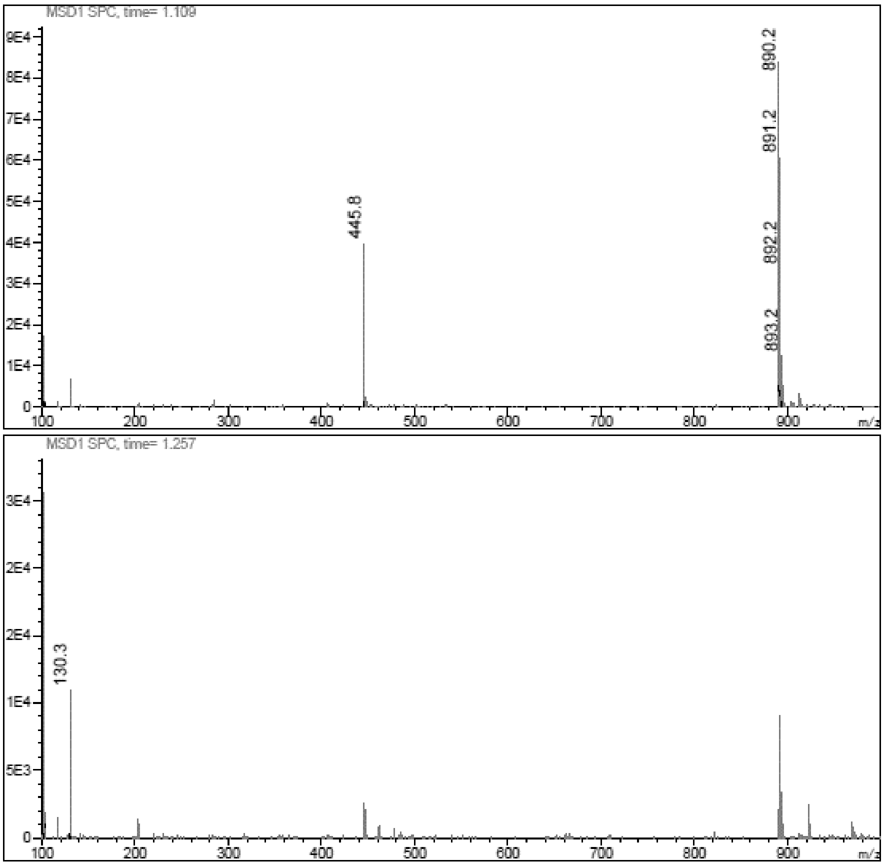

**$^1$H NMR (**1*R*,2*S*,3*R*,5*S*)-3-(6-aminopurin-9-yl)-5-(1,3-benzothiazol-2-ylsulfanylmethyl) cyclopentane-1,2-diol **(26)**

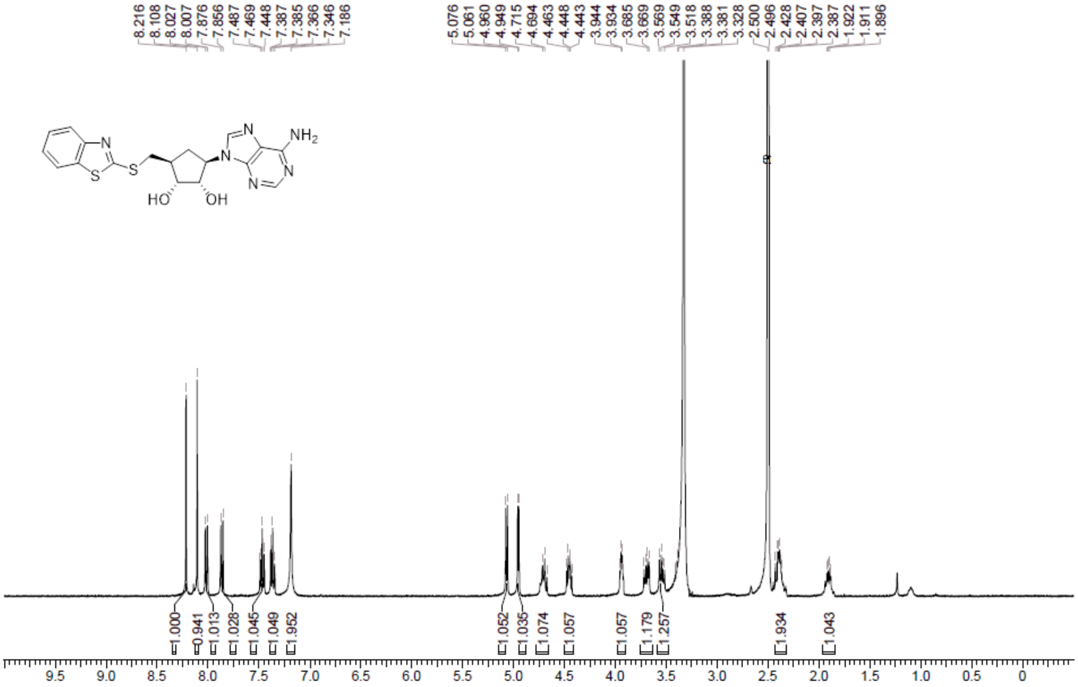

**HPLC** (1*R*,2*S*,3*R*,5*S*)-3-(6-aminopurin-9-yl)-5-(1,3-benzothiazol-2-ylsulfanylmethyl) cyclopentane-1,2-diol **(26)**

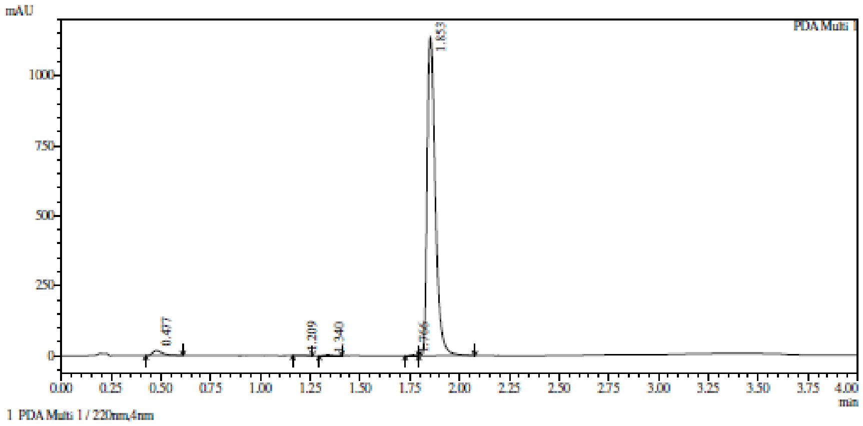

Integration result

PeakTable

PDA Ch1 220nm

| Peak# | Ret. Time | USP Width | Resolution | Height | Area | Area % |
|---|---|---|---|---|---|---|
| 1 | 0.477 | 0.085 | 0.000 | 18134 | 61258 | 1.834 |
| 2 | 1.209 | 0.087 | 8.511 | 972 | 3238 | 0.097 |
| 3 | 1.340 | 0.073 | 1.643 | 3527 | 9774 | 0.293 |
| 4 | 1.766 | 0.150 | 3.809 | 3712 | 9135 | 0.274 |
| 5 | 1.853 | 0.072 | 0.786 | 1139604 | 3256019 | 97.502 |
| Total | | | | 1165949 | 3339423 | 100.000 |

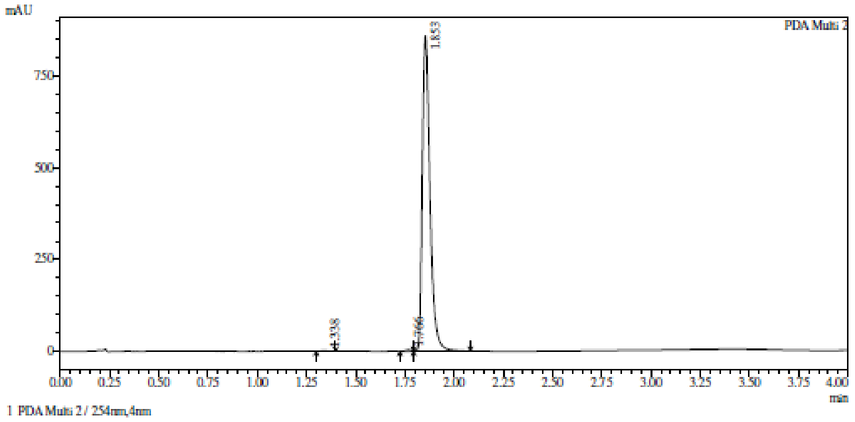

Integration result

PeakTable

PDA Ch2 254nm

| Peak# | Ret. Time | USP Width | Resolution | Height | Area | Area % |
|---|---|---|---|---|---|---|
| 1 | 1.338 | 0.070 | 0.000 | 1558 | 4177 | 0.177 |
| 2 | 1.766 | 0.083 | 5.598 | 3563 | 8857 | 0.375 |
| 3 | 1.853 | 0.069 | 1.142 | 858441 | 2350682 | 99.449 |
| Total | | | | 863562 | 2363716 | 100.000 |

**HRMC (**$1R,2S,3R,5S$)-3-(6-aminopurin-9-yl)-5-(1,3-benzothiazol-2-ylsulfanylmethyl) cyclopentane-1,2-diol **(26)**

**Compound Table**

| Label | Tgt Score | Mass Error (ppm) | Tgt Formula | Obs. RT | Ref. Mass | Obs. Mass |
|---|---|---|---|---|---|---|
| Cpd 1: C18 H18 N6 O2 S2; 1.843 | 98.45 | 3.29 | C18 H18 N6 O2 S2 | 1.843 | 414.0933 | 414.0946 |

| Obs. m/z | Obs. RT | Obs. Mass | Tgt Formula | Tgt Mass | Tgt Mass Error (ppm) | Find Cpds Algorithm |
|---|---|---|---|---|---|---|
| 437.0826 | 1.843 | 414.0946 | C18 H18 N6 O2 S2 | 414.0933 | 3.29 | Find by Formula |

**Compound Chromatograms**

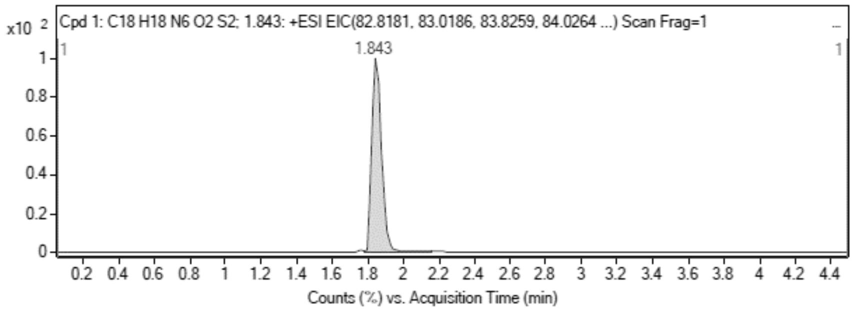

**MS Zoomed Spectrum**

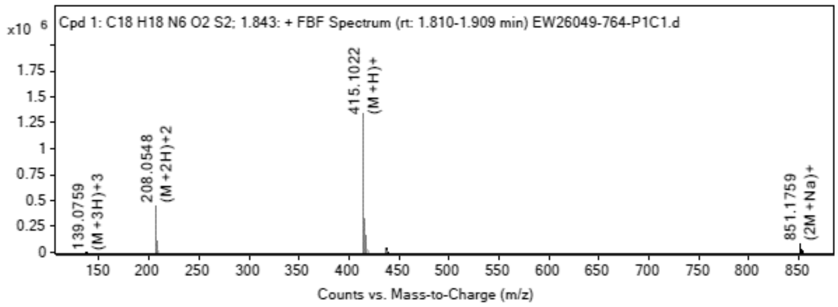

**MS Spectrum Peak List**

| Obs. m/z | Charge | Abund | Ion/Isotope |
|---|---|---|---|
| 208.0548 | 2 | 453201.69 | (M+2H)+2 |
| 208.5559 | 2 | 107129.74 | (M+2H)+2 |
| 209.0536 | 2 | 54589.17 | (M+2H)+2 |
| 415.1022 | 1 | 1340788.38 | (M+H)+ |
| 416.1039 | 1 | 324647.41 | (M+H)+ |
| 417.0994 | 1 | 154169.38 | (M+H)+ |
| 437.0826 | 1 | 42784.64 | (M+Na)+ |
| 851.1759 | 1 | 81445.9 | (2M+Na)+ |
| 852.1782 | 1 | 36897.6 | (2M+Na)+ |
| 853.1748 | 1 | 23124.36 | (2M+Na)+ |

MS Zoomed Spectrum

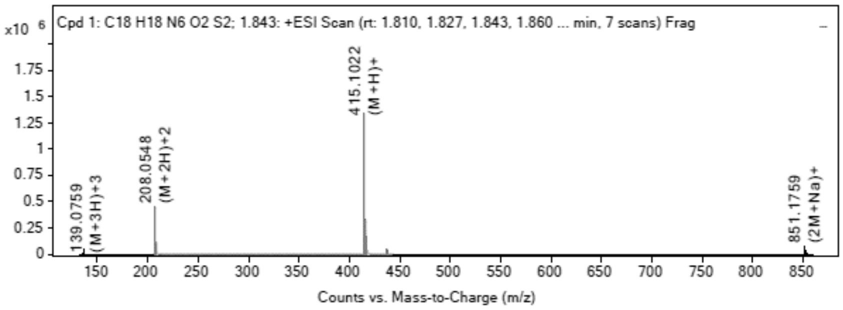

**MS Spectrum Peak List**

| Obs. m/z | Charge | Abund | Ion/Isotope | Tgt Mass Error (ppm) |
|---|---|---|---|---|
| 208.0548 | 2 | 453201.69 | (M+2H)+2 | -4.05 |
| 208.5559 | 2 | 107129.74 | (M+2H)+2 | -3.25 |
| 209.0536 | 2 | 54589.17 | (M+2H)+2 | -2.85 |
| 415.1022 | 1 | 1340788.38 | (M+H)+ | -4 |
| 416.1039 | 1 | 324647.41 | (M+H)+ | -2.04 |
| 417.0994 | 1 | 154169.38 | (M+H)+ | -1.77 |
| 437.0826 | 1 | 42784.64 | (M+Na)+ | -0.35 |
| 851.1759 | 1 | 81445.9 | (2M+Na)+ | -0.17 |
| 852.1782 | 1 | 36897.6 | (2M+Na)+ | 0.07 |
| 853.1748 | 1 | 23124.36 | (2M+Na)+ | 0.44 |

--- End Of Report ---

**$^1$H NMR** (1*R*,2*S*,3*R*,5*S*)-3-(4-Aminopyrrolo[2,3-d]pyrimidin-7-yl)-5-(1,3-benzothiazol-2-ylsulfanylmethyl)cyclopentane-1,2-diol **(28)**

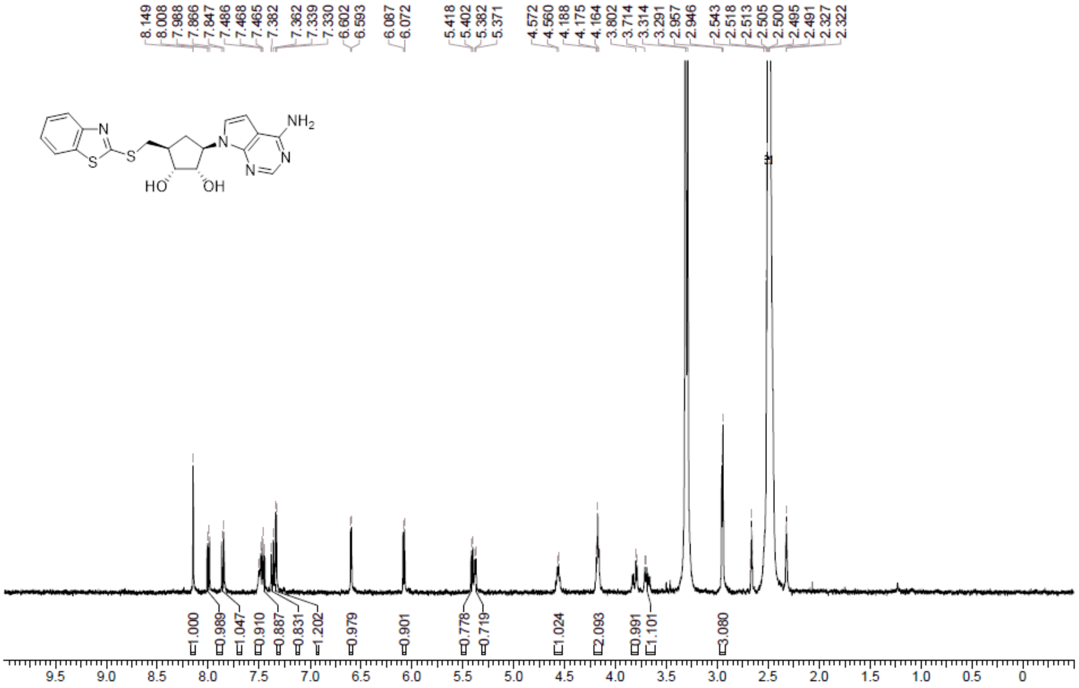

**HPLC** (1*R*,2*S*,3*R*,5*S*)-3-(4-Aminopyrrolo[2,3-d]pyrimidin-7-yl)-5-(1,3-benzothiazol-2-ylsulfanylmethyl)cyclopentane-1,2-diol **(28)**

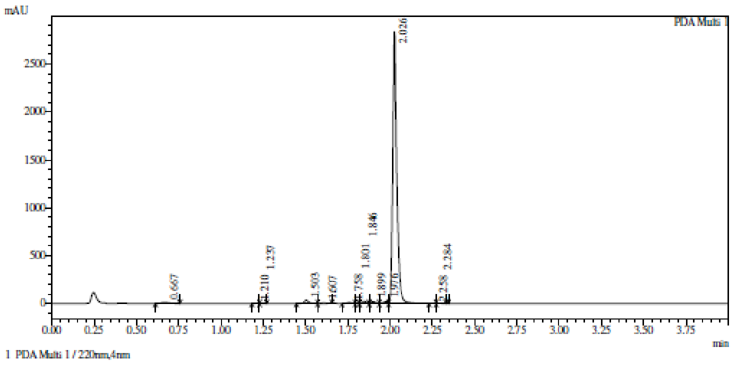

Integration result

PeakTable

PDA Ch1 220nm

| Peak# | Ret. Time | USP Width | Resolution | Height | Area | Area % |
|---|---|---|---|---|---|---|
| 1 | 0.667 | 0.086 | 0.000 | 10725 | 34642 | 0.722 |
| 2 | 1.210 | 0.068 | 7.060 | 1283 | 2158 | 0.045 |
| 3 | 1.237 | 0.310 | 0.144 | 869 | 1163 | 0.024 |
| 4 | 1.503 | 0.047 | 1.487 | 34428 | 61622 | 1.284 |
| 5 | 1.607 | 0.046 | 2.249 | 5755 | 9737 | 0.203 |
| 6 | 1.758 | 0.058 | 2.878 | 9308 | 20233 | 0.422 |
| 7 | 1.801 | 0.221 | 0.311 | 2374 | 3026 | 0.063 |
| 8 | 1.846 | 0.050 | 0.333 | 21108 | 38373 | 0.800 |
| 9 | 1.899 | 0.051 | 1.039 | 19016 | 35022 | 0.730 |
| 10 | 1.976 | 0.070 | 1.288 | 20050 | 37761 | 0.787 |
| 11 | 2.026 | 0.041 | 0.895 | 2835412 | 4545369 | 94.742 |
| 12 | 2.258 | 0.079 | 3.878 | 2674 | 4127 | 0.086 |
| 13 | 2.284 | 0.102 | 0.289 | 2500 | 4376 | 0.091 |
| Total | | | | 2965502 | 4797608 | 100.000 |

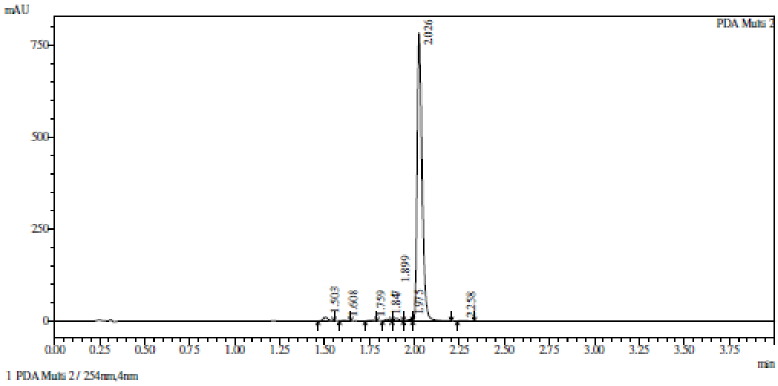

Integration result

PeakTable

PDA Ch2 254nm

| Peak# | Ret. Time | USP Width | Resolution | Height | Area | Area % |
|---|---|---|---|---|---|---|
| 1 | 1.503 | 0.047 | 0.000 | 10837 | 18901 | 1.257 |
| 2 | 1.608 | 0.045 | 2.296 | 1779 | 2827 | 0.188 |
| 3 | 1.759 | 0.050 | 3.178 | 1918 | 3515 | 0.234 |
| 4 | 1.847 | 0.047 | 1.813 | 4866 | 8135 | 0.541 |
| 5 | 1.899 | 0.046 | 1.118 | 6456 | 10951 | 0.728 |
| 6 | 1.975 | 0.076 | 1.243 | 3751 | 7379 | 0.491 |
| 7 | 2.026 | 0.049 | 0.813 | 782552 | 1449925 | 96.431 |
| 8 | 2.258 | 0.076 | 3.738 | 606 | 1949 | 0.130 |
| Total | | | | 812765 | 1503583 | 100.000 |

**HRMC** (1*R*,2*S*,3*R*,5*S*)-3-(4-Aminopyrrolo[2,3-d]pyrimidin-7-yl)-5-(1,3-benzothiazol-2-ylsulfanylmethyl)cyclopentane-1,2-diol **(28)**

**Compound Table**

| Label | Tgt Score | Mass Error (ppm) | Tgt Formula | Obs. RT | Ref. Mass | Obs. Mass |
|---|---|---|---|---|---|---|
| Cpd 1: C19 H19 N5 O3 S2; 1.796 | 90.28 | -4.45 | C19 H19 N5 O3 S2 | 1.796 | 429.0929 | 429.091 |

| Obs. *m/z* | Obs. RT | Obs. Mass | Tgt Formula | Tgt Mass | Tgt Mass Error (ppm) | Find Cpds Algorithm |
|---|---|---|---|---|---|---|
| 430.0987 | 1.796 | 429.091 | C19 H19 N5 O3 S2 | 429.0929 | -4.45 | Find by Formula |

**Compound Chromatograms**

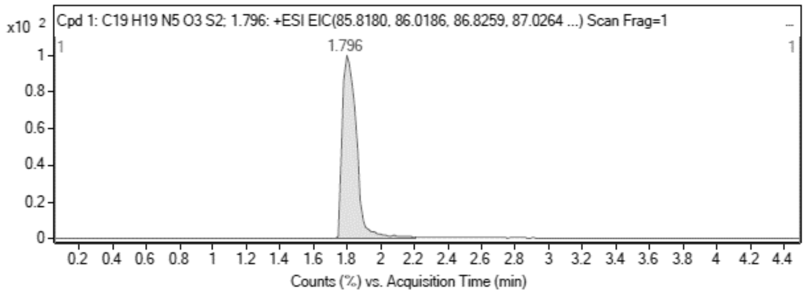

**MS Zoomed Spectrum**

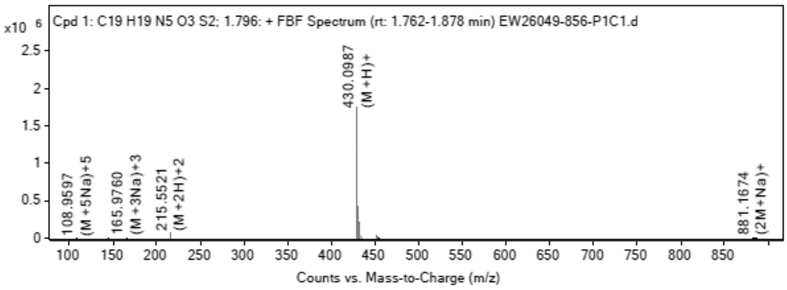

**MS Spectrum Peak List**

| Obs. *m/z* | Charge | Abund | Ion/Isotope |
|---|---|---|---|
| 215.5521 | 2 | 69571.91 | (M+2H)+2 |
| 216.0534 | 2 | 16955.47 | (M+2H)+2 |

| | | | |
|---|---|---|---|
| 216.5517 | 2 | 8361.99 | (M+2H)+2 |
| 430.0987 | 1 | 1749183.63 | (M+H)+ |
| 431.1005 | 1 | 431888.06 | (M+H)+ |
| 432.0961 | 1 | 211083.06 | (M+H)+ |
| 452.0787 | 1 | 29897.41 | (M+Na)+ |
| 453.0816 | 1 | 7462.57 | (M+Na)+ |
| 454.0776 | 1 | 3630.51 | (M+Na)+ |
| 881.1674 | 1 | 5226.82 | (2M+Na)+ |

MS Zoomed Spectrum

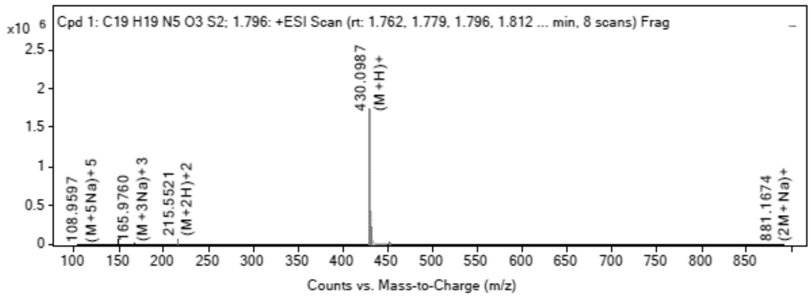

MS Spectrum Peak List

| Obs. *m/z* | Charge | Abund | Ion/Isotope | Tgt Mass Error (ppm) |
|---|---|---|---|---|
| 215.5521 | 2 | 69571.91 | (M+2H)+2 | 7.53 |
| 216.0534 | 2 | 16955.47 | (M+2H)+2 | 7.88 |
| 216.5517 | 2 | 8361.99 | (M+2H)+2 | 5.8 |
| 430.0987 | 1 | 1749183.63 | (M+H)+ | 3.52 |
| 431.1005 | 1 | 431888.06 | (M+H)+ | 5.48 |
| 432.0961 | 1 | 211083.06 | (M+H)+ | 5.65 |
| 452.0787 | 1 | 29897.41 | (M+Na)+ | 7.57 |
| 453.0816 | 1 | 7462.57 | (M+Na)+ | 7.08 |
| 454.0776 | 1 | 3630.51 | (M+Na)+ | 6.48 |
| 881.1674 | 1 | 5226.82 | (2M+Na)+ | 8.76 |

--- End Of Report ---

**$^1$H NMR** [(2*R*,3*R*,4*R*,5*R*)-3,4-Dibenzoyloxy-5-(4-chloropyrrolo[2,3-d] pyrimidin-7-yl)tetrahydrofuran-2-yl]methyl benzoate **(101)**

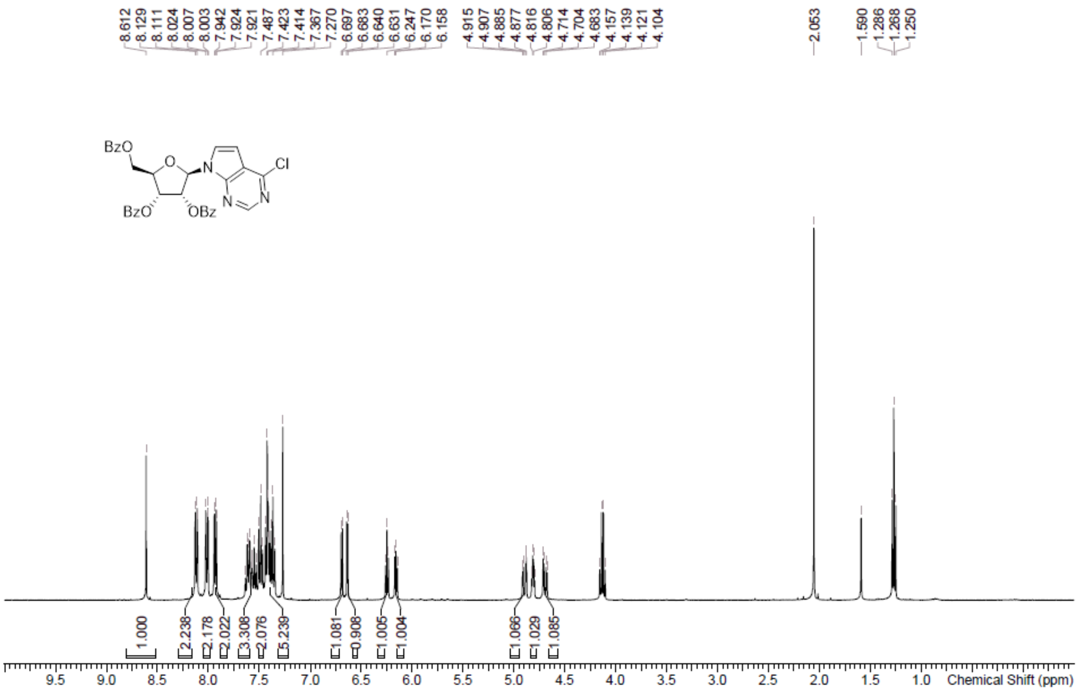

**LCMS** [(2*R*,3*R*,4*R*,5*R*)-3,4-Dibenzoyloxy-5-(4-chloropyrrolo[2,3-d] pyrimidin-7-yl)tetrahydrofuran-2-yl]methyl benzoate **(101)**

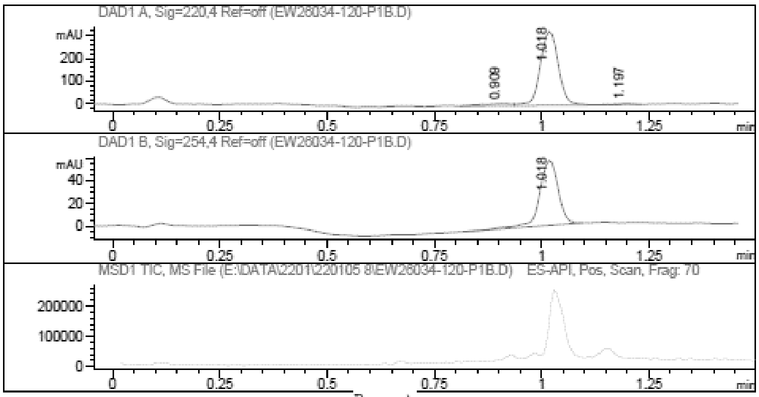

Report

Signal 1 : DAD1 A, Sig=220,4 Ref=off

| # | Meas. Ret. | Height | Width | Area | Area % |
|---|---|---|---|---|---|
| 1 | 0.909 | 9.563 | 0.062 | 44.034 | 4.364 |
| 2 | 1.018 | 325.152 | 0.046 | 950.210 | 94.168 |
| 3 | 1.197 | 4.475 | 0.049 | 14.809 | 1.468 |

Signal 2 : DAD1 B, Sig=254,4 Ref=off

| # | Meas. Ret. | Height | Width | Area | Area % |
|---|---|---|---|---|---|
| 1 | 1.018 | 57.387 | 0.047 | 172.272 | 100.000 |

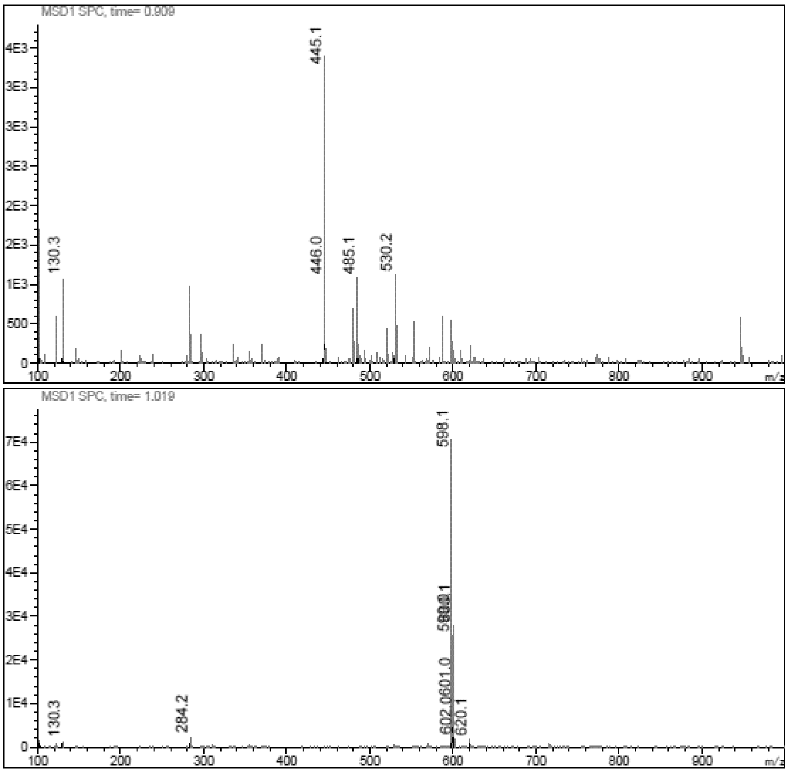

**LCMS** (2*R*,3*R*,4*S*,5*R*)-2-(4-Aminopyrrolo[2,3-d]pyrimidin-7-yl)-5-(hydroxymethyl)tetrahydrofuran-3,4-diol **(102)**

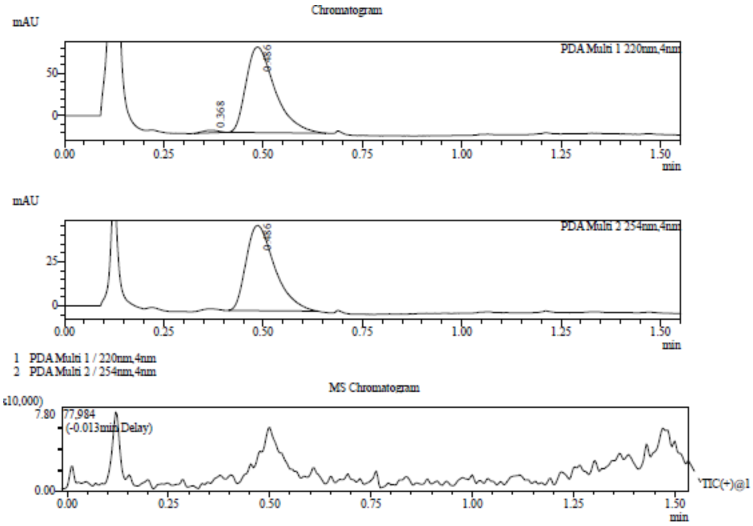

Integration Result

Peak Table

PDA Ch1 220nm

| Peak# | Ret. Time | Height | Height% | USP Width | Area | Area% |
|---|---|---|---|---|---|---|
| 1 | 0.368 | 3132 | 3.016 | 0.080 | 8474 | 1.626 |
| 2 | 0.486 | 100718 | 96.984 | 0.130 | 512634 | 98.374 |

Peak Table

PDA Ch2 254nm

| Peak# | Ret. Time | Height | Height% | USP Width | Area | Area% |
|---|---|---|---|---|---|---|
| 1 | 0.486 | 48023 | 100.000 | 0.130 | 241560 | 100.000 |

**[1]H NMR** 7-[(2*R*,3*R*,4*R*,5*R*)-3,4-Bis[[tert-butyl(dimethyl)silyl]oxy]-5-[[tert-butyl(dimethyl)silyl] oxymethyl]tetrahydrofuran-2-yl]pyrrolo[2,3-d]pyrimidin-4-amine **(103)**

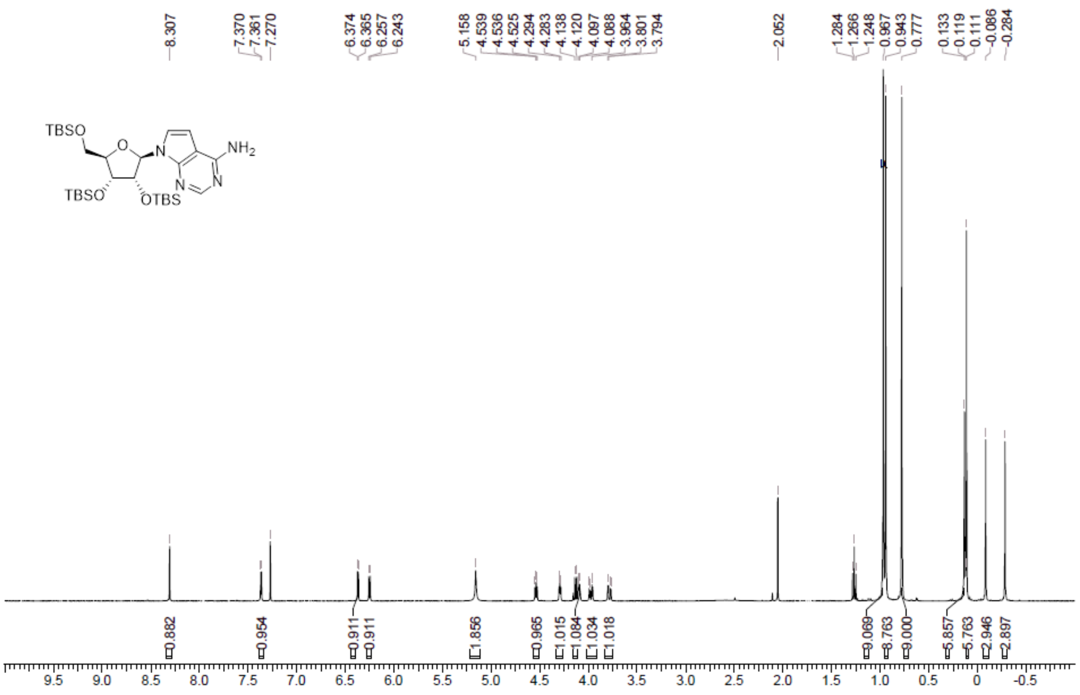

**LCMS** 7-[(2*R*,3*R*,4*R*,5*R*)-3,4-Bis[[tert-butyl(dimethyl)silyl]oxy]-5-[[tert- butyl(dimethyl)silyl] oxymethyl]tetrahydrofuran-2-yl]pyrrolo[2,3-d]pyrimidin-4-amine **(103)**

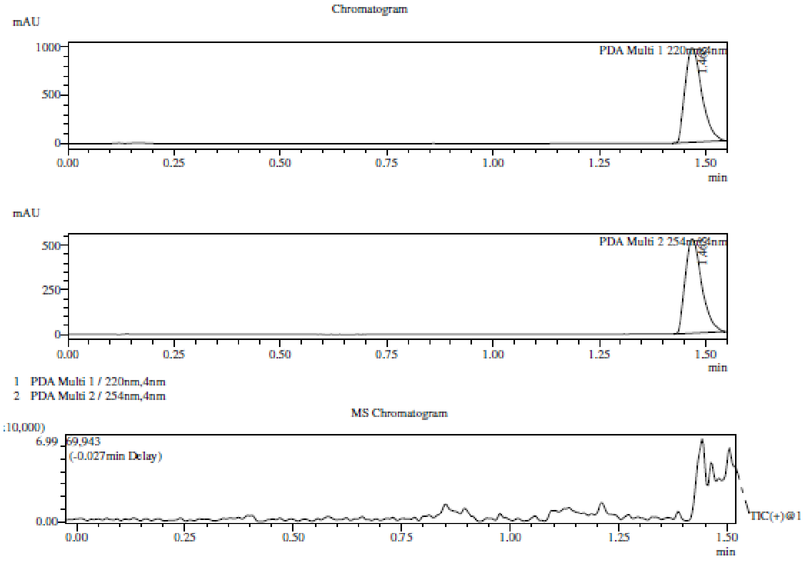

Integration Result

Peak Table

PDA Ch1 220nm

| Peak# | Ret. Time | Height | Height% | USP Width | Area | Area% |
|---|---|---|---|---|---|---|
| 1 | 1.468 | 973077 | 100.000 | 0.072 | 2664939 | 100.000 |

Peak Table

PDA Ch2 254nm

| Peak# | Ret. Time | Height | Height% | USP Width | Area | Area% |
|---|---|---|---|---|---|---|
| 1 | 1.468 | 525919 | 100.000 | 0.071 | 1419429 | 100.000 |

Mass Spectrum

RetTime: 1.470 Datafile: D:\DATA\2022\2201\220121\EW26049-720-P1C1.lcd

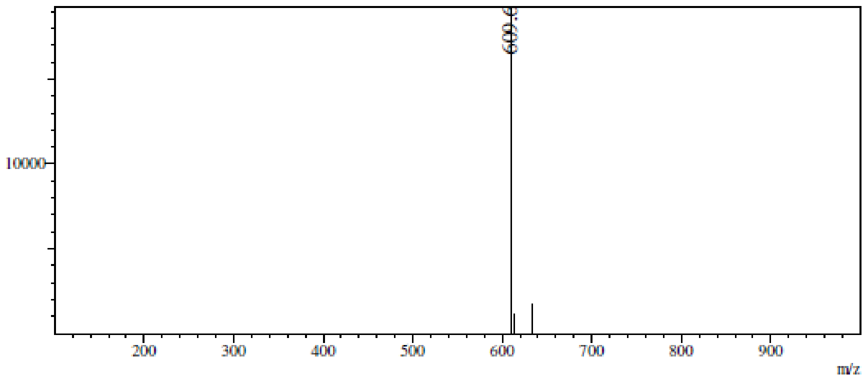

**$^1$H NMR** [(2*R*,3*R*,4*R*,5*R*)-5-(4-Aminopyrrolo[2,3-d]pyrimidin-7-yl)-3,4-bis[[tert-butyl(dimethyl)silyl]oxy]tetrahydrofuran-2-yl]methanol **(104)**

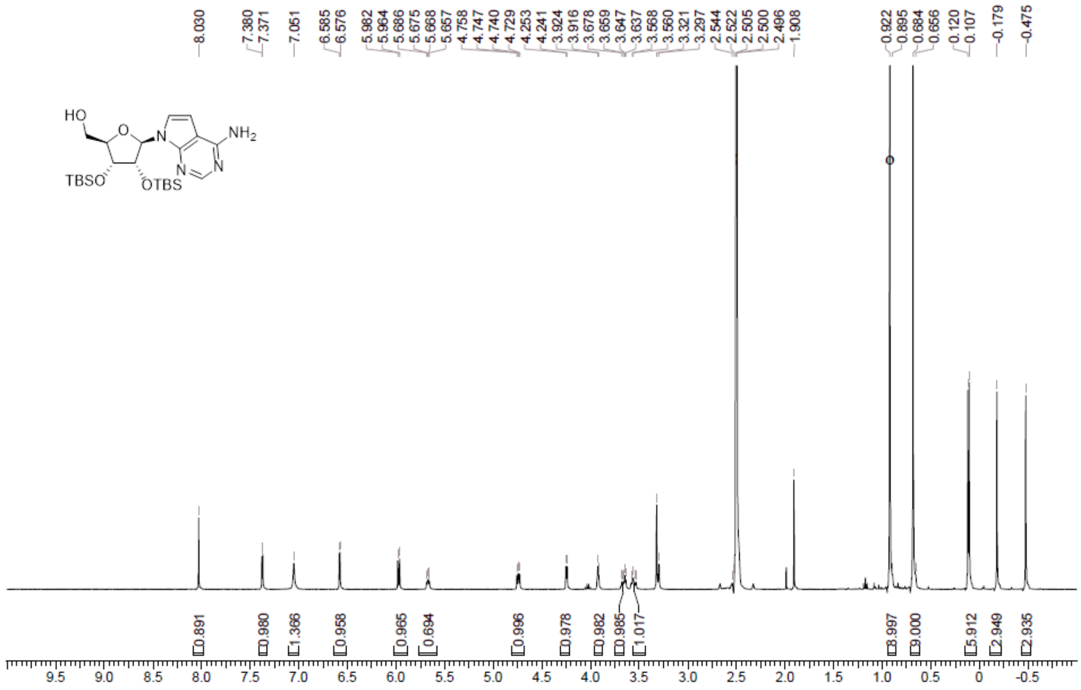

**LCMS** [(2*R*,3*R*,4*R*,5*R*)-5-(4-Aminopyrrolo[2,3-d]pyrimidin-7-yl)-3,4-bis[[tert-butyl(dimethyl)silyl]oxy]tetrahydrofuran-2-yl]methanol **(104)**

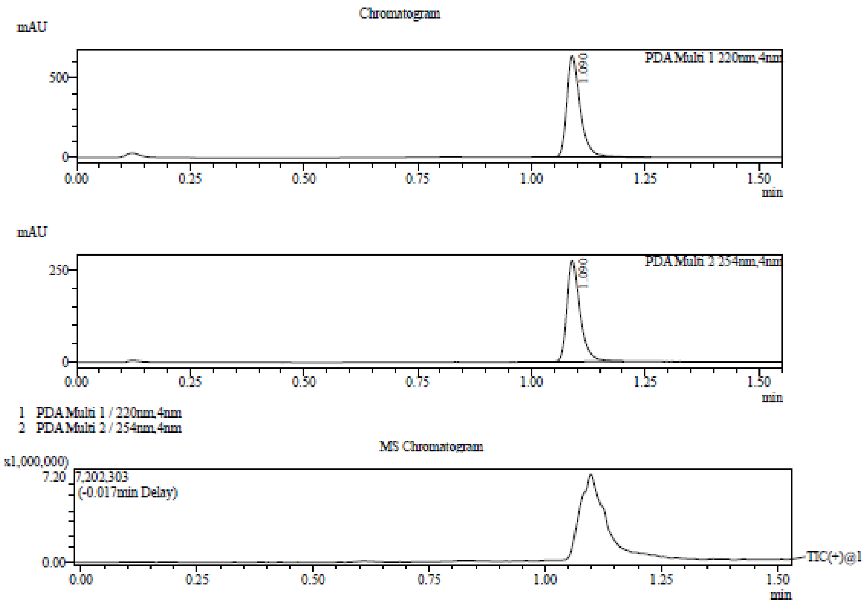

Integration Result

Peak Table

PDA Ch1 220nm

| Peak# | Ret. Time | Height | Height% | USP Width | Area | Area% |
|---|---|---|---|---|---|---|
| 1 | 1.090 | 633282 | 100.000 | 0.051 | 1284214 | 100.000 |

Peak Table

PDA Ch2 254nm

| Peak# | Ret. Time | Height | Height% | USP Width | Area | Area% |
|---|---|---|---|---|---|---|
| 1 | 1.090 | 276360 | 100.000 | 0.051 | 546944 | 100.000 |

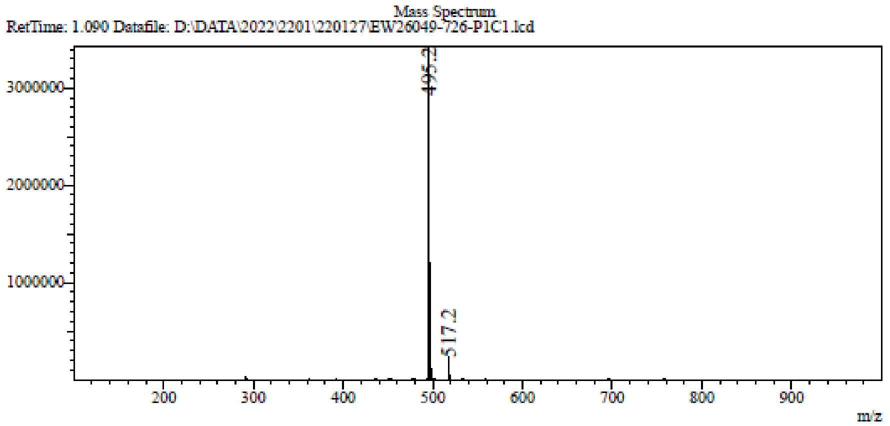

**[1]H NMR** ((2*R*,3*R*,4*R*,5*R*)-5-(4-Amino-7H-pyrrolo[2,3-d]pyrimidin-7-yl)-3,4-bis((tertbutyldimethylsilyl)oxy)tetrahydrofuran-2-yl)methyl methanesulfonate **(105)**

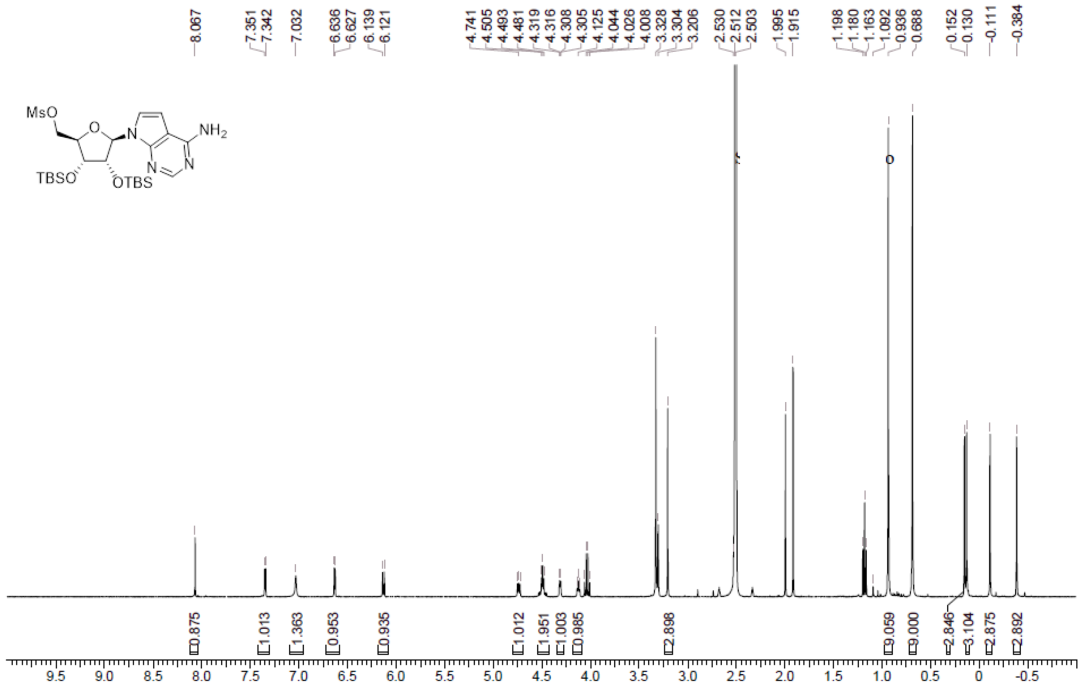

**LCMS** ((2*R*,3*R*,4*R*,5*R*)-5-(4-Amino-7H-pyrrolo[2,3-d]pyrimidin-7-yl)-3,4-bis((tertbutyldimethylsilyl)oxy)tetrahydrofuran-2-yl)methyl methanesulfonate **(105)**

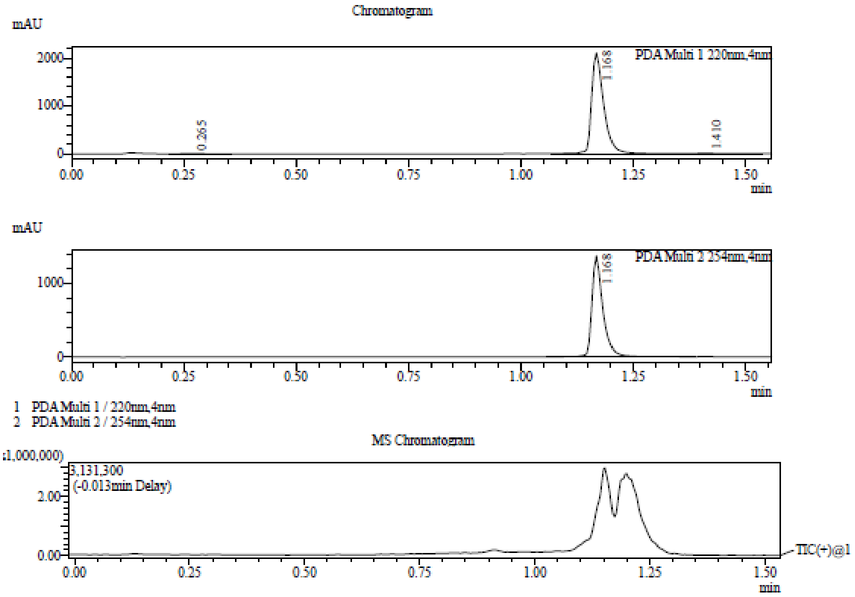

Integration Result

Peak Table

PDA Ch1 220nm

| Peak# | Ret. Time | Height | Height% | USP Width | Area | Area% |
|---|---|---|---|---|---|---|
| 1 | 0.265 | 1812 | 0.085 | 0.109 | 6978 | 0.174 |
| 2 | 1.168 | 2107121 | 99.326 | 0.055 | 3978645 | 99.239 |
| 3 | 1.410 | 12477 | 0.588 | 0.054 | 23546 | 0.587 |

Peak Table

PDA Ch2 254nm

| Peak# | Ret. Time | Height | Height% | USP Width | Area | Area% |
|---|---|---|---|---|---|---|
| 1 | 1.168 | 1375637 | 100.000 | 0.049 | 2339367 | 100.000 |

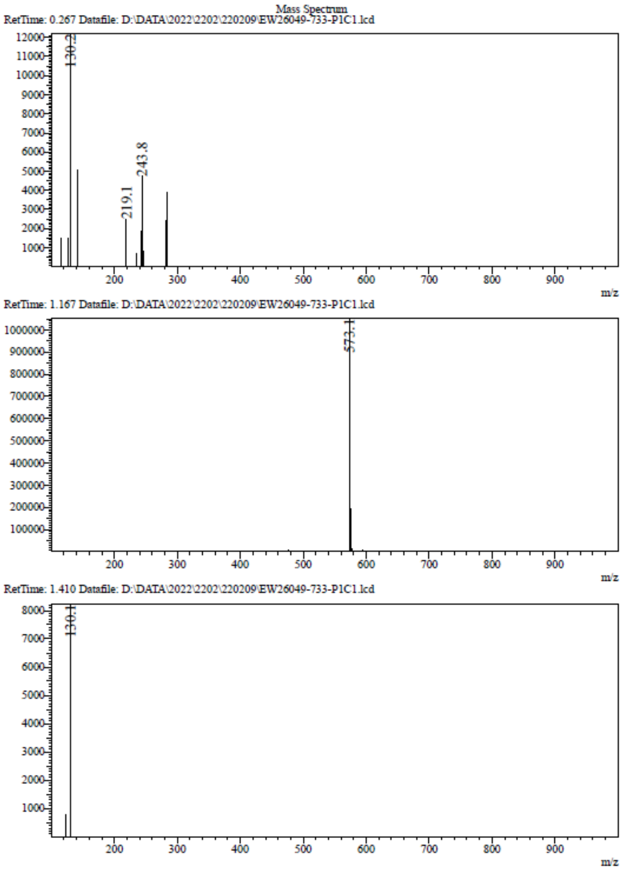

**[1]H NMR** 7-[(2*R*,3*R*,4*R*,5*S*)-5-(1,3-Benzothiazol-2-ylsulfanylmethyl)-3,4-bis [[tert-butyl(dimethyl) silyl]oxy]tetrahydrofuran-2-yl]pyrrolo[2,3-d]pyrimidin-4-amine **(106)**

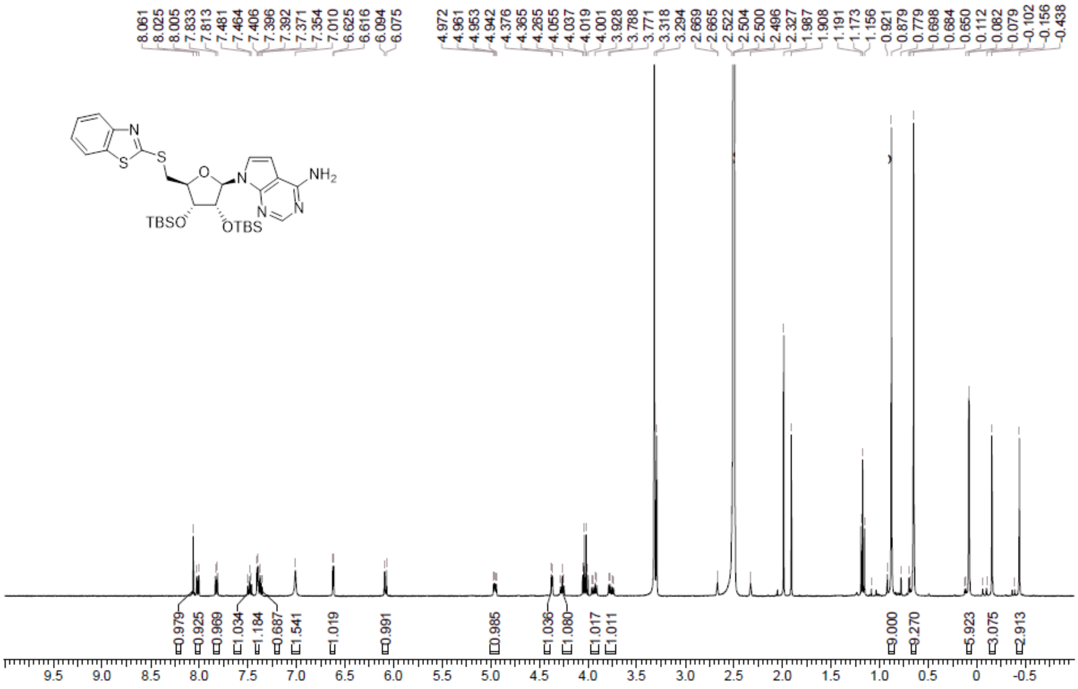

**LCMS** 7-[(2*R*,3*R*,4*R*,5*S*)-5-(1,3-Benzothiazol-2-ylsulfanylmethyl)-3,4-bis [[tert-butyl(dimethyl) silyl]oxy]tetrahydrofuran-2-yl]pyrrolo[2,3-d]pyrimidin-4-amine **(106)**

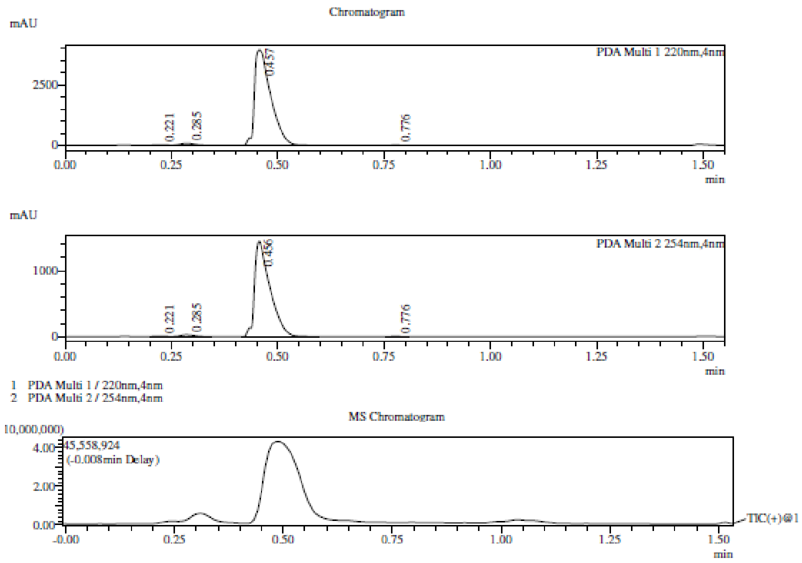

Integration Result

Peak Table

PDA Ch1 220nm

| Peak# | Ret. Time | Height | Height% | USP Width | Area | Area% |
|---|---|---|---|---|---|---|
| 1 | 0.221 | 17634 | 0.437 | 0.046 | 29197 | 0.275 |
| 2 | 0.285 | 75571 | 1.871 | 0.061 | 169299 | 1.596 |
| 3 | 0.457 | 3934164 | 97.394 | 0.063 | 10396308 | 97.986 |
| 4 | 0.776 | 12078 | 0.299 | 0.035 | 15204 | 0.143 |

Peak Table

PDA Ch2 254nm

| Peak# | Ret. Time | Height | Height% | USP Width | Area | Area% |
|---|---|---|---|---|---|---|
| 1 | 0.221 | 5615 | 0.380 | 0.043 | 8592 | 0.233 |
| 2 | 0.285 | 24489 | 1.657 | 0.060 | 53694 | 1.458 |
| 3 | 0.456 | 1442377 | 97.588 | 0.065 | 3612922 | 98.119 |
| 4 | 0.776 | 5546 | 0.375 | 0.035 | 6969 | 0.189 |

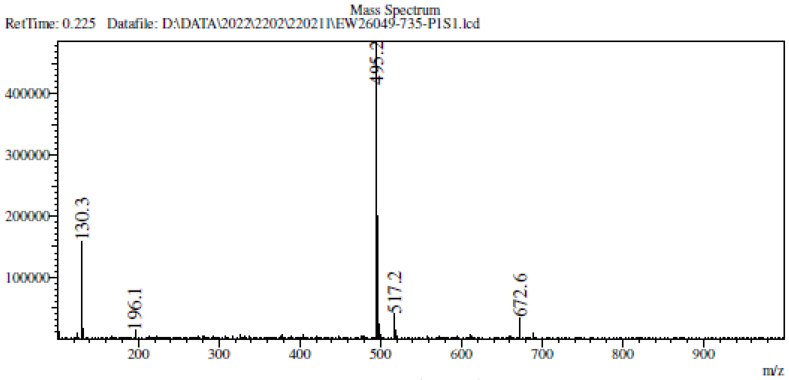
Mass Spectrum
RetTime: 0.225 Datafile: D:\DATA\2022\2202\220211\EW26049-735-P1S1.lcd
495.2
130.3
196.1
517.2
672.6
m/z

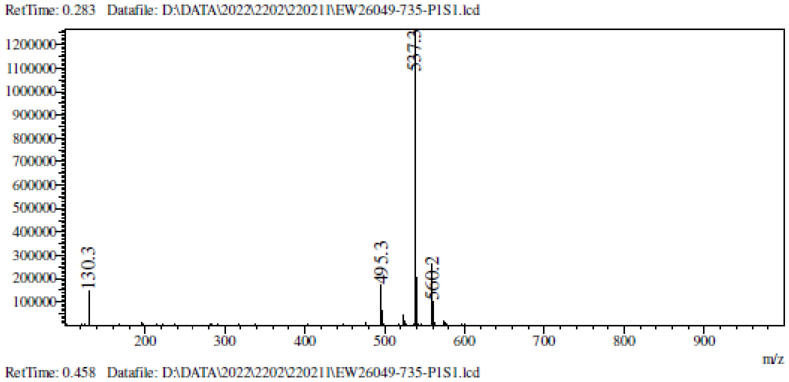
RetTime: 0.283 Datafile: D:\DATA\2022\2202\220211\EW26049-735-P1S1.lcd
537.3
130.3
495.3
560.2
m/z

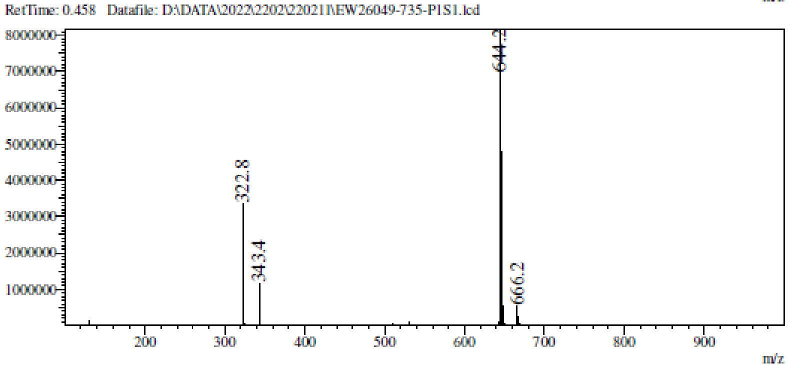
RetTime: 0.458 Datafile: D:\DATA\2022\2202\220211\EW26049-735-P1S1.lcd
644.2
322.8
343.4
666.2
m/z

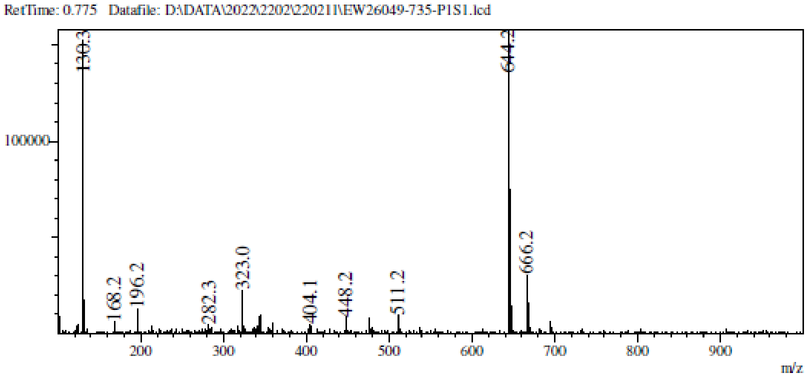
RetTime: 0.775 Datafile: D:\DATA\2022\2202\220211\EW26049-735-P1S1.lcd
130.3
644.2
168.2
196.2
282.3
323.0
404.1
448.2
511.2
666.2
m/z

[1] **H NMR** (2*R*,3*R*,4*S*,5*S*)-2-(4-Aminopyrrolo[2,3-d]pyrimidin-7-yl)-5-(1,3-benzothiazol-2-ylsulfanylmethyl)tetrahydrofuran-3,4-diol **(27)**

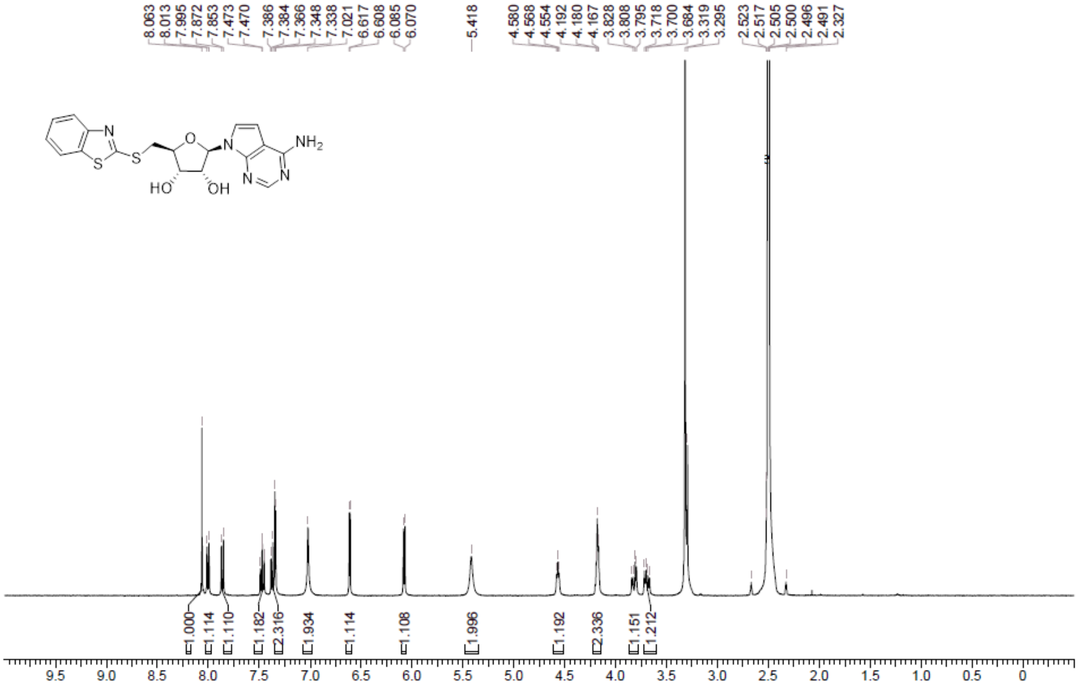

**HPLC** (2*R*,3*R*,4*S*,5*S*)-2-(4-Aminopyrrolo[2,3-d]pyrimidin-7-yl)-5-(1,3-benzothiazol-2-ylsulfanylmethyl)tetrahydrofuran-3,4-diol **(27)**

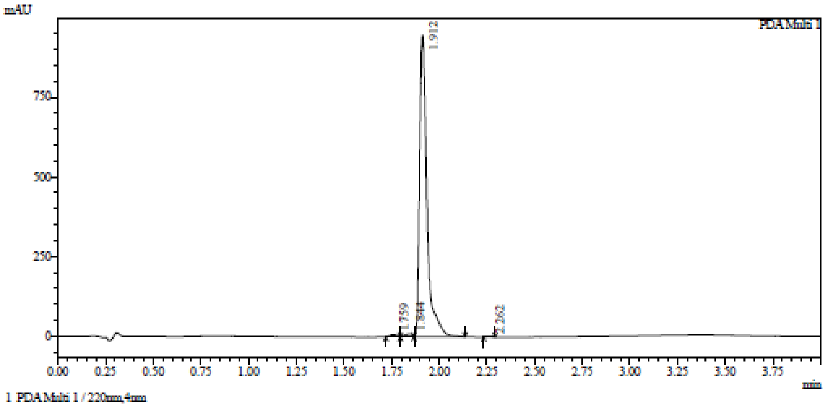

Integration result

PeakTable

PDA Ch1 220nm

| Peak# | Ret. Time | USP Width | Resolution | Height | Area | Area % |
|---|---|---|---|---|---|---|
| 1 | 1.759 | 0.081 | 0.000 | 7310 | 20055 | 0.779 |
| 2 | 1.844 | 0.131 | 0.805 | 9365 | 29156 | 1.133 |
| 3 | 1.912 | 0.065 | 0.695 | 946992 | 2521889 | 98.012 |
| 4 | 2.262 | 0.056 | 5.755 | 914 | 1933 | 0.075 |
| Total | | | | 964581 | 2573033 | 100.000 |

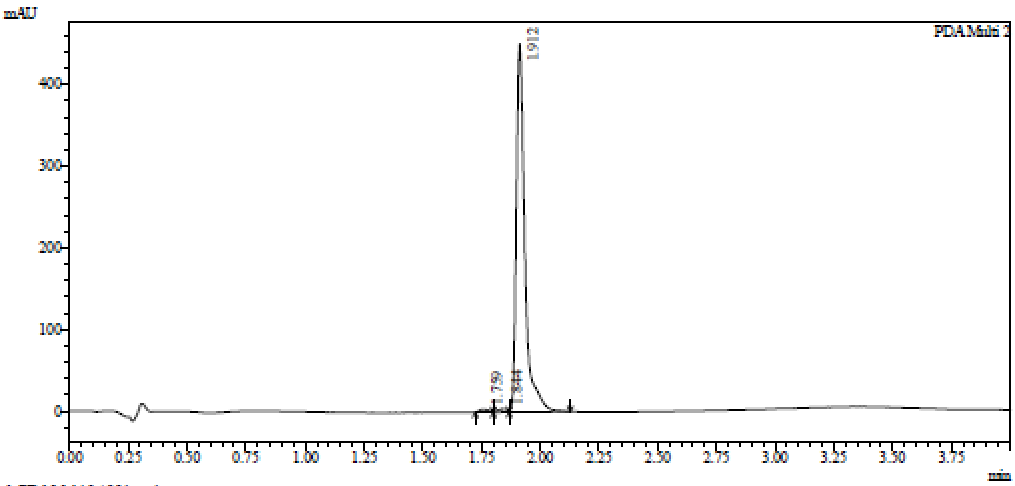

1 PDA Multi 2 / 254nm,4nm

Integration result

PeakTable

PDA Ch2 254nm

| Peak# | Ret. Time | USP Width | Resolution | Height | Area | Area % |
|---|---|---|---|---|---|---|
| 1 | 1.759 | 0.078 | 0.000 | 3714 | 10596 | 0.909 |
| 2 | 1.844 | 0.170 | 0.688 | 3882 | 11969 | 1.027 |
| 3 | 1.912 | 0.063 | 0.582 | 450566 | 1143339 | 98.065 |
| Total | | | | 458161 | 1165904 | 100.000 |

**HRMC** (2*R*,3*R*,4*S*,5*S*)-2-(4-Aminopyrrolo[2,3-d]pyrimidin-7-yl)-5-(1,3-benzothiazol-2-ylsulfanylmethyl)tetrahydrofuran-3,4-diol **(27)**

Compound Table

| Label | Tgt Score | Mass Error (ppm) | Tgt Formula | Obs. RT | Ref. Mass | Obs. Mass |
|---|---|---|---|---|---|---|
| Cpd 1: C18 H17 N5 O3 S2; 1.763 | 97.21 | 0.75 | C18 H17 N5 O3 S2 | 1.763 | 415.0773 | 415.0776 |

| Obs. *m/z* | Obs. RT | Obs. Mass | Tgt Formula | Tgt Mass | Tgt Mass Error (ppm) | Find Cpds Algorithm |
|---|---|---|---|---|---|---|
| 438.0658 | 1.763 | 415.0776 | C18 H17 N5 O3 S2 | 415.0773 | 0.75 | Find by Formula |

Compound Chromatograms

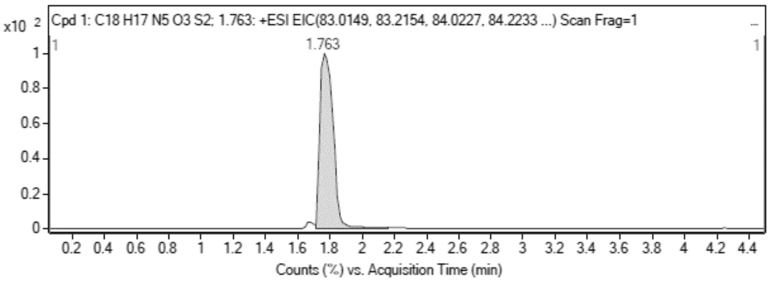

MS Zoomed Spectrum

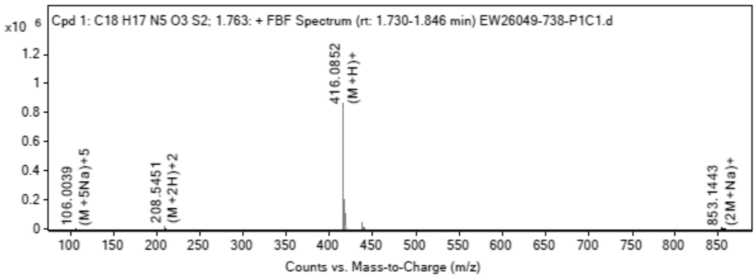

MS Spectrum Peak List

| Obs. *m/z* | Charge | Abund | Ion/Isotope |
|---|---|---|---|
| 208.5451 | 2 | 22678.21 | (M+2H)+2 |
| 209.0469 | 2 | 5371.39 | (M+2H)+2 |

| | | | |
|---|---|---|---|
| 209.5444 | 2 | 2506.16 | (M+2H)+2 |
| 416.0852 | 1 | 863713.56 | (M+H)+ |
| 417.0869 | 1 | 200761.59 | (M+H)+ |
| 418.0825 | 1 | 99680.85 | (M+H)+ |
| 438.0658 | 1 | 42955.25 | (M+Na)+ |
| 439.0681 | 1 | 9944.64 | (M+Na)+ |
| 440.0644 | 1 | 5401.75 | (M+Na)+ |
| 853.1443 | 1 | 4463.23 | (2M+Na)+ |

MS Zoomed Spectrum

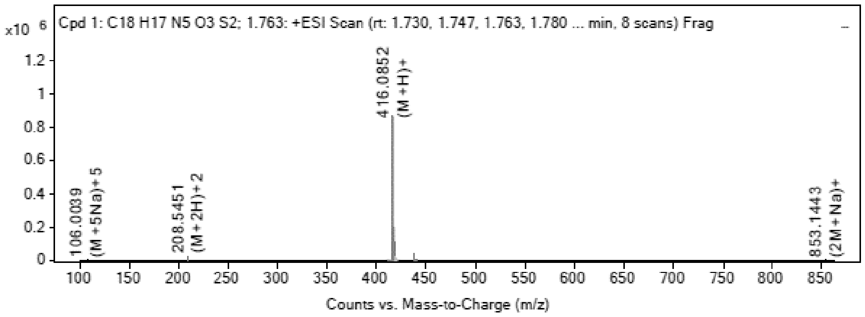

MS Spectrum Peak List

| Obs. m/z | Charge | Abund | Ion/Isotope | Tgt Mass Error (ppm) |
|---|---|---|---|---|
| 208.5451 | 2 | 22678.21 | (M+2H)+2 | 3.85 |
| 209.0469 | 2 | 5371.39 | (M+2H)+2 | 1.74 |
| 209.5444 | 2 | 2506.16 | (M+2H)+2 | 2.86 |
| 416.0852 | 1 | 863713.56 | (M+H)+ | -1.65 |
| 417.0869 | 1 | 200761.59 | (M+H)+ | 0.7 |
| 418.0825 | 1 | 99680.85 | (M+H)+ | 0.67 |
| 438.0658 | 1 | 42955.25 | (M+Na)+ | 1.51 |
| 439.0681 | 1 | 9944.64 | (M+Na)+ | 2.21 |
| 440.0644 | 1 | 5401.75 | (M+Na)+ | 0.72 |
| 853.1443 | 1 | 4463.23 | (2M+Na)+ | -0.62 |

--- End Of Report ---

**$^1$H NMR** (2R,3R,4R,5R)-2-((Benzoyloxy)methyl)-5-(4-chloro-5-iodo-7H-pyrrolo[2,3-d]pyrimidin-7-yl)tetrahydrofuran-3,4-diyl dibenzoate **(64)**

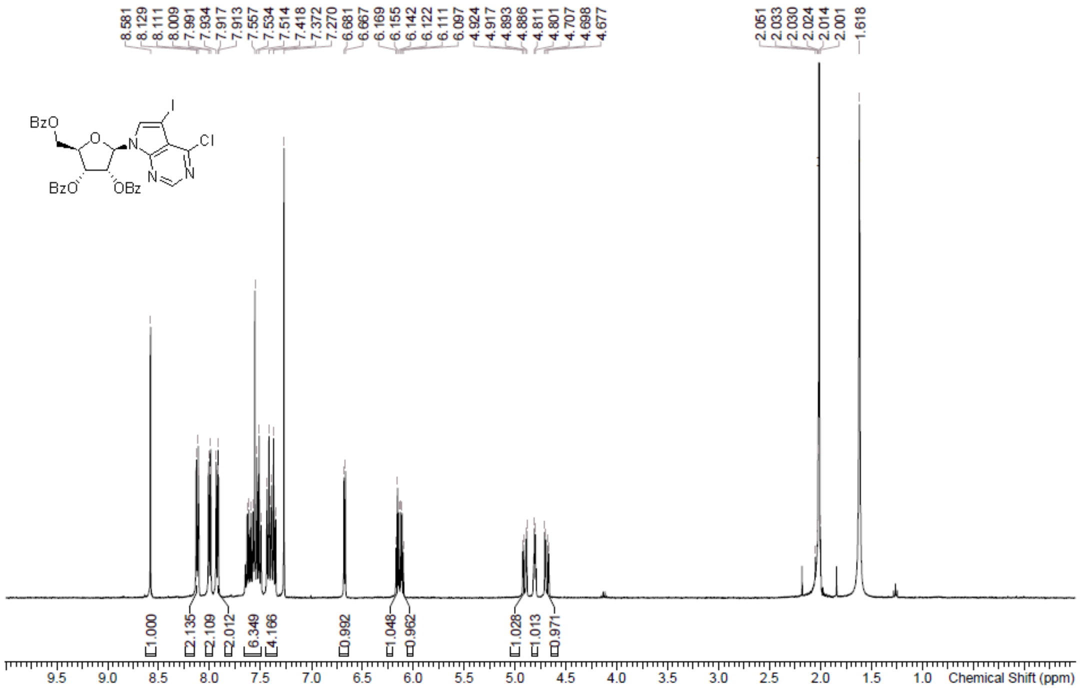

**$^{1}$H NMR** (2R,3R,4S,5R)-2-(4-Amino-5-iodo-7H-pyrrolo[2,3-d]pyrimidin-7-yl)-5-(hydroxymethyl)tetrahydrofuran-3,4-diol **(65)**

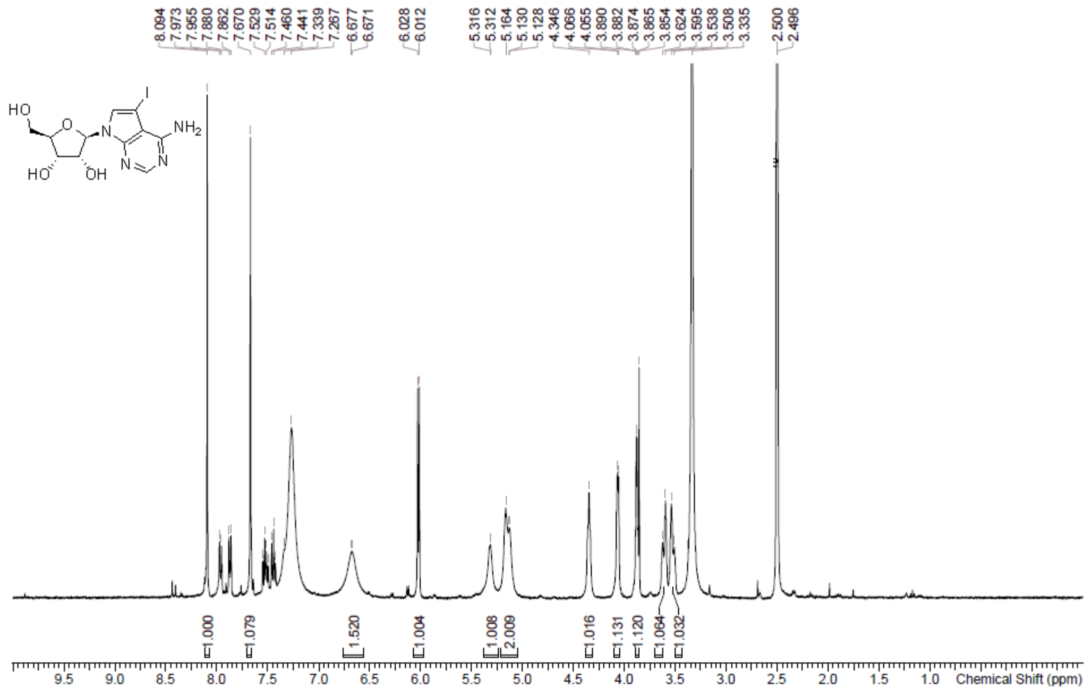

**$^{1}$H NMR** 7-((2R,3R,4R,5R)-3,4-Bis((tert-butyldimethylsilyl)oxy)-5-(((tert-butyldimethylsilyl)oxy)methyl)tetrahydrofuran-2-yl)-5-iodo-7H-pyrrolo[2,3-d]pyrimidin-4-amine **(66)**

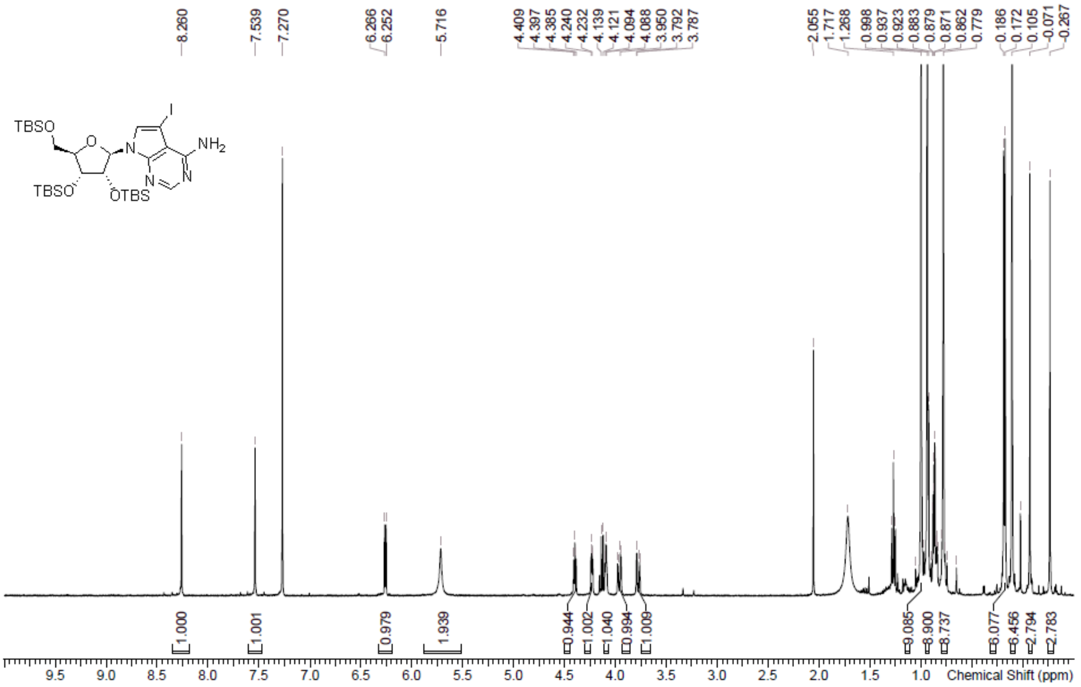

**LCMS** 7-((2R,3R,4R,5R)-3,4-Bis((tert-butyldimethylsilyl)oxy)-5-(((tert-butyldimethylsilyl)oxy)methyl)tetrahydrofuran-2-yl)-5-iodo-7H-pyrrolo[2,3-d]pyrimidin-4-amine **(66)**

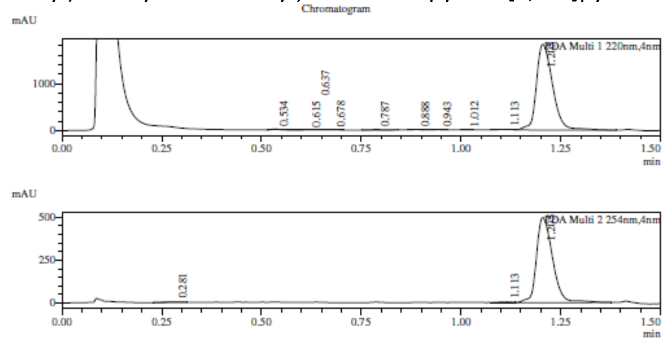

1 PDA Multi 1 / 220nm,4nm
2 PDA Multi 2 / 254nm,4nm

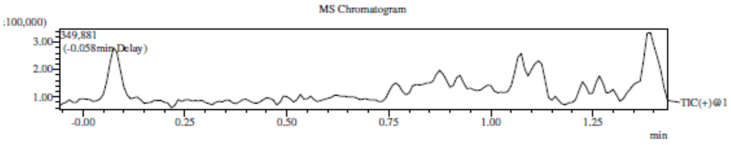

Integration Result

Peak Table

PDA Ch1 220nm

| Peak# | Ret. Time | Height | Height% | USP Width | Area | Area% |
|---|---|---|---|---|---|---|
| 1 | 0.534 | 20708 | 1.089 | 0.038 | 29401 | 0.558 |
| 2 | 0.615 | 6303 | 0.332 | 0.043 | 7490 | 0.142 |
| 3 | 0.637 | 5420 | 0.285 | 0.044 | 5545 | 0.105 |
| 4 | 0.678 | 6402 | 0.337 | 0.043 | 10172 | 0.193 |
| 5 | 0.787 | 10380 | 0.546 | 0.048 | 20047 | 0.380 |
| 6 | 0.888 | 9099 | 0.479 | 0.038 | 13361 | 0.253 |
| 7 | 0.943 | 6160 | 0.324 | 0.037 | 7887 | 0.150 |
| 8 | 1.012 | 3665 | 0.193 | 0.056 | 5244 | 0.099 |
| 9 | 1.113 | 9734 | 0.512 | 0.086 | 24637 | 0.467 |
| 10 | 1.205 | 1823506 | 95.905 | 0.072 | 5149420 | 97.653 |

Peak Table

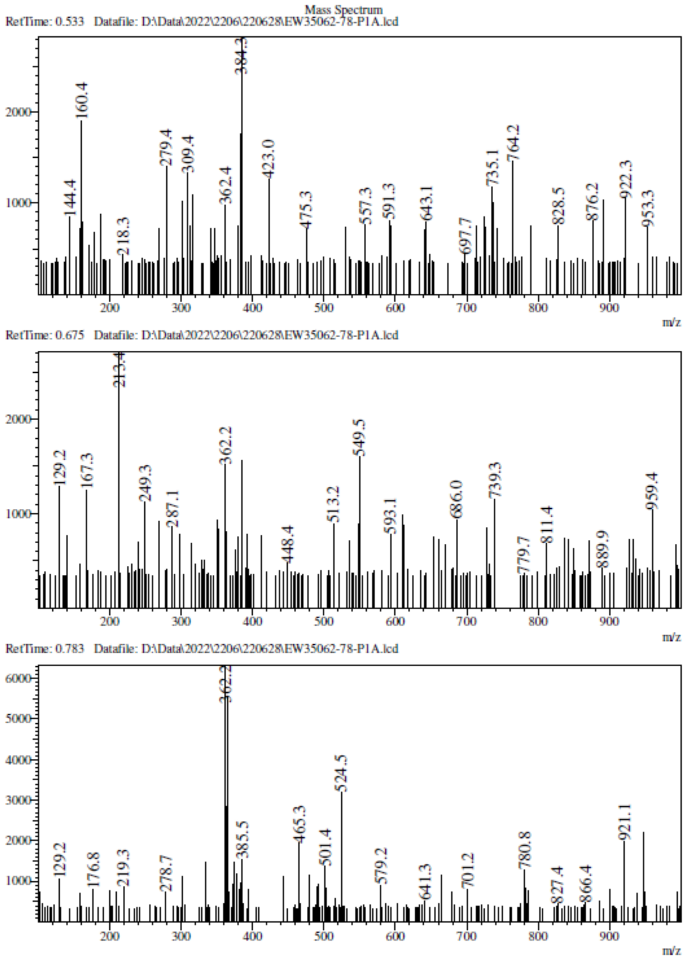

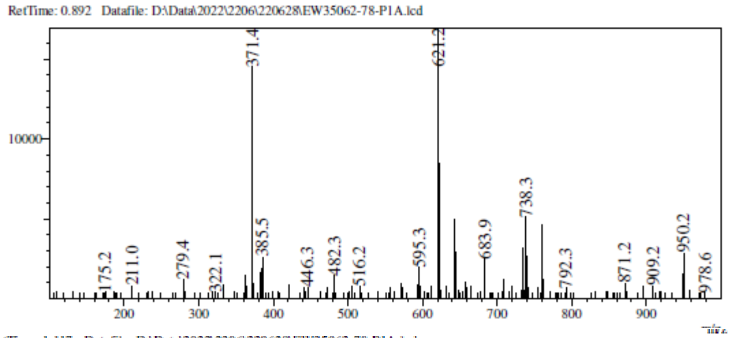

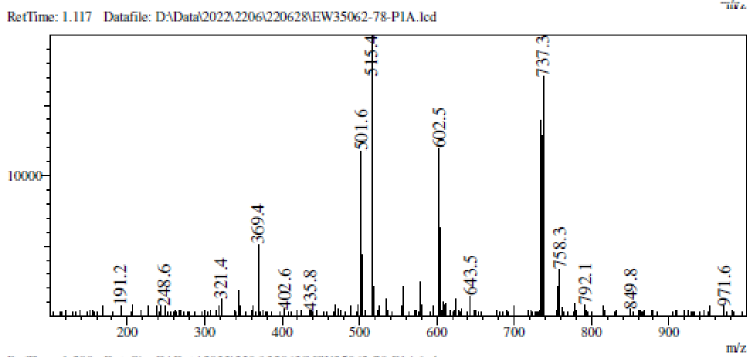

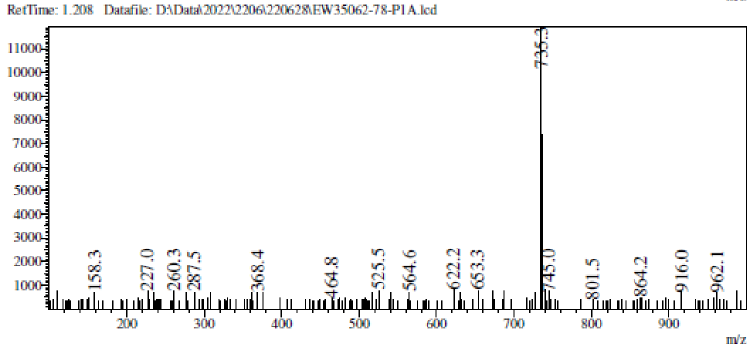

**$^1$H NMR** 1-(5-Benzylsulfanyl-6-fluoro-indazol-1-yl)ethanone **(67)**

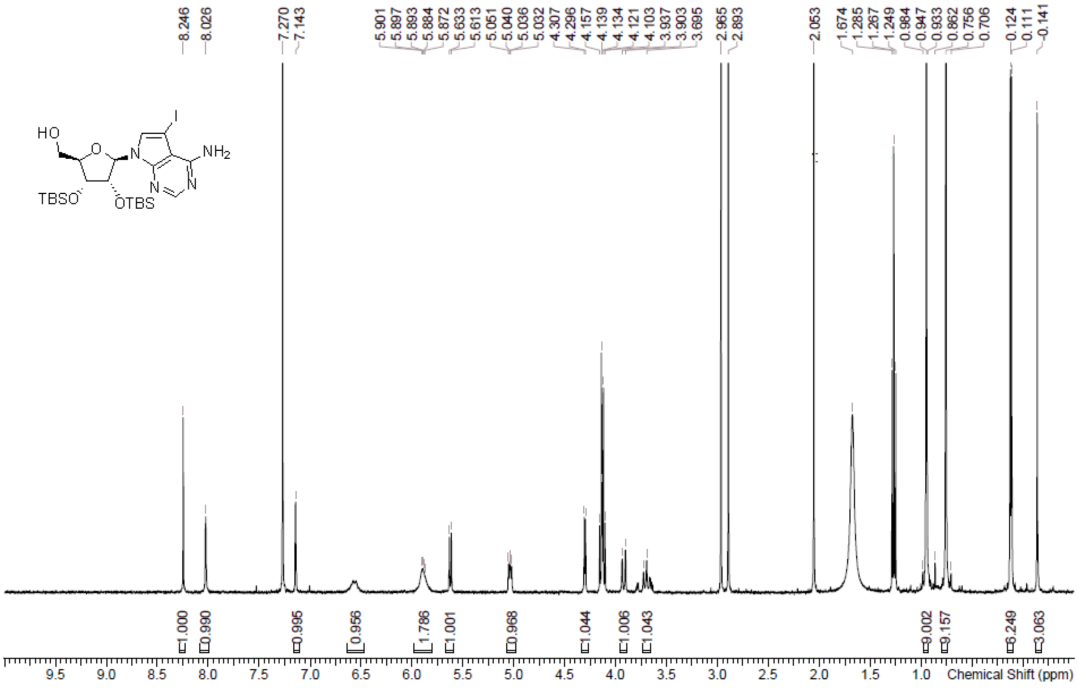

**LCMS** 1-(5-Benzylsulfanyl-6-fluoro-indazol-1-yl)ethanone **(67)**

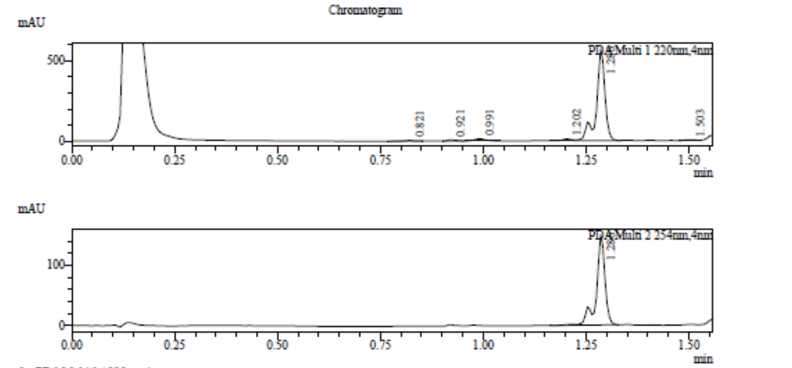

1 PDA Multi 1 / 220nm,4nm
2 PDA Multi 2 / 254nm,4nm

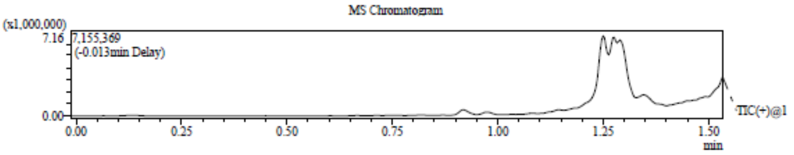

Integration Result

Peak Table

PDA Ch1 220nm

| Peak# | Ret. Time | Height | Height% | USP Width | Area | Area% |
|---|---|---|---|---|---|---|
| 1 | 0.821 | 5537 | 0.902 | 0.027 | 5970 | 0.672 |
| 2 | 0.921 | 7829 | 1.276 | 0.036 | 10278 | 1.157 |
| 3 | 0.991 | 13491 | 2.198 | 0.057 | 23874 | 2.687 |
| 4 | 1.202 | 10525 | 1.715 | 0.048 | 16023 | 1.803 |
| 5 | 1.286 | 573201 | 93.395 | 0.037 | 827403 | 93.115 |
| 6 | 1.503 | 3154 | 0.514 | 0.043 | 5034 | 0.566 |

Peak Table

PDA Ch2 254nm

| Peak# | Ret. Time | Height | Height% | USP Width | Area | Area% |
|---|---|---|---|---|---|---|
| 1 | 1.286 | 150263 | 100.000 | 0.037 | 219703 | 100.000 |

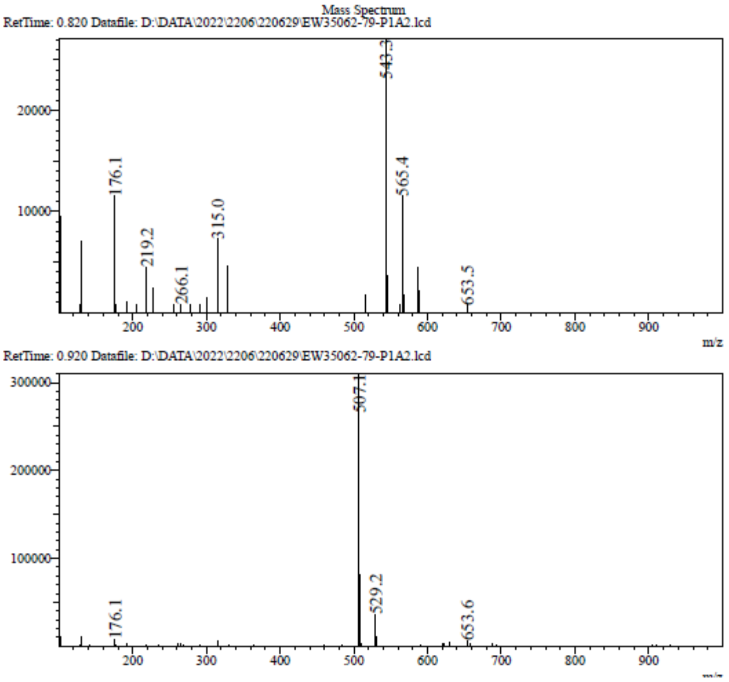

RetTime: 0.990 Datafile: D:\DATA\2022\2206\220629\EW35062-79-P1A2.lcd

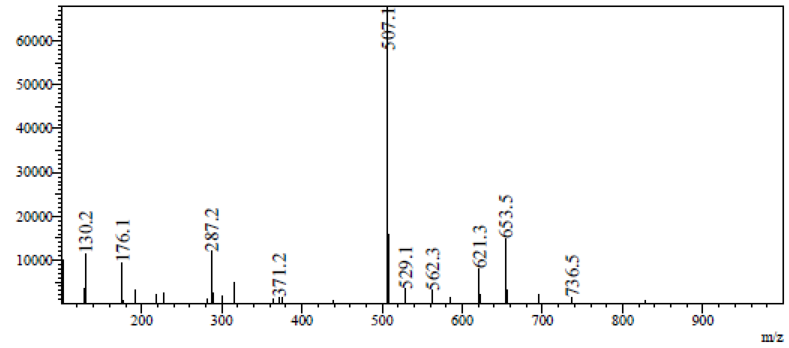

RetTime: 1.203 Datafile: D:\DATA\2022\2206\220629\EW35062-79-P1A2.lcd

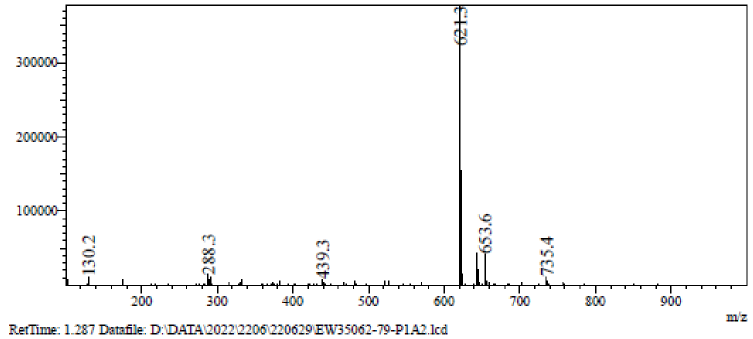

RetTime: 1.287 Datafile: D:\DATA\2022\2206\220629\EW35062-79-P1A2.lcd

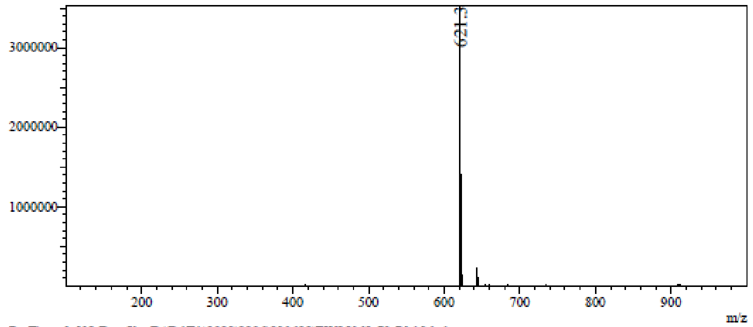

RetTime: 1.503 Datafile: D:\DATA\2022\2206\220629\EW35062-79-P1A2.lcd

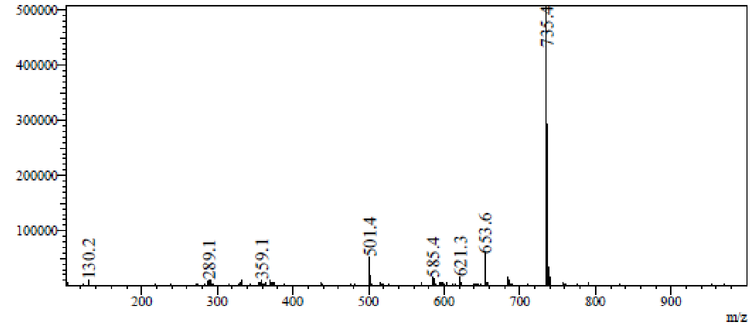

**[1]H NMR** ((2R,3R,4R,5R)-5-(4-Amino-5-iodo-7H-pyrrolo[2,3-d]pyrimidin-7-yl)-3,4-bis((tert-butyldimethylsilyl)oxy)tetrahydrofuran-2-yl)methyl methanesulfonate **(68)**

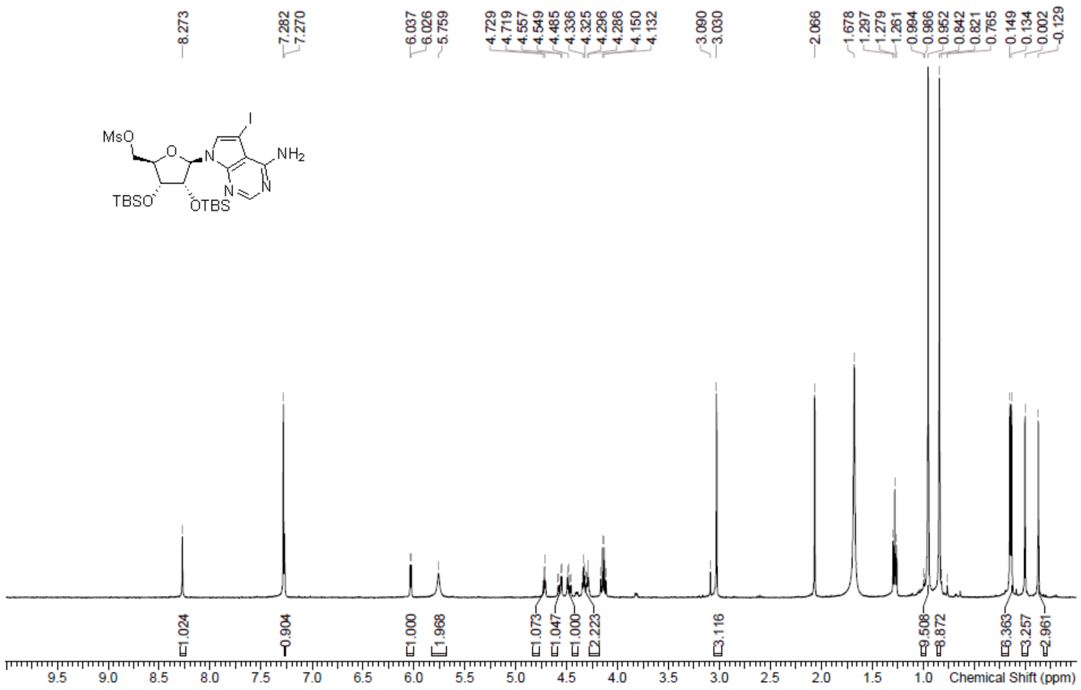

**LCMS** ((2R,3R,4R,5R)-5-(4-Amino-5-iodo-7H-pyrrolo[2,3-d]pyrimidin-7-yl)-3,4-bis((tert-butyldimethylsilyl)oxy)tetrahydrofuran-2-yl)methyl methanesulfonate **(68)**

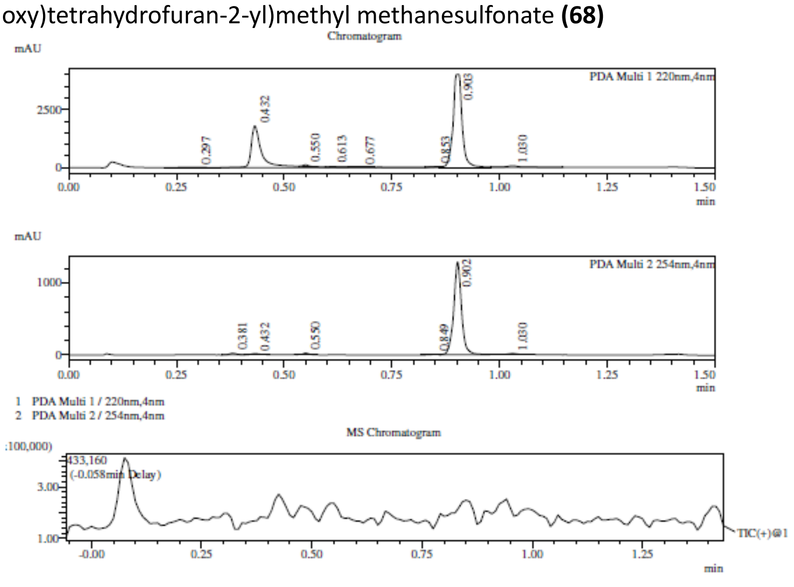

Integration Result

Peak Table

PDA Ch1 220nm

| Peak# | Ret. Time | Height | Height% | USP Width | Area | Area% |
|---|---|---|---|---|---|---|
| 1 | 0.297 | 11446 | 0.192 | 0.096 | 42147 | 0.439 |
| 2 | 0.432 | 1788641 | 30.038 | 0.037 | 3424349 | 35.690 |
| 3 | 0.550 | 73875 | 1.241 | 0.023 | 57594 | 0.600 |
| 4 | 0.613 | 9545 | 0.160 | 0.033 | 13063 | 0.136 |
| 5 | 0.677 | 7360 | 0.124 | 0.043 | 10555 | 0.110 |
| 6 | 0.853 | 5549 | 0.093 | 0.023 | 6445 | 0.067 |
| 7 | 0.903 | 3988438 | 66.981 | 0.034 | 5867584 | 61.155 |
| 8 | 1.030 | 69758 | 1.172 | 0.044 | 172835 | 1.801 |

Peak Table

PDA Ch2 254nm

Mass Spectrum

RetTime: 0.300 Datafile: D:\Data\2022\2207\220706\EW35062-82-P1D.lcd

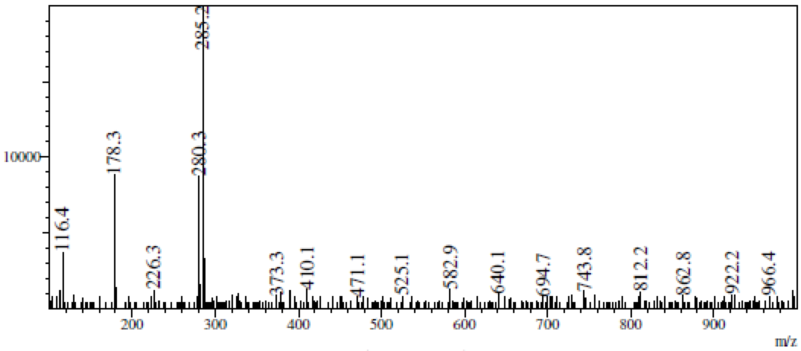

RetTime: 0.433 Datafile: D:\Data\2022\2207\220706\EW35062-82-P1D.lcd

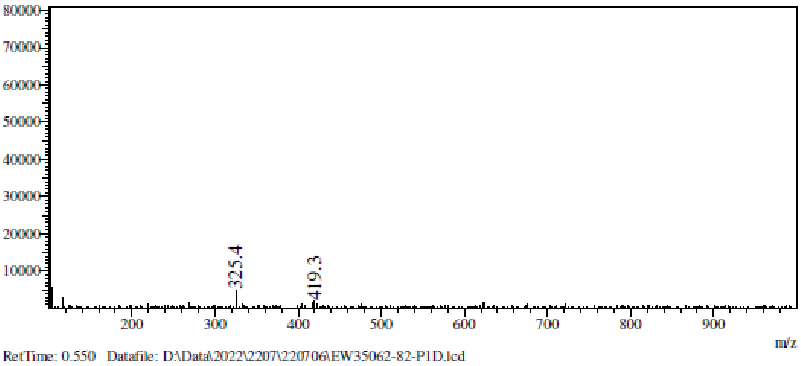

RetTime: 0.550 Datafile: D:\Data\2022\2207\220706\EW35062-82-P1D.lcd

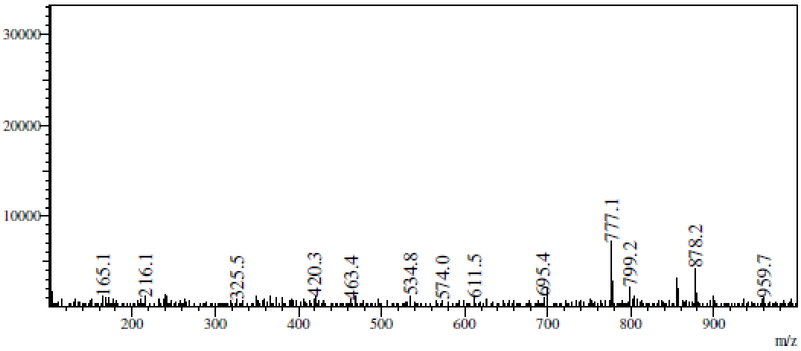

RetTime: 0.617 Datafile: D:\Data\2022\2207\220706\EW35062-82-P1D.lcd

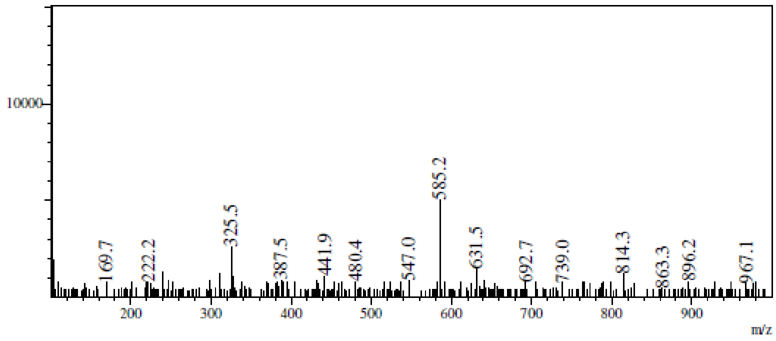

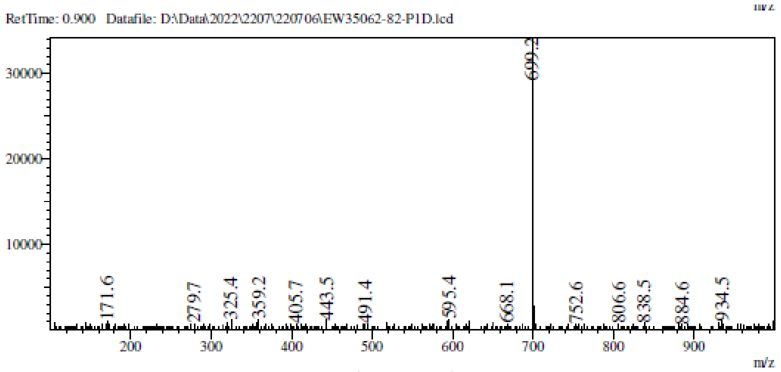

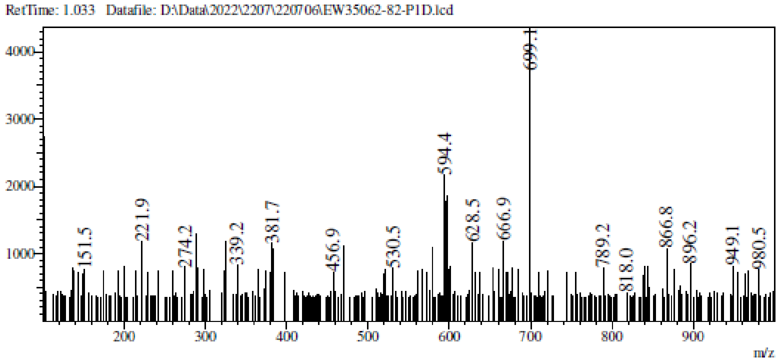

**$^1$H NMR** 7-((2R,3R,4R,5S)-5-((Benzo[d]thiazol-2-ylthio)methyl)-3,4-bis((tert-butyldimethylsilyl)oxy)tetrahydrofuran-2-yl)-5-iodo-7H-pyrrolo[2,3-d]pyrimidin-4-amine **(69)**

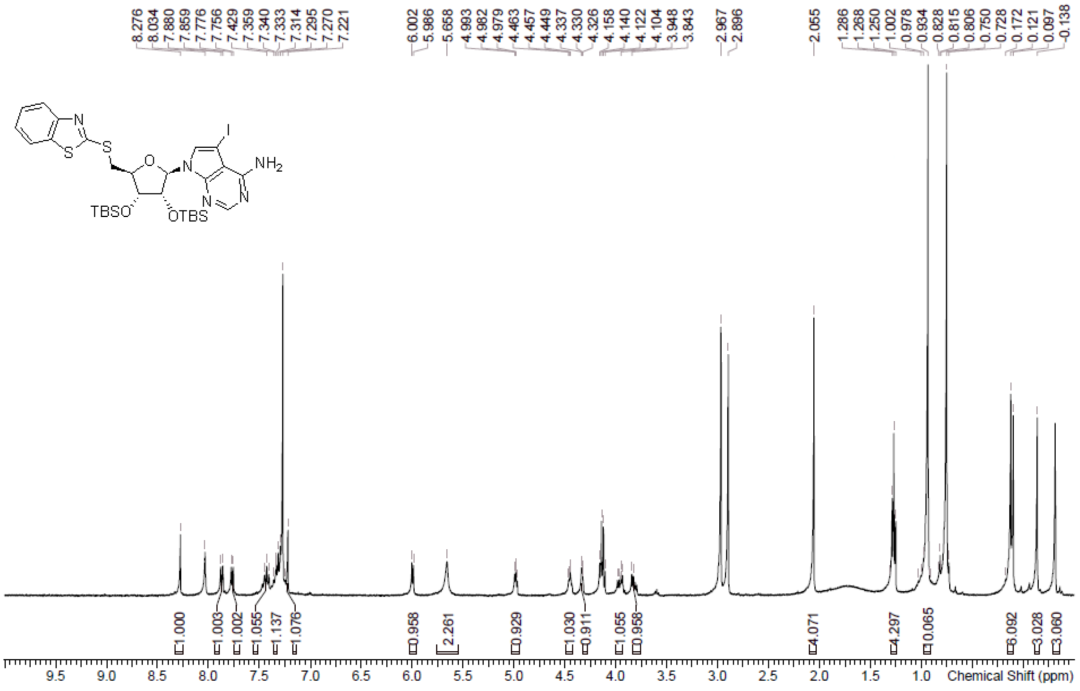

**LCMS** 7-((2R,3R,4R,5S)-5-((Benzo[d]thiazol-2-ylthio)methyl)-3,4-bis((tert-butyldimethylsilyl)oxy)tetrahydrofuran-2-yl)-5-iodo-7H-pyrrolo[2,3-d]pyrimidin-4-amine **(69)**

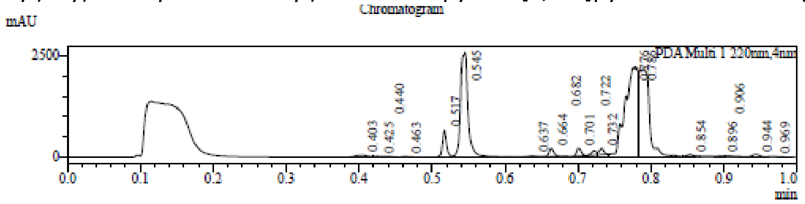

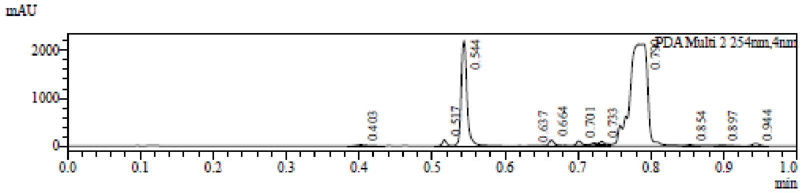

1 PDA Multi 1 / 220nm,4nm
2 PDA Multi 2 / 254nm,4nm

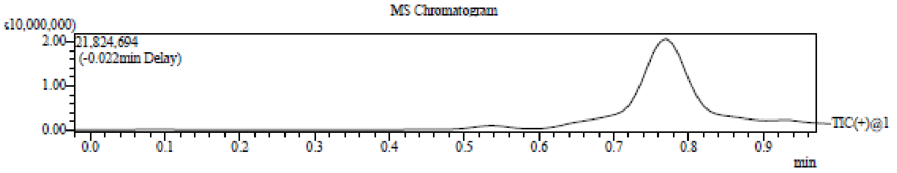

Integration Result

Peak Table

PDA Ch1 220nm

| Peak# | Ret. Time | Height | Height% | USP Width | Area | Area% |
|---|---|---|---|---|---|---|
| 1 | 0.403 | 49851 | 0.580 | 0.033 | 57767 | 0.749 |
| 2 | 0.425 | 21355 | 0.249 | 0.070 | 18263 | 0.237 |
| 3 | 0.440 | 19696 | 0.229 | 0.034 | 11777 | 0.153 |
| 4 | 0.463 | 16567 | 0.193 | 0.016 | 8902 | 0.115 |
| 5 | 0.517 | 669022 | 7.790 | 0.011 | 271116 | 3.517 |
| 6 | 0.545 | 2594224 | 30.206 | 0.018 | 2024636 | 26.266 |
| 7 | 0.637 | 23526 | 0.274 | 0.017 | 13073 | 0.170 |
| 8 | 0.664 | 203254 | 2.367 | 0.014 | 95825 | 1.243 |
| 9 | 0.682 | 17152 | 0.200 | 0.014 | 8167 | 0.106 |
| 10 | 0.701 | 193689 | 2.255 | 0.014 | 85915 | 1.115 |
| 11 | 0.722 | 105327 | 1.226 | 0.019 | 52300 | 0.679 |
| 12 | 0.732 | 157534 | 1.834 | 0.014 | 68516 | 0.889 |
| 13 | 0.776 | 2205382 | 25.678 | 0.078 | 2892441 | 37.525 |
| 14 | 0.786 | 2145964 | 24.986 | 0.063 | 1995864 | 25.893 |

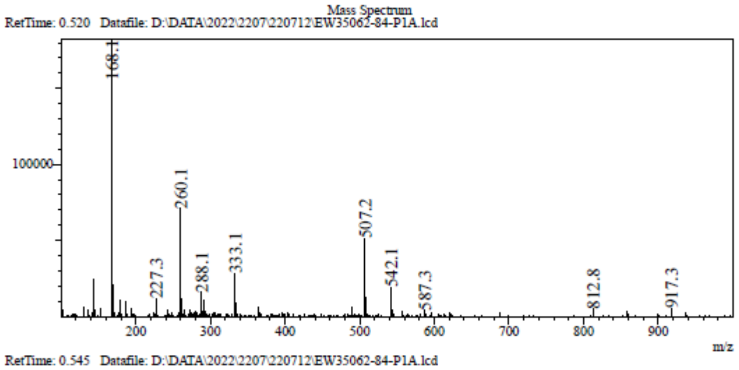

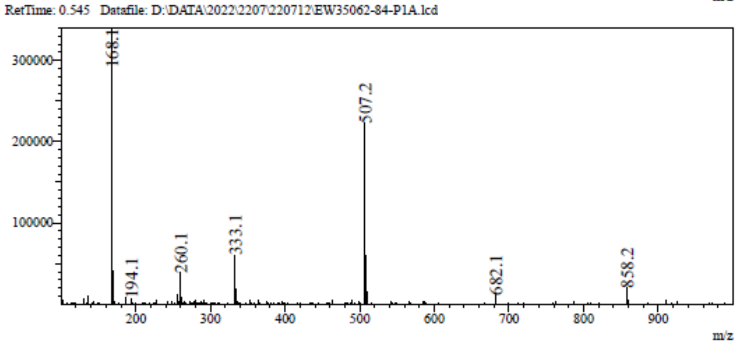

RetTime: 0.662 Datafile: D:\DATA\2022\2207\220712\EW35062-84-P1A.lcd

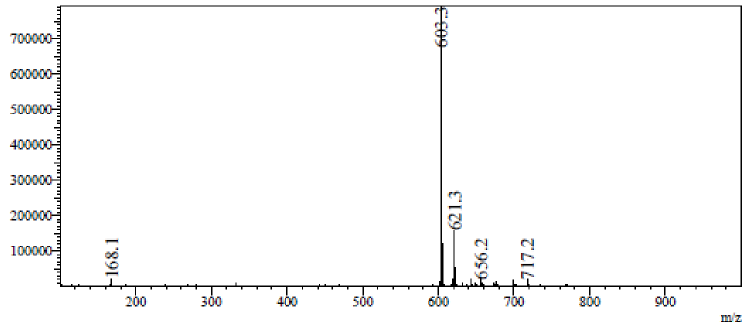

RetTime: 0.703 Datafile: D:\DATA\2022\2207\220712\EW35062-84-P1A.lcd

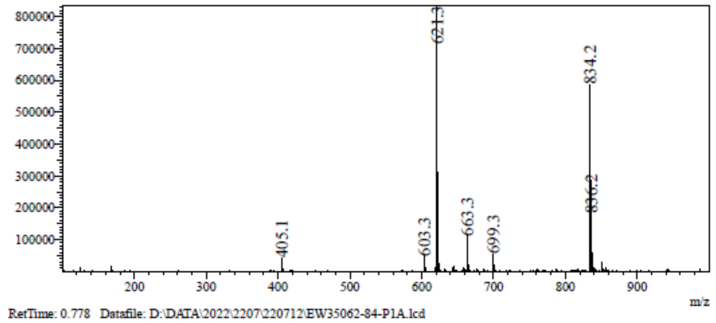

RetTime: 0.778 Datafile: D:\DATA\2022\2207\220712\EW35062-84-P1A.lcd

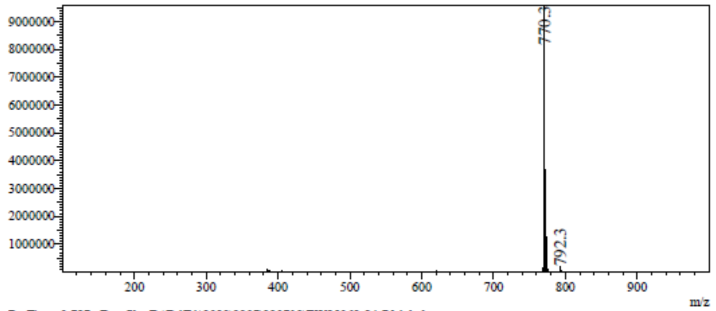

RetTime: 0.787 Datafile: D:\DATA\2022\2207\220712\EW35062-84-P1A.lcd

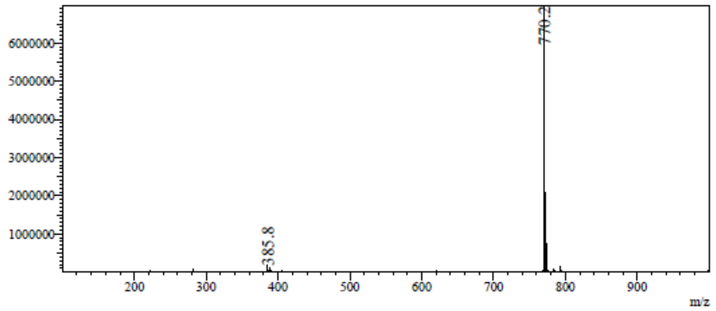

**$^{1}$H NMR** 4-Amino-7-[(2*R*,3*R*,4*R*,5*S*)-5-(1,3-benzothiazol-2-ylsulfanylmethyl)- 3,4-bis[[tert-butyl(dimethyl)silyl]oxy]tetrahydrofuran-2-yl]pyrrolo[2,3-d]pyrimidine-5-carbonitrile **(70)**

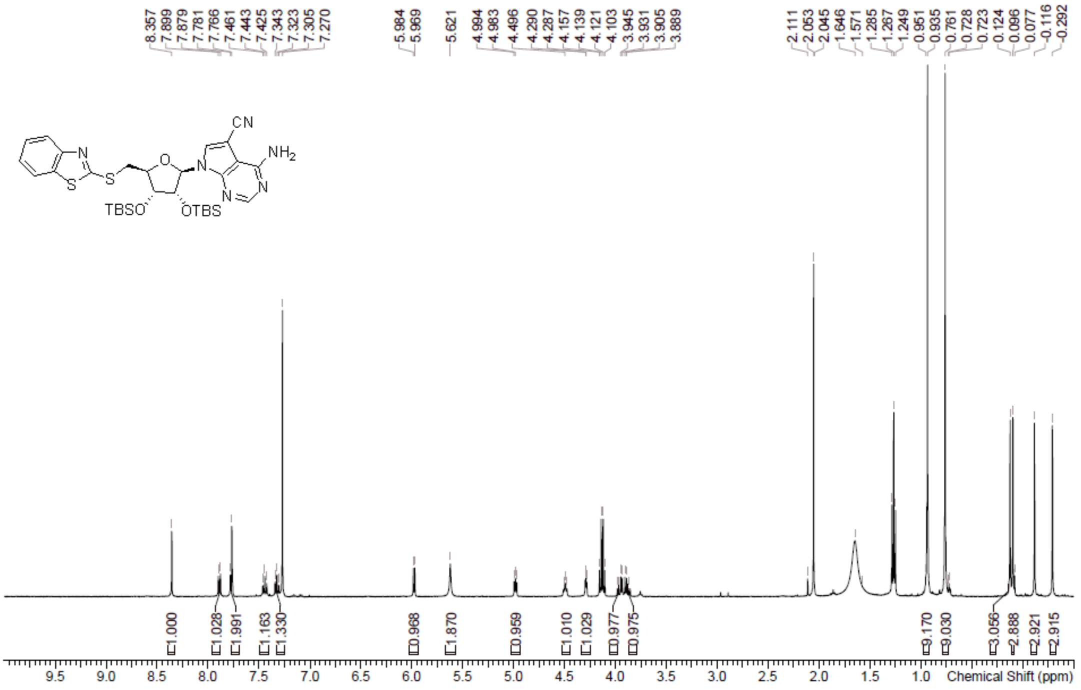

**LCMS** 4-Amino-7-[(2*R*,3*R*,4*R*,5*S*)-5-(1,3-benzothiazol-2-ylsulfanylmethyl)- 3,4-bis[[tert-butyl(dimethyl)silyl]oxy]tetrahydrofuran-2-yl]pyrrolo[2,3-d]pyrimidine-5-carbonitrile **(70)**

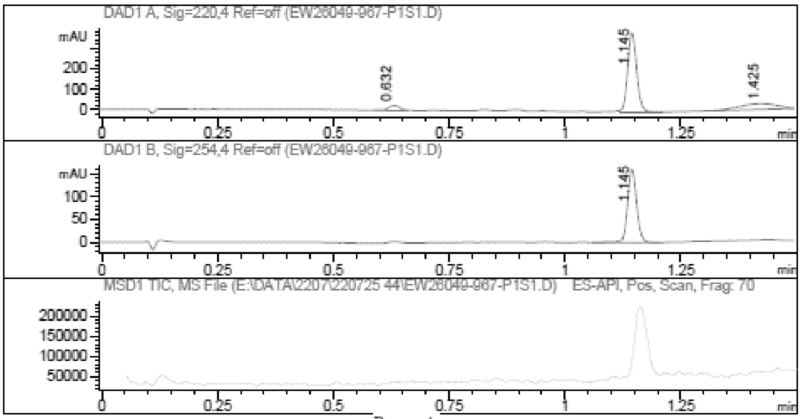

Report

Signal 1 : DAD1 A, Sig=220,4 Ref=off

| # | Meas. Ret. | Height | Width | Area | Area % |
|---|---|---|---|---|---|
| 1 | 0.632 | 23.478 | 0.028 | 40.938 | 5.684 |
| 2 | 1.145 | 388.943 | 0.021 | 520.252 | 72.234 |
| 3 | 1.425 | 28.175 | 0.068 | 159.042 | 22.082 |

Signal 2 : DAD1 B, Sig=254,4 Ref=off

| # | Meas. Ret. | Height | Width | Area | Area % |
|---|---|---|---|---|---|
| 1 | 1.145 | 160.041 | 0.021 | 216.532 | 100.000 |

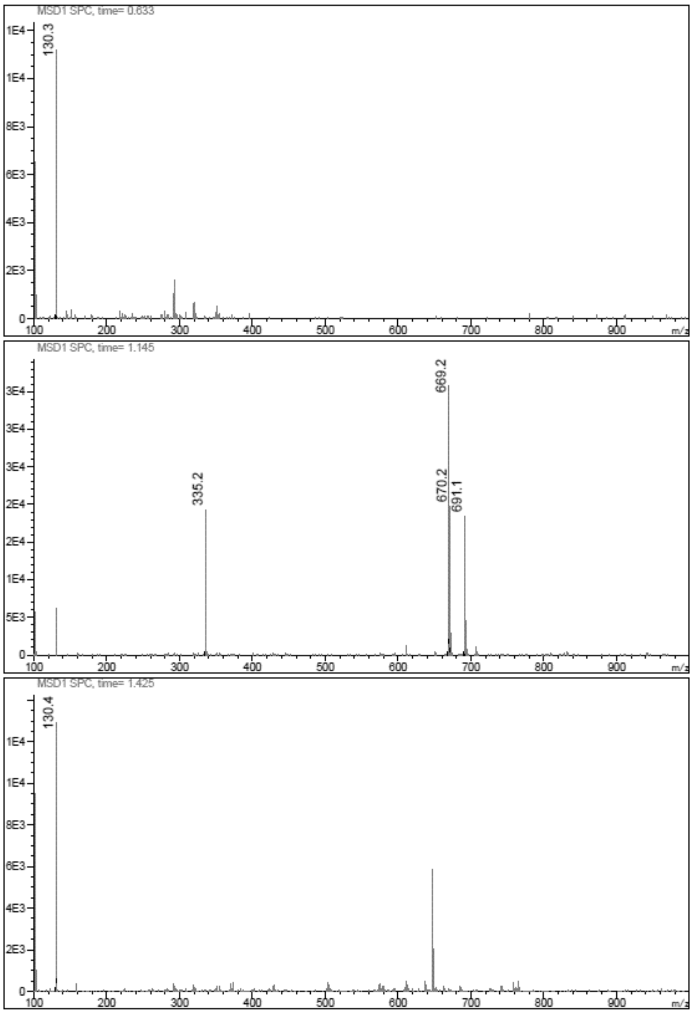

**$^{1}$H NMR** 4-Amino-7-[(2*R*,3*R*,4*S*,5*S*)-5-(1,3-benzothiazol-2-ylsulfanylmethyl) -3,4-dihydroxy-tetrahydrofuran-2-yl]pyrrolo[2,3-d]pyrimidine-5-carbonitrile **(29)**

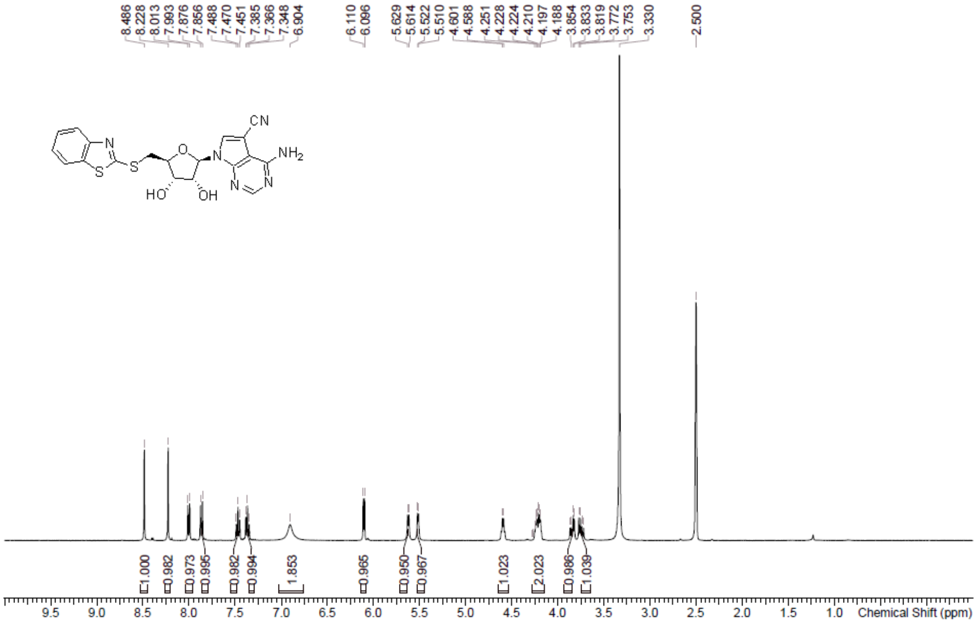

**HPLC** 4-Amino-7-[(2*R*,3*R*,4*S*,5*S*)-5-(1,3-benzothiazol-2-ylsulfanylmethyl) -3,4-dihydroxy-tetrahydrofuran-2-yl]pyrrolo[2,3-d]pyrimidine-5-carbonitrile **(29)**

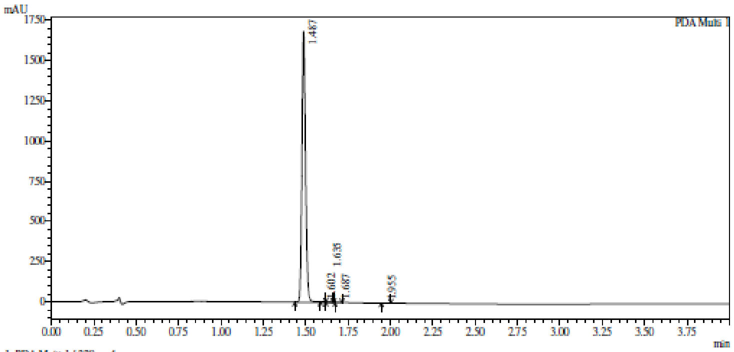

1 PDA Multi 1 / 220nm,4nm

Integration result

PeakTable

PDA Ch1 220nm

| Peak# | Ret. Time | USP Width | Resolution | Height | Area | Area % |
|---|---|---|---|---|---|---|
| 1 | 1.487 | 0.039 | 0.000 | 1684535 | 2406612 | 99.765 |
| 2 | 1.602 | 0.037 | 3.048 | 1499 | 1836 | 0.076 |
| 3 | 1.635 | 0.048 | 0.769 | 1227 | 1595 | 0.066 |
| 4 | 1.687 | 0.035 | 1.261 | 940 | 1093 | 0.045 |
| 5 | 1.955 | 0.051 | 6.204 | 612 | 1145 | 0.047 |
| Total | | | | 1688813 | 2412282 | 100.000 |

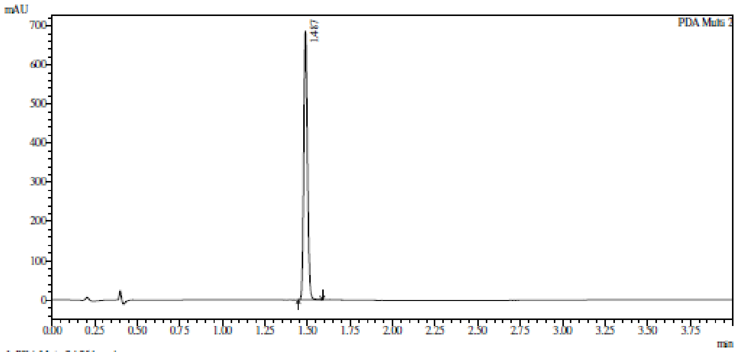

1 PDA Multi 2 / 254nm,4nm

Integration result

PeakTable

PDA Ch2 254nm

| Peak# | Ret. Time | USP Width | Resolution | Height | Area | Area % |
|---|---|---|---|---|---|---|
| 1 | 1.487 | 0.040 | 0.000 | 685292 | 1006213 | 100.000 |
| Total | | | | 685292 | 1006213 | 100.000 |

**HRMC** 4-Amino-7-[(2*R*,3*R*,4*S*,5*S*)-5-(1,3-benzothiazol-2-ylsulfanylmethyl) -3,4-dihydroxy-tetrahydrofuran-2-yl]pyrrolo[2,3-d]pyrimidine-5-carbonitrile **(29)**

**Compound Table**

| Label | Tgt Score | Mass Error (ppm) | Tgt Formula | Obs. RT | Ref. Mass | Obs. Mass |
|---|---|---|---|---|---|---|
| Cpd 1: C19 H16 N6 O3 S2; 1.929 | 97.19 | 3.08 | C19 H16 N6 O3 S2 | 1.929 | 440.0725 | 440.0739 |

| Obs. *m/z* | Obs. RT | Obs. Mass | Tgt Formula | Tgt Mass | Tgt Mass Error (ppm) | Find Cpds Algorithm |
|---|---|---|---|---|---|---|
| 463.0625 | 1.929 | 440.0739 | C19 H16 N6 O3 S2 | 440.0725 | 3.08 | Find by Formula |

**Compound Chromatograms**

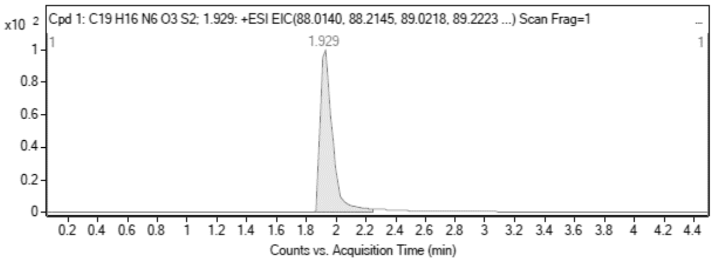

MS Zoomed Spectrum

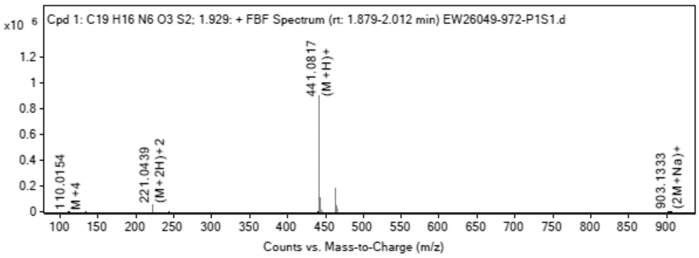

**MS Spectrum Peak List**

| Obs. *m/z* | Charge | Abund | Ion/Isotope |
|---|---|---|---|
| 221.0439 | 2 | 48245.51 | (M+2H)+2 |
| 221.5454 | 2 | 11702.35 | (M+2H)+2 |
| 222.0334 | 2 | 6460.34 | (M+2H)+2 |
| 440.0802 | 1 | 3350.33 | M+ |
| 441.0817 | 1 | 904731.38 | (M+H)+ |
| 442.0832 | 1 | 224744.67 | (M+H)+ |
| 443.079 | 1 | 107433.2 | (M+H)+ |
| 463.0625 | 1 | 183445.64 | (M+Na)+ |
| 464.0646 | 1 | 43487.23 | (M+Na)+ |
| 465.0608 | 1 | 21211.3 | (M+Na)+ |

MS Zoomed Spectrum

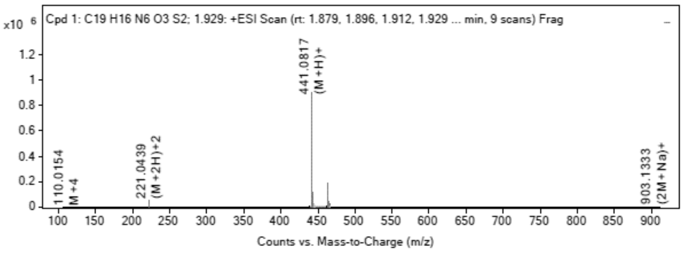

**MS Spectrum Peak List**

| Obs. *m/z* | Charge | Abund | Ion/Isotope | Tgt Mass Error (ppm) |
|---|---|---|---|---|
| 221.0439 | 2 | 48245.51 | (M+2H)+2 | -1.52 |
| 221.5454 | 2 | 11702.35 | (M+2H)+2 | -2.74 |
| 222.0334 | 2 | 6460.34 | (M+2H)+2 | 42.25 |
| 440.0802 | 1 | 3350.33 | M+ | -18.6 |
| 441.0817 | 1 | 904731.38 | (M+H)+ | -4.36 |
| 442.0832 | 1 | 224744.67 | (M+H)+ | -1.98 |
| 443.079 | 1 | 107433.2 | (M+H)+ | -1.81 |
| 463.0625 | 1 | 183445.64 | (M+Na)+ | -1.6 |
| 464.0646 | 1 | 43487.23 | (M+Na)+ | -0.6 |
| 465.0608 | 1 | 21211.3 | (M+Na)+ | -1.35 |

--- End Of Report ---

**$^1$H NMR** 7-[(2*R*,3*R*,4*R*,5*S*)-5-(1,3-benzothiazol-2-ylsulfanylmethyl)-3,4-bis [[tert-butyl(dimethyl)silyl]oxy]tetrahydrofuran-2-yl]-5-(2-trimethylsilylethynyl)pyrrolo[2,3-d]pyrimidin-4-amine **(71)**

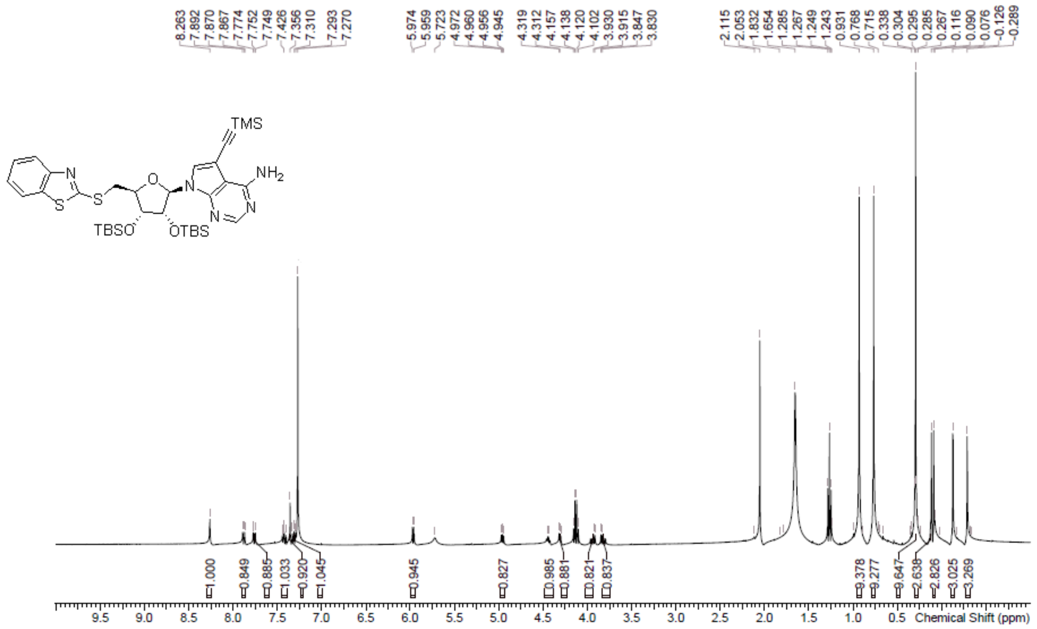

**$^1$H NMR** (2*R*,3*R*,4*S*,5*S*)-2-(4-Amino-5-ethynyl-pyrrolo[2,3-d]pyrimidin-7-yl) -5-(1,3-benzothiazol-2-ylsulfanylmethyl)tetrahydrofuran-3,4-diol **(30)**

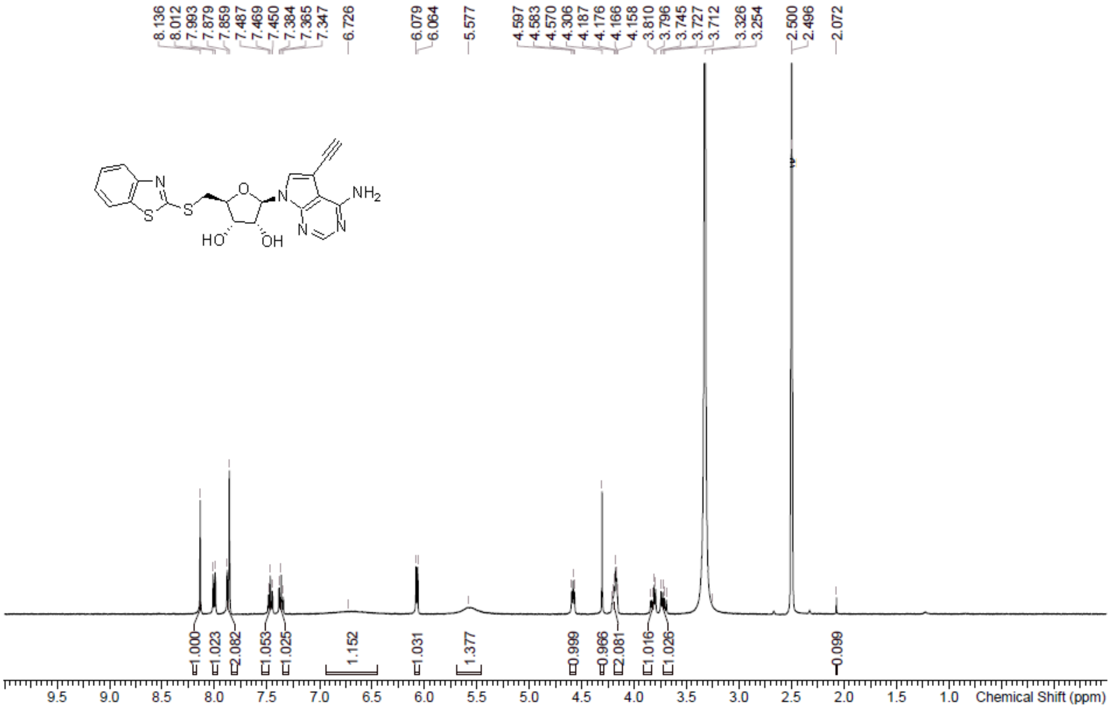

**HPLC** (2*R*,3*R*,4*S*,5*S*)-2-(4-Amino-5-ethynyl-pyrrolo[2,3-d]pyrimidin-7-yl) -5-(1,3-benzothiazol-2-ylsulfanylmethyl)tetrahydrofuran-3,4-diol **(30)**

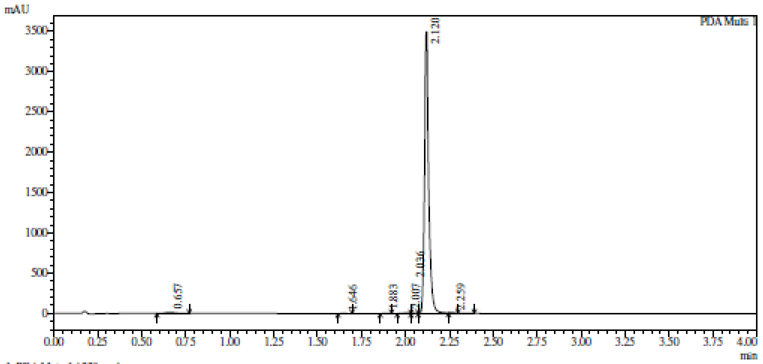

1 PDA Multi 1 / 220nm,4nm

Integration result

PeakTable

PDA Ch1 220nm

| Peak# | Ret. Time | USP Width | Resolution | Height | Area | Area % |
|---|---|---|---|---|---|---|
| 1 | 0.657 | 0.109 | 0.000 | 12357 | 51635 | 0.883 |
| 2 | 1.646 | 0.054 | 12.138 | 1711 | 3279 | 0.056 |
| 3 | 1.883 | 0.047 | 4.719 | 1622 | 2724 | 0.047 |
| 4 | 2.007 | 0.081 | 1.947 | 8785 | 19846 | 0.339 |
| 5 | 2.036 | 1.134 | 0.047 | 6184 | 11813 | 0.202 |
| 6 | 2.120 | 0.038 | 0.143 | 3494132 | 5758487 | 98.421 |
| 7 | 2.259 | 0.044 | 3.413 | 1880 | 3102 | 0.053 |
| Total | | | | 3526670 | 5850885 | 100.000 |

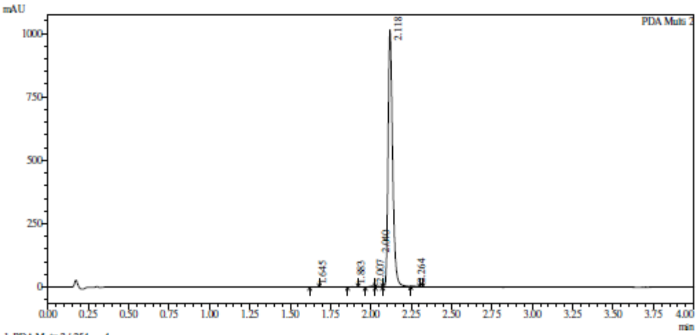

1 PDA Multi 2 / 254nm,4nm

Integration result

PeakTable

PDA Ch2 254nm

| Peak# | Ret. Time | USP Width | Resolution | Height | Area | Area % |
|---|---|---|---|---|---|---|
| 1 | 1.645 | 0.049 | 0.000 | 643 | 1086 | 0.054 |
| 2 | 1.883 | 0.049 | 4.864 | 1097 | 1890 | 0.094 |
| 3 | 2.007 | 0.076 | 1.975 | 3675 | 7550 | 0.377 |
| 4 | 2.040 | 0.300 | 0.178 | 2657 | 5476 | 0.273 |
| 5 | 2.118 | 0.051 | 0.444 | 1013408 | 1985331 | 99.146 |
| 6 | 2.264 | 0.043 | 3.111 | 760 | 1091 | 0.054 |
| Total | | | | 1022240 | 2002424 | 100.000 |

**HRMS** (2*R*,3*R*,4*S*,5*S*)-2-(4-Amino-5-ethynyl-pyrrolo[2,3-d]pyrimidin-7-yl) -5-(1,3-benzothiazol-2-ylsulfanylmethyl)tetrahydrofuran-3,4-diol **(30)**

**Compound Table**

| Label | Tgt Score | Mass Error (ppm) | Tgt Formula | Obs. RT | Ref. Mass | Obs. Mass |
|---|---|---|---|---|---|---|
| Cpd 1: C20 H17 N5 O3 S2; 1.922 | 97.93 | 3.61 | C20 H17 N5 O3 S2 | 1.922 | 439.0773 | 439.0789 |

| Obs. *m/z* | Obs. RT | Obs. Mass | Tgt Formula | Tgt Mass | Tgt Mass Error (ppm) | Find Cpds Algorithm |
|---|---|---|---|---|---|---|
| 462.0668 | 1.922 | 439.0789 | C20 H17 N5 O3 S2 | 439.0773 | 3.61 | Find by Formula |

**Compound Chromatograms**

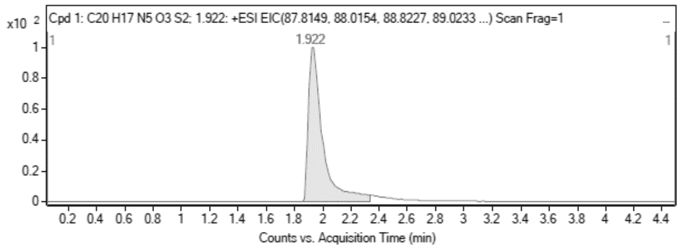

**MS Zoomed Spectrum**

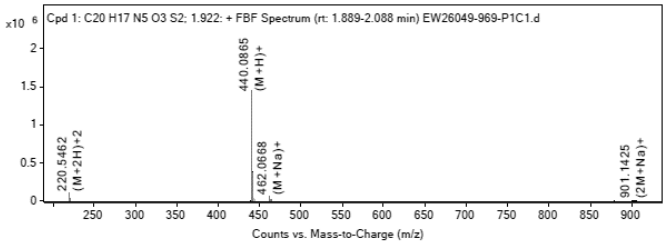

**MS Spectrum Peak List**

| Obs. *m/z* | Charge | Abund | Ion/Isotope |
|---|---|---|---|
| 220.5462 | 2 | 96236.7 | (M+2H)+2 |
| 221.0477 | 2 | 24290.77 | (M+2H)+2 |
| 221.5456 | 2 | 12155.86 | (M+2H)+2 |
| 440.0865 | 1 | 1453699.13 | (M+H)+ |
| 441.0885 | 1 | 383837.03 | (M+H)+ |
| 442.0842 | 1 | 180409.28 | (M+H)+ |
| 462.0668 | 1 | 58040.09 | (M+Na)+ |
| 463.0695 | 1 | 14781.84 | (M+Na)+ |
| 464.0656 | 1 | 7045.21 | (M+Na)+ |
| 901.1425 | 1 | 4713.46 | (2M+Na)+ |

**MS Zoomed Spectrum**

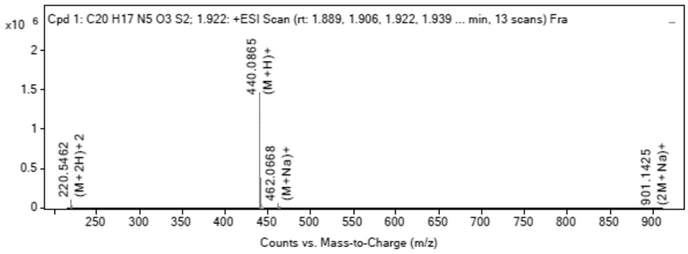

**MS Spectrum Peak List**

| Obs. *m/z* | Charge | Abund | Ion/Isotope | Tgt Mass Error (ppm) |
|---|---|---|---|---|
| 220.5462 | 2 | 96236.7 | (M+2H)+2 | -1.42 |
| 221.0477 | 2 | 24290.77 | (M+2H)+2 | -2.17 |
| 221.5456 | 2 | 12155.86 | (M+2H)+2 | -2.08 |
| 440.0865 | 1 | 1453699.13 | (M+H)+ | -4.47 |
| 441.0885 | 1 | 383837.03 | (M+H)+ | -2.83 |
| 442.0842 | 1 | 180409.28 | (M+H)+ | -2.43 |
| 462.0668 | 1 | 58040.09 | (M+Na)+ | -0.64 |
| 463.0695 | 1 | 14781.84 | (M+Na)+ | -0.59 |
| 464.0656 | 1 | 7045.21 | (M+Na)+ | -1.3 |
| 901.1425 | 1 | 4713.46 | (2M+Na)+ | 1.38 |

--- End Of Report ---

**$^1$H NMR** (2*R*,3*R*,4*S*,5*S*)-2-(4-amino-5-ethyl-pyrrolo[2,3-d]pyrimidin-7-yl) -5-(1,3-benzothiazol-2-ylsulfanylmethyl)tetrahydrofuran-3,4-diol **(31)**

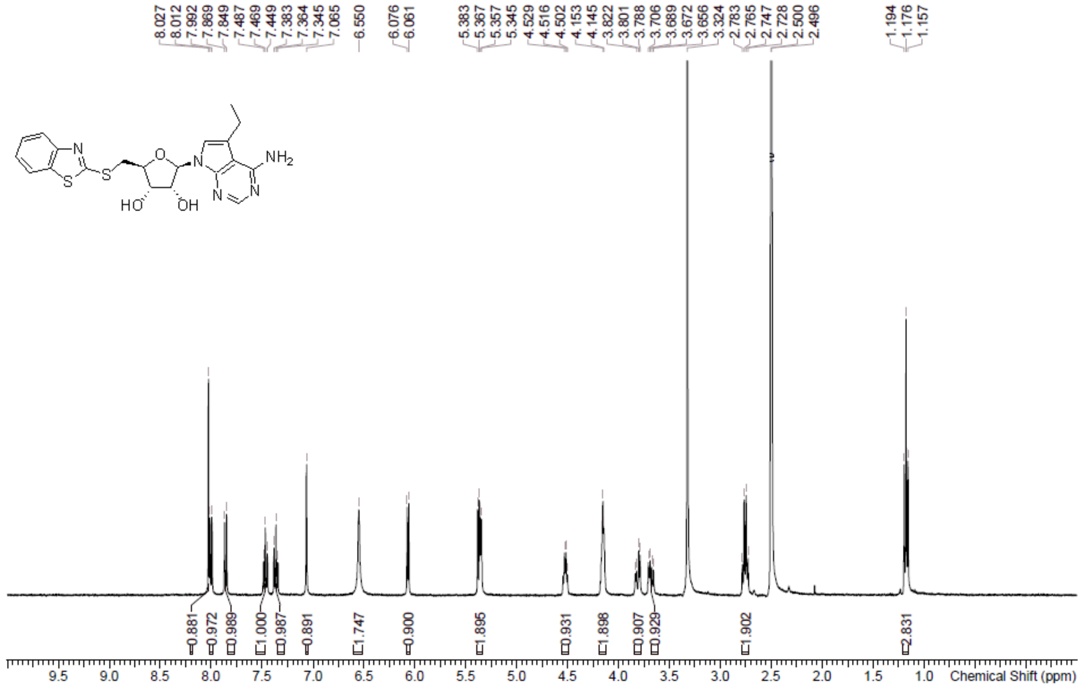

**HPLC** (2*R*,3*R*,4*S*,5*S*)-2-(4-amino-5-ethyl-pyrrolo[2,3-d]pyrimidin-7-yl) -5-(1,3-benzothiazol-2-ylsulfanylmethyl)tetrahydrofuran-3,4-diol **(31)**

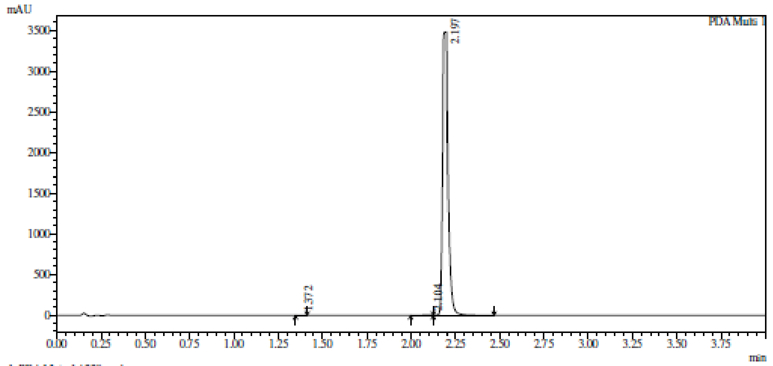

1 PDA Multi 1 / 220nm,4nm

Integration result

PeakTable

PDA Ch1 220nm

| Peak# | Ret. Time | USP Width | Resolution | Height | Area | Area % |
|---|---|---|---|---|---|---|
| 1 | 1.372 | 0.047 | 0.000 | 913 | 1624 | 0.022 |
| 2 | 2.104 | 0.179 | 6.491 | 4171 | 20937 | 0.288 |
| 3 | 2.197 | 0.043 | 0.837 | 3484429 | 7239225 | 99.689 |
| Total | | | | 3489513 | 7261786 | 100.000 |

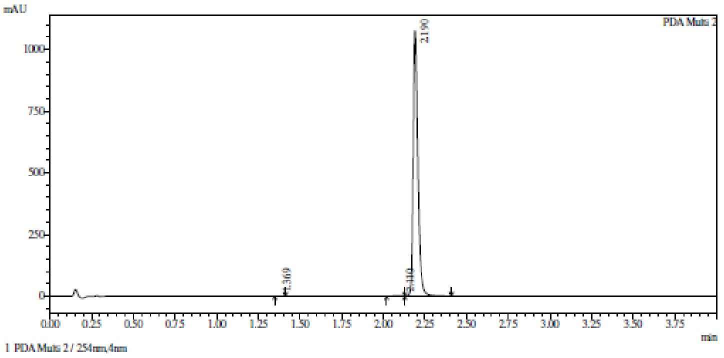

Integration result

PeakTable

PDA Ch2 254nm

| Peak# | Ret. Time | USP Width | Resolution | Height | Area | Area % |
|---|---|---|---|---|---|---|
| 1 | 1.369 | 0.047 | 0.000 | 893 | 1521 | 0.072 |
| 2 | 2.110 | 0.165 | 6.997 | 1561 | 6537 | 0.308 |
| 3 | 2.190 | 0.051 | 0.746 | 1074831 | 2113036 | 99.620 |
| Total | | | | 1077285 | 2121094 | 100.000 |

**HRMC** (2*R*,3*R*,4*S*,5*S*)-2-(4-amino-5-ethyl-pyrrolo[2,3-d]pyrimidin-7-yl) -5-(1,3-benzothiazol-2-ylsulfanylmethyl)tetrahydrofuran-3,4-diol **(31)**

**Compound Table**

| Label | Tgt Score | Mass Error (ppm) | Tgt Formula | Obs. RT | Ref. Mass | Obs. Mass |
|---|---|---|---|---|---|---|
| Cpd 1: C20 H21 N5 O3 S2; 1.953 | 73.38 | -8.84 | C20 H21 N5 O3 S2 | 1.953 | 443.1086 | 443.1047 |

| Obs. *m/z* | Obs. RT | Obs. Mass | Tgt Formula | Tgt Mass | Tgt Mass Error (ppm) | Find Cpds Algorithm |
|---|---|---|---|---|---|---|
| 444.1124 | 1.953 | 443.1047 | C20 H21 N5 O3 S2 | 443.1086 | -8.84 | Find by Formula |

**Compound Chromatograms**

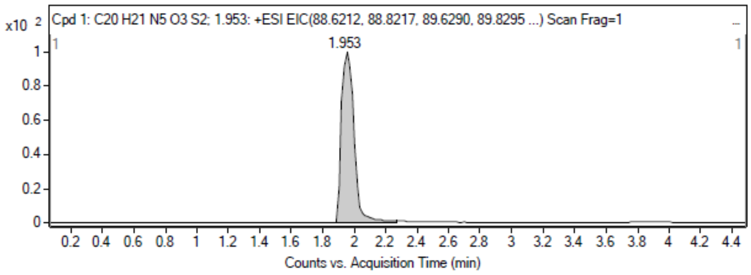

**MS Zoomed Spectrum**

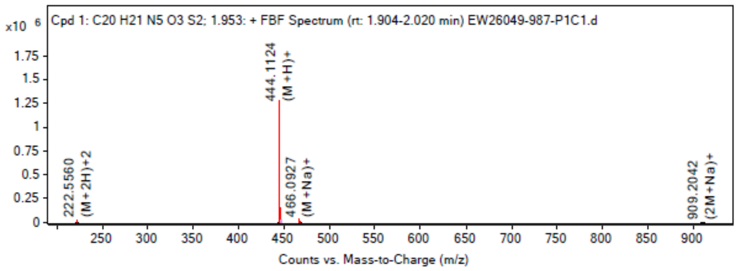

**MS Spectrum Peak List**

| Obs. *m/z* | Charge | Abund | Ion/Isotope |
|---|---|---|---|
| 222.556 | 2 | 28967.12 | (M+2H)+2 |
| 223.0572 | 2 | 7914.94 | (M+2H)+2 |

| | | | |
|---|---|---|---|
| 223.5559 | 2 | 3442.76 | (M+2H)+2 |
| 444.1124 | 1 | 1280260.75 | (M+H)+ |
| 445.1142 | 1 | 335900.38 | (M+H)+ |
| 446.11 | 1 | 157567.98 | (M+H)+ |
| 466.0927 | 1 | 41844.73 | (M+Na)+ |
| 467.0953 | 1 | 10632.84 | (M+Na)+ |
| 468.0918 | 1 | 5172.52 | (M+Na)+ |
| 909.2042 | 1 | 5150.29 | (2M+Na)+ |

MS Zoomed Spectrum

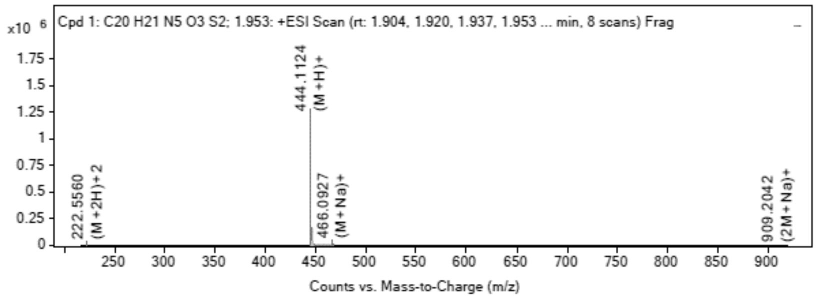

MS Spectrum Peak List

| Obs. m/z | Charge | Abund | Ion/Isotope | Tgt Mass Error (ppm) |
|---|---|---|---|---|
| 222.556 | 2 | 28967.12 | (M+2H)+2 | 24.93 |
| 223.0572 | 2 | 7914.94 | (M+2H)+2 | 25.82 |
| 223.5559 | 2 | 3442.76 | (M+2H)+2 | 22 |
| 444.1124 | 1 | 1280260.75 | (M+H)+ | 7.75 |
| 445.1142 | 1 | 335900.38 | (M+H)+ | 9.7 |
| 446.11 | 1 | 157567.98 | (M+H)+ | 9.91 |
| 466.0927 | 1 | 41844.73 | (M+Na)+ | 10.85 |
| 467.0953 | 1 | 10632.84 | (M+Na)+ | 11.12 |
| 468.0918 | 1 | 5172.52 | (M+Na)+ | 9.69 |
| 909.2042 | 1 | 5150.29 | (2M+Na)+ | 2.44 |

--- End Of Report ---

**$^1$H NMR** 9-[(2*R*,3*R*,4*R*,5*R*)-5-(azidomethyl)-3,4-bis[[tert-butyl(dimethyl)silyl] oxy]tetrahydrofuran-2-yl]purin-6-amine **(107)**

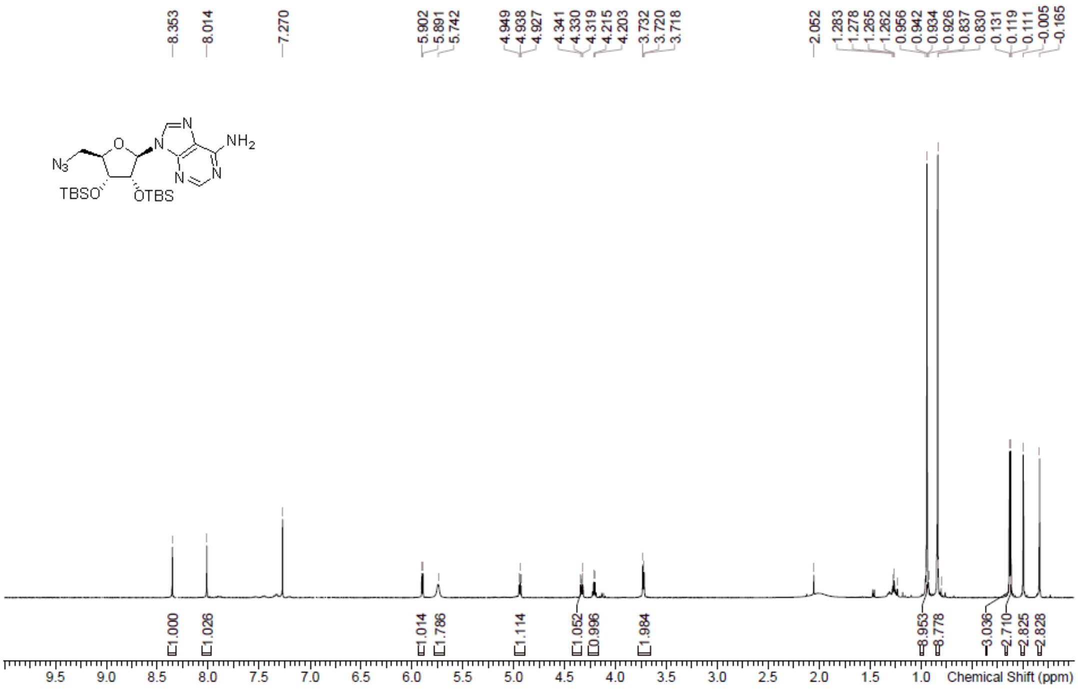

**LCMS** 9-[(2*R*,3*R*,4*R*,5*R*)-5-(azidomethyl)-3,4-bis[[tert-butyl(dimethyl)silyl] oxy]tetrahydrofuran-2-yl]purin-6-amine **(107)**

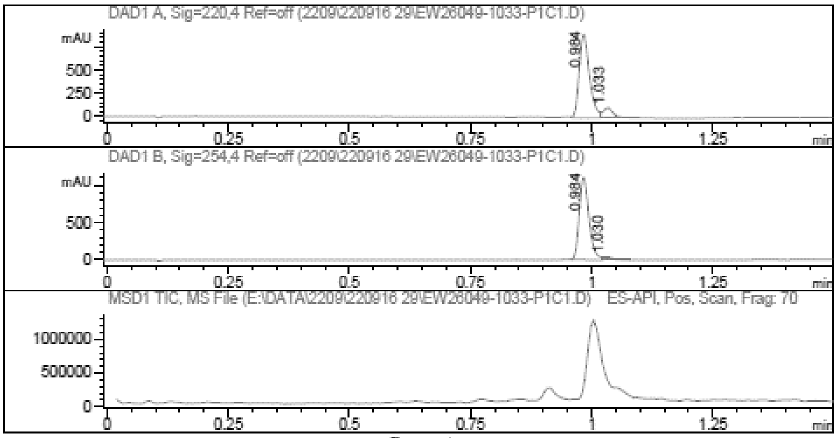

Report

Signal 1 : DAD1 A, Sig=220,4 Ref=off

| # | Meas. Ret. | Height | Width | Area | Area % |
|---|---|---|---|---|---|
| 1 | 0.984 | 897.489 | 0.022 | 1325.514 | 89.852 |
| 2 | 1.033 | 107.453 | 0.021 | 149.712 | 10.148 |

Signal 2 : DAD1 B, Sig=254,4 Ref=off

| # | Meas. Ret. | Height | Width | Area | Area % |
|---|---|---|---|---|---|
| 1 | 0.984 | 1102.536 | 0.021 | 1545.263 | 97.180 |
| 2 | 1.030 | 35.657 | 0.019 | 44.847 | 2.820 |

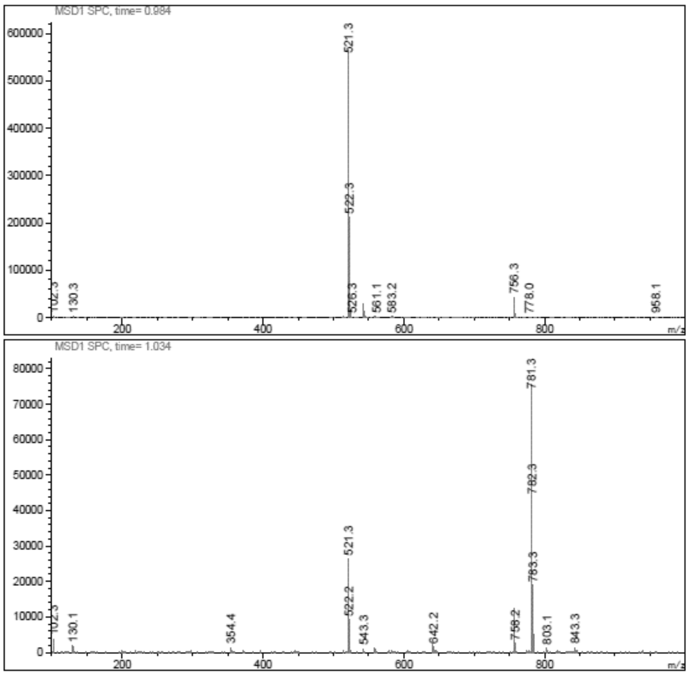

**LCMS** 9-[(2*R*,3*R*,4*R*,5*R)*-5-(Aminomethyl)-3,4-bis[[tert-butyl(dimethyl)silyl] oxy]tetrahydrofuran-2-yl]purin-6-amine **(108)**

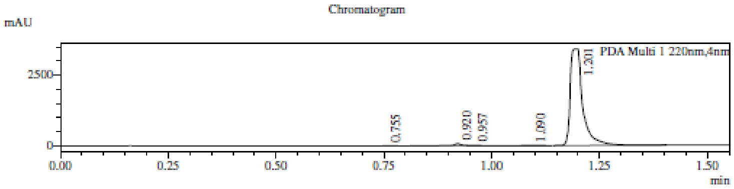

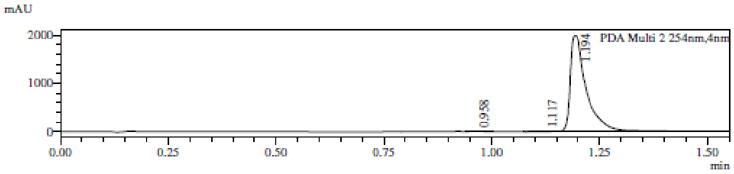

1 PDA Multi 1 / 220nm,4nm
2 PDA Multi 2 / 254nm,4nm

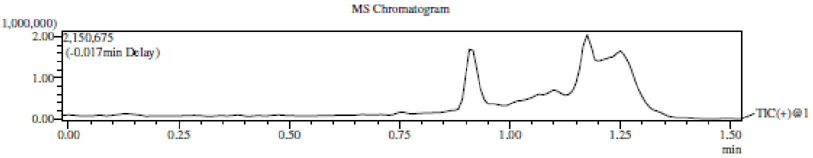

Integration Result

Peak Table

PDA Ch1 220nm

| Peak# | Ret. Time | Height | Height% | USP Width | Area | Area% |
|---|---|---|---|---|---|---|
| 1 | 0.755 | 7103 | 0.203 | 0.038 | 12201 | 0.173 |
| 2 | 0.920 | 73487 | 2.103 | 0.028 | 78367 | 1.113 |
| 3 | 0.957 | 10037 | 0.287 | 0.076 | 17928 | 0.255 |
| 4 | 1.090 | 5979 | 0.171 | 0.057 | 11961 | 0.170 |
| 5 | 1.201 | 3397060 | 97.235 | 0.040 | 6918098 | 98.289 |

Peak Table

PDA Ch2 254nm

| Peak# | Ret. Time | Height | Height% | USP Width | Area | Area% |
|---|---|---|---|---|---|---|
| 1 | 0.958 | 4517 | 0.226 | 0.050 | 7996 | 0.156 |
| 2 | 1.117 | 4908 | 0.246 | 0.042 | 8461 | 0.166 |
| 3 | 1.194 | 1986579 | 99.528 | 0.054 | 5095933 | 99.678 |

Mass Spectrum

RetTime: 0.758 Datafile: D:\DATA\2022\2209\220919\EW26049-1038-P1C1.lcd

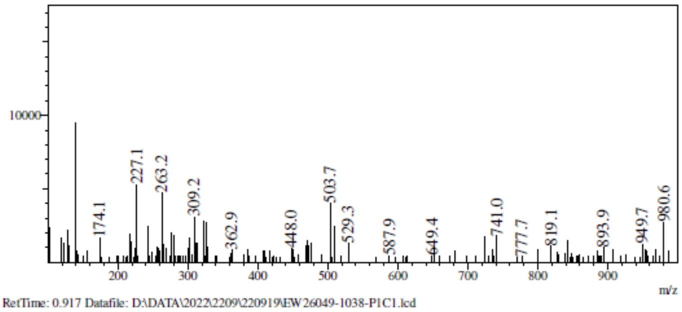

RetTime: 0.917 Datafile: D:\DATA\2022\2209\220919\EW26049-1038-P1C1.lcd

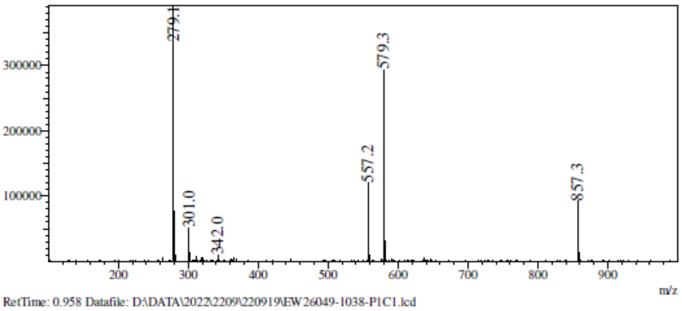

RetTime: 0.958 Datafile: D:\DATA\2022\2209\220919\EW26049-1038-P1C1.lcd

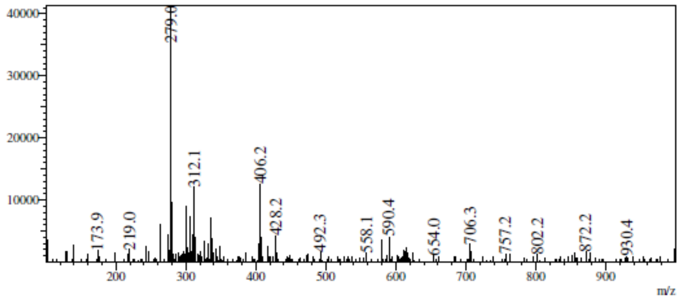

RetTime: 1.092 Datafile: D:\DATA\2022\2209\220919\EW26049-1038-P1C1.lcd

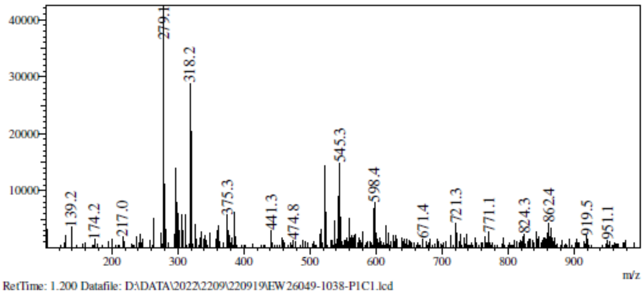

RetTime: 1.200 Datafile: D:\DATA\2022\2209\220919\EW26049-1038-P1C1.lcd

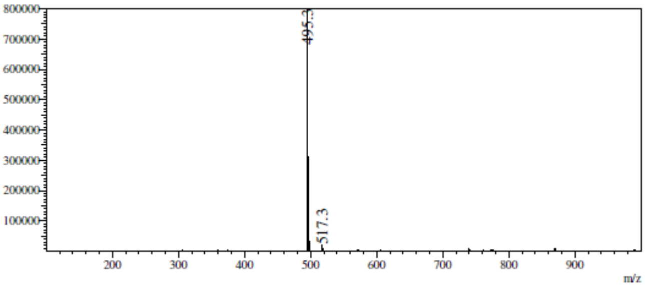

**LCMS** *N*-[[(2*R*,3*R*,4*R*,5*R*)-5-(6-Aminopurin-9-yl)-3,4-bis[[tert-butyl (dimethyl)silyl]oxy]tetrahydrofuran-2-yl]methyl]naphthalene-2-sulfonamide **(109)**

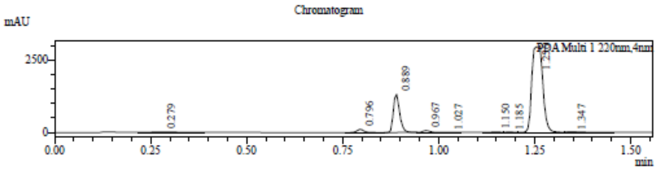

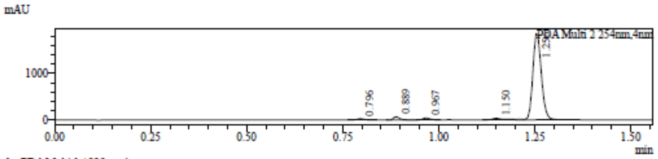

1 PDA Multi 1 / 220nm,4nm
2 PDA Multi 2 / 254nm,4nm

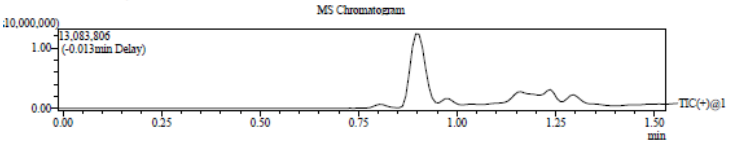

Integration Result

Peak Table

PDA Ch1 220nm

| Peak# | Ret. Time | Height | Height% | USP Width | Area | Area% |
|---|---|---|---|---|---|---|
| 1 | 0.279 | 17989 | 0.398 | 0.118 | 74948 | 0.956 |
| 2 | 0.796 | 108998 | 2.409 | 0.037 | 134718 | 1.718 |
| 3 | 0.889 | 1327751 | 29.348 | 0.034 | 1531207 | 19.529 |
| 4 | 0.967 | 63866 | 1.412 | 0.040 | 83593 | 1.066 |
| 5 | 1.027 | 5921 | 0.131 | 0.043 | 8531 | 0.109 |
| 6 | 1.150 | 15481 | 0.342 | 0.045 | 22584 | 0.288 |
| 7 | 1.185 | 11042 | 0.244 | 0.060 | 17132 | 0.218 |
| 8 | 1.256 | 2967055 | 65.582 | 0.055 | 5960097 | 76.013 |
| 9 | 1.347 | 6096 | 0.135 | 0.042 | 8056 | 0.103 |

Peak Table

PDA Ch2 254nm

Mass Spectrum

RetTime: 0.278 Datafile: D:\DATA\2022\2209\220920\EW26049-1043-P1C1.lcd

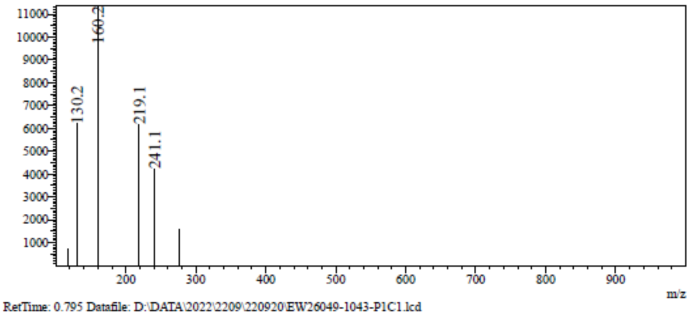

RetTime: 0.795 Datafile: D:\DATA\2022\2209\220920\EW26049-1043-P1C1.lcd

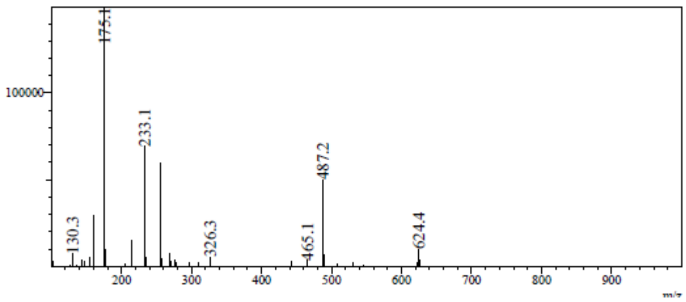

RetTime: 0.887 Datafile: D:\DATA\2022\2209\220920\EW26049-1043-P1C1.lcd

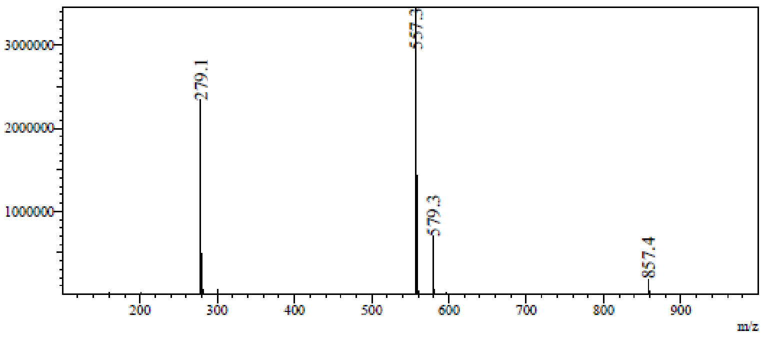

RetTime: 0.970 Datafile: D:\DATA\2022\2209\220920\EW26049-1043-P1C1.lcd

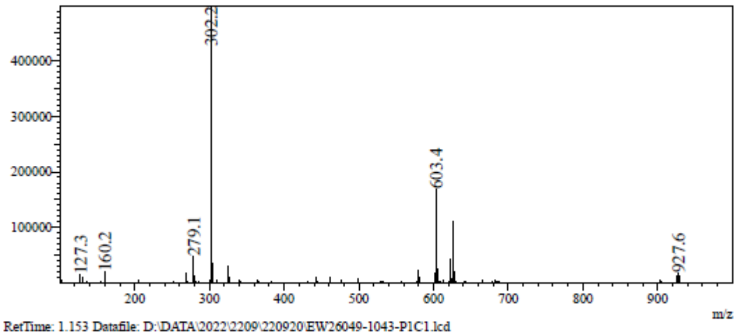

RetTime: 1.153 Datafile: D:\DATA\2022\2209\220920\EW26049-1043-P1C1.lcd

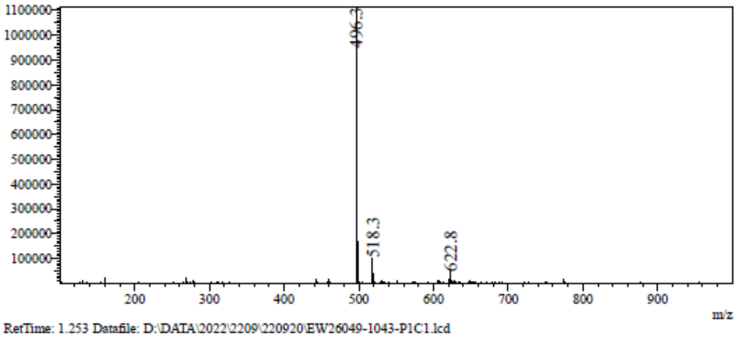

RetTime: 1.253 Datafile: D:\DATA\2022\2209\220920\EW26049-1043-P1C1.lcd

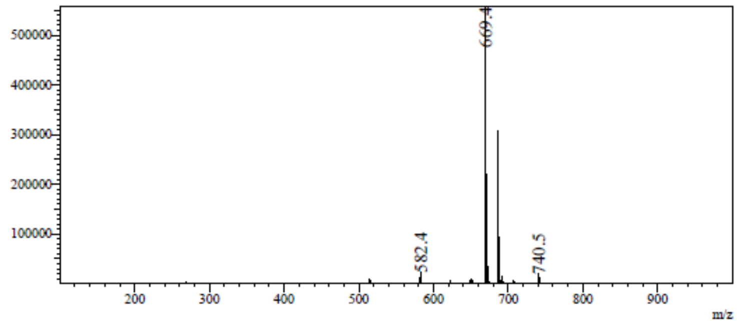

**[1]H NMR** *N*-[[(2*R*,3*S*,4*R*,5*R*)-5-(6-Aminopurin-9-yl)-3,4-dihydroxy-tetrahyd rofuran-2-yl]methyl]naphthalene-2-sulfonamide **(32)**

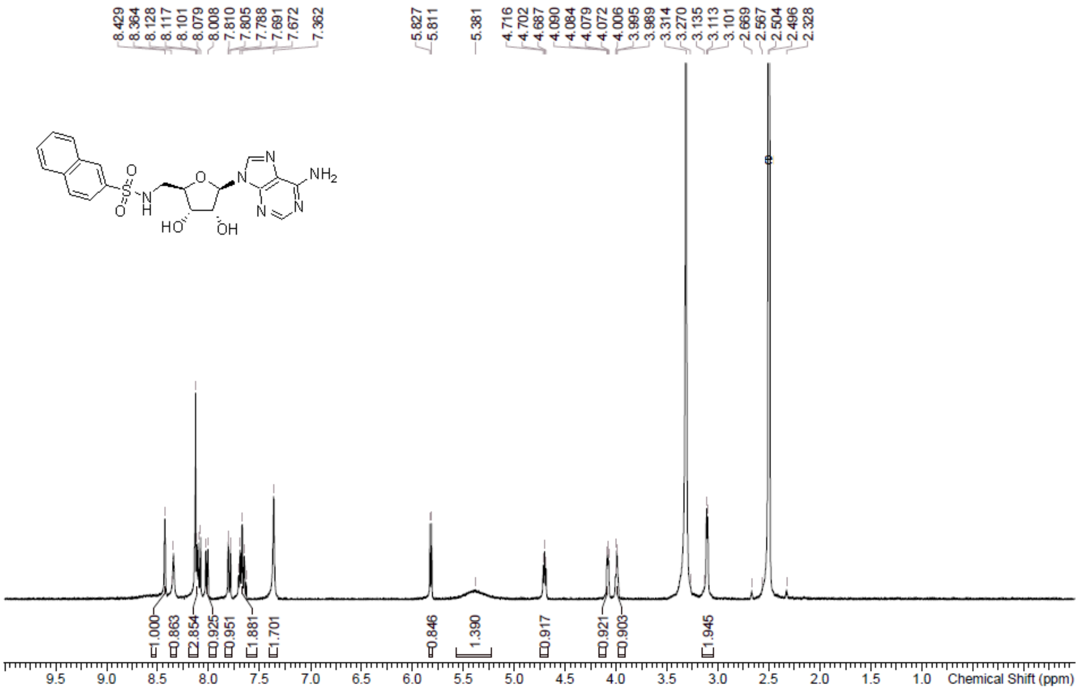

**HPLC** *N*-[[(2*R*,3*S*,4*R*,5*R*)-5-(6-Aminopurin-9-yl)-3,4-dihydroxy-tetrahyd rofuran-2-yl]methyl]naphthalene-2-sulfonamide **(32)**

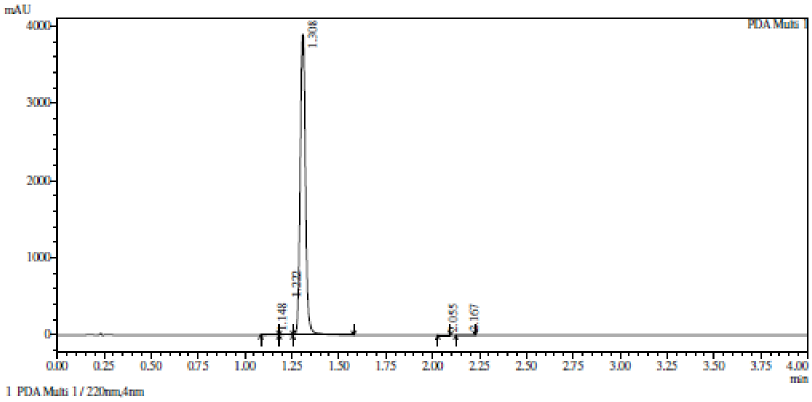

Integration result

PeakTable

PDA Ch1 220nm

| Peak# | Ret. Time | USP Width | Resolution | Height | Area | Area % |
|---|---|---|---|---|---|---|
| 1 | 1.148 | 0.096 | 0.000 | 6594 | 20683 | 0.274 |
| 2 | 1.222 | 0.086 | 0.814 | 7984 | 24836 | 0.329 |
| 3 | 1.308 | 0.050 | 1.275 | 3898289 | 7490537 | 99.306 |
| 4 | 2.055 | 0.043 | 16.227 | 1620 | 2545 | 0.034 |
| 5 | 2.167 | 0.076 | 1.900 | 1548 | 4263 | 0.057 |
| Total | | | | 3916035 | 7542863 | 100.000 |

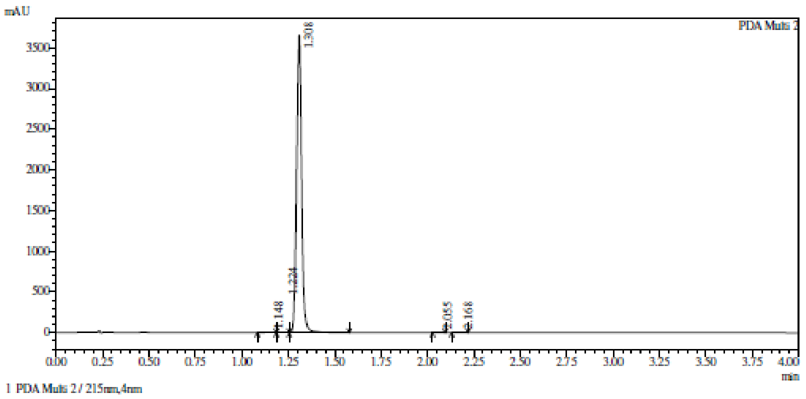

1 PDA Multi 2 / 215nm,4nm

Integration result

PeakTable

PDA Ch2 215nm

| Peak# | Ret. Time | USP Width | Resolution | Height | Area | Area % |
|---|---|---|---|---|---|---|
| 1 | 1.148 | 0.098 | 0.000 | 6377 | 20772 | 0.296 |
| 2 | 1.224 | 0.086 | 0.821 | 7892 | 23905 | 0.341 |
| 3 | 1.308 | 0.050 | 1.238 | 3655501 | 6968253 | 99.282 |
| 4 | 2.055 | 0.042 | 16.225 | 1662 | 2619 | 0.037 |
| 5 | 2.168 | 0.069 | 2.029 | 1232 | 3115 | 0.044 |
| Total | | | | 3672664 | 7018664 | 100.000 |

**HRMC** *N*-[[(2*R*,3*S*,4*R*,5*R*)-5-(6-Aminopurin-9-yl)-3,4-dihydroxy-tetrahyd rofuran-2-yl]methyl]naphthalene-2-sulfonamide **(32)**

Compound Table

| Label | Tgt Score | Mass Error (ppm) | Tgt Formula | Obs. RT | Ref. Mass | Obs. Mass |
|---|---|---|---|---|---|---|
| Cpd 1: C20 H20 N6 O5 S; 1.382 | 55.63 | 19.37 | C20 H20 N6 O5 S | 1.382 | 456.1216 | 456.1304 |

| Obs. m/z | Obs. RT | Obs. Mass | Tgt Formula | Tgt Mass | Tgt Mass Error (ppm) | Find Cpds Algorithm |
|---|---|---|---|---|---|---|
| 479.1183 | 1.382 | 456.1304 | C20 H20 N6 O5 S | 456.1216 | 19.37 | Find by Formula |

Compound Chromatograms

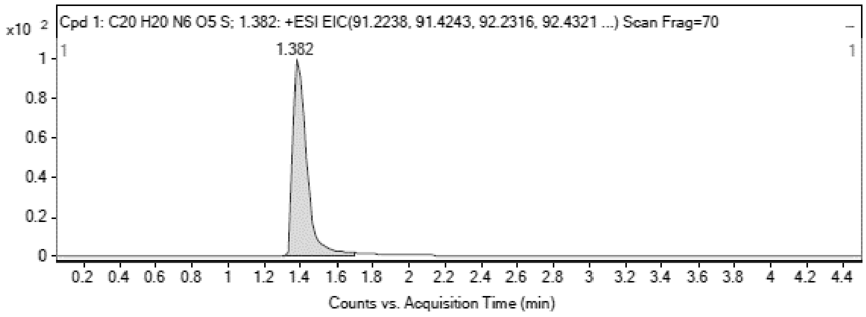

MS Zoomed Spectrum

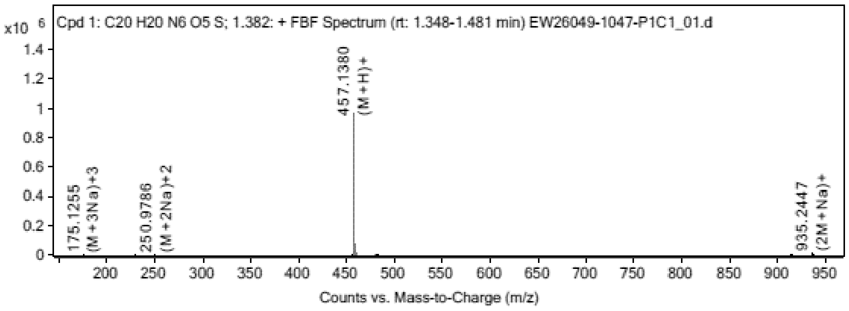

MS Spectrum Peak List

| Obs. m/z | Charge | Abund | Ion/Isotope |
|---|---|---|---|
| 457.138 | 1 | 963884.31 | (M+H)+ |

| | | | |
|---|---|---|---|
| 458.1397 | 1 | 246694.72 | (M+H)+ |
| 459.1368 | 1 | 78569.22 | (M+H)+ |
| 460.138 | 1 | 14403.13 | (M+H)+ |
| 479.1183 | 1 | 8718.25 | (M+Na)+ |
| 480.1208 | 1 | 2249.66 | (M+Na)+ |
| 935.2447 | 1 | 14602.82 | (2M+Na)+ |
| 936.248 | 1 | 7317.42 | (2M+Na)+ |
| 937.2406 | 1 | 4128.79 | (2M+Na)+ |
| 938.2402 | 1 | 1599.48 | (2M+Na)+ |

MS Zoomed Spectrum

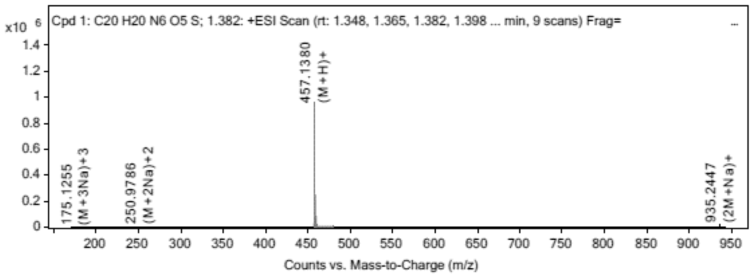

MS Spectrum Peak List

| Obs. m/z | Charge | Abund | Ion/Isotope | Tgt Mass Error (ppm) |
|---|---|---|---|---|
| 457.138 | 1 | 963884.31 | (M+H)+ | -20.01 |
| 458.1397 | 1 | 246694.72 | (M+H)+ | -17.84 |
| 459.1368 | 1 | 78569.22 | (M+H)+ | -16.9 |
| 460.138 | 1 | 14403.13 | (M+H)+ | -16.93 |
| 479.1183 | 1 | 8718.25 | (M+Na)+ | -15.66 |
| 480.1208 | 1 | 2249.66 | (M+Na)+ | -15.12 |
| 935.2447 | 1 | 14602.82 | (2M+Na)+ | -13.16 |
| 936.248 | 1 | 7317.42 | (2M+Na)+ | -13.75 |
| 937.2406 | 1 | 4128.79 | (2M+Na)+ | -7.09 |
| 938.2402 | 1 | 1599.48 | (2M+Na)+ | -5.91 |

--- End Of Report ---

**$^{1}$H NMR** (2*S*,3*R*,4*R*,5*R*)-2-(4-Aminopyrrolo[2,1-f][1,2,4]triazin-7-yl)-3,4- dibenzyloxy-5-(benzyloxymethyl)tetrahydrofuran-2-ol **(111)**

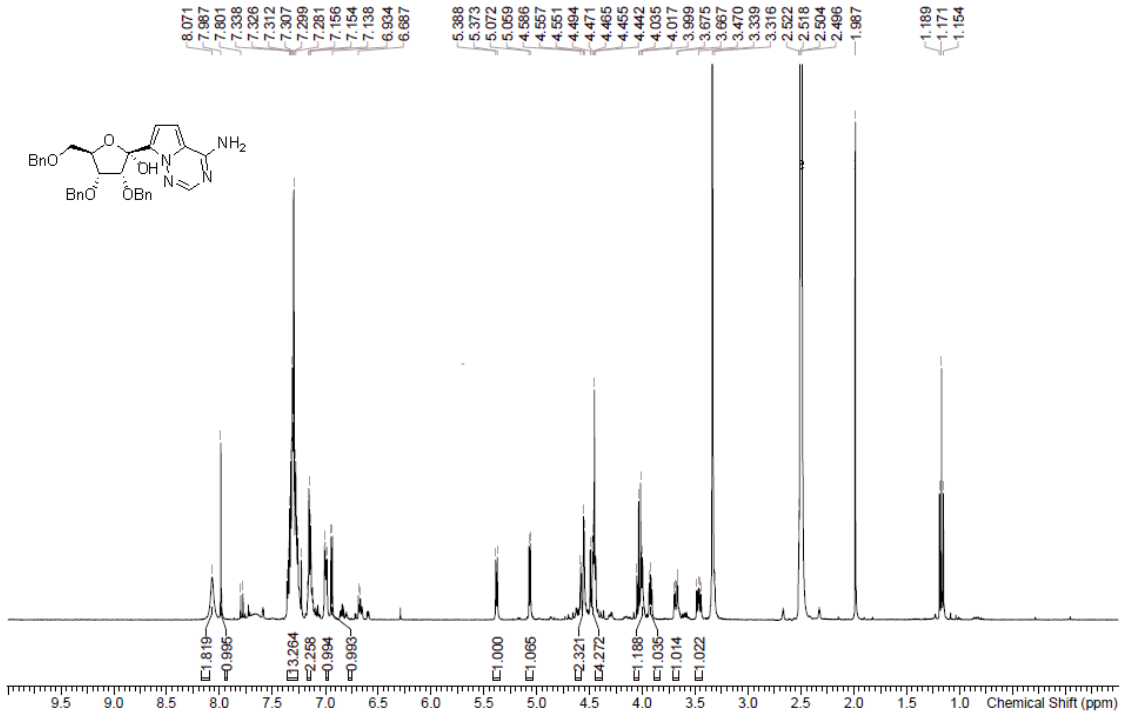

**LCMS** (2*S*,3*R*,4*R*,5*R*)-2-(4-Aminopyrrolo[2,1-f][1,2,4]triazin-7-yl)-3,4- dibenzyloxy-5-(benzyloxymethyl)tetrahydrofuran-2-ol **(111)**

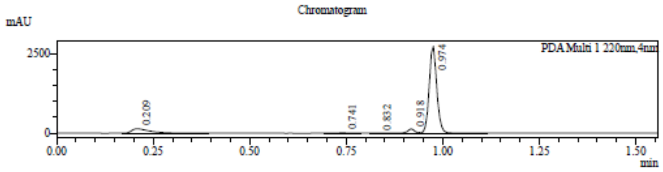

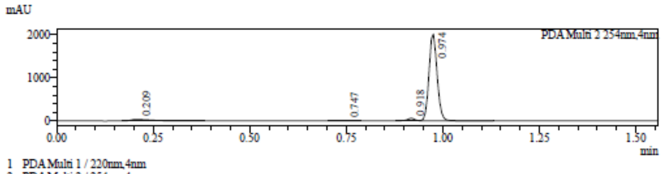

1 PDA Multi 1 / 220nm,4nm
2 PDA Multi 2 / 254nm,4nm

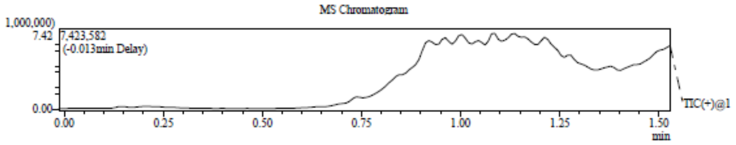

Integration Result

Peak Table

PDA Ch1 220nm

| Peak# | Ret. Time | Height | Height% | USP Width | Area | Area% |
|---|---|---|---|---|---|---|
| 1 | 0.209 | 139773 | 4.625 | 0.091 | 452272 | 10.411 |
| 2 | 0.741 | 17447 | 0.577 | 0.061 | 22276 | 0.513 |
| 3 | 0.832 | 3395 | 0.112 | 0.075 | 5534 | 0.127 |
| 4 | 0.918 | 134929 | 4.464 | 0.036 | 163307 | 3.759 |
| 5 | 0.974 | 2726838 | 90.221 | 0.040 | 3700730 | 85.189 |

Peak Table

PDA Ch2 254nm

| Peak# | Ret. Time | Height | Height% | USP Width | Area | Area% |
|---|---|---|---|---|---|---|
| 1 | 0.209 | 36619 | 1.727 | 0.092 | 119341 | 3.795 |
| 2 | 0.747 | 2360 | 0.111 | 0.052 | 5190 | 0.165 |
| 3 | 0.918 | 61569 | 2.903 | 0.037 | 76247 | 2.424 |

Mass Spectrum

RetTime: 0.212 Datafile: D:\DATA\2022\2210\221019\EW26049-1082-P1E1.lcd

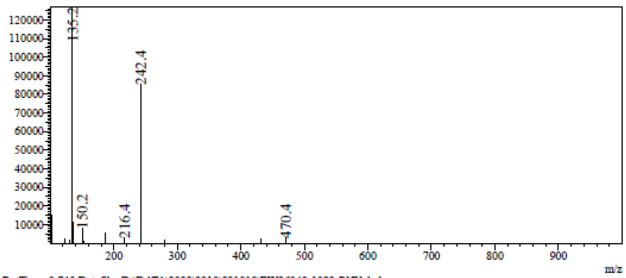

RetTime: 0.745 Datafile: D:\DATA\2022\2210\221019\EW26049-1082-P1E1.lcd

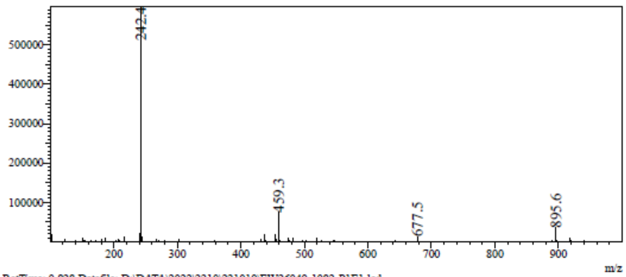

RetTime: 0.828 Datafile: D:\DATA\2022\2210\221019\EW26049-1082-P1E1.lcd

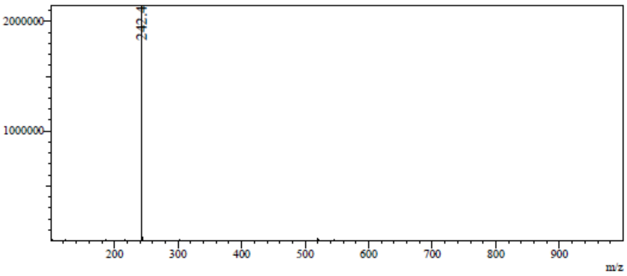

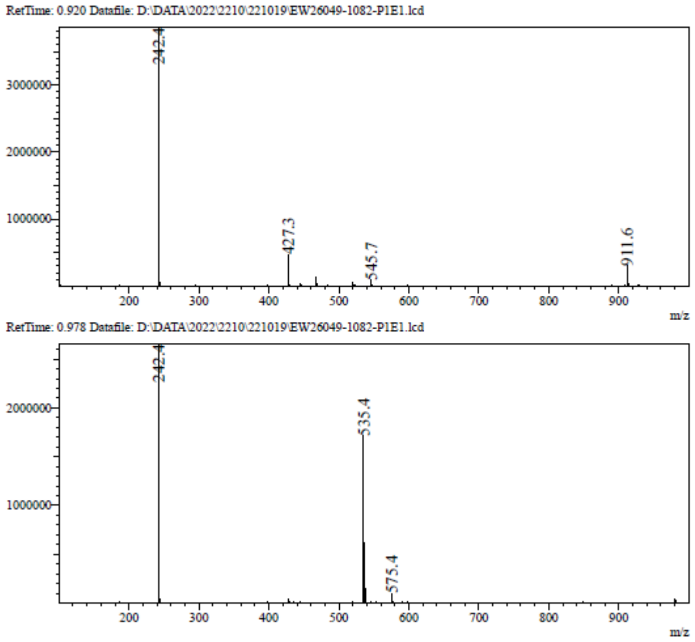

**[1]H NMR** 7-[(2*S*,3*S*,4*R*,5*R*)-3,4-Dibenzyloxy-5-(benzyloxymethyl) tetrahydrofuran-2-yl]pyrrolo[2,1-f][1,2,4]triazin-4-amine **(112)**

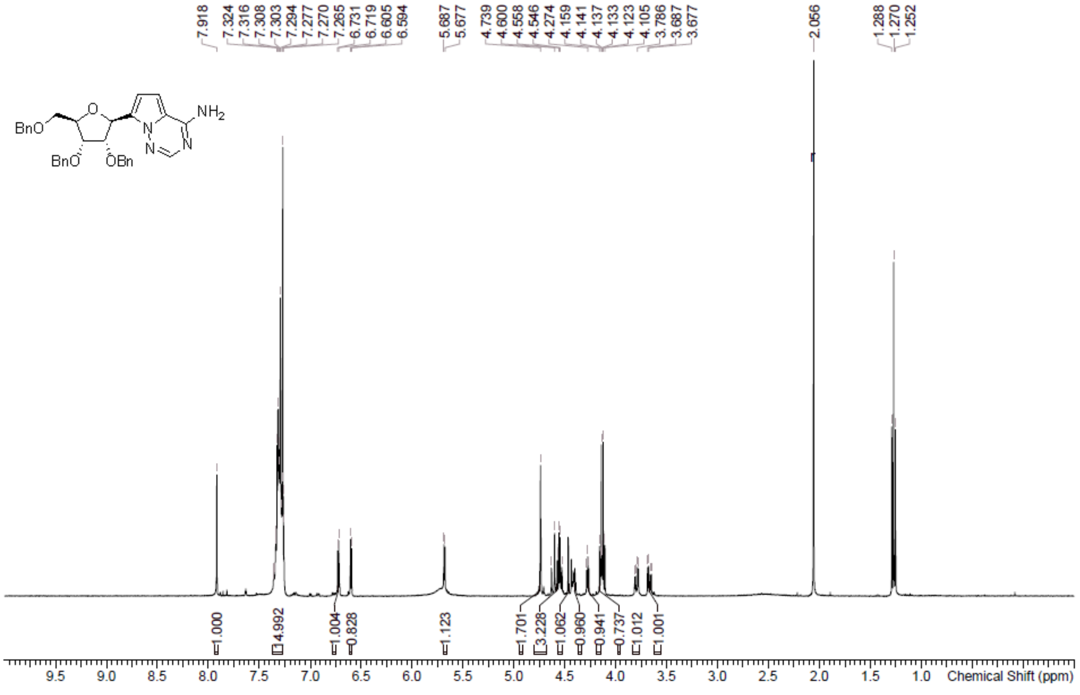

**LCMS** 7-[(2*S*,3*S*,4*R*,5*R*)-3,4-Dibenzyloxy-5-(benzyloxymethyl) tetrahydrofuran-2-yl]pyrrolo[2,1-f][1,2,4]triazin-4-amine **(112)**

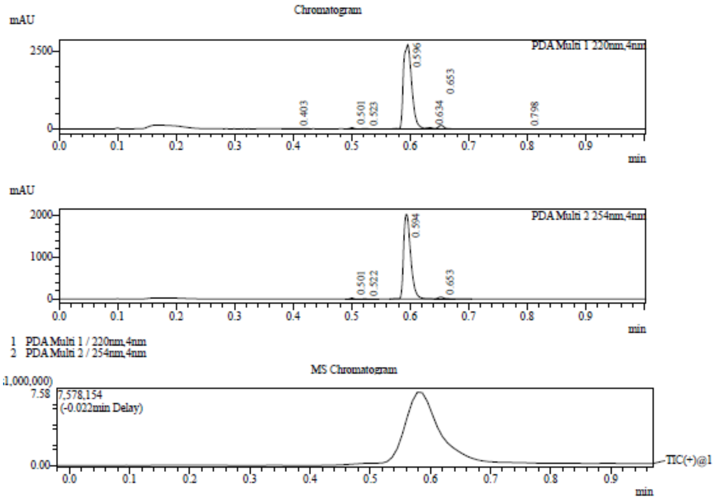

Integration Result

Peak Table

PDA Ch1 220nm

| Peak# | Ret. Time | Height | Height% | USP Width | Area | Area% |
|---|---|---|---|---|---|---|
| 1 | 0.403 | 8122 | 0.276 | 0.050 | 13581 | 0.476 |
| 2 | 0.501 | 39688 | 1.350 | 0.015 | 19264 | 0.676 |
| 3 | 0.523 | 10082 | 0.343 | 0.019 | 6804 | 0.239 |
| 4 | 0.596 | 2736880 | 93.090 | 0.026 | 2715869 | 95.258 |
| 5 | 0.634 | 29710 | 1.011 | 0.014 | 13844 | 0.486 |
| 6 | 0.653 | 113312 | 3.854 | 0.019 | 72626 | 2.547 |
| 7 | 0.798 | 2255 | 0.077 | 0.075 | 9091 | 0.319 |

Peak Table

PDA Ch2 254nm

| Peak# | Ret. Time | Height | Height% | USP Width | Area | Area% |
|---|---|---|---|---|---|---|
| 1 | 0.501 | 22975 | 1.092 | 0.015 | 11201 | 0.633 |
| 2 | 0.522 | 7731 | 0.367 | 0.020 | 5097 | 0.288 |
| 3 | 0.594 | 2028739 | 96.382 | 0.024 | 1721343 | 97.335 |

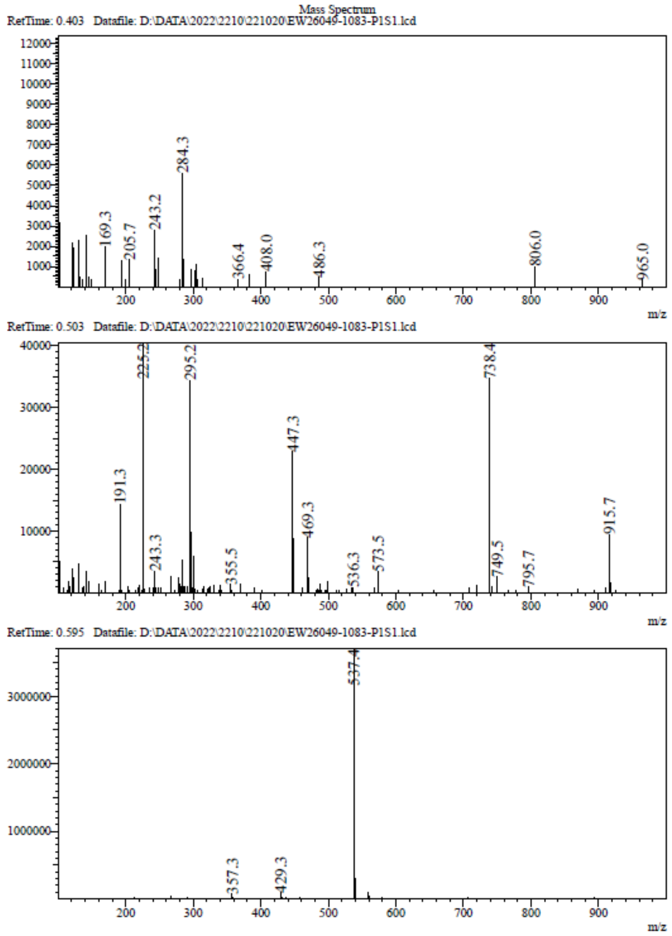

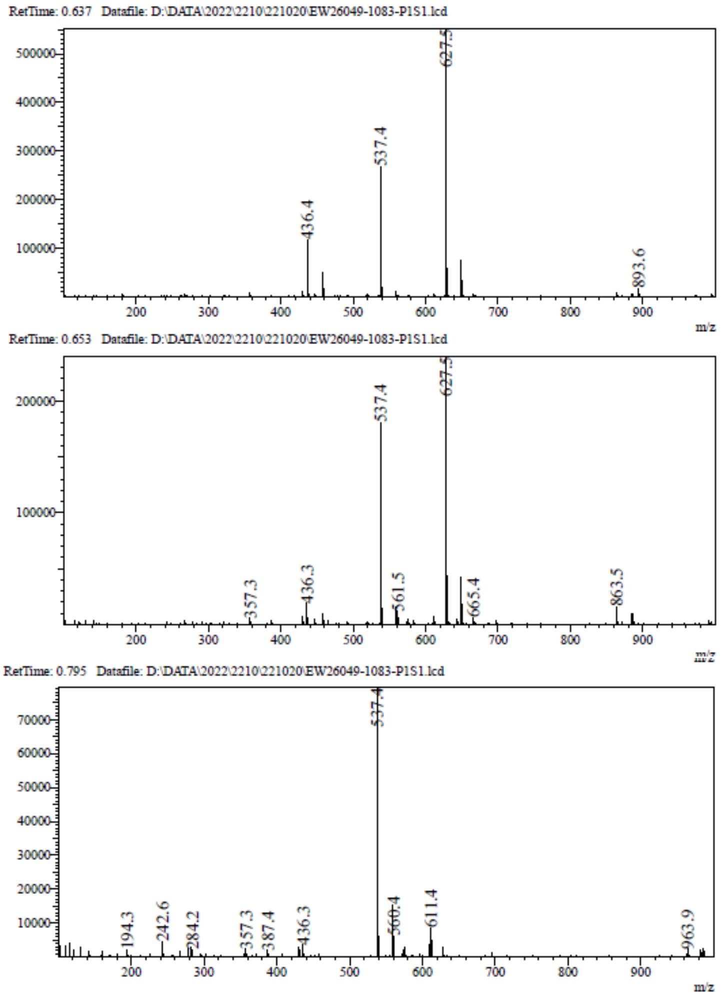

**$^1$H NMR** (2*S*,3*R*,4*S*,5*R*)-2-(4-Aminopyrrolo[2,1-f][1,2,4]triazin-7-yl) -5-(hydroxymethyl)tetrahydrofuran-3,4-diol **(113)**

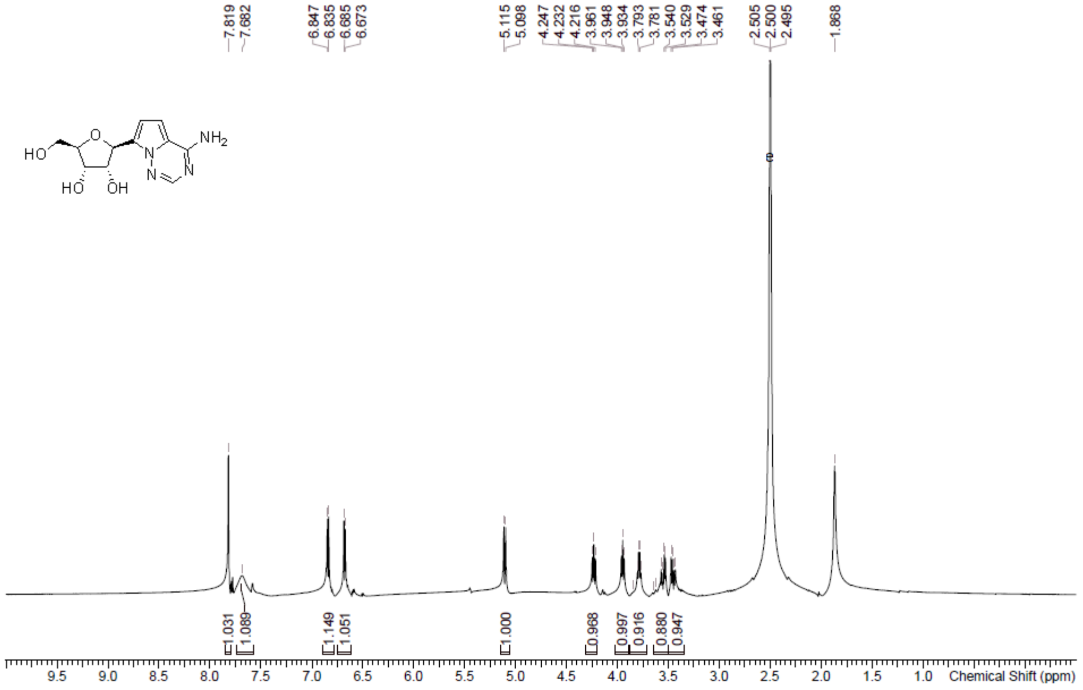

**LCMS** (2*S*,3*R*,4*S*,5*R*)-2-(4-Aminopyrrolo[2,1-f][1,2,4]triazin-7-yl) -5-(hydroxymethyl)tetrahydrofuran-3,4-diol **(113)**

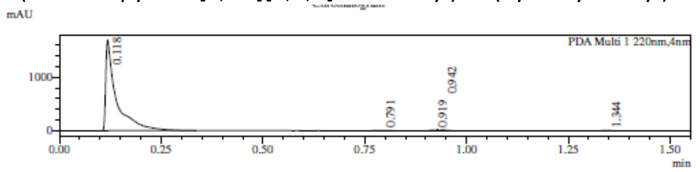

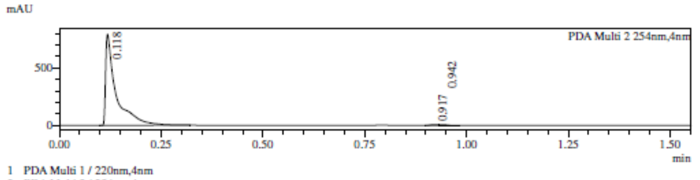

1 PDA Multi 1 / 220nm,4nm
2 PDA Multi 2 / 254nm,4nm

MS Chromatogram

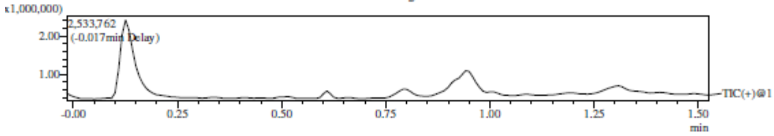

Integration Result

Peak Table

PDA Ch1 220nm

| Peak# | Ret. Time | Height | Height% | USP Width | Area | Area% |
|---|---|---|---|---|---|---|
| 1 | 0.118 | 1700627 | 97.863 | 0.036 | 3080256 | 98.324 |
| 2 | 0.791 | 6785 | 0.390 | 0.047 | 11769 | 0.376 |
| 3 | 0.919 | 11797 | 0.679 | 0.060 | 14638 | 0.467 |
| 4 | 0.942 | 15165 | 0.873 | 0.046 | 20187 | 0.644 |
| 5 | 1.344 | 3386 | 0.195 | 0.051 | 5921 | 0.189 |

Peak Table

PDA Ch2 254nm

| Peak# | Ret. Time | Height | Height% | USP Width | Area | Area% |
|---|---|---|---|---|---|---|
| 1 | 0.118 | 792180 | 98.230 | 0.036 | 1414925 | 98.783 |
| 2 | 0.917 | 6361 | 0.789 | 0.060 | 7453 | 0.520 |
| 3 | 0.942 | 7917 | 0.982 | 0.042 | 9979 | 0.697 |

Mass Spectrum

RetTime: 0.117 Datafile: D:\DATA\2022\2210\221021\EW26049-1086-P1A1.lcd

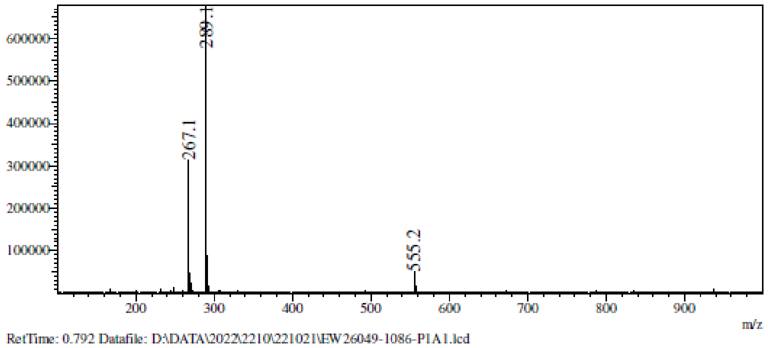

RetTime: 0.792 Datafile: D:\DATA\2022\2210\221021\EW26049-1086-P1A1.lcd

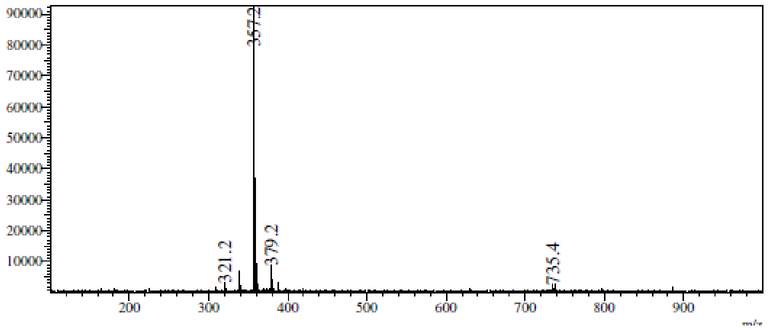

RetTime: 0.917 Datafile: D:\DATA\2022\2210\221021\EW26049-1086-P1A1.lcd

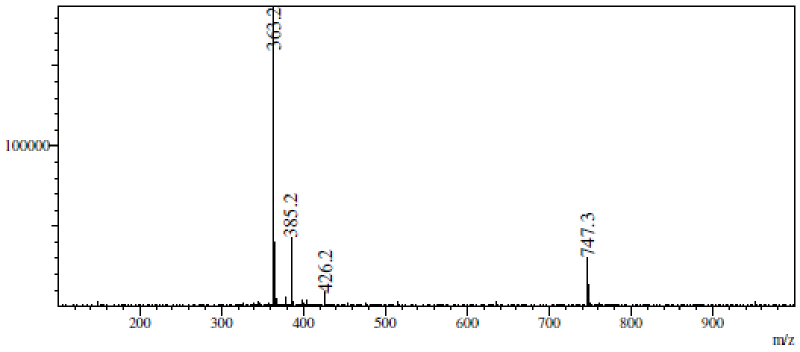

RetTime: 0.942 Datafile: D:\DATA\2022\2210\221021\EW26049-1086-P1A1.lcd

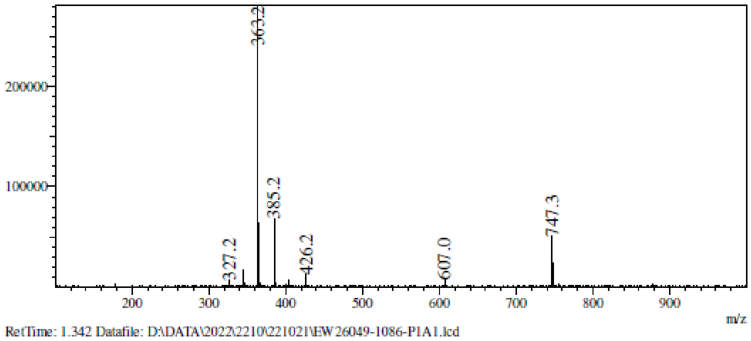

RetTime: 1.342 Datafile: D:\DATA\2022\2210\221021\EW26049-1086-P1A1.lcd

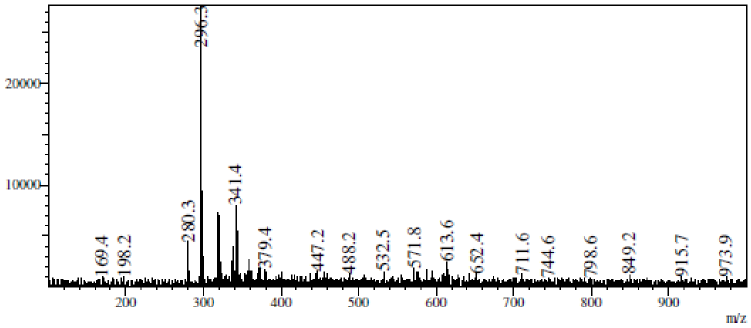

**$^1$H NMR** [(3a*S*,4*S*,6*R*,6a*R*)-4-(4-Aminopyrrolo[2,1-f][1,2,4]triazin-7-yl)-2,2 -dimethyl-3a,4,6,6a-tetrahydrofuro[3,4-d][1,3]dioxol-6-yl]methanol **(114)**

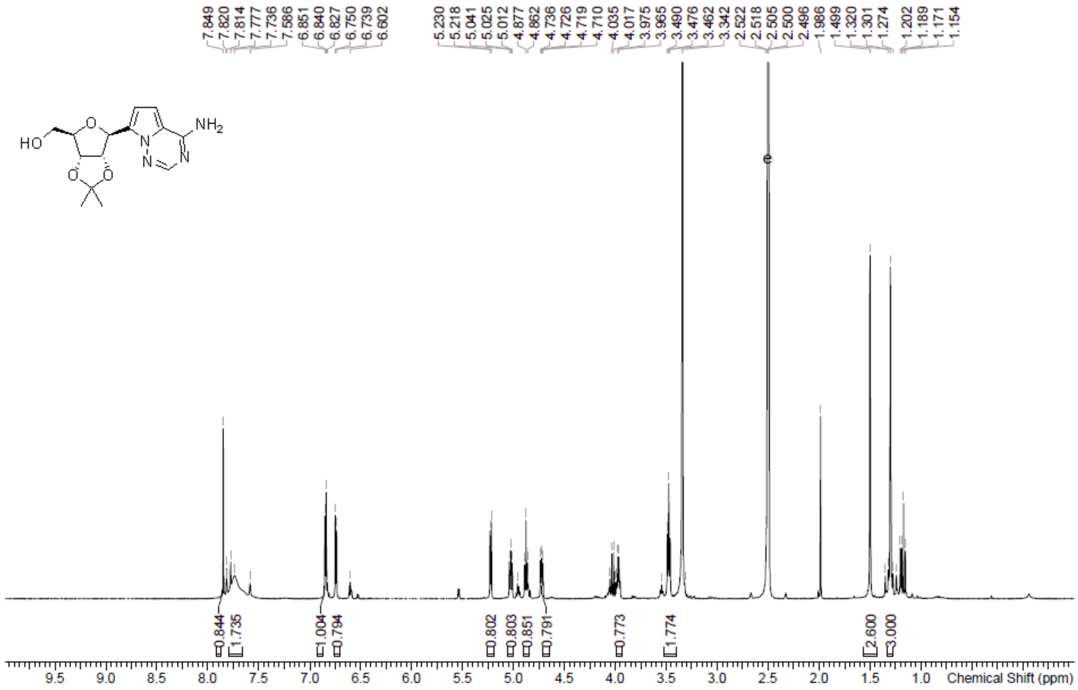

**LCMS** [(3a*S*,4*S*,6*R*,6a*R*)-4-(4-Aminopyrrolo[2,1-f][1,2,4]triazin-7-yl)-2,2 -dimethyl-3a,4,6,6a-tetrahydrofuro[3,4-d][1,3]dioxol-6-yl]methanol **(114)**

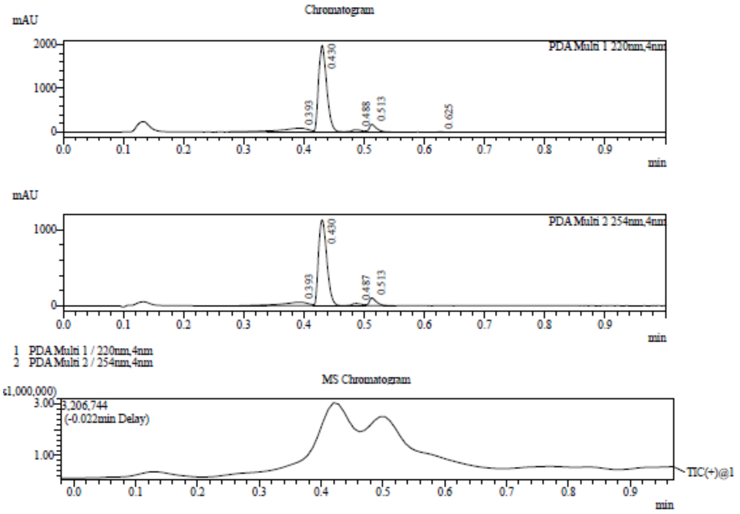

Integration Result

Peak Table

PDA Ch1 220nm

| Peak# | Ret. Time | Height | Height% | USP Width | Area | Area% |
|---|---|---|---|---|---|---|
| 1 | 0.393 | 89246 | 3.873 | 0.093 | 297792 | 12.565 |
| 2 | 0.430 | 1981009 | 85.966 | 0.027 | 1851247 | 78.110 |
| 3 | 0.488 | 53281 | 2.312 | 0.035 | 65594 | 2.768 |
| 4 | 0.513 | 173677 | 7.537 | 0.024 | 149220 | 6.296 |
| 5 | 0.625 | 7198 | 0.312 | 0.025 | 6201 | 0.262 |

Peak Table

PDA Ch2 254nm

| Peak# | Ret. Time | Height | Height% | USP Width | Area | Area% |
|---|---|---|---|---|---|---|
| 1 | 0.393 | 46381 | 3.567 | 0.078 | 150081 | 11.103 |
| 2 | 0.430 | 1123938 | 86.433 | 0.027 | 1080783 | 79.958 |
| 3 | 0.487 | 29100 | 2.238 | 0.034 | 33048 | 2.445 |
| 4 | 0.513 | 100946 | 7.763 | 0.025 | 87773 | 6.494 |

Mass Spectrum

RetTime: 0.395 Datafile: D:\DATA\2022\2210\221026\EW26049-1087-P1C1.lcd

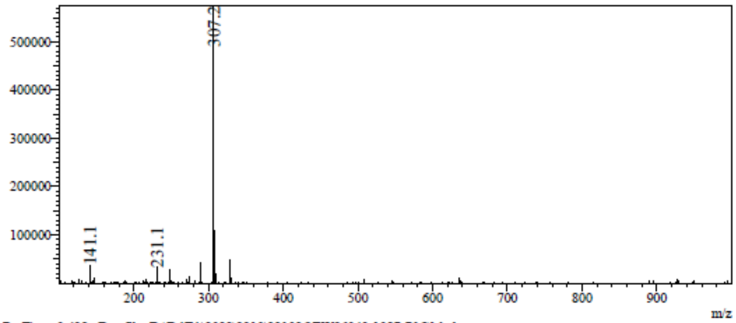

RetTime: 0.428 Datafile: D:\DATA\2022\2210\221026\EW26049-1087-P1C1.lcd

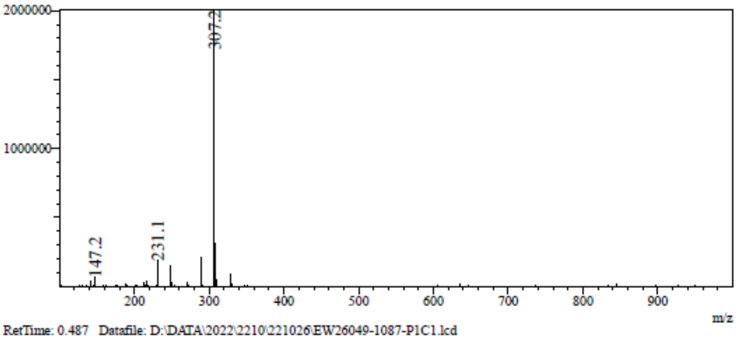

RetTime: 0.487 Datafile: D:\DATA\2022\2210\221026\EW26049-1087-P1C1.lcd

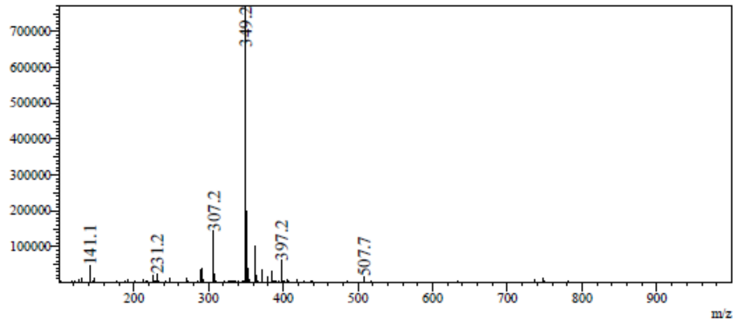

RetTime: 0.512 Datafile: D:\DATA\2022\2210\221026\EW26049-1087-P1C1.lcd

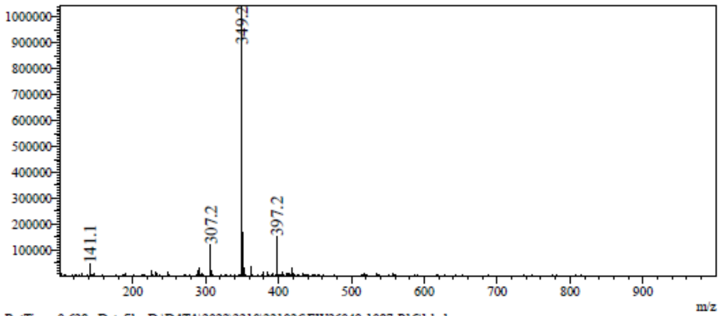

RetTime: 0.628 Datafile: D:\DATA\2022\2210\221026\EW26049-1087-P1C1.lcd

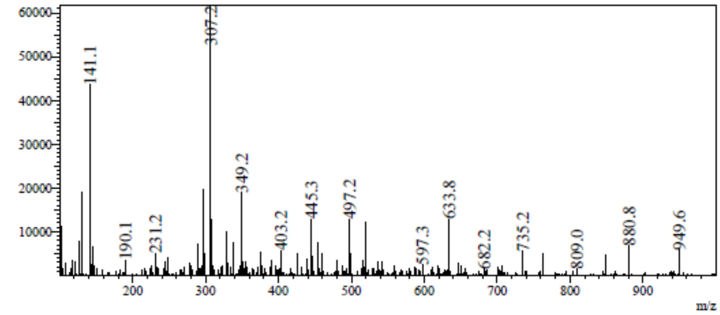

**LCMS** 7-[(3a*S*,4*S*,6*R*,6a*R*)-6-(Azidomethyl)-2,2-dimethyl-3a,4,6,6a-tetrah ydrofuro[3,4-d][1,3]dioxol-4-yl]pyrrolo[2,1-f][1,2,4]triazin-4-amine **(115)**

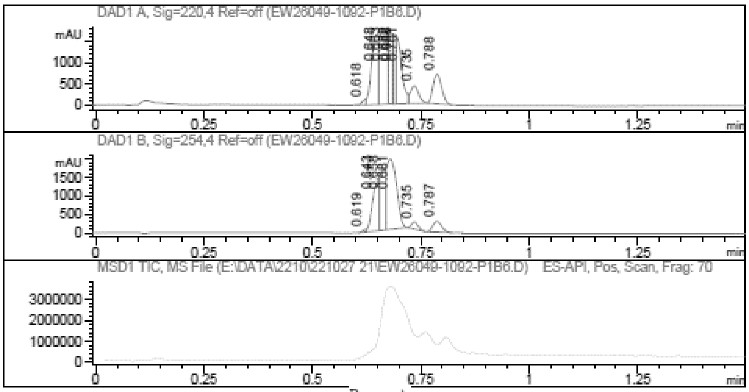

Report

Signal 1 : DAD1 A, Sig=220,4 Ref=off

| # | Meas. Ret. | Height | Width | Area | Area % |
|---|---|---|---|---|---|
| 1 | 0.618 | 106.373 | 0.011 | 85.019 | 0.964 |
| 2 | 0.648 | 1637.115 | 0.017 | 1792.176 | 20.330 |
| 3 | 0.663 | 1720.668 | 0.016 | 2162.343 | 24.529 |
| 4 | 0.682 | 1674.983 | 0.009 | 1145.922 | 12.999 |
| 5 | 0.688 | 1647.396 | 0.007 | 653.476 | 7.413 |
| 6 | 0.701 | 1309.991 | 0.019 | 1464.338 | 16.611 |
| 7 | 0.735 | 421.110 | 0.020 | 546.329 | 6.198 |
| 8 | 0.788 | 694.686 | 0.023 | 965.701 | 10.955 |

Signal 2 : DAD1 B, Sig=254,4 Ref=off

| # | Meas. Ret. | Height | Width | Area | Area % |
|---|---|---|---|---|---|
| 1 | 0.619 | 53.908 | 0.010 | 35.445 | 0.541 |
| 2 | 0.643 | 1081.293 | 0.019 | 1395.774 | 21.306 |
| 3 | 0.658 | 1547.609 | 0.016 | 1446.321 | 22.078 |
| 4 | 0.681 | 1913.257 | 0.025 | 3061.511 | 46.733 |
| 5 | 0.735 | 194.248 | 0.017 | 202.960 | 3.098 |
| 6 | 0.787 | 293.950 | 0.022 | 409.085 | 6.245 |

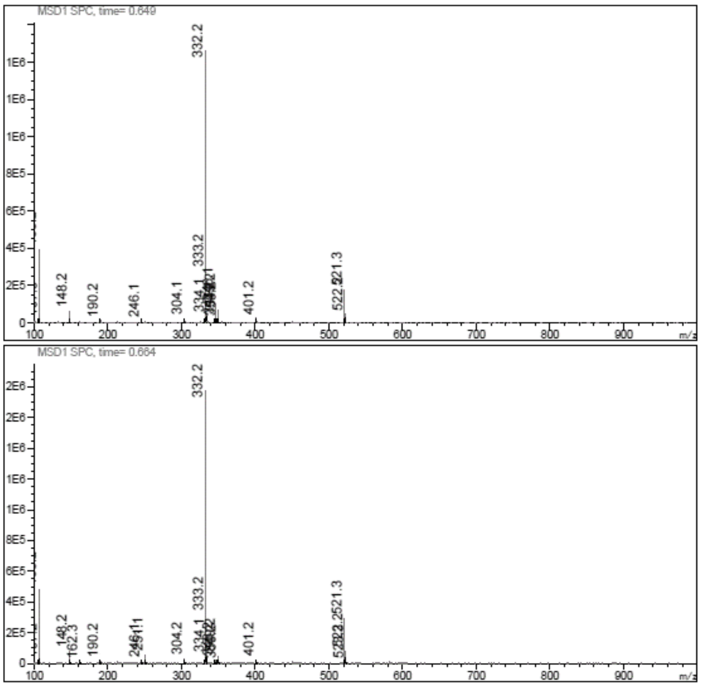

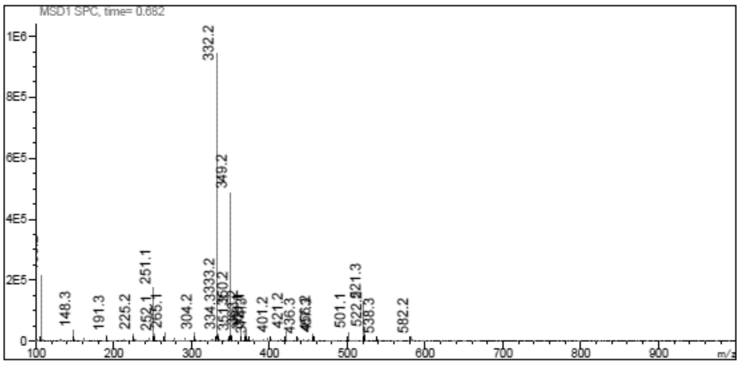
MSD1 SPC, time= 0.682
332.2
349.2
251.1
148.3
191.3
225.2
304.2
401.2
421.2
501.1
538.3
582.2
m/z

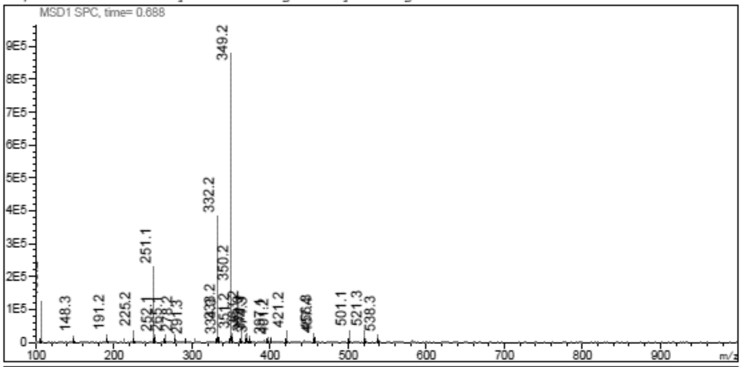
MSD1 SPC, time= 0.688
349.2
332.2
251.1
350.2
148.3
191.2
225.2
421.2
501.1
521.3
538.3
m/z

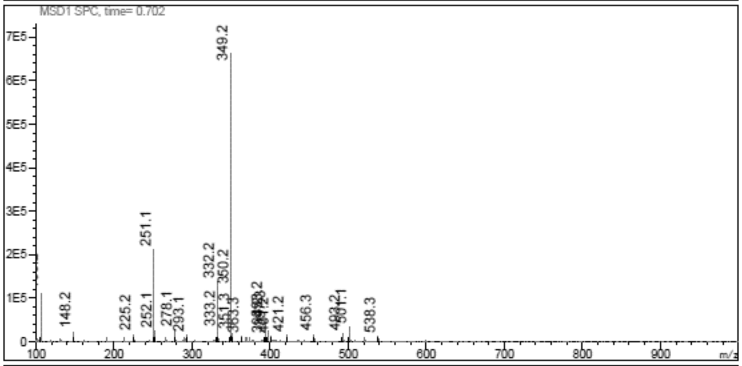
MSD1 SPC, time= 0.702
349.2
251.1
332.2
350.2
148.2
225.2
252.1
278.1
293.1
333.2
421.2
456.3
538.3
m/z

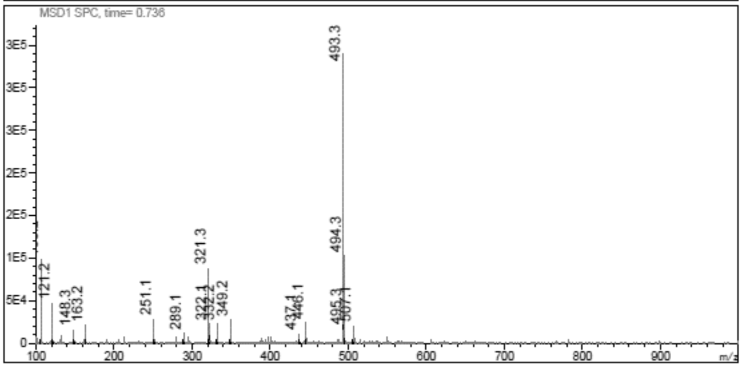
MSD1 SPC, time= 0.736
493.3
494.3
321.3
121.2
148.3
163.2
251.1
289.1
349.2
437.1
m/z

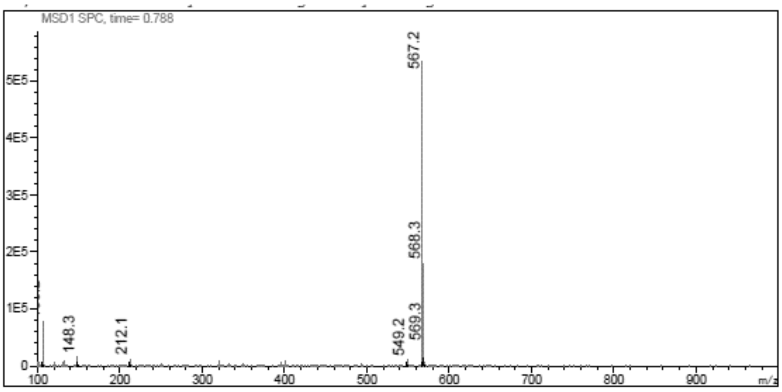

**LCMS** 7-[(3a*S*,4*S*,6*R*,6a*R*)-6-(Aminomethyl)-2,2-dimethyl-3a,4,6,6a-tetrahydrofuro[3,4-d][1,3]dioxol-4-yl]pyrrolo[2,1-f][1,2,4]triazin-4-amine **(116)**

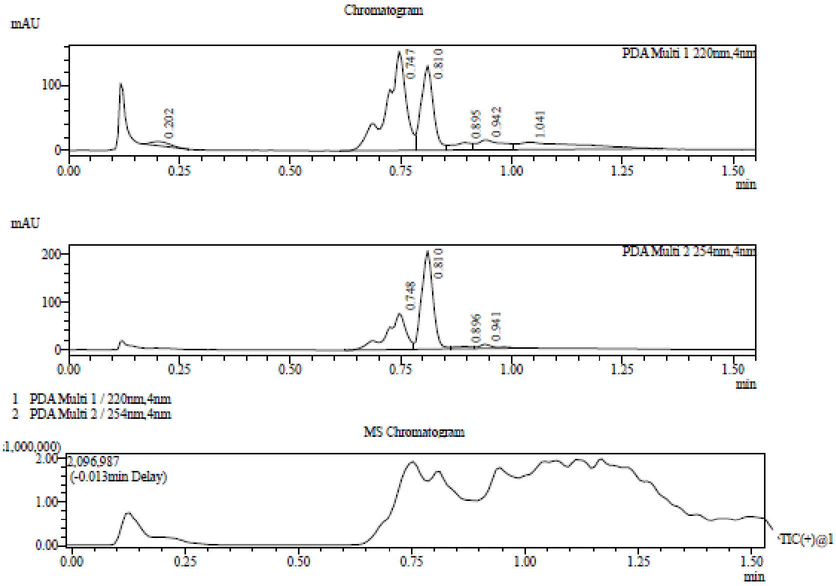

Integration Result

Peak Table

PDA Ch1 220nm

| Peak# | Ret. Time | Height | Height% | USP Width | Area | Area% |
|---|---|---|---|---|---|---|
| 1 | 0.202 | 6335 | 1.933 | 0.082 | 17758 | 1.876 |
| 2 | 0.747 | 152720 | 46.593 | 0.063 | 477623 | 50.468 |
| 3 | 0.810 | 130811 | 39.909 | 0.059 | 241949 | 25.565 |
| 4 | 0.895 | 11397 | 3.477 | 0.189 | 33166 | 3.504 |
| 5 | 0.942 | 15254 | 4.654 | 0.137 | 61302 | 6.477 |
| 6 | 1.041 | 11256 | 3.434 | 0.507 | 114598 | 12.109 |

Peak Table

PDA Ch2 254nm

| Peak# | Ret. Time | Height | Height% | USP Width | Area | Area% |
|---|---|---|---|---|---|---|
| 1 | 0.748 | 75044 | 25.486 | 0.063 | 227406 | 36.494 |
| 2 | 0.810 | 206166 | 70.016 | 0.057 | 363034 | 58.260 |

Mass Spectrum

RetTime: 0.203 Datafile: D:\DATA\2022\2210\221031\EW26049-1093-P1S1.lcd

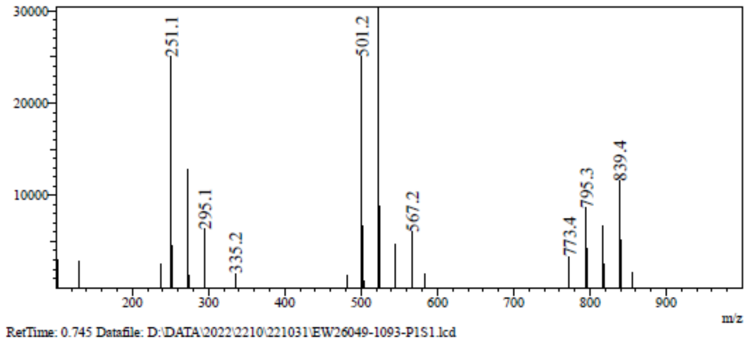

RetTime: 0.745 Datafile: D:\DATA\2022\2210\221031\EW26049-1093-P1S1.lcd

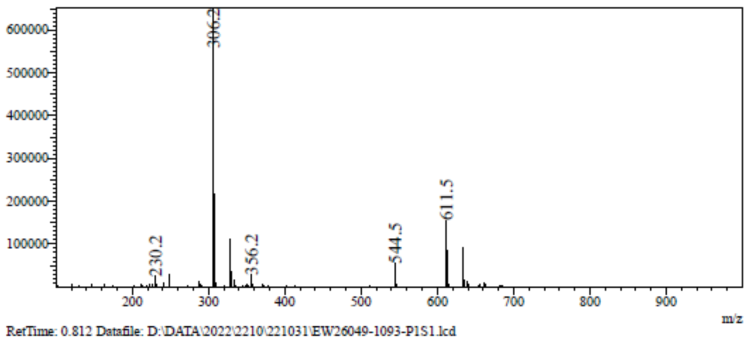

RetTime: 0.812 Datafile: D:\DATA\2022\2210\221031\EW26049-1093-P1S1.lcd

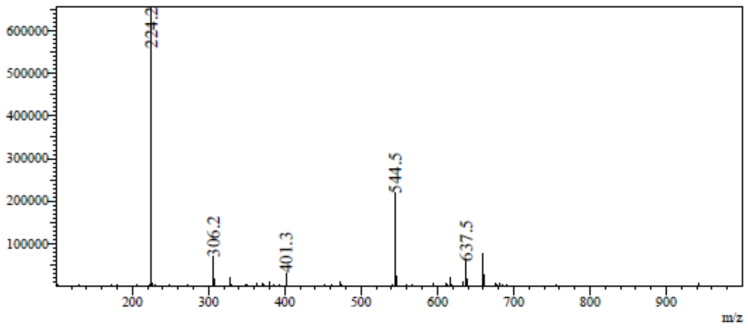

RetTime: 0.895 Datafile: D:\DATA\2022\2210\221031\EW26049-1093-P1S1.lcd

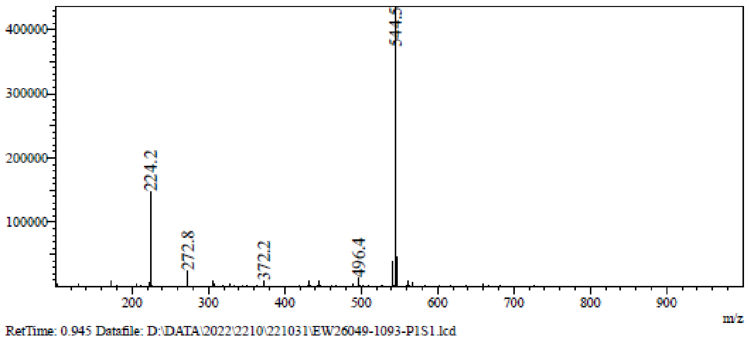

RetTime: 0.945 Datafile: D:\DATA\2022\2210\221031\EW26049-1093-P1S1.lcd

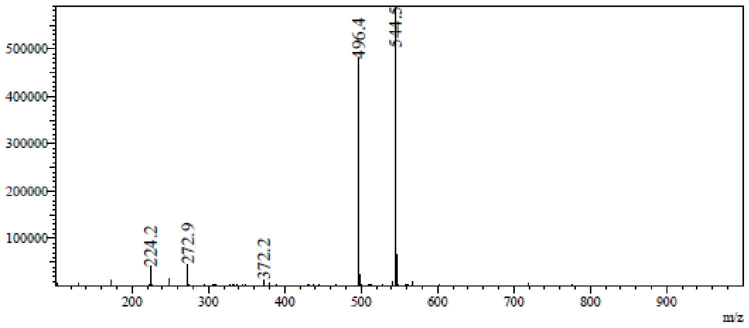

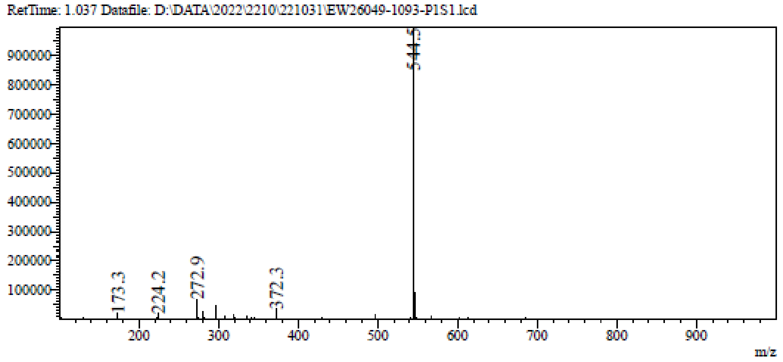

**LCMS** (2*R*,3*S*,4*R*,5*S*)-2-(Aminomethyl)-5-(4-aminopyrrolo[2,1-f][1,2,4] triazin-7-yl)tetrahydrofuran-3,4-diol **(117)**

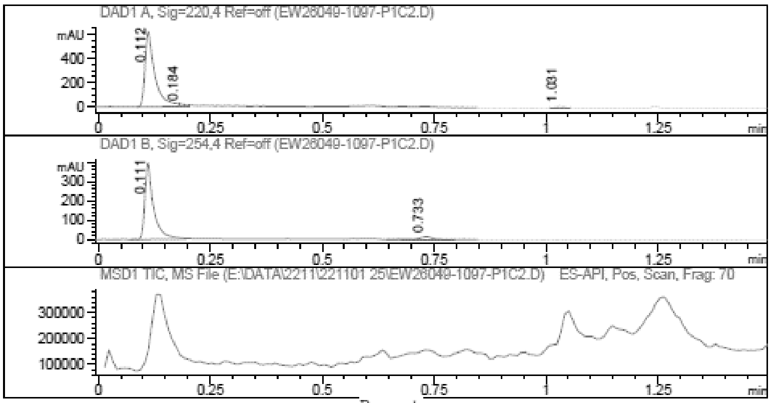

Report

=================================================================

Signal 1 : DAD1 A, Sig=220,4 Ref=off

| # | Meas. Ret. | Height | Width | Area | Area % |
|---|---|---|---|---|---|
| 1 | 0.112 | 619.410 | 0.022 | 946.249 | 95.803 |
| 2 | 0.184 | 19.870 | 0.012 | 17.703 | 1.792 |
| 3 | 1.031 | 19.416 | 0.020 | 23.747 | 2.404 |

Signal 2 : DAD1 B, Sig=254,4 Ref=off

| # | Meas. Ret. | Height | Width | Area | Area % |
|---|---|---|---|---|---|
| 1 | 0.111 | 396.627 | 0.019 | 541.895 | 92.847 |
| 2 | 0.733 | 16.556 | 0.035 | 41.748 | 7.153 |

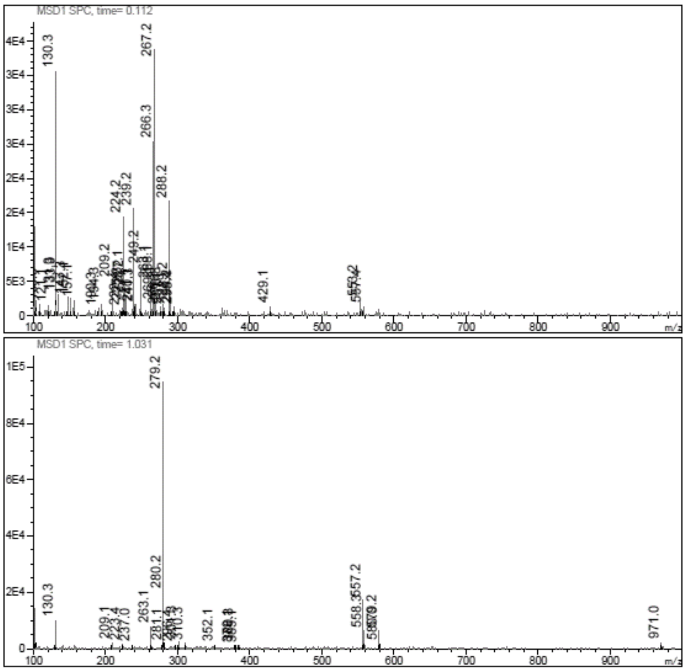

**¹H NMR** *N*-[[(2*R*,3*S*,4*R*,5*S*)-5-(4-Aminopyrrolo[2,1-f][1,2,4]triazin-7-yl)-3,4- dihydroxy-tetrahydrofuran-2-yl]methyl]naphthalene-2-sulfonamide **(34)**

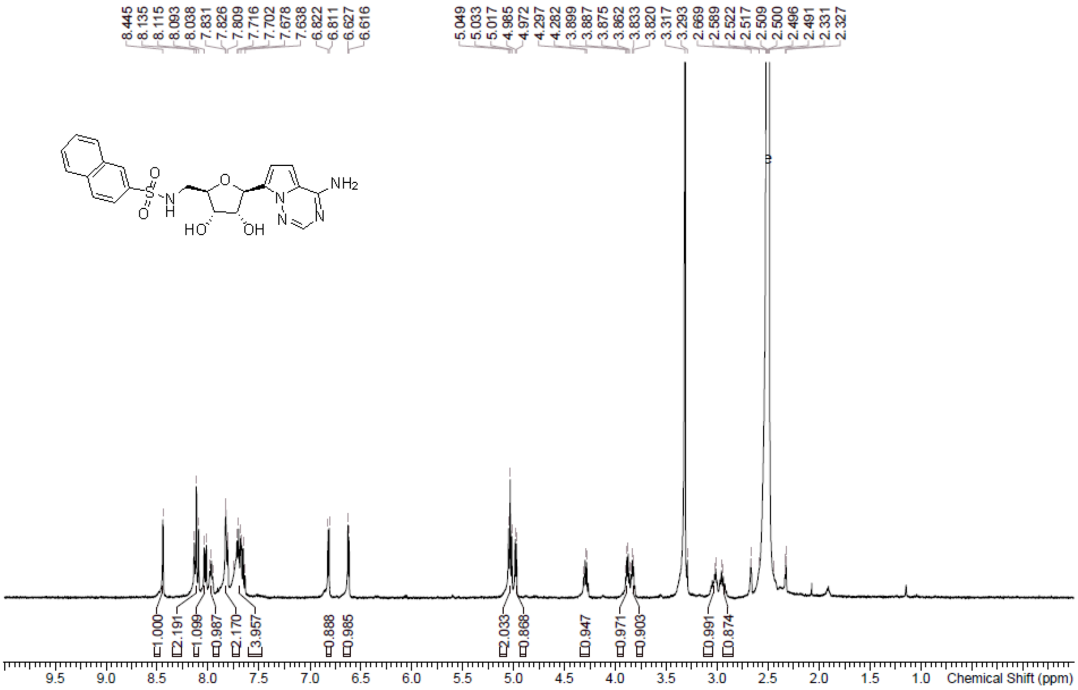

**HPLC** *N*-[[(2*R*,3*S*,4*R*,5*S*)-5-(4-Aminopyrrolo[2,1-f][1,2,4]triazin-7-yl)-3,4- dihydroxy-tetrahydrofuran-2-yl]methyl]naphthalene-2-sulfonamide **(34)**

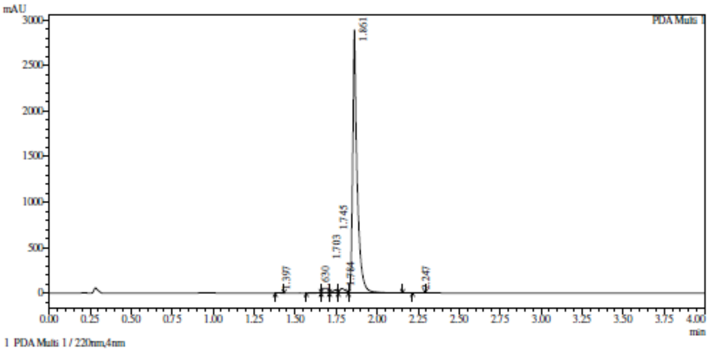

Integration result

PeakTable

PDA Ch1 220nm

| Peak# | Ret. Time | USP Width | Resolution | Height | Area | Area % |
|---|---|---|---|---|---|---|
| 1 | 1.397 | 0.043 | 0.000 | 702 | 1066 | 0.019 |
| 2 | 1.630 | 0.125 | 2.792 | 3571 | 10988 | 0.193 |
| 3 | 1.703 | 0.535 | 0.220 | 7029 | 14769 | 0.260 |
| 4 | 1.745 | 0.079 | 0.137 | 33177 | 68410 | 1.203 |
| 5 | 1.784 | 0.079 | 0.498 | 49412 | 125124 | 2.199 |
| 6 | 1.861 | 0.045 | 1.236 | 2893192 | 5466691 | 96.094 |
| 7 | 2.247 | 0.066 | 6.991 | 796 | 1847 | 0.032 |
| Total | | | | 2987881 | 5688896 | 100.000 |

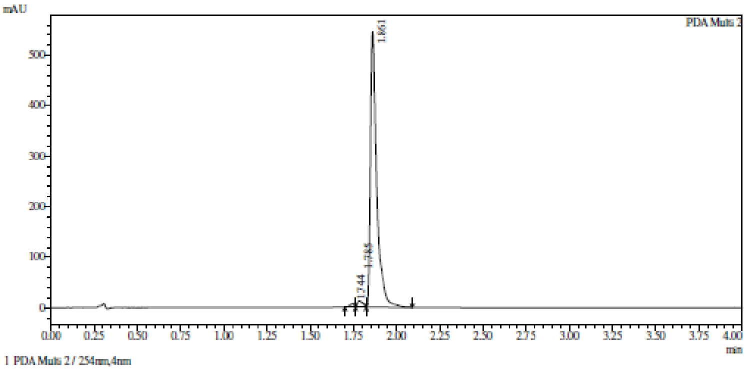

Integration result

PeakTable

PDA Ch2 254nm

| Peak# | Ret. Time | USP Width | Resolution | Height | Area | Area % |
|---|---|---|---|---|---|---|
| 1 | 1.744 | 0.079 | 0.000 | 6956 | 13230 | 0.935 |
| 2 | 1.785 | 0.079 | 0.506 | 12972 | 33163 | 2.345 |
| 3 | 1.861 | 0.060 | 1.090 | 545778 | 1367928 | 96.720 |
| Total | | | | 565707 | 1414320 | 100.000 |

**HRMS** *N*-[[(2*R*,3*S*,4*R*,5*S*)-5-(4-Aminopyrrolo[2,1-f][1,2,4]triazin-7-yl)-3,4- dihydroxy-tetrahydrofuran-2-yl]methyl]naphthalene-2-sulfonamide **(34)**

**Compound Table**

| Label | Tgt Score | Mass Error (ppm) | Tgt Formula | Obs. RT | Ref. Mass | Obs. Mass |
|---|---|---|---|---|---|---|
| Cpd 1: C21 H21 N5 O5 S; 1.763 | 99.38 | -2.2 | C21 H21 N5 O5 S | 1.763 | 455.1263 | 455.1253 |

| Obs. *m/z* | Obs. RT | Obs. Mass | Tgt Formula | Tgt Mass | Tgt Mass Error (ppm) | Find Cpds Algorithm |
|---|---|---|---|---|---|---|
| 933.2426 | 1.763 | 455.1253 | C21 H21 N5 O5 S | 455.1263 | -2.2 | Find by Formula |

**Compound Chromatograms**

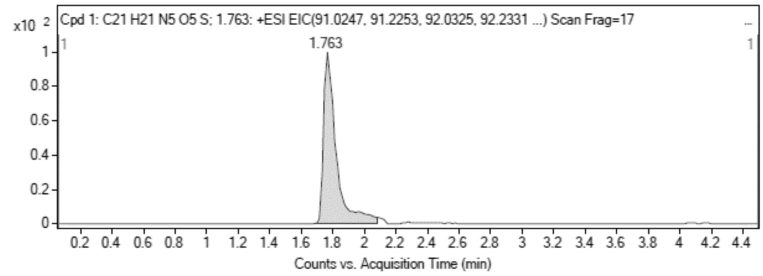

**MS Zoomed Spectrum**

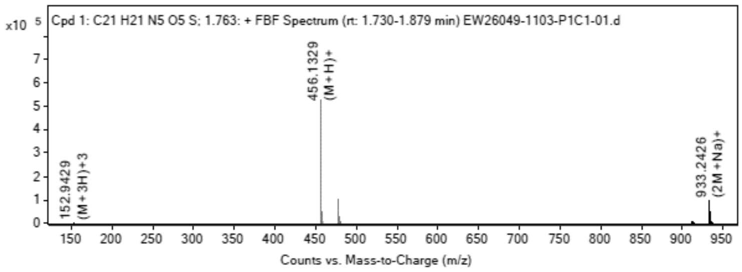

**MS Spectrum Peak List**

| Obs. *m/z* | Charge | Abund | Ion/Isotope |
|---|---|---|---|
| 456.1329 | 1 | 525477.63 | (M+H)+ |
| 457.1349 | 1 | 141160.08 | (M+H)+ |
| 458.1321 | 1 | 43342.77 | (M+H)+ |
| 478.1134 | 1 | 104224.06 | (M+Na)+ |
| 479.1154 | 1 | 27611.89 | (M+Na)+ |
| 480.1133 | 1 | 8924.78 | (M+Na)+ |
| 933.2426 | 1 | 101196.81 | (2M+Na)+ |
| 934.2447 | 1 | 51909.96 | (2M+Na)+ |
| 935.2432 | 1 | 23879.92 | (2M+Na)+ |
| 936.2436 | 1 | 8033.02 | (2M+Na)+ |

**MS Zoomed Spectrum**

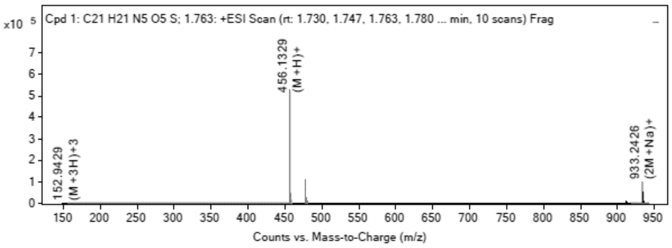

**MS Spectrum Peak List**

| Obs. *m/z* | Charge | Abund | Ion/Isotope | Tgt Mass Error (ppm) |
|---|---|---|---|---|
| 456.1329 | 1 | 525477.63 | (M+H)+ | 1.59 |
| 457.1349 | 1 | 141160.08 | (M+H)+ | 3.46 |
| 458.1321 | 1 | 43342.77 | (M+H)+ | 4.23 |
| 478.1134 | 1 | 104224.06 | (M+Na)+ | 4.52 |
| 479.1154 | 1 | 27611.89 | (M+Na)+ | 6.23 |
| 480.1133 | 1 | 8924.78 | (M+Na)+ | 5.57 |
| 933.2426 | 1 | 101196.81 | (2M+Na)+ | -0.79 |
| 934.2447 | 1 | 51909.96 | (2M+Na)+ | 0.01 |
| 935.2432 | 1 | 23879.92 | (2M+Na)+ | 0.62 |
| 936.2436 | 1 | 8033.02 | (2M+Na)+ | 0.84 |

--- End Of Report ---

**¹H NMR** 7-[(2*S*,3*S*,4*R*,5*R*)-3,4-Bis[[tert-butyl(dimethyl)silyl]oxy]-5-[[tert-butyl(dimethyl)silyl]oxymethyl]tetrahydrofuran-2-yl]pyrrolo[2,1-f][1,2,4]triazin-4-amine **(118)**

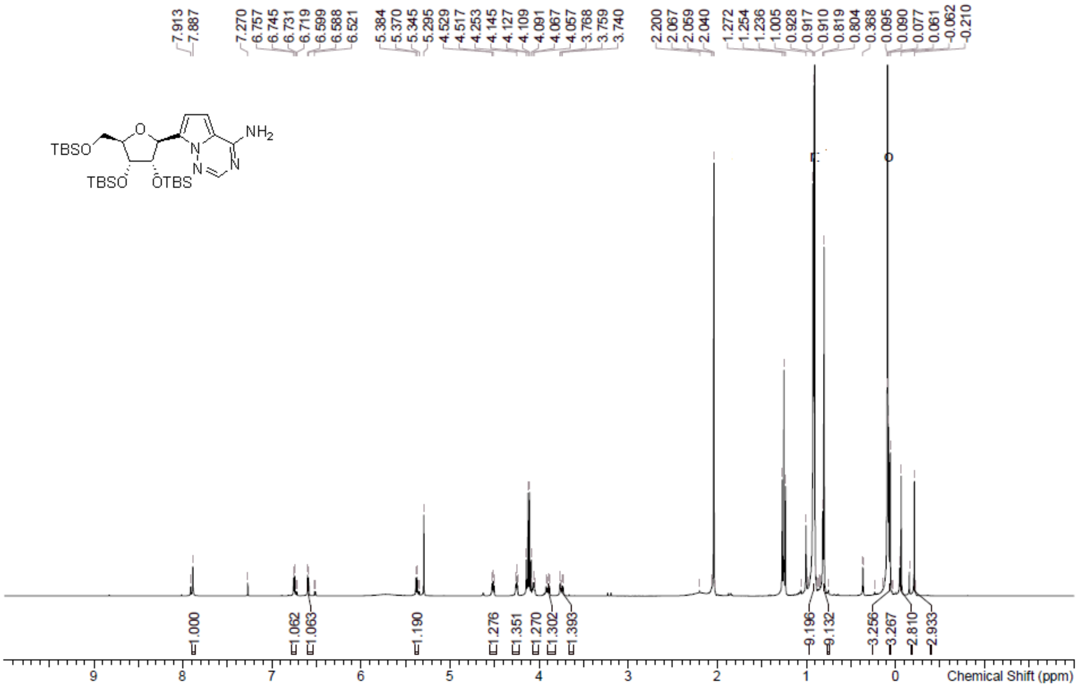

**LCMS** 7-[(2*S*,3*S*,4*R*,5*R*)-3,4-Bis[[tert-butyl(dimethyl)silyl]oxy]-5-[[tert-butyl(dimethyl)silyl]oxymethyl]tetrahydrofuran-2-yl]pyrrolo[2,1-f][1,2,4]triazin-4-amine **(118)**

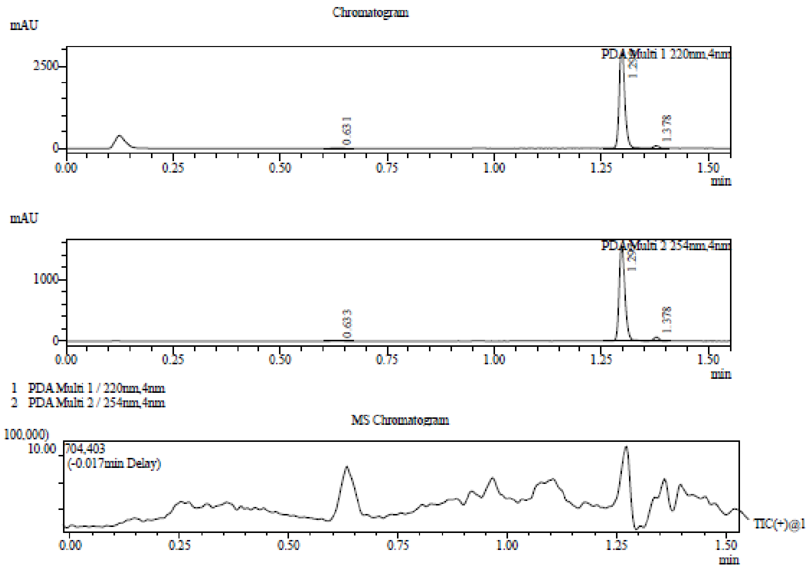

Integration Result

Peak Table

PDA Ch1 220nm

| Peak# | Ret. Time | Height | Height% | USP Width | Area | Area% |
|---|---|---|---|---|---|---|
| 1 | 0.631 | 5178 | 0.173 | 0.052 | 9101 | 0.324 |
| 2 | 1.299 | 2916589 | 97.511 | 0.025 | 2735515 | 97.409 |
| 3 | 1.378 | 69272 | 2.316 | 0.026 | 63658 | 2.267 |

Peak Table

PDA Ch2 254nm

| Peak# | Ret. Time | Height | Height% | USP Width | Area | Area% |
|---|---|---|---|---|---|---|
| 1 | 0.633 | 3275 | 0.202 | 0.047 | 5580 | 0.366 |
| 2 | 1.297 | 1556053 | 96.100 | 0.027 | 1464576 | 96.159 |
| 3 | 1.378 | 59874 | 3.698 | 0.025 | 52920 | 3.475 |

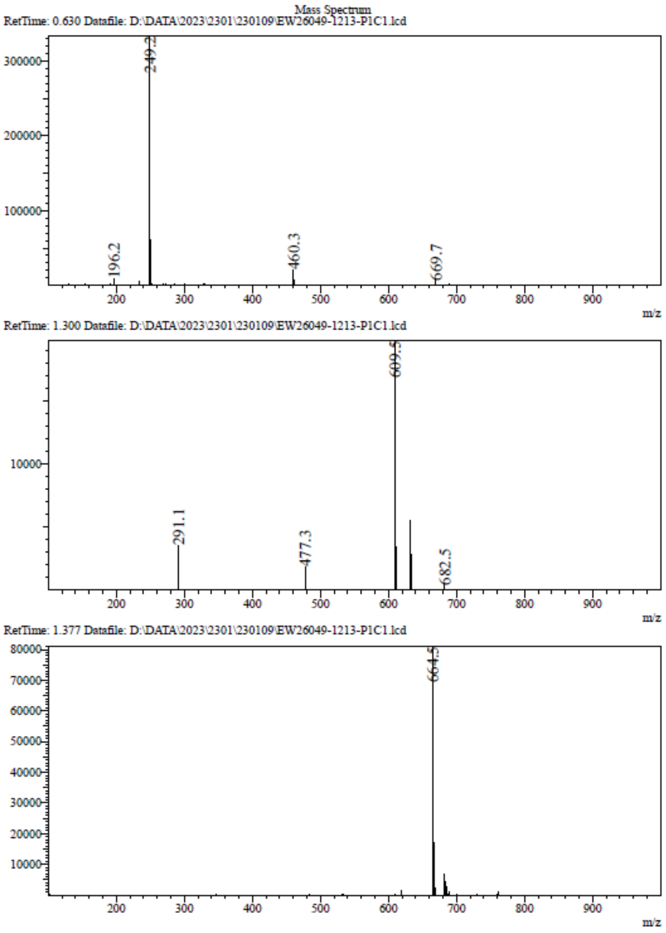

**$^1$H NMR** [(2*R*,3*R*,4*S*,5*S*)-5-(4-Aminopyrrolo[2,1-f][1,2,4]triazin-7-yl) -3,4-bis[[tert-butyl(dimethyl)silyl]oxy]tetrahydrofuran-2-yl]methanol **(119)**

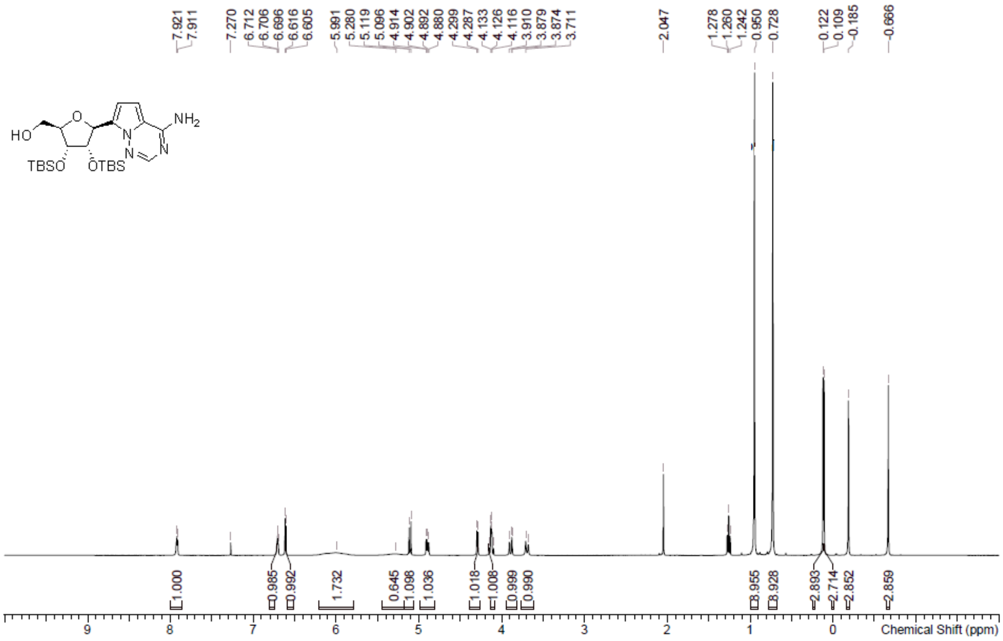

**LCMS** [(2*R*,3*R*,4*S*,5*S*)-5-(4-Aminopyrrolo[2,1-f][1,2,4]triazin-7-yl) -3,4-bis[[tert-butyl(dimethyl)silyl]oxy]tetrahydrofuran-2-yl]methanol **(119)**

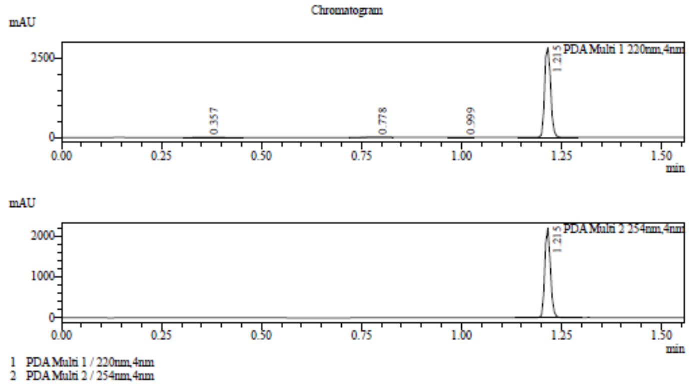

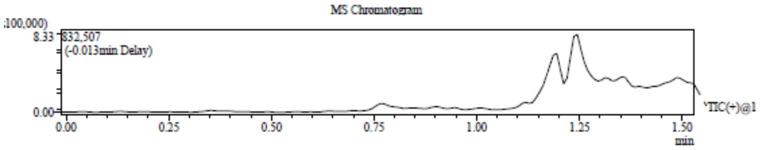

Integration Result

Peak Table

PDA Ch1 220nm

| Peak# | Ret. Time | Height | Height% | USP Width | Area | Area% |
|---|---|---|---|---|---|---|
| 1 | 0.357 | 3051 | 0.107 | 0.107 | 11086 | 0.366 |
| 2 | 0.778 | 2858 | 0.100 | 0.041 | 5460 | 0.180 |
| 3 | 0.999 | 3307 | 0.116 | 0.052 | 5484 | 0.181 |
| 4 | 1.215 | 2838256 | 99.676 | 0.032 | 3006108 | 99.272 |

Peak Table

PDA Ch2 254nm

| Peak# | Ret. Time | Height | Height% | USP Width | Area | Area% |
|---|---|---|---|---|---|---|
| 1 | 1.215 | 2193116 | 100.000 | 0.030 | 2192481 | 100.000 |

Mass Spectrum

RetTime: 0.353 Datafile: D:\DATA\2023\2301\230111\EW26049-1216-P1C1.lcd

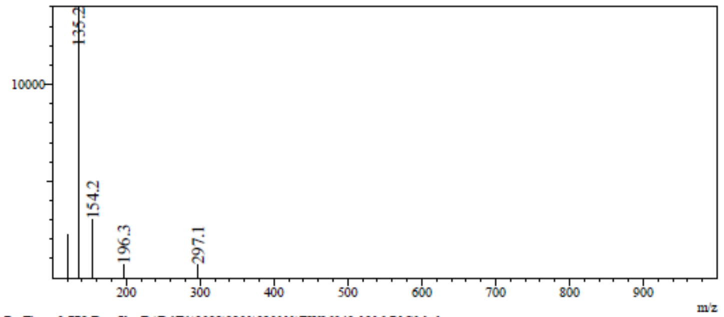

RetTime: 0.778 Datafile: D:\DATA\2023\2301\230111\EW26049-1216-P1C1.lcd

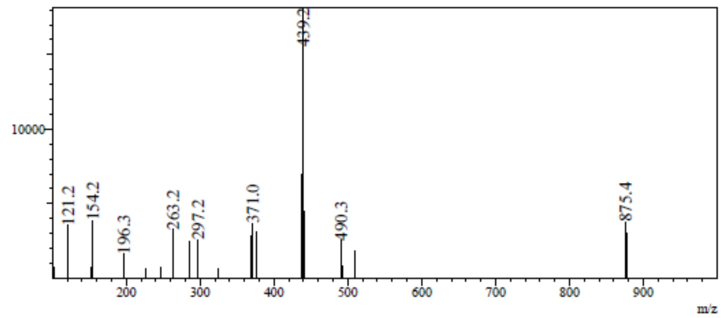

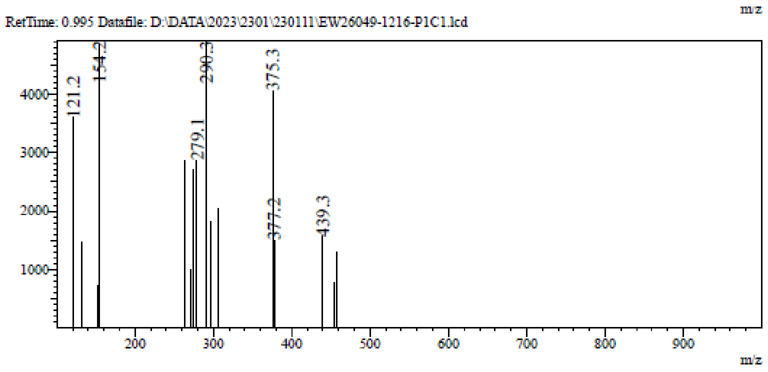

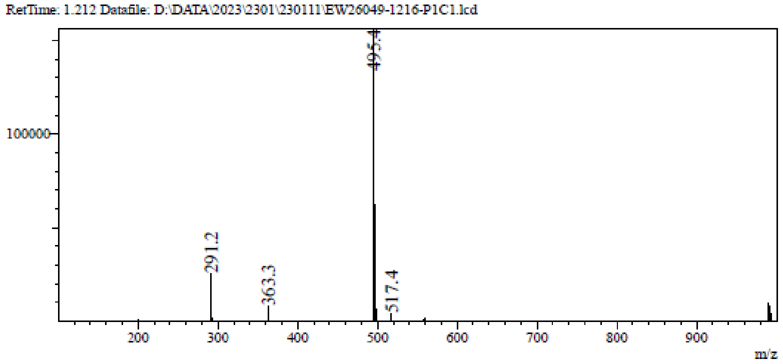

**$^{1}$H NMR** [(2*R*,3*R*,4*S*,5*S*)-5-(4-Aminopyrrolo[2,1-f][1,2,4]triazin-7-yl)-3,4-bis [[tert-butyl(dimethyl)silyl]oxy]tetrahydrofuran-2-yl]methyl methanesulfonate **(120)**

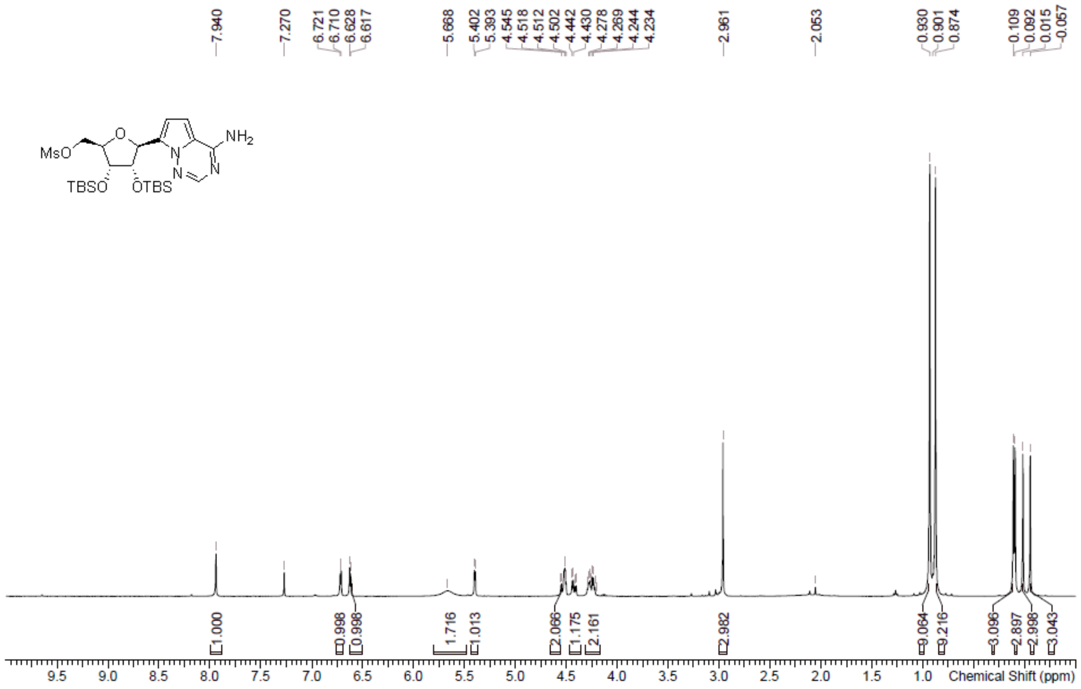

**LCMS** [(2*R*,3*R*,4*S*,5*S*)-5-(4-Aminopyrrolo[2,1-f][1,2,4]triazin-7-yl)-3,4-bis [[tert-butyl(dimethyl)silyl]oxy]tetrahydrofuran-2-yl]methyl methanesulfonate **(120)**

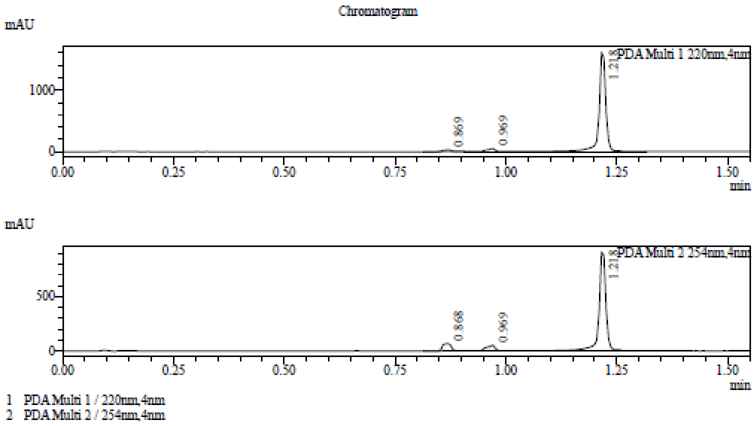

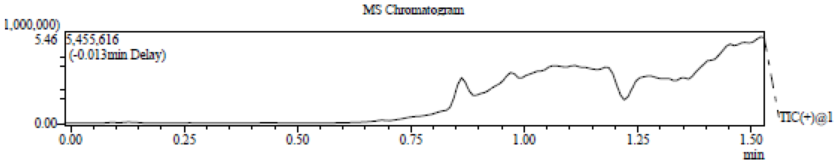

Integration Result

Peak Table

PDA Ch1 220nm

| Peak# | Ret. Time | Height | Height% | USP Width | Area | Area% |
|---|---|---|---|---|---|---|
| 1 | 0.869 | 26688 | 1.585 | 0.038 | 37424 | 2.091 |
| 2 | 0.969 | 49083 | 2.914 | 0.081 | 66412 | 3.710 |
| 3 | 1.218 | 1608495 | 95.501 | 0.201 | 1686154 | 94.199 |

Peak Table

PDA Ch2 254nm

| Peak# | Ret. Time | Height | Height% | USP Width | Area | Area% |
|---|---|---|---|---|---|---|
| 1 | 0.868 | 68327 | 6.610 | 0.036 | 92225 | 8.016 |
| 2 | 0.969 | 52250 | 5.055 | 0.076 | 68228 | 5.930 |
| 3 | 1.218 | 913086 | 88.335 | 0.210 | 990033 | 86.053 |

Mass Spectrum

RetTime: 0.870 Datafile: D:\DATA\2023\2301\230113\EW26049-1221-P1S1.lcd

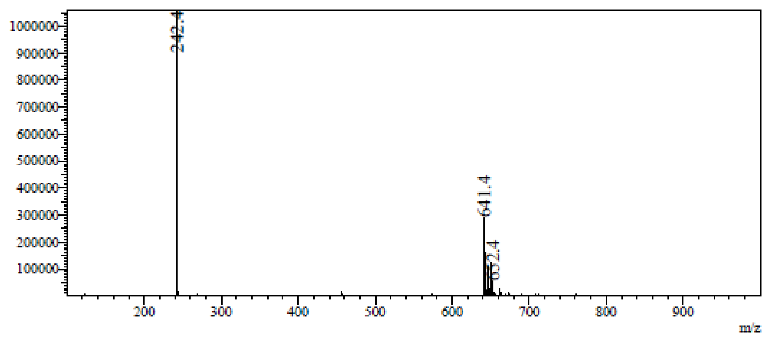

RetTime: 0.970 Datafile: D:\DATA\2023\2301\230113\EW26049-1221-P1S1.lcd

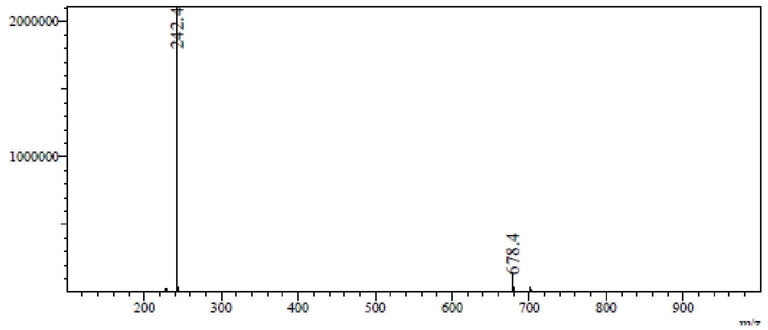

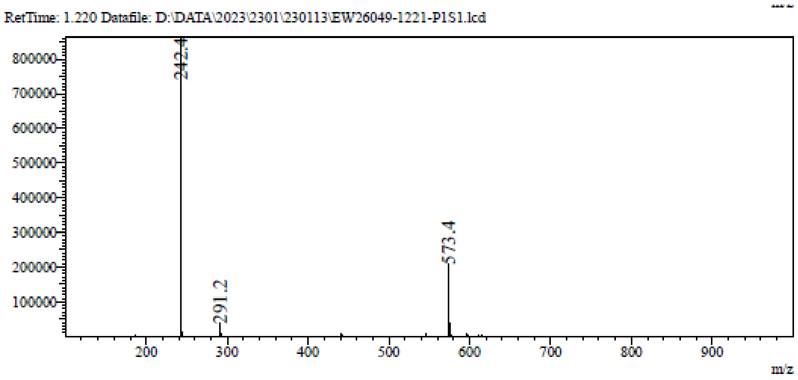

[1]**H NMR** 7-[(2*S*,3*S*,4*R*,5*S*)-5-(1,3-Benzothiazol-2-ylsulfanylmethyl) -3,4-bis[[tert-butyl(dimethyl)silyl]oxy]tetrahydrofuran-2-yl]pyrrolo[2,1-f][1,2,4]triazin-4-amine **(121a)**

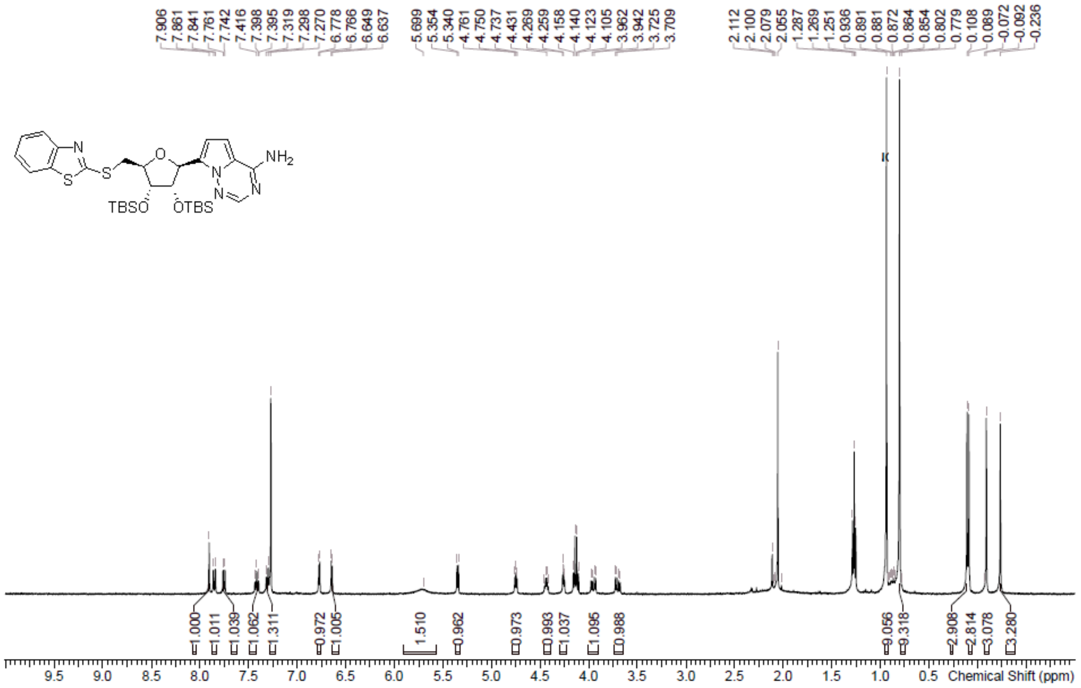

**LCMS** 7-[(2*S*,3*S*,4*R*,5*S*)-5-(1,3-Benzothiazol-2-ylsulfanylmethyl) -3,4-bis[[tert-butyl(dimethyl)silyl]oxy]tetrahydrofuran-2-yl]pyrrolo[2,1-f][1,2,4]triazin-4-amine **(121a)**

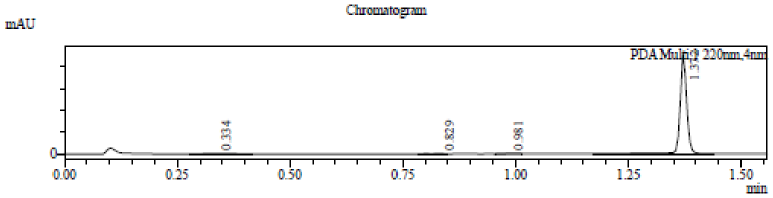

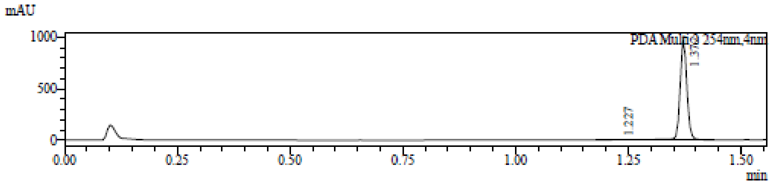

1 PDA Multi 1 / 220nm,4nm
2 PDA Multi 2 / 254nm,4nm

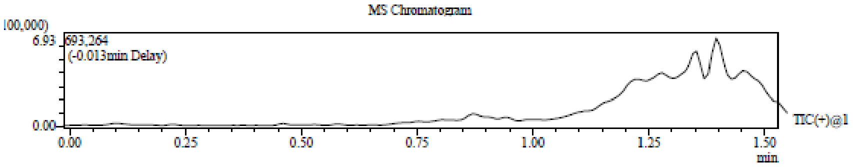

Integration Result

Peak Table

PDA Ch1 220nm

| Peak# | Ret. Time | Height | Height% | USP Width | Area | Area% |
|---|---|---|---|---|---|---|
| 1 | 0.334 | 5086 | 0.217 | 0.102 | 18409 | 0.774 |
| 2 | 0.829 | 2309 | 0.099 | 0.277 | 5476 | 0.230 |
| 3 | 0.981 | 7994 | 0.341 | 0.034 | 9188 | 0.387 |
| 4 | 1.372 | 2325600 | 99.343 | 0.199 | 2343870 | 98.609 |

Peak Table

PDA Ch2 254nm

| Peak# | Ret. Time | Height | Height% | USP Width | Area | Area% |
|---|---|---|---|---|---|---|
| 1 | 1.227 | 2675 | 0.271 | 0.000 | 5723 | 0.540 |
| 2 | 1.372 | 984651 | 99.729 | 0.147 | 1053628 | 99.460 |

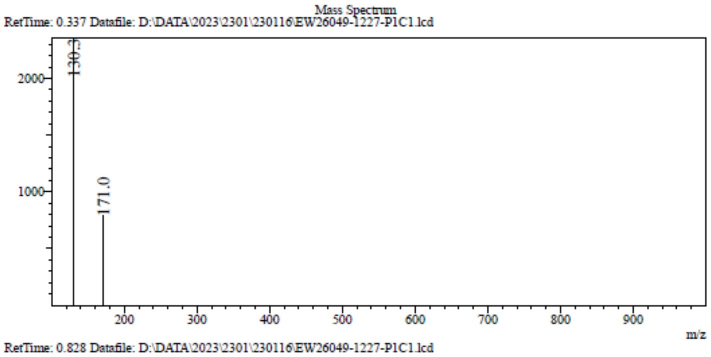

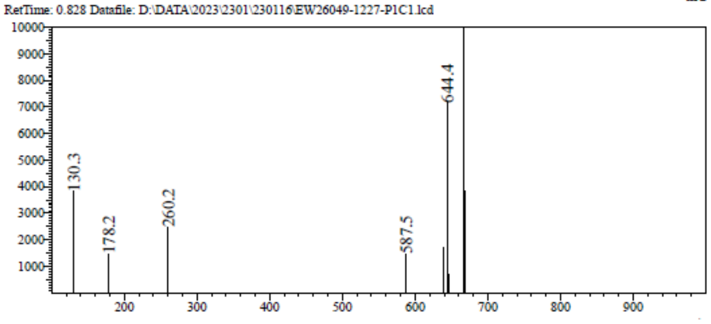

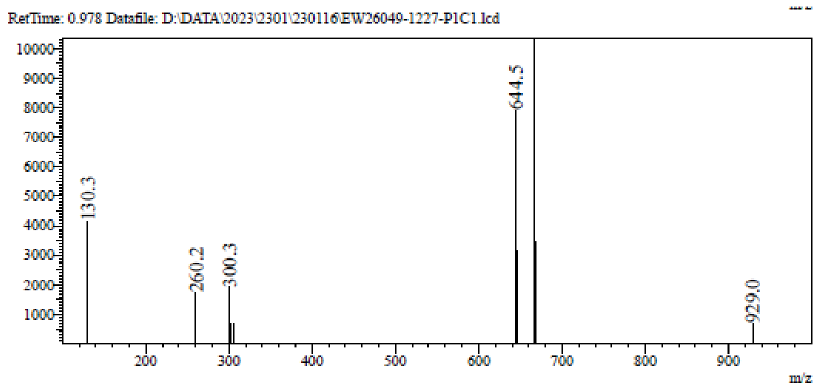

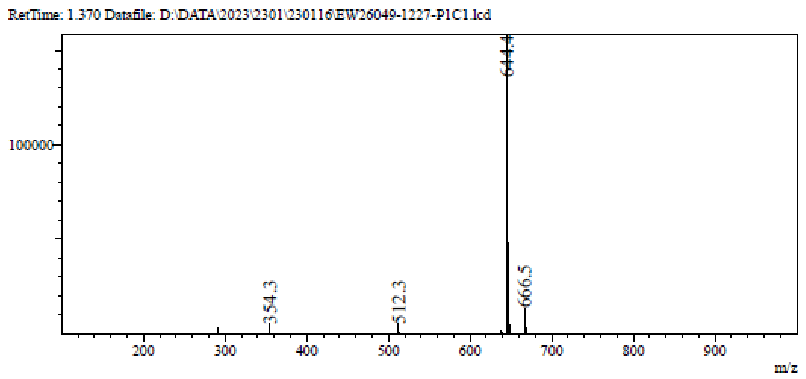

**$^{1}$H NMR** 7-[(2S,3S,4R,5S)-3,4-Bis[[tert-butyl(dimethyl)silyl]oxy]-5-[(6-methyl -1,3-benzothiazol-2-yl)sulfanylmethyl]tetrahydrofuran-2-yl]pyrrolo[2,1-f][1,2,4]triazin-4-amine **(121b)**

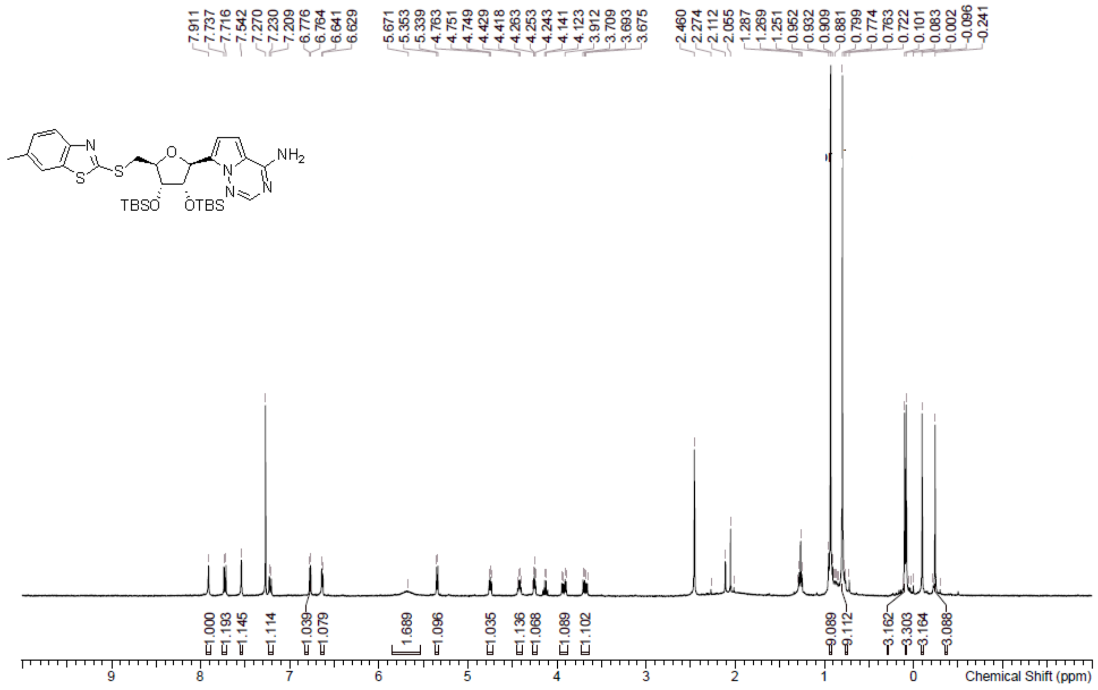

**LCMS** 7-[(2S,3S,4R,5S)-3,4-Bis[[tert-butyl(dimethyl)silyl]oxy]-5-[(6-methyl -1,3-benzothiazol-2-yl)sulfanylmethyl]tetrahydrofuran-2-yl]pyrrolo[2,1-f][1,2,4]triazin-4-amine **(121b)**

Chromatogram

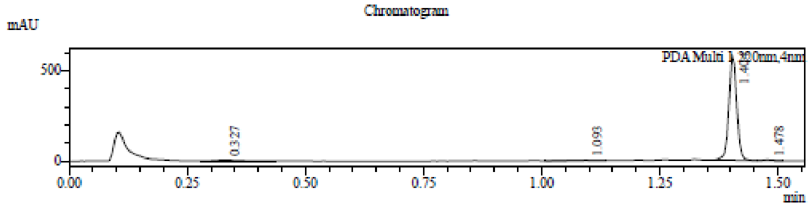

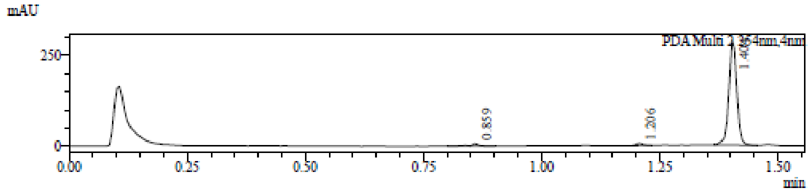

1 PDA Multi 1 / 220nm,4nm
2 PDA Multi 2 / 254nm,4nm

MS Chromatogram

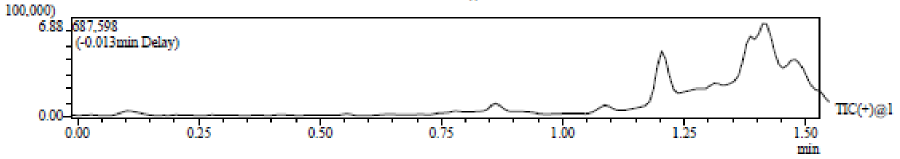

Integration Result

Peak Table

PDA Ch1 220nm

| Peak# | Ret. Time | Height | Height% | USP Width | Area | Area% |
|---|---|---|---|---|---|---|
| 1 | 0.327 | 4505 | 0.758 | 0.108 | 16986 | 2.457 |
| 2 | 1.093 | 2712 | 0.456 | 0.034 | 5561 | 0.804 |
| 3 | 1.404 | 582526 | 98.039 | 0.034 | 662893 | 95.887 |
| 4 | 1.478 | 4435 | 0.746 | 0.043 | 5887 | 0.852 |

Peak Table

PDA Ch2 254nm

| Peak# | Ret. Time | Height | Height% | USP Width | Area | Area% |
|---|---|---|---|---|---|---|
| 1 | 0.859 | 7230 | 2.385 | 0.061 | 8039 | 2.317 |
| 2 | 1.206 | 5876 | 1.938 | 0.028 | 5276 | 1.521 |
| 3 | 1.404 | 290044 | 95.677 | 0.034 | 333646 | 96.162 |

Mass Spectrum

RetTime: 0.328 Datafile: D:\DATA\2023\2301\230116\EW26049-1225-P1C1.lcd

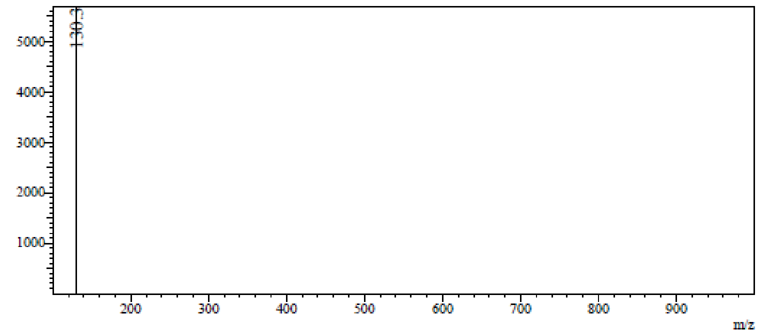

RetTime: 1.095 Datafile: D:\DATA\2023\2301\230116\EW26049-1225-P1C1.lcd

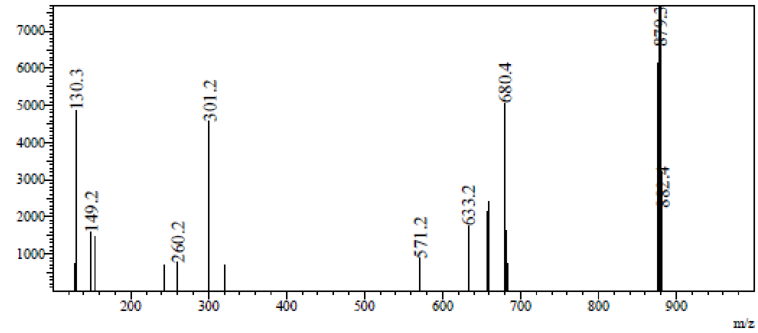

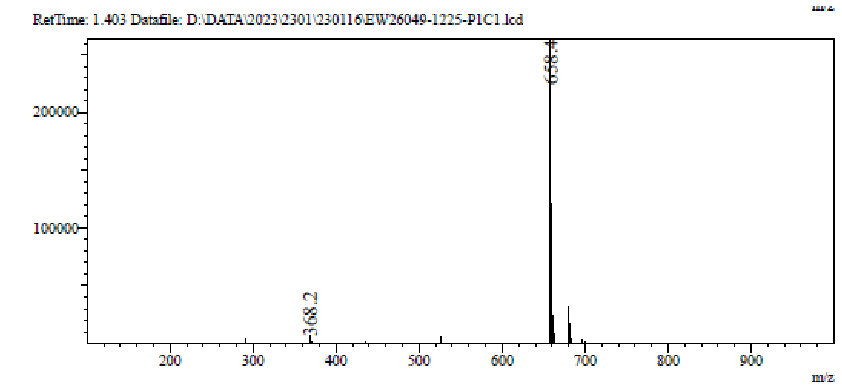

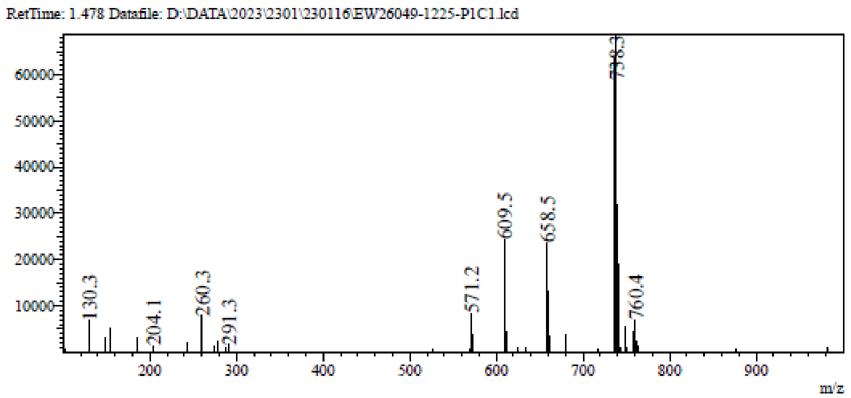

**[1]H NMR** 7-[(2*S*,3*S*,4*R*,5*S*)-3,4-Bis[[tert-butyl(dimethyl)silyl]oxy]-5- (2-naphthylsulfanylmethyl)tetrahydrofuran-2-yl]pyrrolo[2,1-f][1,2,4]triazin-4-amine **(121c)**

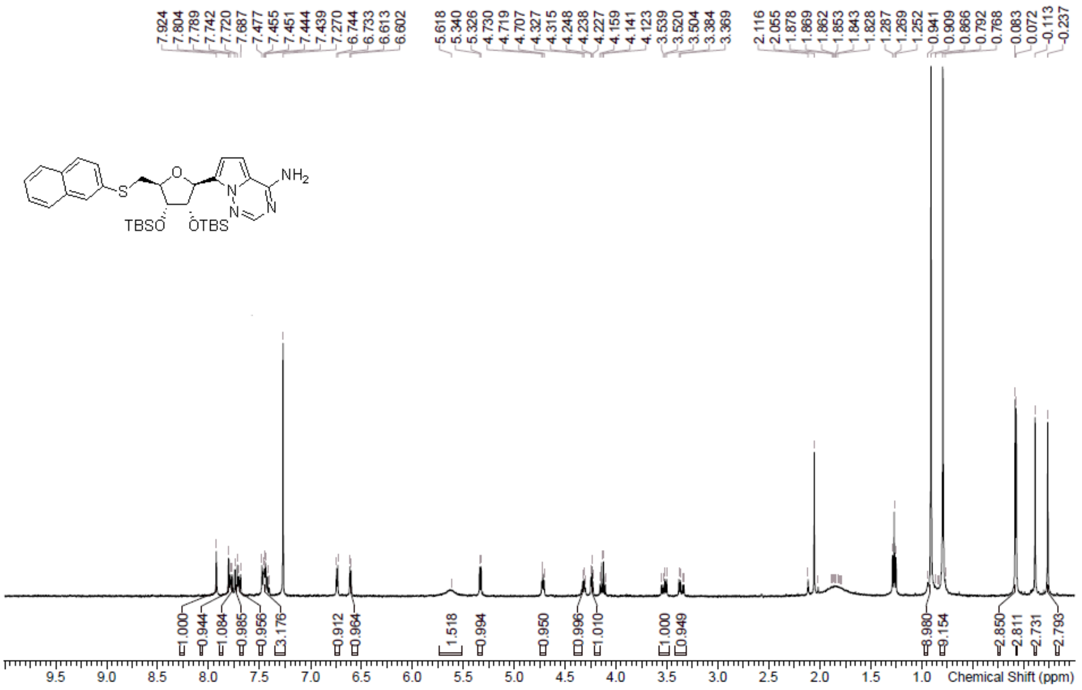

**LCMS** 7-[(2*S*,3*S*,4*R*,5*S*)-3,4-Bis[[tert-butyl(dimethyl)silyl]oxy]-5- (2-naphthylsulfanylmethyl)tetrahydrofuran-2-yl]pyrrolo[2,1-f][1,2,4]triazin-4-amine **(121c)**

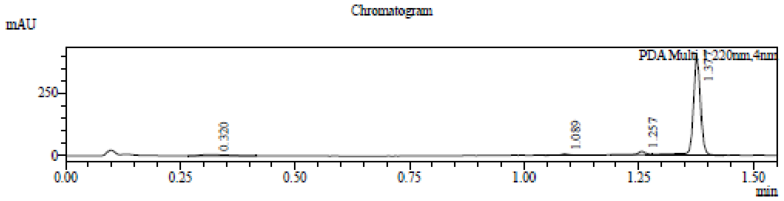

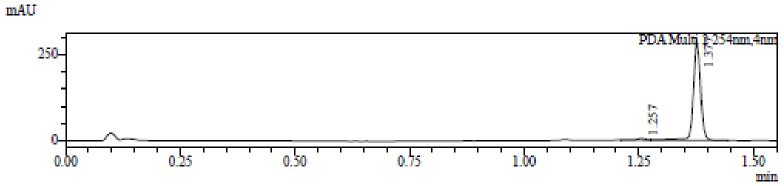

1 PDA Multi 1 / 220nm,4nm
2 PDA Multi 2 / 254nm,4nm

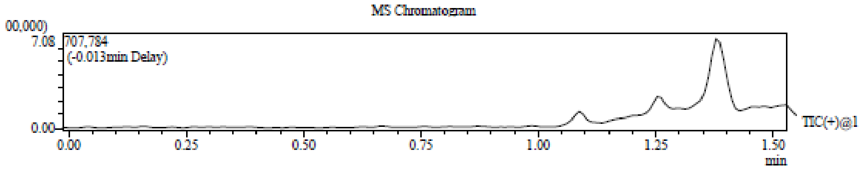

Integration Result

Peak Table

PDA Ch1 220nm

| Peak# | Ret. Time | Height | Height% | USP Width | Area | Area% |
|---|---|---|---|---|---|---|
| 1 | 0.320 | 4646 | 1.074 | 0.109 | 17982 | 3.727 |
| 2 | 1.089 | 5361 | 1.239 | 0.031 | 6016 | 1.247 |
| 3 | 1.257 | 14684 | 3.394 | 0.035 | 20943 | 4.341 |
| 4 | 1.377 | 407918 | 94.293 | 0.156 | 437546 | 90.686 |

Peak Table

PDA Ch2 254nm

| Peak# | Ret. Time | Height | Height% | USP Width | Area | Area% |
|---|---|---|---|---|---|---|
| 1 | 1.257 | 4944 | 1.661 | 0.038 | 6989 | 2.171 |
| 2 | 1.377 | 292657 | 98.339 | 0.139 | 314987 | 97.829 |

Mass Spectrum

RetTime: 0.320 Datafile: D:\DATA\2023\2301\230116\EW26049-1224-P1C1.lcd

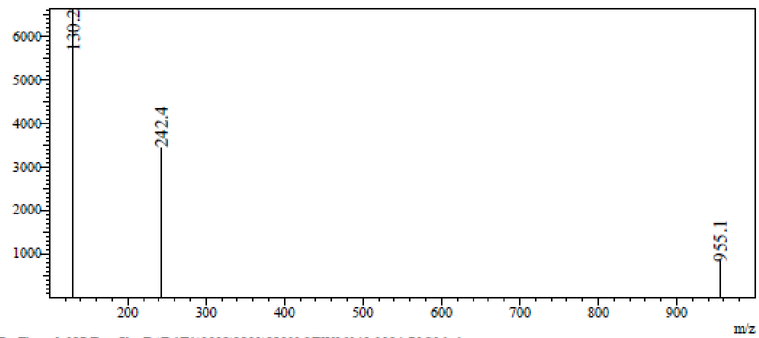

RetTime: 1.087 Datafile: D:\DATA\2023\2301\230116\EW26049-1224-P1C1.lcd

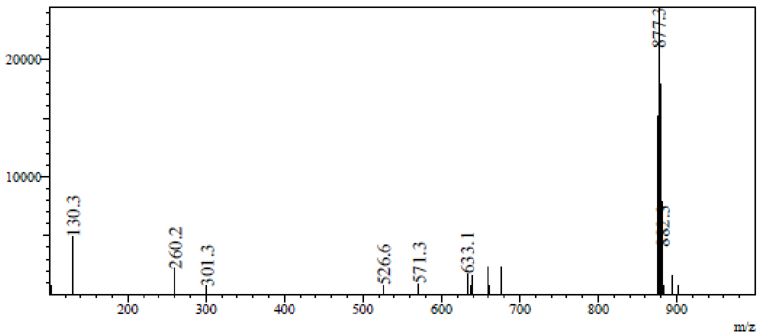

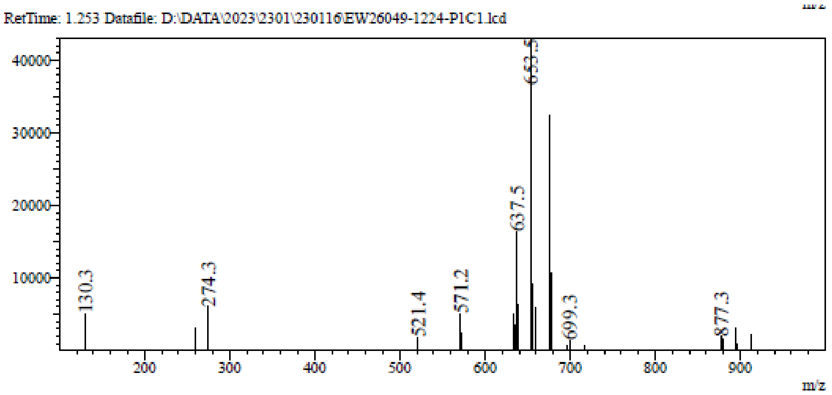

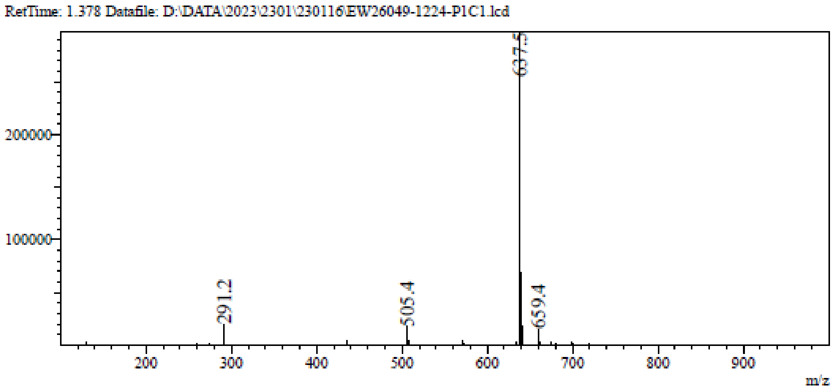

**¹H NMR** 7-[(2*S*,3*S*,4*R*,5*S*)-3,4-Bis[[tert-butyl(dimethyl)silyl]oxy]- 5-(2-quinolylsulfanylmethyl) tetrahydrofuran-2-yl]pyrrolo[2,1-f][1,2,4]triazin-4- amine **(121d)**

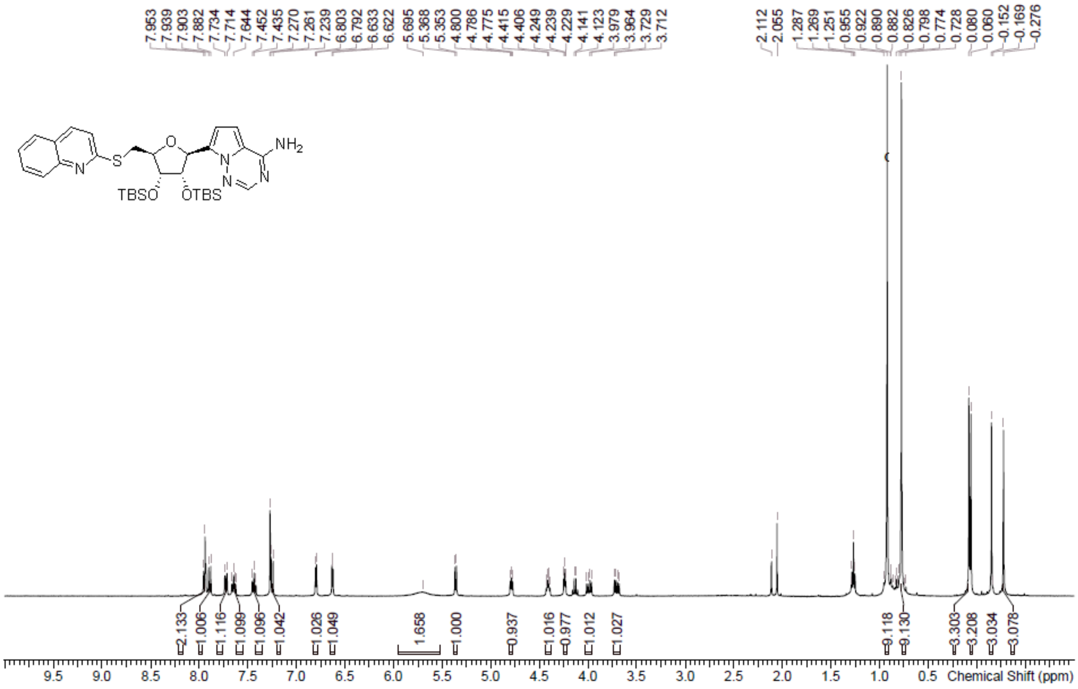

**LCMS** 7-[(2*S*,3*S*,4*R*,5*S*)-3,4-Bis[[tert-butyl(dimethyl)silyl]oxy]- 5-(2-quinolylsulfanylmethyl) tetrahydrofuran-2-yl]pyrrolo[2,1-f][1,2,4]triazin-4- amine **(121d)**

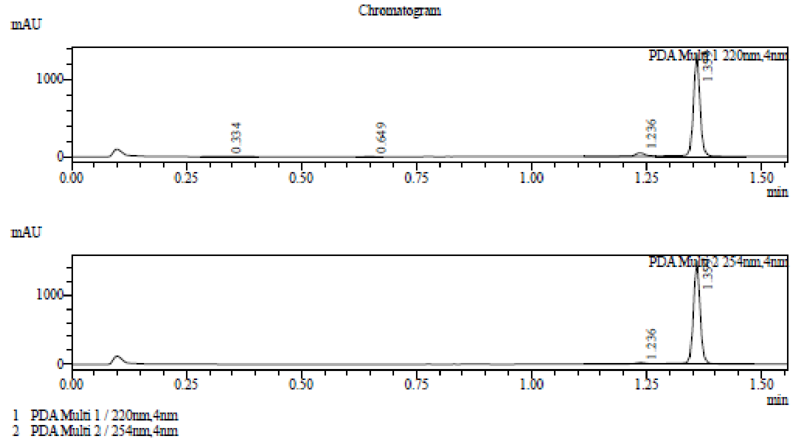

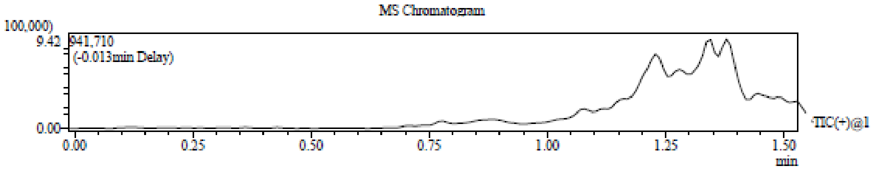

Integration Result

Peak Table

PDA Ch1 220nm

| Peak# | Ret. Time | Height | Height% | USP Width | Area | Area% |
|---|---|---|---|---|---|---|
| 1 | 0.334 | 4335 | 0.312 | 0.104 | 16164 | 1.084 |
| 2 | 0.649 | 4334 | 0.312 | 0.038 | 5566 | 0.373 |
| 3 | 1.236 | 47013 | 3.387 | 0.045 | 84665 | 5.676 |
| 4 | 1.359 | 1332160 | 95.988 | 0.119 | 1385270 | 92.867 |

Peak Table

PDA Ch2 254nm

| Peak# | Ret. Time | Height | Height% | USP Width | Area | Area% |
|---|---|---|---|---|---|---|
| 1 | 1.236 | 23167 | 1.535 | 0.056 | 50256 | 3.046 |
| 2 | 1.359 | 1486112 | 98.465 | 0.032 | 1599631 | 96.954 |

Mass Spectrum

RetTime: 0.337 Datafile: D:\DATA\2023\2301\230116\EW26049-1226-P1C1.lcd

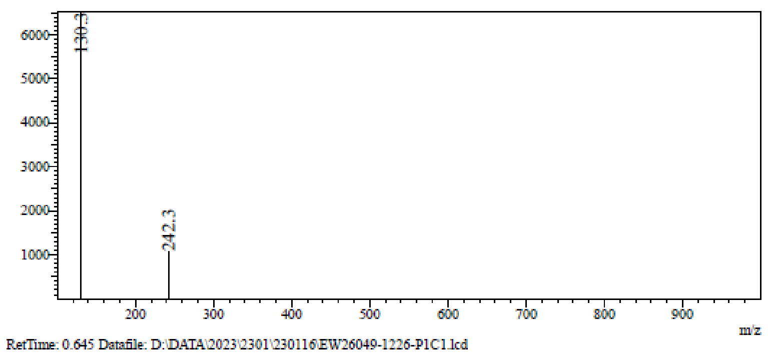

RetTime: 0.645 Datafile: D:\DATA\2023\2301\230116\EW26049-1226-P1C1.lcd

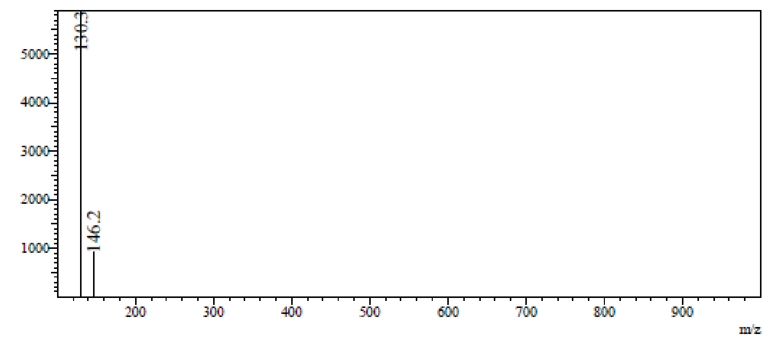

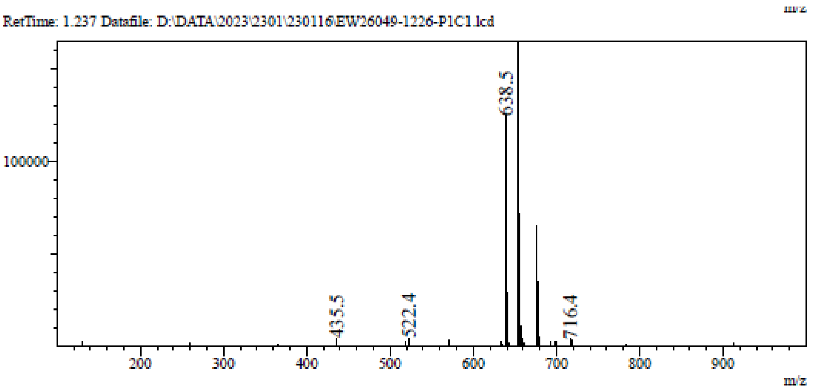

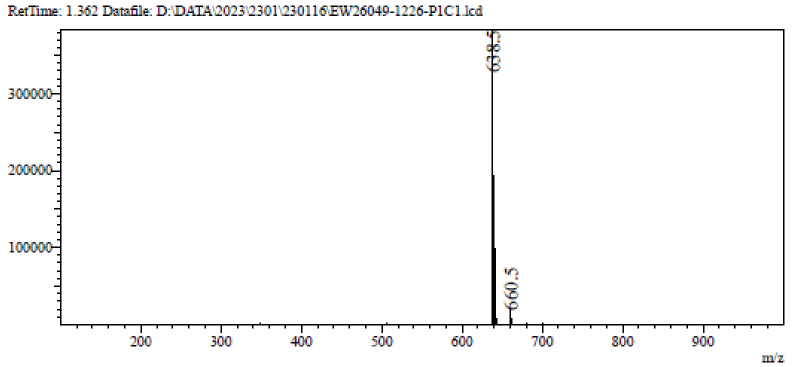

**¹H NMR** (2*S*,3*R*,4*S*,5*S*)-2-(4-Aminopyrrolo[2,1-f][1,2,4]triazin-7-yl)-5- (1,3-benzothiazol-2-ylsulfanylmethyl)tetrahydrofuran-3,4-diol **(35)**

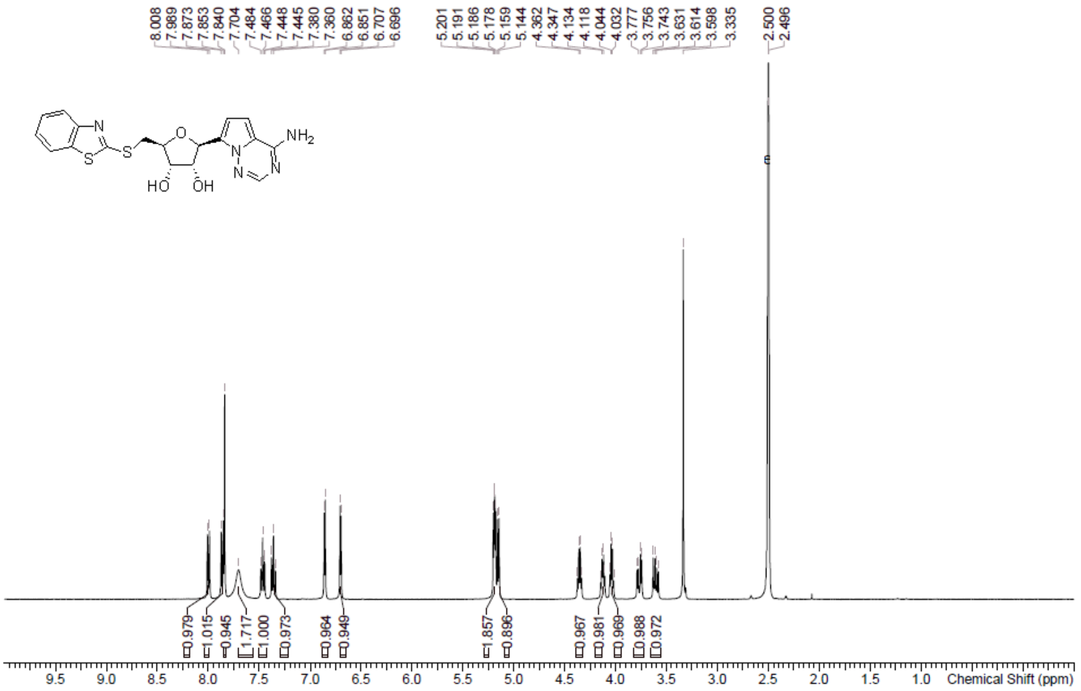

**HPLC** (2*S*,3*R*,4*S*,5*S*)-2-(4-Aminopyrrolo[2,1-f][1,2,4]triazin-7-yl)-5- (1,3-benzothiazol-2-ylsulfanylmethyl)tetrahydrofuran-3,4-diol **(35)**

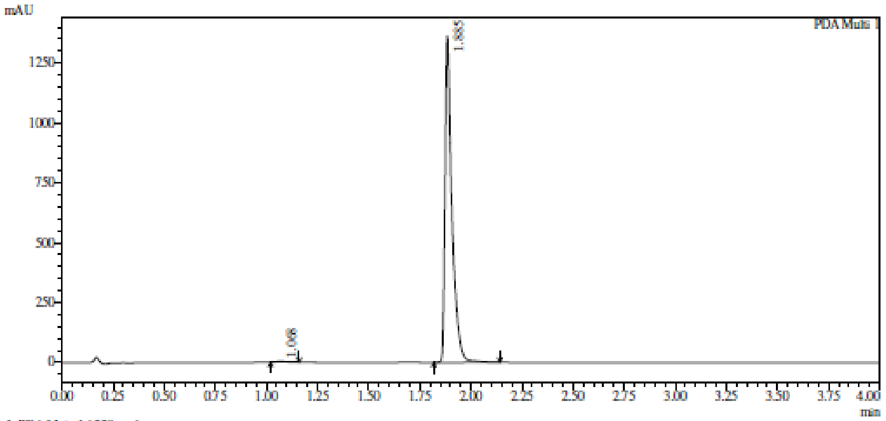

1 PDA Multi 1 / 220nm,4nm

Integration result

PeakTable

PDA Ch1 220nm

| Peak# | Ret. Time | USP Width | Resolution | Height | Area | Area % |
|---|---|---|---|---|---|---|
| 1 | 1.068 | 0.075 | 0.000 | 5252 | 15723 | 0.491 |
| 2 | 1.885 | 0.056 | 12.449 | 1362095 | 3188169 | 99.509 |
| Total | | | | 1367347 | 3203892 | 100.000 |

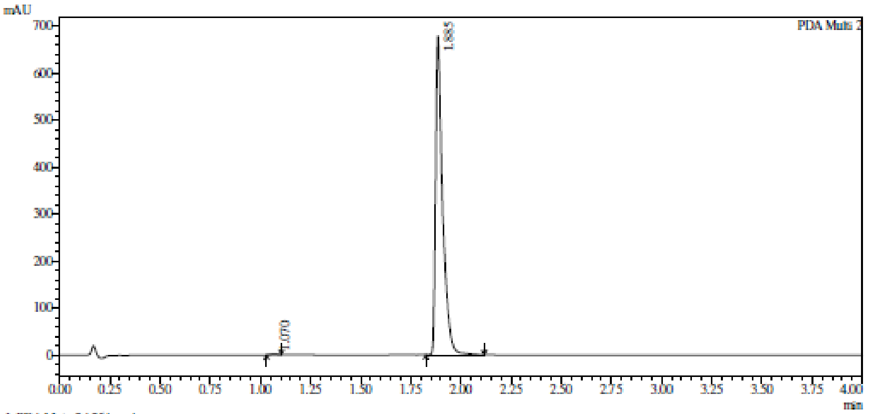

1 PDA Multi 2 / 254nm,4nm

Integration result

PeakTable

PDA Ch2 254nm

| Peak# | Ret. Time | USP Width | Resolution | Height | Area | Area % |
|---|---|---|---|---|---|---|
| 1 | 1.070 | 0.059 | 0.000 | 996 | 2263 | 0.135 |
| 2 | 1.885 | 0.060 | 13.713 | 679112 | 1669071 | 99.865 |
| Total | | | | 680109 | 1671335 | 100.000 |

**HRMS** (2*S*,3*R*,4*S*,5*S*)-2-(4-Aminopyrrolo[2,1-f][1,2,4]triazin-7-yl)-5- (1,3-benzothiazol-2-ylsulfanylmethyl)tetrahydrofuran-3,4-diol **(35)**

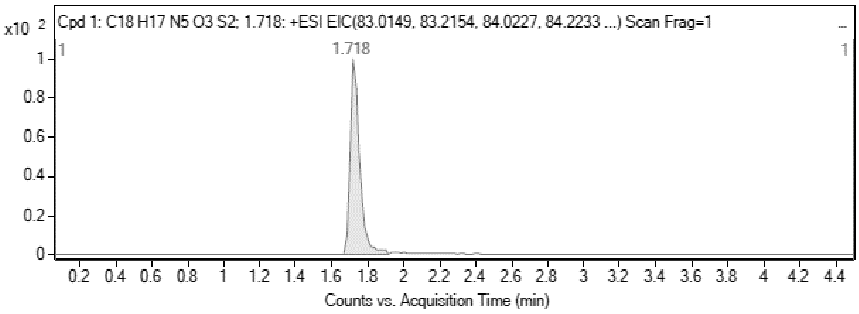

MS Zoomed Spectrum

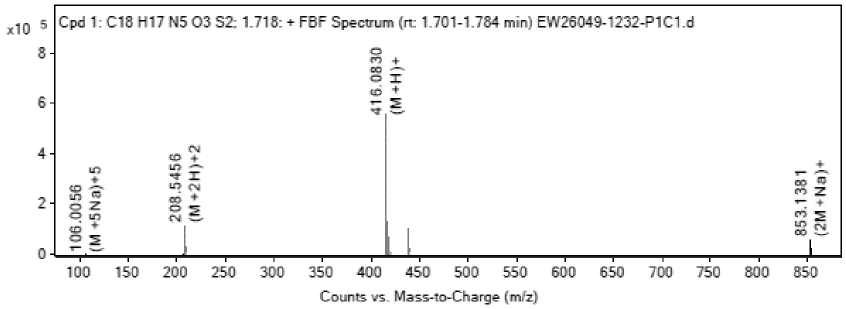

**MS Spectrum Peak List**

| Obs. *m/z* | Charge | Abund | Ion/Isotope |
|---|---|---|---|
| 208.5456 | 2 | 116103.53 | (M+2H)+2 |
| 209.0467 | 2 | 26254.92 | (M+2H)+2 |
| 209.5448 | 2 | 14248.66 | (M+2H)+2 |
| 416.083 | 1 | 559536.25 | (M+H)+ |
| 417.0849 | 1 | 128231.88 | (M+H)+ |
| 418.0803 | 1 | 63988.23 | (M+H)+ |
| 438.0639 | 1 | 101660.52 | (M+Na)+ |
| 439.0662 | 1 | 24114.88 | (M+Na)+ |
| 853.1381 | 1 | 57645.03 | (2M+Na)+ |
| 854.1398 | 1 | 25484.41 | (2M+Na)+ |

MS Zoomed Spectrum

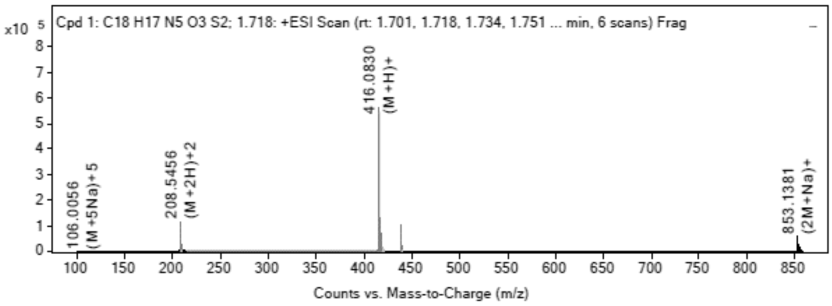

**MS Spectrum Peak List**

| Obs. *m/z* | Charge | Abund | Ion/Isotope | Tgt Mass Error (ppm) |
|---|---|---|---|---|
| 208.5456 | 2 | 116103.53 | (M+2H)+2 | 1.32 |
| 209.0467 | 2 | 26254.92 | (M+2H)+2 | 2.3 |
| 209.5448 | 2 | 14248.66 | (M+2H)+2 | 1.32 |
| 416.083 | 1 | 559536.25 | (M+H)+ | 3.81 |
| 417.0849 | 1 | 128231.88 | (M+H)+ | 5.46 |
| 418.0803 | 1 | 63988.23 | (M+H)+ | 5.87 |
| 438.0639 | 1 | 101660.52 | (M+Na)+ | 5.85 |
| 439.0662 | 1 | 24114.88 | (M+Na)+ | 6.67 |
| 853.1381 | 1 | 57645.03 | (2M+Na)+ | 6.61 |
| 854.1398 | 1 | 25484.41 | (2M+Na)+ | 7.73 |

--- End Of Report ---

**$^1$H NMR** (2*S*,3*R*,4*S*,5*S*)-2-(4-Aminopyrrolo[2,1-f][1,2,4]triazin-7-yl)-5- [(6-methyl-1,3-benzothiazol-2-yl)sulfanylmethyl]tetrahydrofuran-3,4-diol **(36)**

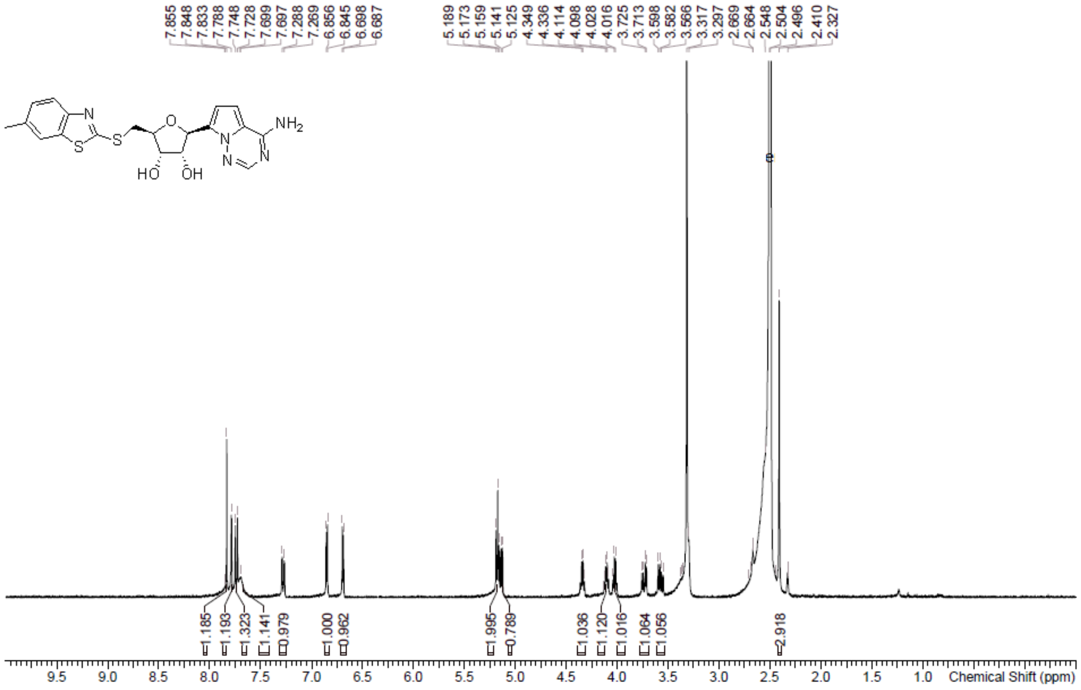

**HPLC** (2*S*,3*R*,4*S*,5*S*)-2-(4-Aminopyrrolo[2,1-f][1,2,4]triazin-7-yl)-5- [(6-methyl-1,3-benzothiazol-2-yl)sulfanylmethyl]tetrahydrofuran-3,4-diol **(36)**

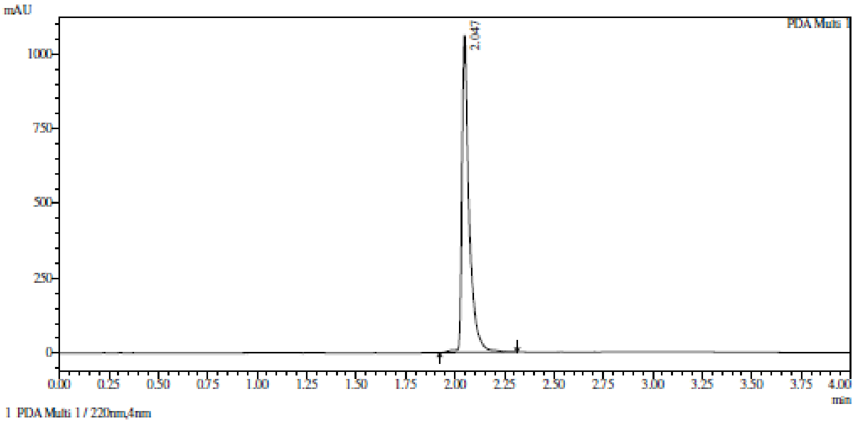

Integration result

PeakTable

PDA Ch1 220nm

| Peak# | Ret. Time | USP Width | Resolution | Height | Area | Area % |
|---|---|---|---|---|---|---|
| 1 | 2.047 | 0.057 | 0.000 | 1060182 | 2646473 | 100.000 |
| Total | | | | 1060182 | 2646473 | 100.000 |

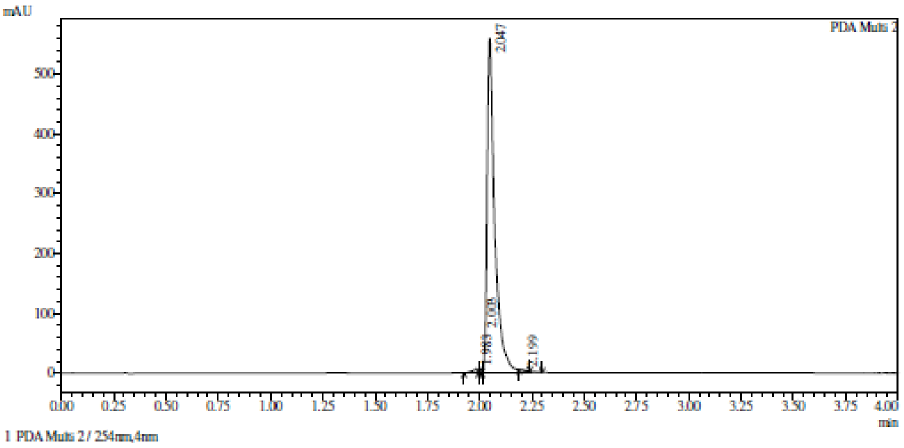

Integration result

PeakTable

PDA Ch2 254nm

| Peak# | Ret. Time | USP Width | Resolution | Height | Area | Area % |
|---|---|---|---|---|---|---|
| 1 | 1.983 | 0.449 | 0.000 | 5882 | 12794 | 0.874 |
| 2 | 2.003 | 0.000 | 0.000 | 6154 | 5875 | 0.401 |
| 3 | 2.047 | 0.060 | 0.000 | 559244 | 1444709 | 98.655 |
| 4 | 2.199 | 0.043 | 2.956 | 644 | 1021 | 0.070 |
| Total | | | | 571924 | 1464399 | 100.000 |

**HRMS** (2*S*,3*R*,4*S*,5*S*)-2-(4-Aminopyrrolo[2,1-f][1,2,4]triazin-7-yl)-5- [(6-methyl-1,3-benzothiazol-2-yl)sulfanylmethyl]tetrahydrofuran-3,4-diol **(36)**

**Compound Table**

| Label | Tgt Score | Mass Error (ppm) | Tgt Formula | Obs. RT | Ref. Mass | Obs. Mass |
|---|---|---|---|---|---|---|
| Cpd 1: C19 H19 N5 O3 S2; 1.859 | 97.39 | 0.46 | C19 H19 N5 O3 S2 | 1.859 | 429.0929 | 429.0931 |

| Obs. *m/z* | Obs. RT | Obs. Mass | Tgt Formula | Tgt Mass | Tgt Mass Error (ppm) | Find Cpds Algorithm |
|---|---|---|---|---|---|---|
| 430.1006 | 1.859 | 429.0931 | C19 H19 N5 O3 S2 | 429.0929 | 0.46 | Find by Formula |

**Compound Chromatograms**

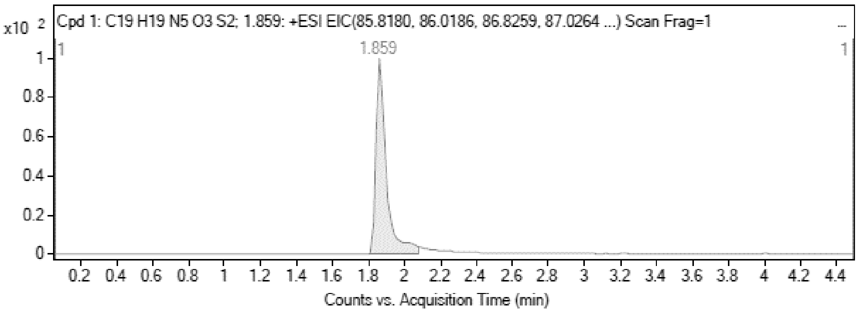

**MS Zoomed Spectrum**

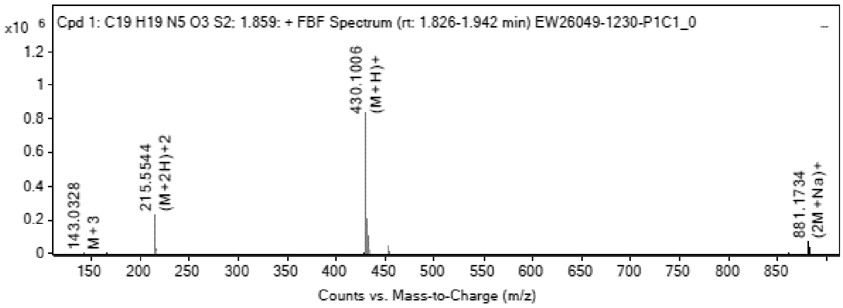

**MS Spectrum Peak List**

| Obs. *m/z* | Charge | Abund | Ion/Isotope |
|---|---|---|---|
| 215.5544 | 2 | 228431.22 | (M+2H)+2 |

| | | | |
|---|---|---|---|
| 216.0555 | 2 | 55924.73 | (M+2H)+2 |
| 216.5535 | 2 | 28726.35 | (M+2H)+2 |
| 430.1006 | 1 | 834947.69 | (M+H)+ |
| 431.1025 | 1 | 204448.72 | (M+H)+ |
| 432.098 | 1 | 99319.41 | (M+H)+ |
| 452.0811 | 1 | 48844.35 | (M+Na)+ |
| 453.0836 | 1 | 12057.81 | (M+Na)+ |
| 881.1734 | 1 | 69624.05 | (2M+Na)+ |
| 882.1762 | 1 | 33133.84 | (2M+Na)+ |

MS Zoomed Spectrum

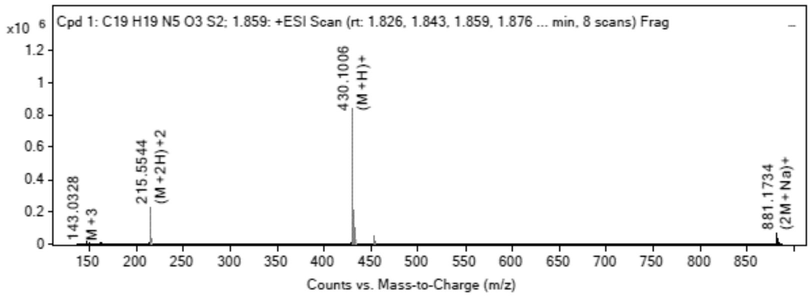

MS Spectrum Peak List

| Obs. m/z | Charge | Abund | Ion/Isotope | Tgt Mass Error (ppm) |
|---|---|---|---|---|
| 215.5544 | 2 | 228431.22 | (M+2H)+2 | -3.06 |
| 216.0555 | 2 | 55924.73 | (M+2H)+2 | -2.03 |
| 216.5535 | 2 | 28726.35 | (M+2H)+2 | -2.75 |
| 430.1006 | 1 | 834947.69 | (M+H)+ | -0.81 |
| 431.1025 | 1 | 204448.72 | (M+H)+ | 0.87 |
| 432.098 | 1 | 99319.41 | (M+H)+ | 1.33 |
| 452.0811 | 1 | 48844.35 | (M+Na)+ | 2.35 |
| 453.0836 | 1 | 12057.81 | (M+Na)+ | 2.73 |
| 881.1734 | 1 | 69624.05 | (2M+Na)+ | 1.91 |
| 882.1762 | 1 | 33133.84 | (2M+Na)+ | 1.69 |

--- End Of Report ---

**$^1$H NMR** (2*S*,3*R*,4*S*,5*S*)-2-(4-Aminopyrrolo[2,1-f][1,2,4]triazin-7-yl)- 5-(2-naphthylsulfanylmethyl) tetrahydrofuran-3,4-diol **(37)**

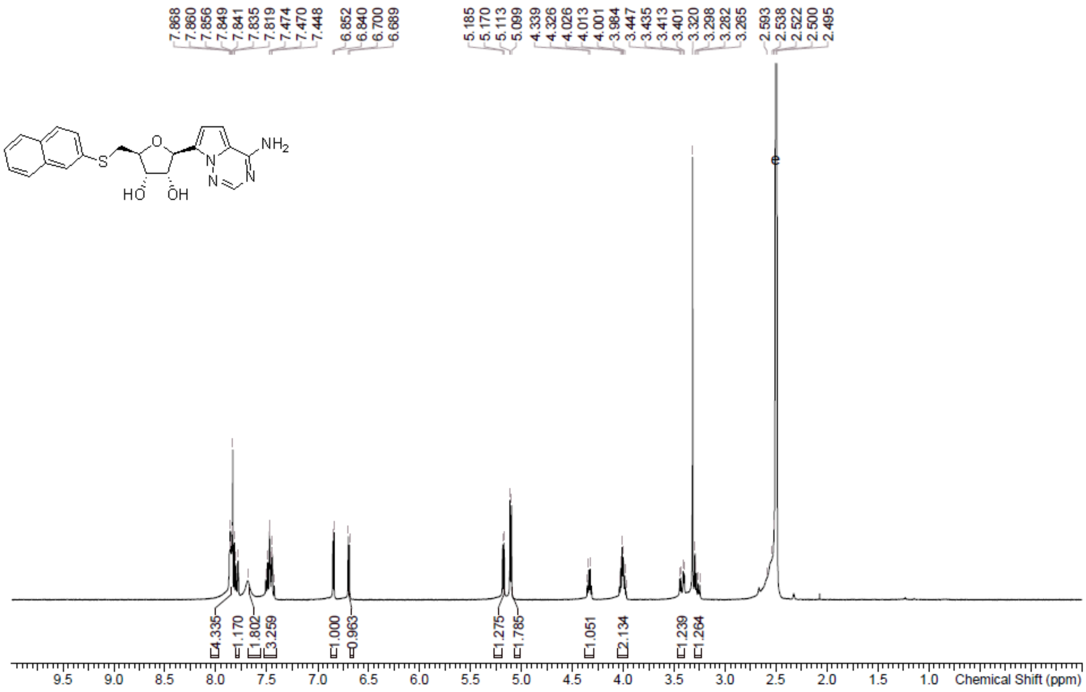

**HPLC** (2*S*,3*R*,4*S*,5*S*)-2-(4-Aminopyrrolo[2,1-f][1,2,4]triazin-7-yl)- 5-(2-naphthylsulfanylmethyl)tetrahydrofuran-3,4-diol **(37)**

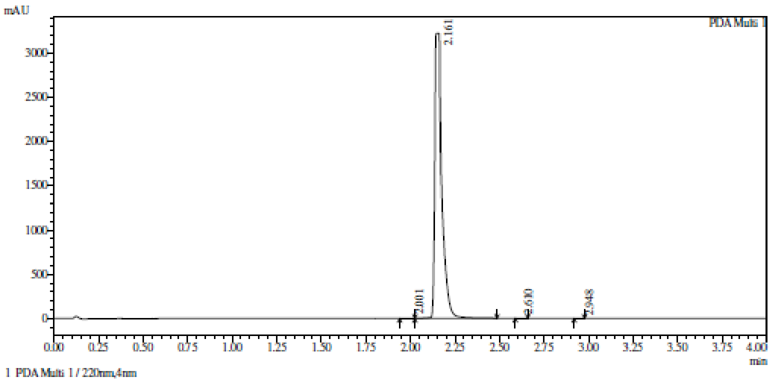

Integration result

PeakTable

PDA Ch1 220nm

| Peak# | Ret. Time | USP Width | Resolution | Height | Area | Area % |
|---|---|---|---|---|---|---|
| 1 | 2.001 | 0.103 | 0.000 | 1210 | 3918 | 0.045 |
| 2 | 2.161 | 0.050 | 2.106 | 3225123 | 8687268 | 99.921 |
| 3 | 2.610 | 0.052 | 8.854 | 889 | 1692 | 0.019 |
| 4 | 2.948 | 0.050 | 6.654 | 702 | 1232 | 0.014 |
| Total | | | | 3227925 | 8694109 | 100.000 |

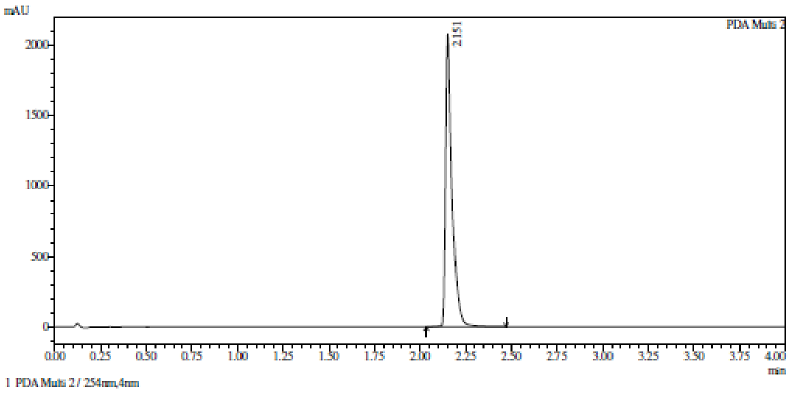

Integration result

PeakTable

PDA Ch2 254nm

| Peak# | Ret. Time | USP Width | Resolution | Height | Area | Area % |
|---|---|---|---|---|---|---|
| 1 | 2.151 | 0.060 | 0.000 | 2076332 | 5202085 | 100.000 |
| Total | | | | 2076332 | 5202085 | 100.000 |

**HRMS** (2*S*,3*R*,4*S*,5*S*)-2-(4-Aminopyrrolo[2,1-f][1,2,4]triazin-7-yl)- 5-(2-naphthylsulfanylmethyl)tetrahydrofuran-3,4-diol **(37)**

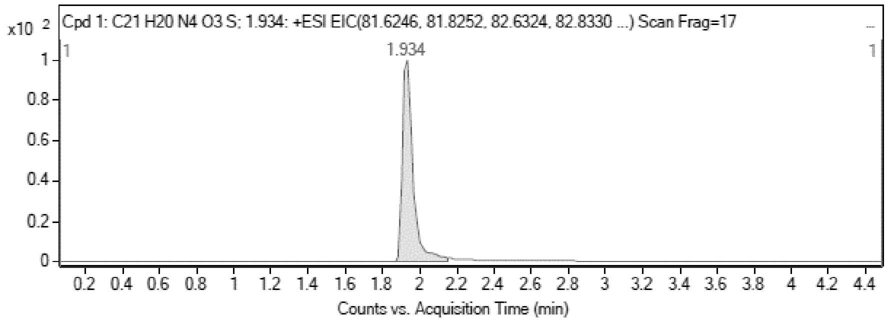

MS Zoomed Spectrum

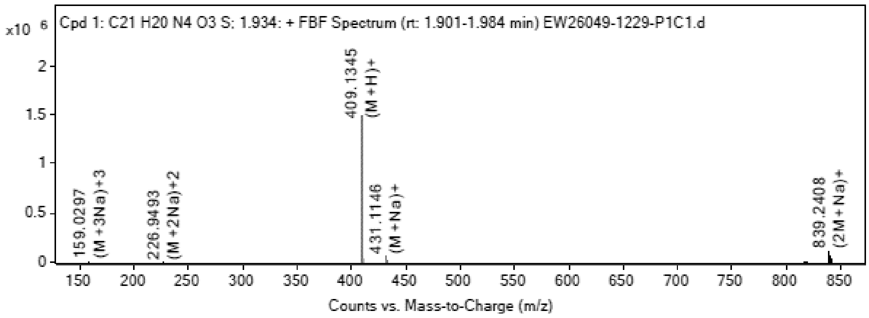

MS Spectrum Peak List

| Obs. *m/z* | Charge | Abund | Ion/Isotope |
|---|---|---|---|
| 409.1345 | 1 | 1475529.13 | (M+H)+ |
| 410.1366 | 1 | 389048.5 | (M+H)+ |
| 411.1338 | 1 | 117840.38 | (M+H)+ |
| 431.1146 | 1 | 57345.16 | (M+Na)+ |
| 432.1174 | 1 | 14351.31 | (M+Na)+ |
| 816.2443 | 1 | 3766.16 | 2M+ |
| 839.2408 | 1 | 110424.12 | (2M+Na)+ |
| 840.2431 | 1 | 56908.58 | (2M+Na)+ |
| 841.2425 | 1 | 24947.69 | (2M+Na)+ |
| 842.2418 | 1 | 8018.12 | (2M+Na)+ |

MS Zoomed Spectrum

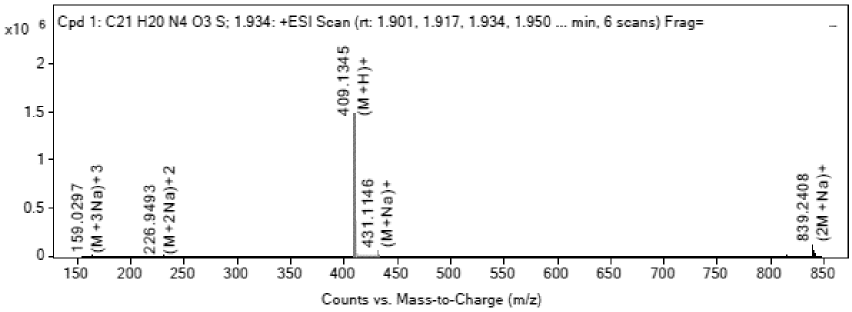

MS Spectrum Peak List

| Obs. *m/z* | Charge | Abund | Ion/Isotope | Tgt Mass Error (ppm) |
|---|---|---|---|---|
| 409.1345 | 1 | 1475529.13 | (M+H)+ | -3.88 |
| 410.1366 | 1 | 389048.5 | (M+H)+ | -1.96 |
| 411.1338 | 1 | 117840.38 | (M+H)+ | -1.76 |
| 431.1146 | 1 | 57345.16 | (M+Na)+ | 0.52 |
| 432.1174 | 1 | 14351.31 | (M+Na)+ | 0.76 |
| 816.2443 | 1 | 3766.16 | 2M+ | 7.8 |
| 839.2408 | 1 | 110424.12 | (2M+Na)+ | -0.38 |
| 840.2431 | 1 | 56908.58 | (2M+Na)+ | 0.26 |
| 841.2425 | 1 | 24947.69 | (2M+Na)+ | -0.36 |
| 842.2418 | 1 | 8018.12 | (2M+Na)+ | 1.17 |

--- End Of Report ---

**$^1$H NMR** (2*S*,3*R*,4*S*,5*S*)-2-(4-Aminopyrrolo[2,1-f][1,2,4]triazin-7-yl)-5- (2-quinolylsulfanylmethyl)tetrahydrofuran-3,4-diol **(38)**

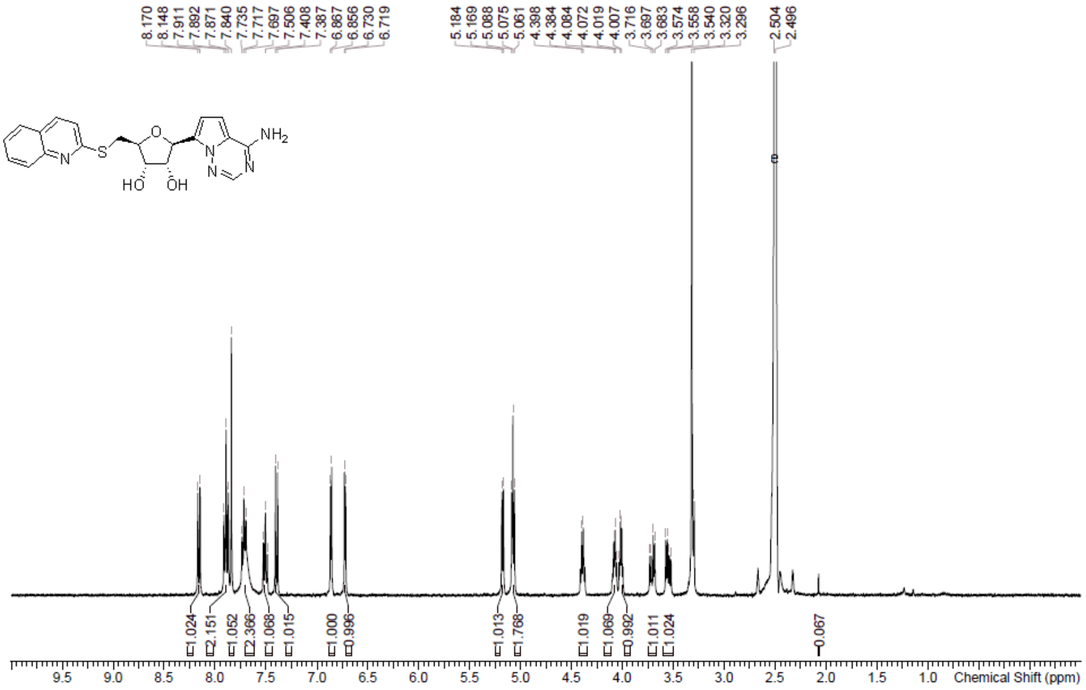

**HPLC** (2*S*,3*R*,4*S*,5*S*)-2-(4-Aminopyrrolo[2,1-f][1,2,4]triazin-7-yl)-5- (2-quinolylsulfanylmethyl)tetrahydrofuran-3,4-diol **(38)**

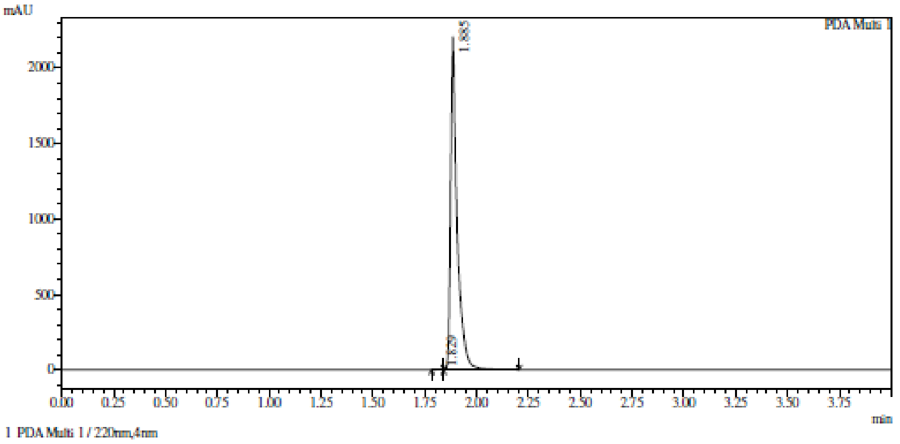

Integration result

PeakTable

PDA Ch1 220nm

| Peak# | Ret. Time | USP Width | Resolution | Height | Area | Area % |
|---|---|---|---|---|---|---|
| 1 | 1.829 | 0.360 | 0.000 | 2722 | 5049 | 0.105 |
| 2 | 1.885 | 0.052 | 0.270 | 2207403 | 4826089 | 99.895 |
| Total | | | | 2210125 | 4831138 | 100.000 |

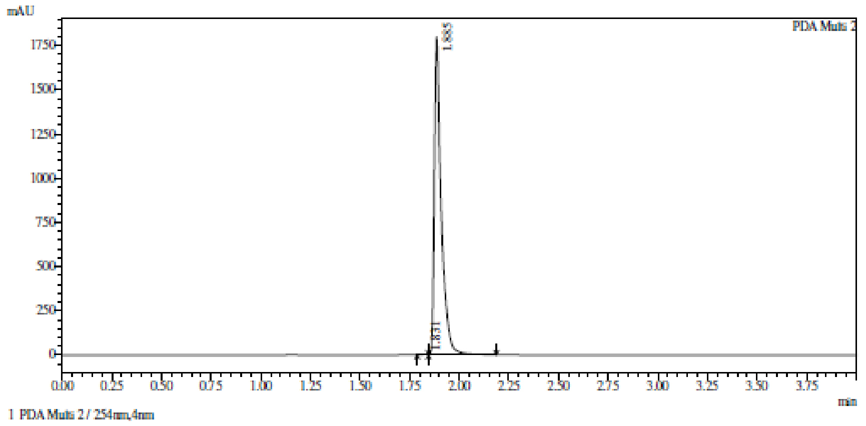

Integration result

PeakTable

PDA Ch2 254nm

| Peak# | Ret. Time | USP Width | Resolution | Height | Area | Area % |
|---|---|---|---|---|---|---|
| 1 | 1.831 | 0.189 | 0.000 | 3022 | 5723 | 0.131 |
| 2 | 1.885 | 0.058 | 0.443 | 1799424 | 4367031 | 99.869 |
| Total | | | | 1802446 | 4372754 | 100.000 |

**HRMS** (2*S*,3*R*,4*S*,5*S*)-2-(4-Aminopyrrolo[2,1-f][1,2,4]triazin-7-yl)-5- (2-quinolylsulfanylmethyl)tetrahydrofuran-3,4-diol **(38)**

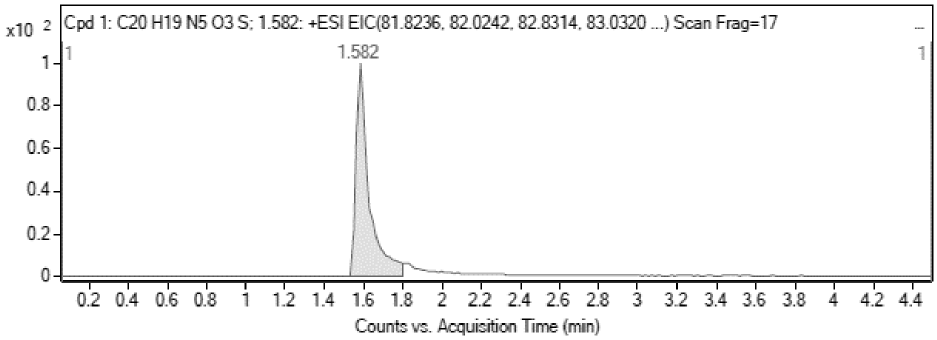

MS Zoomed Spectrum

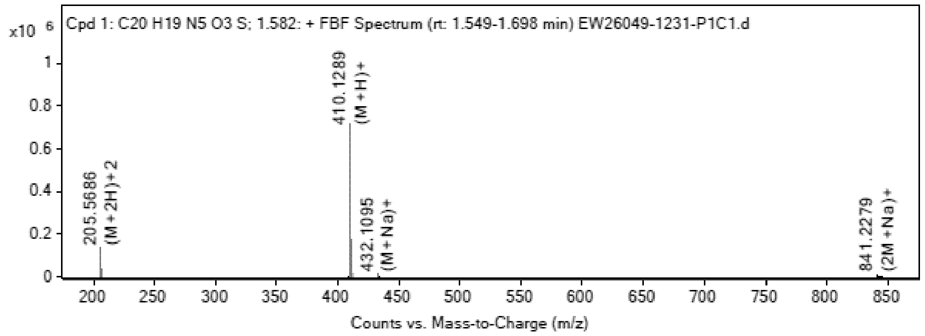

MS Spectrum Peak List

| Obs. *m/z* | Charge | Abund | Ion/Isotope |
|---|---|---|---|
| 205.5686 | 2 | 131317.02 | (M+2H)+2 |
| 206.0698 | 2 | 34125.49 | (M+2H)+2 |

| | | | |
|---|---|---|---|
| 206.5683 | 2 | 10675.34 | (M+2H)+2 |
| 410.1289 | 1 | 716498 | (M+H)+ |
| 411.1309 | 1 | 176433.63 | (M+H)+ |
| 412.1281 | 1 | 54705.14 | (M+H)+ |
| 432.1095 | 1 | 11102.03 | (M+Na)+ |
| 433.1122 | 1 | 2709.34 | (M+Na)+ |
| 841.2279 | 1 | 8108.4 | (2M+Na)+ |
| 842.2323 | 1 | 4070.93 | (2M+Na)+ |

MS Zoomed Spectrum

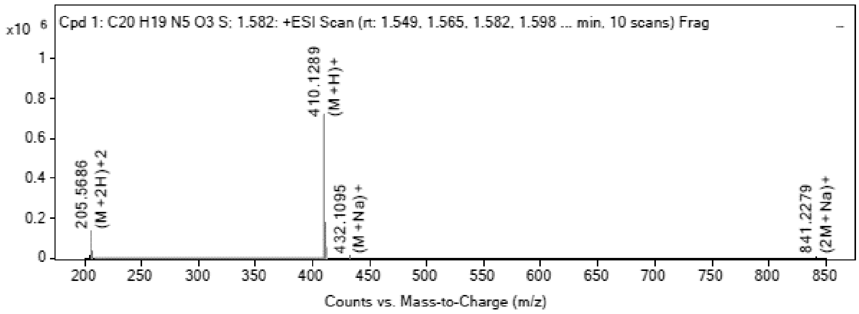

**MS Spectrum Peak List**

| Obs. *m/z* | Charge | Abund | Ion/Isotope | Tgt Mass Error (ppm) |
|---|---|---|---|---|
| 205.5686 | 2 | 131317.02 | (M+2H)+2 | -4.52 |
| 206.0698 | 2 | 34125.49 | (M+2H)+2 | -3.51 |
| 206.5683 | 2 | 10675.34 | (M+2H)+2 | -2.84 |
| 410.1289 | 1 | 716498 | (M+H)+ | -1.86 |
| 411.1309 | 1 | 176433.63 | (M+H)+ | 0.03 |
| 412.1281 | 1 | 54705.14 | (M+H)+ | -0.05 |
| 432.1095 | 1 | 11102.03 | (M+Na)+ | 1.4 |
| 433.1122 | 1 | 2709.34 | (M+Na)+ | 1.65 |
| 841.2279 | 1 | 8108.4 | (2M+Na)+ | 3.56 |
| 842.2323 | 1 | 4070.93 | (2M+Na)+ | 1.65 |

--- End Of Report ---

[1]**H NMR** (2S,3R,4S,5S)-5-(4-Aminopyrrolo[2,1-f][1,2,4]triazin-7-yl)-3,4-bis [[tert-butyl(dimethyl)silyl]oxy]tetrahydrofuran-2-carbaldehyde **(122)**

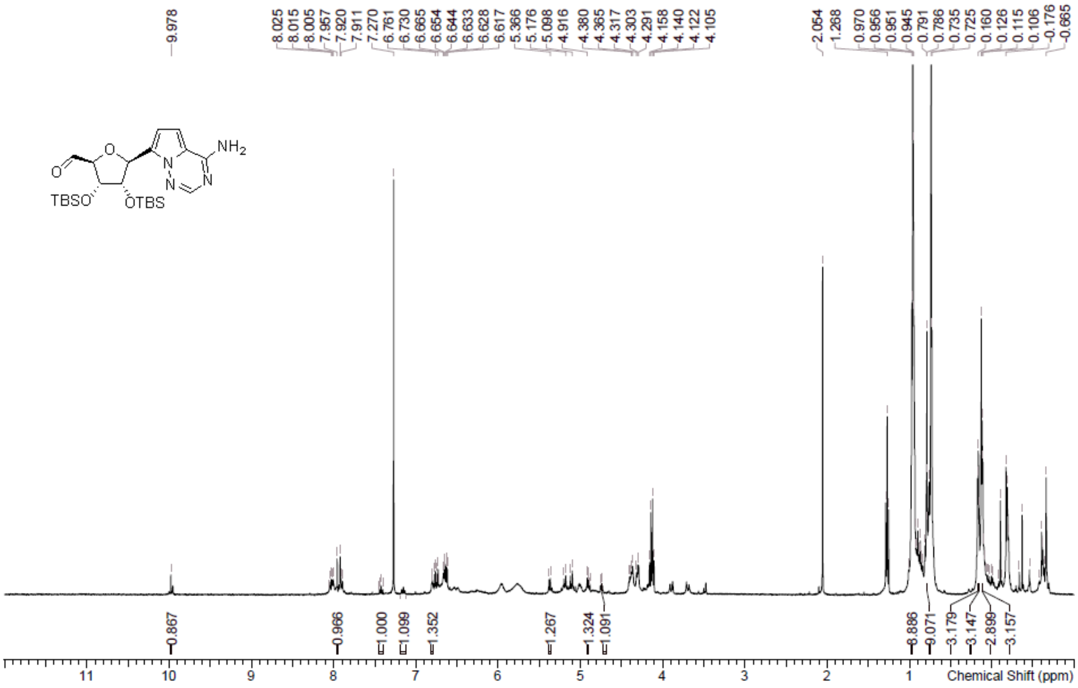

**LCMS** (2S,3R,4S,5S)-5-(4-Aminopyrrolo[2,1-f][1,2,4]triazin-7-yl)-3,4-bis [[tert-butyl(dimethyl)silyl]oxy]tetrahydrofuran-2-carbaldehyde **(122)**

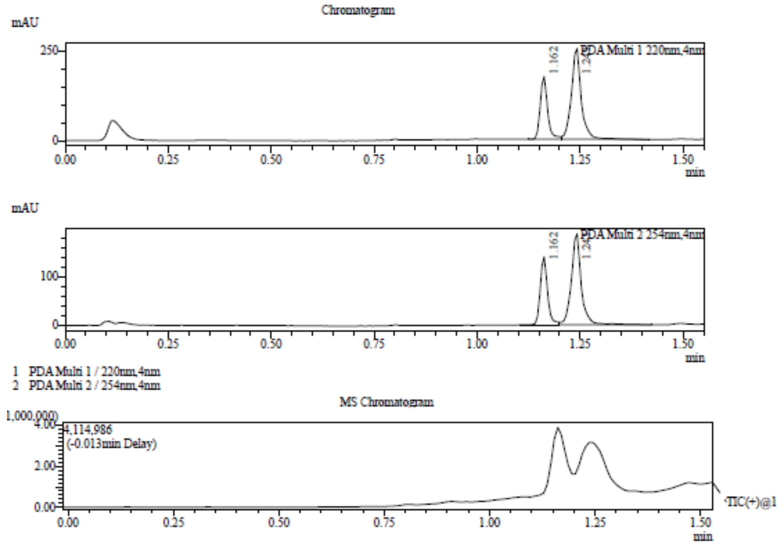

Integration Result

Peak Table

PDA Ch1 220nm

| Peak# | Ret. Time | Height | Height% | USP Width | Area | Area% |
|---|---|---|---|---|---|---|
| 1 | 1.162 | 175202 | 40.862 | 0.036 | 216793 | 33.969 |
| 2 | 1.241 | 253567 | 59.138 | 0.046 | 421417 | 66.031 |

Peak Table

PDA Ch2 254nm

| Peak# | Ret. Time | Height | Height% | USP Width | Area | Area% |
|---|---|---|---|---|---|---|
| 1 | 1.162 | 140119 | 42.848 | 0.036 | 172054 | 35.564 |
| 2 | 1.241 | 186894 | 57.152 | 0.046 | 311728 | 64.436 |

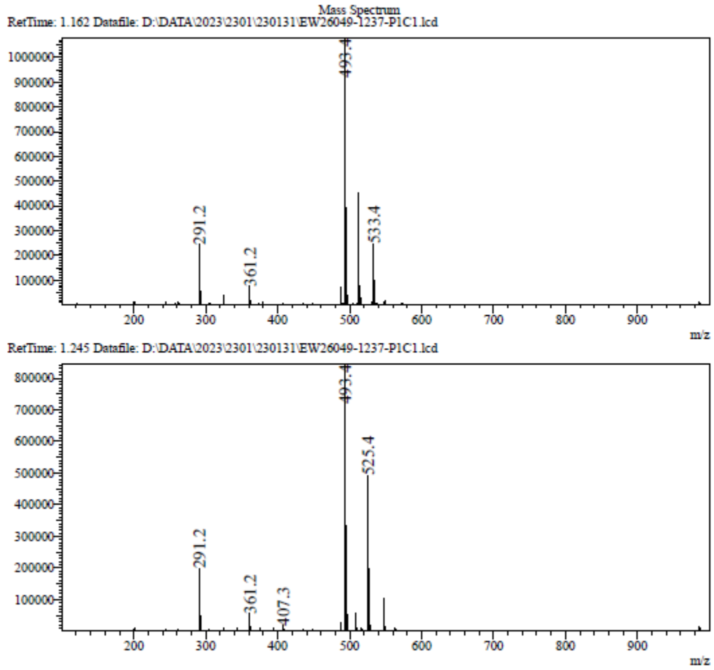

**$^1$H NMR** 7-[(2*S*,3*S*,4*R*,5*R*)-5-[(*E*)-2-(1,3-Benzothiazol-2-yl)vinyl]-3,4-bis [[tert-butyl(dimethyl)silyl]oxy]tetrahydrofuran-2-yl]pyrrolo[2,1-f][1,2,4]triazin-4-amine **(123)**

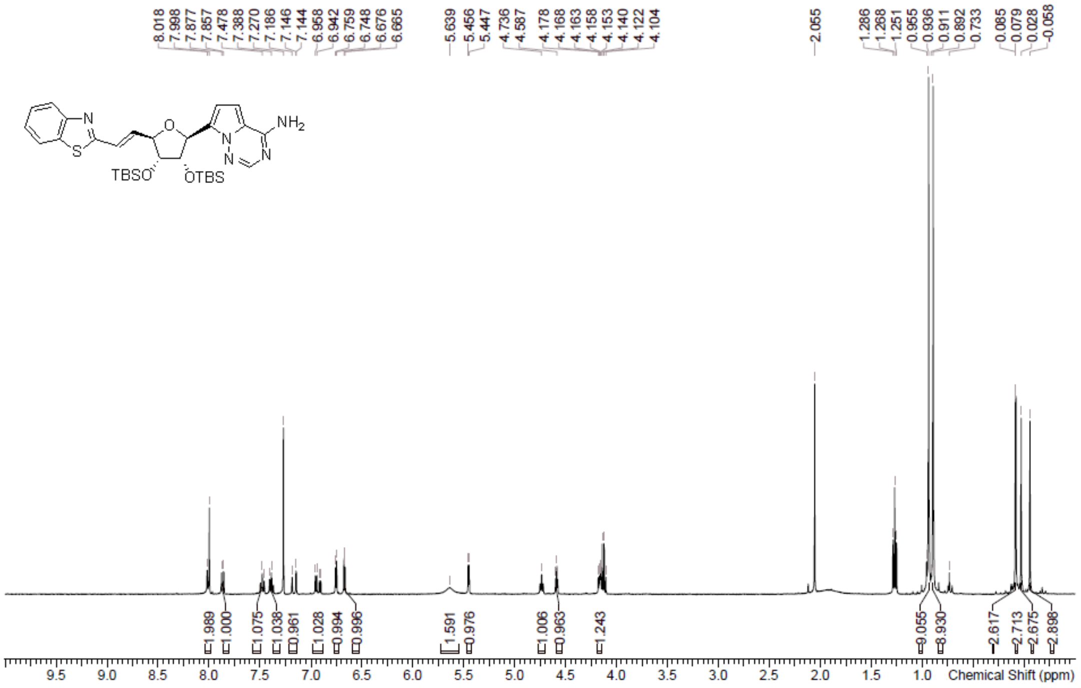

**LCMS** 7-[(2*S*,3*S*,4*R*,5*R*)-5-[(*E*)-2-(1,3-Benzothiazol-2-yl)vinyl]-3,4-bis [[tert-butyl(dimethyl)silyl]oxy]tetrahydrofuran-2-yl]pyrrolo[2,1-f][1,2,4]triazin-4-amine **(123)**

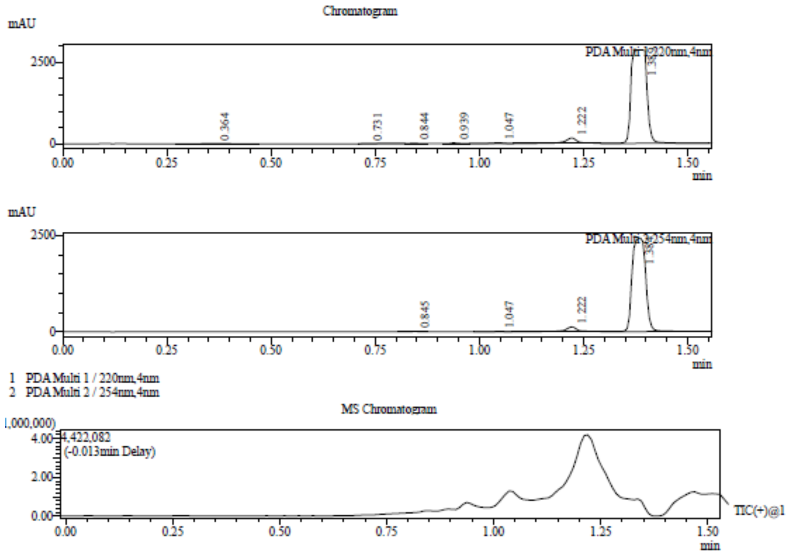

Integration Result

Peak Table

PDA Ch1 220nm

| Peak# | Ret. Time | Height | Height% | USP Width | Area | Area% |
|---|---|---|---|---|---|---|
| 1 | 0.364 | 5345 | 0.173 | 0.123 | 22903 | 0.289 |
| 2 | 0.731 | 3391 | 0.110 | 0.311 | 8368 | 0.106 |
| 3 | 0.844 | 18885 | 0.610 | 0.037 | 22529 | 0.284 |
| 4 | 0.939 | 19380 | 0.626 | 0.035 | 22579 | 0.285 |
| 5 | 1.047 | 23326 | 0.754 | 0.052 | 50926 | 0.643 |
| 6 | 1.222 | 168259 | 5.436 | 0.054 | 370884 | 4.683 |
| 7 | 1.389 | 2856913 | 92.292 | 0.062 | 7421135 | 93.709 |

Peak Table

PDA Ch2 254nm

| Peak# | Ret. Time | Height | Height% | USP Width | Area | Area% |
|---|---|---|---|---|---|---|
| 1 | 0.845 | 7004 | 0.273 | 0.035 | 8121 | 0.142 |

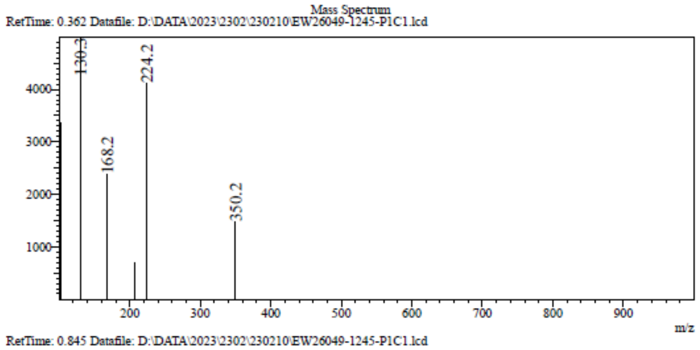
Mass Spectrum
RetTime: 0.362 Datafile: D:\DATA\2023\2302\230210\EW26049-1245-P1C1.lcd
130.3
224.2
168.2
350.2
m/z

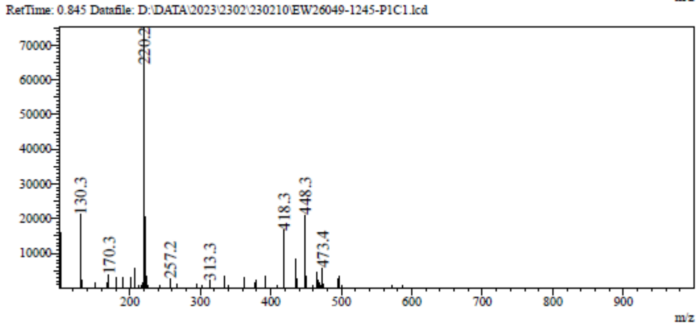
RetTime: 0.845 Datafile: D:\DATA\2023\2302\230210\EW26049-1245-P1C1.lcd
220.2
130.3
170.3
257.2
313.3
418.3
448.3
473.4
m/z

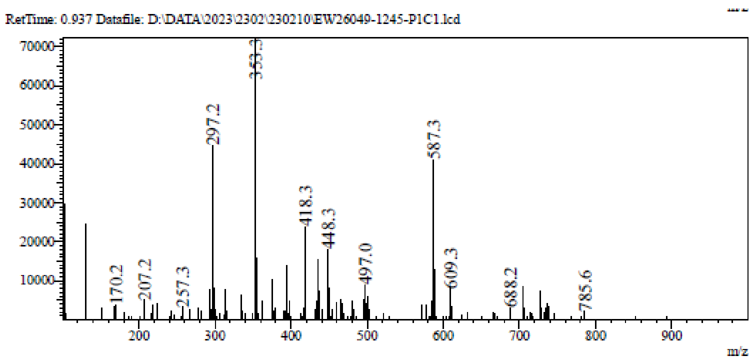
RetTime: 0.937 Datafile: D:\DATA\2023\2302\230210\EW26049-1245-P1C1.lcd
353.3
297.2
587.3
418.3
448.3
497.0
609.3
170.2
207.2
257.3
688.2
785.6
m/z

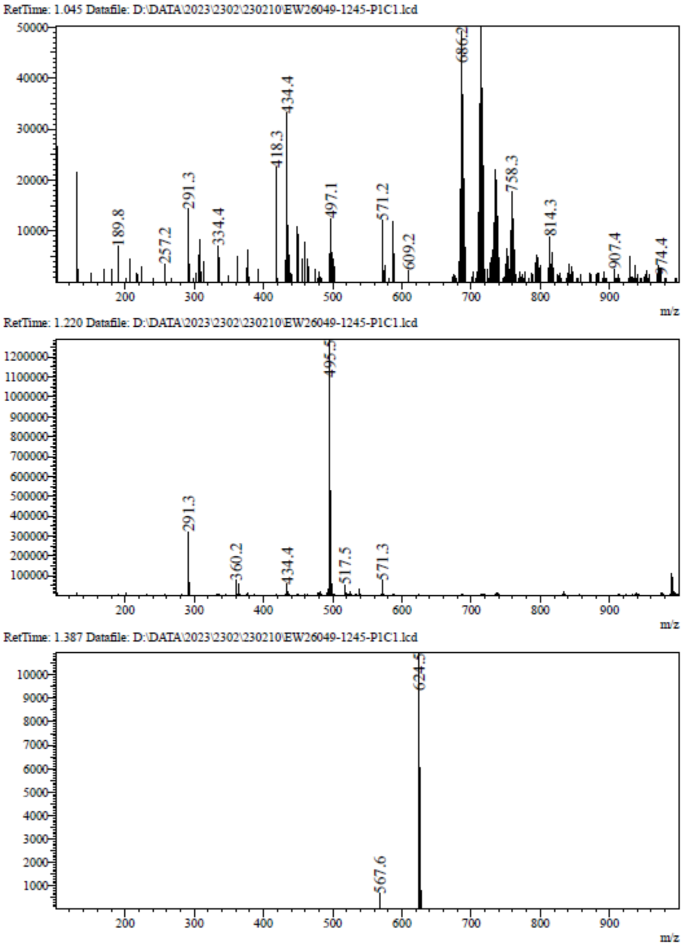

**[1]H NMR** (2*S*,3*R*,4*S*,5*R*)-2-(4-aminopyrrolo[2,1-f][1,2,4]triazin-7-yl)-5-[(*E*)-2-(1,3-benzothiazol-2-yl)vinyl]tetrahydrofuran-3,4-diol **(40)**

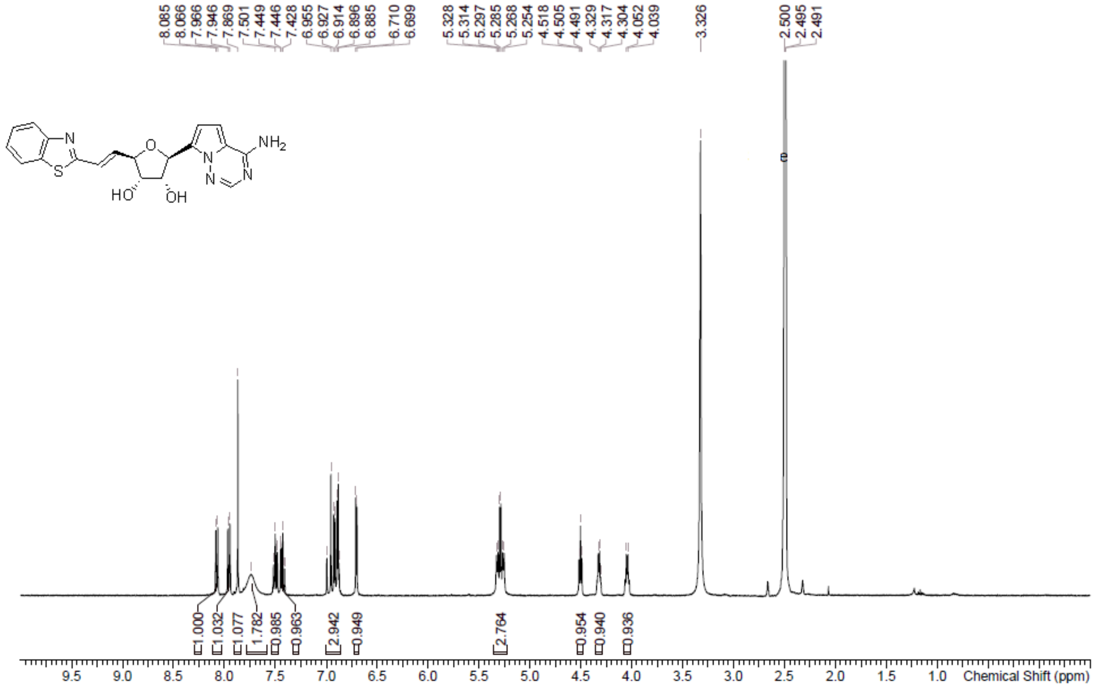

**HPLC** (2*S*,3*R*,4*S*,5*R*)-2-(4-aminopyrrolo[2,1-f][1,2,4]triazin-7-yl)-5-[(*E*)-2- (1,3-benzothiazol-2-yl)vinyl]tetrahydrofuran-3,4-diol **(40)**

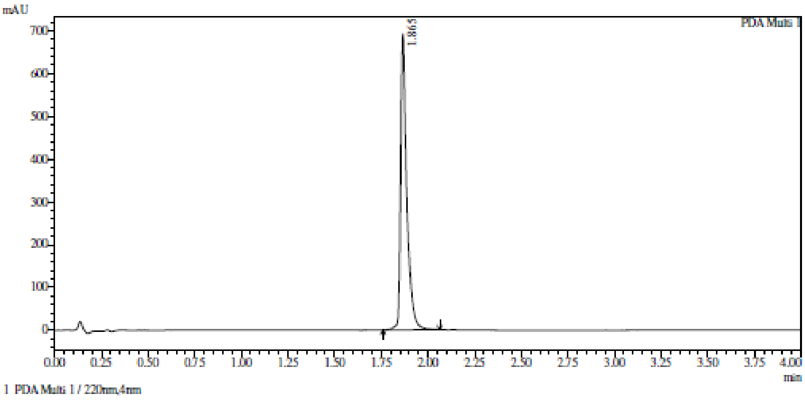

Integration result

PeakTable

PDA Ch1 220nm

| Peak# | Ret. Time | USP Width | Resolution | Height | Area | Area % |
|---|---|---|---|---|---|---|
| 1 | 1.865 | 0.055 | 0.000 | 692840 | 1597361 | 100.000 |
| Total | | | | 692840 | 1597361 | 100.000 |

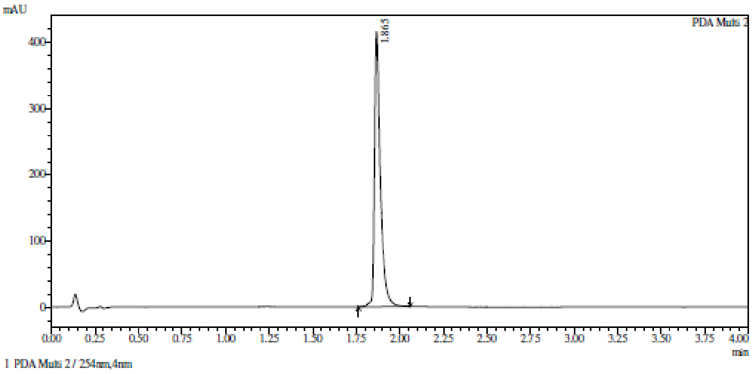

Integration result

PeakTable

PDA Ch2 254nm

| Peak# | Ret. Time | USP Width | Resolution | Height | Area | Area % |
|---|---|---|---|---|---|---|
| 1 | 1.865 | 0.056 | 0.000 | 415790 | 978971 | 100.000 |
| Total | | | | 415790 | 978971 | 100.000 |

**HRMC** (2*S*,3*R*,4*S*,5*R*)-2-(4-aminopyrrolo[2,1-f][1,2,4]triazin-7-yl)-5-[(*E*)-2- (1,3-benzothiazol-2-yl)vinyl]tetrahydrofuran-3,4-diol **(40)**

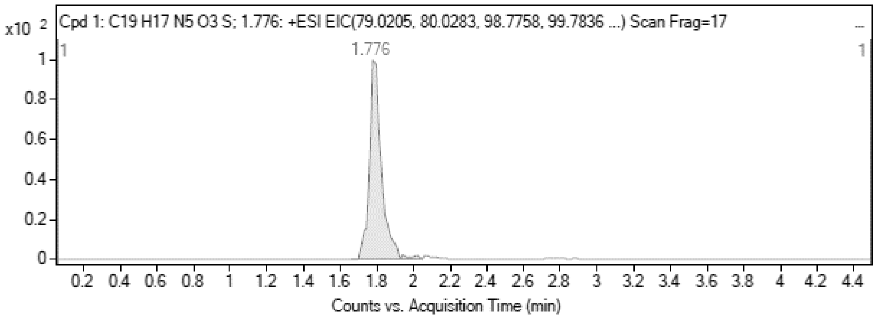

MS Zoomed Spectrum

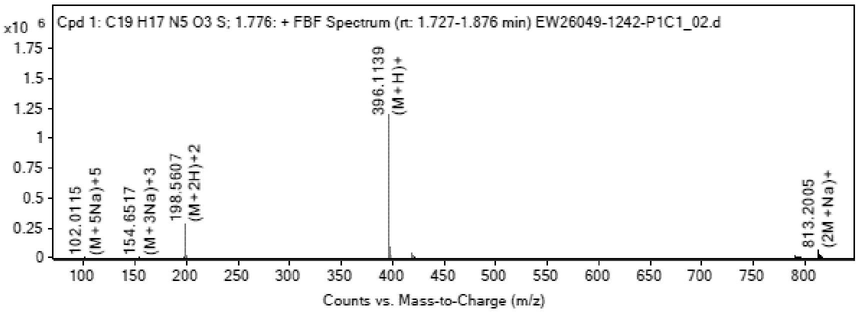

**MS Spectrum Peak List**

| Obs. *m/z* | Charge | Abund | Ion/Isotope |
|---|---|---|---|
| 198.5607 | 2 | 273549.53 | (M+2H)+2 |
| 199.0618 | 2 | 66969.58 | (M+2H)+2 |
| 199.5605 | 2 | 21584.39 | (M+2H)+2 |
| 396.1139 | 1 | 1196108.75 | (M+H)+ |
| 397.116 | 1 | 286005.56 | (M+H)+ |
| 398.113 | 1 | 87600.13 | (M+H)+ |
| 418.0943 | 1 | 42684.21 | (M+Na)+ |
| 813.2005 | 1 | 55999.89 | (2M+Na)+ |
| 814.2029 | 1 | 26855.9 | (2M+Na)+ |
| 815.2003 | 1 | 12059.15 | (2M+Na)+ |

MS Zoomed Spectrum

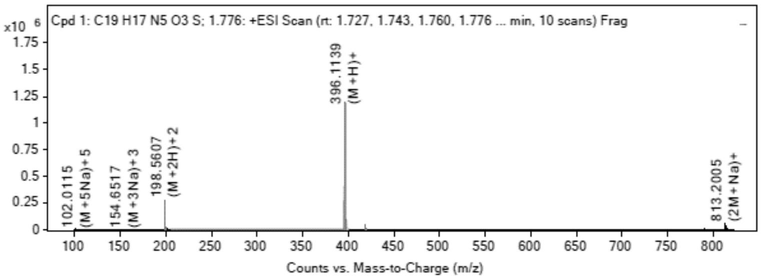

**MS Spectrum Peak List**

| Obs. *m/z* | Charge | Abund | Ion/Isotope | Tgt Mass Error (ppm) |
|---|---|---|---|---|
| 198.5607 | 2 | 273549.53 | (M+2H)+2 | -4.18 |
| 199.0618 | 2 | 66969.58 | (M+2H)+2 | -2.62 |
| 199.5605 | 2 | 21584.39 | (M+2H)+2 | -3.51 |
| 396.1139 | 1 | 1196108.75 | (M+H)+ | -3.56 |
| 397.116 | 1 | 286005.56 | (M+H)+ | -2.02 |
| 398.113 | 1 | 87600.13 | (M+H)+ | -1.76 |
| 418.0943 | 1 | 42684.21 | (M+Na)+ | 0.24 |
| 813.2005 | 1 | 55999.89 | (2M+Na)+ | -1.04 |
| 814.2029 | 1 | 26855.9 | (2M+Na)+ | -0.55 |
| 815.2003 | 1 | 12059.15 | (2M+Na)+ | 0.78 |

--- End Of Report ---

**$^1$H NMR** 7-((2*S*,3*S*,4*R*,5*R*)-5-(2-(benzo[d]thiazol-2-yl)ethyl)-3,4-bis((tert-butyl dimethylsilyl)oxy)tetrahydrofuran-2-yl)pyrrolo[2,1-f][1,2,4]triazin-4-amine **(124)**

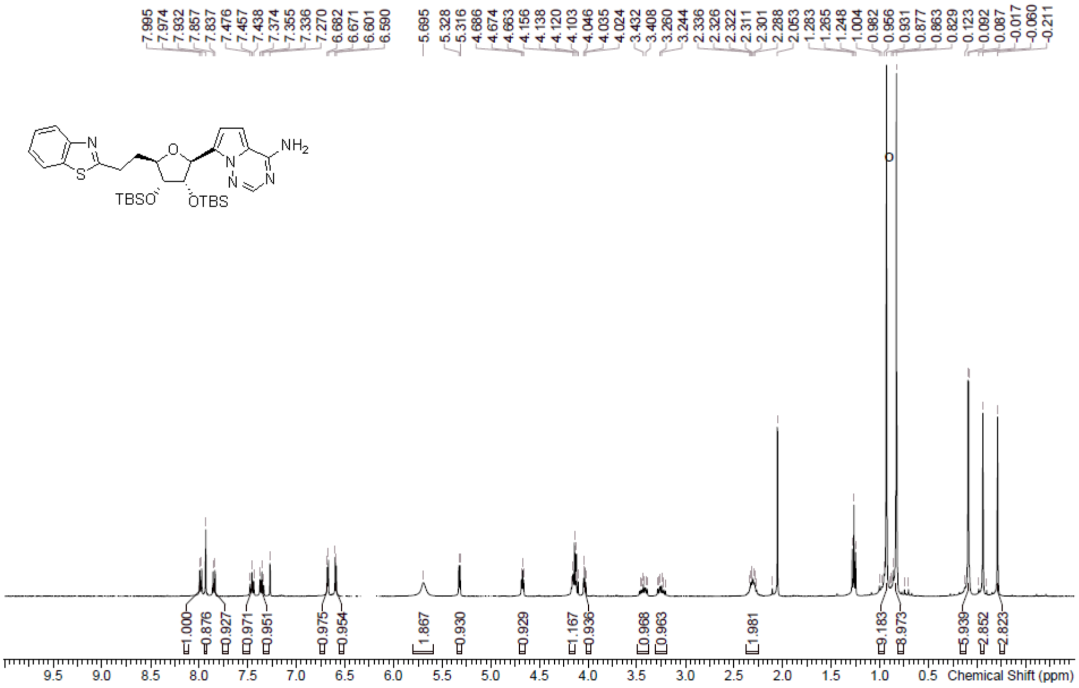

**LCMS** 7-((2*S*,3*S*,4*R*,5*R*)-5-(2-(benzo[d]thiazol-2-yl)ethyl)-3,4-bis((tert-butyl dimethylsilyl)oxy)tetrahydrofuran-2-yl)pyrrolo[2,1-f][1,2,4]triazin-4-amine **(124)**

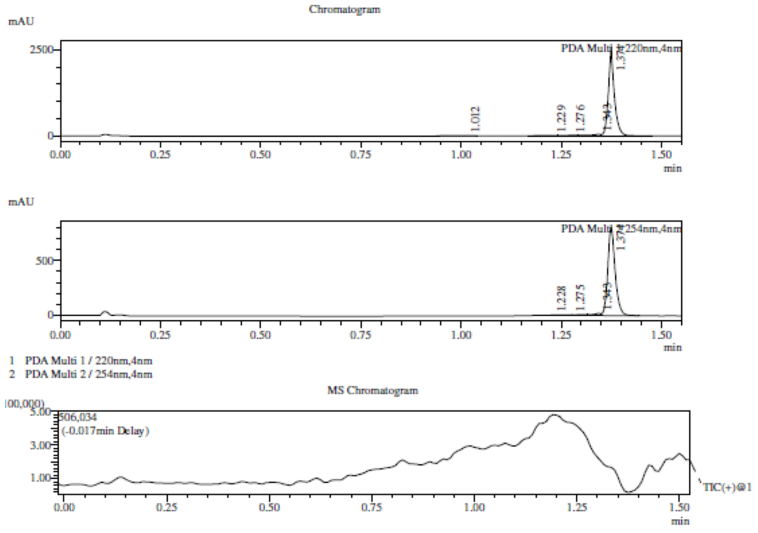

Integration Result

Peak Table

PDA Ch1 220nm

| Peak# | Ret. Time | Height | Height% | USP Width | Area | Area% |
|---|---|---|---|---|---|---|
| 1 | 1.012 | 2280 | 0.085 | 0.092 | 8860 | 0.314 |
| 2 | 1.229 | 8294 | 0.309 | 0.789 | 20726 | 0.735 |
| 3 | 1.276 | 14669 | 0.546 | 0.000 | 37473 | 1.329 |
| 4 | 1.343 | 48085 | 1.789 | 0.045 | 88300 | 3.132 |
| 5 | 1.374 | 2614459 | 97.272 | 0.137 | 2663576 | 94.489 |

Peak Table

PDA Ch2 254nm

| Peak# | Ret. Time | Height | Height% | USP Width | Area | Area% |
|---|---|---|---|---|---|---|
| 1 | 1.228 | 2554 | 0.304 | 0.149 | 7953 | 0.730 |
| 2 | 1.275 | 5432 | 0.646 | 0.654 | 18625 | 1.711 |
| 3 | 1.343 | 20173 | 2.399 | 0.049 | 26077 | 2.395 |
| 4 | 1.374 | 812831 | 96.652 | 0.035 | 1036189 | 95.164 |

Mass Spectrum
RetTime: 1.008 Datafile: D:\DATA\2023\2302\230214\EW26049-1252-P1C1.lcd

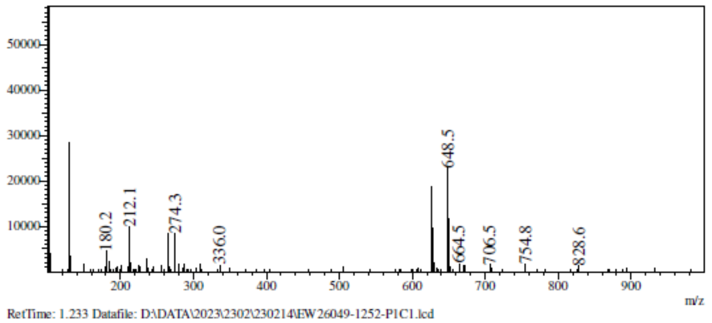

RetTime: 1.233 Datafile: D:\DATA\2023\2302\230214\EW26049-1252-P1C1.lcd

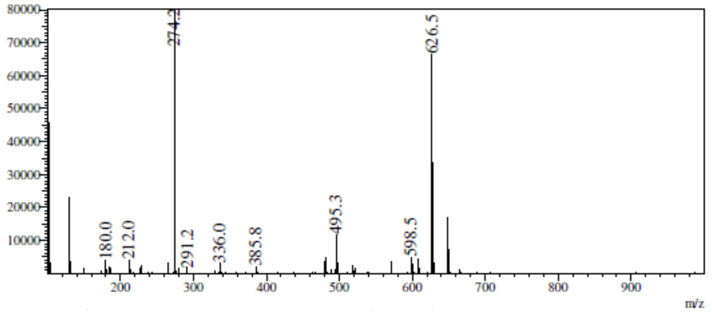

RetTime: 1.275 Datafile: D:\DATA\2023\2302\230214\EW26049-1252-P1C1.lcd

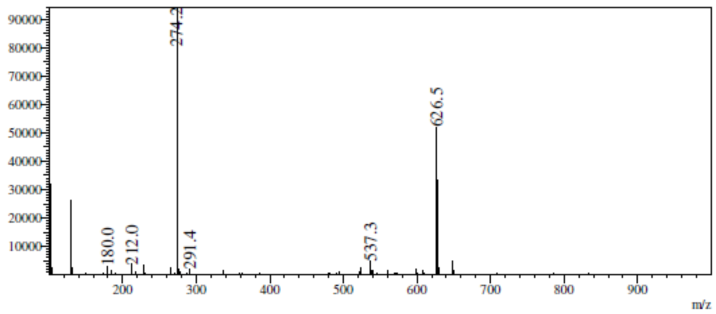

RetTime: 1.342 Datafile: D:\DATA\2023\2302\230214\EW26049-1252-P1C1.lcd

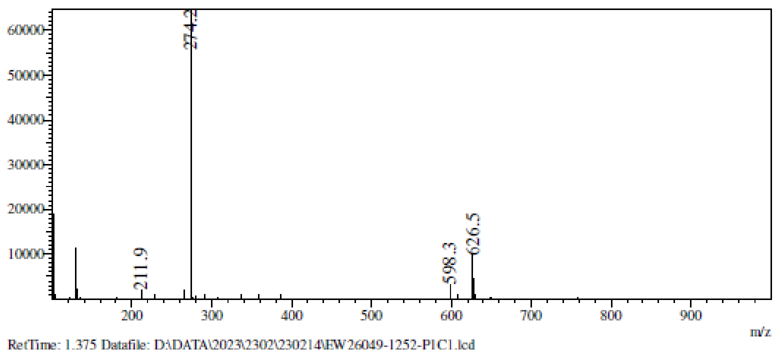

RetTime: 1.375 Datafile: D:\DATA\2023\2302\230214\EW26049-1252-P1C1.lcd

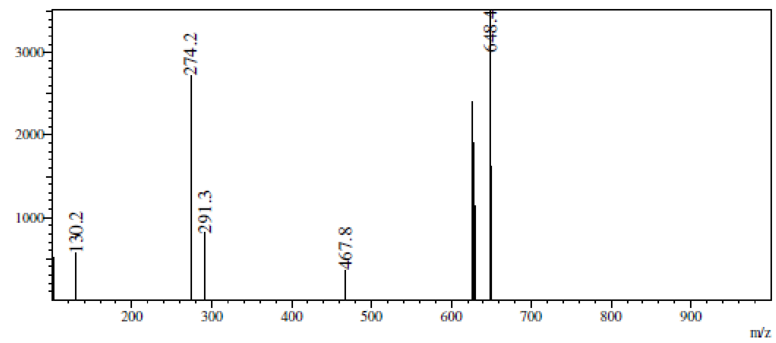

**[1]H NMR** (2*S*,3*R*,4*S*,5*R*)-2-(4-Aminopyrrolo[2,1-f][1,2,4]triazin-7-yl)-5- [2-(1,3-benzothiazol-2-yl)ethyl]tetrahydrofuran-3,4-diol **(39)**

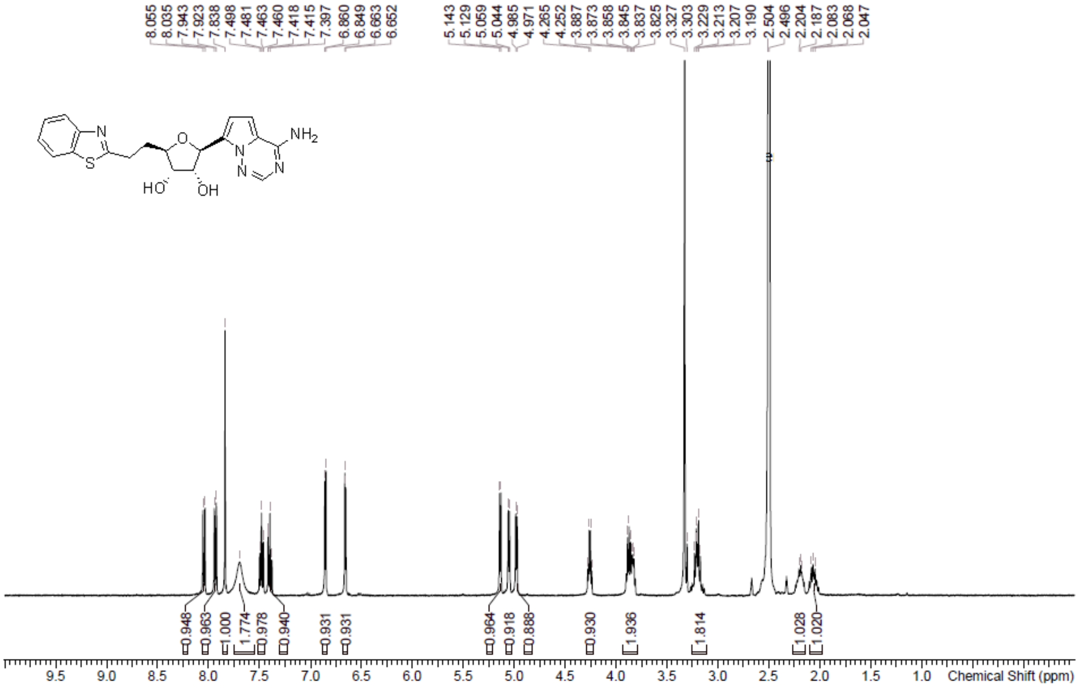

**HPLC** (2*S*,3*R*,4*S*,5*R*)-2-(4-Aminopyrrolo[2,1-f][1,2,4]triazin-7-yl)-5- [2-(1,3-benzothiazol-2-yl)ethyl]tetrahydrofuran-3,4-diol **(39)**

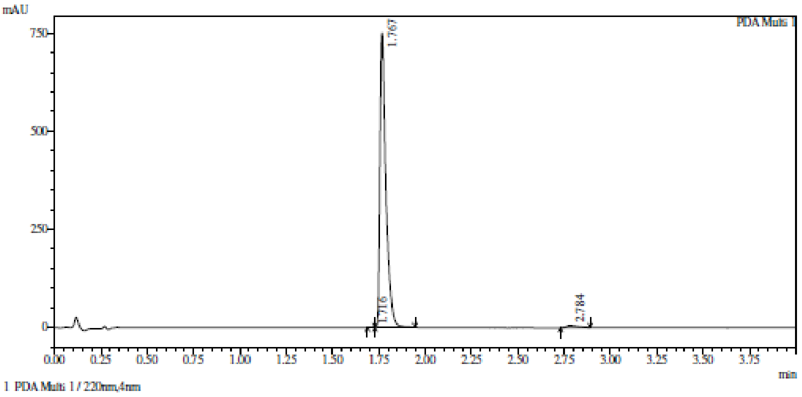

Integration result

PeakTable

PDA Ch1 220nm

| Peak# | Ret. Time | USP Width | Resolution | Height | Area | Area % |
|---|---|---|---|---|---|---|
| 1 | 1.716 | 0.000 | 0.000 | 999 | 1596 | 0.096 |
| 2 | 1.767 | 0.054 | 0.000 | 749570 | 1640868 | 98.651 |
| 3 | 2.784 | 0.110 | 12.387 | 5120 | 20847 | 1.253 |
| Total | | | | 755689 | 1663311 | 100.000 |

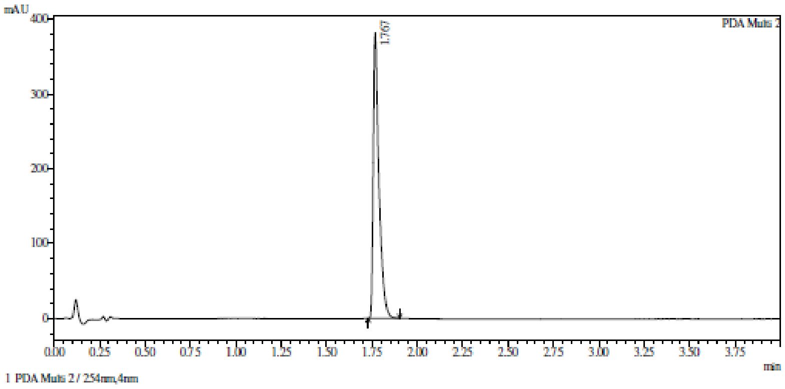

Integration result

PeakTable

PDA Ch2 254nm

| Peak# | Ret. Time | USP Width | Resolution | Height | Area | Area % |
|---|---|---|---|---|---|---|
| 1 | 1.767 | 0.056 | 0.000 | 381870 | 856790 | 100.000 |
| Total | | | | 381870 | 856790 | 100.000 |

**HRMS** (2*S*,3*R*,4*S*,5*R*)-2-(4-Aminopyrrolo[2,1-f][1,2,4]triazin-7-yl)-5- [2-(1,3-benzothiazol-2-yl)ethyl]tetrahydrofuran-3,4-diol **(39)**

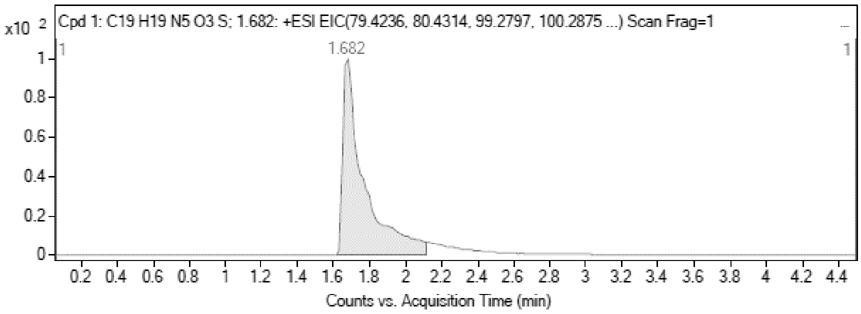

MS Zoomed Spectrum

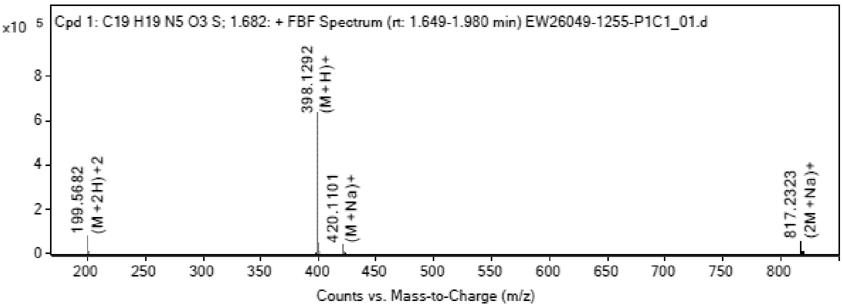

**MS Spectrum Peak List**

| Obs. *m/z* | Charge | Abund | Ion/Isotope |
|---|---|---|---|
| 199.5682 | 2 | 75111.74 | (M+2H)+2 |

| | | | |
|---|---|---|---|
| 200.0695 | 2 | 18732.5 | (M+2H)+2 |
| 398.1292 | 1 | 633819.06 | (M+H)+ |
| 399.1313 | 1 | 152739.59 | (M+H)+ |
| 400.1286 | 1 | 46530.06 | (M+H)+ |
| 420.1101 | 1 | 41638.2 | (M+Na)+ |
| 421.1129 | 1 | 10103.08 | (M+Na)+ |
| 817.2323 | 1 | 54105.62 | (2M+Na)+ |
| 818.2347 | 1 | 26256.98 | (2M+Na)+ |
| 819.2327 | 1 | 11576.3 | (2M+Na)+ |

**MS Zoomed Spectrum**

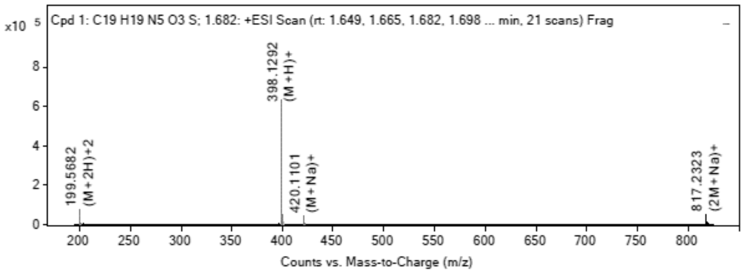

**MS Spectrum Peak List**

| Obs. *m/z* | Charge | Abund | Ion/Isotope | Tgt Mass Error (ppm) |
|---|---|---|---|---|
| 199.5682 | 2 | 75111.74 | (M+2H)+2 | -2.59 |
| 200.0695 | 2 | 18732.5 | (M+2H)+2 | -1.85 |
| 398.1292 | 1 | 633819.06 | (M+H)+ | -2.6 |
| 399.1313 | 1 | 152739.59 | (M+H)+ | -1.07 |
| 400.1286 | 1 | 46530.06 | (M+H)+ | -1.74 |
| 420.1101 | 1 | 41638.2 | (M+Na)+ | -0.06 |
| 421.1129 | 1 | 10103.08 | (M+Na)+ | -0.1 |
| 817.2323 | 1 | 54105.62 | (2M+Na)+ | -1.61 |
| 818.2347 | 1 | 26256.98 | (2M+Na)+ | -1.2 |
| 819.2327 | 1 | 11576.3 | (2M+Na)+ | -0.56 |

--- End Of Report ---
